# Strong evidence for multiple horizontal transfers of immune genes between teleost fishes

**DOI:** 10.1101/2024.12.17.628919

**Authors:** Maxime Policarpo, Walter Salzburger, Florian Maumus, Clément Gilbert

## Abstract

Horizontal gene transfer (HGT) is less frequent in eukaryotes than in prokaryotes, yet can have strong functional implications and was proposed as causal factor for major adaptations in several eukaryotic lineages. Most cases of eukaryote HGT reported to date are inter-domain transfers and only few studies have investigated eukaryote-to-eukaryote HGTs. Furthermore, the extent to which HGT has contributed to the gene repertoire of some lineages such as vertebrates remains unclear. Here, we performed a large-scale survey of HGT among 242 species of ray-finned fishes. We found strong evidence supporting 18 teleost-to-teleost HGT events that involve 17 different genes in 11 teleost fish orders. The genes involved in these transfers show lower synonymous divergence than expected under vertical transmission, their phylogeny is inconsistent with that of teleost fishes, and high nucleotide similarity on these genes between donor and recipient lineages extends to non-coding regions. The distribution of HGT events in the teleost tree is heterogenous, with eight of the 18 transfers occurring between the same two orders (Osmeriformes and Clupeiformes). Besides the previously reported transfer of an antifreeze protein, most transferred genes play roles in immunity, suggesting that such genes are more likely than others to confer strong selective advantage to the recipient species. Overall, our work shows that teleost-to-teleost HGT has occurred on multiple occasions and it will be worth further quantifying these transfers and evaluating their impact on teleost evolution as more genomes are being sequenced.

## Introduction

Horizontal gene transfer (HGT) is the passage of genes between organisms through ways other than sexual or asexual reproduction (Soucy, Huang, and Gogarten 2015). In archaea and bacteria, where several mechanisms and vectors are specifically dedicated to such transfers of genetic material (Haudiquet et al. 2022), HGTs occur frequently and are key to their rapid adaptation to changing environmental conditions (Arnold, Huang, and Hanage 2022). By contrast, the extent to which HGT has shaped eukaryotic genomes and their evolution remains debated (Martin 2017; 2018; Roger 2018; Leger et al. 2018). A consensual trend is that the frequency of HGT is much lower in eukaryotes than in prokaryotes. In unicellular eukaryotes, in which horizontal transfer has been more thoroughly studied, only one to a few percent of genes have been acquired through HGT (Van Etten and Bhattacharya 2020), which is several folds lower compared to prokaryotes. The proportion of foreign genes is even lower in most multicellular eukaryotes investigated so far (but see (Wilson et al. 2024)). That HGT is rarer in eukaryotes than in prokaryotes may be in part explained by the differences in cell organization and metabolism between the two groups, which implies that the kind of genetic variation on which selection operates is less easily obtained by HGT in eukaryotes (changes in complex developmental pathways) than in prokaryotes (metabolic enzymes) (Keeling 2024). However, although rare, it is likely that HGT substantially impacted eukaryote evolution on several occasions, and was even instrumental to the adaptation of multiple lineages to new ecological niches (Keeling 2009; Danchin 2016). For example, many microbial eukaryotes have horizontally acquired genes allowing them to survive in extreme environments, such as icy water (Raymond and Kim 2012), high-salt (Harding, Roger, and Simpson 2017), high-temperature (Schönknecht et al. 2013) or anaerobic conditions (Eme et al. 2017; Van Etten and Bhattacharya 2020). In arthropods, HGT from bacteria and other sources likely facilitated the replicate evolution towards herbivory in mites and in species belonging to at least five insect orders (Kirsch et al. 2022; Xia et al. 2021; Gilbert and Maumus 2022; Wybouw et al. 2016; Wybouw, Van Leeuwen, and Dermauw 2018). Major HGT episodes, again mainly from bacterial sources, may have also facilitated the colonization of land by plants (J. Ma et al. 2022). Another remarkable example is that of the interphotoreceptor retinoid-binding protein (IRBP) gene, which is involved in the formation of the vertebrate eye, and was acquired by the last common ancestor of vertebrates through HGT from bacteria (Kalluraya et al. 2023).

HGT detection is difficult to automate in eukaryotes, and problems linked to contamination have long plagued the field (Bemm et al. 2016; Salzberg 2017). In addition, hypotheses alternative to HGT such as multiple gene losses and/or variation in evolutionary rates among lineages are difficult to exclude in many cases, especially when taxon sampling is sparse (Cote-L’Heureux, Maurer-Alcalá, and Katz 2022). A good illustration of this is that several initial reports of HGT in human have later been shown to be due to technical artefacts (Salzberg et al. 2001; Salzberg 2017; Willerslev et al. 2002). In spite of these limitations, several large-scale studies have reported hundreds of solid cases of HGT in multiple eukaryotic lineages such as microbial eukaryotes (Van Etten and Bhattacharya 2020; Cote-L’Heureux, Maurer-Alcalá, and Katz 2022), plants (J. Ma et al. 2022; Mahelka et al. 2017; Hibdige et al. 2021), fungi (Ciach et al. 2024; Sahu et al. 2023; Marcet-Houben and Gabaldón 2010; Cote-L’Heureux, Maurer-Alcalá, and Katz 2022), and insects (Li et al. 2022). However, most of these studies tended to focus on inter-domain HGT rather than within-eukaryote transfers, because the power to infer HGT (and discard alternative hypotheses) increases as the phylogenetic distance between donor and receiving lineages augments (Adato et al. 2015). Thus, although cases of eukaryote-to-eukaryote HGT have been inferred (Mahelka et al. 2017; Hibdige et al. 2021; Ciach et al. 2024; Sahu et al. 2023; Szöllősi et al. 2015; Marcet-Houben and Gabaldón 2010; Gasmi et al. 2015; Gilbert and Maumus 2022; Mishina et al. 2023), this type of HGT is rarely the subject of large-scale, multi-taxa analyses.

To start filling this gap, we performed a systematic search for HGT events within a vertebrate lineage, ray-finned fishes. We focused on this group because vertebrates in general, and fishes in particular, have never been the subject of large-scale HGT surveys, numerous well-annotated fish genomes are available, ray-finned fishes have incurred massive horizontal transfer of transposable elements (TEs) (H.-H. Zhang et al. 2020), and the only known cases of vertebrate-to-vertebrate HGT reported so far involve two fish lineages (Graham and Davies 2021; Han, Xu, and Gao 2023).

## Results

### A multipronged approach to detect candidate horizontally transferred genes between teleost fishes

To identify HGT between teleosts, we took advantage of 225 high-quality ray-finned fishes genome assemblies from 48 orders, with available annotations on RefSeq or GenBank. We further improved the phylogenetic representativity of this dataset by annotating 17 additional genomes (Fig. 1, Fig. S1, Supplementary files 1 and 2), belonging to 14 orders for which no annotated genomes were available. We then applied a multipronged approach to this dataset, largely inspired from published HGT detection pipelines and metrics. In agreement with the hypothesis proposed by Cote-L’heureux et al. (2022), we postulated that in instances where multiple gene losses mimic a pattern that may be erroneously interpreted as HGT, gene divergence should be consistent with vertical transmission. In contrast, horizontally transferred genes should show a lower divergence than most vertically transmitted genes. To identify the least diverged genes, we classified 5,859,286 genes extracted from the 242 fish genomes in 75,363 hierarchical orthologous groups (HOG) (97% of the 6,035,924 annotated teleost genes). Among each HOG, dS (number of synonymous substitutions per synonymous site) and coding sequence identities were computed between all possible pairs of sequences aligned over at least 100 codons. HOGs appeared to be accurate, as mean dS and quantile values computed from these were highly similar to the values computed using BUSCO groups, which include only single-copy and ultra-conserved genes, that are most likely always vertically transmitted (Fig. S2-A,B). Then, for each possible species pair, we produced a dS distribution, consisting of all dS values computed between the two species across all HOGs. Considering that the dS values of horizontally transferred genes would be located on the left tails of these distributions, we extracted the quantile 0.5% (q0.5). As expected, mean and q0.5 dS values were strongly correlated to divergence times between species, but variances were high, likely reflecting heterogeneity in mutation rates across lineages (Fig. S2-C,D). At this stage, we kept 2,558,049 gene pairs, belonging to 6,367 distinct HOGs, which had a dS value below q0.5.

**Figure 1:**
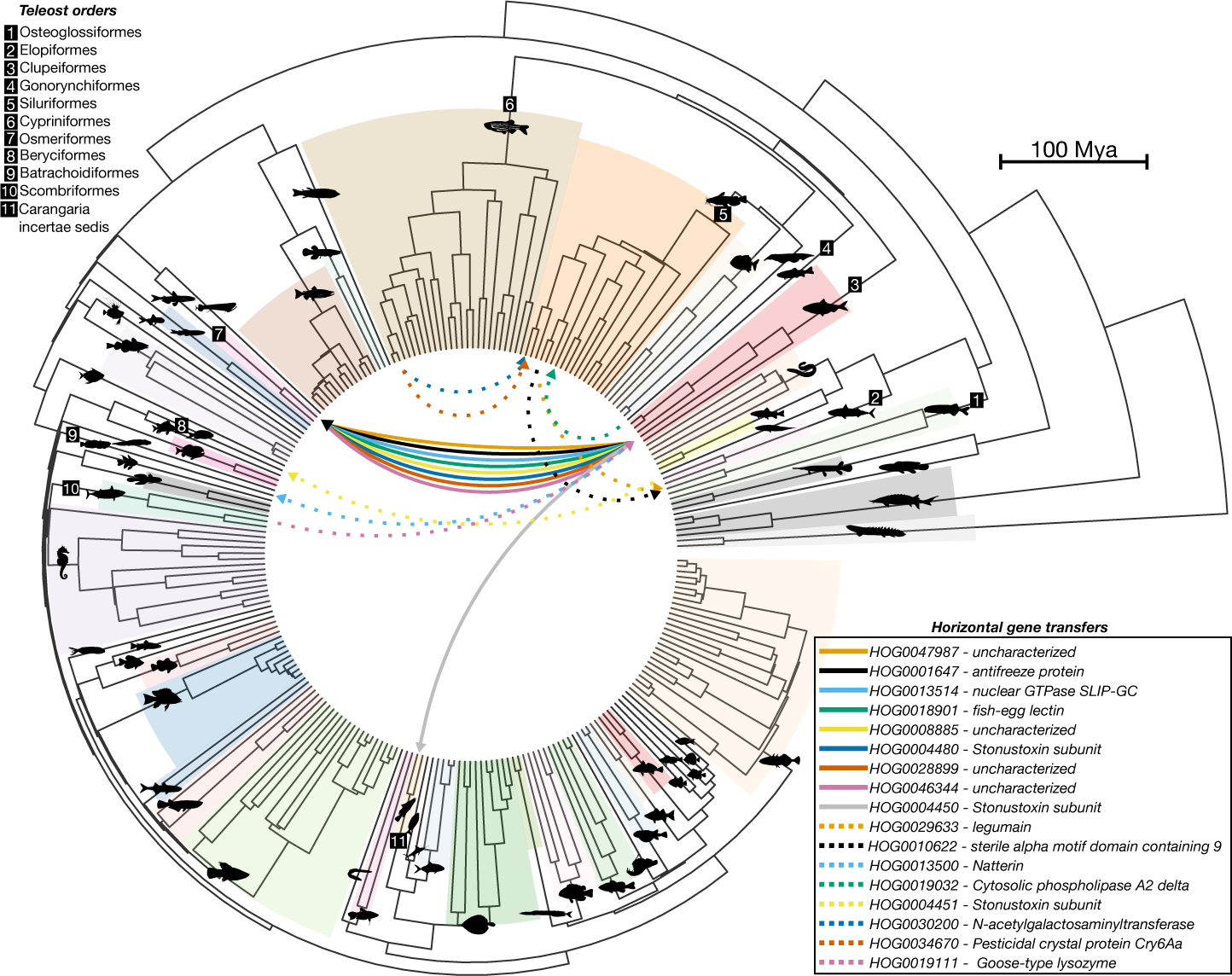
HGTs between teleosts. Phylogeny of annotated ray-finned fishes inferred using ASTRAL-III (with 3,584 BUSCO genes phylogenies) and dated using the least square dating method. The same species tree with species name and annotations sources can be found in Supplementary Figure 1. Alternating shades of color, and silhouettes, represent the different species orders. Orders involved in putative HGTs are annotated with numbers. Branches with no color shades represent orders with only one species in our dataset. Species silhouettes were retrieved in PhyloPic.org (Supplementary file 2). Arrows inside the tree represent putative HGTs, and are drawn from one representative species of the donor clade to one representative species of the recipient clade. Arrows line type and colors are defined based on the HOG. HOG numbers and names (based on the best HHpred match) are indicated.

Next, following (Adato et al. 2015), we reasoned that it would be highly unlikely for a horizontally transferred gene to be located at the same locus, i.e., between the same orthologous genes, in the donor and recipient species. For each pair of species, and for each gene retained above based on low dS and belonging to the same HOG, we counted how many of the ten genes located upstream and the ten genes located downstream of the gene in question in the two species belonged to an identical HOG. This enabled us to compute a micro-syntenic score (akin to the synteny index of Adato et al. 2015), which represents the number of genes belonging to an identical HOG among the 20 flanking genes. A score of 0 indicates the complete lack of micro-synteny for a given gene, while a score of 20 indicates a complete micro-synteny conservation (Fig. S3-A). As expected, the mean micro-syntenic score was highly correlated to divergence time, i.e, the more distantly related species are, the less conserved micro-synteny blocks become (Fig. S3-B). The annotation method also had an impact, as genomes with RefSeq annotations (representing high-quality annotations) had significantly higher mean micro-syntenic scores than other genomes (Fig. S3-B,C). Note that these micro-syntenic scores are slightly biased, as two genes belonging to the same HOG (and are hence orthologs at the level of teleosts most recent common ancestor [MRCA]) are not necessarily orthologs at the level of the MRCA of two species but could represent paralogs (if one or more duplications have occurred anywhere between the teleost MRCA and the MRCA of these two species). We discarded gene pairs that had a non-null micro-syntenic scores, which narrowed down the list of candidate HGTs to 594,324 gene pairs, belonging to 2,381 distinct HOGs.

In parallel to this parametric method, we also used similarity-based approaches, in ways largely inspired by previous studies (Koutsovoulos et al. 2022; Gladyshev, Meselson, and Arkhipova 2008; Boschetti et al. 2012). For this, we performed all-versus-all gene BLASTN searches and the results were filtered using three strategies (detailed in Material and Methods): (i) mismatch between a query corresponding species order and the best-match order (BLASTN-A), (ii) best-match divergence time higher than expected (BLASTN-B), (iii) matches more similar to the query than expected, using the recently described HGTindex alien metrics (Yuan et al. 2023) (BLASTN-C). Again, after discarding gene pairs with a non-null micro-syntenic score, these methods retained 1,000, 1,177 and 522 gene pairs, belonging to 579, 612 and 283 distinct HOGs, respectively.

A total of 3,213 HOGs were retrieved with all the above procedures. 139 HOGs with at least one candidate transfer were retrieved by all three BLASTN strategies, while only 56 HOGs were found in common between our parametric method relying on dS values and all three BLASTN strategies (Fig. 2-A). The largest intersection was found between the BLASTN-A and the BLASTN-B methods, with 270 retained HOGs in common.

**Figure 2:**
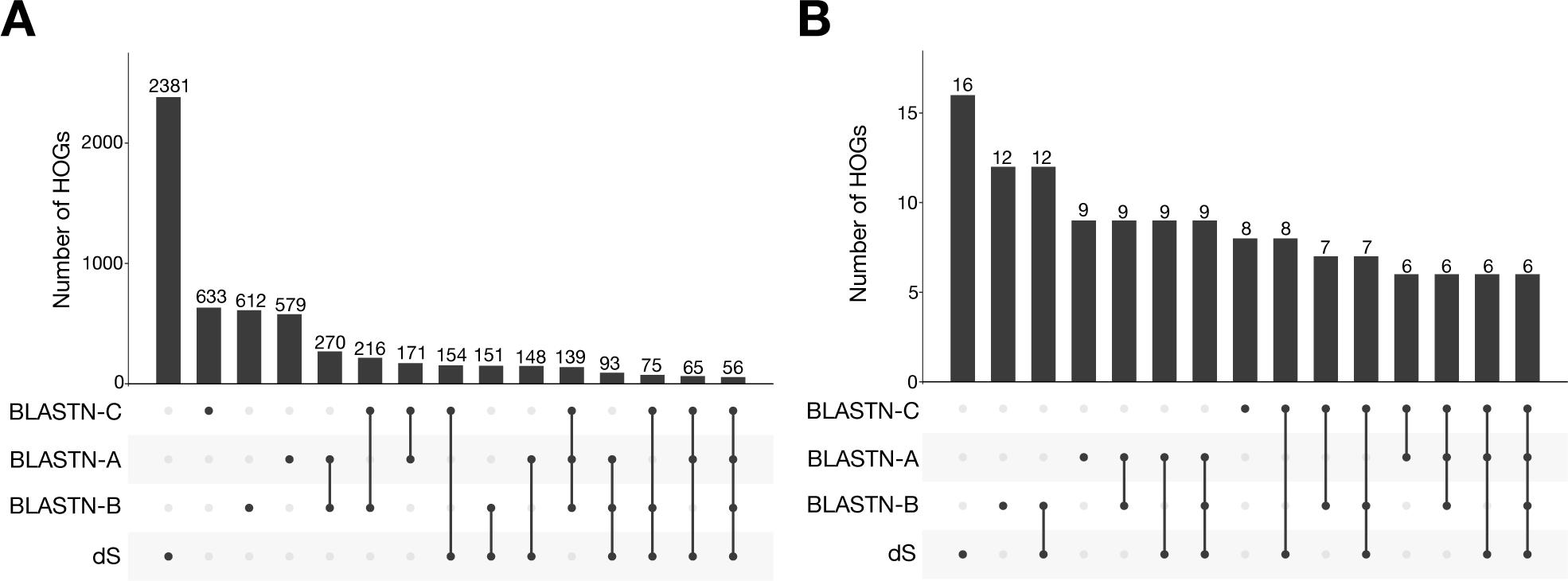
HGTs detection methods. Upset plots showing the number of HOGs retained by the parametric procedure (based on dS) and by the three BLAST procedures (A, B, C). A- Number of unique HOG with at least one candidate HGT retained before manual inspection. B- Number of unique HOG with at least one HGT discussed in this study (after manual curation).

Finally, we investigated whether we could use coding sequences compositions to infer transferred genes across teleost species, as in bacteria (Ravenhall et al. 2015; Dutton and Reiter 2023). We observed large overlaps in the GC content of genes from teleost species, preventing our ability to detect outlier genes in each genome (Fig. S4-A). The same limitation arose regarding codon usage, as all teleost species have a very weak codon usage bias, with ENCprime values comprised between 52.4 and 58.3 (mean = 57) (Fig. S4-B). Furthermore, codon usage profiles, defined by relative synonymous codon usage (RSCU) values of all codons, were highly similar between species (Fig. S4-C).

### Phylogenetic analysis and manual curation of candidate teleost-to-teleost HGTs

As the phylogeny of horizontally transferred genes is generally inconsistent with that of the host taxa (Ravenhall et al. 2015), we computed maximum likelihood phylogenies for the 3,213 retained HOGs and we manually inspected the resulting trees, as is commonly done in HGT studies (Jaramillo, Sukno, and Thon 2015; Shen et al. 2018; Li et al. 2022). We excluded ladder-shape trees (Jaramillo, Sukno, and Thon 2015) and selected trees consistent with a HGT scenario, i.e., trees in which genes from paraphyletic species in the teleost phylogeny form a strongly supported monophyletic group (Fig. 3).

**Figure 3:**
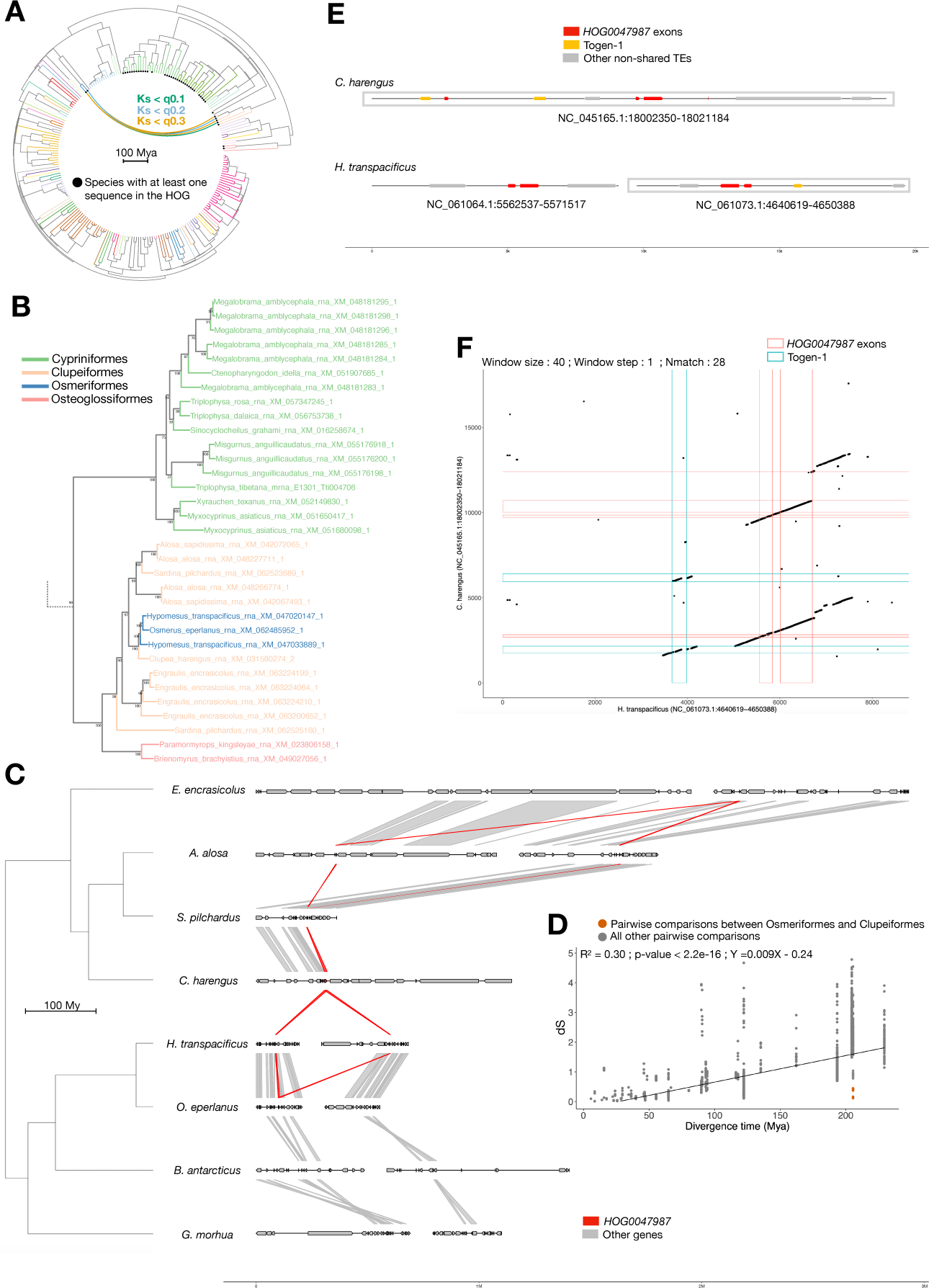
Multiple lines of evidence supporting HGT in the orthogroup HOG0047987. A- Teleost species tree. Lines inside the tree connect two species if they share at least one gene which is less divergent than expected under vertical transmission, i.e., its dS falls below quantile thresholds of the dS distribution computed on all genes that could be aligned between the two species. B- Maximum likelihood phylogeny of the HOG0047987 orthogroup. The tree was pruned to retain the clade with the transfer, as well as close branches. The complete tree can be found in Supplementary file 1. Branches are colored according to the species order. Bootstrap values are shown at each node. Here, the tree topology indicates a transfer from Clupea harengus to the ancestor of Osmeriformes, followed by a duplication, and a subsequent loss of one duplicate in Osmerus eperlanus. C- Micro-synteny plot, showing the region of the transferred genes in the donor and recipient clades (C. harengus and Osmeriformes respectively), as well as in closely related species. Red arrows represent the transferred genes, while gray arrows represent neighboring genes. Genes are linked between species if they belong to the same HOG. D- Relationship between dS values, computed between all HOG0047987 pairs of genes, and divergence times between corresponding species. The linear regression is represented by a black line, and its R^2^, p-value and equation are indicated. Orange dots correspond to pairwise comparisons between C. harengus and Osmeriformes genes. E- Zoom on the regions harboring the transferred gene in C. harengus and Hypomesus transpacificus (Osmeriformes). Transposable elements (TEs) found in only one of these regions are indicated by gray arrows, while those found in both species’ regions are indicated by yellow arrows. Here, one TE, Togen-1, was found in common. Red arrows correspond to the transferred gene exons. Gray boxes indicate regions aligned in F. F- Dot-matrix plot between C. harengus and H. transpacificus regions. Black lines are indicative of conserved regions. Regions between red lines are HOG0047987 exons, while the regions between blue lines correspond to Togen-1. Introns and flanking regions show strong conservations, and, when searched on all teleost genomes, only matched to Osmeriformes and Clupeiformes species. Such multiple lines of evidence, as well as additional analysis (e.g. correction of genes annotations, genes re-annotations and inspection of alternative genome assemblies), are provided for all 18 HGT events discussed in this study, in Supplementary file 1.

In total, we retained 17 HGTs (HGTs) involving 16 distinct HOGs (Fig. 2-B, Table 1, Supplementary file 1). We ensured that the tree topologies of the genes involved in transfer remained consistent with an HGT scenario when based on the same alignment but trimmed to keep the most conserved sites. We further verified that the HGT scenario still holds when reconstructing a phylogeny based on an independent alignment, performed with the nearest sequences found in the UniProt database (The UniProt Consortium 2021) and in a database of ray-finned fishes’ proteins (Supplementary file 1). These phylogenetic analyses allowed us to infer the direction of the transfer (Fig. 3; Supplementary File 1) based on the premise that the receiving clade should appear embedded in the donor clade in the gene tree. While three out of the 17 HGTs were only detected through our dS-based approach, the remaining 14 HGTs were detected by at least two methods (Fig. 2-B, Table 1). Seven HGTs, in six distinct HOGs, were detected by all the methods. In addition to passing the q0.5 threshold established for all HOGs at the level of each pair of species (Fig. 3-A and Supplementary File 1), the dS of HGT candidates are also very low in the dS distribution of genes from the same HOG for all pairs of species (Fig 3-D, Supplementary File 1). Thus, the lower-than-expected dS of these genes cannot be explained by a peculiar selection regime acting on some HOGs. Furthermore, we were able to rule out contamination as an alternative scenario to horizontal transfer as all transferred genes were found either in two different species, for both the donor and recipient clades, or in two independent genome assemblies of the species involved.

**Table 1:** Summary of HGTs between teleosts. Gene names are attributed based on the best HHpred match, along with the associated probability. For each hierarchical orthogroup (HOG), one representative sequence of the transfer was used as input in HHpred (see Material and Methods). “Method” indicates with which method(s) was the HGT detected. “A”, “B” and “C” correspond to the three BLASTN filtering strategies. dS: minimum dS quantile threshold below which pairwise dS values between donor and recipient clades were found. CDS identity: maximum nucleotide identity of the transferred gene between the donor and the recipient clades. The length (in base pairs, bp) of the alignment (denoted “aln”) used to compute the CDS identity, as well as the dS, is indicated between squared brackets. Additional evidence for a horizontal transfer, other than based on the coding sequence, are indicated. I = nucleotide-level similarity in intron(s); F = nucleotide-level similarity in flanking region(s). TE = presence of highly similar transposable elements in the gene flanking regions. The probability of null micro-synteny is given for any gene between the donor and recipient species. It corresponds to the proportion of gene pairs with a micro-syntenic score of 0 between representative species of the donor and recipient clades. For example, there are 19.4% chances that one gene from the same HOG drawn from Clupea harengus and Osmeriformes have a null micro-syntenic score, i.e, with a complete lack of micro-synteny. “Number of losses” indicate the minimum number of losses required in the species tree in the alternative scenario assuming vertical transmission.

| N5 HOG | Name<br>(HHpred<br>probability) | Method | dS | CDS<br>identity<br>[aln length (bp)] | Donor | Recipient | Additional<br>evidences | Probability<br>null<br>micro-syteny | Number of loss<br>(alternative<br>hypothesis) |
| --- | --- | --- | --- | --- | --- | --- | --- | --- | --- |
| HOG0047987 | GCN1 (57.28%) | dS + A + B +<br>C | q0.1 | 88.5%<br>[972] | *Clupea harengus* | Osmeriformes | I + F + TE | 19.4% | 6 |
| HOG0001647 | C type lectin [AFP]<br>(99.79%) | dS + A + B | q0.1 | 91.6%<br>[438] | *Clupea harengus* | Osmeriformes | I + F + TE | 19.4% | 6 |
| HOG0013514 | nuclear GTPase<br>SLIP-GC (99.96%) | dS + A + B<br>+ C | q0.1 | 92.1%<br>[1425] | *Clupea harengus* | Osmeriformes | I + F + TE | 19.4% | 6 |
| HOG0018901 | fish-egg lectin<br>(99.86%) | dS + A + B | q0.3 | 91.7%<br>[675] | *Clupea harengus* | Osmeriformes | I + F + TE | 19.4% | 6 |
| HOG0008885 | Tax1-binding<br>protein 1 (56.97%) | dS + A + B | q0.2 | 81.6%<br>[1584] | *Clupea harengus* | Osmeriformes | F + TE | 19.4% | 6 |
| HOG0004480 | Stonustoxin<br>subunit alpha (100%) | dS | q0.1 | 95.2%<br>[3804] | *Clupea harengus* | Osmeriformes | F | 19.4% | 6 |
| HOG0028899 | De novo<br>design protein<br>(7.07%) | dS + B | q0.5 | 72.6%<br>[336] | Clupeiformes | Osmeriformes | I + F | 19.4% | 5 |
| HOG0004450 | Stonustoxin<br>subunit alpha (100%) | dS | q0.1 | 82.1%<br>[1851] | Clupeiformes | Carangaria | F | 17.4% | 17 |
| HOG0029633 | legumain (100%) | dS + B | q0.1 | 91.1%<br>[585] | Siluriformes | *Paramormyrops*<br>spp. | I + F | 18.9% | 9 |
| HOG0046344 | MifM-stalling<br>(18.43%) | None | q0.6 | 59.4%<br>[591] | Siluriformes | *Paramormyrops*<br>spp. | I + F | 18.9% | 9 |
| HOG0010622 | sterile alpha<br>motif domain<br>containing 9<br>(99.97%) | dS + A + B<br>+ C | q0.1 | 84.8%<br>[4461] | Siluriformes | *Paramormyrops*<br>spp. | None | 18.9% | 9 |
| HOG0013500 | Natterin<br>(99.94%) | dS | q0.2 | 74.6%<br>[1206] | *Clupea harengus* | Batrachoididae | None | 19% | 12 |
| HOG0019032 | Cytosolic<br>phospholipase<br>A2 delta (100%) | dS + A + B<br>+ C | q0.1 | 84.7%<br>[1272] | *Chanos chanos* | *Neoarius* spp. | I + F + TE | 12% | 8 |
| HOG0004451 | Stonustoxin<br>subunit alpha<br>(100%) | dS + C | q0.1 | 81.2%<br>[1413] | *Megalops*<br>*cyprinoides* | Berycidae | F | 10% | 11 |
| HOG0030200 | N-acetylgalactos<br>-aminyltransferase<br>(99.76%) | dS + A + B<br>+ C | q0.1 | 93.8%<br>[1446] | Cypriniformes | Pangasiidae | I + F | 13.7% | 8 |
| HOG0034670 | Pesticidal crystal<br>protein Cry6Aa<br>(99.46%) | dS + B + C | q0.1 | 88.6%<br>[1206] | Cypriniformes | Pangasiidae | I + F + TE | 13.7% | 8 |
| HOG0019111 | Goose-type<br>lysozyme (99.24%) | dS + A + B<br>+ C | q0.1 | 78.3%<br>[456] | Scombridae | Clupeiformes | I + F | 21% | 16 |
| HOG0019111 | Goose-type<br>lysozyme (99.24%) | dS + A + B<br>+ C | q0.2 | 90.2%<br>[471] | *Clupea harengus* | Osmeriformes | I + F | 21% | 16 |

Among these HGTs, we retrieved the only vertebrate-to-vertebrate HGT event described to date: the transfer of an anti-freeze protein (AFP) from *Clupea harengus* (Clupeiformes) to Osmeriformes (*HOG0001647*) (Graham et al. 2008; Graham and Davies 2021). Surprisingly, eight HGTs – about half of all HGTs detected in this study – occurred between these two clades: *HOG0001647*, *HOG0047987*, *HOG0013514*, *HOG0018901*, *HOG0008885*, *HOG0004480*, *HOG0028899* and *HOG0019111* (Fig. 1, Fig. 3, Table 1, Supplementary file 1). There is no indication that these genes were transferred together, as they do not colocalize in the donor and recipient clades (in the two representative highly continuous genomes, *Clupea harengus* and *Hypomesus transpacificus* respectively) (Fig. S5). Interestingly, the *HOG0019111* phylogeny suggests that Clupeiformes firstly acquired this gene from Scombridae before its subsequent transfer to Osmeriformes (Supplementary File 1, Table 1).

Besides these candidate transfers between Osmeriformes and Clupeiformes, we found two HGTs from Cypriniformes to Pangasiidae (*HOG0030200* and *HOG0034670*), one from Clupeiformes to Carangaria (*HOG0004450*), one from Clupeiformes to Batrachoididae (*HOG0013500*), one from *Chanos chanos* to *Neoarius spp*. (*HOG0019032*) and one from *Megalops cyprinoides* to Berycidae species (*HOG0004451*). In addition, we observed a putative transfer of *HOG0029633* from catfishes (Siluriformes) to *Paramormyrops* species (Osteoglossiformes). While investigating the flanking regions of this gene, we found a second likely co-transferred HOG, *HOG0046344*, for which the gene topology was also consistent with an HGT scenario, and for which pairwise dS values were slightly above the q0.5 threshold (< q0.6). Despite these two genes being next to each other in both the donor and recipient clades, null micro-syntenic scores were observed for *HOG0029633,* due to the absence of *HOG0046344* in several catfish species. Finally, one independent HGT (*HOG0010622*) was additionally retrieved between those two clades. Thus, a total of 18 HGTs were found across teleost species.

Additional evidence supporting these HGTs was retrieved and is summarized for each HOG in Supplementary file 1, with the example of *HOG0047987* shown in Fig. 3. First, for most HGTs (16 out of 18), we were able to observe high sequence similarity at the nucleotide level in flanking regions and/or introns, as reported for the AFP gene (Graham et al. 2008; Graham and Davies 2021). We used these regions as queries to perform similarity searches in all the genome assemblies of our dataset and obtained significant hits only in the corresponding donor and recipient clades. Thus, these regions do not correspond to teleost conserved non-coding elements (CNEs), in which one would expect high conservation in many species, through purifying selection. Furthermore, for seven HGTs (including the AFP HGT reported by Graham et al., here *HOG*0001647), we found that the donor and recipient regions harbored at least one common and highly similar TE copy. As reported for the AFP transfer, the TEs found in the vicinity of two additional HGTs (*HOG0047987 and HOG0018901*) occurring between *C. harengus* and *Osmeriformes* further supported the Cluperiformes-to-Osmeriformes direction of the transfer initially inferred based on the HOG topology. Indeed, for each of these three cases, we found one TE (*crack-7*, *togen-1* and *L1-19* respectively), present in many copies across the whole genome of *C. harengus*, but present as single copies and only next to the transferred genes in Osmeriformes.

We next estimated that scenarios alternative to horizontal transfer – that would involve vertical transmission of the 18 HGTs followed by repeated losses – would have required between five and sixteen independent losses across the teleost phylogeny depending on the gene. These numbers were computed assuming that the gene appeared in the MRCA of the donor and recipient species and corresponded to the minimum number of losses needed to retrieve these species in a monophyletic gene clade, assuming zero duplication event, and without considering the rest of the HOG phylogeny.

Finally, we investigated the coherence of these HGTs in terms of species habitats. We found that HGTs always occurred between fresh-water species or between marine species, but never between fresh and marine species. Furthermore, there were always overlaps, of various sizes, in the distributions of donors and recipients (Fig. S6). One limitation to reliably point out the location where these transfers occurred, is that these distributions correspond to extant species, and they may vary from those of ancestor lineages in which the HGTs occurred.

### Function of putative transferred genes

We then investigated the functions of the 17 transferred genes, by first searching for known homologous sequences, using HHpred and profile hidden Markov models (HMMs) (Table 1). As mentioned above, we retrieved the type II antifreeze protein (*HOG0001647*), allowing fish species to survive in icy seawater (Graham and Davies 2021; DeVries 1971). In addition, we found that almost half of all identified HGTs (i.e., 7 genes; Table 1, Fig. 4) are homologous to genes known to be involved in vertebrate immunity. The *HOG0018901* gene is homologous to the fish-egg lectin (*FEL*) which, in several teleost species, is mainly expressed in eggs and larvae, binds bacteria and promotes their phagocytosis by macrophages (Wang et al. 2016; Kai Zhang et al. 2020; Qiao et al. 2022). The *HOG*0019111 gene is homologous to the goose type lysozyme (*LyzG*), an antibacterial enzyme able to break down bacteria (Liu et al. 2022; Tullio, Spaccapelo, and Polimeni 2015). The remaining immune genes homologous to the horizontally transferred genes uncovered here include (i) the nuclear GTPase SLIP-GC (*NUGGC, HOG0013514*), which enhances genome stability of B cells (Richter et al. 2009; 2012), (ii) the cytosolic phospholipase A2 (*PLA2, HOG0019032*), involved in the initiation and amplification of inflammatory responses (Sun et al. 2009; Schilke et al. 2020; Lehr 2001) but also in the defense against pathogens (Pungerčar et al. 2021), (iii) the protease legumain (*HOG0029633*), involved in the presentation of antigens to the MHCII complex (Dall and Brandstetter 2016; Manoury et al. 1998), (iv) the sterile alpha motif domain containing 9 (*SAMD9, HOG0010622*), playing roles in tumor suppressions and in the response to viral infections (Levraud et al. 2019; Nounamo et al. 2017; Q. Ma et al. 2014; Peng et al. 2022), and (v) the beta-1,4 N-acetylgalactosaminyltransferase 2 (*B4GALNT2, HOG0030200*), involved in cancer cells regulation (Groux-Degroote et al. 2014; Furukawa et al. 2014; Pucci, Malagolini, and Dall’Olio 2021).

**Figure 4:**
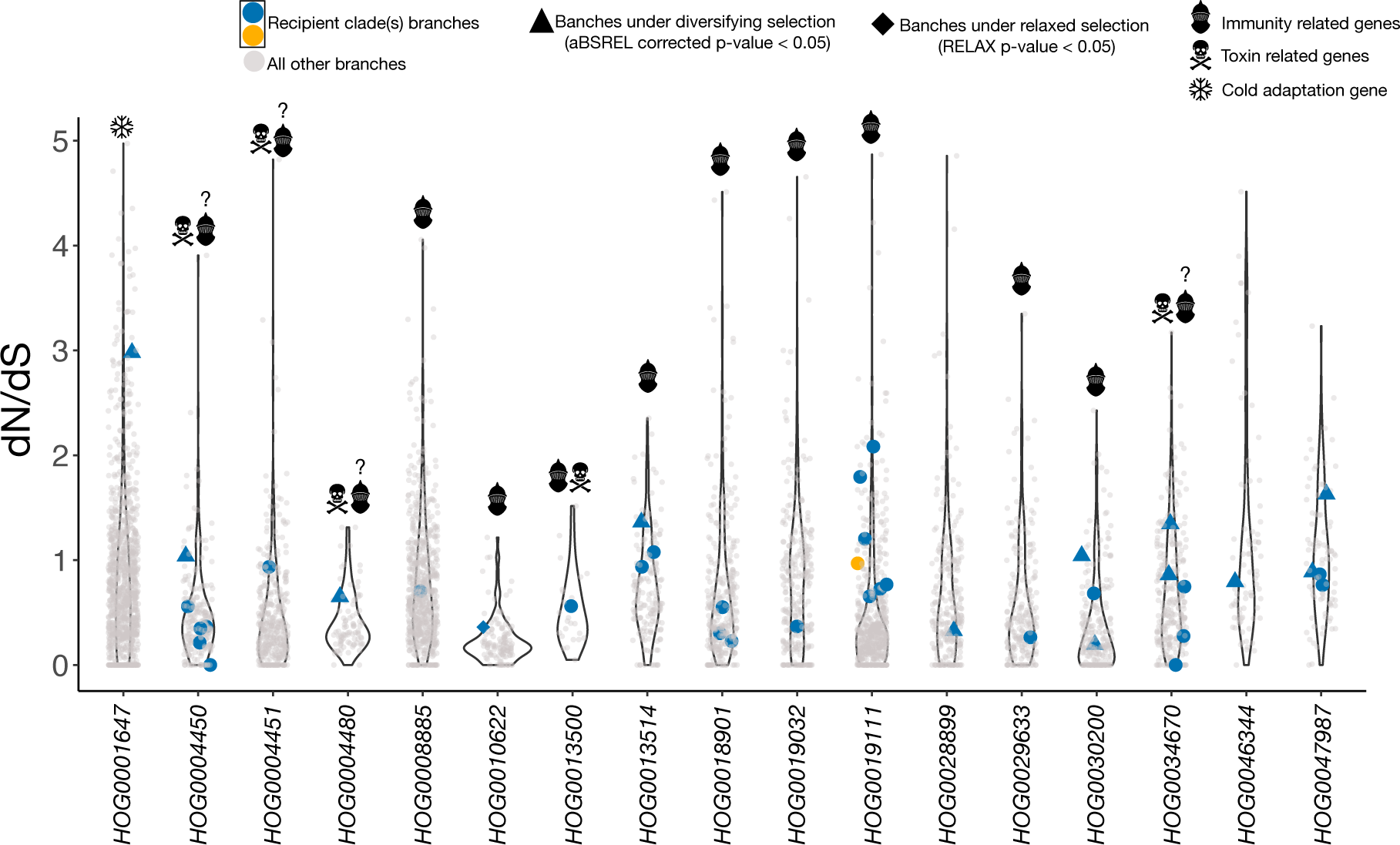
Putative functional category and selective pressures acting on transferred genes. For each transferred HOG, dN/dS of every extant and ancestral sequences (terminal and internal branches, respectively), were computed with HyPhy. dN/dS values computed on branches corresponding to the recipient clades are shown with colored symbols. For HOG0019111, where two transfers occurred, dN/dS values on Clupeiformes branches are indicated in blue, while the unique Osmeriformes branch is shown in orange. Triangles represent sequences evolving under diversifying selection while diamonds represent sequences evolving under relaxed selection. Symbols indicate genes with known roles in immunity, in toxicity or in adaptation to icy waters. For HOG corresponding to pore-forming proteins, the question mark indicates that their role in the immune system is putative, based on knowledges from other PFPs, rather than directly observed for these genes (except for natterins for which immune functions have been shown previously (Seni-Silva et al. 2022; Jia et al. 2016)). Branches with a saturation of substitutions (dS or dN > 1) or not enough substitutions (dS < 0.01) were discarded.

Five of the remaining transferred genes were pore-forming proteins (PFPs). These include one natterin (*HOG0013500*, Table 1, Fig. 4), transferred from *Clupea harengus to Thalassophryne nattereri*, three stonustoxin subunits (*HOG0004480, HOG0004450, HOG0004451*), and one Cry6Aa homolog (*HOG0034670*), the latter having been mainly studied in the bacteria *Bacillus thuringiensis (Dementiev et al. 2016)*. While stonustoxins and natterins have initially been described and studied in toxic fishes (Ghadessy et al. 1996; Magalhães et al. 2006), there is growing evidence for their role in defense against pathogens in fishes, probably preluding their use in venoms (Ellisdon et al. 2015; Lima et al. 2021; Seni-Silva et al. 2022), which is the also the case for many other PFPs (Peraro and van der Goot 2016; Verma et al. 2021; Xiang et al. 2014; Galinier et al. 2013). Interestingly, some natterins have already been proposed to be horizontally transferred between two distant clades of eukaryotes, from fungi to corals (Gacesa et al. 2020). No homologous genes were retrieved for four genes: HOG0047987, HOG0008885, HOG0028899 and HOG0046344.

We then predicted the structures of HGT candidate proteins using AlphaFold3 (Abramson et al. 2024) and searched for similar structures in the AFDP-SwissProt and PDB100 databases using FoldSeek (van Kempen et al. 2024) (Fig. S7). We found that the genes *HOG0008885* and *HOG0028899*, were structurally similar to IL-40 (interleukin 40; probability = 0.66; e-value = 0.02) and CILP1 (cartilage intermediate layer protein 1; probability = 1; e-value = 2.2e-9) respectively. IL-40 is yet another immune gene, being involved in B cells development and immunoglobulin production (Catalan-Dibene, McIntyre, and Zlotnik 2018, 40; Dabbagh-Gorjani 2024). On the other hand, CILP1 is mainly expressed in the extracellular matrix of cartilage tissues, likely involved in its structural integrity (Lorenzo, Bayliss, and Heinegård 1998). The putative function of *HOG0047987* and *HOG0046344* remained elusive, as no structural similarities were retrieved (Fig. S7).

Finally, we computed a dN/dS ratio for every branches of HOGs containing HGTs and used branch-site models to assess the evolutionary regime of transferred genes in the receiving lineage (Fig. 4). First, when looking at dN/dS ratio distributions, we observed that transferred sequences did not appear to have particularly low or high ratios compared to other sequences from the same HOG. For most branches tested (28/41), no sign of diversifying nor relaxed selection was found. For nine HOGs, at least one branch of the recipient clade was evolving under diversifying selection. Furthermore, across nine HOGs constituted of only one recipient branch, only four were evolving under diversifying selection. Importantly, only SAMD9 (*HOG0010622*) was found to evolve under relaxed selection upon reception in the genome of *Paramormyrops kingsleyae (*K= 0.42; p-value = 3.5e-5). Thus, these results indicate that there is, in some cases, an accelerated evolution of the gene upon reception, but this is far from being systematic.

### Pseudoparalogs and post-transfer duplications

While, among the 17 transferred genes, 14 were completely new to the recipient species, three (*HOG0019111*, *HOG0028899* and *HOG0008885*) likely represented pseudoparalogs (Makarova et al. 2005), as the recipient species already possessed at least one other non-transferred gene in the same HOG. Also, in accordance to what has been described in other eukaryotes such as nematodes (Paganini et al. 2012) and algae (Schönknecht, Weber, and Lercher 2014), we found that half (9/18) of HGT events were followed by lineage-specific gene duplications in the recipient clade (Supplementary File 1). This concerns three HGT events between Clupeiformes and Osmeriformes: (i) HOG0047987 with one duplication in the MRCA of Osmeriformes (Figure 3), (ii) HOG0013514 with one duplication specific to *H. transpacificus* and two duplications specific to *O*. eperlanus, (iii) *HOG0018901* with three duplications specific to *H. transpacificus.* The *HOG0004450* transferred gene *was* also duplicated several times after the transfer, one time in *Lates calcarifer* and two in times in *Caranx melampygus.* In addition, there was one post-transfer duplication of *HOG0013500 in Thalassophryne amazonica, one of HOG0030200* in *Pangasius djambal, two of HOG0019032* in *Neoarius graeffei, two of HOG0019111* in *Clupea harengus, and three of HOG0034670 in the MRCA of Pangasianodon species.* These duplication events might have contributed to expand the newly acquired beneficial functional gain (Schönknecht, Weber, and Lercher 2014).

## Discussion

### Strong support for multiple fish-to-fish HGTs

While increasing numbers of HGTs involving eukaryotes are being reported, only few studies have investigated eukaryote-to-eukaryote gene transfers in general and even fewer studies dealt with such type of HGT between vertebrates. Here we surveyed 242 ray-finned fishes’ species and uncovered 18 fish-to-fish HGT events supported by multiple lines of evidence and involving 17 different genes and 11 fish lineages. Among the transferred genes is the AFP protein previously shown to have undergone a HGT event between Clupeiformes and Osmeriformes (Graham et al. 2008; Graham and Davies 2021). All transferred genes were found in multiple independent genomes from the donor and recipient fish clades, ruling out contamination.

Phylogenetic analyses of these genes grouped distantly related fish species in strongly supported monophyletic groups, there was no conserved synteny around transferred genes between the donor and recipient lineages, and most transfers were also recovered using BLAST alien matches approaches (Koutsovoulos et al. 2022; Yuan et al. 2023). Manual curation revealed that in most instances, the high nucleotide identity in coding sequences between donor and recipient lineages extended to intronic and/or flanking sequences, which sometimes even included shared TEs. It is unlikely that these non-coding regions are conserved non-coding elements (CNEs) on the ground that they were not recovered in any other ray-finned fishes, ruling out the possibility that the high similarity detected in these regions between HGT donors and recipient is due to purifying selection on regulatory regions. We even found two cases where, as reported earlier for the AFP transfer (Graham and Davies 2021), the shared TE present in the gene flanking regions was present as a single copy in one species and in multiple copies in a second species. This pattern is consistent with the gene tree topology in strongly suggesting that the second species is the donor, in which the TE generated multiple copies of itself, while the first species is the recipient, in which the TE has not transposed from the horizontally acquired region. Finally, we took special care in ruling out scenario alternative to HGT that would involve multiple gene losses. First, we found that between at least five and 17 independent such losses would be required, depending on the gene, to explain the topology of their phylogeny. These numbers are to be contrasted with the fact that genes showing high similarity between species at the sequence level (both in the coding sequence, introns and flanking regions) are rather expected to show the lowest loss rates (Waterhouse, Zdobnov, and Kriventseva 2010). In fact, we show that the 17 transferred genes uncovered in this study are characterized by extremely low dS values, not only when compared to the distributions of dS values computed on all the genes between the species involved, but also when compared to dS values computed between all the genes among these HOGs (Figure 3, Supplementary Figure 1). Thus, the lower than expected divergence of these genes is inconsistent with them being present in the common ancestors of fishes and enduring recurrent losses during fish evolution (Cote-L’Heureux, Maurer-Alcalá, and Katz 2022).

### Low frequency and heterogenous distribution of fish-to-fish HGTs

Our results are consistent with previous studies of HGT in eukaryotes, which all suggest that the numbers of horizontally acquired genes is many folds lower than in prokaryotes (Van Etten and Bhattacharya 2020; Martin 2017; Danchin 2016; Keeling 2024).Yet, as a first step in evaluating HGT among relatively closely related eukaryote lineages (i.e., among vertebrates), we used stringent criteria and favored a manual curation to retain only strongly supported transfers. Thus, it is likely that the 18 HGTs we inferred only represent a fraction of the total number of gene transfers that have occurred among the fish species included in our dataset. Furthermore, we have not investigated HGT between fish and non-fish species. Our study included 233 out of the >30,000 known teleost species (Near et al. 2012) and the inclusion of many more genomes will likely allow developing more powerful, unsupervised automatic methods to detect HGT. However, the extent to which numbers of HGT among fish will inflate as more genomic data are available is difficult to predict as the distribution of these transfers appears highly heterogeneous in the fish phylogeny. Among the 58 teleost orders included in our study, only 11 were found to be involved in HGT. Strikingly, almost half of the transfers we detected occurred between two of these orders (Clupeiformes and Osmeriformes), with no support from the genomic location of transferred genes in favor of a single co-transfer (Fig. S5). We have no good explanation for this pattern and can only speculate that shedding light on the mechanisms involved in HGT may help resolve it.

### Mechanisms of HGT between fishes

The number of fish-to-fish HGT reported here is low, but finding that such transfers have occurred multiple times is interesting, begging among other questions that of the underlying mechanisms. It has previously been proposed that the mode of reproduction used by most fish species, namely external fertilization, may render their germline more susceptible to be colonized by foreign DNA (Huang 2013; Graham and Davies 2021). All species found involved in HGT in this study use external fertilization. However, at least 12 independent transitions from external to internal fertilization have occurred in fish (Benun Sutton and Wilson 2019), such that as more fish genomes become available, it will be possible to test whether transfers occur more often in external versus internal fertilizers. To increase the power of such an analysis, horizontal transfers of TEs, which seemingly occurred more frequently than HGT among fishes, should be included (H.-H. Zhang et al. 2020). Another hypothesis regularly discussed in the literature posits that viruses may act as vectors of HGT among eukaryotes, as they do in prokaryotes (Gilbert and Cordaux 2017). Much like other animals, fishes are infected by a large diversity of viruses (Costa and Holmes 2024; Costa et al. 2024). These include large dsDNA viruses of the Iridoviridae family (Leiva-Rebollo et al. 2024), which are considered good candidate vectors of horizontal transfer because they can capture host genes (Yoxsimer et al. 2024; Filée, Pouget, and Chandler 2008) and TEs (Piégu et al. 2014; Loiseau et al. 2021) and they are able to infect species diverged since hundreds of million years (Chuang et al. 2022). Other potentially good candidate HGT vectors are the Teratorn endogenous viral elements (EVEs), which result from the fusion of Alloherpesviridae genomes and piggybac DNA transposons (Inoue and Takeda 2023). These elements are widespread in teleost fishes, are able to transpose and produce multiple copies of themselves in a given genome, and their phylogeny suggests they undergo recurrent cross-species transmission (Inoue and Takeda 2023). However, our manual inspection of genomic regions flanking horizontally transferred genes did not reveal any evidence of EVEs. In any case, the HGT events inferred here suggest that the species involved in these transfers are or have been in direct or indirect contact (e.g., through shared viruses or other parasites), which is consistent with the fact that these species currently live in similar environments (marine or fresh-waters) and overlap, to some extent, in their geographical distribution (Fig. S6).

### Opportunity for transfer, retention rate and functional impact of HGT in teleosts

According to Keeling (2024), opportunities for HGT may not be that rare in eukaryotes but it is rather the very low likelihood for a horizontally transferred gene to be retained in the receiving host that may largely explain the lower HGT rates observed in eukaryotes compared to prokaryotes. Such a low retention rate in eukaryotes could be due to the lower probability for a gene acquired through HGT to be beneficial in eukaryotes compared to prokaryotes. The number of HGTs that we can observe today in eukaryotes would thus correspond to a small fraction of a wider foreign gene pool, the majority of which would bring no benefit to the receiving host and be lost through drift or purifying selection. In apparent agreement with this prediction, the number of teleost-to-teleost transfers of TEs (910 among 64 species; (H.-H. Zhang et al. 2020)) is much larger than that of genes (18 among 247 species; this study), indicating that the number of opportunities for transfer might be much higher than what the number of HGTs uncovered in this study suggests. While the capacity of TEs to transpose might, to some extent, result in a higher number of transfer opportunities for these elements compared to genes, there is no doubt that the capacity of TEs to generate multiple copies of themselves strongly increases their retention rate once transferred. In any case, following Keeling (2024)’s argument, horizontally acquired genes are either the result of a recent transfer and did not have time yet to decay (which could be the case of SAMD9, evolving under relaxed selective constraints in the unique recipient species, relative to other fishes), or they were retained because of the strong selective advantage they conferred to the recipient species. Here, we found that most transferred genes were likely involved in fishes’ immunity. Indeed, we observed the transfer of eight genes with clearly established roles in the vertebrate immune system, as well as five PFPs, also likely involved in the innate immunity of fishes (Figure 4). This clear enrichment for immune genes suggests that they are preferentially retained after HGT, and thus more often beneficial to the host than other genes, enhancing and/or diversifying the species defense repertoire. This is in line with many recent studies showing that, across vertebrates, genes under positive selection are often enriched for immune functions, highlighting their rapid evolution and impact on species’ fitness (Xiao et al. 2015; Limborg et al. 2012; Rincon-Sandoval et al. 2024; Vinkler et al. 2023). This also fits well with previous HGTs studies in eukaryotes, in which transferred genes were often found to be involved in the immune system (Ciach et al. 2024; Keeling 2024; Li et al. 2022; Tarnopol et al. 2024; Bai et al. 2016), but also in host-parasite interaction (Gasmi et al. 2021; 2015; Yang et al. 2016; Xia et al. 2021; Gilbert and Maumus 2023; Keeling 2024).

## Materials and Methods

### Genome data and annotations

230 annotated (annotations from RefSeq or GenBank) and 1,200 non-annotated ray-finned fish genomes were downloaded using genome_updater (https://github.com/pirovc/genome_updater) with the option -T “7898” (taxonomic identifier of ray-finned fishes), as of March 9, 2024. We used BUSCO v5.1.274 (Manni et al. 2021) to assess the completeness of each genome (Supplementary File 2). 842 genomes with a BUSCO score ≥ 80% were retained, among which 225 had previously been annotated. The species order of these 842 genomes were extracted using Entrez (Maglott et al. 2007) and the NCBI taxonomy database (Schoch et al. 2020) (Supplementary File 2). For each retained annotated genome, we used agat (Dainat 2022) to keep only the longest transcript per gene and to extract the corresponding coding sequence. Coding sequences with at least one internal stop codon were discarded. All coding sequences from these 225 species were combined and translated into proteins using EMBOSS transeq (Rice, Longden, and Bleasby 2000) to build a ray-finned fishes protein database.

We identified 14 orders for which no annotated genomes were present, but which contained at least one non-annotated genome with a BUSCO score ≥ 80%. We annotated 17 representative species from these 14 orders as follows. First, for each of these species, a *de novo* repeat library was generated using RepeatModeler2 (Flynn et al. 2020). This library was combined with ray-finned fishes repeats extracted from Dfam (Storer et al. 2021), and used as input to soft-mask the corresponding genome assembly using RepeatMasker (Smit, Hubley, and Green 2013). In addition, we used the SRA toolkit to download RNA-seq reads from twelve species with at least one RNA sequencing run present in the SRA database. Finally, these 17 soft-masked genome assemblies were annotated with BRAKER (Gabriel et al. 2024), using as input: (i) our ray-finned fishes protein database, (ii) representative ray-finned fishes proteins extracted from OrthoDBv11 (Kuznetsov et al. 2022), (iii) RNA-seq reads when available (Supplementary File 2). Resulting gene models not supported by any hints were discarded using the “selectSupportedSubsets.py” script provided in BRAKER. We further filtered BRAKER annotations by retaining only the longest transcript per gene and discarding genes with at least one internal stop codon. Thus, our final species dataset comprised 242 annotated ray-finned fish species, among which 233 were teleosts and nine were non-teleosts.

### Species phylogeny

3,584 BUSCO proteins present in at least half of annotated species (≥ 121) were aligned using MUSCLE v5.1 (Edgar 2022) and trimmed using trimAl (Capella-Gutiérrez, Silla-Martínez, and Gabaldón 2009) with the option “-automated1”. A maximum likelihood phylogeny was reconstructed for each alignment using IQ-TREE2 (Minh et al. 2020), with the optimal model found by ModelFinder (Kalyaanamoorthy et al. 2017) and with 1000 ultrafast bootstraps to evaluate the robustness of the nodes (Hoang et al. 2018). Nodes with a low support (< 10% bootstrap support) were collapsed using the R package ape v5.0 (Paradis and Schliep 2019). The 3,584 unrooted gene trees were used as input in ASTRAL-III (C. Zhang et al. 2018) to compute an unrooted species tree. This tree was then rooted and dated using the least square dating method (To et al. 2016) implemented in IQ-TREE2, with three calibration dates, on deep and strongly supported nodes (Near et al. 2012), retrieved on TimeTree.org (Kumar et al. 2022): (i) 250 million years (My) for the split between *Danio rerio* and *Megalops atlanticus*; (ii) 396 My between *Danio rerio* and *Polypterus senegalus*; (iii) 224 My between *Danio rerio* and *Takifugu bimaculatus*.

### Orthologous groups identification, alignment and pairwise metrics

We used OrthoFinder (Emms and Kelly 2019) with (i) the proteomes of our 242 species (for a total of 6,263,181 ray-finned fish protein sequences), (ii) the rooted species tree and (iii) non-teleost species as outgroups. Hierarchical orthologous groups (HOG), as defined in this study, correspond to groups of genes which share a unique origin in the most recent common ancestor (MRCA) of teleost species (node 5 of the species tree, N5). For each HOG, protein sequences were aligned using MUSCLE v5.1 and the alignments were trimmed and back translated to codon alignments using trimAl and the options “-automated1” and “-backtrans”. We then computed four metrics, using the R (R Core Team 2018) packages “seqinr” (Charif and Lobry 2007), “bio3d” (Grant et al. 2006) and “MSA2dist” (Ullrich 2024), between every possible pairs of sequences among each orthogroup: (i) dS, (ii) dN, (iii) number of aligned nucleotides, (iv) sequence identity. In addition, the same alignment procedure and metrics computation were used for BUSCO genes.

### Genes GC content and codon usage bias

For each teleost species, we computed the effective number of codons used per gene, while accounting for the nucleotide composition, thereafter called ENCprime (Novembre 2002). The ENCprime value per species was computed as the mean ENCprime across all genes, by first discarding coding sequences smaller than 100 codons. If each amino acid is encoded by each of its corresponding codons at an equal frequency, ENCprime would have a value of 61. On the other hand, if each amino acid is encoded by a single codon, ENCprime would be equal to 20. It is generally assumed than ENCprime values above 35 indicate a weak codon usage preference, while ENCprime values below 35 indicate codon usage preferences (Gao et al. 2024; Kun Zhang et al. 2024).

We then selected a set of seven representative teleost species across our species tree: *Clupea harengus, Paramormyrops kingsleyae, Megalops cyprinoides, Danio rerio, Hypomesus transpacificus, Pangasianodon hypophthalmus,* and *Oreochromis niloticus* (five of these species were involved in HGTs described in this study, except the zebrafish, *D. rerio,* and the Nile tilapia, *O. niloticus*). For each species, we computed the GC3 content per gene using seqinr (GC content of third-codon position), by first discarding sequences smaller than 100 codons. For each of these seven species, we computed the relative synonymous codon usage (RSCU) of each codon (Sharp and Li 1986), using seqinr. This allowed us to build a codon usage bias profile per species, defined by 64 RSCU values. Codon usage bias profiles between species were compared using pairwise Pearson’s correlations.

### Candidate horizontal gene transfers identification using dS values

For each possible teleost species pair (27,028 pairs), we computed a distribution of all dS values obtained between genes from these two species (only if the genes belonged to the same HOG), by first discarding (i) genes matching to any transposable element present in the RepeatPeps library of RepeatMasker or RepBase (Bao, Kojima, and Kohany 2015) (identified using BLASTP implemented in BLAST+ (Camacho et al. 2009) or Diamond (Buchfink, Reuter, and Drost 2021)), (ii) genes for which less than 100 codons were aligned, (iii) saturated dS values (value of 9.999). Using these distributions, we then computed the 0.5, 0.4, 0.3, 0.2 and 0.1% dS quantiles using the “stat” package in R (R Core Team 2018), as well as the mean and median dS values (Fig. S8). The same procedure was used to compute dS quantiles, mean and median from BUSCO genes.

In addition, for each species pair, and for every genes belonging to the same HOG, we computed micro-syntenic scores, similar to the k synteny index previously described by (Adato et al. 2015). Considering the orthologous gene G_i_ (defined by its presence in the same HOG) present in two species, S_a_ and S_b_, the micro-syntenic score represents the number of orthologous genes among the 20 flanking genes between S_a_ and S_b_ (ten genes upstream and ten genes downstream G_i_). Thus, each pair of genes had a score between zero (if no common HOG were retrieved among flanking genes) and twenty (if all the flanking HOGs were in common). Micro-synteny plots presented in this study were generated using gggenomes (Hackl, J. Ankenbrand, and van Adrichem 2024).

We then removed gene pairs with a micro-syntenic scores ≥ 1, as well as genes present on a scaffold containing only that gene. This led to a final dataset of 594,324 gene pairs belonging to 2,381 distinct orthogroups. Maximum likelihood phylogenies were computed for these 2,381 orthogroups, using IQ-TREE2, and the optimal model found by ModelFinder (using both non-trimmed and trimmed alignments). These trees were plotted using ggtree (Yu et al. 2017) and manually inspected to find topologies consistent with a horizontal gene transfer scenario. We also discarded HOGs for which multiple gene pairs were consistently showing low dS values and for which the phylogenetic tree was ladder-shaped, suggesting selection on synonymous sites and a lack of informative sites (Fig. S9, (Jaramillo, Sukno, and Thon 2015)).

Finally, for each retained genes, we ensured the robustness of the potential HGT topology by performing two BLASTP, using one protein sequence representative of the transfer as query, against (i) a database of all ray-finned fishes proteins and (ii) the Uniprot database devoid of ray-finned fishes proteins (The UniProt Consortium 2021). The 50 and 10 best matches were retained respectively. These protein sequences were aligned using MUSCLE v5, trimmed using trimAl, and a maximum likelihood phylogeny was computed with IQTREE2.

### Further inspections of putative horizontal gene transfers

We further manually inspected putative horizontal gene transfers. First, we verified the quality of protein alignments using the “msaplot” function in ggtree. Then we ensured that among each orthogroup, where one transfer event was inferred, there was a positive correlation between pairwise genes dS values and the divergence time between corresponding species. As expected, the dS values between transferred genes always appeared as outliers, i.e, showed low dS values compared to other pairwise comparisons with the same divergence time.

Furthermore, we selected one representative species from the two clades involved in the gene transfer. For these two species, the transferred gene(s) and flanking regions (5,000 bp upstream and 5,000 bp downstream) were extracted. These regions were aligned using dotter-plot matrixes and the seqinr R package, allowing the identification of similarity at the nucleotide level in introns or in flanking regions. Any similar non-coding region identified on these dotter-plot matrixes were extracted from the genomes using SAMtools (Danecek et al. 2021) and aligned using Needle (Rice, Longden, and Bleasby 2000). These regions were then searched in the genome assemblies of all teleost species using BLASTN. Transposable elements present in these regions were identified through a TBLASTN using a transposable element protein database (combination of RepeatMasker and RepBase databases) as query. Redundancy of this TE database was first reduced by clustering similar proteins with cd-hit (Fu et al. 2012) and a similarity threshold of 80%. Finally, non-overlapping best-hit regions were extracted and used as queries in a BLASTX against the TE database, and these regions were assigned to a transposable element based on the best BLASTX match. TEs found in common between the donor and recipient species were used as queries in a TBLASTN against these species genome assemblies, as well as against the genome assemblies of closely related species. Again, non-overlapping best-hit regions were extracted and the best TE match against these regions was identified using BLASTX. Donor and recipient clades were assigned based on the HOG tree topology and were, in some cases, further supported by neighboring transposable elements, which were present in multiple copies around the donor genome while appeared only in one copy near the transferred gene in the recipient genome.

To rule out contamination as an alternative hypothesis to HGT, we ensured that transferred genes were present in at least two donor and two recipient species. If not, we ensured that these genes were present in another and independent genome assembly of these species. Finally, we systematically verified that the transferred genes were completely absent from species closely related to the donor and recipient clades, using a combination of TBLASTN and Exonerate (Slater and Birney 2005). In some cases, the same strategy was used to correct or complete partial annotations of genes. Detailed analysis results, for each HGT, can be found in Supplementary file 1.

Geographical distributions of species involved in HGT events were extracted from the OBIS and GBIF databases, accessed through the R packages “robis” (Provoost and Bosch 2022) and “rgbif” (Chamberlain and Boettiger 2017). Locations of transferred genes in the genome of *Clupea harengus* and *Hypomesus transpacificus* were represented using the R package “circlize” (Gu et al. 2014).

### Candidate horizontal gene transfers identification using BLAST

We assessed if we could retrieve potential transfers identified using our dS procedure (see above) with another and independent method, relying on BLAST and which is commonly used to find horizontal transfers across the tree of life (Fig. S10; (Koutsovoulos et al. 2022; Yuan et al. 2023; Dutton and Reiter 2023)). First, each coding sequence of each teleost species was used as a query in a BLASTN against all coding sequences from other teleost species, and we discarded genes matching to transposable elements. We then only retained matches if the BLAST alignment had a length ≥ 300 nucleotides and ≥ 85% of the query length.

For the first BLASTN filtering strategy (referred as strategy A in this study), we retained 44,360 queries for which the best match (defined by the lowest e-value), did not belong to the same species order, which we only applied to orders with at least two species.

A second filtering strategy was applied (strategy B). For each species, only the best blast match (defined by the lowest e-value) per gene was retained. Then, for each species, we computed an inverse cumulative distribution of divergence times (which are pseudo-continuous) between this species and species corresponding to the best blast matches. We retained divergence time values for which ≤ 10% of blast match were found, allowing us to identify outliers, i.e., best blast match for which the target species was more distant than expected to the query species.

Finally, for the strategy C, we adapted the HGTindex metric decribed in (Yuan et al. 2023), which was originally designed to detect HGT events between highly distant species, i.e., between life domains or between phyla. For each BLASTN query, we fist retained the best match belonging to the same order, thereafter called BM_ingroup_, as well as the best match belonging to another order, thereafter called BM_outgroup_. Then, we computed the HGTindex per query, defined as the BM_outgroup_ bitscore divided by the BM_ingroup_ bitscore. For each species, we only retained queries with a HGTindex ≥ 0.85 and extracted the species corresponding to the BM_outgroup_, thereafter called outgroup species.

Then, for each query species, we computed the distributions of HGTindex values between this species and all retained outgroup species (one HGTindex distribution per outgroup species). Each of these HGTindex distributions was computed retaining the best matches belonging to the same order (BM_ingroups_) and the best matches belonging to the corresponding outgroup species (i.e. constraining BM_outgroups_ to belong to this species). As described above, HGTindexes were defined as the BM_outgroups_ bitscores divided by the BM_ingroups_ bitscores. Finally, genes were kept if meeting two criteria: (i) passed the initial HGTindex treshhold (≥ 0.85) and (ii) exceeded the 99.5th percentile of the HGTindex distribution for the corresponding outgroup species.

Again, for these three strategies, we discarded gene pairs (pair of a query and its best match) with a micro-syntenic score ≥ 1 as well as genes present on a scaffold containing only that gene. Maximum likelihood phylogenies of corresponding HOGs were computed with IQTREE2, as described above.

### Genes functions and selection analysis

For each transferred gene, we performed homology searches, through profile hidden Markov models, using HHpred (Söding 2005; Zimmermann et al. 2018) and three databases, PDB, Pfam and SMART. One representative protein sequence of each transfer was used: *HOG0046344*, XM_023822671; *HOG0010622*, XM_023831520; *HOG0047987*, XM_047020147; *HOG0013514*, XM_047045767; *HOG0018901*, XM_047023826; *HOG0008885*, XM_047048755; *HOG0004480*, XM_062453720; *HOG0028899*, XM_047031800; *HOG0004450*, XM_042704194; *HOG0029633*, XM_046853148; *HOG0013500*, XM_034166331; *HOG0019032*, XM_060938829; *HOG0004451*, XM_036520634; *HOG0030200*, XM_026945431; *HOG0034670*, XM_053236679; *HOG0019111*, XM_047049897; *HOG0001647*, XM_04703247. Gene names were attributed based on the best hit (highest probability). The 3D structure of the same proteins were predicted using AlphaFold3, on the AlphaFold server (Abramson et al. 2024). Resulting crystallographic information files (CIF) were used as input in FoldSeek (van Kempen et al. 2024) (AFDP-SwissProt and PDB100 databases) to find structurally similar proteins, and the best-hit was retained (highest probability).

For each HOG, we used the standard MG94 fit program of HyPhy, with the coding sequence alignment and the unrooted maximum likelihood phylogeny as input, to compute one dN/dS value per branch (Kosakovsky Pond et al. 2019). Branches with a dN or dS saturation (≥1) or with too few synonymous mutations (dS < 0.01) were discarded. We then used aBSREL (Smith et al. 2015) to infer if internal and terminal branches of recipient clades were evolving under diversifying selection. Each branch for which aBSREL detected a signal of diversifying selection (Holm-Bonferroni corrected p-value < 0.05), were further assigned as test branch in RELAX (Wertheim et al. 2015). A branch was considered under relaxed selection if the RELAX p-value was inferior to 0.05 and if the parameter K (relaxation parameter) was inferior to 1.

## Acknowledgments

We would like to thank Matthieu Haudiquet, as well as the members of the Salzburger laboratory for enriching discussions about this study. All calculations were performed at sciCORE (http://scicore.unibas.ch/), the center of scientific computing at University of Basel (with support by the SIB/Swiss Institute of Bioinformatics). This work was funded by the Swiss National Science Foundation (SNSF; grants 189970 and 208002) and by Agence Nationale de la Recherche (project ANR-24-CE02-1004-01 VIRHOZFER).

## Data availability

All scripts and data needed to reproduce the results of this study are available on GitHub (https://github.com/MaximePolicarpo/HGT_Teleostei) and Zenodo (https://doi.org/10.5281/zenodo.14011660; https://doi.org/10.5281/zenodo.14012814; https://doi.org/10.5281/zenodo.14012963).

## Contributions

M.P., F.M. and C.G. conceived and designed this study, with input from W.S. M.P. performed all data analyses. C.G. and M.P. drafted the manuscript with input and revisions from F.M. and W.S.

## Competing Interest Statement

The authors declare no competing interest.

## Notes

https://github.com/MaximePolicarpo/HGT_Teleostei

https://doi.org/10.5281/zenodo.14011660

https://doi.org/10.5281/zenodo.14012814

https://doi.org/10.5281/zenodo.14012963

### Competing Interest Statement

The authors have declared no competing interest.

## References

1. Abramson, Josh, Jonas Adler, Jack Dunger, Richard Evans, Tim Green, Alexander Pritzel, Olaf Ronneberger, et al. 2024. “Accurate Structure Prediction of Biomolecular Interactions with AlphaFold 3.” Nature 630 (8016): 493–500. 10.1038/s41586-024-07487-w.

2. Adato, Orit, Noga Ninyo, Uri Gophna, and Sagi Snir. 2015. “Detecting Horizontal Gene Transfer between Closely Related Taxa.” PLOS Computational Biology 11 (10): e1004408. 10.1371/journal.pcbi.1004408.

3. Arnold, Brian J., I.-Ting Huang, and William P. Hanage. 2022. “Horizontal Gene Transfer and Adaptive Evolution in Bacteria.” Nature Reviews Microbiology 20 (4): 206–18. 10.1038/s41579-021-00650-4.

4. Bai, Lina, Mu Qiao, Rong Zheng, Changyan Deng, Shuqi Mei, and Wanping Chen. 2016. “Phylogenomic Analysis of Transferrin Family from Animals and Plants.” Comparative Biochemistry and Physiology Part D: Genomics and Proteomics 17 (March):1–8. 10.1016/j.cbd.2015.11.002.

5. Bao, Weidong, Kenji K. Kojima, and Oleksiy Kohany. 2015. “Repbase Update, a Database of Repetitive Elements in Eukaryotic Genomes.” Mobile DNA 6 (1): 11. 10.1186/s13100-015-0041-9.

6. Bemm, Felix, Clemens Leonard Weiß, Jörg Schultz, and Frank Förster. 2016. “Genome of a Tardigrade: Horizontal Gene Transfer or Bacterial Contamination?” Proceedings of the National Academy of Sciences 113 (22): E3054–56. 10.1073/pnas.1525116113.

7. Benun Sutton, Frieda, and Anthony B. Wilson. 2019. “Where Are All the Moms? External Fertilization Predicts the Rise of Male Parental Care in Bony Fishes.” Evolution 73 (12): 2451–60. 10.1111/evo.13846.

8. Boschetti, Chiara, Adrian Carr, Alastair Crisp, Isobel Eyres, Yuan Wang-Koh, Esther Lubzens, Timothy G. Barraclough, Gos Micklem, and Alan Tunnacliffe. 2012. “Biochemical Diversification through Foreign Gene Expression in Bdelloid Rotifers.” PLOS Genetics 8 (11): e1003035. 10.1371/journal.pgen.1003035.

9. Buchfink, Benjamin, Klaus Reuter, and Hajk-Georg Drost. 2021. “Sensitive Protein Alignments at Tree-of-Life Scale Using DIAMOND.” Nature Methods 18 (4): 366–68. 10.1038/s41592-021-01101-x.

10. Camacho, Christiam, George Coulouris, Vahram Avagyan, Ning Ma, Jason Papadopoulos, Kevin Bealer, and Thomas L Madden. 2009. “BLAST+: Architecture and Applications.” BMC Bioinformatics 10 (December):421. 10.1186/1471-2105-10-421.

11. Capella-Gutiérrez, Salvador, José M. Silla-Martínez, and Toni Gabaldón. 2009. “trimAl: A Tool for Automated Alignment Trimming in Large-Scale Phylogenetic Analyses.” Bioinformatics 25 (15): 1972–73. 10.1093/bioinformatics/btp348.

12. Catalan-Dibene, Jovani, Laura L. McIntyre, and Albert Zlotnik. 2018. “Interleukin 30 to Interleukin 40.” Journal of Interferon & Cytokine Research 38 (10): 423. 10.1089/jir.2018.0089.

13. Chamberlain, Scott A., and Carl Boettiger. 2017. “R Python, and Ruby Clients for GBIF Species Occurrence Data.” e3304v1. PeerJ Inc. 10.7287/peerj.preprints.3304v1.

14. Charif, Delphine, and Jean R. Lobry. 2007. “SeqinR 1.0-2: A Contributed Package to the R Project for Statistical Computing Devoted to Biological Sequences Retrieval and Analysis.” In Structural Approaches to Sequence Evolution: Molecules, Networks, Populations, edited by Ugo Bastolla, Markus Porto, H. Eduardo Roman, and Michele Vendruscolo, 207–32. Biological and Medical Physics, Biomedical Engineering. Berlin, Heidelberg: Springer. 10.1007/978-3-540-35306-5_10.

15. Chuang, Hsiang-Chieh, Tah-Wei Chu, Ann-Chang Cheng, Nai-Yu Chen, and Yu-Shen Lai. 2022. “Iridovirus Isolated from Marine Giant Sea Perch Causes Infection in Freshwater Ornamental Fish.” Aquaculture 548 (February):737588. 10.1016/j.aquaculture.2021.737588.

16. Ciach, Michał Aleksander, Julia Pawłowska, Paweł Górecki, and Anna Muszewska. 2024. “The Interkingdom Horizontal Gene Transfer in 44 Early Diverging Fungi Boosted Their Metabolic, Adaptive, and Immune Capabilities.” Evolution Letters 8 (4): 526–38. 10.1093/evlett/qrae009.

17. Costa, Vincenzo A., and Edward C. Holmes. 2024. “Diversity, Evolution, and Emergence of Fish Viruses.” Journal of Virology 98 (6): e00118. 10.1128/jvi.00118-24.

18. Costa, Vincenzo A., Fabrizia Ronco, Jonathon C. O. Mifsud, Erin Harvey, Walter Salzburger, and Edward C. Holmes. 2024. “Host Adaptive Radiation Is Associated with Rapid Virus Diversification and Cross-Species Transmission in African Cichlid Fishes.” Current Biology 34 (6): 1247–1257.e3. 10.1016/j.cub.2024.02.008.

19. Cote-L’Heureux, Auden, Xyrus X. Maurer-Alcalá, and Laura A. Katz. 2022. “Old Genes in New Places: A Taxon-Rich Analysis of Interdomain Lateral Gene Transfer Events.” PLOS Genetics 18 (6): e1010239. 10.1371/journal.pgen.1010239.

20. Dabbagh-Gorjani, Feryal. 2024. “A Comprehensive Review on the Role of Interleukin-40 as a Biomarker for Diagnosing Inflammatory Diseases.” Autoimmune Diseases 2024 (1): 3968767. 10.1155/2024/3968767.

21. Dainat, Jacques. 2022. “AGAT.” Perl. NBIS - National Bioinformatics Infrastructure Sweden. 10.5281/zenodo.3552717.

22. Dall, Elfriede, and Hans Brandstetter. 2016. “Structure and Function of Legumain in Health and Disease.” Biochimie, A potpourri of proteases and inhibitors: from molecular toolboxes to signaling scissors, 122 (March):126–50. 10.1016/j.biochi.2015.09.022.

23. Danchin, Etienne G. J. 2016. “Lateral Gene Transfer in Eukaryotes: Tip of the Iceberg or of the Ice Cube?” BMC Biology 14 (1): 101. 10.1186/s12915-016-0330-x.

24. Danecek, Petr, James K Bonfield, Jennifer Liddle, John Marshall, Valeriu Ohan, Martin O Pollard, Andrew Whitwham, et al. 2021. “Twelve Years of SAMtools and BCFtools.” GigaScience 10 (2): giab008. 10.1093/gigascience/giab008.

25. Dementiev, Alexey, Jason Board, Anand Sitaram, Timothy Hey, Matthew S. Kelker, Xiaoping Xu, Yan Hu, et al. 2016. “The Pesticidal Cry6Aa Toxin from Bacillus Thuringiensis Is Structurally Similar to HlyE-Family Alpha Pore-Forming Toxins.” BMC Biology 14 (1): 71. 10.1186/s12915-016-0295-9.

26. DeVries, A. L. 1971. “Glycoproteins as Biological Antifreeze Agents in Antarctic Fishes.” Science (New York, N.Y.) 172 (3988): 1152–55. 10.1126/science.172.3988.1152.

27. Dutton, Rachel J., and Taylor Reiter. 2023. “PreHGT: A Scalable Workflow That Screens for Horizontal Gene Transfer within and between Kingdoms.” Arcadia Science, July. 10.57844/arcadia-jfbp-7p11.

28. Edgar, Robert C. 2022. “Muscle5: High-Accuracy Alignment Ensembles Enable Unbiased Assessments of Sequence Homology and Phylogeny.” Nature Communications 13 (November):6968. 10.1038/s41467-022-34630-w.

29. Ellisdon, Andrew M., Cyril F. Reboul, Santosh Panjikar, Kitmun Huynh, Christine A. Oellig, Kelly L. Winter, Michelle A. Dunstone, et al. 2015. “Stonefish Toxin Defines an Ancient Branch of the Perforin-like Superfamily.” Proceedings of the National Academy of Sciences of the United States of America 112 (50): 15360. 10.1073/pnas.1507622112.

30. Eme, Laura, Eleni Gentekaki, Bruce Curtis, John M. Archibald, and Andrew J. Roger. 2017. “Lateral Gene Transfer in the Adaptation of the Anaerobic Parasite Blastocystis to the Gut.” Current Biology: CB 27 (6): 807–20. 10.1016/j.cub.2017.02.003.

31. Emms, David M., and Steven Kelly. 2019. “OrthoFinder: Phylogenetic Orthology Inference for Comparative Genomics.” Genome Biology 20 (1): 238. 10.1186/s13059-019-1832-y.

32. Filée, Jonathan, Noëlle Pouget, and Mick Chandler. 2008. “Phylogenetic Evidence for Extensive Lateral Acquisition of Cellular Genes by Nucleocytoplasmic Large DNA Viruses.” BMC Evolutionary Biology 8 (1): 320. 10.1186/1471-2148-8-320.

33. Flynn, Jullien M., Robert Hubley, Clément Goubert, Jeb Rosen, Andrew G. Clark, Cédric Feschotte, and Arian F. Smit. 2020. “RepeatModeler2 for Automated Genomic Discovery of Transposable Element Families.” Proceedings of the National Academy of Sciences 117 (17): 9451–57. 10.1073/pnas.1921046117.

34. Fu, Limin, Beifang Niu, Zhengwei Zhu, Sitao Wu, and Weizhong Li. 2012. “CD-HIT: Accelerated for Clustering the next-Generation Sequencing Data.” Bioinformatics 28 (23): 3150–52. 10.1093/bioinformatics/bts565.

35. Furukawa, Koichi, Keiko Furukawa, Yuhsuke Ohmi, Yuki Ohkawa, Yoshio Yamauchi, Noboru Hashimoto, and Orie Tajima. 2014. “Beta-1,4 N-Acetylgalactosaminyltransferase 1,2 (B4GALNT1,2).” In Handbook of Glycosyltransferases and Related Genes, edited by Naoyuki Taniguchi, Koichi Honke, Minoru Fukuda, Hisashi Narimatsu, Yoshiki Yamaguchi, and Takashi Angata, 417–28. Tokyo: Springer Japan. 10.1007/978-4-431-54240-7_34.

36. Gabriel, Lars, Tomáš Brůna, Katharina J. Hok, Matthis Ebel, Alexandre Lomsadze, Mark Borodovsky, and Mario Stanke. 2024. “BRAKER3: Fully Automated Genome Annotation Using RNA-Seq and Protein Evidence with GeneMark-ETP, AUGUSTUS, and TSEBRA.” Genome Research 34 (5): 769–77. 10.1101/gr.278090.123.

37. Gacesa, Ranko, Julia Yun-hsuan Hung, David G. Bourne, and Paul F. Long. 2020. “Horizontal Transfer of a Natterin-like Toxin Encoding Gene within the Holobiont of the Reef Building Coral Acropora Digitifera (Cnidaria: Anthozoa: Scleractinia) and across Multiple Animal Linages.” Journal of Venom Research 10 (April):7.

38. Galinier, Richard, Julien Portela, Yves Moné, Jean François Allienne, Hélène Henri, Stéphane Delbecq, Guillaume Mitta, Benjamin Gourbal, and David Duval. 2013. “Biomphalysin, a New β Pore-Forming Toxin Involved in Biomphalaria Glabrata Immune Defense against Schistosoma Mansoni.” PLoS Pathogens 9 (3): e1003216. 10.1371/journal.ppat.1003216.

39. Gao, Wei, Xiaodie Chen, Jing He, Ajia Sha, Yingyong Luo, Wenqi Xiao, Zhuang Xiong, and Qiang Li. 2024. “Intraspecific and Interspecific Variations in the Synonymous Codon Usage in Mitochondrial Genomes of 8 Pleurotus Strains.” BMC Genomics 25 (May):456. 10.1186/s12864-024-10374-3.

40. Gasmi, Laila, Helene Boulain, Jeremy Gauthier, Aurelie Hua-Van, Karine Musset, Agata K. Jakubowska, Jean-Marc Aury, et al. 2015. “Recurrent Domestication by Lepidoptera of Genes from Their Parasites Mediated by Bracoviruses.” PLOS Genetics 11 (9): e1005470. 10.1371/journal.pgen.1005470.

41. Gasmi, Laila, Edyta Sieminska, Shohei Okuno, Rie Ohta, Cathy Coutu, Mohammad Vatanparast, Stephanie Harris, et al. 2021. “Horizontally Transmitted Parasitoid Killing Factor Shapes Insect Defense to Parasitoids.” Science 373 (6554): 535–41. 10.1126/science.abb6396.

42. Ghadessy, Farid John, Desong Chen, R. Manjunatha Kini, Maxey C. M. Chung, Kandiah Jeyaseelan, Hoon Eng Khoo, and Raymond Yuen. 1996. “Stonustoxin Is a Novel Lethal Factor from Stonefish (*Synanceja Horrida*) Venom.” Journal of Biological Chemistry 271 (41): 25575–81. 10.1074/jbc.271.41.25575.

43. Gilbert, Clément, and Richard Cordaux. 2017. “Viruses as Vectors of Horizontal Transfer of Genetic Material in Eukaryotes.” Current Opinion in Virology, Animal models for viral diseases • Paleovirology, 25 (August):16–22. 10.1016/j.coviro.2017.06.005.

44. Gilbert, Clément, and Florian Maumus. 2022. “Multiple Horizontal Acquisitions of Plant Genes in the Whitefly Bemisia Tabaci.” Genome Biology and Evolution 14 (10): evac141. 10.1093/gbe/evac141.

45. Gilbert, Clément, and Florian Maumus. 2023. “Sidestepping Darwin: Horizontal Gene Transfer from Plants to Insects.” Current Opinion in Insect Science 57 (June):101035. 10.1016/j.cois.2023.101035.

46. Gladyshev, Eugene A., Matthew Meselson, and Irina R. Arkhipova. 2008. “Massive Horizontal Gene Transfer in Bdelloid Rotifers.” Science 320 (5880): 1210–13. 10.1126/science.1156407.

47. Graham, Laurie A., and Peter L. Davies. 2021. “Horizontal Gene Transfer in Vertebrates: A Fishy Tale.” Trends in Genetics 37 (6): 501–3. 10.1016/j.tig.2021.02.006.

48. Graham, Laurie A., Stephen C. Lougheed, K. Vanya Ewart, and Peter L. Davies. 2008. “Lateral Transfer of a Lectin-Like Antifreeze Protein Gene in Fishes.” PLoS ONE 3 (7): e2616. 10.1371/journal.pone.0002616.

49. Grant, Barry J., Ana P. C. Rodrigues, Karim M. ElSawy, J. Andrew McCammon, and Leo S. D. Caves. 2006. “Bio3d: An R Package for the Comparative Analysis of Protein Structures.” Bioinformatics 22 (21): 2695–96. 10.1093/bioinformatics/btl461.

50. Groux-Degroote, Sophie, Cindy Wavelet, Marie-Ange Krzewinski-Recchi, Lucie Portier, Marlène Mortuaire, Adriana Mihalache, Marco Trinchera, et al. 2014. “*B4GALNT2* Gene Expression Controls the Biosynthesis of Sda and Sialyl Lewis X Antigens in Healthy and Cancer Human Gastrointestinal Tract.” The International Journal of Biochemistry & Cell Biology 53 (August):442–49. 10.1016/j.biocel.2014.06.009.

51. Gu, Zuguang, Lei Gu, Roland Eils, Matthias Schlesner, and Benedikt Brors. 2014. “Circlize Implements and Enhances Circular Visualization in R.” Bioinformatics 30 (19): 2811–12. 10.1093/bioinformatics/btu393.

52. Hackl, Thomas, Markus J. Ankenbrand, and Bart van Adrichem. 2024. “Gggenomes.” R. https://github.com/thackl/gggenomes.

53. Han, Zhiqiang, Shengyong Xu, and Tianxiang Gao. 2023. “Unexpected Complex Horizontal Gene Transfer in Teleost Fish.” Current Zoology 69 (2): 222–23. 10.1093/cz/zoac032.

54. Harding, Tommy, Andrew J. Roger, and Alastair G. B. Simpson. 2017. “Adaptations to High Salt in a Halophilic Protist: Differential Expression and Gene Acquisitions through Duplications and Gene Transfers.” Frontiers in Microbiology 8 (May). 10.3389/fmicb.2017.00944.

55. Haudiquet, Matthieu, Jorge Moura de Sousa, Marie Touchon, and Eduardo P. C. Rocha. 2022. “Selfish, Promiscuous and Sometimes Useful: How Mobile Genetic Elements Drive Horizontal Gene Transfer in Microbial Populations.” Philosophical Transactions of the Royal Society B: Biological Sciences 377 (1861): 20210234. 10.1098/rstb.2021.0234.

56. Hibdige, Samuel G. S., Pauline Raimondeau, Pascal-Antoine Christin, and Luke T. Dunning. 2021. “Widespread Lateral Gene Transfer among Grasses.” New Phytologist 230 (6): 2474–86. 10.1111/nph.17328.

57. Hoang, Diep Thi, Olga Chernomor, Arndt von Haeseler, Bui Quang Minh, and Le Sy Vinh. 2018. “UFBoot2: Improving the Ultrafast Bootstrap Approximation.” Molecular Biology and Evolution 35 (2): 518–22. 10.1093/molbev/msx281.

58. Huang, Jinling. 2013. “Horizontal Gene Transfer in Eukaryotes: The Weak-Link Model.” BioEssays 35 (10): 868–75. 10.1002/bies.201300007.

59. Inoue, Yusuke, and Hiroyuki Takeda. 2023. “Teratorn and Its Relatives – a Cross-Point of Distinct Mobile Elements, Transposons and Viruses.” Frontiers in Veterinary Science 10 (April). 10.3389/fvets.2023.1158023.

60. Jaramillo, Vinicio D. Armijos, Serenella A. Sukno, and Michael R. Thon. 2015. “Identification of Horizontally Transferred Genes in the Genus Colletotrichum Reveals a Steady Tempo of Bacterial to Fungal Gene Transfer.” BMC Genomics 16 (1): 2. 10.1186/1471-2164-16-2.

61. Jia, Ning, Nan Liu, Wang Cheng, Yong-Liang Jiang, Hui Sun, Lan-Lan Chen, Junhui Peng, et al. 2016. “Structural Basis for Receptor Recognition and Pore Formation of a Zebrafish Aerolysin-like Protein.” EMBO Reports 17 (2): 235–48. 10.15252/embr.201540851.

62. Kalluraya, Chinmay A., Alexander J. Weitzel, Brian V. Tsu, and Matthew D. Daugherty. 2023. “Bacterial Origin of a Key Innovation in the Evolution of the Vertebrate Eye.” Proceedings of the National Academy of Sciences 120 (16): e2214815120. 10.1073/pnas.2214815120.

63. Kalyaanamoorthy, Subha, Bui Quang Minh, Thomas K. F. Wong, Arndt von Haeseler, and Lars S. Jermiin. 2017. “ModelFinder: Fast Model Selection for Accurate Phylogenetic Estimates.” Nature Methods 14 (6): 587–89. 10.1038/nmeth.4285.

64. Keeling, Patrick J. 2009. “Functional and Ecological Impacts of Horizontal Gene Transfer in Eukaryotes.” Current Opinion in Genetics & Development, Genomes and evolution, 19 (6): 613–19. 10.1016/j.gde.2009.10.001.

65. Keeling, Patrick J. 2024. “Horizontal Gene Transfer in Eukaryotes: Aligning Theory with Data.” Nature Reviews Genetics 25 (6): 416–30. 10.1038/s41576-023-00688-5.

66. Kempen, Michel van, Stephanie S. Kim, Charlotte Tumescheit, Milot Mirdita, Jeongjae Lee, Cameron L. M. Gilchrist, Johannes Söding, and Martin Steinegger. 2024. “Fast and Accurate Protein Structure Search with Foldseek.” Nature Biotechnology 42 (2): 243–46. 10.1038/s41587-023-01773-0.

67. Kirsch, Roy, Yu Okamura, Wiebke Haeger, Heiko Vogel, Grit Kunert, and Yannick Pauchet. 2022. “Metabolic Novelty Originating from Horizontal Gene Transfer Is Essential for Leaf Beetle Survival.” Proceedings of the National Academy of Sciences 119 (40): e2205857119. 10.1073/pnas.2205857119.

68. Kosakovsky Pond, Sergei L, Art F Y Poon, Ryan Velazquez, Steven Weaver, N Lance Hepler, Ben Murrell, Stephen D Shank, et al. 2019. “HyPhy 2.5—A Customizable Platform for Evolutionary Hypothesis Testing Using Phylogenies.” Molecular Biology and Evolution 37 (1): 295–99. 10.1093/molbev/msz197.

69. Koutsovoulos, Georgios D., Solène Granjeon Noriot, Marc Bailly-Bechet, Etienne G. J. Danchin, and Corinne Rancurel. 2022. “AvP: A Software Package for Automatic Phylogenetic Detection of Candidate Horizontal Gene Transfers.” PLOS Computational Biology 18 (11): e1010686. 10.1371/journal.pcbi.1010686.

70. Kumar, Sudhir, Michael Suleski, Jack M Craig, Adrienne E Kasprowicz, Maxwell Sanderford, Michael Li, Glen Stecher, and S Blair Hedges. 2022. “TimeTree 5: An Expanded Resource for Species Divergence Times.” Molecular Biology and Evolution 39 (8): msac174. 10.1093/molbev/msac174.

71. Kuznetsov, Dmitry, Fredrik Tegenfeldt, Mosè Manni, Mathieu Seppey, Matthew Berkeley, Evgenia V Kriventseva, and Evgeny M Zdobnov. 2022. “OrthoDB V11: Annotation of Orthologs in the Widest Sampling of Organismal Diversity.” Nucleic Acids Research 51 (D1): D445–51. 10.1093/nar/gkac998.

72. Leger, Michelle M., Laura Eme, Courtney W. Stairs, and Andrew J. Roger. 2018. “Demystifying Eukaryote Lateral Gene Transfer (Response to Martin 2017 DOI: 10.1002/Bies.201700115).” BioEssays 40 (5): 1700242. 10.1002/bies.201700242.

73. Lehr, Matthias. 2001. “Phospholipase A2 Inhibitors in Inflammation.” Expert Opinion on Therapeutic Patents 11 (7): 1123–36. 10.1517/13543776.11.7.1123.

74. Leiva-Rebollo, Rocío, Alejandro M. Labella, Juan Gémez-Mata, Dolores Castro, and Juan J. Borrego. 2024. “Fish Iridoviridae: Infection, Vaccination and Immune Response.” Veterinary Research 55 (1): 88. 10.1186/s13567-024-01347-1.

75. Levraud, Jean-Pierre, Luc Jouneau, Valérie Briolat, Valerio Laghi, and Pierre Boudinot. 2019. “IFN-Stimulated Genes in Zebrafish and Humans Define an Ancient Arsenal of Antiviral Immunity.” Journal of Immunology (Baltimore, Md.: 1950) 203 (12): 3361–73. 10.4049/jimmunol.1900804.

76. Li, Yang, Zhiguo Liu, Chao Liu, Zheyi Shi, Lan Pang, Chuzhen Chen, Yun Chen, et al. 2022. “HGT Is Widespread in Insects and Contributes to Male Courtship in Lepidopterans.” Cell 185 (16): 2975–2987.e10. 10.1016/j.cell.2022.06.014.

77. Lima, Carla, Geonildo Rodrigo Disner, Maria Alice Pimentel Falcão, Ana Carolina Seni-Silva, Adolfo Luis Almeida Maleski, Milena Marcolino Souza, Mayara Cristina Reis Tonello, and Monica Lopes-Ferreira. 2021. “The Natterin Proteins Diversity: A Review on Phylogeny, Structure, and Immune Function.” Toxins 13 (8): 538. 10.3390/toxins13080538.

78. Limborg, Morten T., Scott M. Blankenship, Sewall F. Young, Fred M. Utter, Lisa W. Seeb, Mette H. H. Hansen, and James E. Seeb. 2012. “Signatures of Natural Selection among Lineages and Habitats in Oncorhynchus Mykiss.” Ecology and Evolution 2 (1): 1. 10.1002/ece3.59.

79. Liu, Yingying, Haidong Zha, Shanshan Yu, Jiniao Zhong, Xueqin Liu, Hui Yang, and Qian Zhu. 2022. “Molecular Characterization and Antibacterial Activities of a Goose-Type Lysozyme Gene from Roughskin Sculpin (*Trachidermus Fasciatus*).” Fish & Shellfish Immunology 127 (August):1079–87. 10.1016/j.fsi.2022.07.053.

80. Loiseau, Vincent, Jean Peccoud, Clémence Bouzar, Sandra Guillier, Jiangbin Fan, Gianpiero Gueli Alletti, Carine Meignin, et al. 2021. “Monitoring Insect Transposable Elements in Large Double-Stranded DNA Viruses Reveals Host-to-Virus and Virus-to-Virus Transposition.” Molecular Biology and Evolution 38 (9): 3512–30. 10.1093/molbev/msab198.

81. Lorenzo, Pilar, Michael T. Bayliss, and Dick Heinegård. 1998. “A Novel Cartilage Protein (CILP) Present in the Mid-Zone of Human Articular Cartilage Increases with Age*.” Journal of Biological Chemistry 273 (36): 23463–68. 10.1074/jbc.273.36.23463.

82. Ma, Jianchao, Shuanghua Wang, Xiaojing Zhu, Guiling Sun, Guanxiao Chang, Linhong Li, Xiangyang Hu, et al. 2022. “Major Episodes of Horizontal Gene Transfer Drove the Evolution of Land Plants.” Molecular Plant 15 (5): 857–71. 10.1016/j.molp.2022.02.001.

83. Ma, Qing, Tao Yu, Yao-Yao Ren, Ting Gong, and Dian-Sheng Zhong. 2014. “Overexpression of SAMD9 Suppresses Tumorigenesis and Progression during Non Small Cell Lung Cancer.” Biochemical and Biophysical Research Communications 454 (1): 157–61. 10.1016/j.bbrc.2014.10.054.

84. Magalhães, G. S., I. L. M. Junqueira-de-Azevedo, M. Lopes-Ferreira, D. M. Lorenzini, P. L. Ho, and A. M. Moura-da-Silva. 2006. “Transcriptome Analysis of Expressed Sequence Tags from the Venom Glands of the Fish Thalassophryne Nattereri.” Biochimie 88 (6): 693–99. 10.1016/j.biochi.2005.12.008.

85. Maglott, Donna, Jim Ostell, Kim D. Pruitt, and Tatiana Tatusova. 2007. “Entrez Gene: Gene-Centered Information at NCBI.” Nucleic Acids Research 35 (Database issue): D26–31. 10.1093/nar/gkl993.

86. Mahelka, Václav, Karol Krak, David Kopecký, Judith Fehrer, Jan Šafář, Jan Bartoš, Roman Hobza, Nicolas Blavet, and Frank R. Blattner. 2017. “Multiple Horizontal Transfers of Nuclear Ribosomal Genes between Phylogenetically Distinct Grass Lineages.” Proceedings of the National Academy of Sciences 114 (7): 1726–31. 10.1073/pnas.1613375114.

87. Makarova, Kira S., Yuri I. Wolf, Sergey L. Mekhedov, Boris G. Mirkin, and Eugene V. Koonin. 2005. “Ancestral Paralogs and Pseudoparalogs and Their Role in the Emergence of the Eukaryotic Cell.” Nucleic Acids Research 33 (14): 4626. 10.1093/nar/gki775.

88. Manni, Mosè, Matthew R Berkeley, Mathieu Seppey, Felipe A Simão, and Evgeny M Zdobnov. 2021. “BUSCO Update: Novel and Streamlined Workflows along with Broader and Deeper Phylogenetic Coverage for Scoring of Eukaryotic, Prokaryotic, and Viral Genomes.” Molecular Biology and Evolution 38 (10): 4647–54. 10.1093/molbev/msab199.

89. Manoury, Bénédicte, Eric W. Hewitt, Nick Morrice, Pam M. Dando, Alan J. Barrett, and Colin Watts. 1998. “An Asparaginyl Endopeptidase Processes a Microbial Antigen for Class II MHC Presentation.” Nature 396 (6712): 695–99. 10.1038/25379.

90. Marcet-Houben, Marina, and Toni Gabaldón. 2010. “Acquisition of Prokaryotic Genes by Fungal Genomes.” Trends in Genetics: TIG 26 (1): 5–8. 10.1016/j.tig.2009.11.007.

91. Martin, William F. 2017. “Too Much Eukaryote LGT.” BioEssays 39 (12): 1700115. 10.1002/bies.201700115.

92. Martin, William F. 2018. “Eukaryote Lateral Gene Transfer Is Lamarckian.” Nature Ecology & Evolution 2 (5): 754–754. 10.1038/s41559-018-0521-7.

93. Minh, Bui Quang, Heiko A Schmidt, Olga Chernomor, Dominik Schrempf, Michael D Woodhams, Arndt von Haeseler, and Robert Lanfear. 2020. “IQ-TREE 2: New Models and Efficient Methods for Phylogenetic Inference in the Genomic Era.” Molecular Biology and Evolution 37 (5): 1530–34. 10.1093/molbev/msaa015.

94. Mishina, Tappei, Ming-Chung Chiu, Yasuyuki Hashiguchi, Sayumi Oishi, Atsunari Sasaki, Ryuichi Okada, Hironobu Uchiyama, et al. 2023. “Massive Horizontal Gene Transfer and the Evolution of Nematomorph-Driven Behavioral Manipulation of Mantids.” Current Biology 33 (22): 4988–4994.e5. 10.1016/j.cub.2023.09.052.

95. Near, Thomas J., Ron I. Eytan, Alex Dornburg, Kristen L. Kuhn, Jon A. Moore, Matthew P. Davis, Peter C. Wainwright, Matt Friedman, and W. Leo Smith. 2012. “Resolution of Ray-Finned Fish Phylogeny and Timing of Diversification.” Proceedings of the National Academy of Sciences 109 (34): 13698–703. 10.1073/pnas.1206625109.

96. Nounamo, Bernice, Yibo Li, Peter O’Byrne, Aoife M. Kearney, Amir Khan, and Jia Liu. 2017. “An Interaction Domain in Human SAMD9 Is Essential for Myxoma Virus Host-Range Determinant M062 Antagonism of Host Anti-Viral Function.” Virology 503 (January):94. 10.1016/j.virol.2017.01.004.

97. Novembre, John A. 2002. “Accounting for Background Nucleotide Composition When Measuring Codon Usage Bias.” Molecular Biology and Evolution 19 (8): 1390–94. 10.1093/oxfordjournals.molbev.a004201.

98. Paganini, Julien, Amandine Campan-Fournier, Martine Da Rocha, Philippe Gouret, Pierre Pontarotti, Eric Wajnberg, Pierre Abad, and Etienne G. J. Danchin. 2012. “Contribution of Lateral Gene Transfers to the Genome Composition and Parasitic Ability of Root-Knot Nematodes.” PLoS ONE 7 (11): e50875. 10.1371/journal.pone.0050875.

99. Paradis, Emmanuel, and Klaus Schliep. 2019. “Ape 5.0: An Environment for Modern Phylogenetics and Evolutionary Analyses in R.” Bioinformatics 35 (3): 526–28. 10.1093/bioinformatics/bty633.

100. Peng, Shuxia, Xiangzhi Meng, Fushun Zhang, Prabhat Kumar Pathak, Juhi Chaturvedi, Jaime Coronado, Marisol Morales, et al. 2022. “Structure and Function of an Effector Domain in Antiviral Factors and Tumor Suppressors SAMD9 and SAMD9L.” Proceedings of the National Academy of Sciences 119 (4): e2116550119. 10.1073/pnas.2116550119.

101. Peraro, Matteo Dal, and F. Gisou van der Goot. 2016. “Pore-Forming Toxins: Ancient, but Never Really out of Fashion.” Nature Reviews Microbiology 14 (2): 77–92. 10.1038/nrmicro.2015.3.

102. Piégu, Benoît, Sébastien Guizard, Tan Yeping, Corinne Cruaud, Sassan Asgari, Dennis K. Bideshi, Brian A. Federici, and Yves Bigot. 2014. “Genome Sequence of a Crustacean Iridovirus, IIV31, Isolated from the Pill Bug, Armadillidium Vulgare.” Journal of General Virology 95 (7): 1585–90. 10.1099/vir.0.066076-0.

103. Provoost, Pieter, and Samuel Bosch. 2022. “Robis: Ocean Biodiversity Information System (OBIS) Client.” https://github.com/iobis/robis.

104. Pucci, Michela, Nadia Malagolini, and Fabio Dall’Olio. 2021. “Glycosyltransferase B4GALNT2 as a Predictor of Good Prognosis in Colon Cancer: Lessons from Databases.” International Journal of Molecular Sciences 22 (9): 4331. 10.3390/ijms22094331.

105. Pungerčar, Jože, Franck Bihl, Gérard Lambeau, and Igor Križaj. 2021. “What Do Secreted Phospholipases A2 Have to Offer in Combat against Different Viruses up to SARS-CoV-2?” Biochimie 189 (October):40–50. 10.1016/j.biochi.2021.05.017.

106. Qiao, Hongye, Yunyang Wang, Xianjuan Zhang, Ran Lu, Junyun Niu, Fulong Nan, Dingxin Ke, Zhou Zeng, Yashuo Wang, and Bin Wang. 2022. “Cross-Species Opsonic Activity of Zebrafish Fish-Egg Lectin on Mouse Macrophages.” Developmental & Comparative Immunology 129 (April):104332. 10.1016/j.dci.2021.104332.

107. R Core Team. 2018. “R: A Language and Environment for Statistical Computing.” Vienna, Austria: Foundation for Statistical Computing. https://www.r-project.org/.

108. Ravenhall, Matt, Nives Škunca, Florent Lassalle, and Christophe Dessimoz. 2015. “Inferring Horizontal Gene Transfer.” PLOS Computational Biology 11 (5): e1004095. 10.1371/journal.pcbi.1004095.

109. Raymond, James A., and Hak Jun Kim. 2012. “Possible Role of Horizontal Gene Transfer in the Colonization of Sea Ice by Algae.” PLOS ONE 7 (5): e35968. 10.1371/journal.pone.0035968.

110. Rice, P., I. Longden, and A. Bleasby. 2000. “EMBOSS: The European Molecular Biology Open Software Suite.” Trends in Genetics: TIG 16 (6): 276–77. 10.1016/s0168-9525(00)02024-2.

111. Richter, Kathleen, Sukhdev Brar, Madhumita Ray, Prapaporn Pisitkun, Silvia Bolland, Laurent Verkoczy, and Marilyn Diaz. 2009. “Speckled-like Pattern in the Germinal Center (SLIP-GC), a Nuclear GTPase Expressed in Activation-Induced Deaminase-Expressing Lymphomas and Germinal Center B Cells.” The Journal of Biological Chemistry 284 (44): 30652–61. 10.1074/jbc.M109.014506.

112. Richter, Kathleen, Lauranell Burch, Frank Chao, David Henke, Chuancang Jiang, Janssen Daly, Ming-Lang Zhao, Grace Kissling, and Marilyn Diaz. 2012. “Altered Pattern of Immunoglobulin Hypermutation in Mice Deficient in *Slip-GC* Protein*.” Journal of Biological Chemistry 287 (38): 31856–65. 10.1074/jbc.M112.340661.

113. Rincon-Sandoval, Melissa, Rishi De-Kayne, Stephen D. Shank, Stacy Pirro, Alfred Ko’ou, Linelle Abueg, Alan Tracey, et al. 2024. “Ecological Diversification of Sea Catfishes Is Accompanied by Genome-Wide Signatures of Positive Selection.” Nature Communications 15 (1): 10040. 10.1038/s41467-024-54184-3.

114. Roger, Andrew J. 2018. “Reply to ‘Eukaryote Lateral Gene Transfer Is Lamarckian.’” Nature Ecology & Evolution 2 (5): 755–755. 10.1038/s41559-018-0522-6.

115. Sahu, Neha, Boris Indic, Johanna Wong-Bajracharya, Zsolt Merényi, Huei-Mien Ke, Steven Ahrendt, Tori-Lee Monk, et al. 2023. “Vertical and Horizontal Gene Transfer Shaped Plant Colonization and Biomass Degradation in the Fungal Genus Armillaria.” Nature Microbiology 8 (9): 1668–81. 10.1038/s41564-023-01448-1.

116. Salzberg, Steven L. 2017. “Horizontal Gene Transfer Is Not a Hallmark of the Human Genome.” Genome Biology 18 (1): 85. 10.1186/s13059-017-1214-2.

117. Salzberg, Steven L., Owen White, Jeremy Peterson, and Jonathan A. Eisen. 2001. “Microbial Genes in the Human Genome: Lateral Transfer or Gene Loss?” Science 292 (5523): 1903–6. 10.1126/science.1061036.

118. Schilke, Robert M., Cassidy M. R. Blackburn, Temitayo T. Bamgbose, and Matthew D. Woolard. 2020. “Interface of Phospholipase Activity, Immune Cell Function, and Atherosclerosis.” Biomolecules 10 (10): 1449. 10.3390/biom10101449.

119. Schoch, Conrad L., Stacy Ciufo, Mikhail Domrachev, Carol L. Hotton, Sivakumar Kannan, Rogneda Khovanskaya, Detlef Leipe, et al. 2020. “NCBI Taxonomy: A Comprehensive Update on Curation, Resources and Tools.” Database: The Journal of Biological Databases and Curation 2020 (January):baaa062. 10.1093/database/baaa062.

120. Schönknecht, Gerald, Wei-Hua Chen, Chad M. Ternes, Guillaume G. Barbier, Roshan P. Shrestha, Mario Stanke, Andrea Bräutigam, et al. 2013. “Gene Transfer from Bacteria and Archaea Facilitated Evolution of an Extremophilic Eukaryote.” Science 339 (6124): 1207–10. 10.1126/science.1231707.

121. Schönknecht, Gerald, Andreas P. M. Weber, and Martin J. Lercher. 2014. “Horizontal Gene Acquisitions by Eukaryotes as Drivers of Adaptive Evolution.” BioEssays 36 (1): 9–20. 10.1002/bies.201300095.

122. Seni-Silva, Ana Carolina, Adolfo Luis Almeida Maleski, Milena Marcolino Souza, Maria Alice Pimentel Falcao, Geonildo Rodrigo Disner, Monica Lopes-Ferreira, and Carla Lima. 2022. “Natterin-like Depletion by CRISPR/Cas9 Impairs Zebrafish (Danio Rerio) Embryonic Development.” BMC Genomics 23 (1): 123. 10.1186/s12864-022-08369-z.

123. Sharp, Paul M., and Wen-Hsiung Li. 1986. “An Evolutionary Perspective on Synonymous Codon Usage in Unicellular Organisms.” Journal of Molecular Evolution 24 (1): 28–38. 10.1007/BF02099948.

124. Shen, Xing-Xing, Dana A. Opulente, Jacek Kominek, Xiaofan Zhou, Jacob L. Steenwyk, Kelly V. Buh, Max A. B. Haase, et al. 2018. “Tempo and Mode of Genome Evolution in the Budding Yeast Subphylum.” Cell 175 (6): 1533–1545.e20. 10.1016/j.cell.2018.10.023.

125. Slater, Guy St C., and Ewan Birney. 2005. “Automated Generation of Heuristics for Biological Sequence Comparison.” BMC Bioinformatics 6 (1): 31. 10.1186/1471-2105-6-31.

126. Smit, Arian FA, Robert Hubley, and Phil Green. 2013. “RepeatMasker Open-4.0.” http://www.repeatmasker.org.

127. Smith, Martin D., Joel O. Wertheim, Steven Weaver, Ben Murrell, Konrad Scheffler, and Sergei L. Kosakovsky Pond. 2015. “Less Is More: An Adaptive Branch-Site Random Effects Model for Efficient Detection of Episodic Diversifying Selection.” Molecular Biology and Evolution 32 (5): 1342–53. 10.1093/molbev/msv022.

128. Söding, Johannes. 2005. “Protein Homology Detection by HMM–HMM Comparison.” Bioinformatics 21 (7): 951–60. 10.1093/bioinformatics/bti125.

129. Soucy, Shannon M., Jinling Huang, and Johann Peter Gogarten. 2015. “Horizontal Gene Transfer: Building the Web of Life.” Nature Reviews Genetics 16 (8): 472–82. 10.1038/nrg3962.

130. Storer, Jessica, Robert Hubley, Jeb Rosen, Travis J. Wheeler, and Arian F. Smit. 2021. “The Dfam Community Resource of Transposable Element Families, Sequence Models, and Genome Annotations.” Mobile DNA 12 (January):2. 10.1186/s13100-020-00230-y.

131. Sun, Grace Y., Phullara B. Shelat, Michael B. Jensen, Yan He, Albert Y. Sun, and Agnes Simonyi. 2009. “Phospholipases A2 and Inflammatory Responses in the Central Nervous System.” Neuromolecular Medicine 12 (2): 133. 10.1007/s12017-009-8092-z.

132. Szöllősi, Gergely J., Adrián Arellano Davín, Eric Tannier, Vincent Daubin, and Bastien Boussau. 2015. “Genome-Scale Phylogenetic Analysis Finds Extensive Gene Transfer among Fungi.” Philosophical Transactions of the Royal Society B: Biological Sciences 370 (1678): 20140335. 10.1098/rstb.2014.0335.

133. Tarnopol, Rebecca L., Josephine Tamsil, Gyöngyi Cinege, Ji Heon Ha, Kirsten I. Verster, Edit Ábrahám, Lilla B. Magyar, et al. 2024. “Retracing the Horizontal Transfer of a Novel Innate Immune Factor in Drosophila.” bioRxiv. 10.1101/2024.05.29.596511.

134. The UniProt Consortium. 2021. “UniProt: The Universal Protein Knowledgebase in 2021.” Nucleic Acids Research 49 (D1): D480–89. 10.1093/nar/gkaa1100.

135. To, Thu-Hien, Matthieu Jung, Samantha Lycett, and Olivier Gascuel. 2016. “Fast Dating Using Least-Squares Criteria and Algorithms.” Systematic Biology 65 (1): 82–97. 10.1093/sysbio/syv068.

136. Tullio, Vivian, Roberta Spaccapelo, and Manuela Polimeni. 2015. “Lysozymes in the Animal Kingdom.” In Human and Mosquito Lysozymes: Old Molecules for New Approaches Against Malaria, edited by Mauro Prato, 45–57. Cham: Springer International Publishing. 10.1007/978-3-319-09432-8_3.

137. Ullrich, Kristian K. 2024. “MSA2dist: MSA2dist Calculates Pairwise Distances between All Sequences of a DNAStringSet or a AAStringSet Using a Custom Score Matrix and Conducts Codon Based Analysis.” https://gitlab.gwdg.de/mpievolbio-it/MSA2dist.

138. Van Etten, Julia, and Debashish Bhattacharya. 2020. “Horizontal Gene Transfer in Eukaryotes: Not If, but How Much?” Trends in Genetics 36 (12): 915–25. 10.1016/j.tig.2020.08.006.

139. Verma, Pratima, Shraddha Gandhi, Kusum Lata, and Kausik Chattopadhyay. 2021. “Pore-Forming Toxins in Infection and Immunity.” Biochemical Society Transactions 49 (1): 455–65. 10.1042/BST20200836.

140. Vinkler, Michal, Steven R. Fiddaman, Martin Těšický, Emily A. O’Connor, Anna E. Savage, Tobias L. Lenz, Adrian L. Smith, et al. 2023. “Understanding the Evolution of Immune Genes in Jawed Vertebrates.” Journal of Evolutionary Biology 36 (6): 847. 10.1111/jeb.14181.

141. Wang, Yashuo, Lingzhen Bu, Lili Yang, Hongyan Li, and Shicui Zhang. 2016. “Identification and Functional Characterization of Fish-Egg Lectin in Zebrafish.” Fish & Shellfish Immunology 52 (May):23–30. 10.1016/j.fsi.2016.03.016.

142. Waterhouse, Robert M., Evgeny M. Zdobnov, and Evgenia V. Kriventseva. 2010. “Correlating Traits of Gene Retention, Sequence Divergence, Duplicability and Essentiality in Vertebrates, Arthropods, and Fungi.” Genome Biology and Evolution 3 (December):75. 10.1093/gbe/evq083.

143. Wertheim, Joel O., Ben Murrell, Martin D. Smith, Sergei L. Kosakovsky Pond, and Konrad Scheffler. 2015. “RELAX: Detecting Relaxed Selection in a Phylogenetic Framework.” Molecular Biology and Evolution 32 (3): 820–32. 10.1093/molbev/msu400.

144. Willerslev, Eske, Tobias Mourier, Anders J. Hansen, Bent Christensen, Ian Barnes, and Steven L. Salzberg. 2002. “Contamination in the Draft of the Human Genome Masquerades As Lateral Gene Transfer.” DNA Sequence 13 (2): 75–76. 10.1080/10425170290023392.

145. Wilson, Christopher G., Tymoteusz Pieszko, Reuben W. Nowell, and Timothy G. Barraclough. 2024. “Recombination in Bdelloid Rotifer Genomes: Asexuality, Transfer and Stress.” Trends in Genetics 40 (5): 422–36. 10.1016/j.tig.2024.02.001.

146. Wybouw, N., T. Van Leeuwen, and W. Dermauw. 2018. “A Massive Incorporation of Microbial Genes into the Genome of Tetranychus Urticae, a Polyphagous Arthropod Herbivore.” Insect Molecular Biology 27 (3): 333–51. 10.1111/imb.12374.

147. Wybouw, Nicky, Yannick Pauchet, David G. Heckel, and Thomas Van Leeuwen. 2016. “Horizontal Gene Transfer Contributes to the Evolution of Arthropod Herbivory.” Genome Biology and Evolution 8 (6): 1785–1801. 10.1093/gbe/evw119.

148. Xia, Jixing, Zhaojiang Guo, Zezhong Yang, Haolin Han, Shaoli Wang, Haifeng Xu, Xin Yang, et al. 2021. “Whitefly Hijacks a Plant Detoxification Gene That Neutralizes Plant Toxins.” Cell 184 (7): 1693–1705.e17. 10.1016/j.cell.2021.02.014.

149. Xiang, Yang, Chao Yan, Xiaolong Guo, Kaifeng Zhou, Sheng’an Li, Qian Gao, Xuan Wang, et al. 2014. “Host-Derived, Pore-Forming Toxin–like Protein and Trefoil Factor Complex Protects the Host against Microbial Infection.” Proceedings of the National Academy of Sciences 111 (18): 6702–7. 10.1073/pnas.1321317111.

150. Xiao, Jun, Huan Zhong, Zhen Liu, Fan Yu, Yongju Luo, Xi Gan, and Yi Zhou. 2015. “Transcriptome Analysis Revealed Positive Selection of Immune-Related Genes in Tilapia.” Fish & Shellfish Immunology 44 (1): 60–65. 10.1016/j.fsi.2015.01.022.

151. Yang, Zhenzhen, Yeting Zhang, Eric K. Wafula, Loren A. Honaas, Paula E. Ralph, Sam Jones, Christopher R. Clarke, et al. 2016. “Horizontal Gene Transfer Is More Frequent with Increased Heterotrophy and Contributes to Parasite Adaptation.” Proceedings of the National Academy of Sciences 113 (45): E7010–19. 10.1073/pnas.1608765113.

152. Yoxsimer, Alyssa M., Emma G. Okenberg, Austin Wolfgang Katzer, Michael A. Bell, Robert L. Massengill, and David M. Kingsley. 2024. “Genomic Sequence of the Threespine Stickleback Iridovirus (TSIV) from Wild Gasterosteus Aculeatus in Stormy Lake, Alaska.” Viruses 16 (11): 1663. 10.3390/v16111663.

153. Yu, Guangchuang, David K. Smith, Huachen Zhu, Yi Guan, and Tommy Tsan-Yuk Lam. 2017. “Ggtree: An r Package for Visualization and Annotation of Phylogenetic Trees with Their Covariates and Other Associated Data.” Methods in Ecology and Evolution 8 (1): 28–36. 10.1111/2041-210X.12628.

154. Yuan, Le, Hongzhong Lu, Feiran Li, Jens Nielsen, and Eduard J. Kerkhoven. 2023. “HGTphyloDetect: Facilitating the Identification and Phylogenetic Analysis of Horizontal Gene Transfer.” Briefings in Bioinformatics 24 (2): bbad035. 10.1093/bib/bbad035.

155. Zhang, Chao, Maryam Rabiee, Erfan Sayyari, and Siavash Mirarab. 2018. “ASTRAL-III: Polynomial Time Species Tree Reconstruction from Partially Resolved Gene Trees.” BMC Bioinformatics 19 (6): 153. 10.1186/s12859-018-2129-y.

156. Zhang, Hua-Hao, Jean Peccoud, Min-Rui-Xuan Xu, Xiao-Gu Zhang, and Clément Gilbert. 2020. “Horizontal Transfer and Evolution of Transposable Elements in Vertebrates.” Nature Communications 11 (1): 1362. 10.1038/s41467-020-15149-4.

157. Zhang, Kai, Xiaobing Liu, Xuemei Li, Yuxiang Liu, Haiyang Yu, Jinxiang Liu, and Quanqi Zhang. 2020. “Antibacterial Functions of a Novel Fish-Egg Lectin from Spotted Knifejaw (*Oplegnathus Punctatus*) during Host Defense Immune Responses.” Developmental & Comparative Immunology 111 (October):103758. 10.1016/j.dci.2020.103758.

158. Zhang, Kun, Yiheng Wang, Yue Zhang, and Xiaofei Shan. 2024. “Codon Usage Characterization and Phylogenetic Analysis of the Mitochondrial Genome in Hemerocallis Citrina.” BMC Genomic Data 25 (January):6. 10.1186/s12863-024-01191-4.

159. Zimmermann, Lukas, Andrew Stephens, Seung-Zin Nam, David Rau, Jonas Kübler, Marko Lozajic, Felix Gabler, Johannes Söding, Andrei N. Lupas, and Vikram Alva. 2018. “A Completely Reimplemented MPI Bioinformatics Toolkit with a New HHpred Server at Its Core.” Journal of Molecular Biology, Computation Resources for Molecular Biology, 430 (15): 2237–43. 10.1016/j.jmb.2017.12.007.

