## Supplementary Figures for "Strong evidence for multiple horizontal transfers of immune genes between teleost fishes"

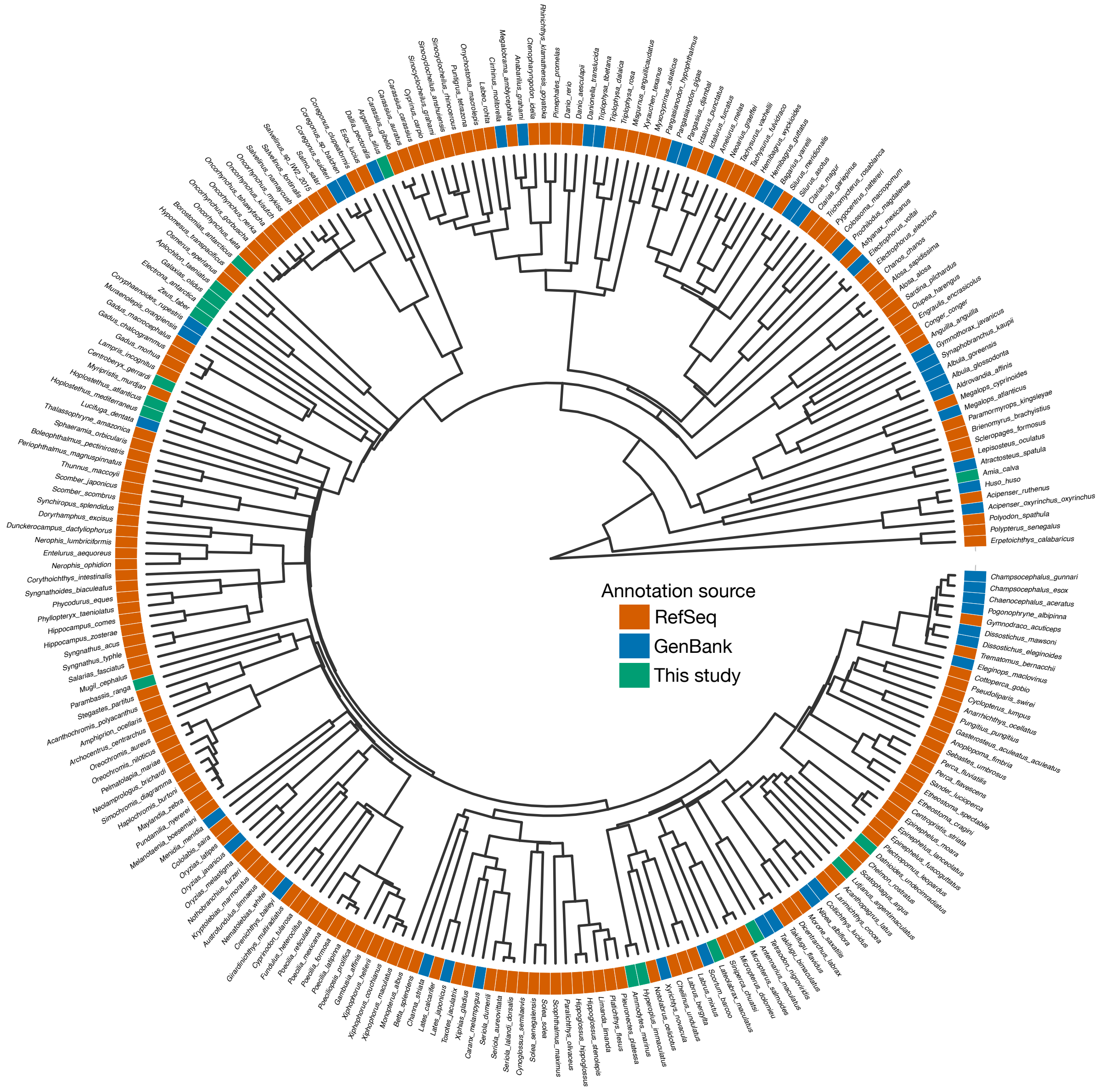


**Supplementary figure 1**. Annotated species of the dataset. Phylogeny of ray-finned fishes, computed from 3,584 BUSCO genes using ASTRAL-III, and dated using IQ-TREE2 and the least square dating method. For each species, the annotation source is indicated: (i) "RefSeq" correspond to NCBI RefSeq annotations, (ii) "GenBank" correspond to user submitted annotation files in GenBank and (iii) "This study" correspond to genome annotated for this study, using BRAKER2 or BRAKER3.

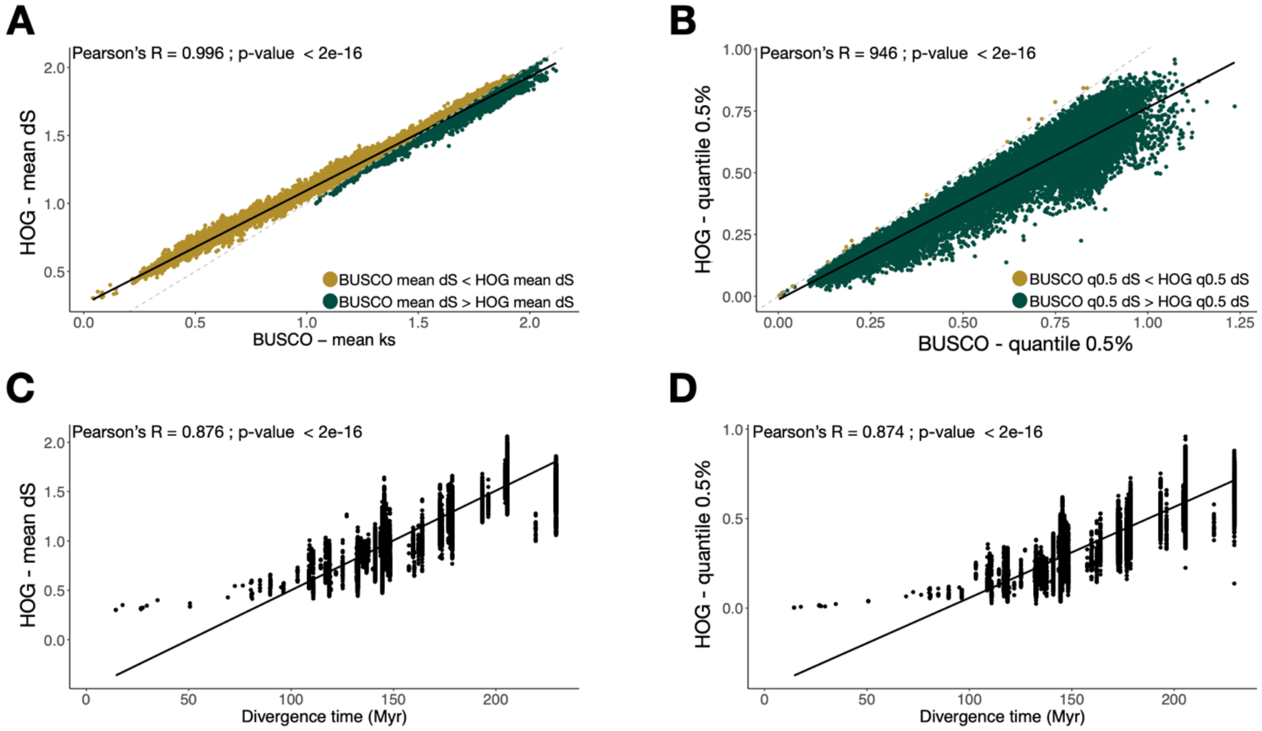


**Supplementary figure 2**. dS statistics. Pearson correlations of A- the mean and B- the 0.5% quantile of dS values, computed for every pair of species, between all the genes (every HOG) and every BUSCO groups. The dashed line corresponds to X=Y. Pearson correlations between A- the mean dS and B- the 0.5% quantile of dS values and the divergence time between species.

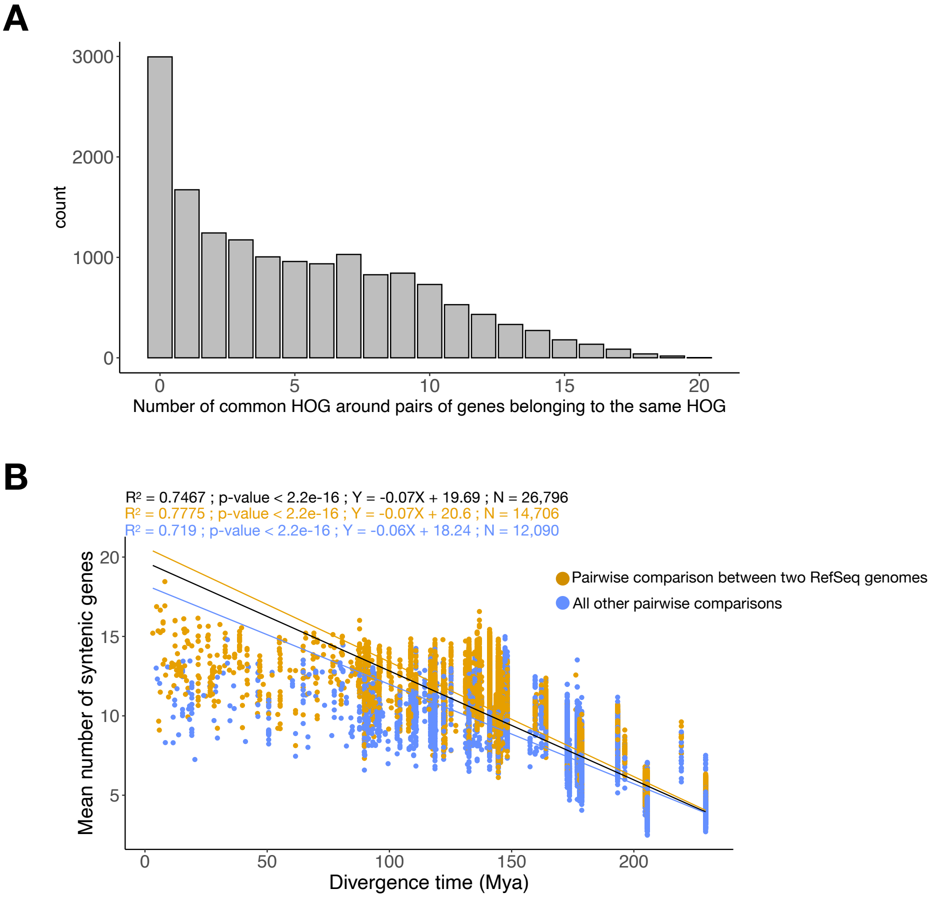


**C**

| Response | Predictor(s) | Adjusted $R^2$ | p-value | AIC |
|---|---|---|---|---|
| Mean syntenic score | Divergence time | 0.7467 | < 2.2e-16 | 100467.00 |
| Mean syntenic score | Divergence time + Annotation method | 0.7585 | < 2.2e-16 | 99195.59 |

**Supplementary figure 3**. Micro-synteny analysis. A- Example of micro-syntenic scores, computed between each pair of genes belonging to the same HOG, between two species. B- Relationship between the mean micro-syntenic scores (computed for every pair of species) and the divergence time. To have similar comparisons between pairs of species, we only kept pairs for which the micro-syntenic scores were computed on at least 100 genes (first discarding genes for which there were less than 10 genes upstream or downstream). Three linear regressions were computed: (i) on the whole dataset (black), (ii) only keeping species pairs which both had RefSeq annotations (orange) and (iii) discarding species pairs which both had RefSeq annotations (blue). C- Multiple linear regression result. The simple model (mean micro-syntenic score ~ divergence time) was compared to the multiple model (mean micro-syntenic score ~ divergence time + annotation method). The annotation method was either “RefSeq”, if both species had RefSeq annotations, or “other” for every other species pairs. The multiple model had a higher adjusted $R^2$ and a much lower AIC, with a tendency of RefSeq comparisons to have higher mean micro-syntenic scores.

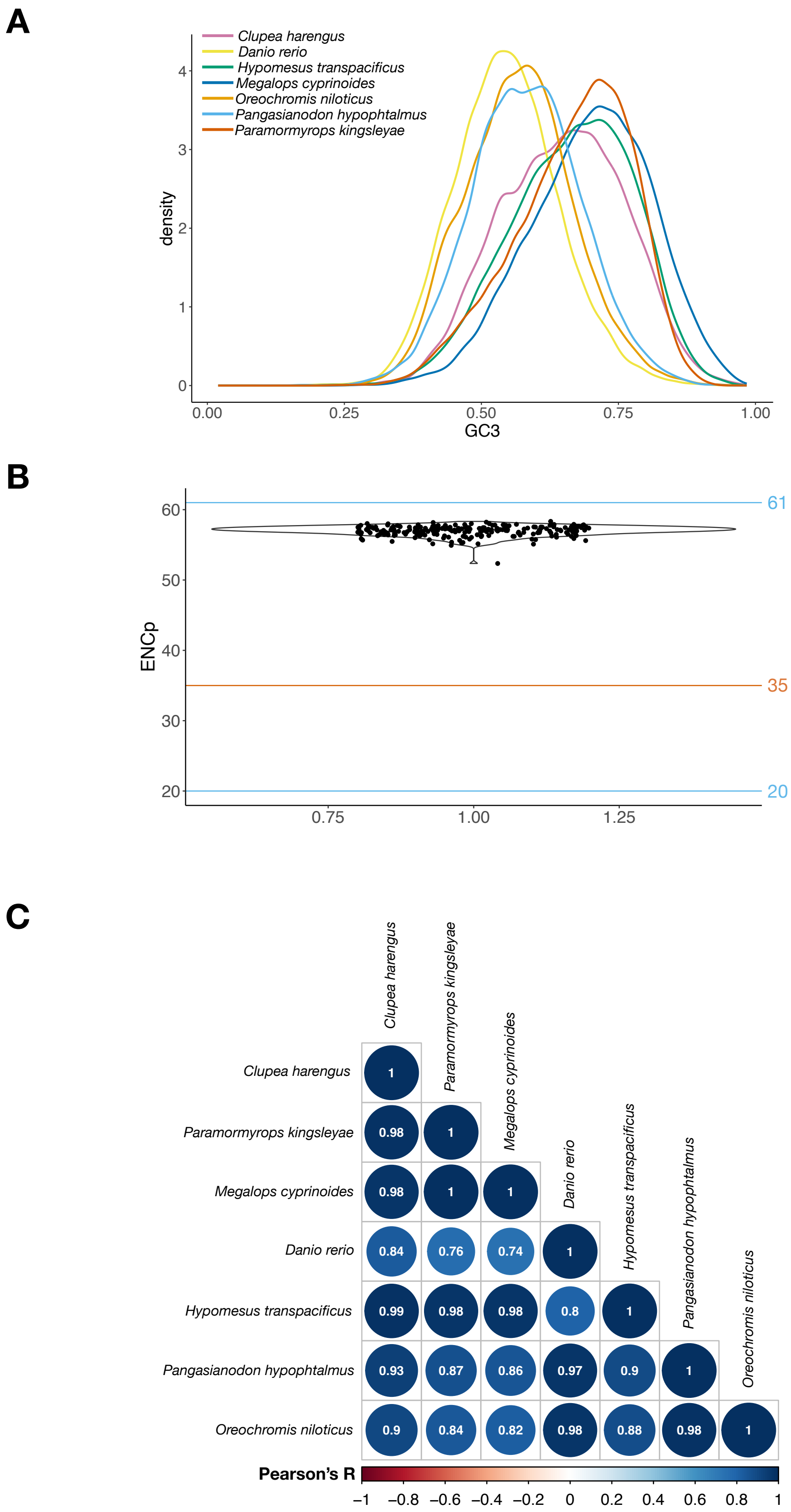


**Supplementary figure 4**. Gene sequences composition. A- Distribution of the GC3 proportion per coding sequence (CDS), for all the CDS of seven phylogenetically representative species. B- Mean ENCprime for each teleost species in the dataset, computed over every coding sequences of the genome. C- Pairwise Pearson correlation coefficients of RSCU profiles between seven phylogenetically representative species.

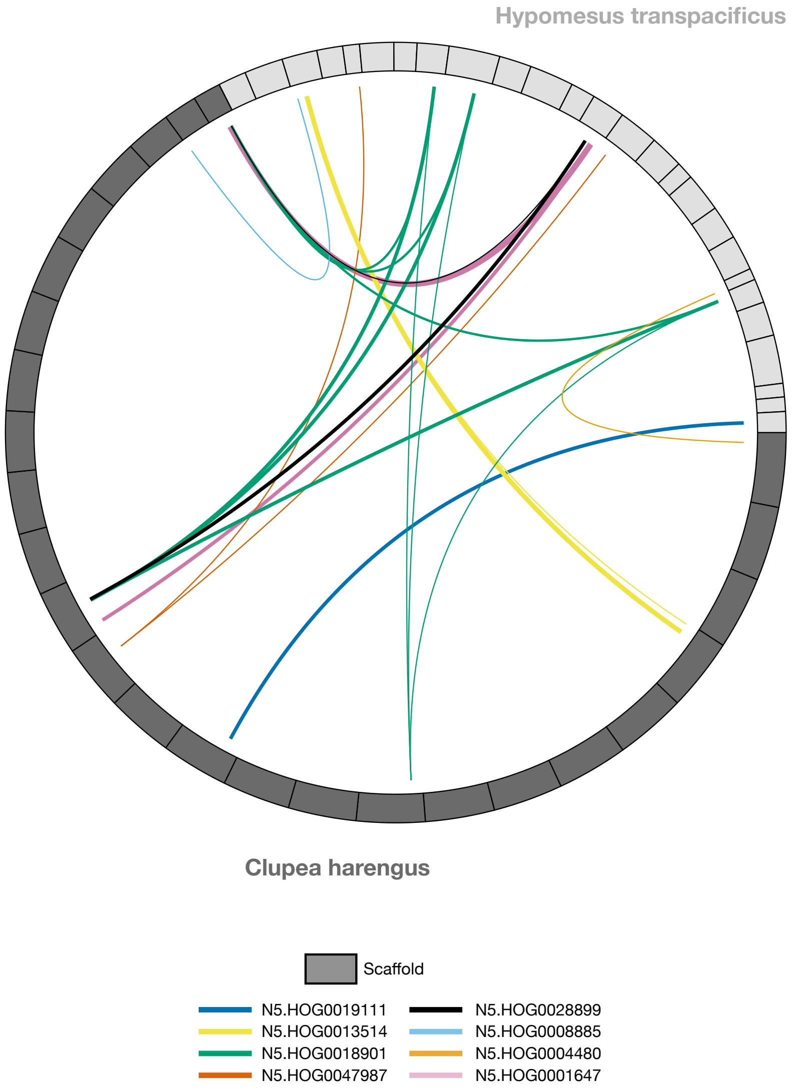


**Supplementary figure 5**. Location of transferred genes in *C. harengus* and *H. transpacificus*. The location of transferred genes in the genome of these two species are represented in a circos plot, with links between each gene belonging to the same HOG. Multiple links for the same HOG indicate species specific duplications.

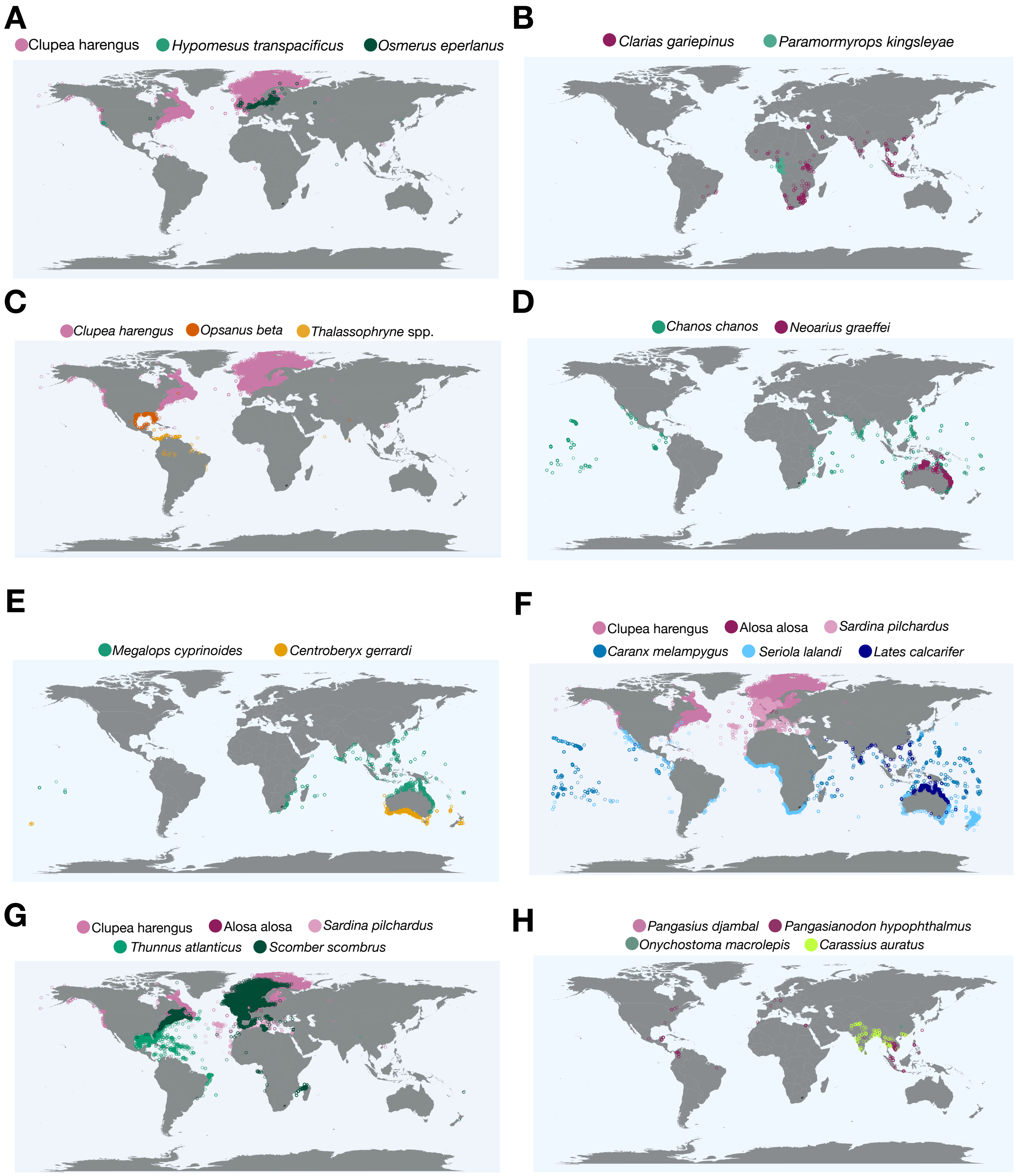


**Supplementary figure 6. Overlapping distributions of species involved in HGTs.** Geographical distributions of donor and recipient species for A- HOG0047987, HOG0001647, HOG0013514, HOG0018901, HOG0008885, HOG0004480, HOG0028899, HOG0019111, B- HOG0029633, HOG0046344, HOG0010622, C- HOG0013500, D- HOG0019032, E- HOG0004451, F- HOG0004450, G- HOG0019111 and H-HOG0030200, HOG0034670.

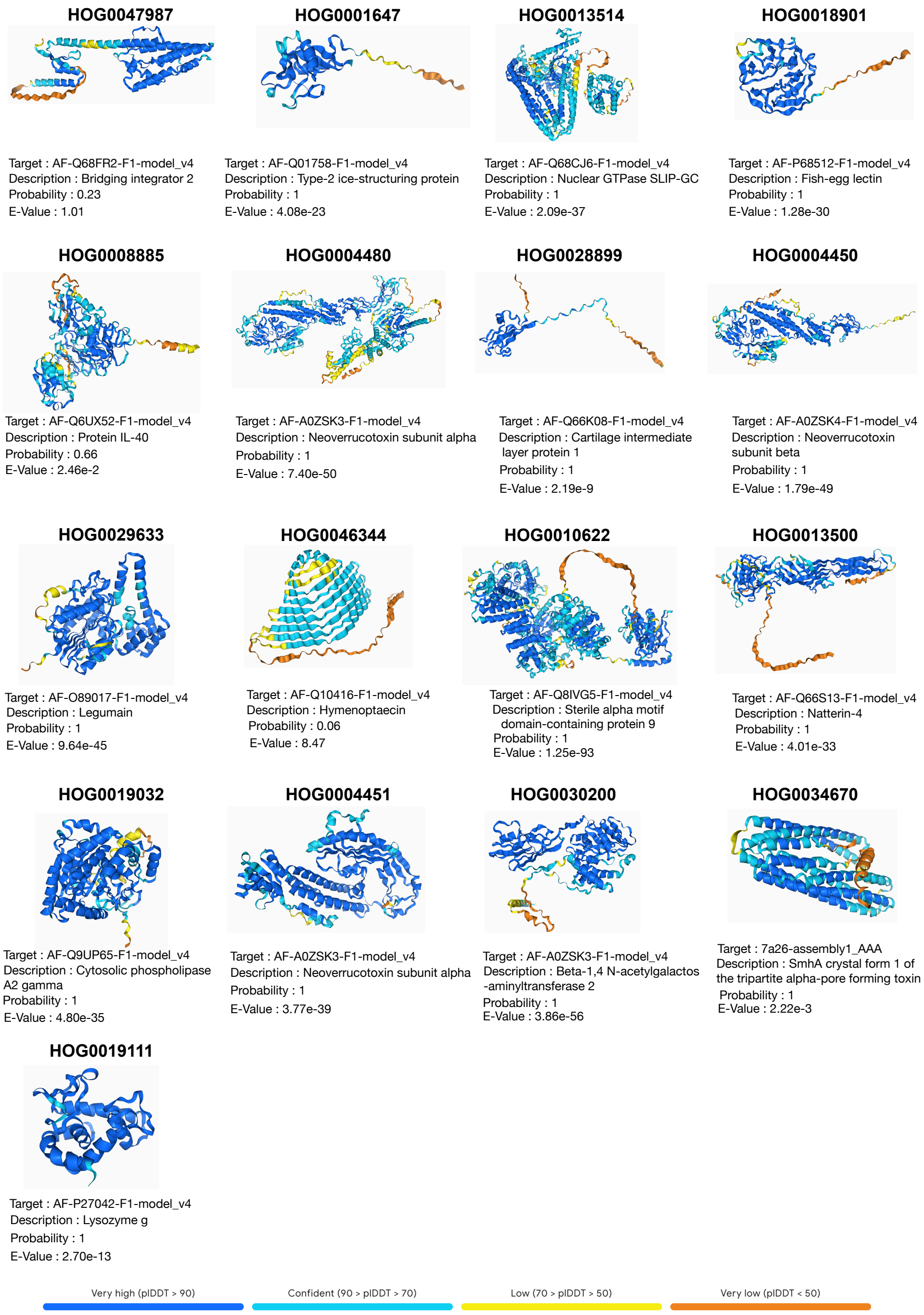


**Supplementary figure 7**. Structural similarities results. For each HGT, the 3D structure of one representative protein sequence was predicted using AlphaFold3. HOG0046344 : XM_023822671 ; HOG0010622 : XM_023831520 ; HOG0047987 : XM_047020147 ; HOG0013514 : XM_047045767 ; HOG0018901 : XM_047023826 ; HOG0008885 : XM_047048755 ; HOG0004480 : XM_062453720 ; HOG0028899 : XM_047031800 ; HOG0004450 : XM_042704194 ; HOG0029633 : XM_046853148 ; HOG0013500 : XM_034166331 ; HOG0019032 : XM_060938829 ; HOG0004451 : XM_036520634 ; HOG0030200 : XM_026945431 ; HOG0034670 : XM_053236679 ; HOG0019111 : XM_047049897 ; HOG0001647 : XM_047032478. These 3D structures were used as input in a structural similarity search using FoldSeek. FoldSeek best targets are indicated, as well as their descriptions, probabilities and e-values.

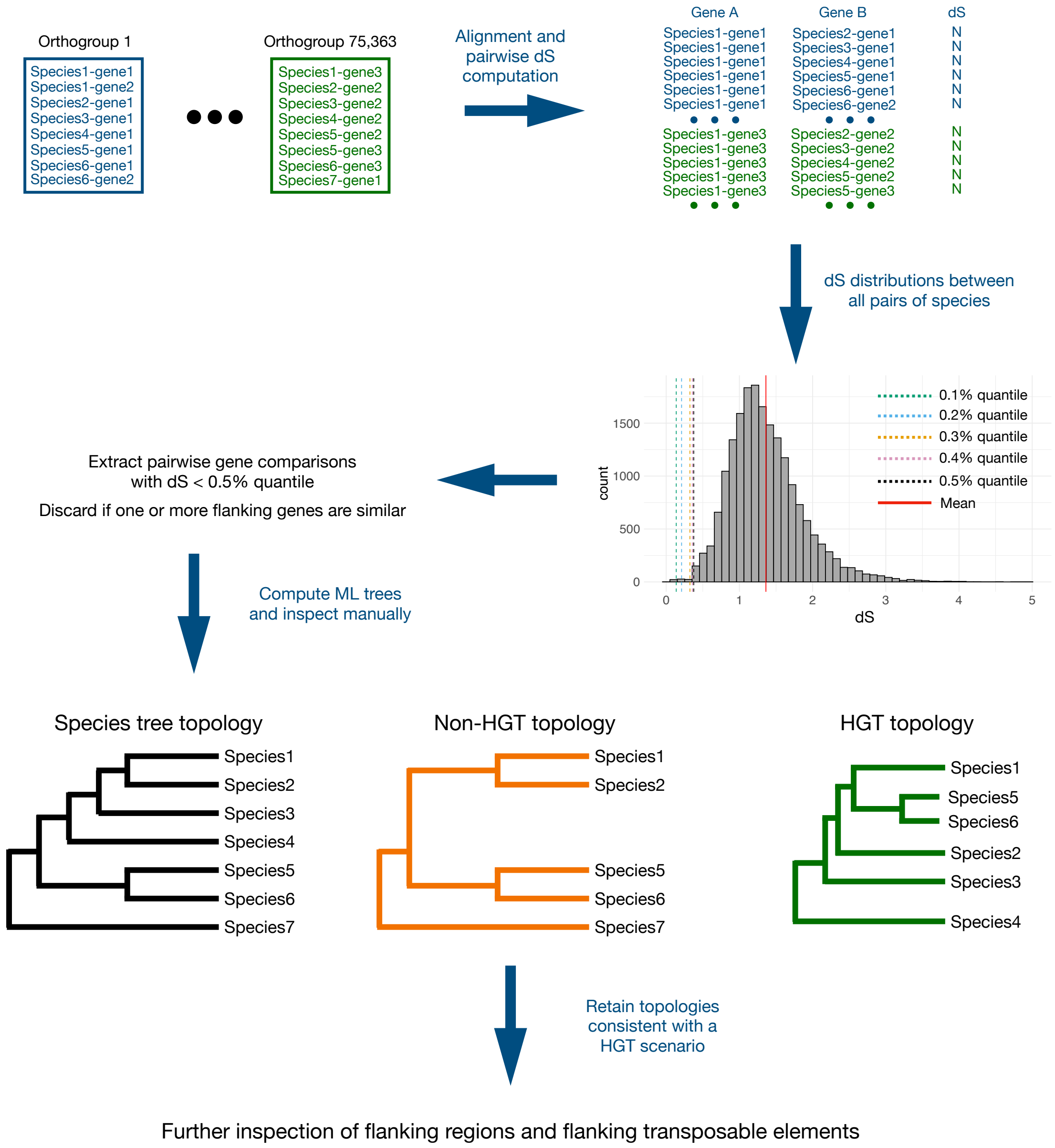


**Supplementary figure 8**. Overview of the dS workflow to detect horizontal gene transfers between teleosts.

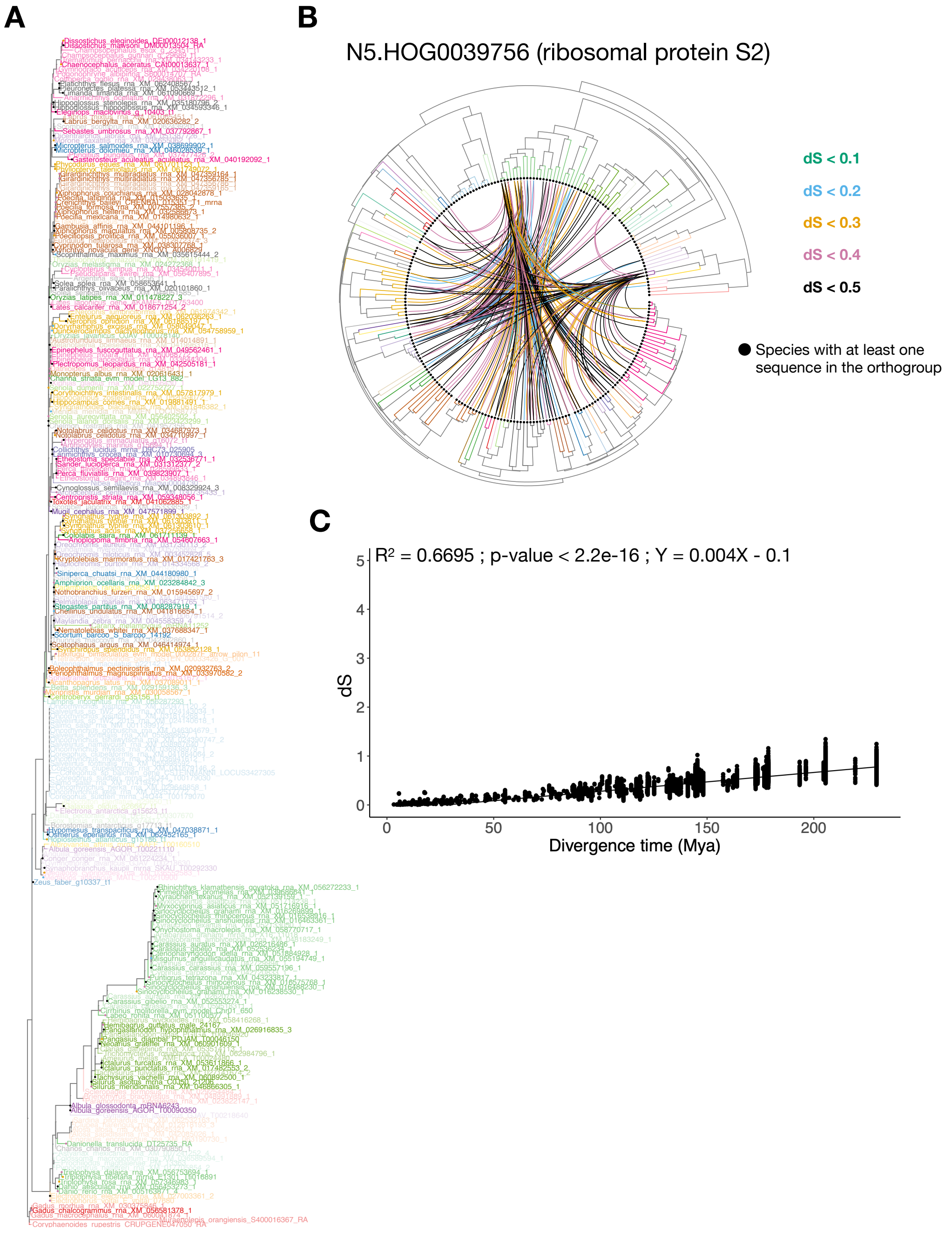


**Supplementary figure 9**. Example of a gene with a ladder-shaped tree topology and likely under purifying selection on synonymous sites. A- Phylogenetic tree of HOG0039756. Terminal branches with a black dot represent sequences which pass the dS threshold (q0.5) with at least one other sequence of the HOG. B- Teleost species tree showing the pair of species with at least one gene pair below the dS threshold. C- Relationship between pairwise dS values and the divergence time. The dS between every pair of genes is extremely low, even for high divergence times.

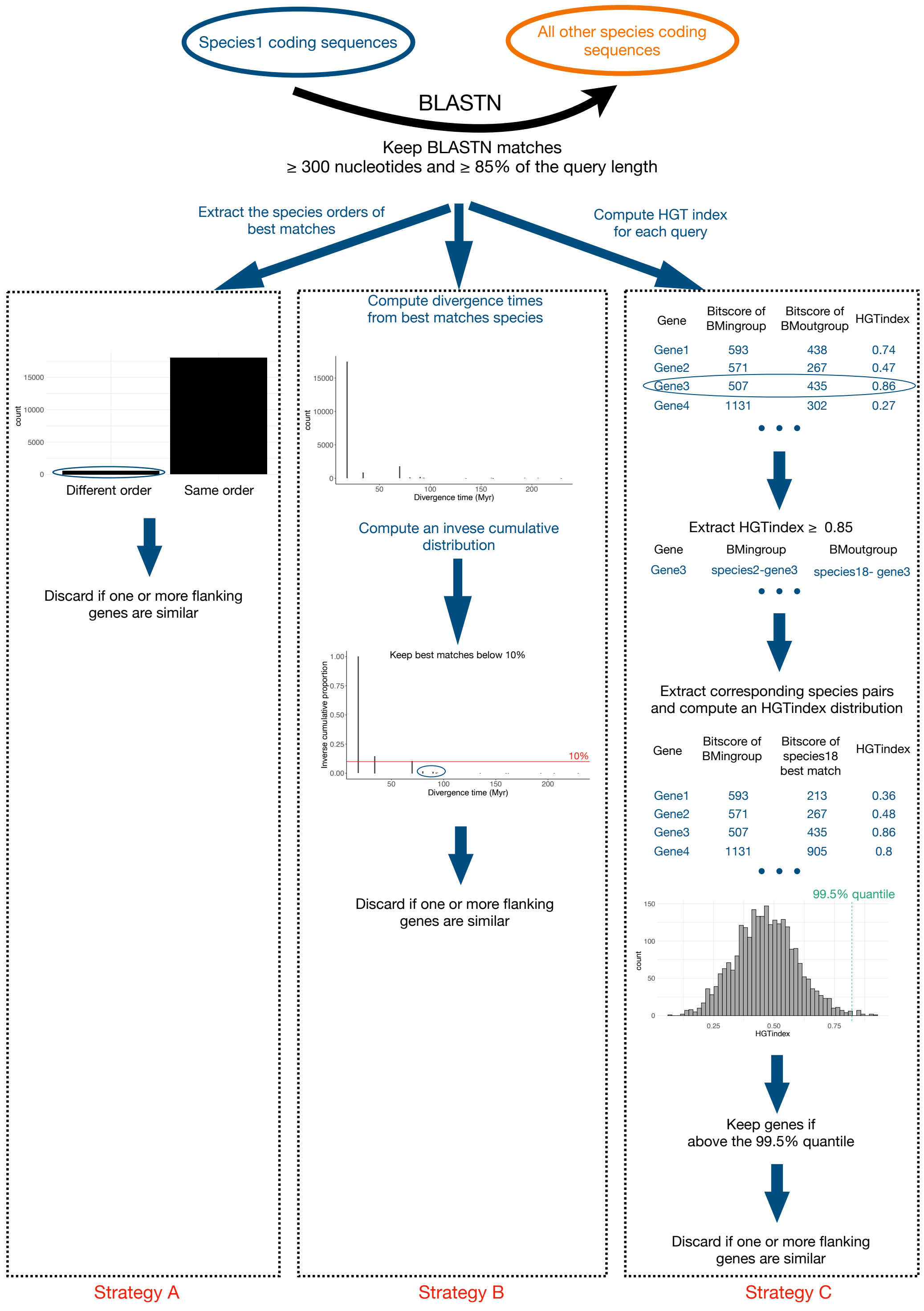


**Supplementary figure 10**. Overview of the three BLAST workflows to detect horizontal gene transfers between teleosts.
