## Supplementary File 1 for "Strong evidence for multiple horizontal transfers of immune genes between teleost fishes"

N5.HOG0047987

N5.HOG0047987

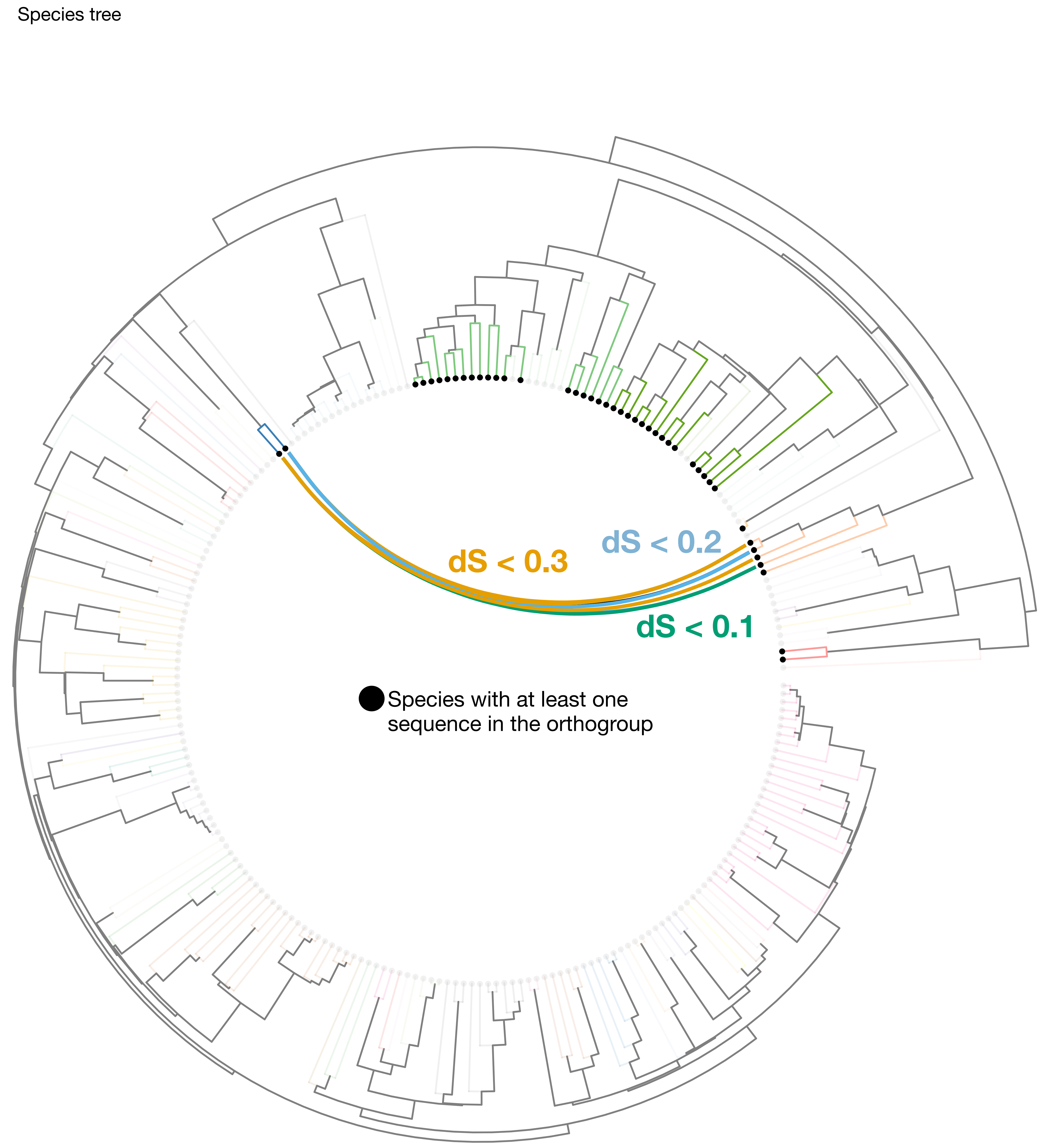

### N5.HOG0047987

Orthogroup phylogeny and protein alignment

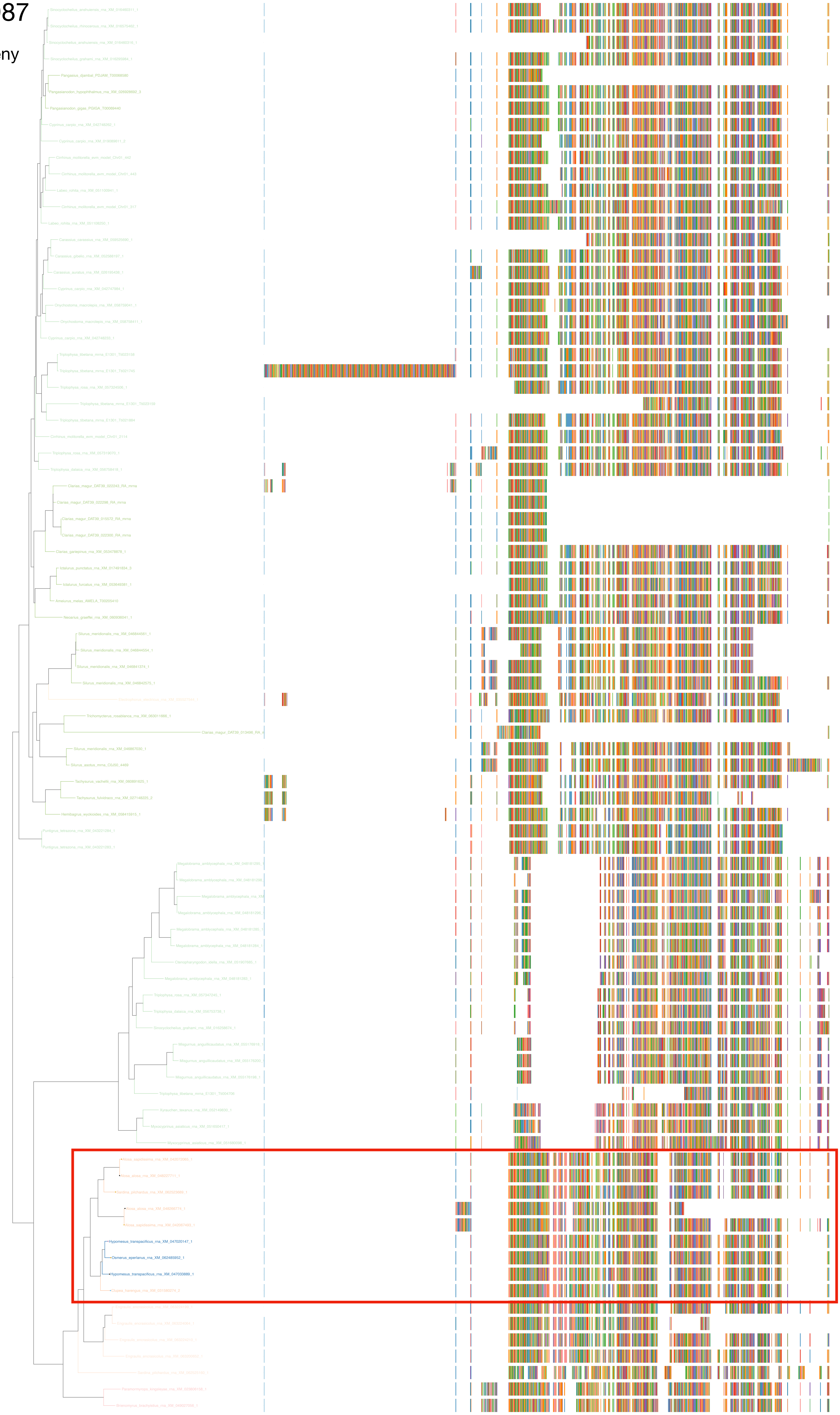

N5.HOG0047987

Zoom on HGT clade with bootstrap values

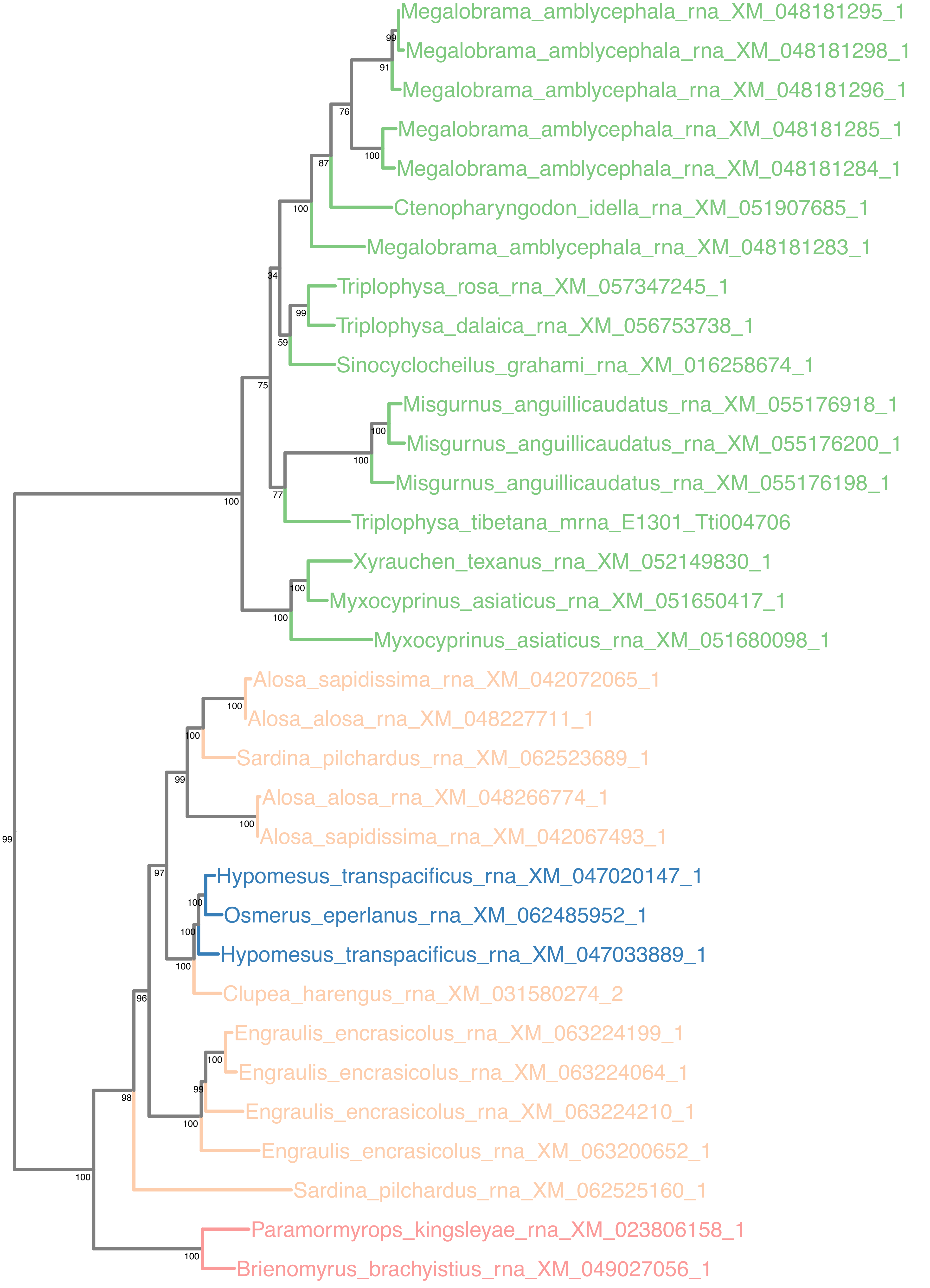

### N5.HOG0047987

Orthogroup phylogeny and trimmed protein alignment

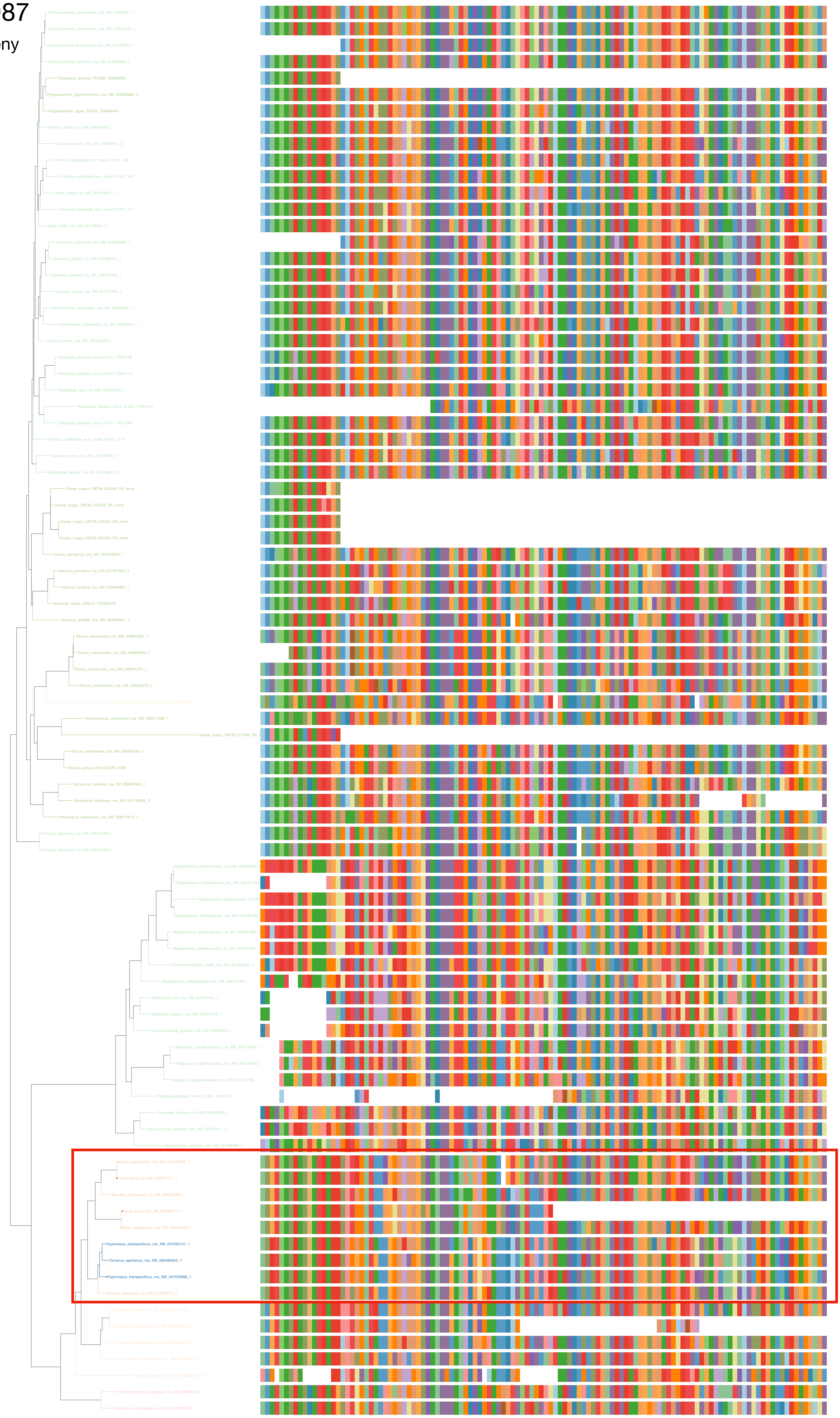

### N5.HOG0047987

« Best-match » phylogeny
and
protein alignment

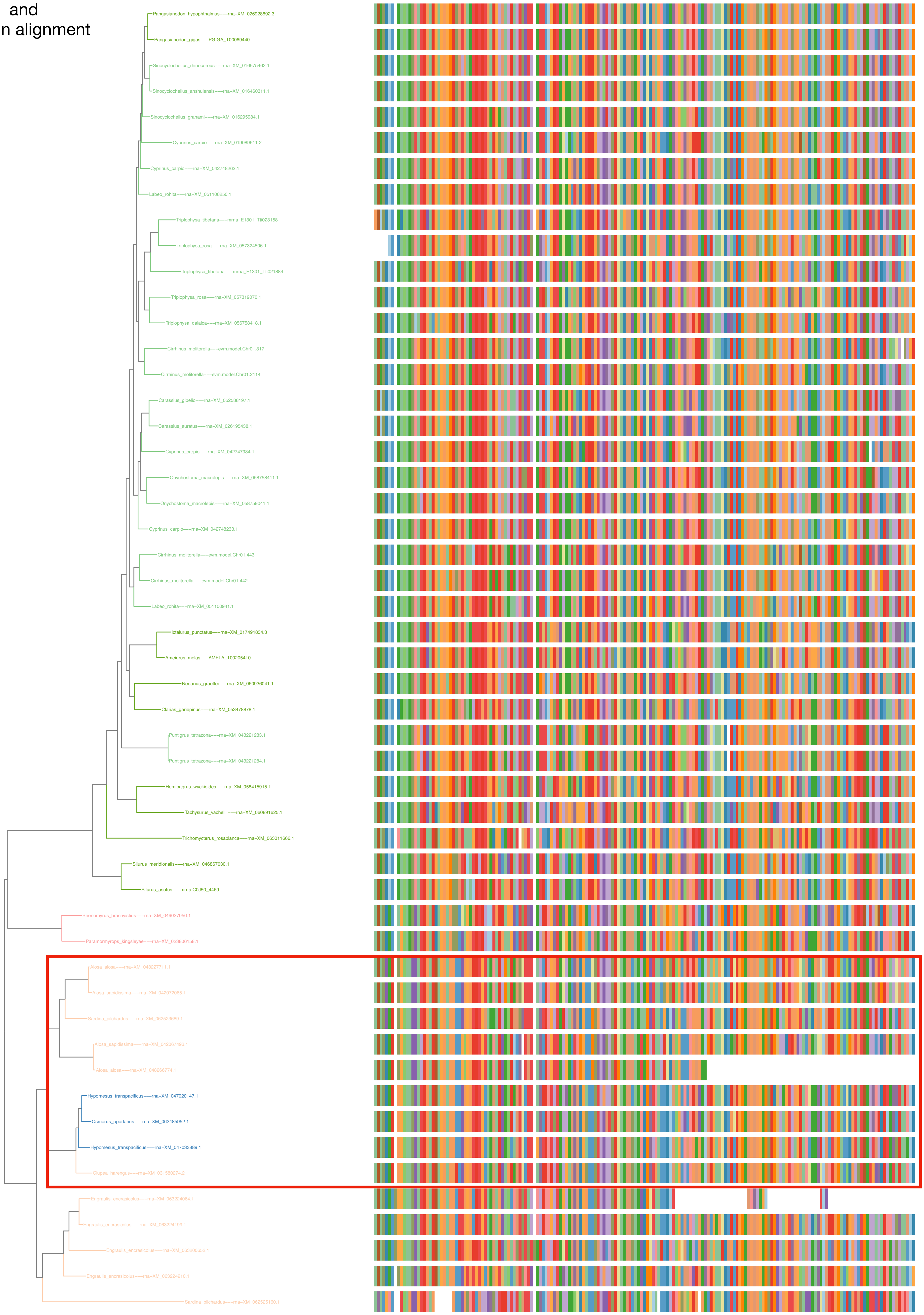

N5.HOG0047987

dS distribution

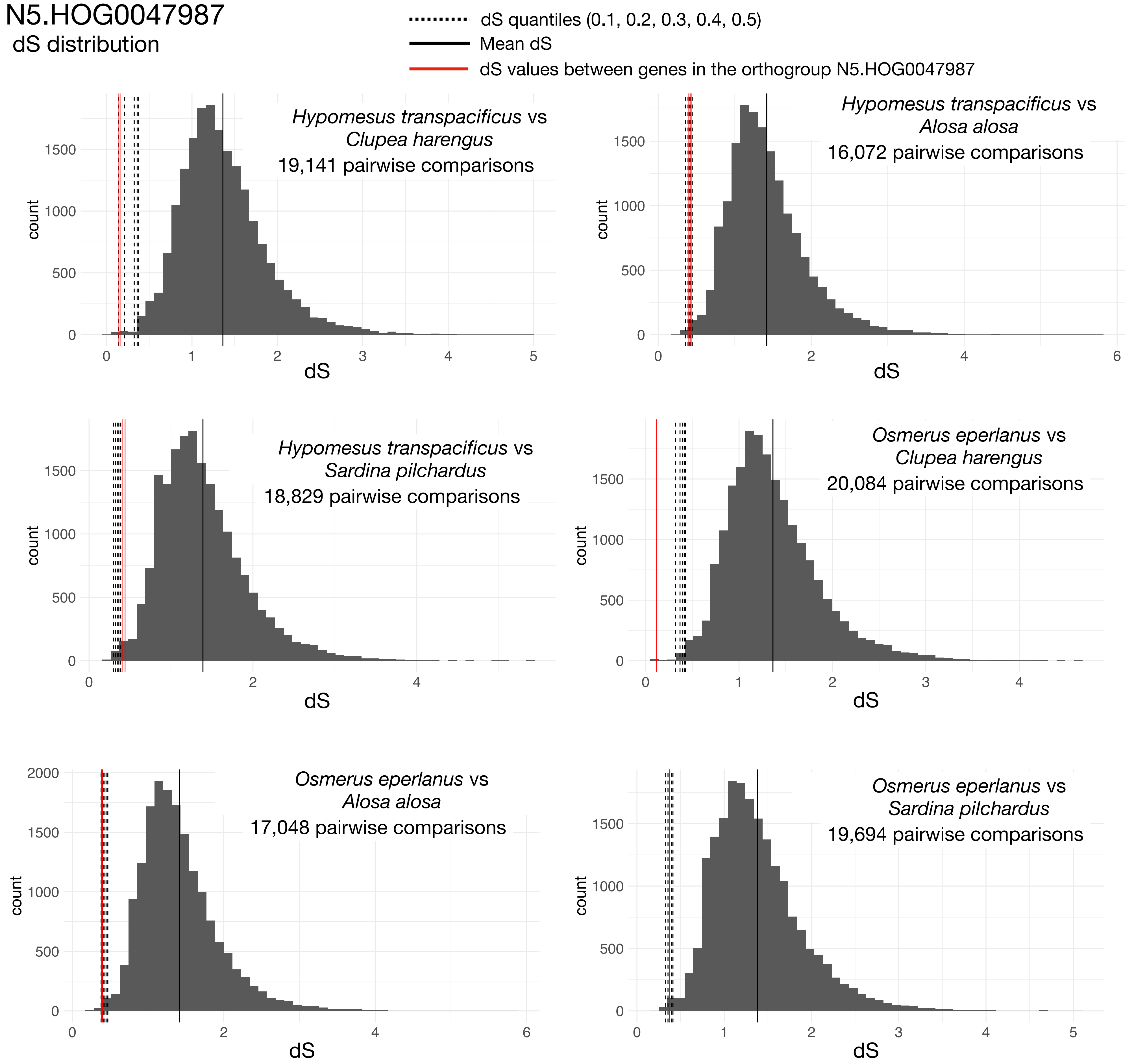


Intra-orthogroup dS

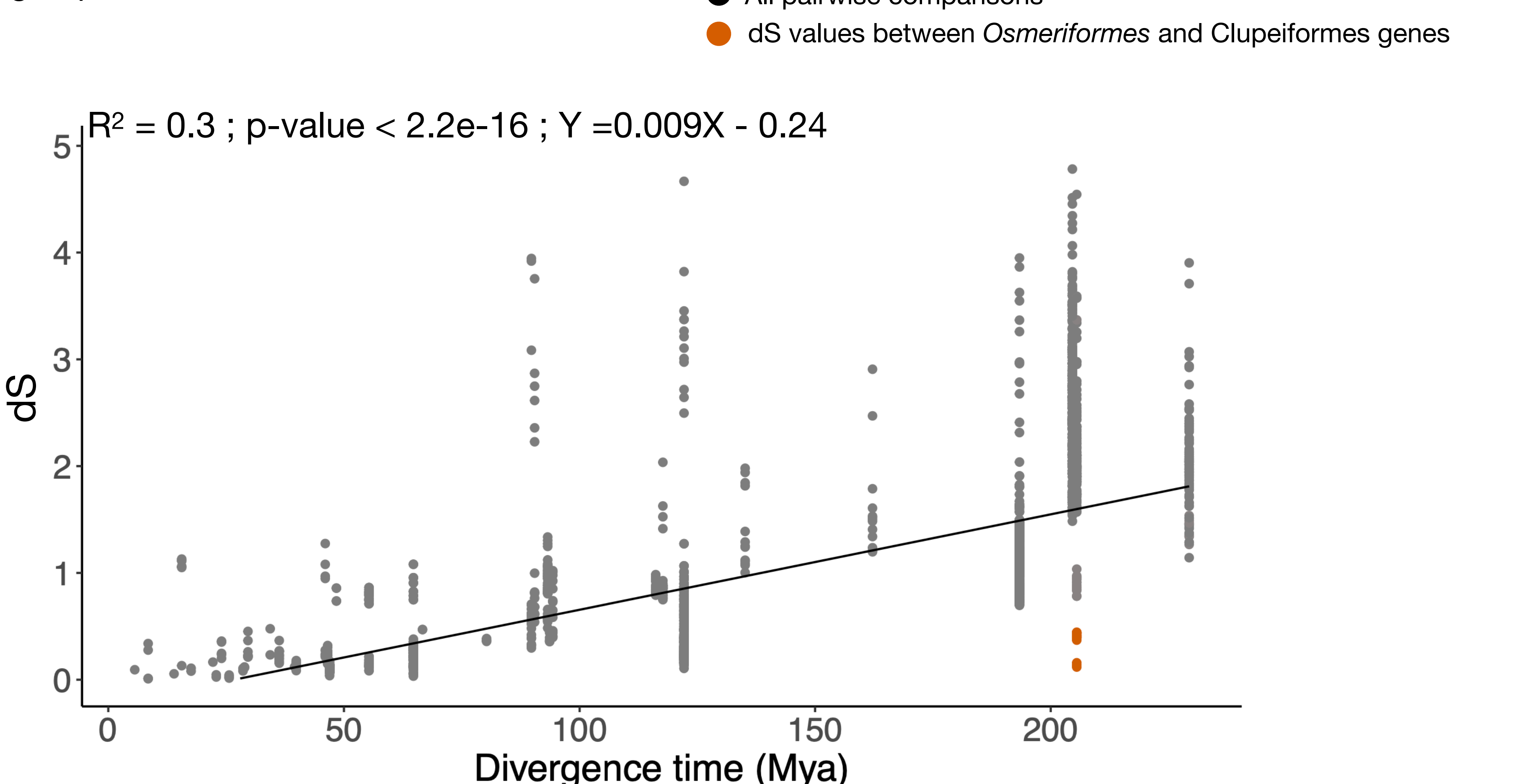

### N5.HOG0047987

#### Micro-synteny

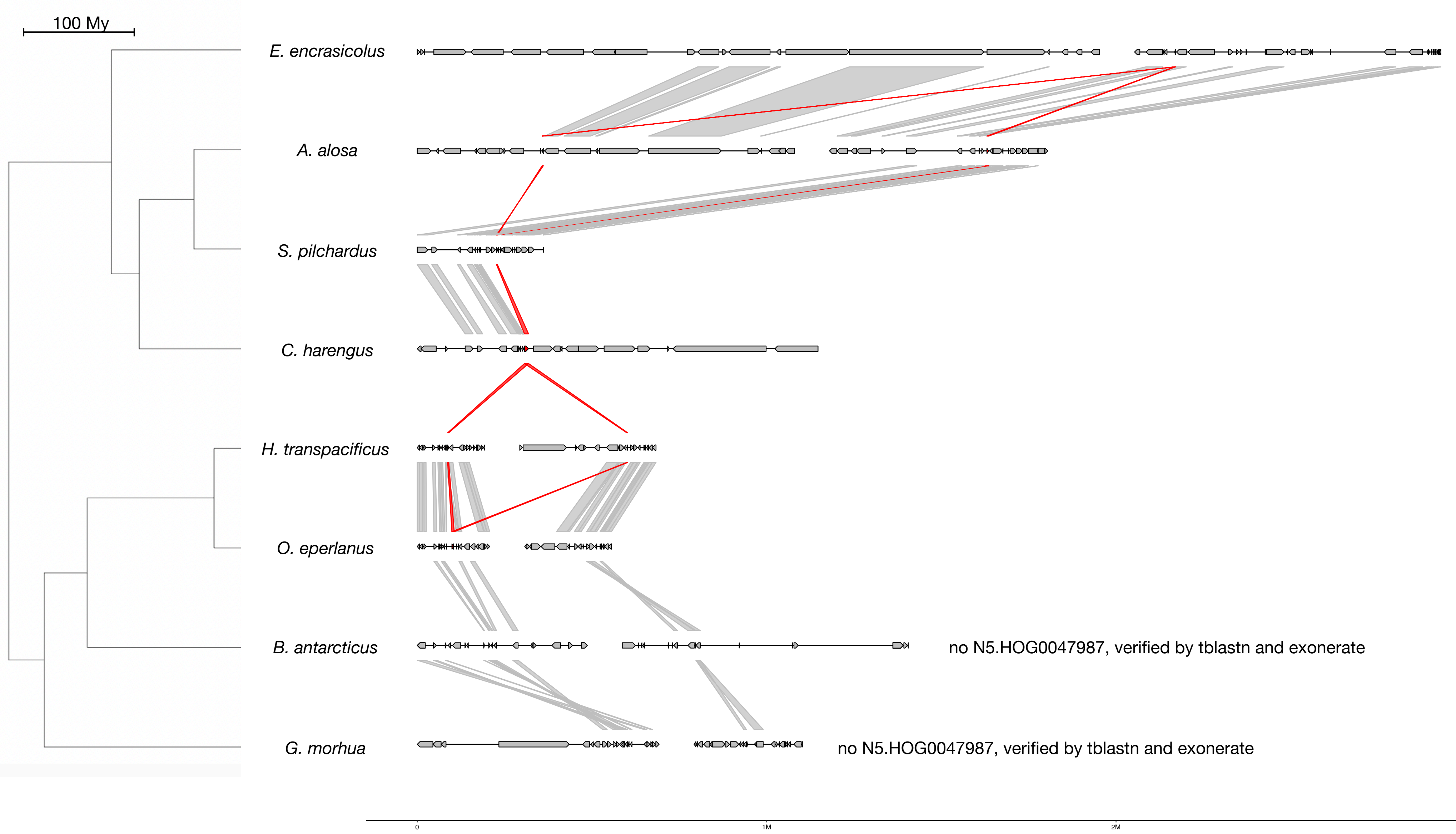


#### Zoom on *C. harengus* and *H. transpacificus*

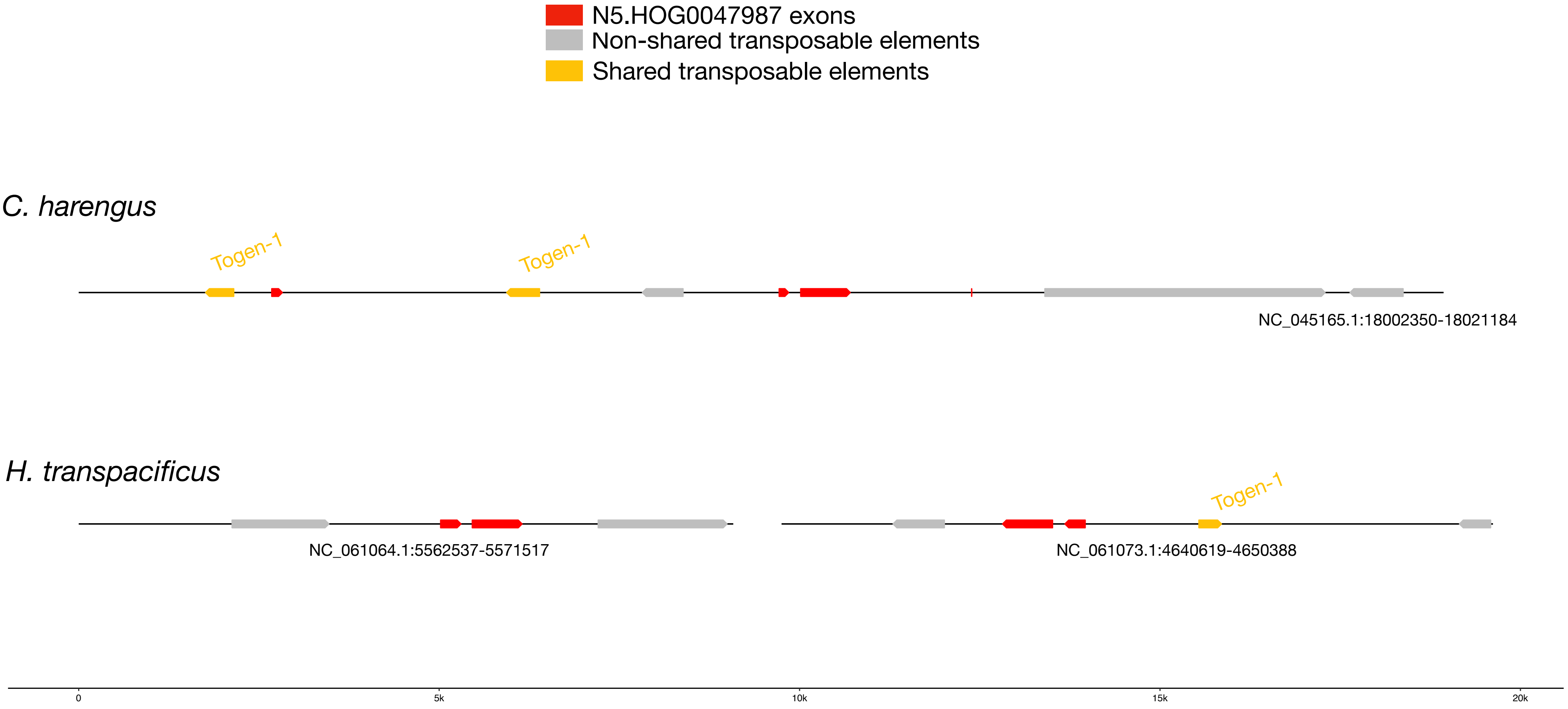


#### Togen-1 :

| | |
|---|---|
| *E. encrasicolus* | *1035 copies scattered in the genome* |
| *A. alosa* | *466 copies scattered in the genome* |
| *S. pilchardus* | *426 copies scattered in the genome* |
| *C. harengus* | *262 copies scattered in the genome* |
| *H. transpacificus* | *1 copy downstream N5.HOG0047987* |
| *O. eperlanus* | *1 copy on an unplaced scaffold* |
| *B. antarcticus* | *344 copies scattered in the genome* |
| *G. morhua* | *269 copies scattered in the genome* |

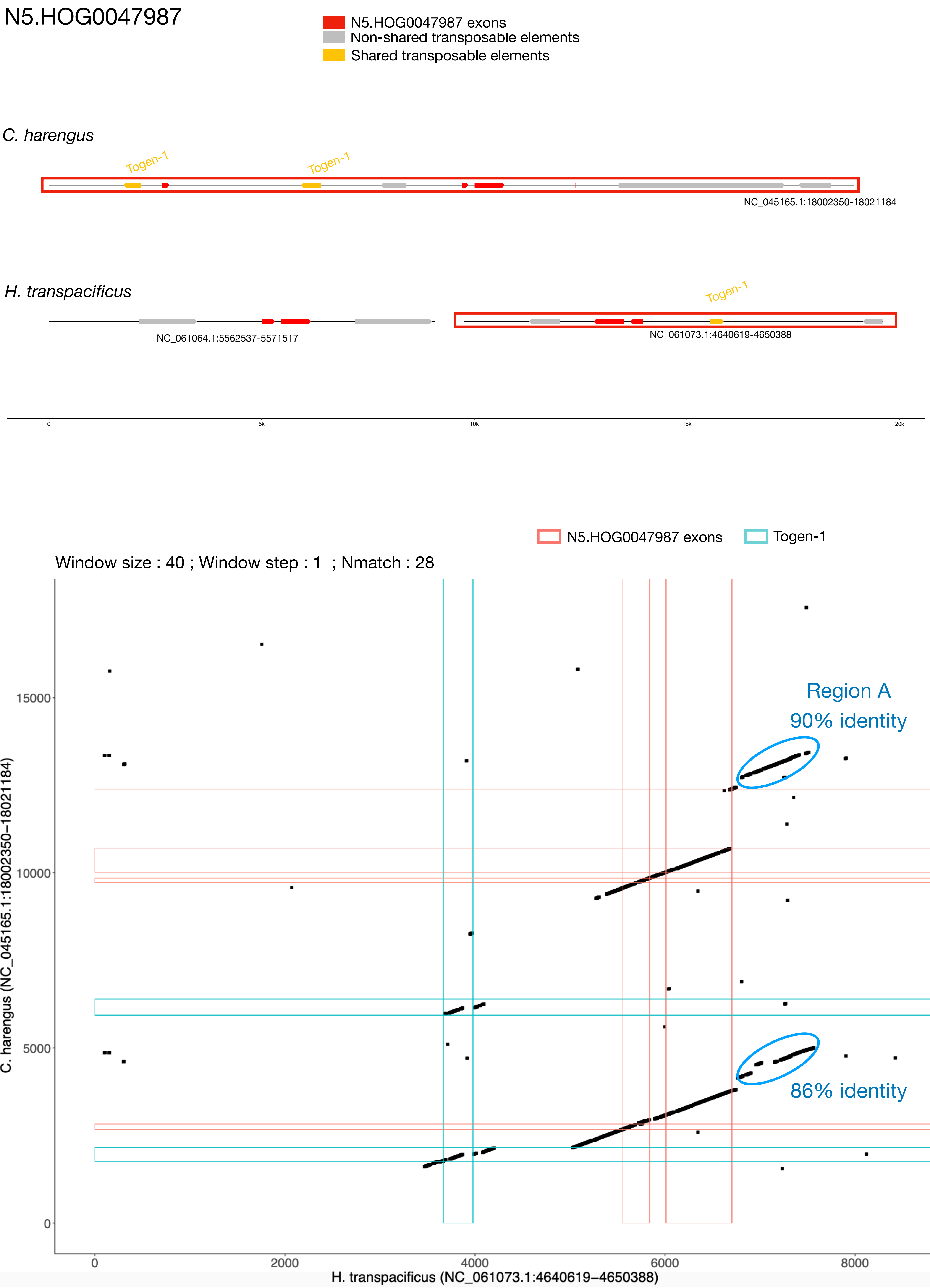

N5.HOG0047987
N5.HOG0047987 exons
Non-shared transposable elements
Shared transposable elements
C. harengus
Togen-1
Togen-1
NC_045165.1:18002350-18021184
H. transpacificus
Togen-1
NC_061064.1:5562537-5571517
NC_061073.1:4640619-4650388
0
5k
10k
15k
20k
N5.HOG0047987 exons
Togen-1
Window size : 40 ; Window step : 1 ; Nmatch : 28
Region A
90% identity
86% identity
C. harengus (NC_045165.1:18002350−18021184)
0
5000
10000
15000
0
2000
4000
6000
8000
H. transpacificus (NC_061073.1:4640619−4650388)

### N5.HOG0047987 Blastn of conserved regions against all genome assemblies

#### Region A

| sseqid | pident | length | mismatch | gapopen | qstart | qend | sstart | send | evalue | bitscore |
|---|---|---|---|---|---|---|---|---|---|---|
| Hypomesus_transpacificus-NC_061073.1 | 100.000 | 898 | 0 | 0 | 1 | 898 | 4643686 | 4642789 | 0.0 | 1659 |
| Osmerus_eperlanus-NC_085036.1 | 92.541 | 429 | 18 | 9 | 29 | 450 | 6386253 | 6386674 | 5.84e-168 | 603 |
| Osmerus_eperlanus-NC_085036.1 | 100.000 | 63 | 0 | 0 | 1 | 63 | 6386205 | 6386267 | 9.28e-22 | 117 |
| Clupea_harengus-NC_045165.1 | 82.945 | 686 | 64 | 21 | 93 | 735 | 18015066 | 18015741 | 5.92e-158 | 569 |
| Clupea_harengus-NC_045165.1 | 81.955 | 399 | 31 | 14 | 469 | 842 | 18006960 | 18007342 | 8.58e-77 | 300 |
| Hypomesus_transpacificus-NC_061064.1 | 87.609 | 460 | 22 | 17 | 457 | 889 | 5568865 | 5569316 | 2.19e-137 | 501 |
| Hypomesus_transpacificus-NC_061064.1 | 94.898 | 98 | 5 | 0 | 29 | 126 | 5568745 | 5568842 | 7.07e-33 | 154 |

N5.HOG0047987

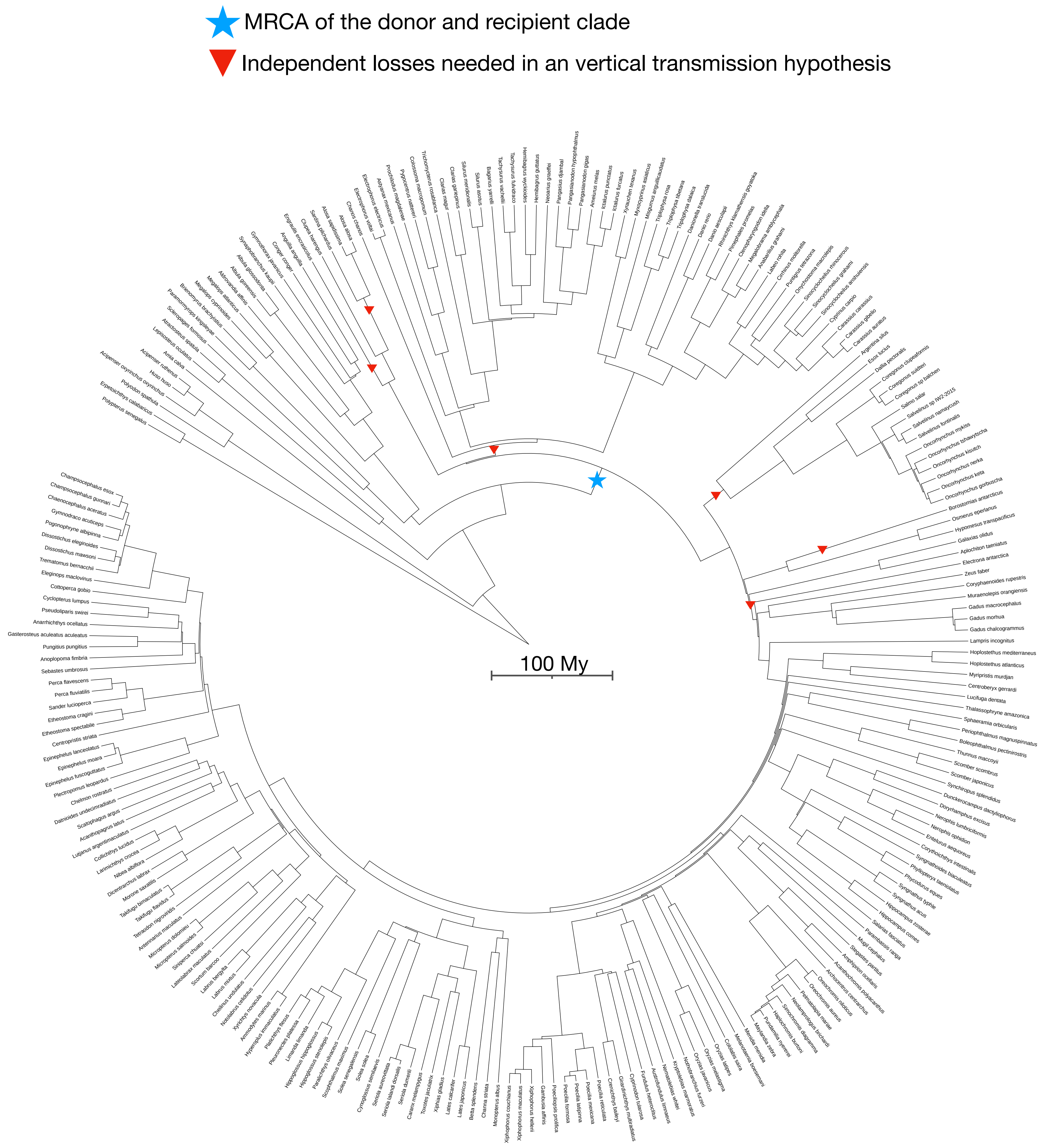

N5.HOG0001647

### N5.HOG0001647

Species tree

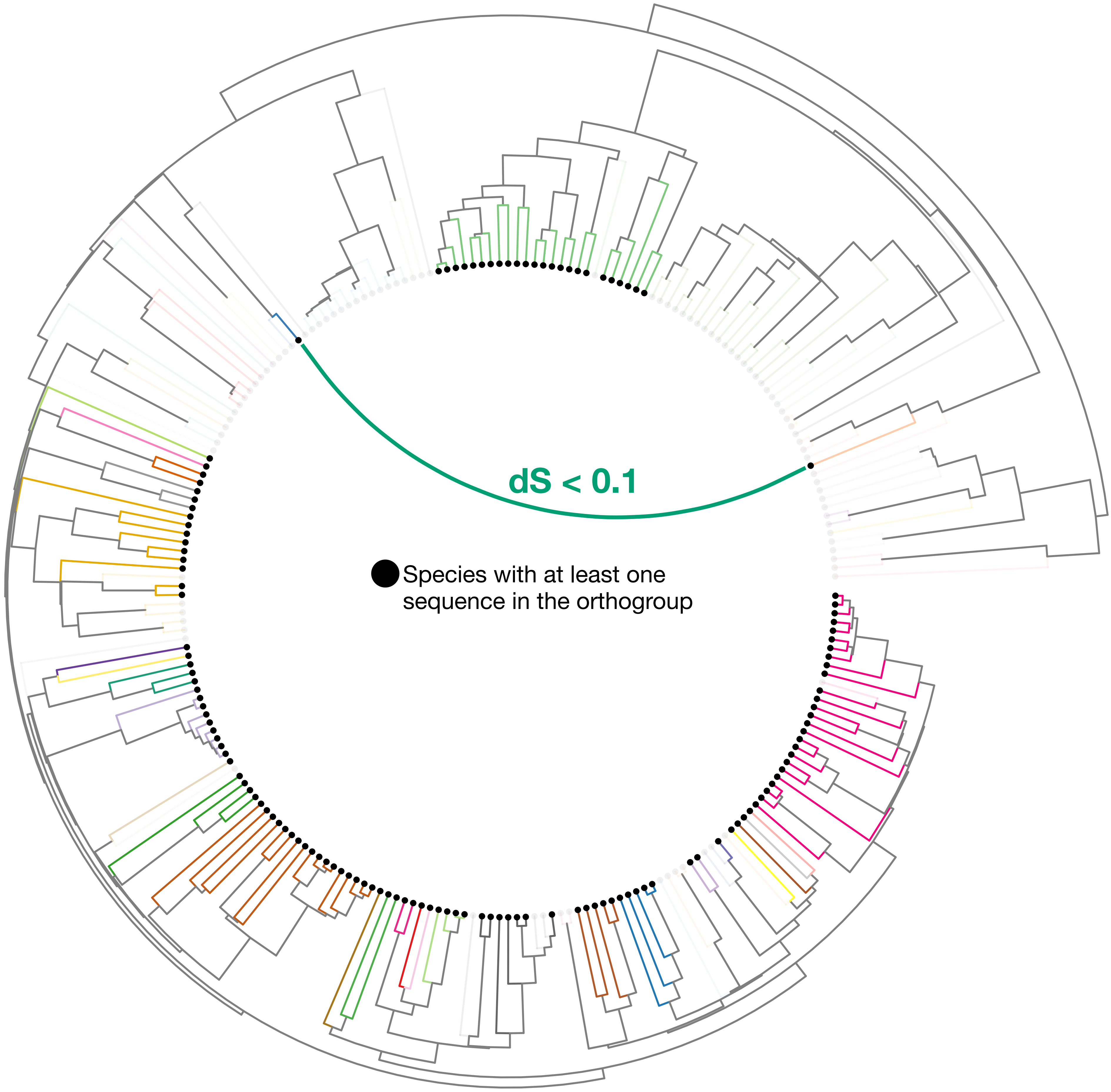

### N5.HOG0001647

Orthogroup phylogeny
and
protein alignment

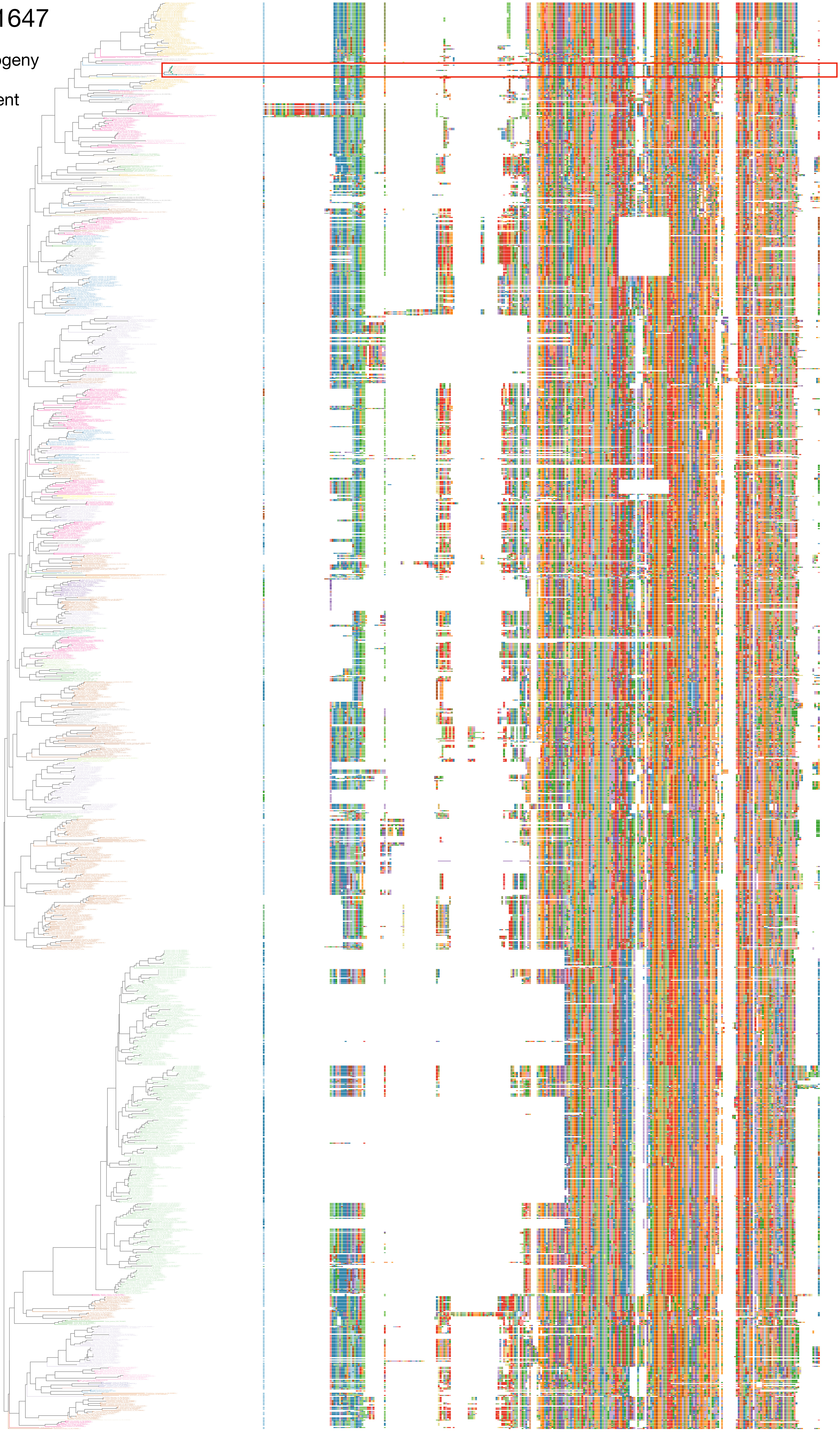

### N5.HOG0001647

Zoom on HGT clade with bootstrap values

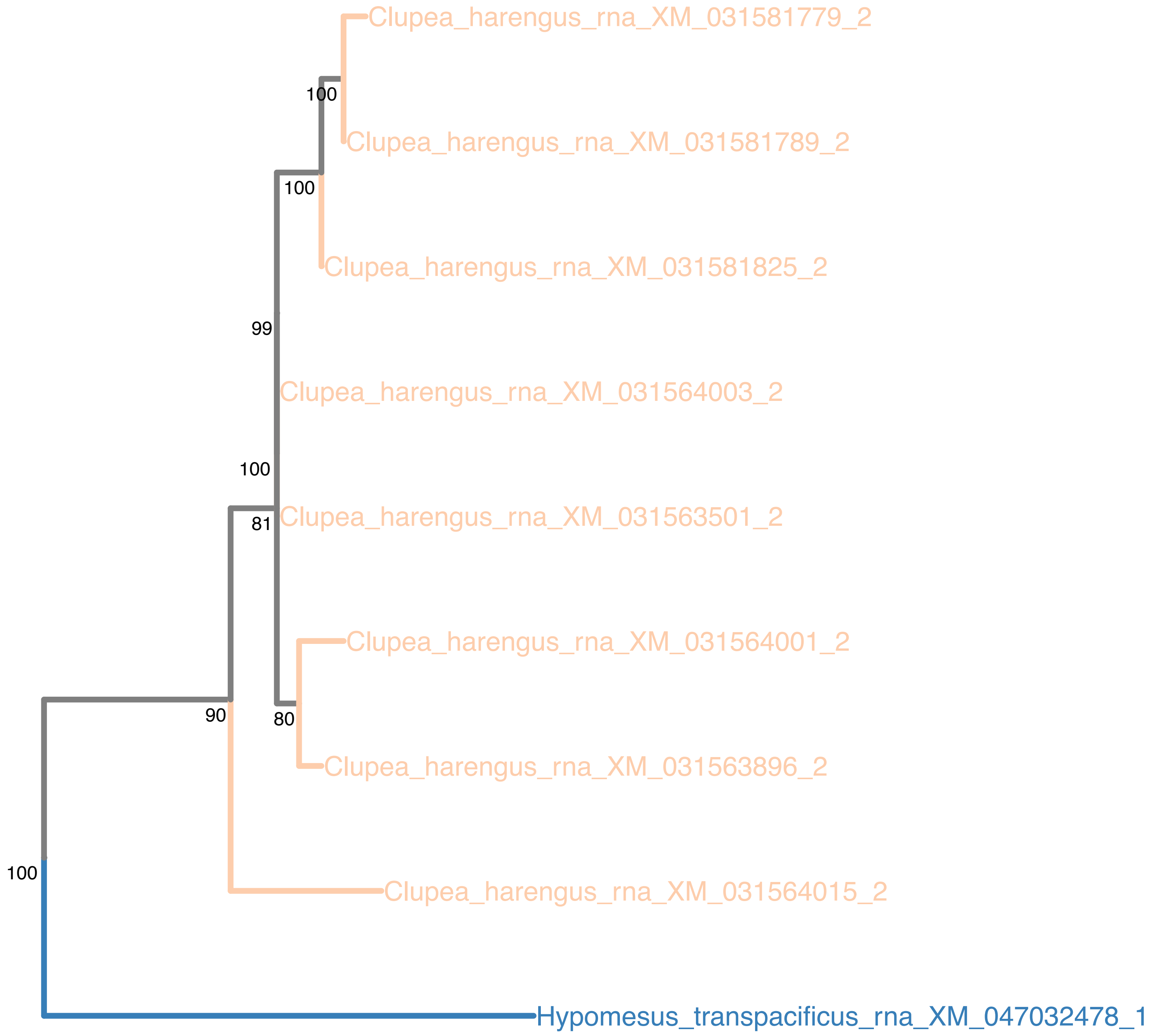

### N5.HOG0001647

Orthogroup phylogeny
and
trimmed
protein alignment

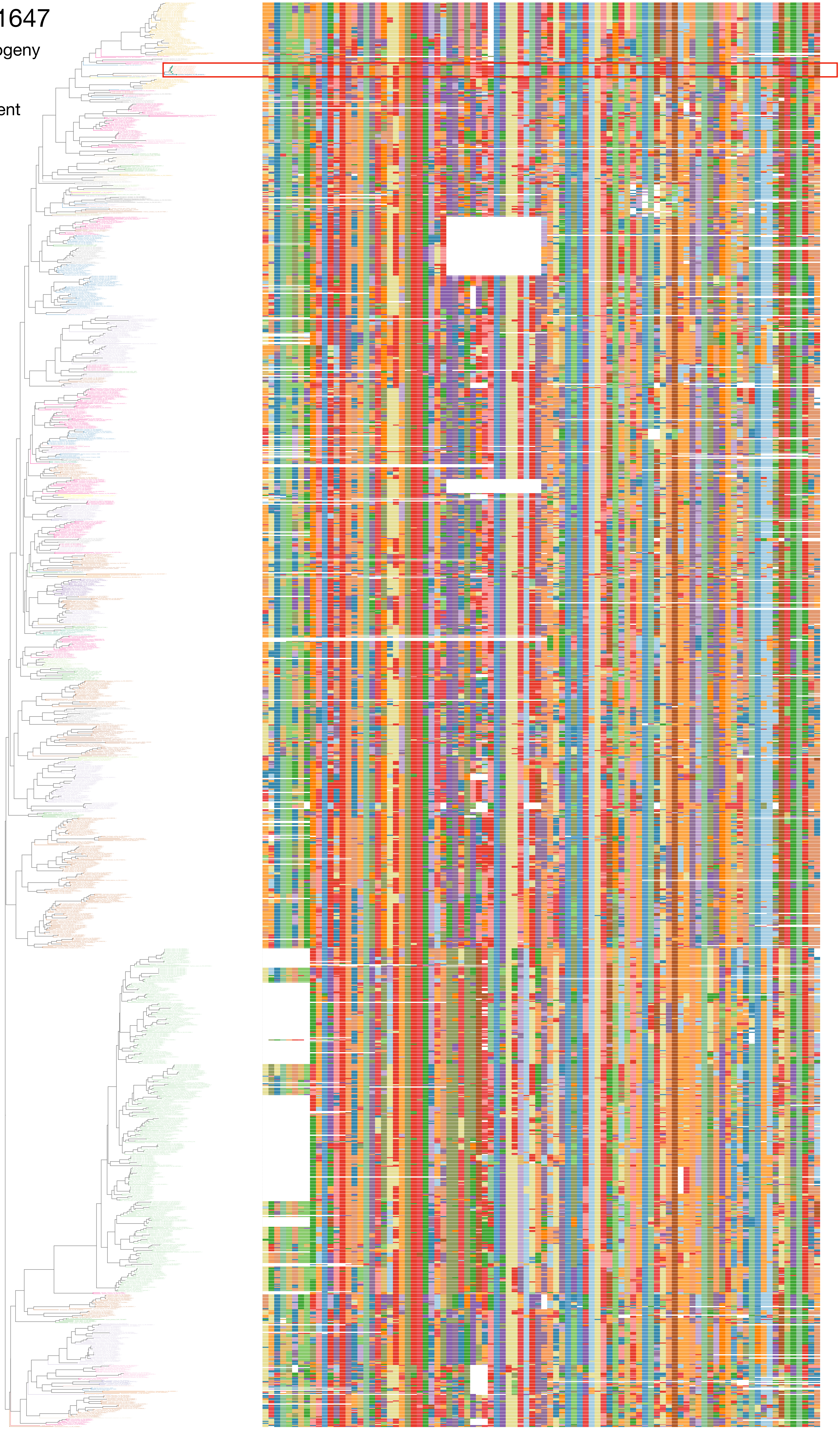

### N5.HOG0001647

« Best-match » phylogeny
and
protein alignment

50 best match against ray-finned fishes protein databases
10 best match against non-fish proteins from UniProt

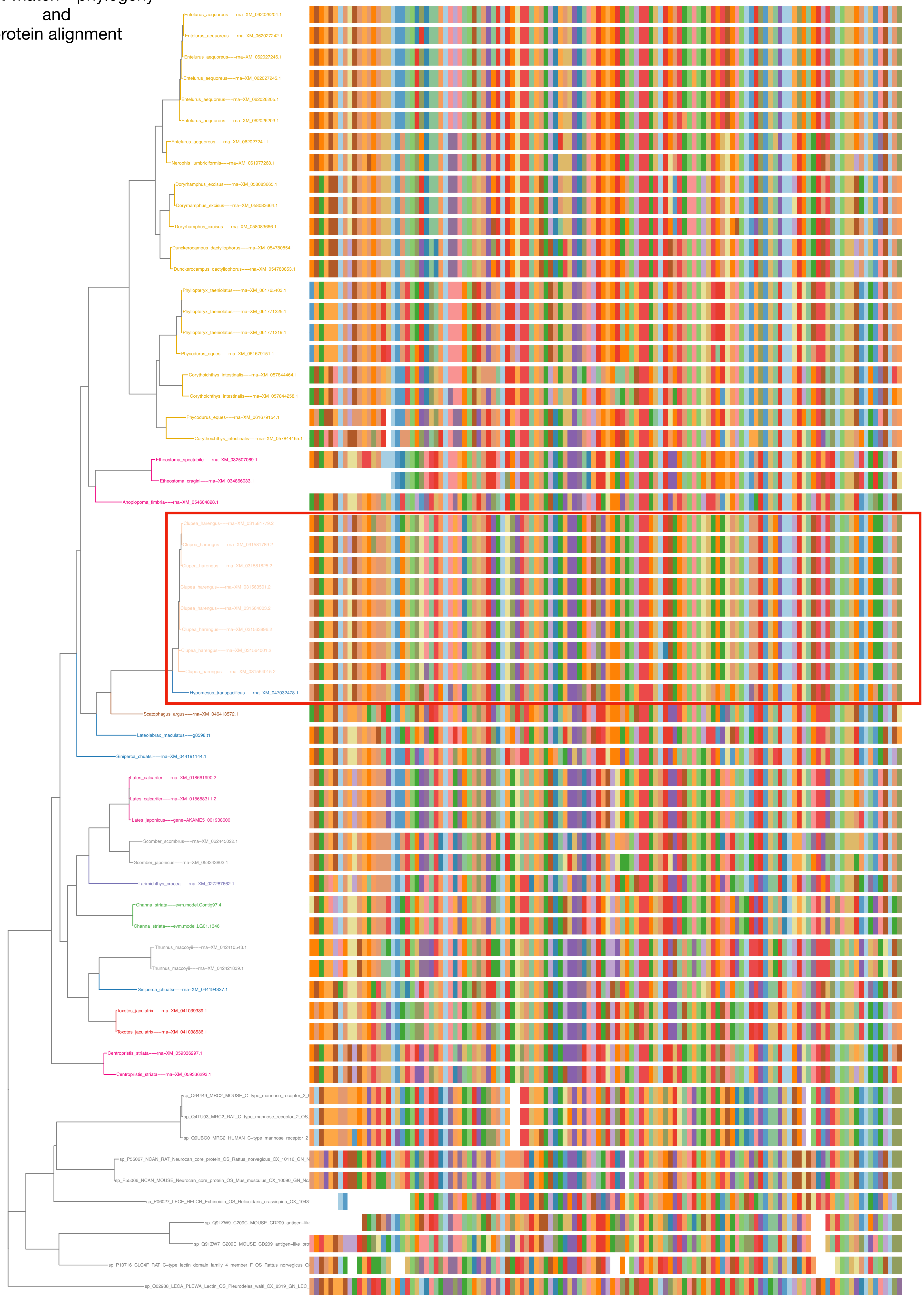

### N5.HOG0001647

dS distribution

Inter-species dS distribution

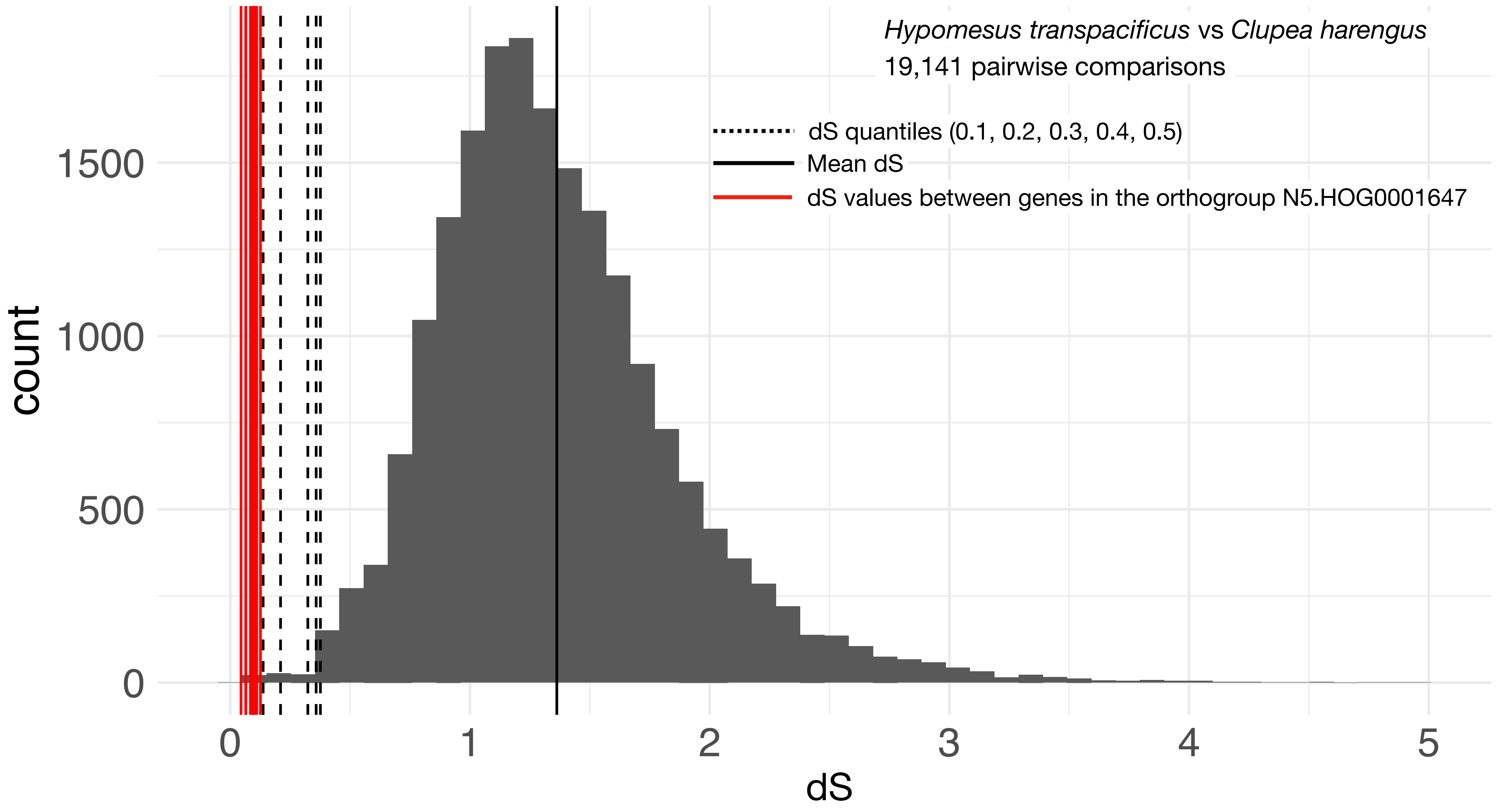


Intra-orthogroup dS

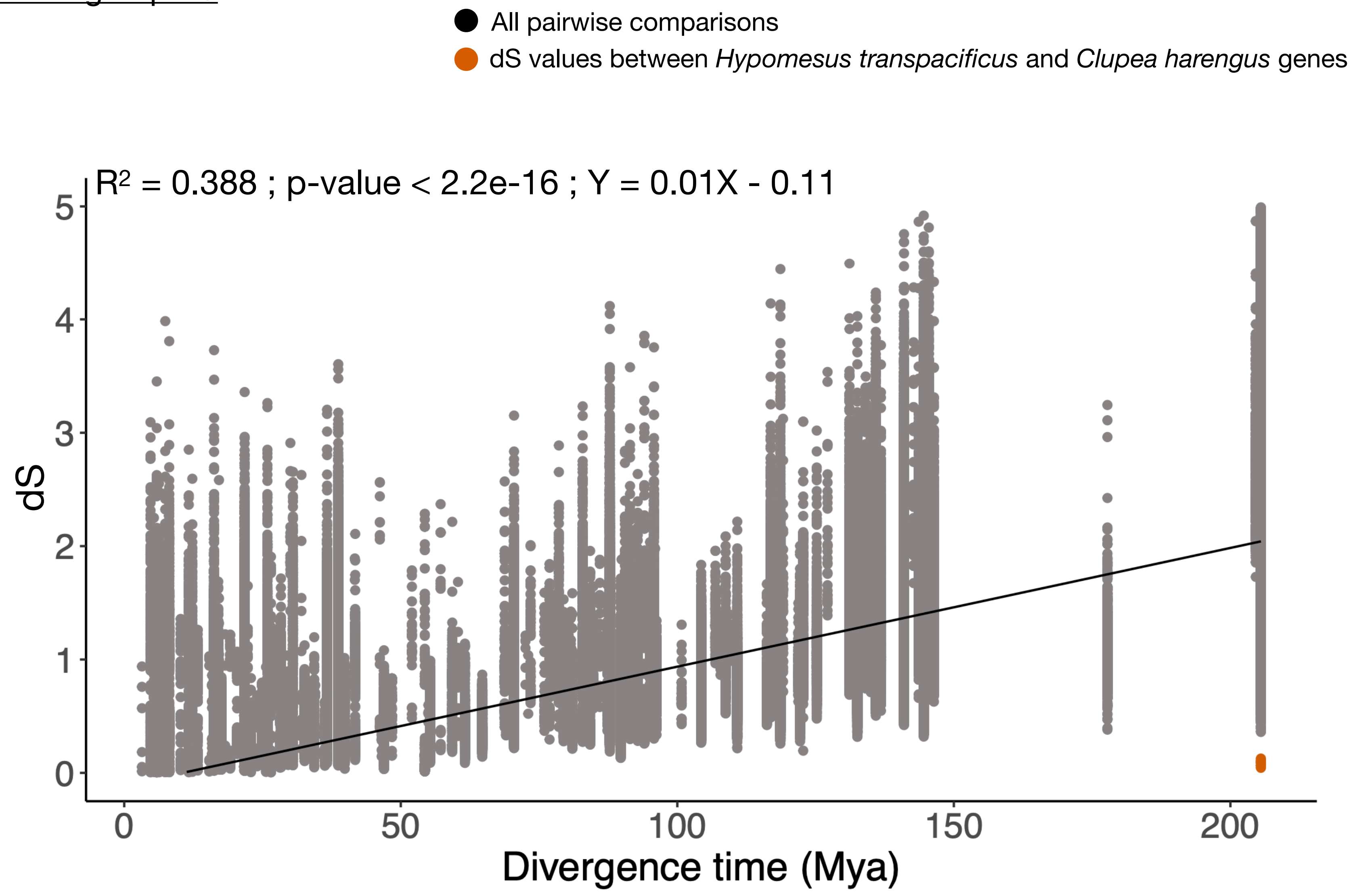

### Osmerus eperlanus have a pseudogenized version of N5.HOG0001647
### Only the 4 last exons remains intact

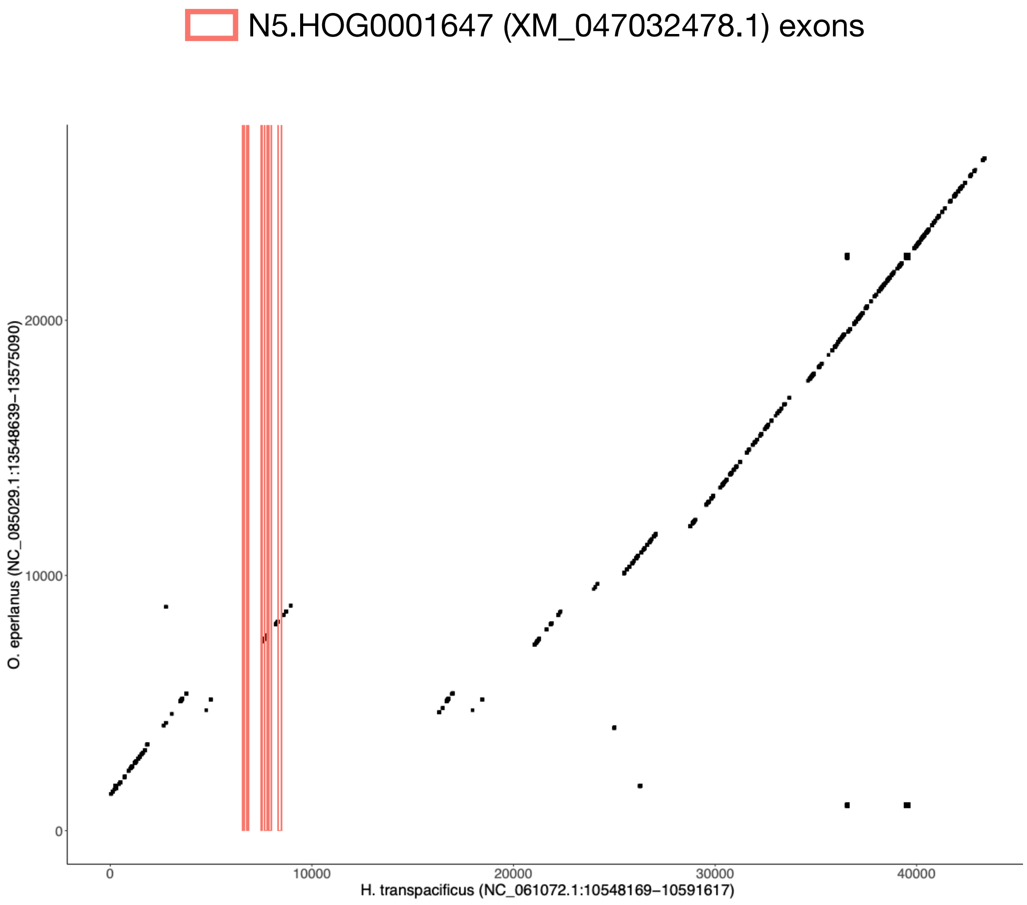


```
         Query: Hypomesus_transpacificus---rna-XM_047032478.1
        Target: NC_085029.1:13548639-13575090 [revcomp]
         Model: protein2genome:local
     Raw score: 586
   Query range: 20 -> 145
  Target range: 19118 -> 18198

   21 : CysProThrAspTrpLysValPheAsnGlyThrCysTyrArgPheAsnProSerValLeu :    40
        ||||||||||||||||||:!!||||||||||  !||||!:!! !||||||||||! !  !|||
        CysProThrAspTrpLysMetPheAsnGlyArgCysPheLeuPheAsnProLeuGlnLeu
19118 : TGTCCCACTGATTGGAAGATGTTCAATGGTCGCTGTTTCCTTTTTAATCCATTACAATTG : 19061

   41 : HisTrpAlaAspAlaGln  >>>> Target Intron 1 >>>>  GluSerCysMetL :    51
        ||||||||| !!|||||||            104 bp           |||||||||||||
        HisTrpAlaHisAlaGln++                         ++GluSerCysMetL
19060 : CATTGGGCTCACGCTCAGgt.........................agGAAAGCTGTATGA : 18924

   52 : ysGluGlyAlaAsnLeuAlaSerIleHisSerLeuGluGluTyrLysPheValLysGluL :    71
        ||||||||||||||||||||||||||||||||||||||||||||  !||||||||||||||
        ysGluGlyAlaAsnLeuAlaSerIleHisSerLeuGluGluTyrAlaPheValLysGluL
18923 : AGGAGGGGGCAAACCTTGCATCCATACACAGCCTTGAAGAGTATGCGTTTGTTAAGGAGC : 18864

   72 : euThrThrAlaAlaLeuThrProSerTrpLeuGlyGlyGlyAspCysGlnVal  >>>>  :    89
        ||||||||||||! !||||! !||||||||||:!!|||||||  !|||||||||||||
        euThrThrAlaAspLeuIleProSerTrpIleGlyGlyThrAspCysGlnVal++
18863 : TGACAACTGCAGACTTAATACCATCATGGATTGGAGGCACAGATTGCCAAGTGgt..... : 18808

   90 : Target Intron 2 >>>>  SerThrArgTrpPheTrpMetAspAlaThrArgTrpAs :   101
             104 bp           ||||||  !|||||||||||||||||::!||||!!!  !||
                            ++SerThrTyrTrpPheTrpMetAspSerThrSerMetAs
18807 : ....................agTCAACATATTGGTTTTGGATGGATAGTACAAGTATGGA : 18670

  102 : pPheThrAspTrpCysTyrAlaGlnProAspThrThrLeuThrGluCysCysLeuGlnMe :   121
        ||||.!!|||||||||||||||||||||||||  !||||||||||||||||||:!!||||||
        pPheAlaAspTrpCysTyrAlaGlnProAspPheThrLeuThrGluCysCysIleGlnMe
18669 : TTTTGCTGACTGGTGCTATGCACAACCTGATTTTACCTTAACTGAGTGCTGCATACAGAT : 18610

  122 : tAsnValGly{V}  >>>> Target Intron 3 >>>>  {al}GlyLysCysTrpAs :   130
        ||||||||||{:}            337 bp           {!!}||||||||||||||
        tAsnValGly{I}++                         ++{le}GlyLysCysTrpAs
18609 : GAATGTTGGA{A}gt.........................ag{TT}GGAAAATGCTGGAA : 18246

  131 : nAspThrProCysThrHisLeuHisAlaSerIleCysAlaLysPro :   145
        ||||||||||||||||||||||||||:!!||||||||||||||||||
        nAspThrProCysThrHisLeuHisSerSerIleCysAlaLysPro
18245 : TGACACACCTTGTACGCATCTTCATTCATCAATCTGCGCCAAGCCA : 18199
```

### N5.HOG0001647

#### Micro-synteny

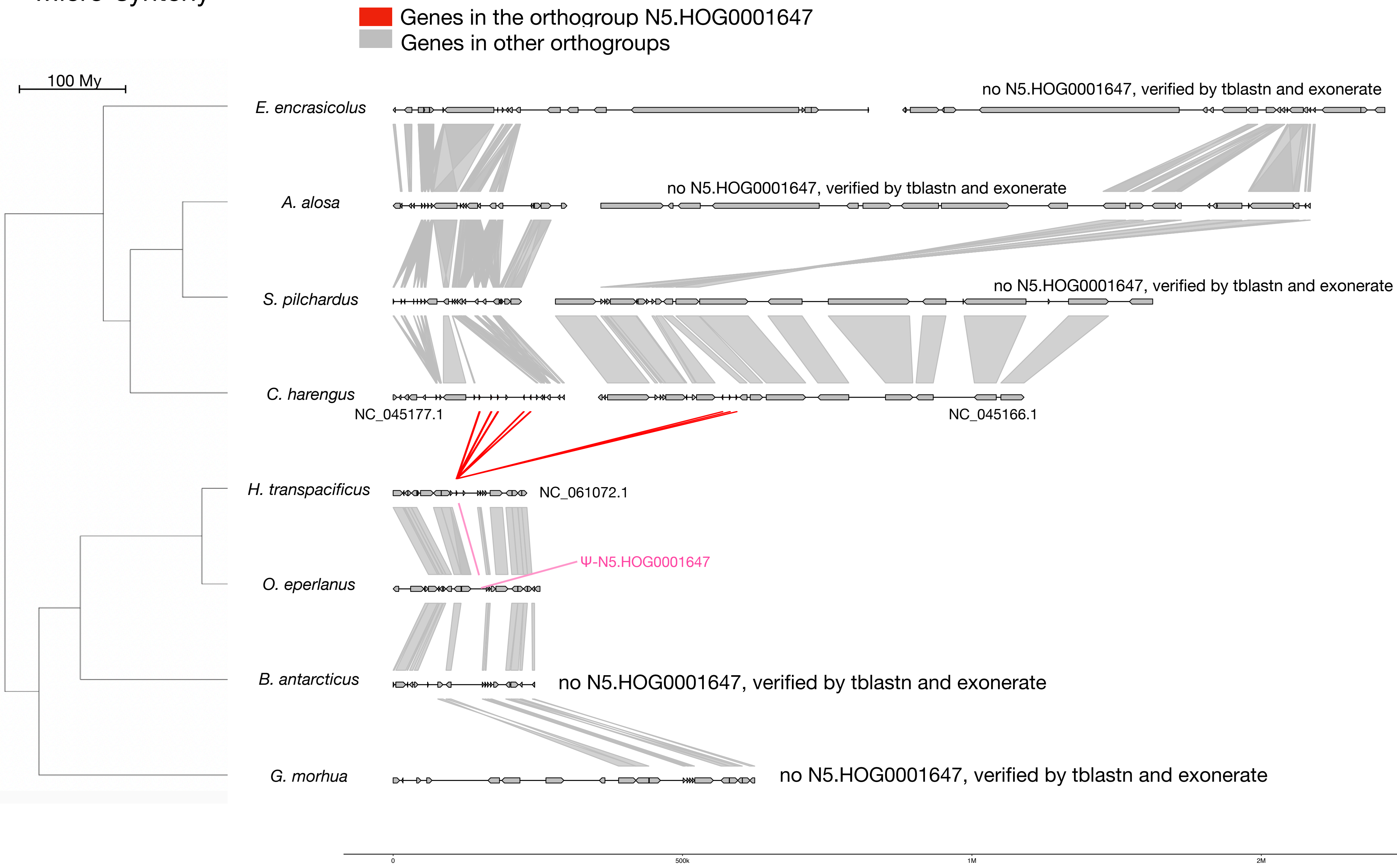


#### Zoom on *C. harengus* and *H. transpacificus*

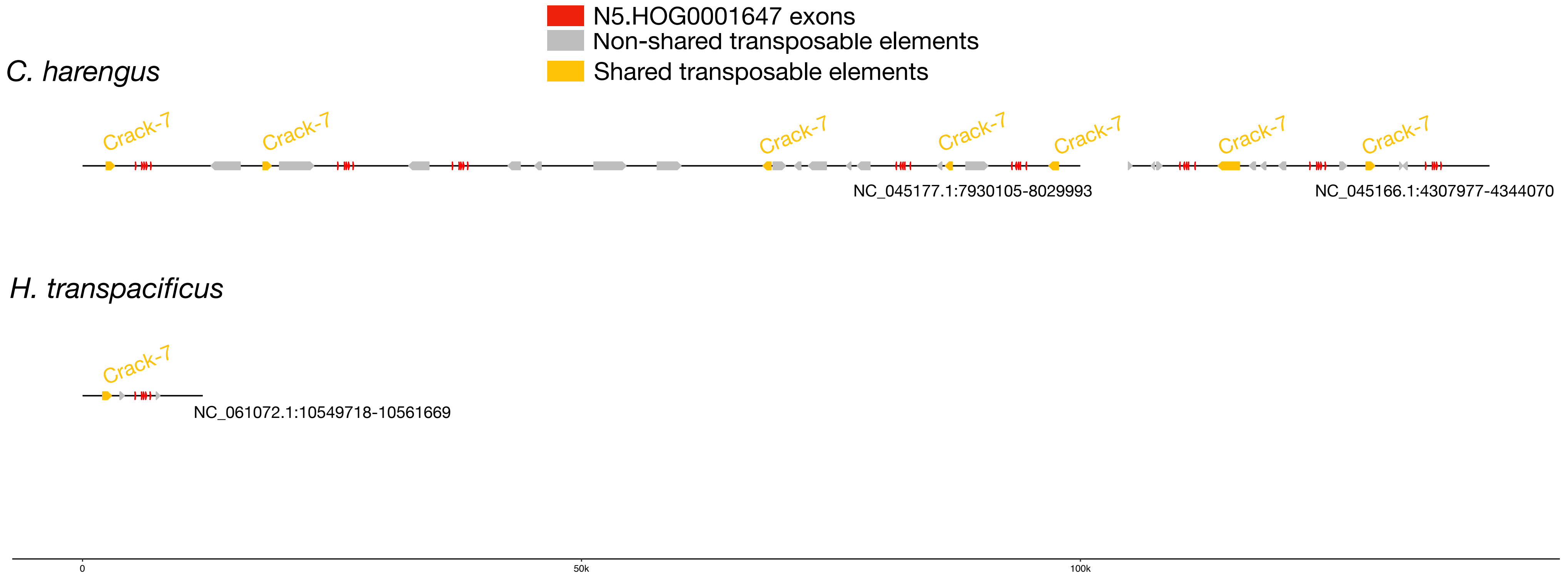


#### Crack-7 :

| | |
|---|---|
| *E. encrasicolus* | *253 copies scattered in the genome* |
| *A. alosa* | *108 copies scattered in the genome* |
| *S. pilchardus* | *101 copies scattered in the genome* |
| *C. harengus* | *231 copies scattered in the genome* |
| *H. transpacificus* | *1 copy upstream N5.HOG0001647* |
| *O. eperlanus* | *1 copy upstream N5.HOG0001647* |
| *B. antarcticus* | *711 copies scattered in the genome* |
| *G. morhua* | *24 copies scattered in the genome* |

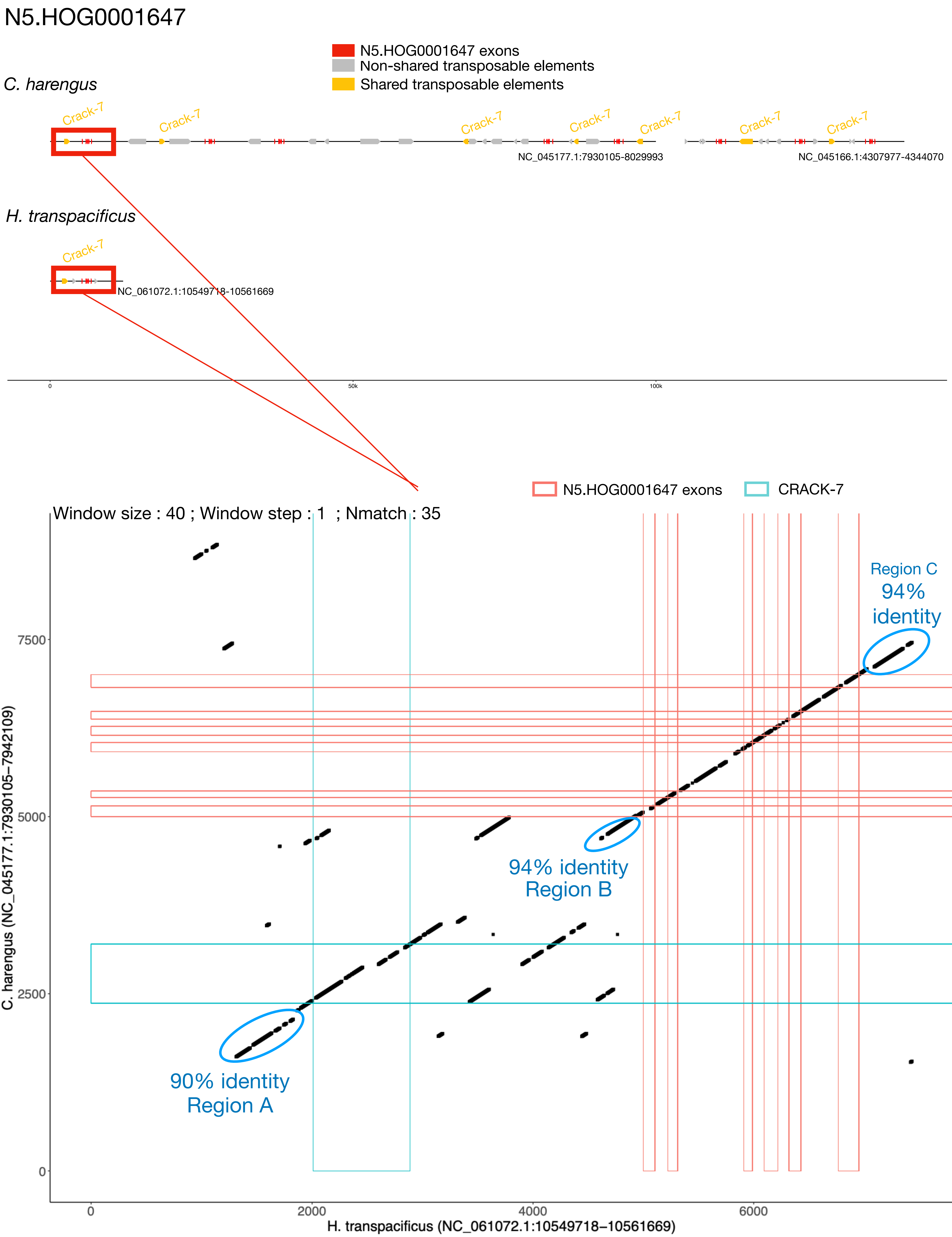

N5.HOG0001647
N5.HOG0001647 exons
Non-shared transposable elements
Shared transposable elements
C. harengus
Crack-7
NC_045177.1:7930105-8029993
NC_045166.1:4307977-4344070
H. transpacificus
NC_061072.1:10549718-10561669
0
50k
100k
N5.HOG0001647 exons
CRACK-7
Window size : 40 ; Window step : 1 ; Nmatch : 35
Region C
94% identity
94% identity
Region B
90% identity
Region A
7500
5000
2500
0
2000
4000
6000
C. harengus (NC_045177.1:7930105–7942109)
H. transpacificus (NC_061072.1:10549718–10561669)

### N5.HOG0001647 Blastn of conserved regions against all genome assemblies

#### Region A

| sseqid | pident | length | mismatch | gapopen | qstart | qend | sstart | send | evalue | bitscore |
|---|---|---|---|---|---|---|---|---|---|---|
| Hypomesus_transpacificus-NC_061072.1 | 100.000 | 701 | 0 | 0 | 1 | 701 | 10551017 | 10551717 | 0.0 | 1295 |
| Hypomesus_transpacificus-NC_061072.1 | 96.050 | 481 | 15 | 4 | 222 | 701 | 10564441 | 10564918 | 0.0 | 780 |
| Osmerus_eperlanus-NC_085029.1 | 88.667 | 600 | 34 | 14 | 18 | 602 | 13570716 | 13570136 | 0.0 | 701 |
| Osmerus_eperlanus-NC_085029.1 | 93.491 | 169 | 8 | 2 | 534 | 701 | 13570112 | 13569946 | 2.43e-61 | 248 |
| Clupea_harengus-NC_045177.1 | 88.732 | 568 | 36 | 9 | 15 | 561 | 7931715 | 7932275 | 0.0 | 669 |
| Clupea_harengus-NC_045177.1 | 89.755 | 449 | 31 | 9 | 18 | 451 | 7999562 | 7999114 | 2.75e-155 | 560 |
| Clupea_harengus-NC_045177.1 | 93.617 | 188 | 11 | 1 | 252 | 438 | 7942735 | 7942922 | 8.63e-71 | 279 |
| Clupea_harengus-NC_045177.1 | 93.617 | 188 | 11 | 1 | 252 | 438 | 7962543 | 7962730 | 8.63e-71 | 279 |
| Clupea_harengus-NC_045177.1 | 88.426 | 216 | 11 | 4 | 15 | 216 | 7959519 | 7959734 | 2.43e-61 | 248 |
| Clupea_harengus-NC_045177.1 | 91.860 | 172 | 11 | 2 | 273 | 444 | 8017327 | 8017159 | 5.27e-58 | 237 |
| Clupea_harengus-NC_045177.1 | 93.793 | 145 | 9 | 0 | 18 | 162 | 8029995 | 8029851 | 1.91e-52 | 219 |
| Clupea_harengus-NC_045177.1 | 88.636 | 176 | 10 | 3 | 534 | 701 | 7932333 | 7932506 | 1.49e-48 | 206 |
| Clupea_harengus-NC_045177.1 | 87.571 | 177 | 11 | 4 | 534 | 701 | 8028012 | 8027838 | 3.21e-45 | 195 |
| Clupea_harengus-NC_045177.1 | 94.495 | 109 | 6 | 0 | 292 | 400 | 7953680 | 7953788 | 1.95e-37 | 169 |
| Clupea_harengus-NC_045177.1 | 94.495 | 109 | 6 | 0 | 292 | 400 | 7965177 | 7965285 | 1.95e-37 | 169 |
| Clupea_harengus-NC_045177.1 | 94.495 | 109 | 6 | 0 | 292 | 400 | 8026676 | 8026568 | 1.95e-37 | 169 |
| Clupea_harengus-NC_045177.1 | 93.578 | 109 | 7 | 0 | 292 | 400 | 7998064 | 7997956 | 9.07e-36 | 163 |
| Clupea_harengus-NC_045177.1 | 92.523 | 107 | 8 | 0 | 294 | 400 | 8015179 | 8015073 | 5.46e-33 | 154 |
| Clupea_harengus-NC_045177.1 | 91.743 | 109 | 7 | 2 | 449 | 556 | 12020161 | 12020268 | 7.06e-32 | 150 |
| Clupea_harengus-NC_045166.1 | 92.661 | 109 | 8 | 0 | 292 | 400 | 4316863 | 4316755 | 4.22e-34 | 158 |
| Clupea_harengus-NC_045166.1 | 92.661 | 109 | 8 | 0 | 292 | 400 | 4332915 | 4333023 | 4.22e-34 | 158 |
| Clupea_harengus-NC_045166.1 | 92.661 | 109 | 8 | 0 | 292 | 400 | 4333464 | 4333572 | 4.22e-34 | 158 |
| Clupea_harengus-NC_045166.1 | 86.620 | 142 | 14 | 5 | 446 | 586 | 7687769 | 7687906 | 1.96e-32 | 152 |
| Clupea_harengus-NC_045161.1 | 85.897 | 156 | 12 | 10 | 449 | 602 | 10703794 | 10703941 | 4.22e-34 | 158 |
| Clupea_harengus-NC_045161.1 | 85.256 | 156 | 13 | 10 | 449 | 602 | 10697665 | 10697812 | 1.96e-32 | 152 |
| Clupea_harengus-NC_045161.1 | 90.351 | 114 | 9 | 2 | 449 | 561 | 10698341 | 10698453 | 2.54e-31 | 148 |
| Clupea_harengus-NC_045160.1 | 85.535 | 159 | 13 | 10 | 446 | 602 | 18963533 | 18963683 | 4.22e-34 | 158 |
| Clupea_harengus-NW_024880432.1 | 85.806 | 155 | 12 | 10 | 449 | 602 | 3936 | 3791 | 1.52e-33 | 156 |
| Clupea_harengus-NC_045164.1 | 92.661 | 109 | 7 | 1 | 449 | 557 | 19556421 | 19556314 | 1.52e-33 | 156 |
| Clupea_harengus-NC_045164.1 | 90.598 | 117 | 9 | 2 | 446 | 561 | 19696149 | 19696264 | 5.46e-33 | 154 |
| Clupea_harengus-NC_045164.1 | 85.161 | 155 | 13 | 10 | 450 | 602 | 5904684 | 5904830 | 7.06e-32 | 150 |
| Clupea_harengus-NC_045164.1 | 84.713 | 157 | 13 | 11 | 449 | 602 | 18888121 | 18887973 | 9.13e-31 | 147 |
| Clupea_harengus-NC_045172.1 | 86.429 | 140 | 17 | 2 | 445 | 583 | 4917278 | 4917416 | 1.96e-32 | 152 |
| Clupea_harengus-NC_045172.1 | 90.517 | 116 | 10 | 1 | 446 | 561 | 17356316 | 17356430 | 1.96e-32 | 152 |
| Clupea_harengus-NC_045172.1 | 84.810 | 158 | 12 | 12 | 449 | 603 | 20788760 | 20788612 | 2.54e-31 | 148 |
| Clupea_harengus-NC_045167.1 | 87.407 | 135 | 12 | 5 | 449 | 581 | 12722539 | 12722408 | 7.06e-32 | 150 |
| Clupea_harengus-NC_045152.1 | 84.472 | 161 | 15 | 10 | 444 | 602 | 27779543 | 27779695 | 7.06e-32 | 150 |
| Clupea_harengus-NC_045157.1 | 88.618 | 123 | 12 | 2 | 449 | 570 | 9836960 | 9836839 | 2.54e-31 | 148 |
| Clupea_harengus-NC_045154.1 | 91.667 | 108 | 7 | 2 | 449 | 555 | 25653264 | 25653158 | 2.54e-31 | 148 |
| Clupea_harengus-NC_045174.1 | 84.277 | 159 | 15 | 10 | 446 | 602 | 14935948 | 14936098 | 9.13e-31 | 147 |

### N5.HOG0001647 Blastn of conserved regions against all genome assemblies

#### Region B

| sseqid | pident | length | mismatch | gapopen | qstart | qend | sstart | send | evalue | bitscore |
|---|---|---|---|---|---|---|---|---|---|---|
| Hypomesus_transpacificus-NC_061072.1 | 100.000 | 301 | 0 | 0 | 1 | 301 | 10554417 | 10554717 | 1.43e-154 | 556 |
| Hypomesus_transpacificus-NC_061072.1 | 97.690 | 303 | 4 | 3 | 1 | 301 | 10566748 | 10567049 | 6.73e-143 | 518 |
| Hypomesus_transpacificus-NC_061072.1 | 98.394 | 249 | 2 | 2 | 1 | 248 | 10553289 | 10553536 | 1.94e-118 | 436 |
| Clupea_harengus-NC_045177.1 | 95.203 | 271 | 13 | 0 | 1 | 271 | 7966638 | 7966908 | 3.25e-116 | 429 |
| Clupea_harengus-NC_045177.1 | 94.834 | 271 | 14 | 0 | 1 | 271 | 7934882 | 7935152 | 1.51e-114 | 424 |
| Clupea_harengus-NC_045177.1 | 94.096 | 271 | 16 | 0 | 1 | 271 | 7955150 | 7955420 | 3.27e-111 | 412 |
| Clupea_harengus-NC_045177.1 | 92.620 | 271 | 19 | 1 | 1 | 271 | 8013628 | 8013359 | 5.51e-104 | 388 |
| Clupea_harengus-NC_045177.1 | 93.798 | 258 | 15 | 1 | 1 | 258 | 7996421 | 7996165 | 1.98e-103 | 387 |
| Clupea_harengus-NC_045177.1 | 88.462 | 260 | 11 | 6 | 12 | 271 | 8025187 | 8024947 | 3.43e-76 | 296 |
| Clupea_harengus-NC_045166.1 | 86.038 | 265 | 18 | 4 | 8 | 271 | 4325640 | 4325886 | 2.69e-67 | 267 |
| Clupea_harengus-NC_045166.1 | 84.906 | 265 | 21 | 4 | 8 | 271 | 4337226 | 4337472 | 2.71e-62 | 250 |

#### Region C

| sseqid | pident | length | mismatch | gapopen | qstart | qend | sstart | send | evalue | bitscore |
|---|---|---|---|---|---|---|---|---|---|---|
| Hypomesus_transpacificus-NC_061072.1 | 100.000 | 531 | 0 | 0 | 1 | 531 | 10556687 | 10557217 | 0.0 | 981 |
| Hypomesus_transpacificus-NC_061072.1 | 90.335 | 269 | 5 | 5 | 1 | 269 | 10570317 | 10570564 | 4.86e-87 | 333 |
| Clupea_harengus-NC_045166.1 | 95.030 | 503 | 21 | 1 | 1 | 499 | 4312955 | 4312453 | 0.0 | 787 |
| Clupea_harengus-NC_045166.1 | 94.235 | 503 | 25 | 1 | 1 | 499 | 4327900 | 4328402 | 0.0 | 765 |
| Clupea_harengus-NC_045166.1 | 94.235 | 503 | 25 | 1 | 1 | 499 | 4339493 | 4339995 | 0.0 | 765 |
| Clupea_harengus-NC_045177.1 | 94.831 | 503 | 22 | 1 | 1 | 499 | 7957408 | 7957910 | 0.0 | 782 |
| Clupea_harengus-NC_045177.1 | 94.212 | 501 | 24 | 4 | 1 | 499 | 8023049 | 8022552 | 0.0 | 760 |
| Clupea_harengus-NC_045177.1 | 93.849 | 504 | 26 | 2 | 1 | 499 | 7968889 | 7969392 | 0.0 | 754 |
| Clupea_harengus-NC_045177.1 | 93.988 | 499 | 20 | 4 | 1 | 499 | 7994298 | 7993810 | 0.0 | 747 |
| Clupea_harengus-NC_045177.1 | 93.455 | 382 | 21 | 1 | 122 | 499 | 7937217 | 7937598 | 1.58e-156 | 564 |
| Clupea_harengus-NC_045177.1 | 96.698 | 212 | 7 | 0 | 6 | 217 | 7985928 | 7986139 | 3.73e-93 | 353 |
| Clupea_harengus-NC_045177.1 | 96.386 | 166 | 6 | 0 | 52 | 217 | 8010910 | 8010745 | 2.99e-69 | 274 |
| Clupea_harengus-NC_045177.1 | 92.683 | 123 | 5 | 1 | 381 | 499 | 7959368 | 7959490 | 3.12e-39 | 174 |
| Clupea_harengus-NC_045177.1 | 91.870 | 123 | 6 | 1 | 381 | 499 | 8030149 | 8030027 | 1.45e-37 | 169 |
| Osmerus_eperlanus-NC_085029.1 | 94.821 | 502 | 19 | 3 | 2 | 499 | 13566701 | 13566203 | 0.0 | 776 |
| Osmerus_eperlanus-NC_085029.1 | 91.870 | 123 | 6 | 1 | 381 | 499 | 13570870 | 13570748 | 1.45e-37 | 169 |

### N5.HOG0001647

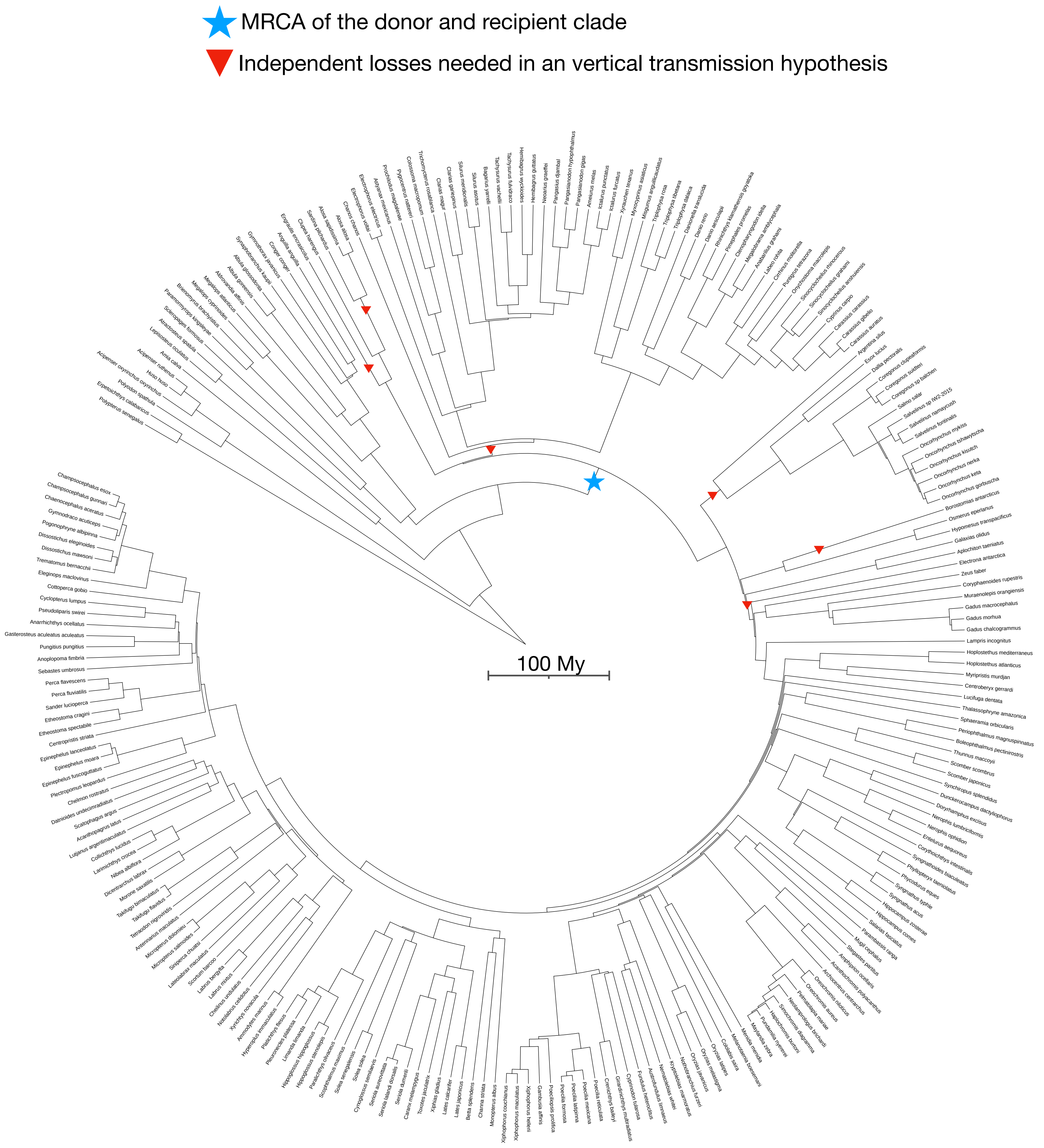

N5.HOG0013514

### N5.HOG0013514

Species tree

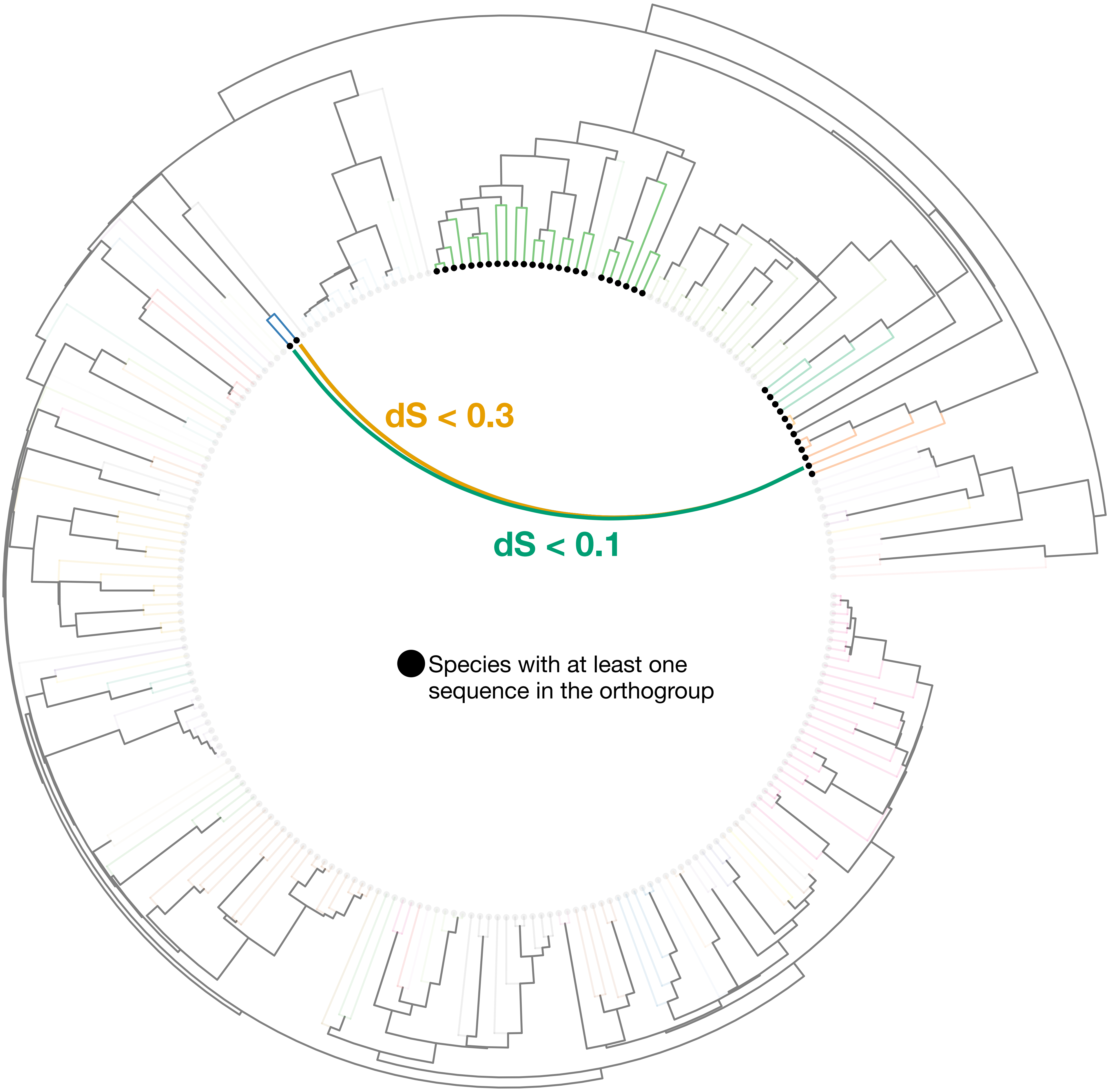

### N5.HOG0013514

Orthogroup phylogeny
and
protein alignment

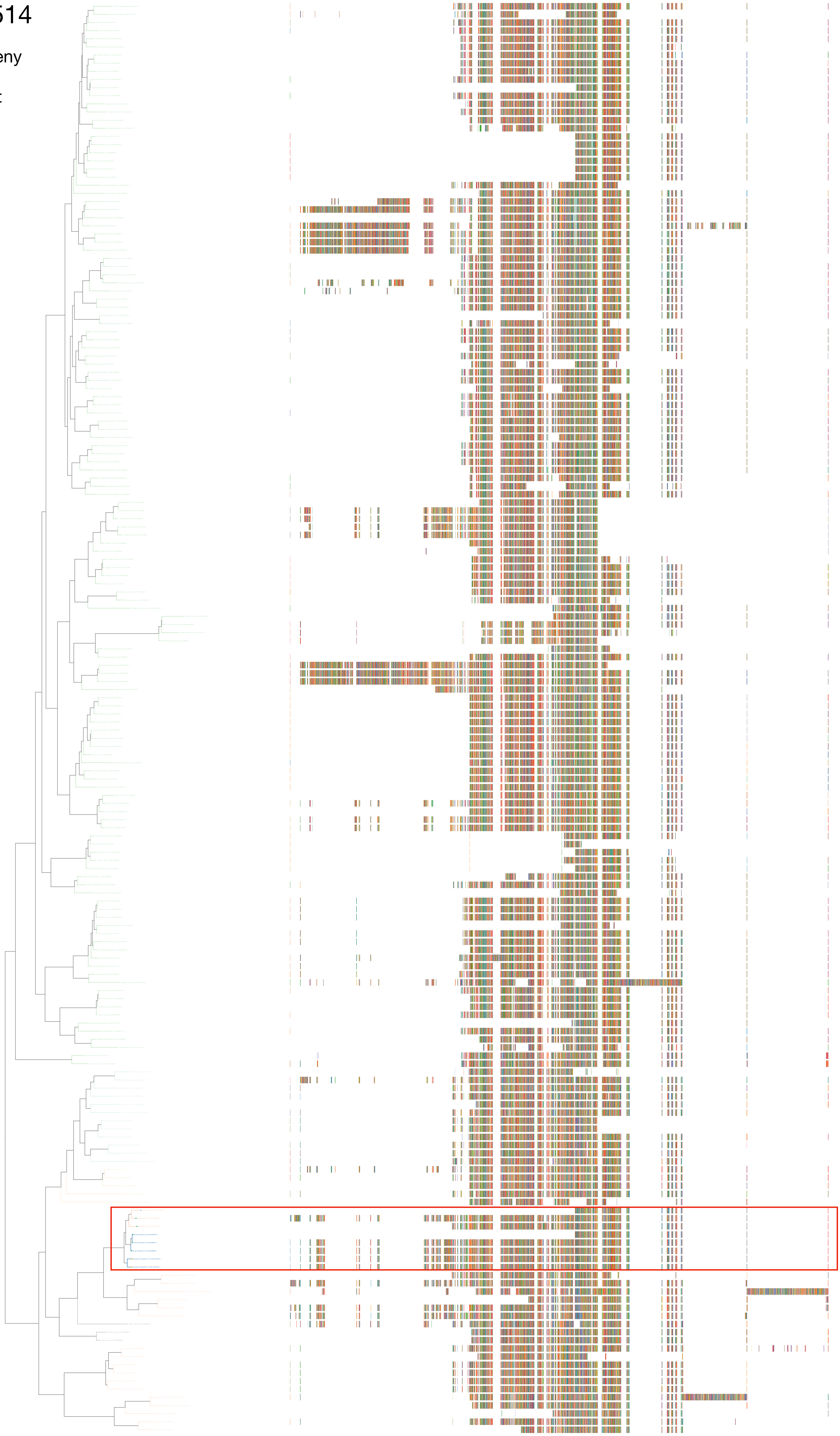

N5.HOG0013514

Zoom on HGT clade with bootstrap values

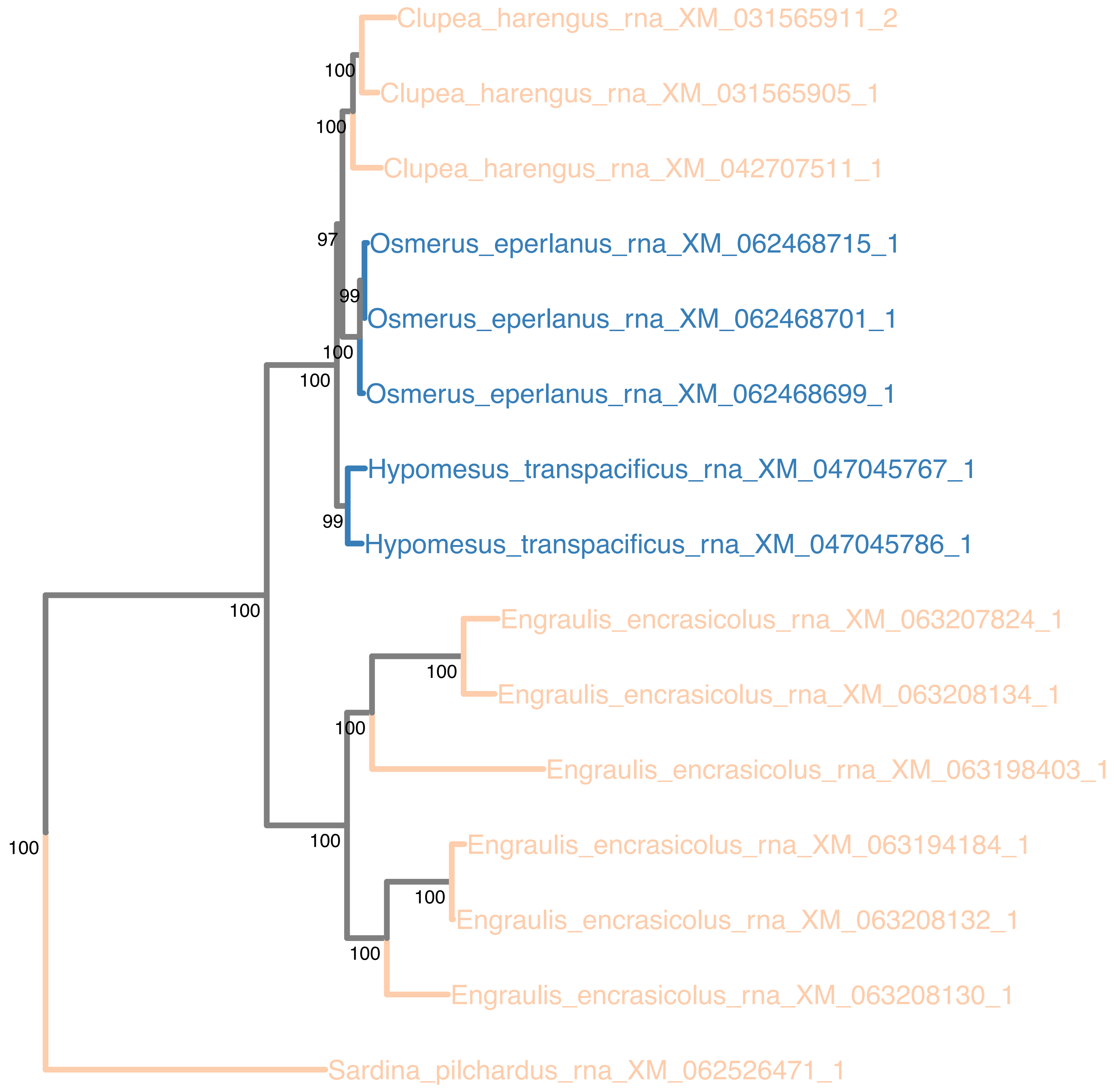

### N5.HOG0013514

Orthogroup phylogeny
and
trimmed
protein alignment

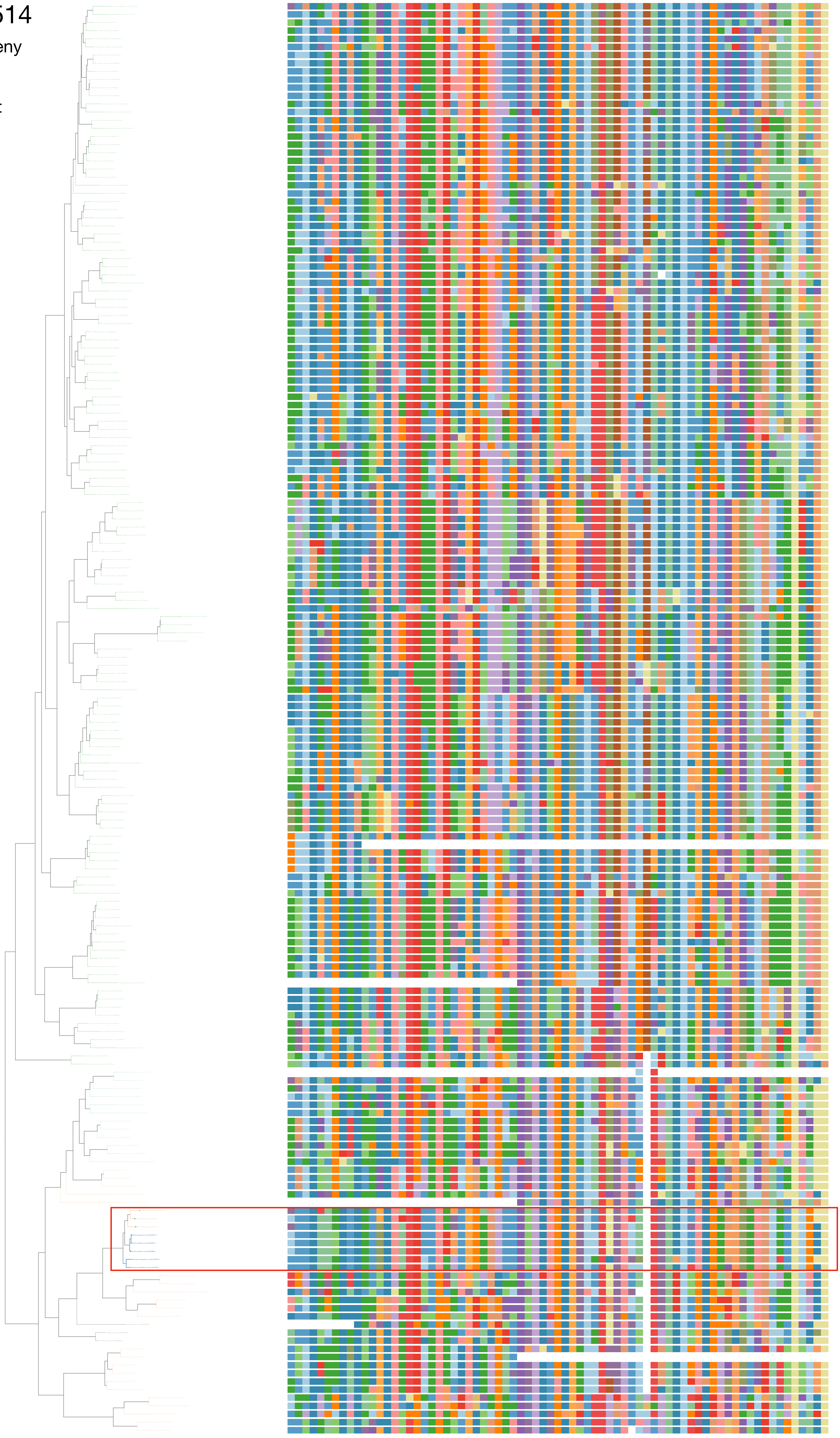

### N5.HOG0013514

« Best-match » phylogeny
and
protein alignment

50 best match against ray-finned fishes protein databases
10 best match against non-fish proteins from UniProt

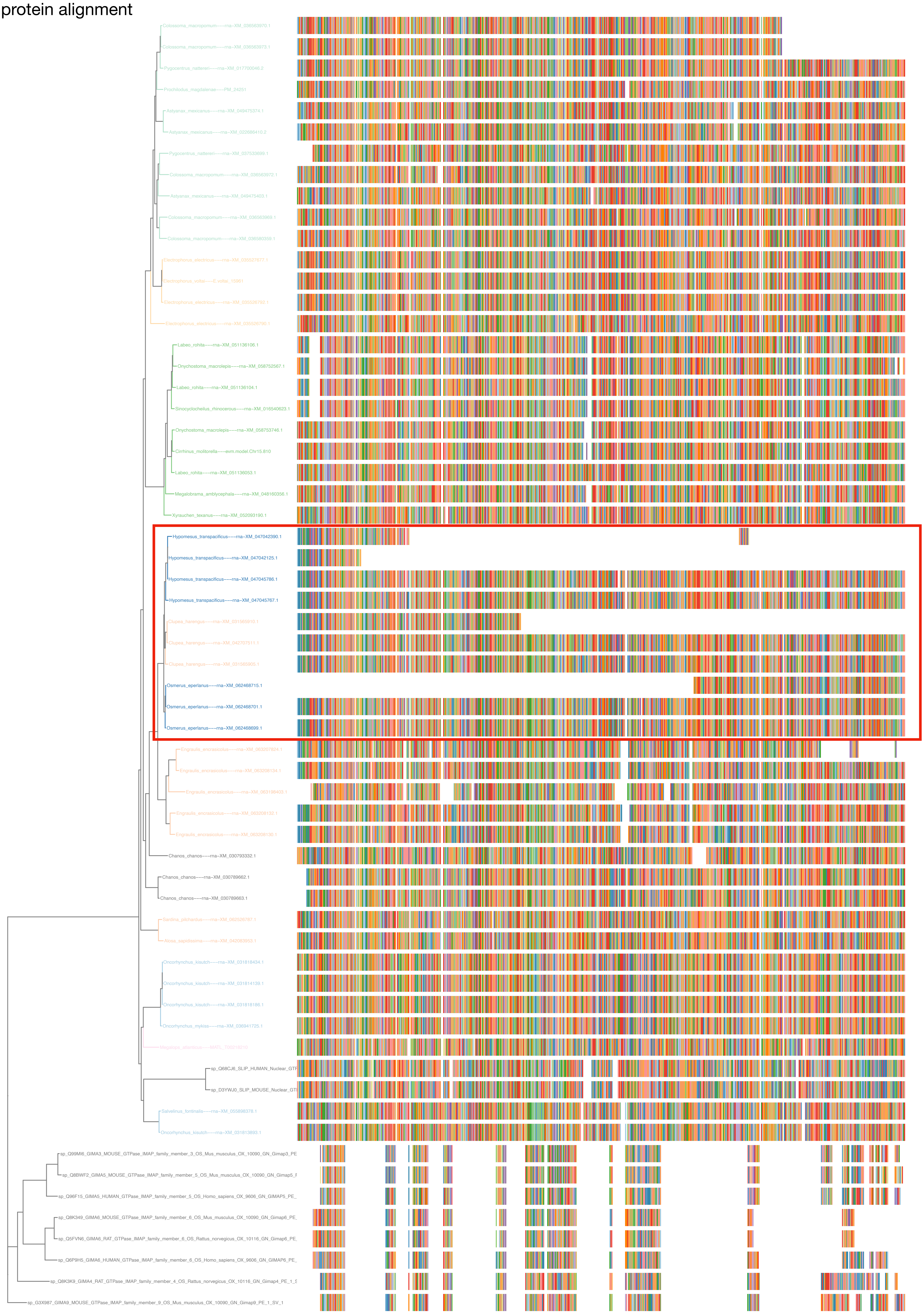

### N5.HOG0013514

#### dS distribution

##### Inter-species dS distribution

dS quantiles (0.1, 0.2, 0.3, 0.4, 0.5)
Mean dS
dS values between genes in the orthogroup N5.HOG0013514

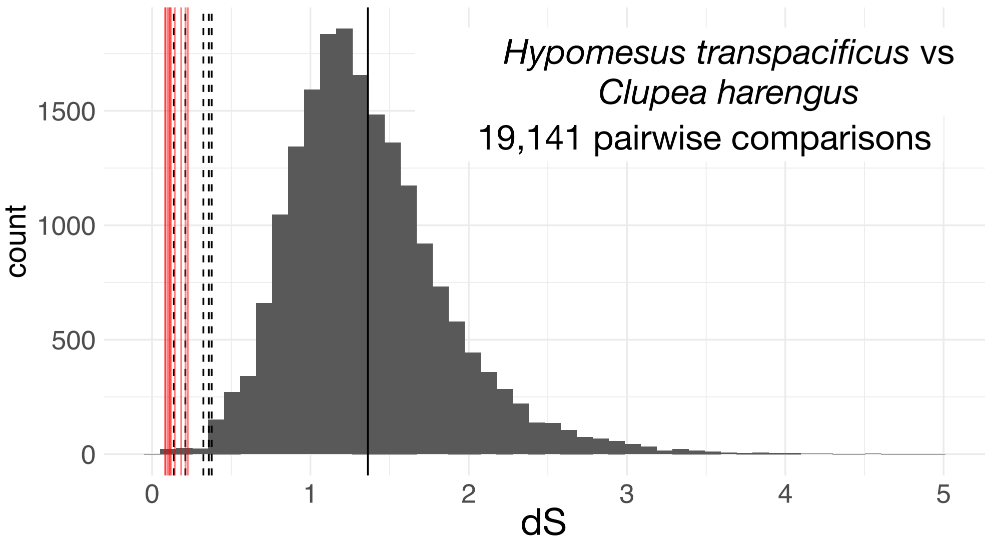


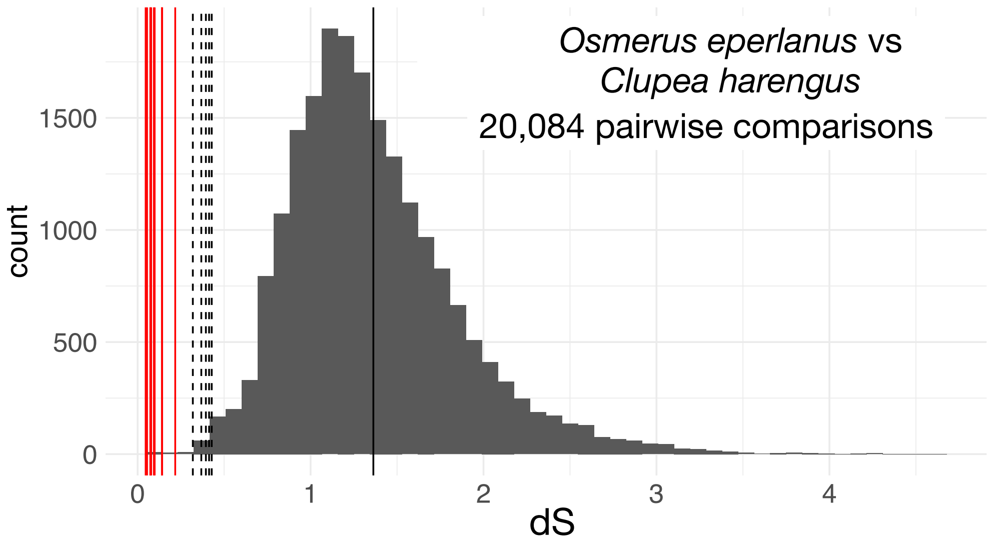


##### Intra-orthogroup dS

All pairwise comparisons
dS values between *Hypomesus transpacificus* and *Clupea harengus* genes
dS values between *Osmerus eperlanus* and *Clupea harengus* genes

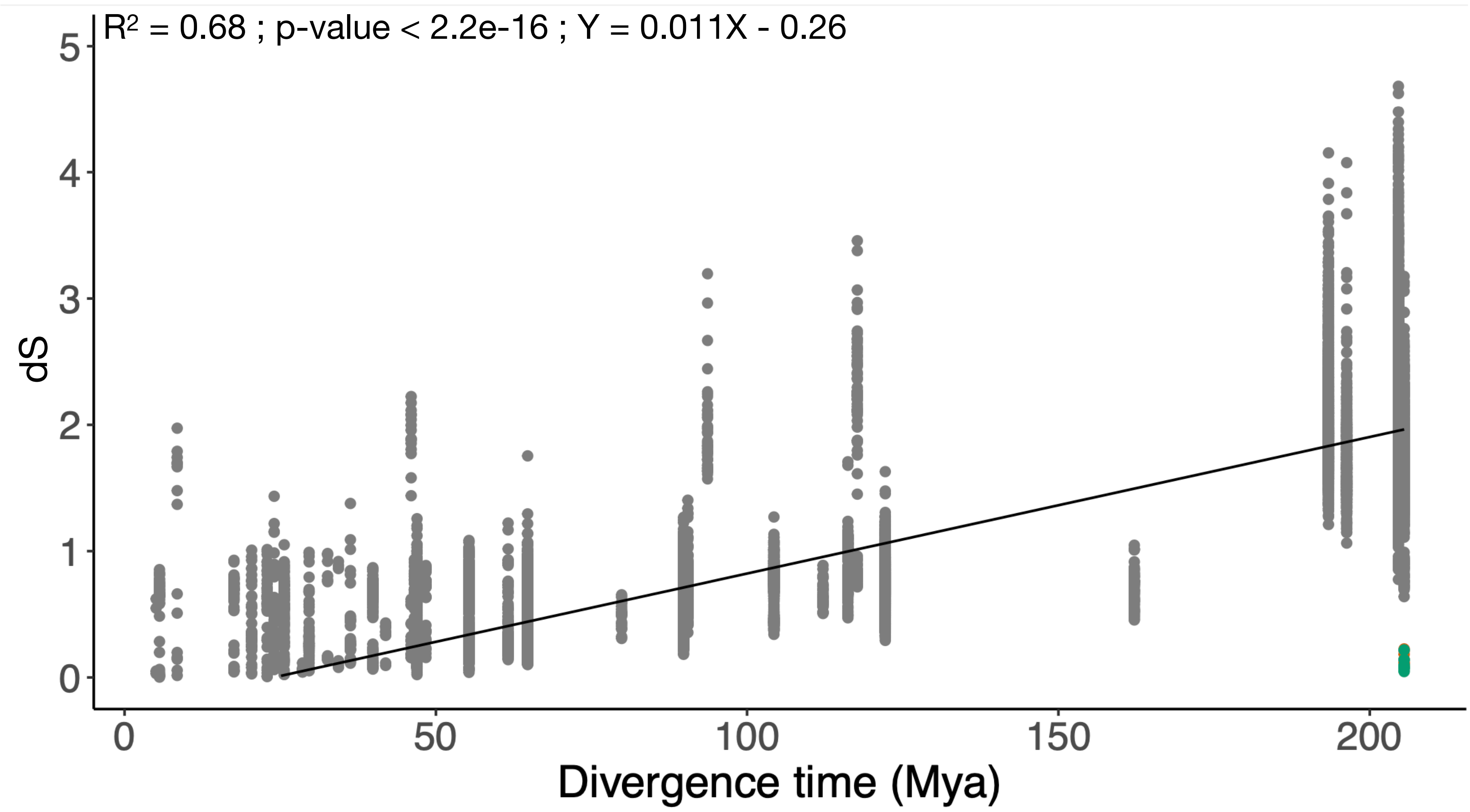

### N5.HOG0013514

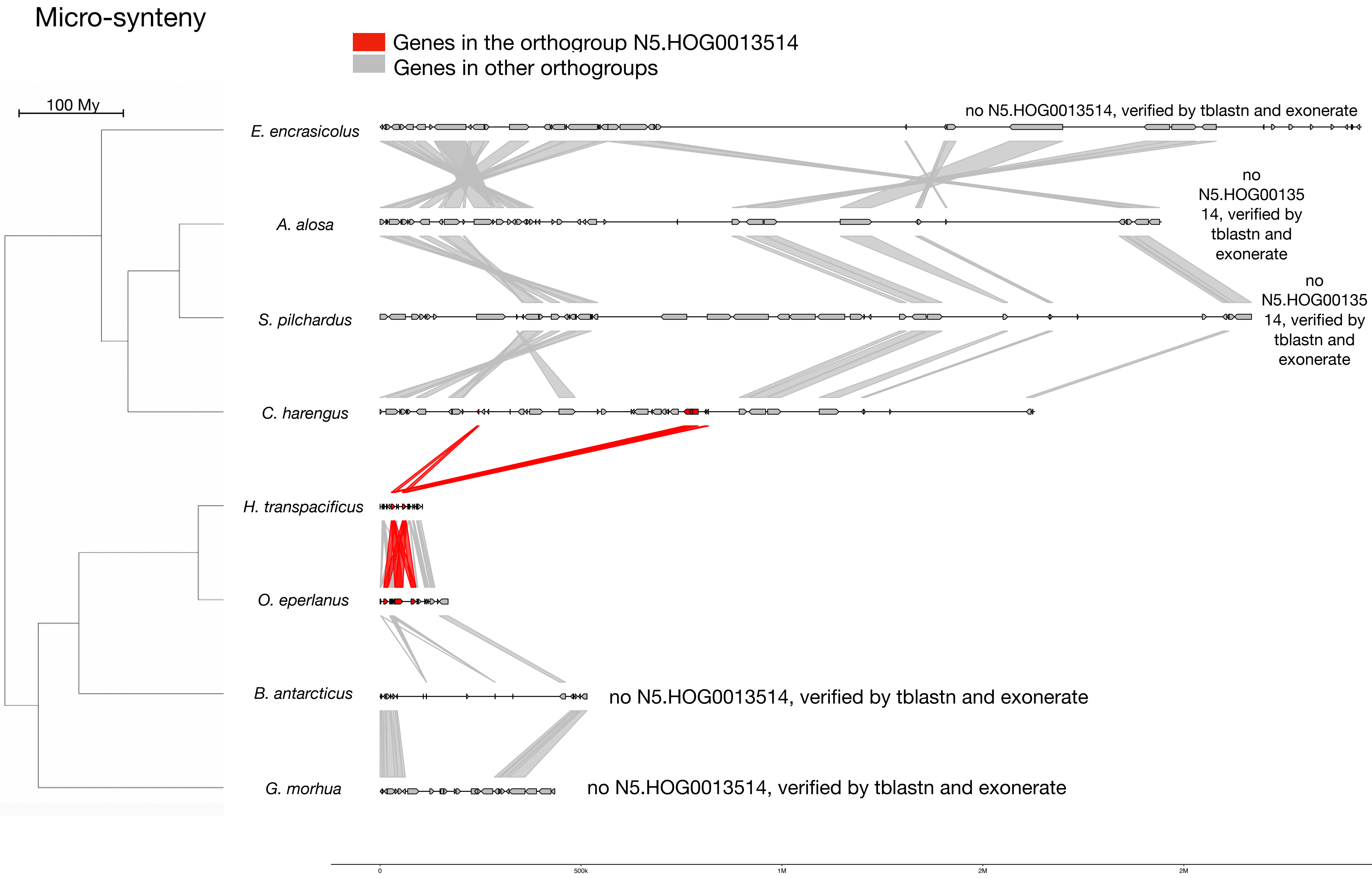


#### Zoom on *C. harengus* and *H. transpacificus*

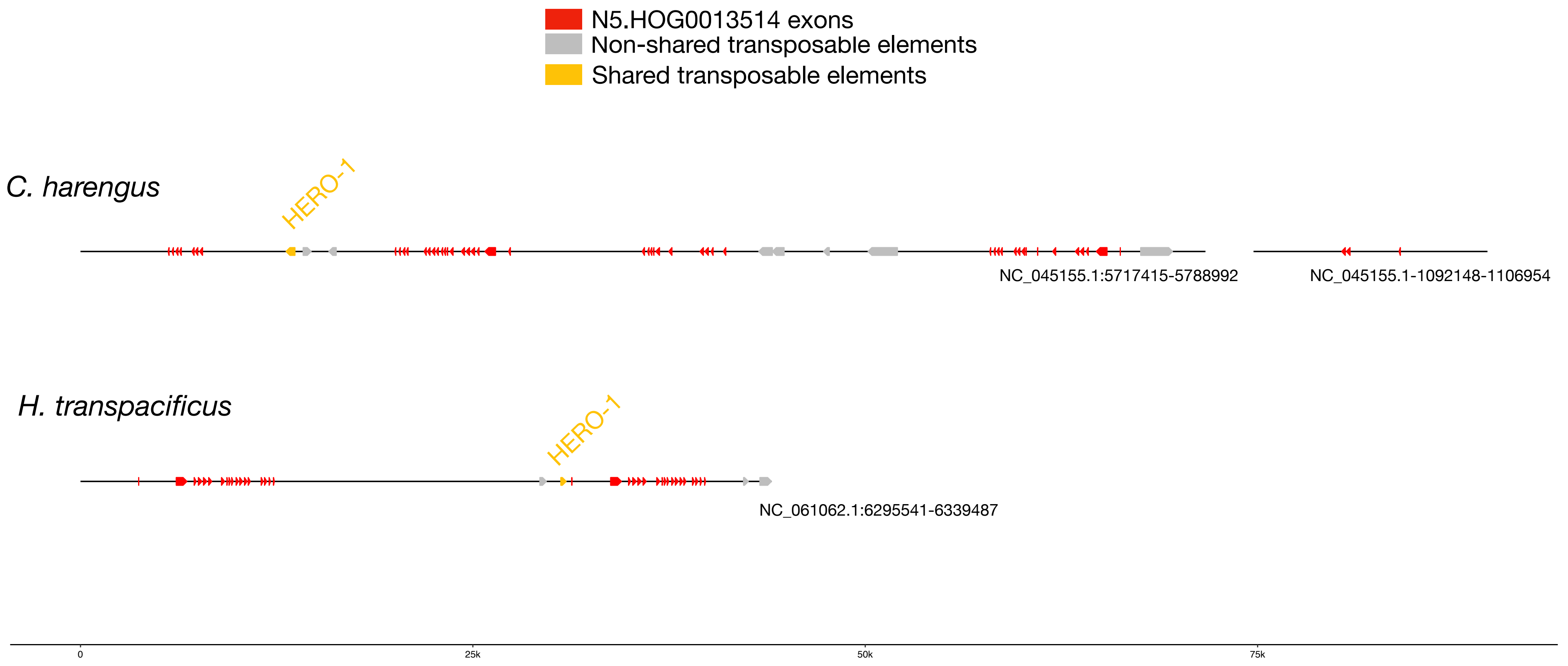


#### HERO-1 :

| | |
|---|---|
| *E. encrasicolus* | *741 copies scattered in the genome* |
| *A. alosa* | *410 copies scattered in the genome* |
| *S. pilchardus* | *266 copies scattered in the genome* |
| *C. harengus* | *1095 copies scattered in the genome* |
| *H. transpacificus* | *17 copies scattered in the genome* |
| *O. eperlanus* | *14 copies scattered in the genome* |
| *B. antarcticus* | *128 copies scattered in the genome* |
| *G. morhua* | *33 copies scattered in the genome* |

### N5.HOG0013514

#### Micro-synteny

##### Zoom on *C. harengus* and *H. transpacificus* (-5000/+5000 bp around N5.HOG0013514)

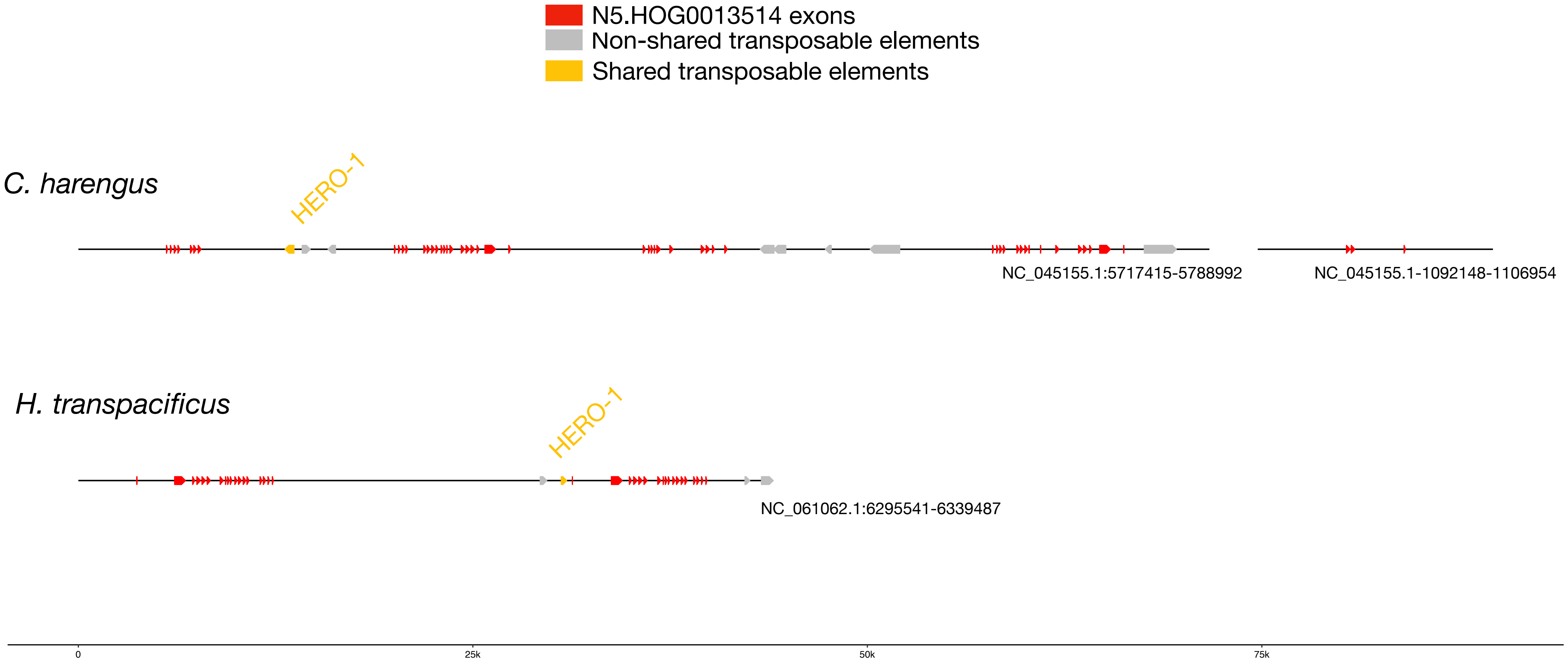


*Clupea harengus* genes are only partially annotated: some exons are missing in the annotations, but retrieved when performing an exonerate search. Exonerate allowed to retrieve complete sequences of N5.HOG0013514 in *C. harengus*

N5.HOG0013514 full length versions in *Clupea harengus*

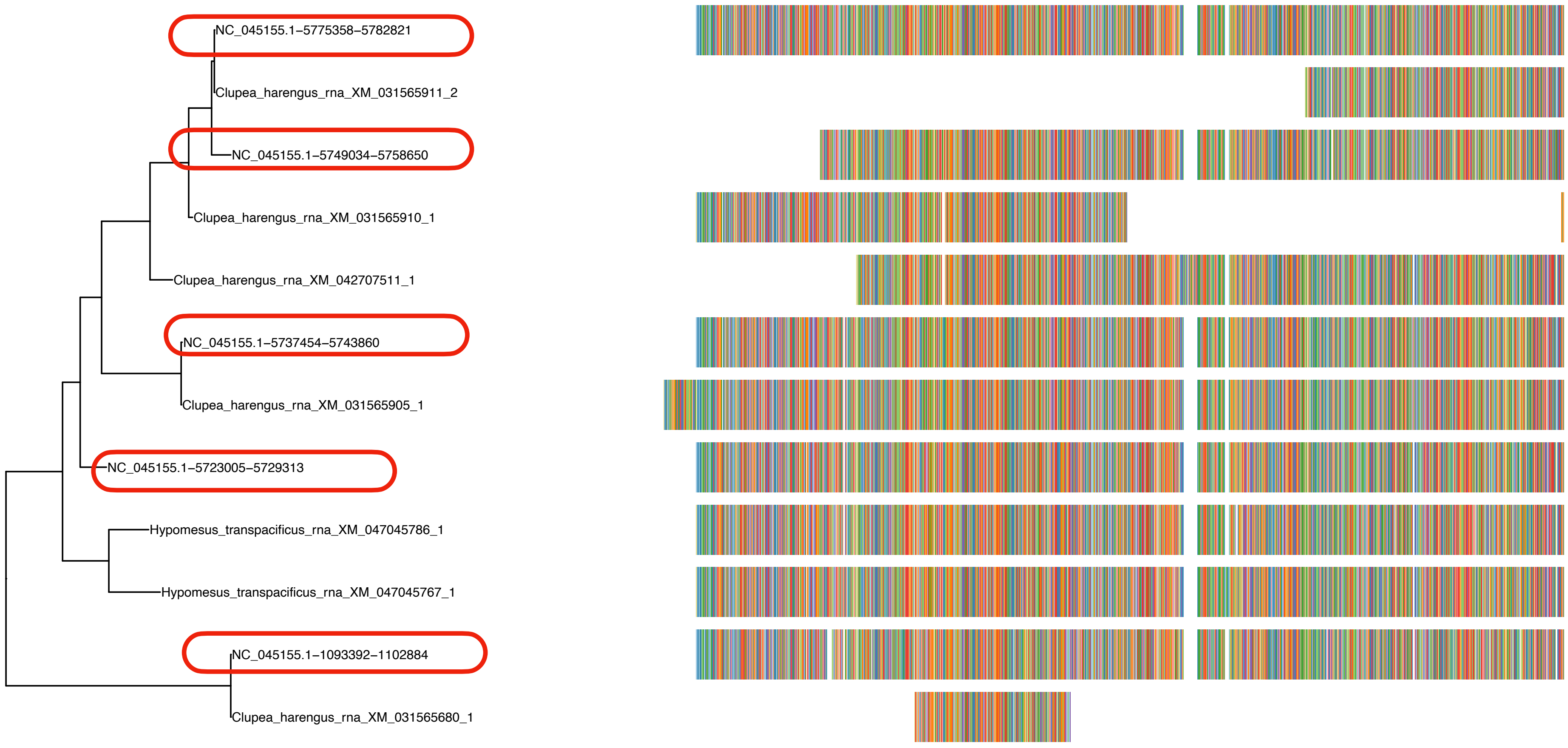

N5.HOG0013514

dS are similar with the full length versions of *C. harengus* genes

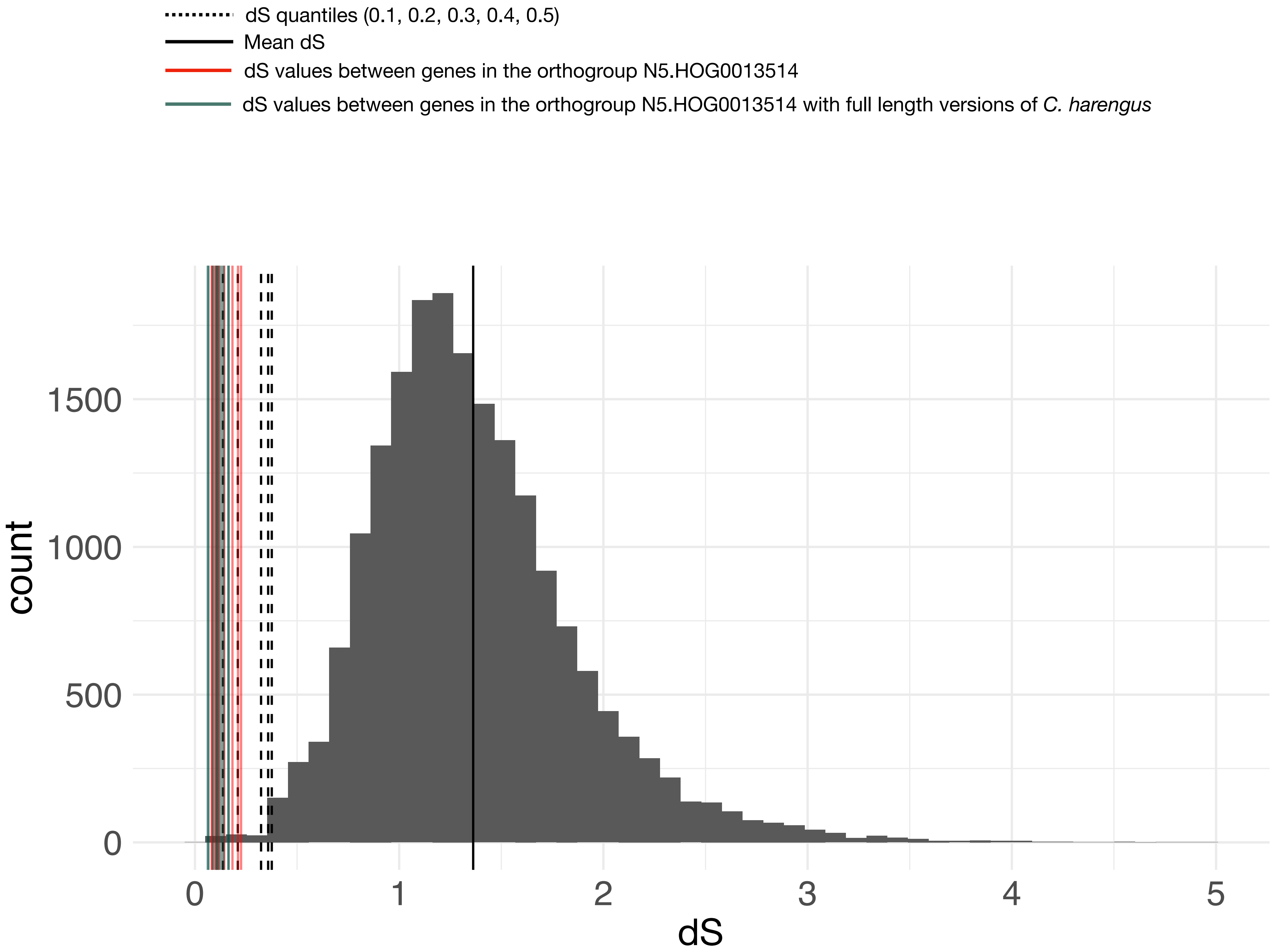

### N5.HOG0013514

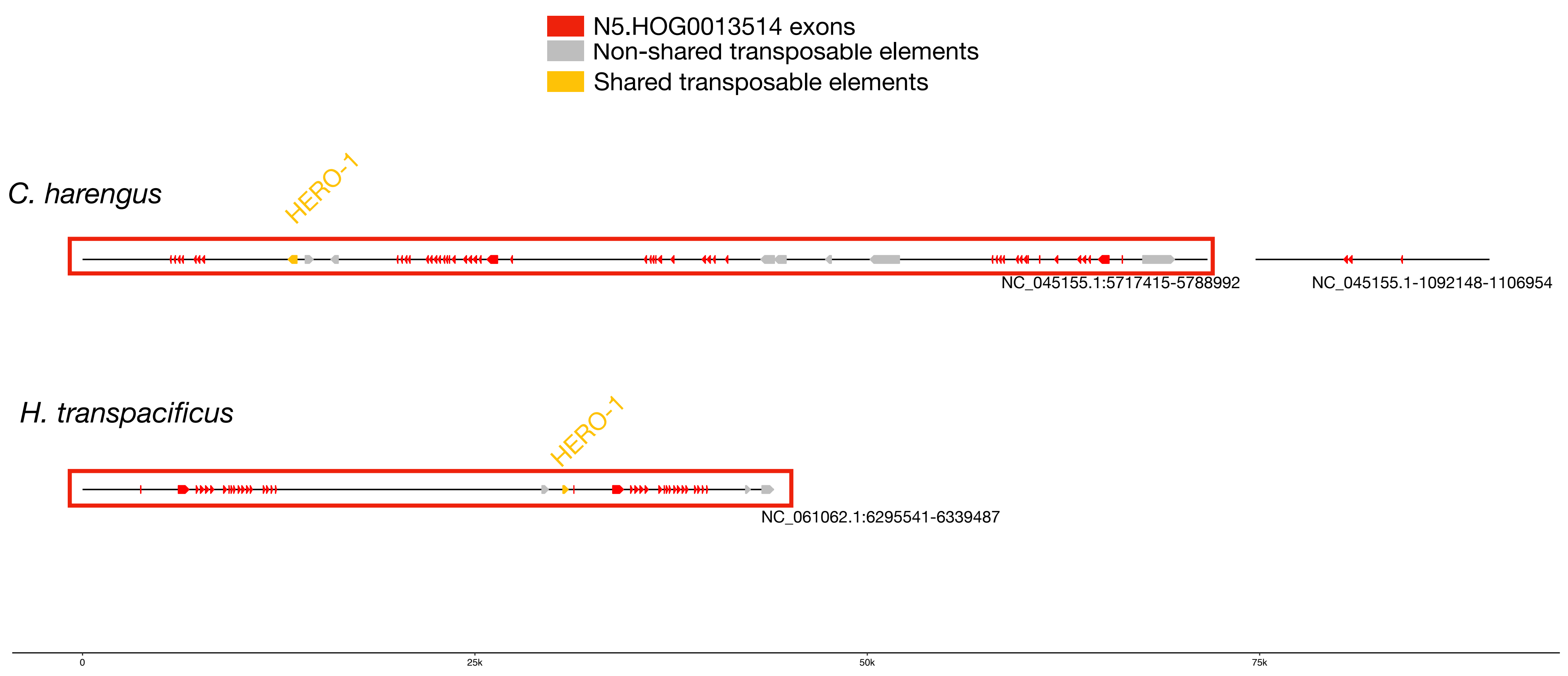


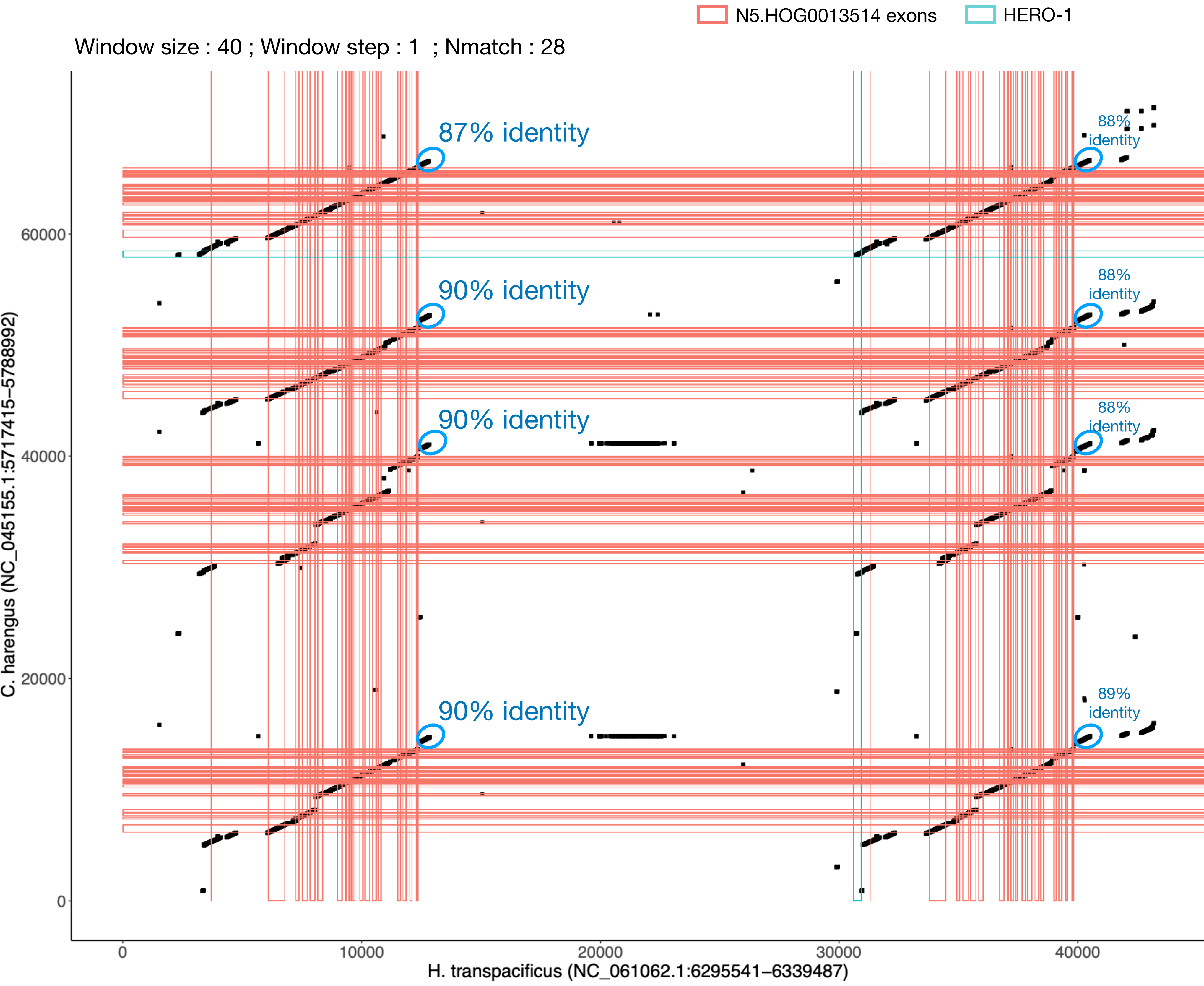

### N5.HOG0013514

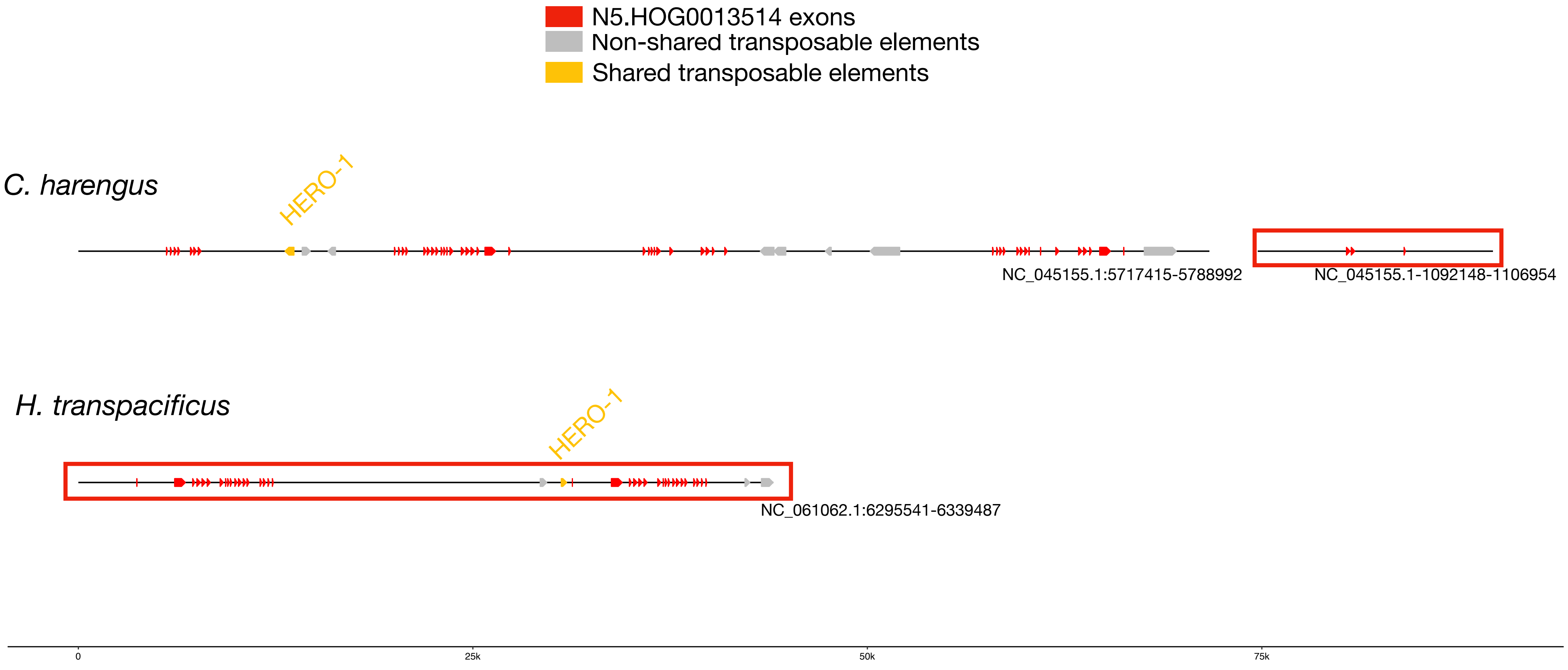


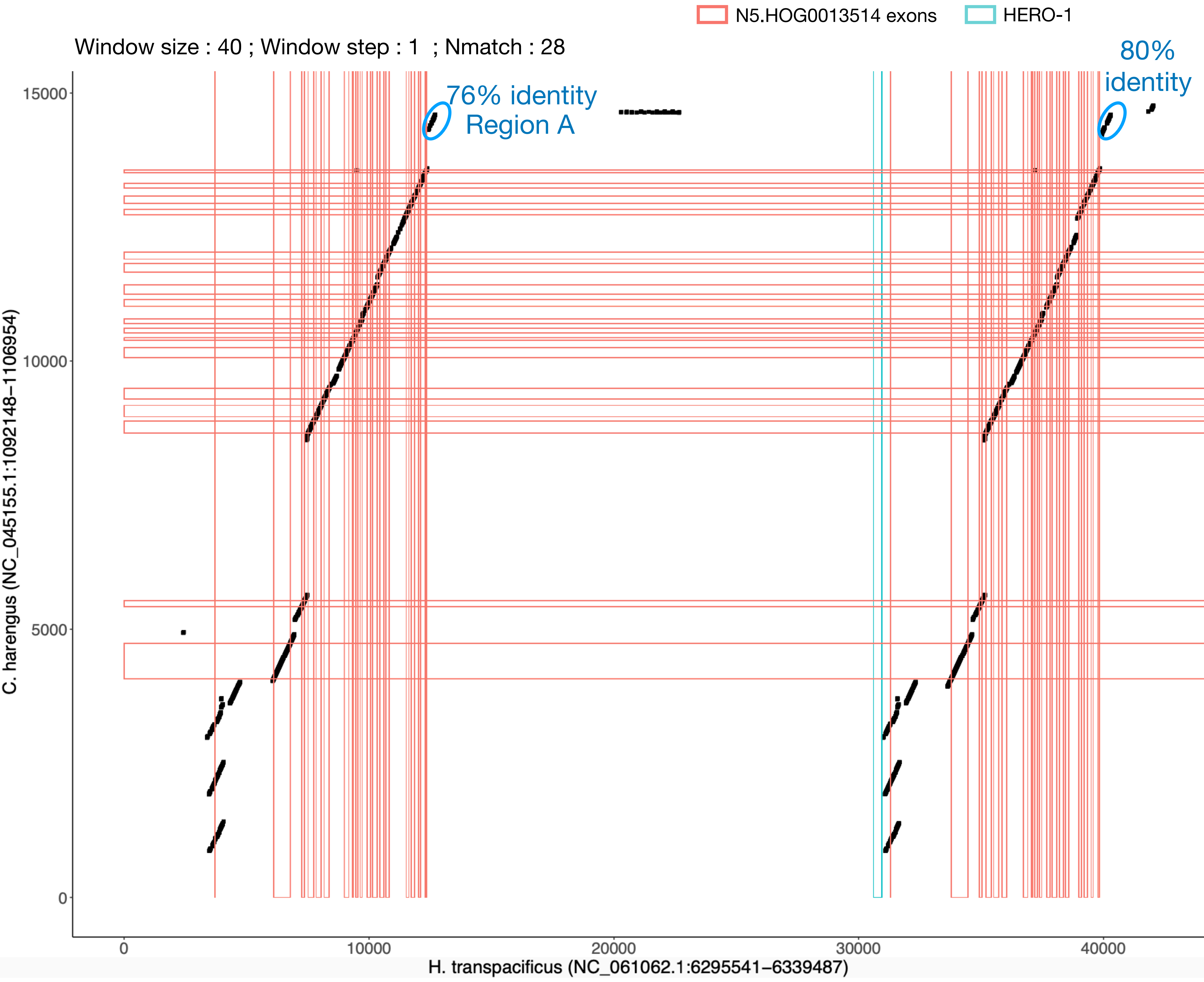

### N5.HOG0013514 Blastn of conserved regions against all genome assemblies

#### Region A

| sseqid | pident | length | mismatch | gapopen | qstart | qend | sstart | send | evalue | bitscore |
|---|---|---|---|---|---|---|---|---|---|---|
| Hypomesus_transpacificus-NC_061062.1 | 100.000 | 1001 | 0 | 0 | 1 | 1001 | 6307892 | 6308892 | 0.0 | 1849 |
| Hypomesus_transpacificus-NC_061062.1 | 94.694 | 490 | 16 | 4 | 520 | 1001 | 6346209 | 6345722 | 0.0 | 752 |
| Hypomesus_transpacificus-NC_061062.1 | 94.444 | 396 | 22 | 0 | 520 | 915 | 6353732 | 6353337 | 3.91e-170 | 610 |
| Hypomesus_transpacificus-NC_061062.1 | 90.435 | 460 | 21 | 7 | 82 | 521 | 6335549 | 6336005 | 2.37e-162 | 584 |
| Hypomesus_transpacificus-NC_061062.1 | 93.315 | 359 | 22 | 2 | 520 | 876 | 6344756 | 6344398 | 1.13e-145 | 529 |
| Hypomesus_transpacificus-NC_061062.1 | 91.304 | 391 | 26 | 5 | 520 | 909 | 6351640 | 6351257 | 4.05e-145 | 527 |
| Hypomesus_transpacificus-NC_061062.1 | 89.731 | 409 | 37 | 5 | 594 | 998 | 6273456 | 6273863 | 2.44e-142 | 518 |
| Hypomesus_transpacificus-NC_061062.1 | 92.201 | 359 | 23 | 5 | 520 | 876 | 6278623 | 6278978 | 6.82e-138 | 503 |
| Hypomesus_transpacificus-NC_061062.1 | 90.782 | 358 | 30 | 2 | 520 | 876 | 6309836 | 6310191 | 1.49e-129 | 475 |
| Hypomesus_transpacificus-NC_061062.1 | 88.515 | 357 | 22 | 5 | 520 | 876 | 6231525 | 6231188 | 3.29e-111 | 414 |
| Hypomesus_transpacificus-NC_061062.1 | 97.143 | 140 | 4 | 0 | 520 | 659 | 6295968 | 6296107 | 7.64e-58 | 237 |
| Hypomesus_transpacificus-NC_061062.1 | 95.714 | 140 | 6 | 0 | 520 | 659 | 6323794 | 6323933 | 1.65e-54 | 226 |
| Hypomesus_transpacificus-NC_061062.1 | 98.667 | 75 | 1 | 0 | 1 | 75 | 6335363 | 6335437 | 1.03e-26 | 134 |
| Hypomesus_transpacificus-NC_061062.1 | 97.059 | 68 | 2 | 0 | 520 | 587 | 6274986 | 6275053 | 3.73e-21 | 115 |
| Osmerus_eperlanus-NC_085026.1 | 90.062 | 483 | 36 | 8 | 520 | 1001 | 4898318 | 4897847 | 8.40e-172 | 616 |
| Osmerus_eperlanus-NC_085026.1 | 94.264 | 401 | 20 | 3 | 520 | 918 | 4900933 | 4900534 | 3.91e-170 | 610 |
| Osmerus_eperlanus-NC_085026.1 | 92.893 | 394 | 24 | 2 | 520 | 912 | 4903038 | 4902648 | 6.63e-158 | 569 |
| Osmerus_eperlanus-NC_085026.1 | 92.132 | 394 | 27 | 2 | 520 | 912 | 4896840 | 4896450 | 6.68e-153 | 553 |
| Osmerus_eperlanus-NC_085026.1 | 93.296 | 358 | 23 | 1 | 520 | 876 | 4805564 | 4805921 | 4.05e-145 | 527 |
| Osmerus_eperlanus-NC_085026.1 | 93.296 | 358 | 23 | 1 | 520 | 876 | 4842652 | 4843009 | 4.05e-145 | 527 |
| Osmerus_eperlanus-NC_085026.1 | 93.296 | 358 | 23 | 1 | 520 | 876 | 4874810 | 4875167 | 4.05e-145 | 527 |
| Osmerus_eperlanus-NC_085026.1 | 91.460 | 363 | 24 | 4 | 520 | 876 | 4729340 | 4728979 | 1.48e-134 | 492 |
| Osmerus_eperlanus-NC_085026.1 | 91.365 | 359 | 23 | 4 | 520 | 876 | 4783128 | 4783480 | 2.47e-132 | 484 |
| Osmerus_eperlanus-NC_085026.1 | 89.106 | 358 | 22 | 4 | 520 | 876 | 4774237 | 4774578 | 1.17e-115 | 429 |
| Osmerus_eperlanus-NC_085026.1 | 90.698 | 301 | 25 | 3 | 617 | 915 | 4731344 | 4731045 | 3.31e-106 | 398 |
| Osmerus_eperlanus-NC_085026.1 | 91.289 | 287 | 23 | 2 | 520 | 804 | 4894746 | 4894460 | 5.54e-104 | 390 |
| Osmerus_eperlanus-NC_085026.1 | 86.971 | 307 | 22 | 9 | 82 | 374 | 4856001 | 4856303 | 1.23e-85 | 329 |
| Osmerus_eperlanus-NC_085026.1 | 86.645 | 307 | 23 | 10 | 82 | 374 | 4823835 | 4824137 | 5.70e-84 | 324 |
| Osmerus_eperlanus-NC_085026.1 | 87.413 | 286 | 18 | 7 | 85 | 362 | 4801106 | 4801381 | 1.23e-80 | 313 |
| Osmerus_eperlanus-NC_085026.1 | 87.413 | 286 | 18 | 7 | 85 | 362 | 4838194 | 4838469 | 1.23e-80 | 313 |
| Osmerus_eperlanus-NC_085026.1 | 87.413 | 286 | 18 | 7 | 85 | 362 | 4870556 | 4870831 | 1.23e-80 | 313 |
| Hypomesus_transpacificus-NC_061078.1 | 89.706 | 408 | 33 | 5 | 594 | 998 | 15313803 | 15314204 | 1.13e-140 | 512 |
| Hypomesus_transpacificus-NC_061078.1 | 95.714 | 140 | 6 | 0 | 520 | 659 | 15328849 | 15328988 | 1.65e-54 | 226 |
| Hypomesus_transpacificus-NC_061078.1 | 93.571 | 140 | 9 | 0 | 520 | 659 | 15269402 | 15269263 | 1.67e-49 | 209 |
| Hypomesus_transpacificus-NC_061078.1 | 97.059 | 68 | 2 | 0 | 520 | 587 | 15315327 | 15315394 | 3.73e-21 | 115 |
| Clupea_harengus-NC_045155.1 | 85.381 | 472 | 32 | 8 | 84 | 519 | 5736799 | 5736329 | 1.94e-123 | 455 |
| Clupea_harengus-NC_045155.1 | 85.350 | 471 | 32 | 10 | 84 | 519 | 5774715 | 5774247 | 6.97e-123 | 453 |
| Clupea_harengus-NC_045155.1 | 83.624 | 458 | 33 | 19 | 85 | 504 | 5748377 | 5747924 | 1.54e-104 | 392 |
| Clupea_harengus-NC_045155.1 | 83.886 | 422 | 45 | 9 | 82 | 484 | 5722826 | 5722409 | 3.33e-101 | 381 |
| Clupea_harengus-NC_045155.1 | 83.010 | 206 | 14 | 12 | 200 | 393 | 109092 | 109288 | 1.02e-36 | 167 |
| Clupea_harengus-NC_045155.1 | 82.902 | 193 | 21 | 5 | 200 | 382 | 1092517 | 1092327 | 1.31e-35 | 163 |
| Clupea_harengus-NC_045155.1 | 82.090 | 201 | 23 | 8 | 204 | 393 | 23010068 | 23009870 | 1.70e-34 | 159 |
| Clupea_harengus-NC_045170.1 | 82.704 | 318 | 36 | 11 | 87 | 389 | 24193888 | 24193575 | 3.50e-66 | 265 |
| Clupea_harengus-NC_045170.1 | 82.222 | 315 | 33 | 9 | 87 | 382 | 24161302 | 24160992 | 9.81e-62 | 250 |
| Clupea_harengus-NC_045160.1 | 78.571 | 476 | 35 | 22 | 84 | 521 | 2043897 | 2043451 | 2.73e-62 | 252 |
| Clupea_harengus-NW_024880083.1 | 80.000 | 325 | 32 | 14 | 85 | 389 | 36064 | 35753 | 1.67e-49 | 209 |

N5.HOG0013514

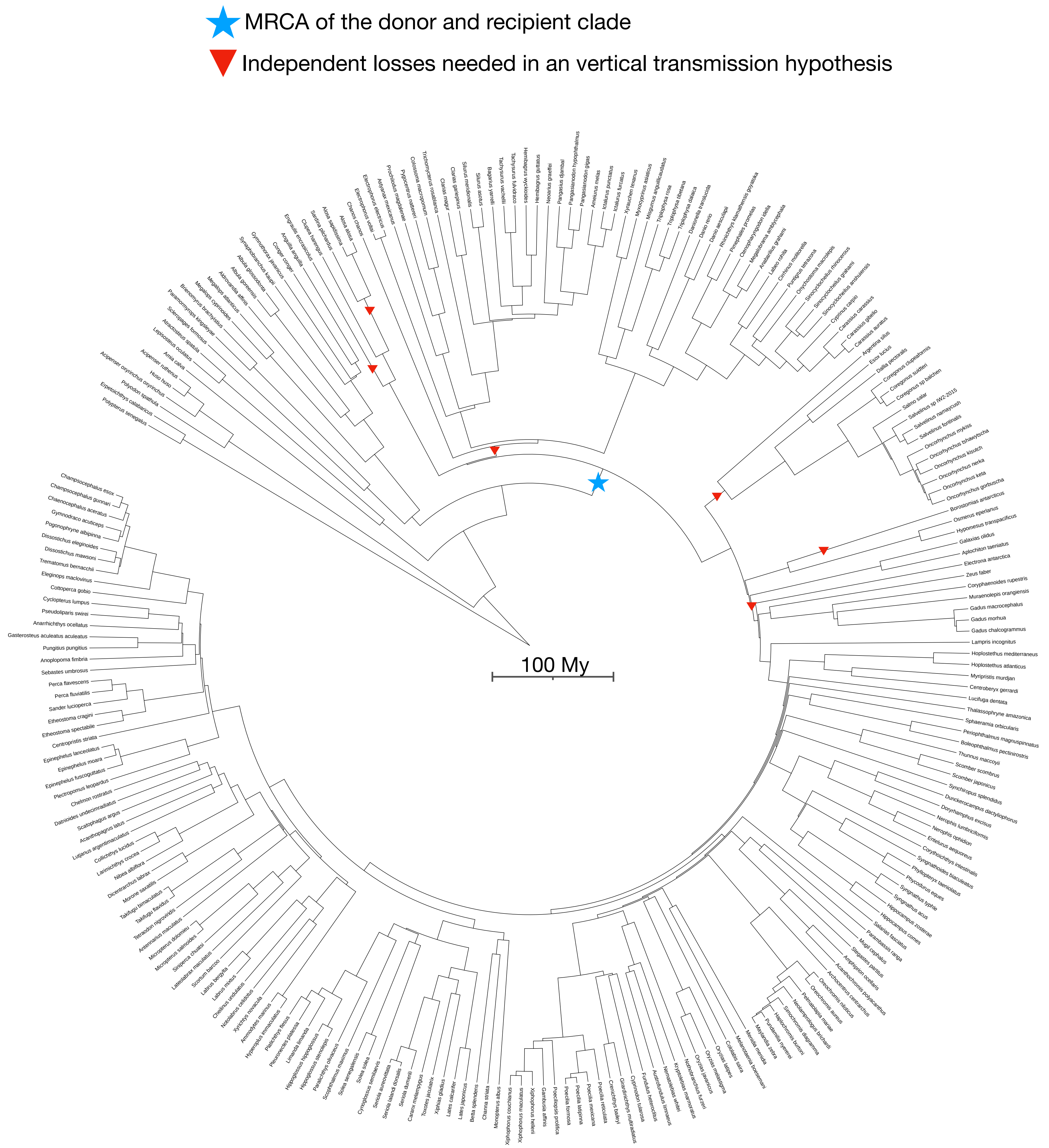

N5.HOG0018901

N5.HOG0018901

Species tree

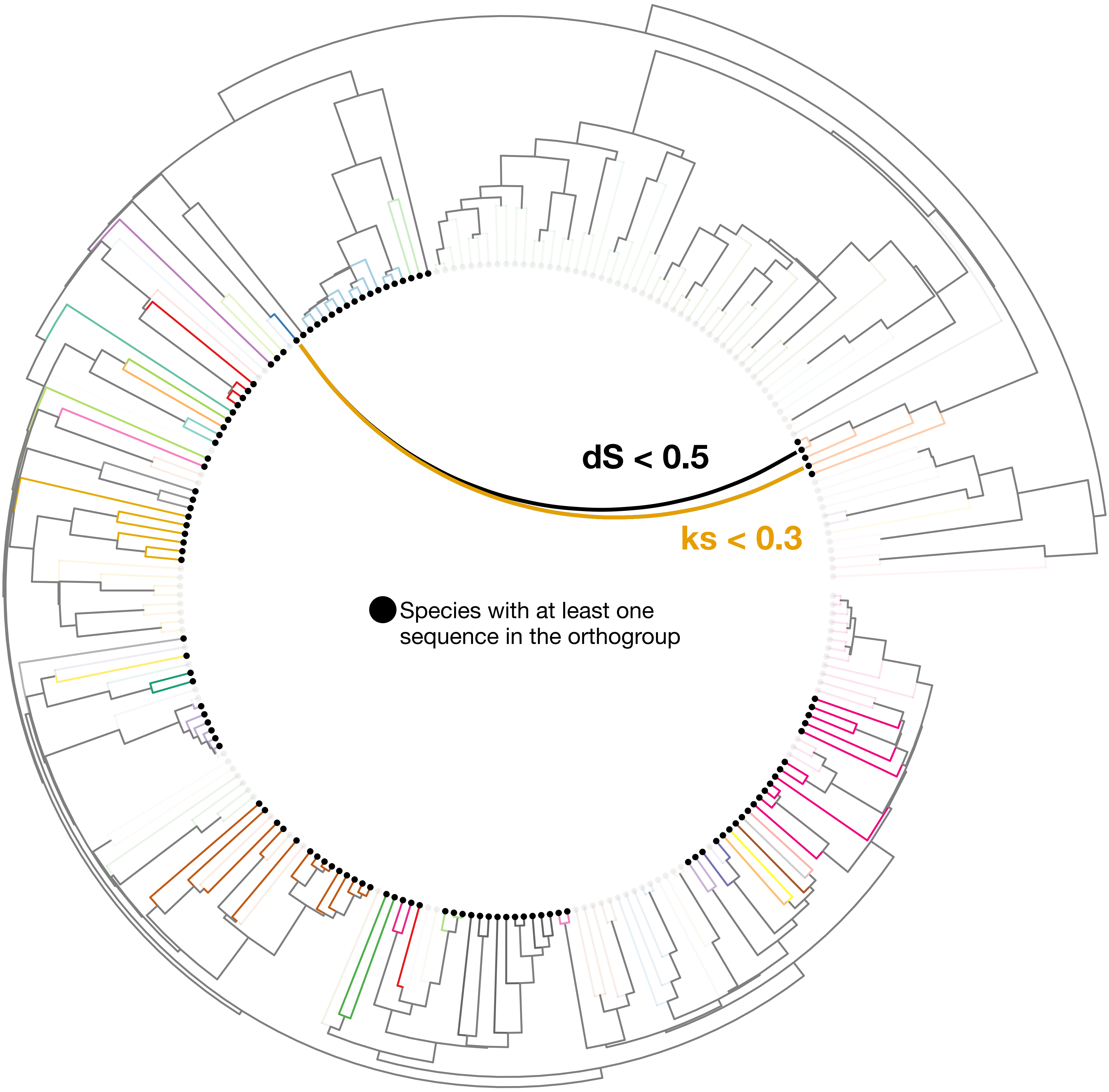

### N5.HOG0018901

Orthogroup phylogeny
and
protein alignment

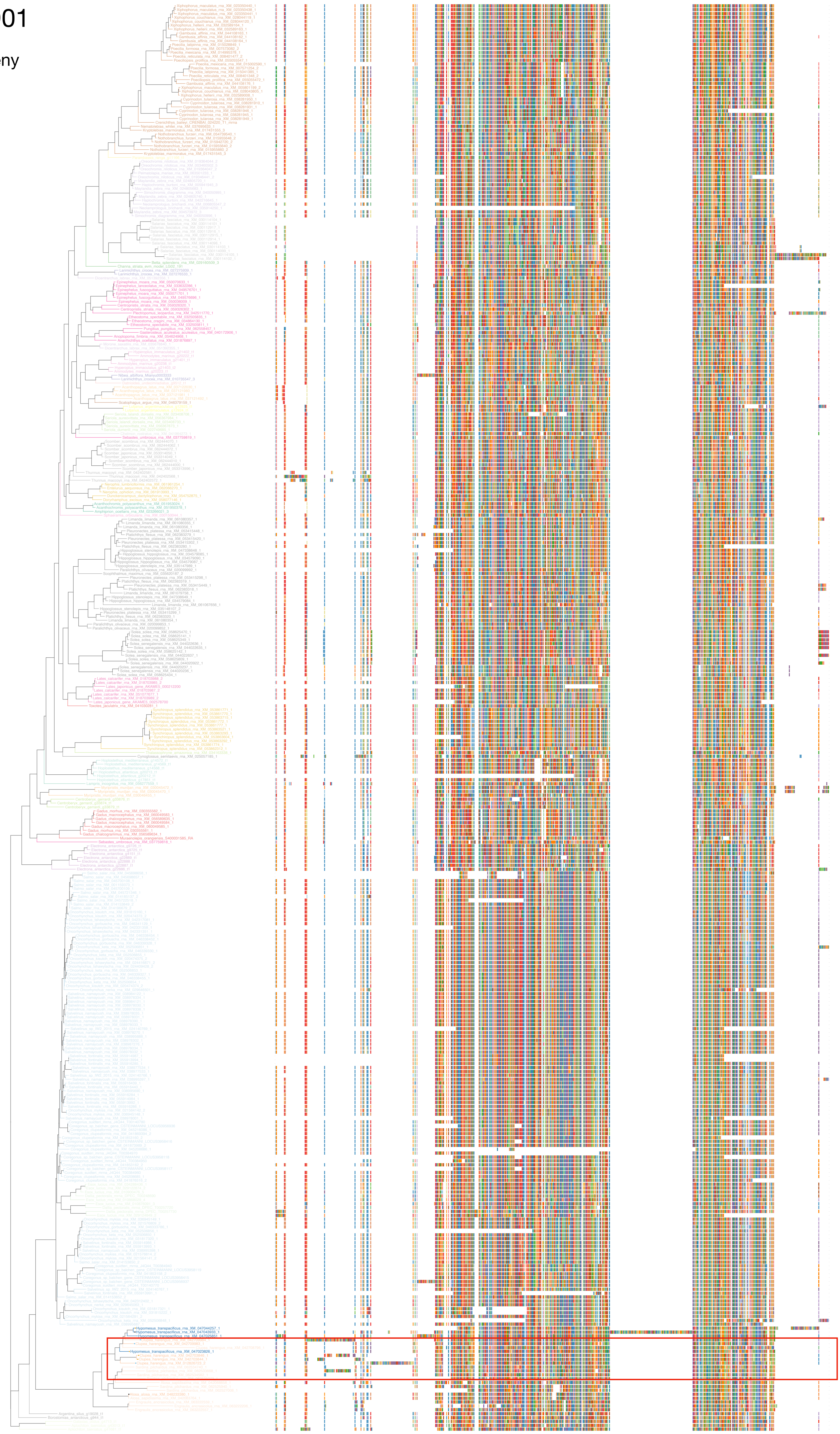

### N5.HOG0018901

Zoom on HGT clade with bootstrap values

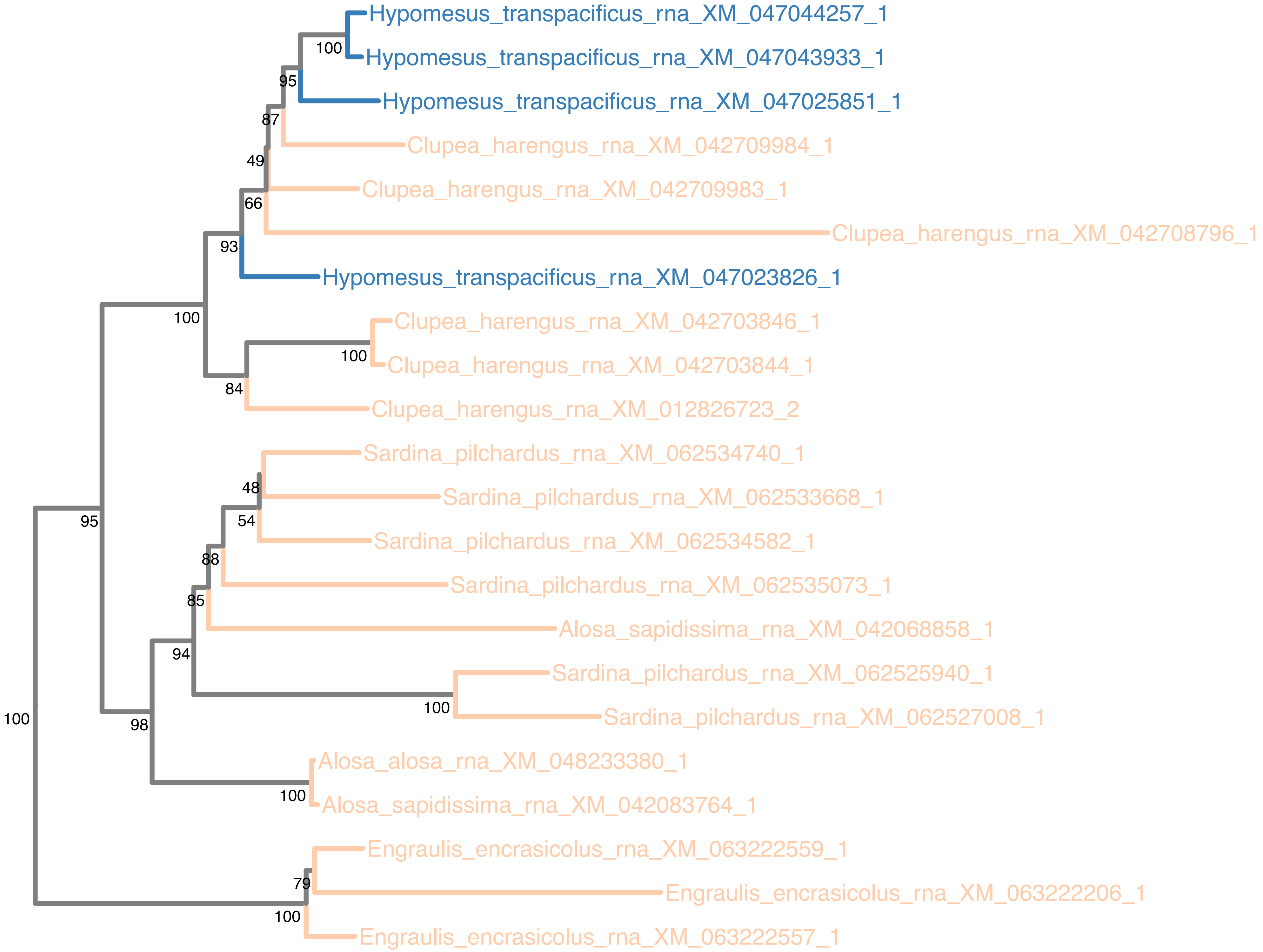

### N5.HOG0018901

Orthogroup phylogeny
and
trimmed
protein alignment

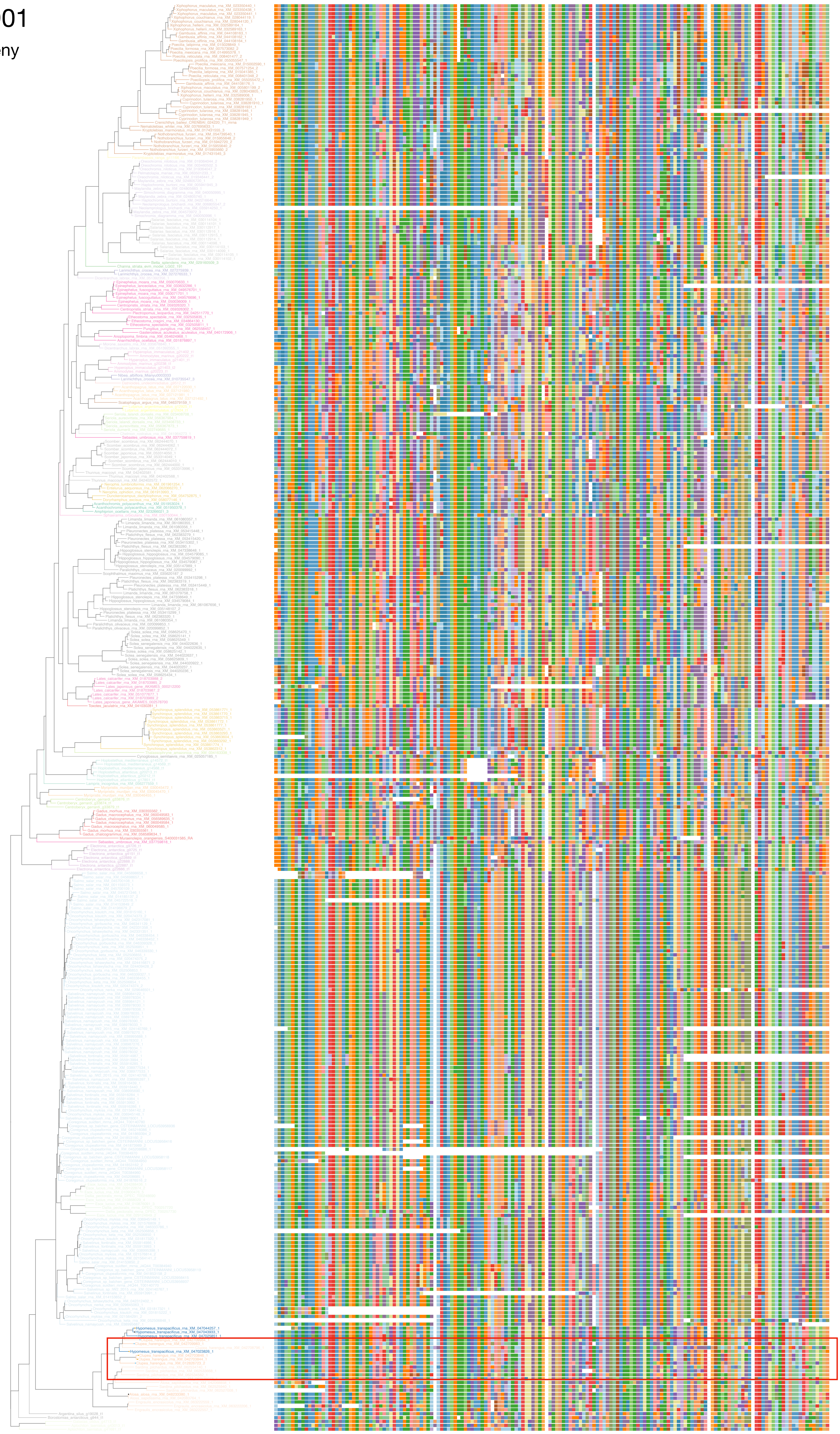

### N5.HOG0018901

« Best-match » phylogeny
and
protein alignment

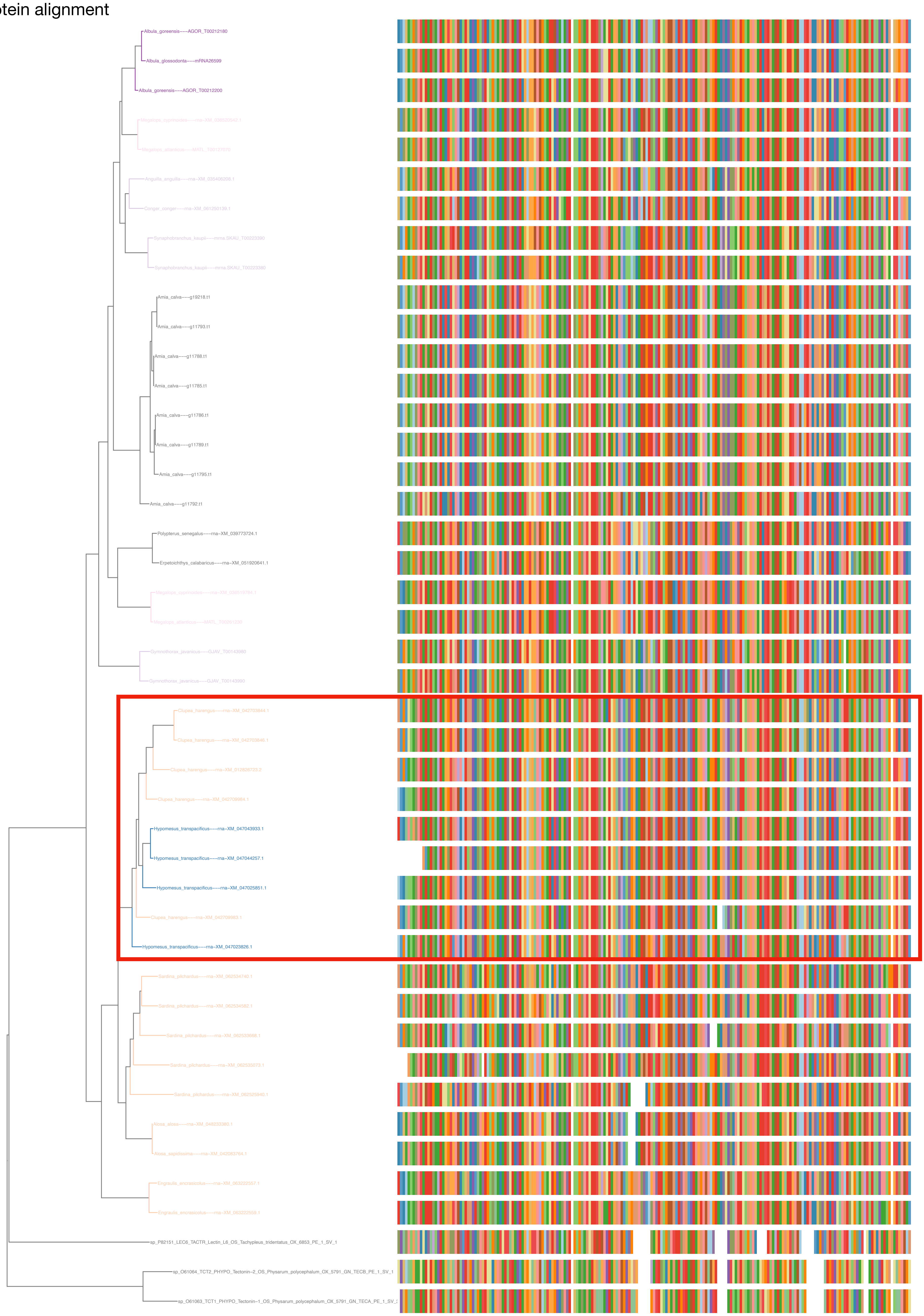

N5.HOG0018901

dS distribution

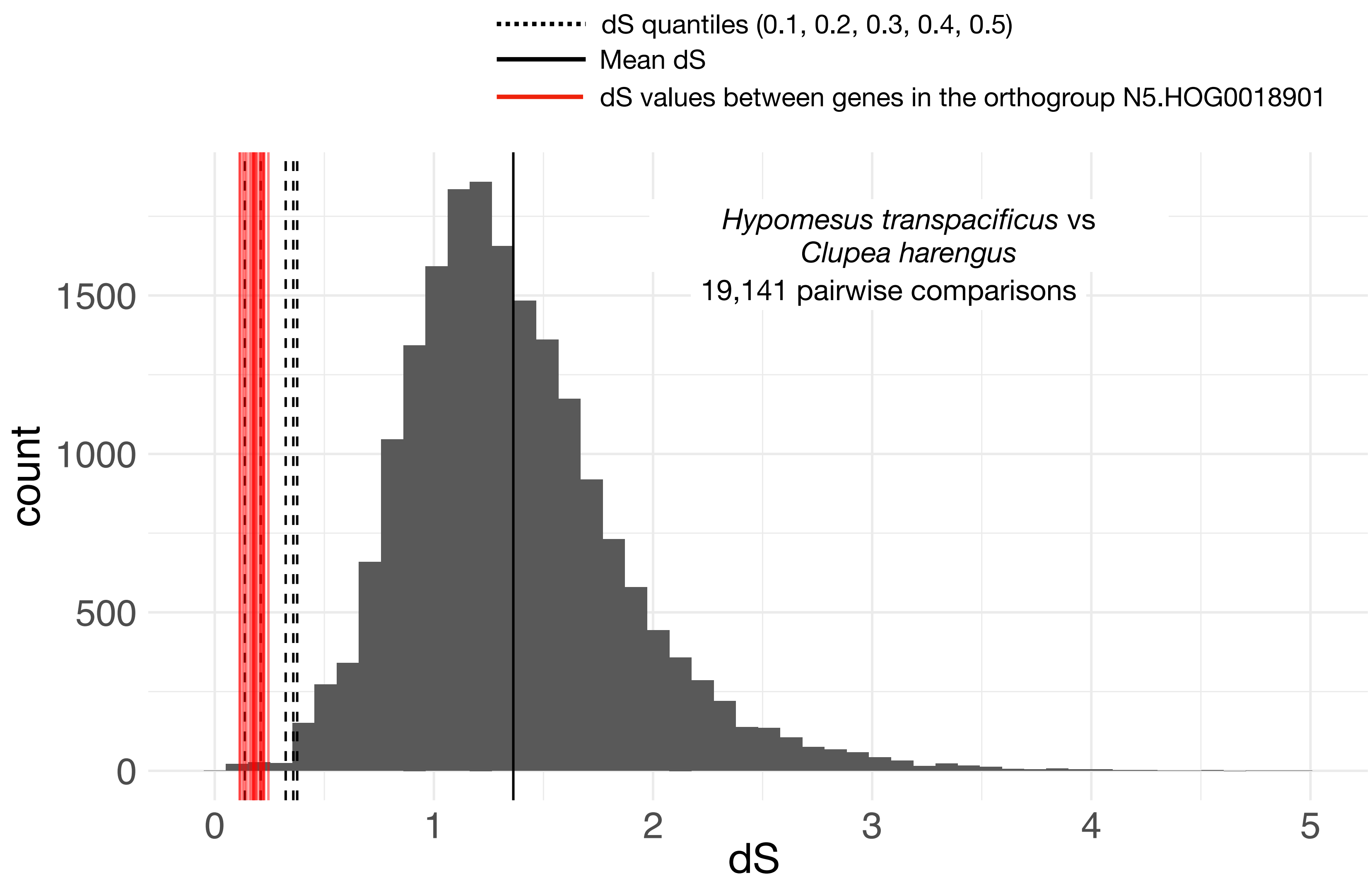


Intra-orthogroup Ks

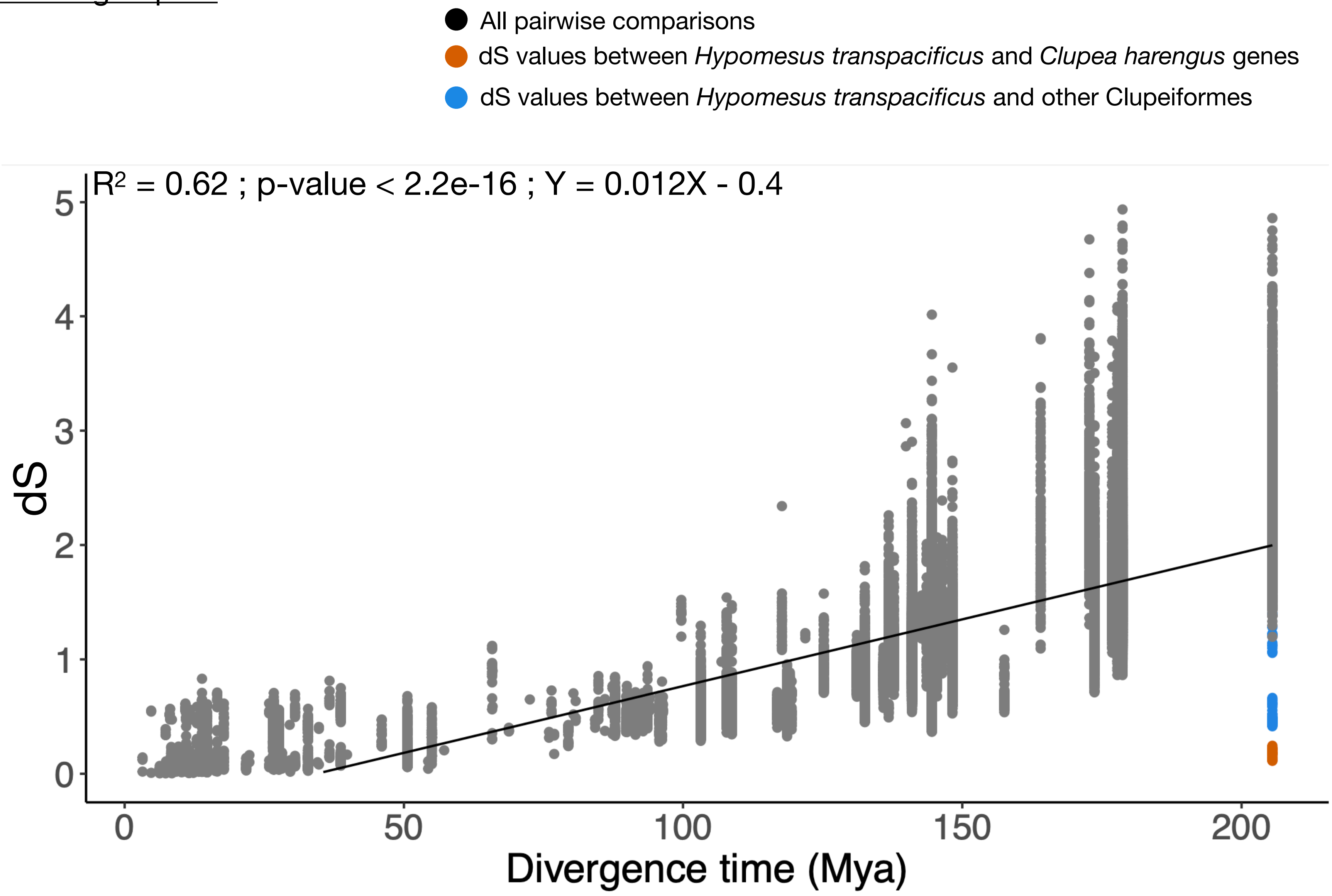

### N5.HOG0018901

#### Micro-synteny

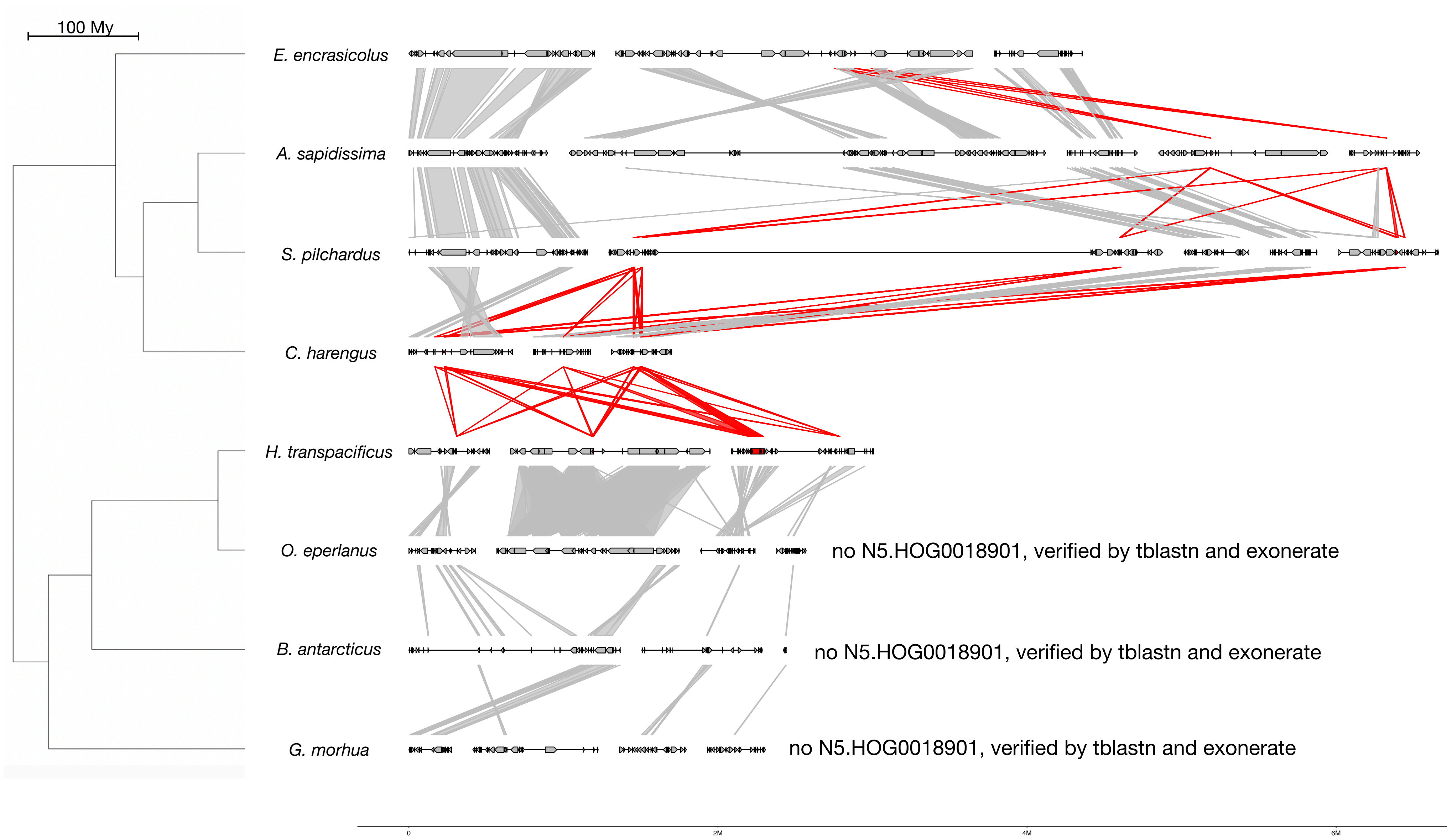


#### Zoom on *C. harengus* and *H. transpacificus*

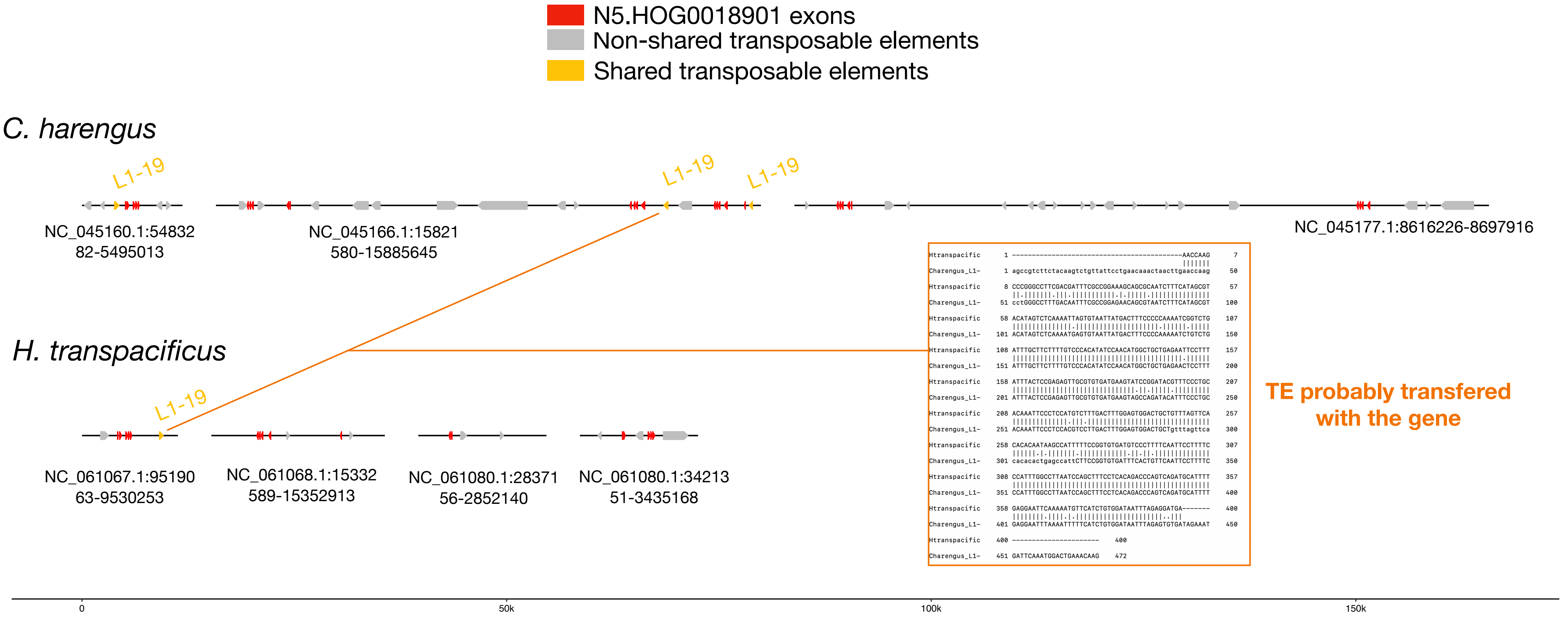


## L1-19 :

| | |
|---|---|
| *E. encrasicolus* | *86 copies scattered in the genome* |
| *A. sapidissima* | *25 copies scattered in the genome* |
| *S. pilchardus* | *30 copies scattered in the genome* |
| *C. harengus* | *31 copies scattered in the genome* |
| *H. transpacificus* | *1 copy upstream N5.HOG0018901* |
| *O. eperlanus* | *10 copies scattered in the genome* |
| *B. antarcticus* | *451 copies scattered in the genome* |
| *G. morhua* | *54 copies scattered in the genome* |

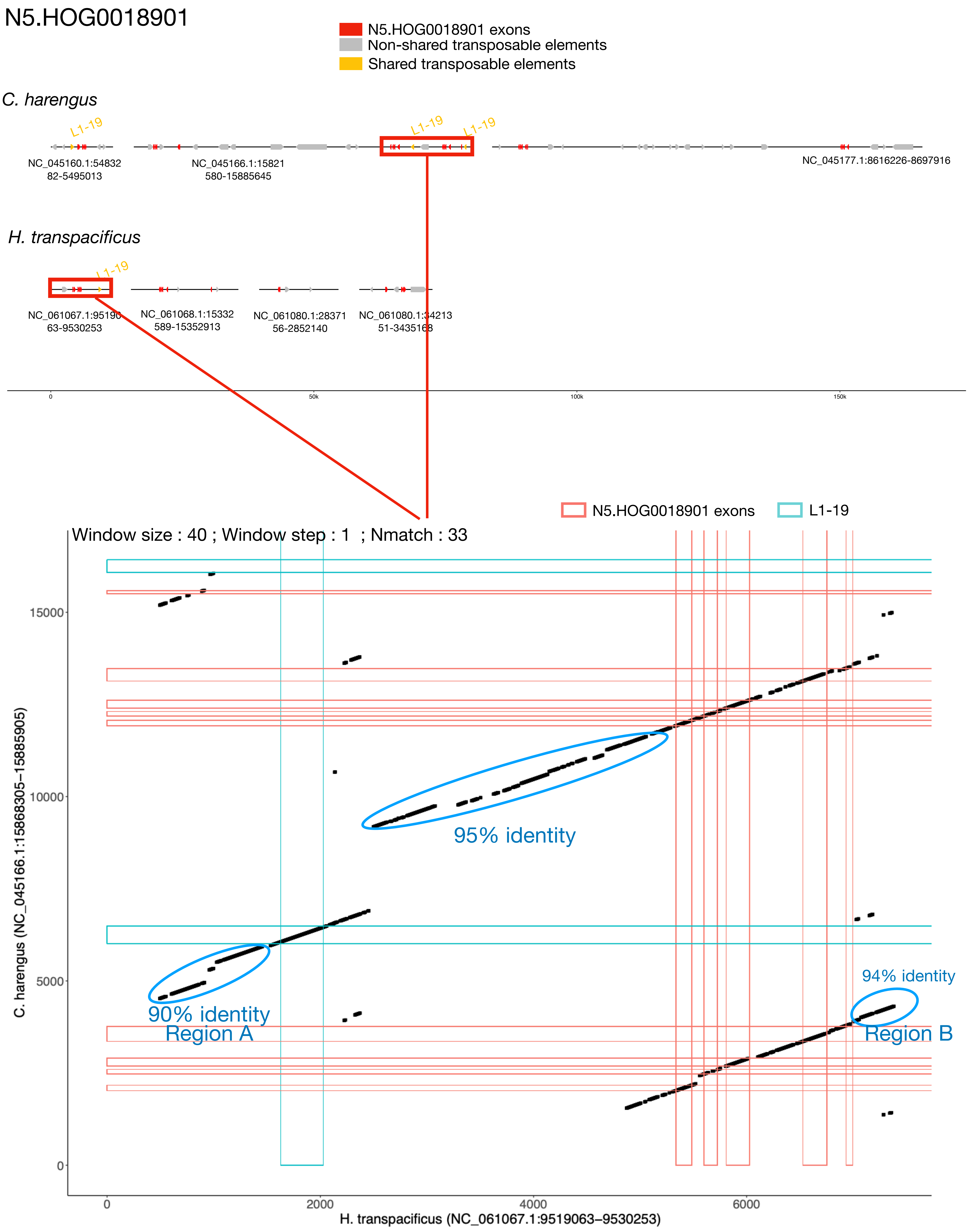

N5.HOG0018901
N5.HOG0018901 exons
Non-shared transposable elements
Shared transposable elements
C. harengus
L1-19
NC_045160.1:5483282-5495013
NC_045166.1:15821580-15885645
NC_045177.1:8616226-8697916
H. transpacificus
NC_061067.1:9519063-9530253
NC_061068.1:15332589-15352913
NC_061080.1:2837156-2852140
NC_061080.1:3421351-3435168
0
50k
100k
150k
N5.HOG0018901 exons
L1-19
Window size : 40 ; Window step : 1 ; Nmatch : 33
95% identity
90% identity
Region A
94% identity
Region B
C. harengus (NC_045166.1:15868305–15885905)
H. transpacificus (NC_061067.1:9519063–9530253)
15000
10000
5000
0
2000
4000
6000

### N5.HOG0018901 Blastn of conserved regions against all genome assemblies

#### Region A

| sseqid | pident | length | mismatch | gapopen | qstart | qend | sstart | send | evalue | bitscore |
|---|---|---|---|---|---|---|---|---|---|---|
| Hypomesus_transpacificus-NC_061067.1 | 100.000 | 1129 | 0 | 0 | 1 | 1129 | 9529754 | 9528626 | 0.0 | 2085 |
| Clupea_harengus-NC_045166.1 | 93.187 | 455 | 30 | 1 | 538 | 992 | 15873820 | 15874273 | 0.0 | 667 |
| Clupea_harengus-NC_045166.1 | 92.375 | 459 | 23 | 7 | 2 | 448 | 15872835 | 15873293 | 4.36e-180 | 643 |
| Clupea_harengus-NC_045166.1 | 84.255 | 470 | 54 | 13 | 672 | 1129 | 18134856 | 18134395 | 6.14e-119 | 440 |
| Clupea_harengus-NC_045166.1 | 82.129 | 498 | 56 | 13 | 641 | 1129 | 13026043 | 13025570 | 1.35e-105 | 396 |
| Clupea_harengus-NC_045166.1 | 82.129 | 498 | 56 | 13 | 641 | 1129 | 13029473 | 13029000 | 1.35e-105 | 396 |
| Clupea_harengus-NC_045166.1 | 82.863 | 461 | 37 | 21 | 2 | 448 | 15883519 | 15883951 | 1.76e-99 | 375 |
| Clupea_harengus-NC_045166.1 | 80.097 | 412 | 62 | 14 | 732 | 1129 | 17348054 | 17348459 | 2.35e-73 | 289 |
| Clupea_harengus-NC_045166.1 | 93.860 | 114 | 7 | 0 | 1016 | 1129 | 7304491 | 7304604 | 2.47e-38 | 172 |
| Clupea_harengus-NC_045166.1 | 97.753 | 89 | 2 | 0 | 1041 | 1129 | 15874275 | 15874363 | 8.95e-33 | 154 |
| Clupea_harengus-NC_045170.1 | 89.961 | 508 | 41 | 5 | 629 | 1129 | 25842221 | 25841717 | 0.0 | 647 |
| Clupea_harengus-NC_045170.1 | 85.556 | 450 | 50 | 9 | 689 | 1129 | 592411 | 591968 | 6.09e-124 | 457 |
| Clupea_harengus-NC_045170.1 | 84.255 | 470 | 53 | 14 | 672 | 1129 | 19218528 | 19218068 | 2.21e-118 | 438 |
| Clupea_harengus-NC_045170.1 | 84.043 | 470 | 53 | 14 | 672 | 1129 | 22258789 | 22258330 | 1.03e-116 | 433 |
| Clupea_harengus-NC_045170.1 | 79.918 | 488 | 71 | 16 | 662 | 1128 | 26513345 | 26512864 | 1.07e-86 | 333 |
| Clupea_harengus-NC_045170.1 | 92.661 | 109 | 8 | 0 | 1021 | 1129 | 1102117 | 1102009 | 6.92e-34 | 158 |
| Clupea_harengus-NC_045165.1 | 93.703 | 397 | 24 | 1 | 733 | 1129 | 8690704 | 8691099 | 4.45e-165 | 593 |
| Clupea_harengus-NC_045165.1 | 85.141 | 498 | 59 | 8 | 641 | 1129 | 13277792 | 13277301 | 1.29e-135 | 496 |
| Clupea_harengus-NC_045165.1 | 85.141 | 498 | 59 | 8 | 641 | 1129 | 13281220 | 13280729 | 1.29e-135 | 496 |
| Clupea_harengus-NC_045160.1 | 86.891 | 534 | 45 | 17 | 612 | 1129 | 21780708 | 21780184 | 1.61e-159 | 575 |
| Clupea_harengus-NC_045160.1 | 83.144 | 528 | 60 | 15 | 612 | 1129 | 30145162 | 30144654 | 2.19e-123 | 455 |
| Clupea_harengus-NC_045160.1 | 87.871 | 371 | 38 | 6 | 759 | 1129 | 11575344 | 11575707 | 1.33e-115 | 429 |
| Clupea_harengus-NC_045160.1 | 81.853 | 529 | 51 | 28 | 612 | 1129 | 10041066 | 10041560 | 8.05e-108 | 403 |
| Clupea_harengus-NC_045160.1 | 77.719 | 377 | 27 | 31 | 755 | 1129 | 30467238 | 30467559 | 5.31e-40 | 178 |
| Clupea_harengus-NC_045173.1 | 90.796 | 402 | 30 | 7 | 729 | 1129 | 22666974 | 22667369 | 3.54e-146 | 531 |
| Clupea_harengus-NC_045173.1 | 84.951 | 412 | 36 | 11 | 731 | 1129 | 20499657 | 20499259 | 4.85e-105 | 394 |
| Clupea_harengus-NC_045173.1 | 80.414 | 531 | 62 | 21 | 611 | 1129 | 5151390 | 5151890 | 1.06e-96 | 366 |
| Clupea_harengus-NC_045172.1 | 86.694 | 481 | 50 | 12 | 612 | 1088 | 7472984 | 7472514 | 2.13e-143 | 521 |
| Clupea_harengus-NC_045172.1 | 78.277 | 534 | 67 | 25 | 609 | 1129 | 14564461 | 14564958 | 3.91e-76 | 298 |
| Clupea_harengus-NC_045172.1 | 87.283 | 173 | 18 | 3 | 204 | 374 | 3173729 | 3173899 | 5.28e-45 | 195 |
| Clupea_harengus-NC_045158.1 | 84.457 | 534 | 59 | 14 | 608 | 1129 | 25900799 | 25900278 | 2.15e-138 | 505 |
| Clupea_harengus-NC_045158.1 | 83.083 | 532 | 68 | 17 | 611 | 1129 | 15764655 | 15764133 | 3.64e-126 | 464 |
| Clupea_harengus-NC_045158.1 | 83.166 | 499 | 56 | 12 | 641 | 1129 | 29523932 | 29524412 | 3.69e-116 | 431 |
| Clupea_harengus-NC_045158.1 | 82.966 | 499 | 58 | 11 | 641 | 1129 | 29520711 | 29521192 | 1.72e-114 | 425 |
| Clupea_harengus-NC_045158.1 | 81.633 | 539 | 70 | 19 | 611 | 1129 | 9360451 | 9360980 | 8.00e-113 | 420 |
| Clupea_harengus-NC_045158.1 | 81.132 | 530 | 70 | 20 | 612 | 1129 | 13838374 | 13837863 | 3.75e-106 | 398 |
| Clupea_harengus-NC_045158.1 | 80.377 | 530 | 62 | 21 | 612 | 1129 | 9404490 | 9404989 | 3.80e-96 | 364 |
| Clupea_harengus-NC_045158.1 | 80.074 | 271 | 27 | 16 | 612 | 860 | 10296382 | 10296647 | 1.91e-39 | 176 |
| Clupea_harengus-NC_045158.1 | 87.755 | 147 | 7 | 5 | 731 | 867 | 27311991 | 27311846 | 5.35e-35 | 161 |
| Clupea_harengus-NC_045158.1 | 89.831 | 118 | 10 | 2 | 1012 | 1129 | 13301840 | 13301955 | 1.16e-31 | 150 |
| Clupea_harengus-NC_045158.1 | 95.238 | 84 | 4 | 0 | 1046 | 1129 | 9400749 | 9400832 | 1.17e-26 | 134 |
| Clupea_harengus-NC_045158.1 | 94.048 | 84 | 5 | 0 | 1046 | 1129 | 30219079 | 30218996 | 5.43e-25 | 128 |
| Clupea_harengus-NW_024879608.1 | 84.328 | 536 | 58 | 15 | 608 | 1129 | 144790 | 145313 | 2.78e-137 | 501 |
| Clupea_harengus-NW_024879593.1 | 85.481 | 489 | 56 | 8 | 650 | 1129 | 167561 | 168043 | 1.29e-135 | 496 |
| Clupea_harengus-NC_045154.1 | 85.141 | 498 | 59 | 8 | 641 | 1129 | 13488756 | 13489247 | 1.29e-135 | 496 |
| Clupea_harengus-NC_045154.1 | 85.846 | 325 | 37 | 7 | 808 | 1129 | 24919410 | 24919728 | 8.28e-88 | 337 |
| Clupea_harengus-NW_024879761.1 | 83.521 | 534 | 64 | 14 | 608 | 1129 | 22447 | 22968 | 4.68e-130 | 477 |
| Clupea_harengus-NC_045152.1 | 84.728 | 478 | 57 | 9 | 662 | 1129 | 10700800 | 10701271 | 3.64e-126 | 464 |
| Clupea_harengus-NC_045152.1 | 84.255 | 470 | 53 | 14 | 672 | 1129 | 17212998 | 17213458 | 2.21e-118 | 438 |
| Clupea_harengus-NW_024879990.1 | 84.342 | 479 | 58 | 10 | 662 | 1129 | 41824 | 41352 | 7.88e-123 | 453 |
| Clupea_harengus-NC_045163.1 | 82.739 | 533 | 69 | 17 | 610 | 1129 | 6591359 | 6591881 | 7.88e-123 | 453 |
| Clupea_harengus-NC_045163.1 | 82.093 | 497 | 55 | 14 | 641 | 1129 | 22823320 | 22823790 | 4.85e-105 | 394 |
| Clupea_harengus-NC_045163.1 | 82.093 | 497 | 55 | 14 | 641 | 1129 | 22926623 | 22926153 | 4.85e-105 | 394 |
| Clupea_harengus-NC_045163.1 | 85.714 | 245 | 24 | 9 | 890 | 1129 | 2417287 | 2417049 | 3.99e-61 | 248 |
| Clupea_harengus-NC_045171.1 | 82.707 | 532 | 66 | 21 | 611 | 1129 | 22189213 | 22188695 | 1.02e-121 | 449 |
| Clupea_harengus-NC_045171.1 | 84.043 | 470 | 53 | 14 | 672 | 1129 | 19106061 | 19105602 | 1.03e-116 | 433 |
| Clupea_harengus-NC_045171.1 | 84.043 | 470 | 53 | 14 | 672 | 1129 | 19109442 | 19108983 | 1.03e-116 | 433 |
| Clupea_harengus-NC_045171.1 | 85.993 | 307 | 23 | 7 | 731 | 1037 | 12016004 | 12015718 | 5.02e-80 | 311 |
| Clupea_harengus-NC_045161.1 | 84.468 | 470 | 53 | 13 | 672 | 1129 | 13834911 | 13834450 | 1.32e-120 | 446 |
| Clupea_harengus-NC_045161.1 | 80.851 | 470 | 48 | 20 | 672 | 1129 | 26923869 | 26923430 | 3.85e-86 | 331 |
| Clupea_harengus-NC_045161.1 | 80.851 | 470 | 48 | 20 | 672 | 1129 | 26927121 | 26926682 | 3.85e-86 | 331 |
| Clupea_harengus-NC_045155.1 | 83.925 | 479 | 61 | 10 | 662 | 1129 | 22511603 | 22512076 | 4.74e-120 | 444 |
| Clupea_harengus-NC_045155.1 | 86.967 | 399 | 42 | 10 | 734 | 1129 | 4921479 | 4921088 | 6.14e-119 | 440 |
| Clupea_harengus-NC_045155.1 | 83.830 | 470 | 56 | 13 | 672 | 1129 | 12005907 | 12006368 | 1.33e-115 | 429 |
| Clupea_harengus-NC_045155.1 | 84.146 | 410 | 47 | 16 | 732 | 1129 | 21769847 | 21770250 | 3.77e-101 | 381 |
| Clupea_harengus-NC_045155.1 | 79.849 | 531 | 63 | 23 | 612 | 1129 | 12037108 | 12037607 | 3.83e-91 | 348 |
| Clupea_harengus-NC_045155.1 | 90.217 | 184 | 17 | 1 | 731 | 913 | 25007344 | 25007527 | 2.40e-58 | 239 |
| Clupea_harengus-NC_045155.1 | 83.333 | 186 | 21 | 8 | 944 | 1129 | 27204536 | 27204361 | 1.49e-35 | 163 |
| Clupea_harengus-NW_024880258.1 | 83.891 | 478 | 61 | 10 | 662 | 1129 | 21866 | 21395 | 1.71e-119 | 442 |
| Clupea_harengus-NC_045177.1 | 82.397 | 534 | 68 | 17 | 612 | 1129 | 9097696 | 9098219 | 1.71e-119 | 442 |
| Clupea_harengus-NC_045177.1 | 89.441 | 161 | 15 | 1 | 959 | 1119 | 604928 | 605086 | 3.15e-47 | 202 |
| Clupea_harengus-NC_045159.1 | 82.364 | 533 | 71 | 17 | 610 | 1129 | 17420567 | 17420045 | 1.71e-119 | 442 |
| Clupea_harengus-NC_045159.1 | 83.099 | 497 | 60 | 9 | 641 | 1129 | 30168924 | 30169404 | 3.69e-116 | 431 |
| Clupea_harengus-NC_045159.1 | 83.099 | 497 | 60 | 9 | 641 | 1129 | 30172157 | 30172637 | 3.69e-116 | 431 |
| Clupea_harengus-NC_045159.1 | 83.299 | 479 | 60 | 14 | 662 | 1129 | 5668294 | 5667825 | 6.18e-114 | 424 |
| Clupea_harengus-NC_045164.1 | 84.255 | 470 | 54 | 13 | 672 | 1129 | 13910912 | 13911373 | 6.14e-119 | 440 |
| Clupea_harengus-NC_045164.1 | 83.830 | 470 | 56 | 13 | 672 | 1129 | 4166568 | 4166107 | 1.33e-115 | 429 |
| Clupea_harengus-NC_045164.1 | 90.323 | 186 | 16 | 1 | 944 | 1129 | 29780581 | 29780764 | 1.86e-59 | 243 |
| Clupea_harengus-NC_045153.1 | 83.857 | 477 | 62 | 9 | 662 | 1129 | 1222201 | 1221731 | 6.14e-119 | 440 |
| Clupea_harengus-NC_045153.1 | 81.714 | 525 | 62 | 23 | 612 | 1123 | 23831766 | 23832269 | 6.23e-109 | 407 |
| Clupea_harengus-NC_045153.1 | 80.566 | 530 | 61 | 21 | 612 | 1129 | 10564739 | 10564240 | 8.17e-98 | 370 |
| Clupea_harengus-NC_045153.1 | 80.414 | 531 | 61 | 22 | 612 | 1129 | 10561532 | 10561032 | 3.80e-96 | 364 |
| Clupea_harengus-NC_045153.1 | 80.962 | 499 | 52 | 16 | 641 | 1129 | 4773175 | 4772710 | 2.29e-93 | 355 |
| Clupea_harengus-NC_045153.1 | 83.077 | 260 | 35 | 8 | 869 | 1127 | 11914314 | 11914565 | 5.20e-55 | 228 |
| Clupea_harengus-NC_045174.1 | 83.099 | 497 | 60 | 9 | 641 | 1129 | 9101939 | 9102419 | 3.69e-116 | 431 |
| Clupea_harengus-NC_045174.1 | 83.099 | 497 | 60 | 9 | 641 | 1129 | 9105190 | 9105670 | 3.69e-116 | 431 |
| Clupea_harengus-NC_045174.1 | 88.660 | 97 | 11 | 0 | 204 | 300 | 16251387 | 16251291 | 3.27e-22 | 119 |
| Clupea_harengus-NC_045167.1 | 83.099 | 497 | 60 | 9 | 641 | 1129 | 9601845 | 9602325 | 3.69e-116 | 431 |
| Clupea_harengus-NC_045167.1 | 83.099 | 497 | 60 | 9 | 641 | 1129 | 9605260 | 9605740 | 3.69e-116 | 431 |
| Clupea_harengus-NC_045167.1 | 82.897 | 497 | 61 | 9 | 641 | 1129 | 23102596 | 23102116 | 1.72e-114 | 425 |
| Clupea_harengus-NC_045167.1 | 82.897 | 497 | 61 | 9 | 641 | 1129 | 23106011 | 23105531 | 1.72e-114 | 425 |
| Clupea_harengus-NC_045167.1 | 80.847 | 543 | 61 | 23 | 612 | 1129 | 1759809 | 1759285 | 8.11e-103 | 387 |
| Clupea_harengus-NC_045167.1 | 79.167 | 480 | 57 | 22 | 662 | 1129 | 17465823 | 17466271 | 1.82e-74 | 292 |
| Clupea_harengus-NC_045167.1 | 88.235 | 238 | 21 | 7 | 894 | 1129 | 7229754 | 7229522 | 5.09e-70 | 278 |
| Clupea_harengus-NC_045167.1 | 90.955 | 199 | 13 | 5 | 932 | 1129 | 7214290 | 7214484 | 1.43e-65 | 263 |
| Clupea_harengus-NW_024880153.1 | 83.034 | 501 | 56 | 13 | 641 | 1129 | 4769 | 5252 | 4.78e-115 | 427 |
| Clupea_harengus-NW_024880153.1 | 83.830 | 470 | 53 | 15 | 672 | 1129 | 1548 | 2006 | 1.72e-114 | 425 |
| Clupea_harengus-NC_045168.1 | 82.897 | 497 | 60 | 11 | 641 | 1129 | 15185205 | 15185684 | 6.18e-114 | 424 |
| Clupea_harengus-NC_045168.1 | 80.189 | 530 | 63 | 21 | 612 | 1129 | 23950196 | 23949697 | 1.77e-94 | 359 |
| Clupea_harengus-NC_045168.1 | 80.189 | 530 | 63 | 21 | 612 | 1129 | 23954352 | 23953853 | 1.77e-94 | 359 |
| Clupea_harengus-NW_024879911.1 | 83.125 | 480 | 61 | 14 | 662 | 1129 | 7244 | 6773 | 8.00e-113 | 420 |
| Clupea_harengus-NC_045157.1 | 83.054 | 478 | 65 | 10 | 662 | 1129 | 12381122 | 12381593 | 8.00e-113 | 420 |
| Clupea_harengus-NC_045157.1 | 81.046 | 459 | 48 | 18 | 682 | 1129 | 9020057 | 9020487 | 1.39e-85 | 329 |
| Clupea_harengus-NC_045175.1 | 85.222 | 406 | 46 | 10 | 734 | 1129 | 17353123 | 17352722 | 2.24e-108 | 405 |
| Clupea_harengus-NC_045175.1 | 78.026 | 537 | 79 | 22 | 608 | 1120 | 14857971 | 14858492 | 3.02e-77 | 302 |
| Clupea_harengus-NC_045175.1 | 85.789 | 190 | 21 | 5 | 940 | 1129 | 16037831 | 16037648 | 1.47e-45 | 196 |
| Clupea_harengus-NW_024879659.1 | 81.928 | 498 | 62 | 20 | 641 | 1129 | 101352 | 100874 | 1.35e-105 | 396 |
| Clupea_harengus-NC_045162.1 | 82.227 | 467 | 57 | 11 | 672 | 1129 | 5645894 | 5646343 | 1.36e-100 | 379 |
| Clupea_harengus-NC_045162.1 | 82.227 | 467 | 57 | 11 | 672 | 1129 | 5649256 | 5649705 | 1.36e-100 | 379 |
| Clupea_harengus-NC_045162.1 | 81.197 | 468 | 50 | 17 | 672 | 1129 | 16722191 | 16722630 | 1.78e-89 | 342 |
| Clupea_harengus-NC_045162.1 | 80.123 | 488 | 71 | 15 | 662 | 1129 | 3241536 | 3242017 | 6.40e-89 | 340 |
| Clupea_harengus-NC_045162.1 | 87.356 | 174 | 17 | 4 | 956 | 1129 | 16724609 | 16724777 | 5.28e-45 | 195 |
| Clupea_harengus-NC_045162.1 | 80.645 | 217 | 35 | 7 | 204 | 415 | 12055106 | 12055320 | 5.35e-35 | 161 |
| Clupea_harengus-NC_045162.1 | 94.667 | 75 | 4 | 0 | 1055 | 1129 | 29278133 | 29278207 | 1.17e-21 | 117 |
| Clupea_harengus-NC_045169.1 | 80.377 | 530 | 62 | 21 | 612 | 1129 | 20683031 | 20683530 | 3.80e-96 | 364 |
| Clupea_harengus-NC_045169.1 | 78.554 | 401 | 48 | 17 | 731 | 1129 | 26577882 | 26578246 | 1.45e-55 | 230 |
| Clupea_harengus-NC_045156.1 | 79.852 | 541 | 74 | 21 | 612 | 1129 | 23625995 | 23625467 | 1.37e-95 | 363 |
| Clupea_harengus-NC_045176.1 | 81.935 | 465 | 48 | 18 | 685 | 1129 | 462351 | 461903 | 4.92e-95 | 361 |
| Clupea_harengus-NC_045176.1 | 87.500 | 128 | 16 | 0 | 213 | 340 | 7999221 | 7999094 | 4.17e-31 | 148 |
| Clupea_harengus-NW_024879574.1 | 79.952 | 414 | 39 | 28 | 733 | 1129 | 11580 | 11194 | 3.96e-66 | 265 |
| Clupea_harengus-NW_024880278.1 | 81.735 | 219 | 32 | 7 | 756 | 970 | 23791 | 24005 | 1.91e-39 | 176 |

### N5.HOG0018901 Blastn of conserved regions against all genome assemblies

#### Region B

| sseqid | pident | length | mismatch | gapopen | qstart | qend | sstart | send | evalue | bitscore |
|---|---|---|---|---|---|---|---|---|---|---|
| Hypomesus_transpacificus-NC_061067.1 | 100.000 | 301 | 0 | 0 | 1 | 301 | 9523254 | 9522954 | 1.43e-154 | 556 |
| Clupea_harengus-NC_045166.1 | 91.518 | 224 | 15 | 1 | 78 | 301 | 15872317 | 15872536 | 5.71e-79 | 305 |
| Clupea_harengus-NC_045166.1 | 92.391 | 92 | 7 | 0 | 6 | 97 | 15872209 | 15872300 | 1.02e-26 | 132 |

N5.HOG0018901

The gene is found in an alernative assembly of *H. transpacificus*

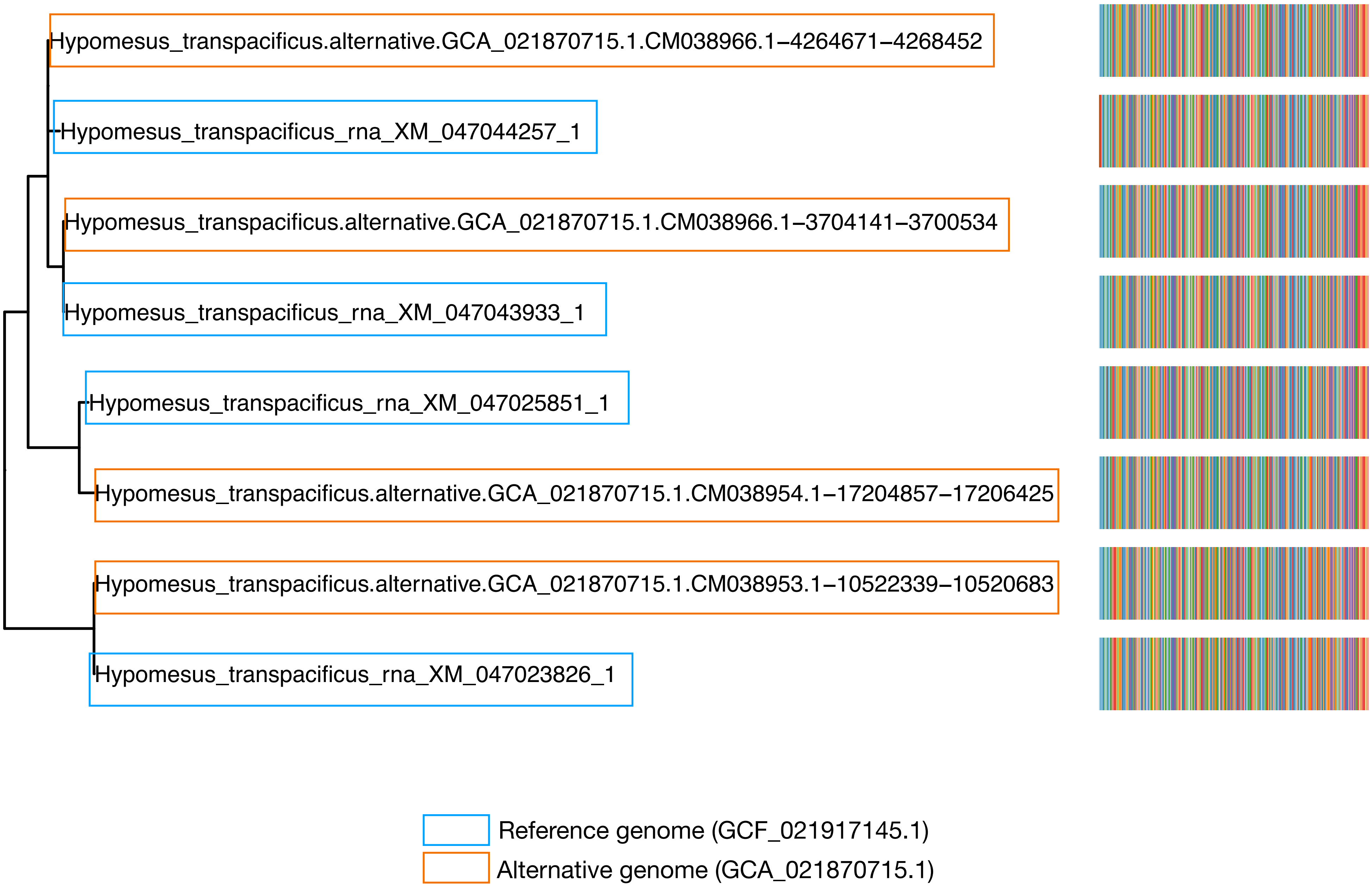

N5.HOG0018901

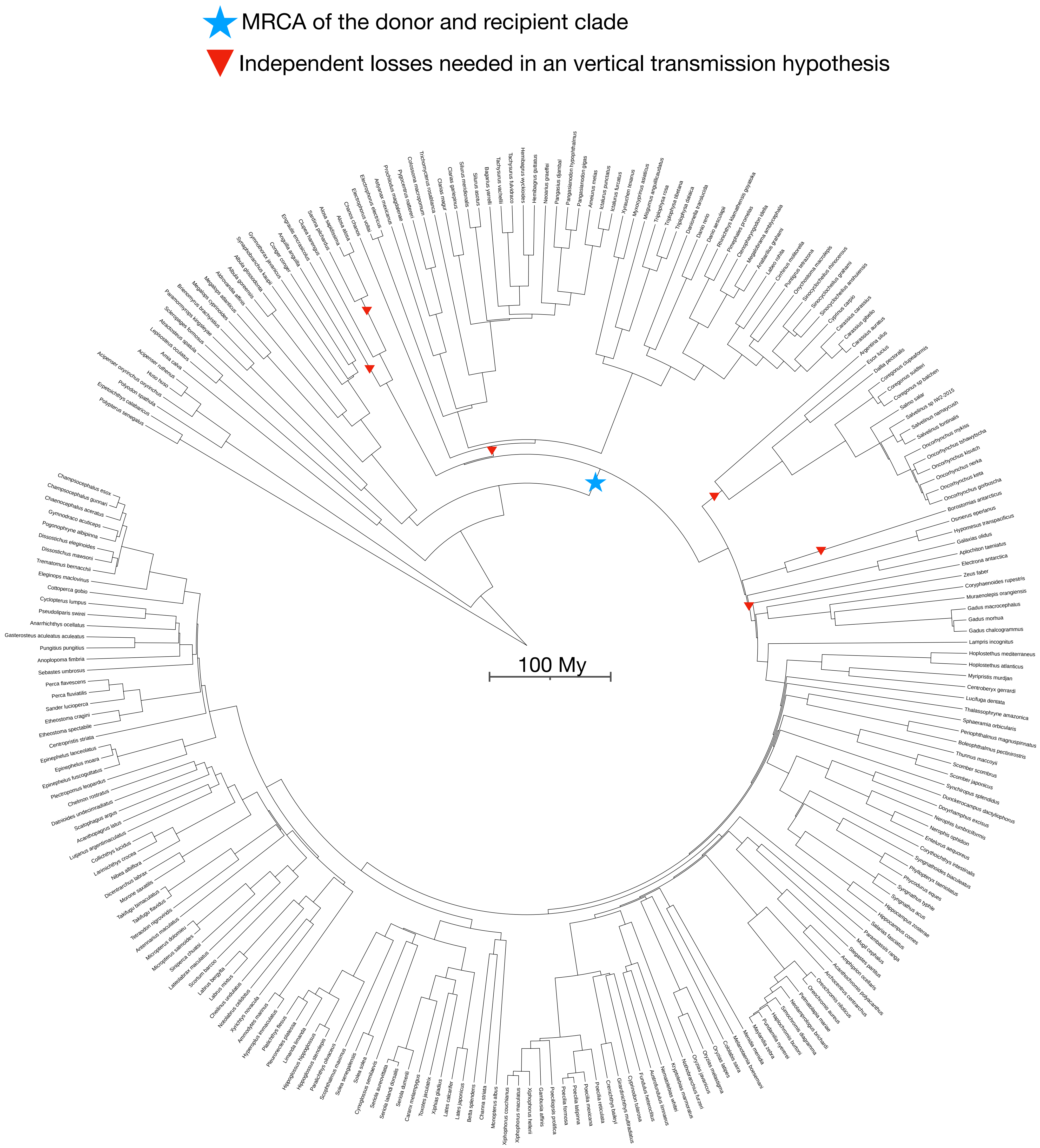

N5.HOG0008885

### N5.HOG0008885

Species tree

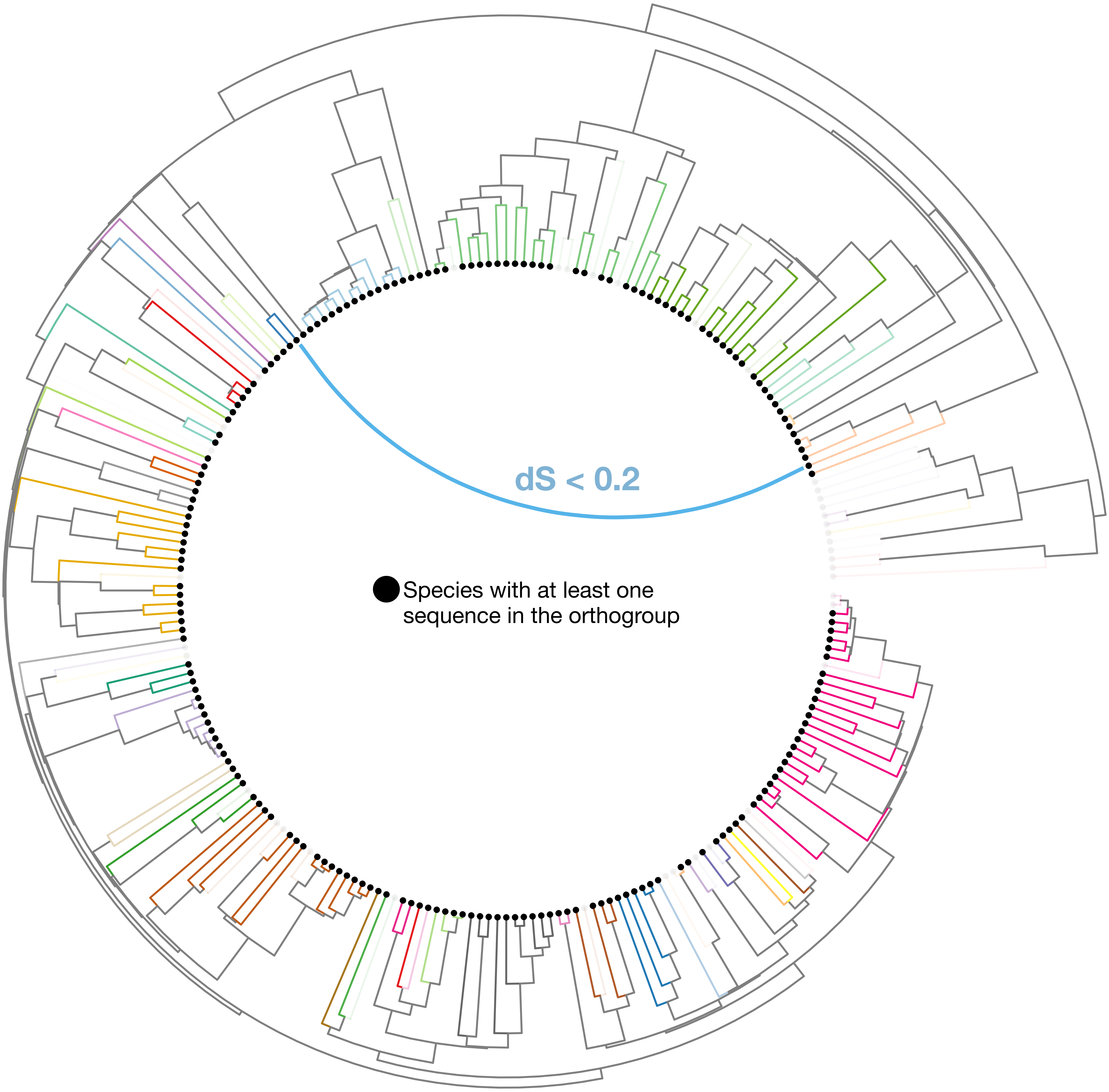

### N5.HOG0008885

Orthogroup phylogeny
and
protein alignment

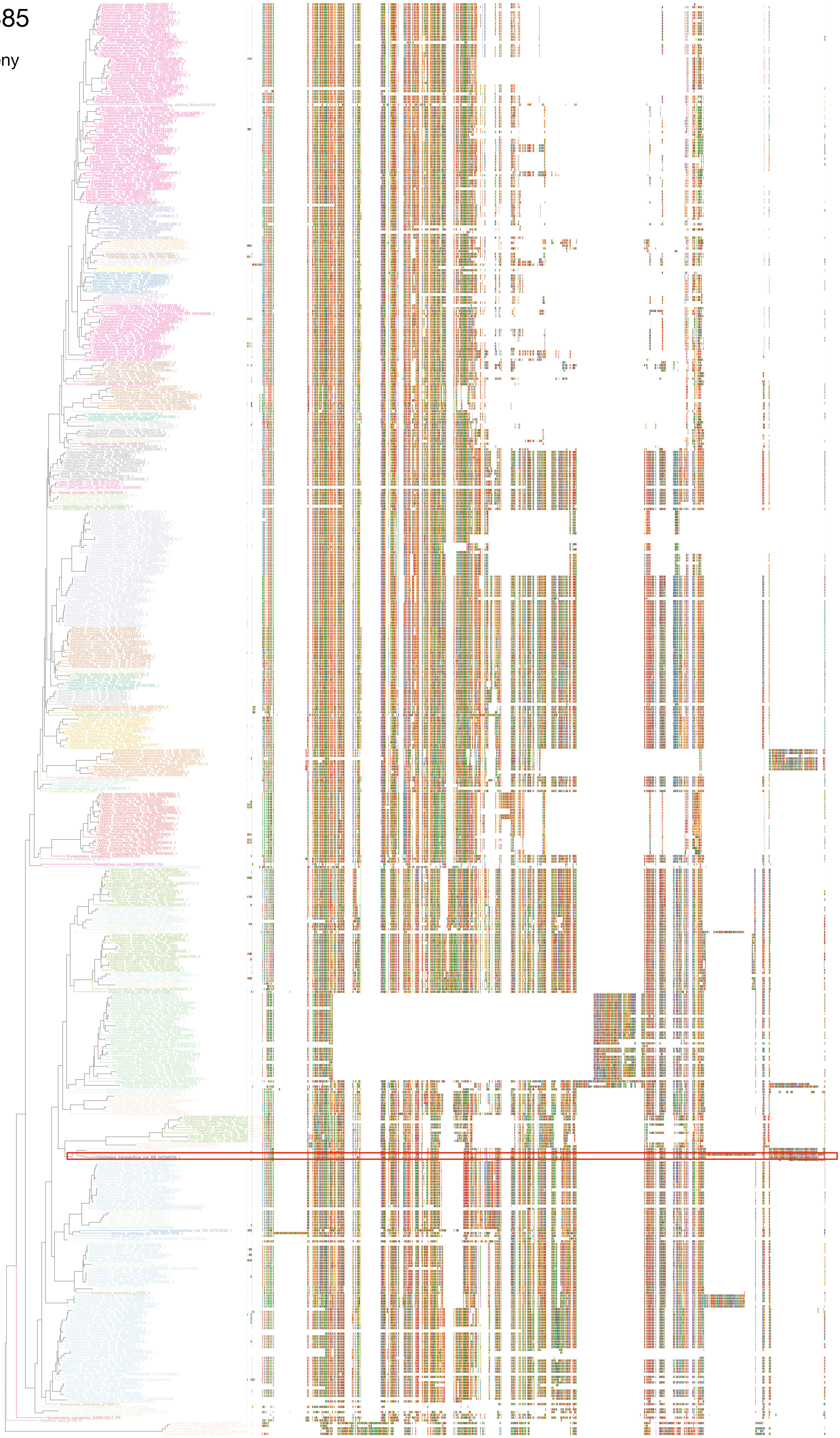

### N5.HOG0008885

Zoom on HGT clade with bootstrap values

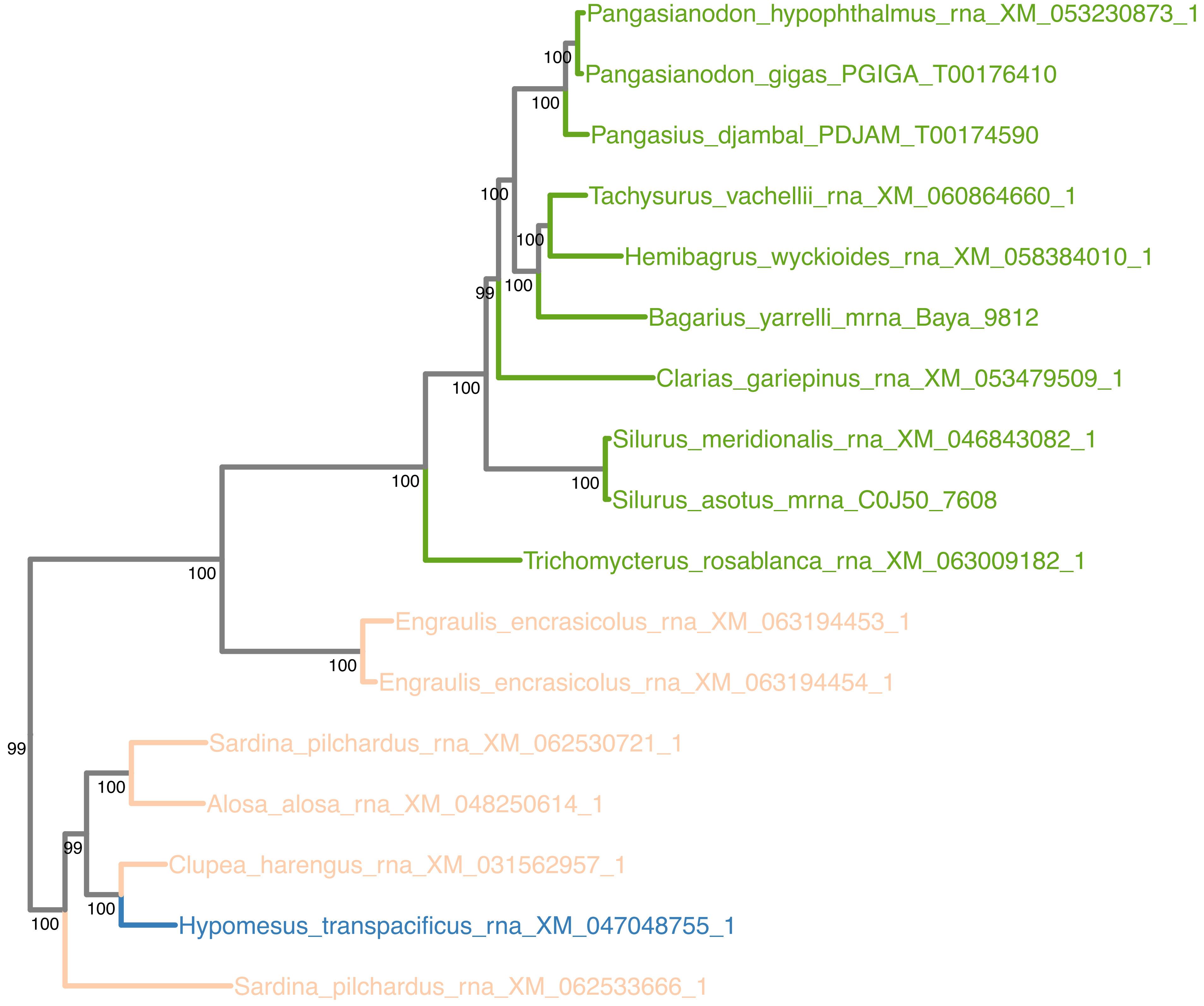

### N5.HOG0008885

Orthogroup phylogeny
and
trimmed
protein alignment

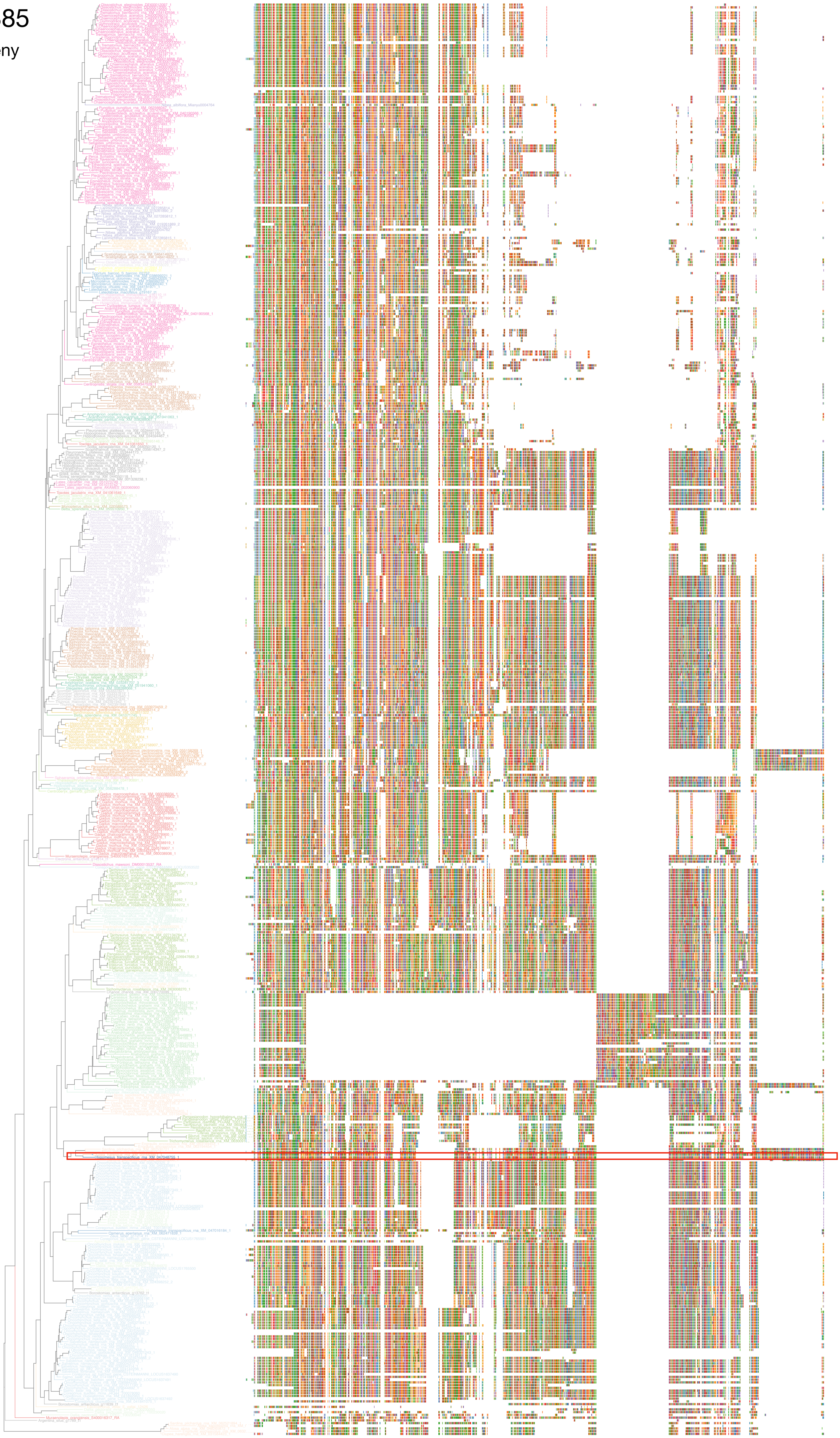

### N5.HOG0008885

« Best-match » phylogeny
and
protein alignment

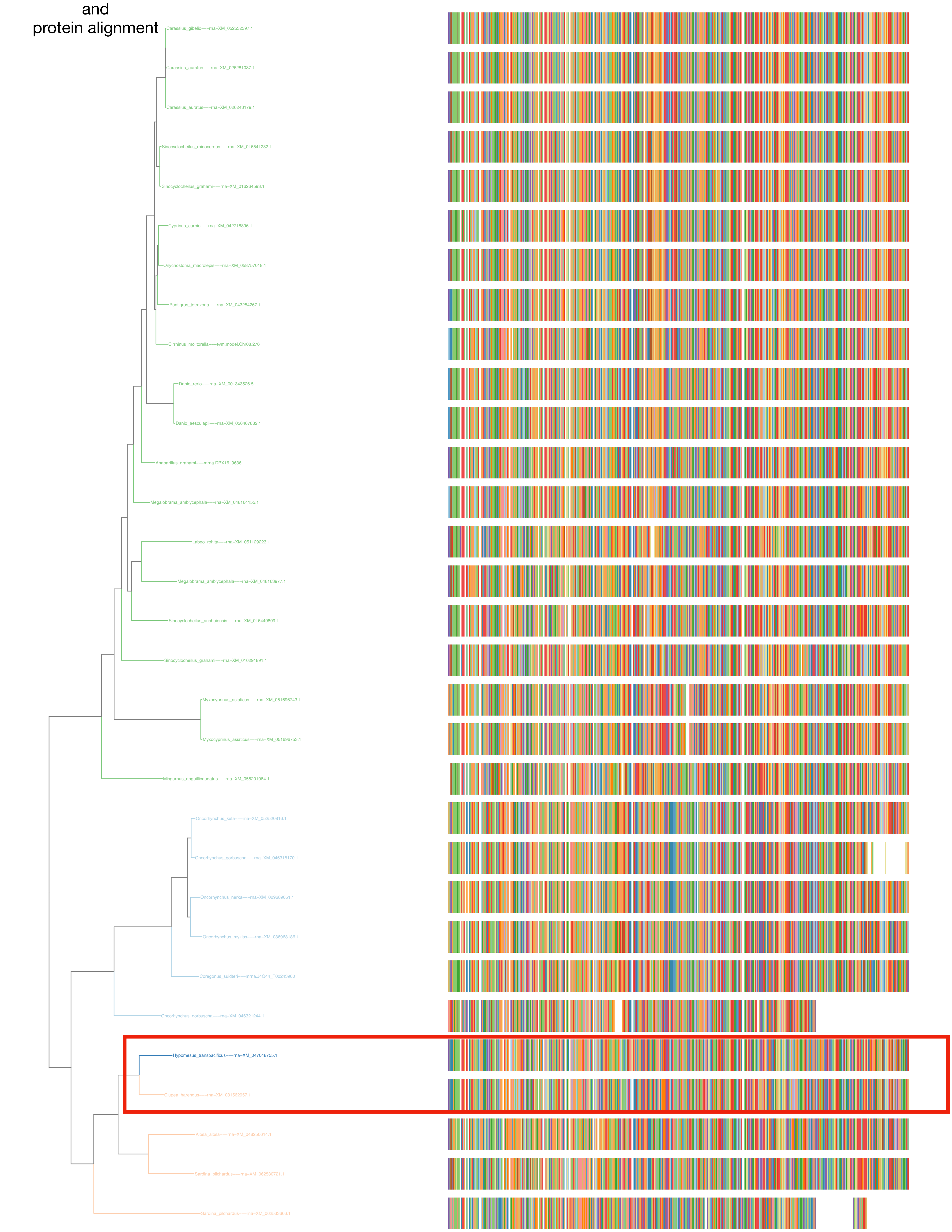

N5.HOG0008885
dS distribution

Inter-species dS distribution

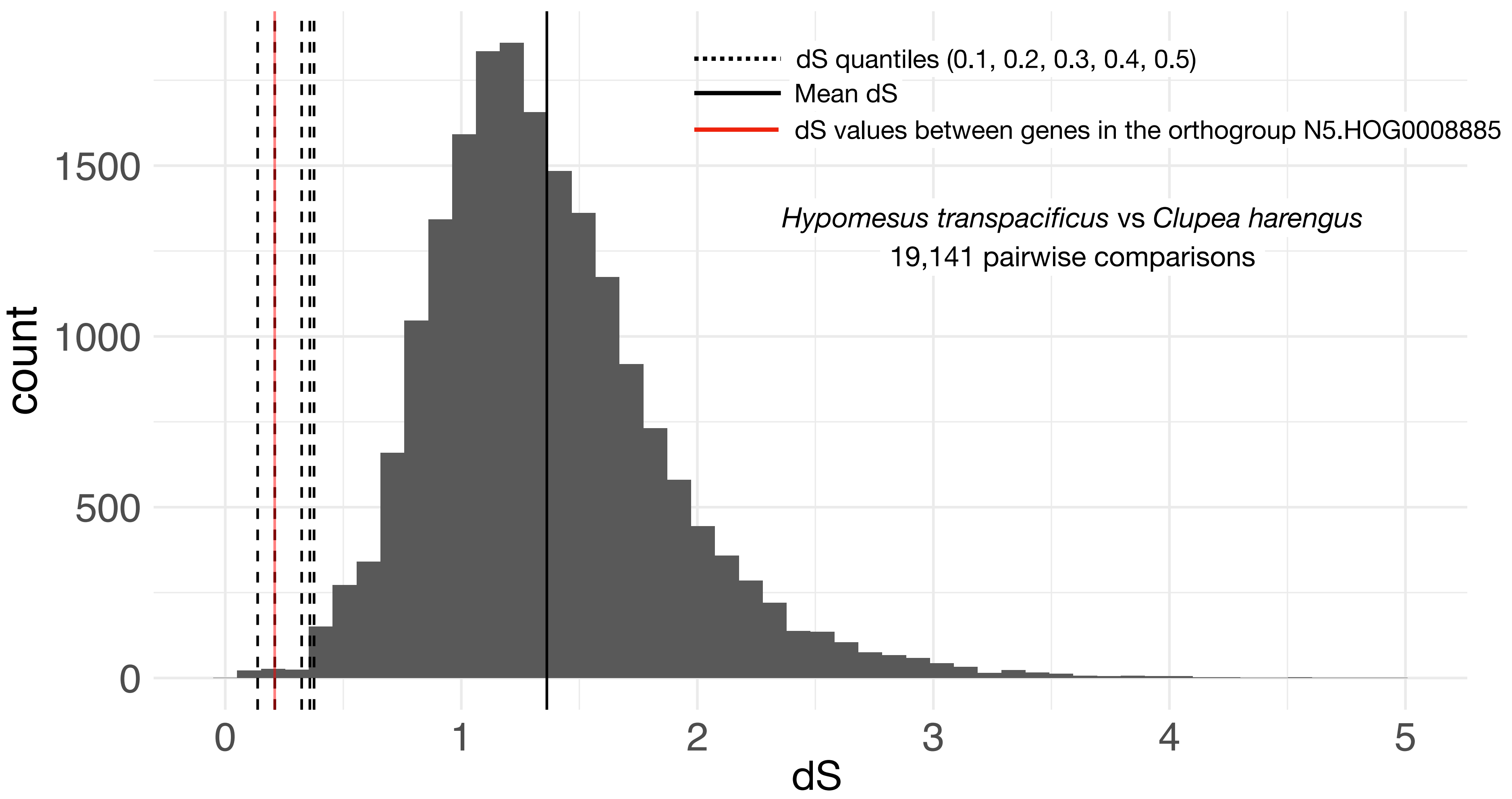


Intra-orthogroup dS

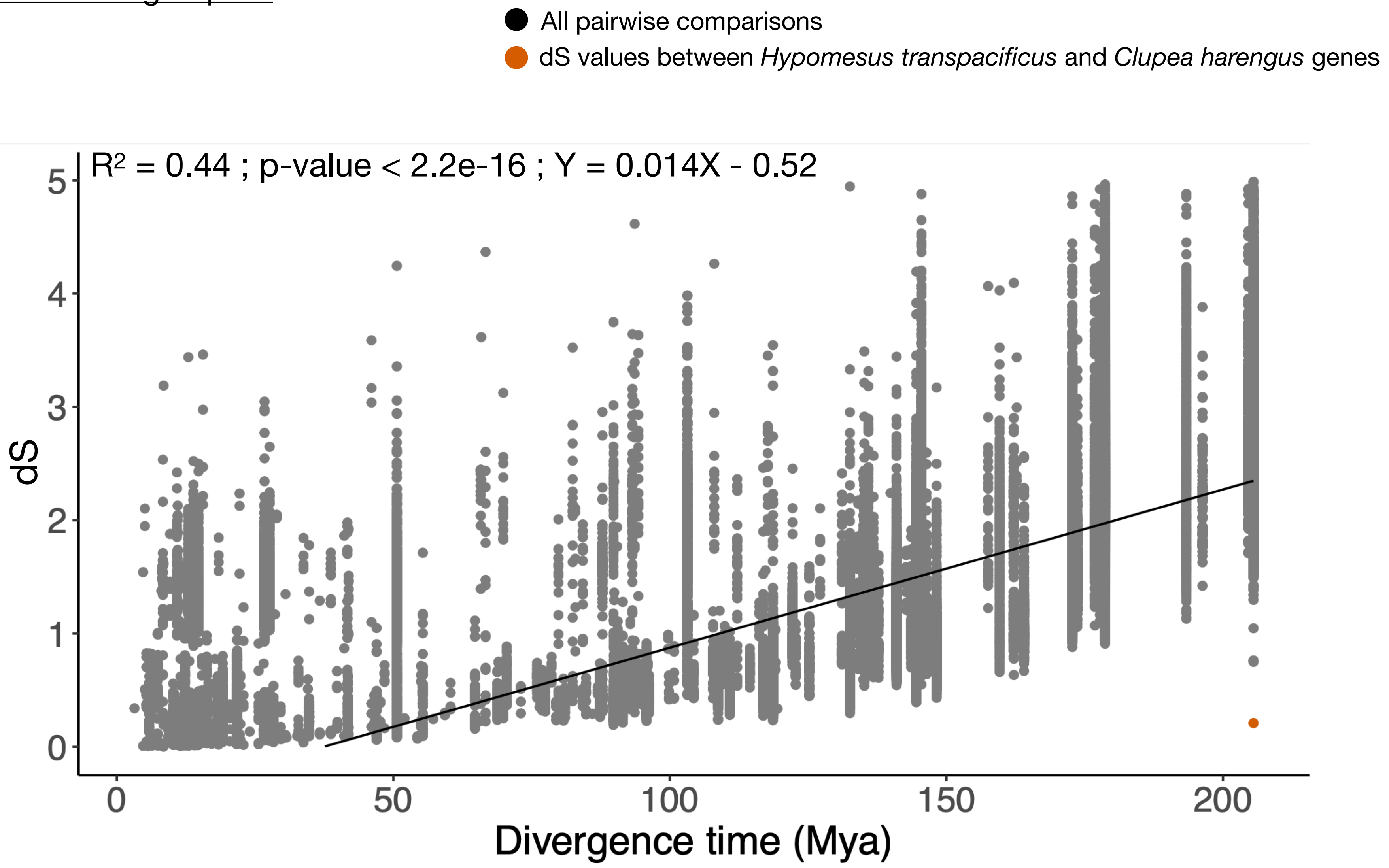

### N5.HOG0008885

#### Micro-synteny

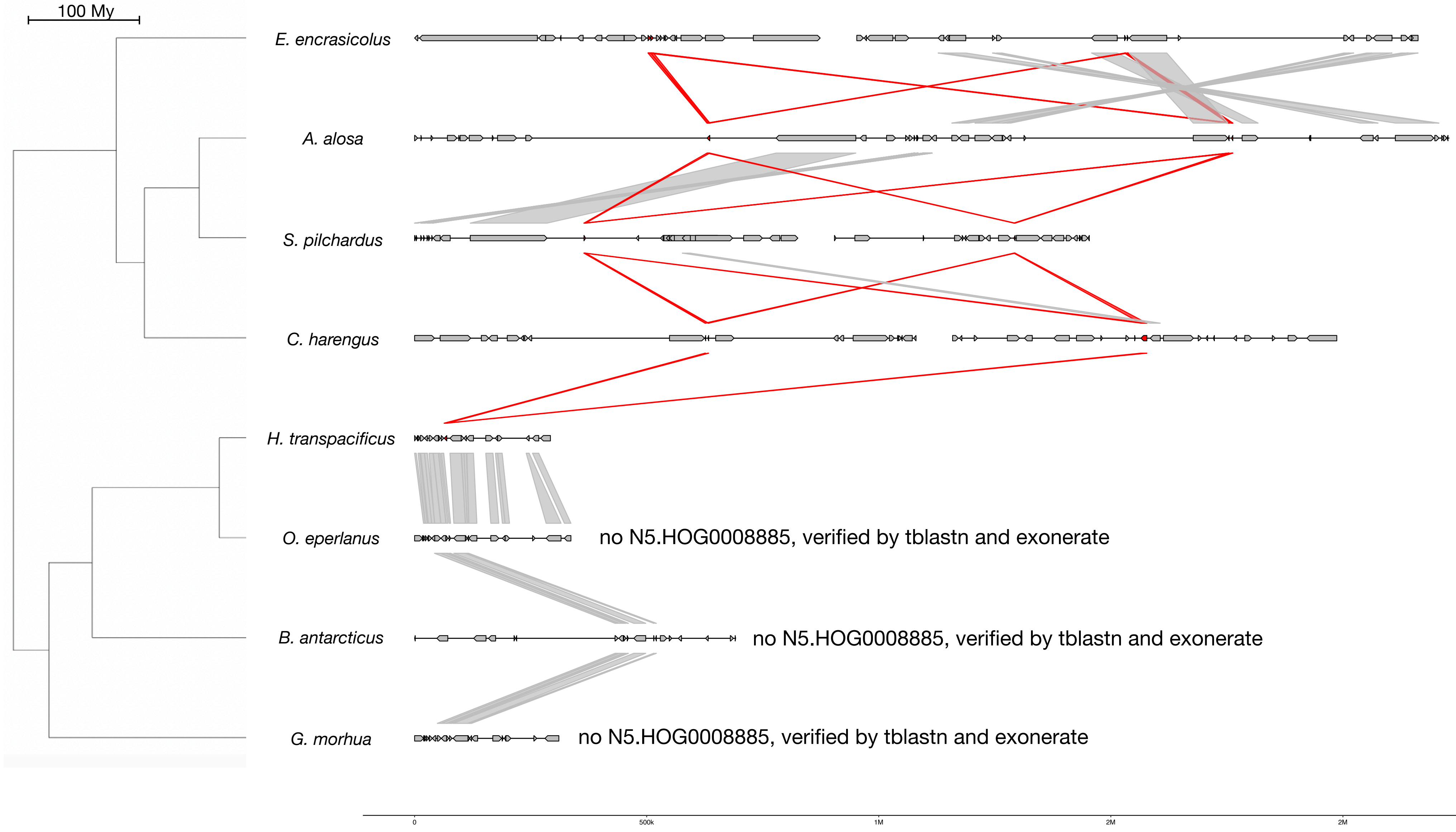


#### Zoom on *C. harengus* and *H. transpacificus*

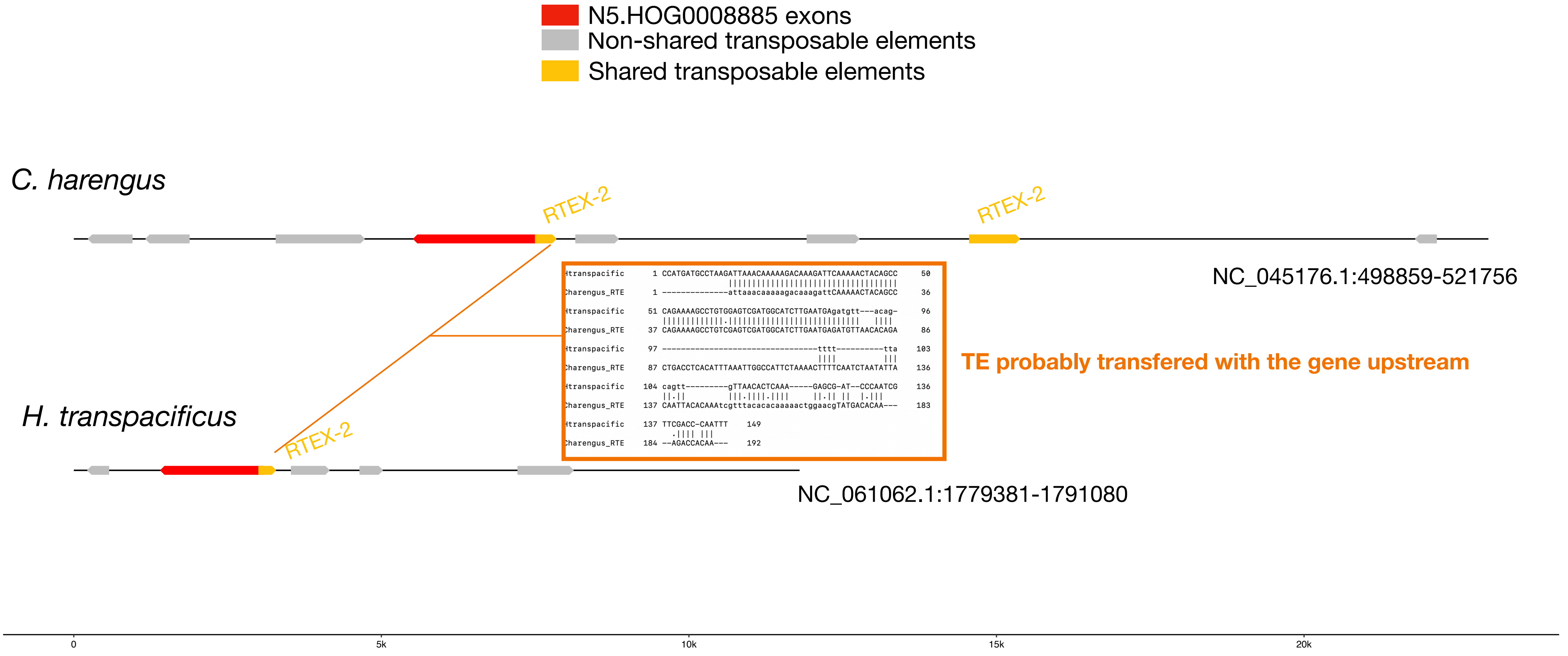


#### RTEX-2 :

| | |
|---|---|
| *E. encrasicolus* | *33 copies scattered in the genome* |
| *A. alosa* | *338 copies scattered in the genome* |
| *S. pilchardus* | *204 copies scattered in the genome* |
| *C. harengus* | *338 copies scattered in the genome* |
| *H. transpacificus* | *16 copies scattered in the genome* |
| *O. eperlanus* | *64 copies scattered in the genome* |
| *B. antarcticus* | *834 copies scattered in the genome* |
| *G. morhua* | *97 copies scattered in the genome* |

N5.HOG0008885

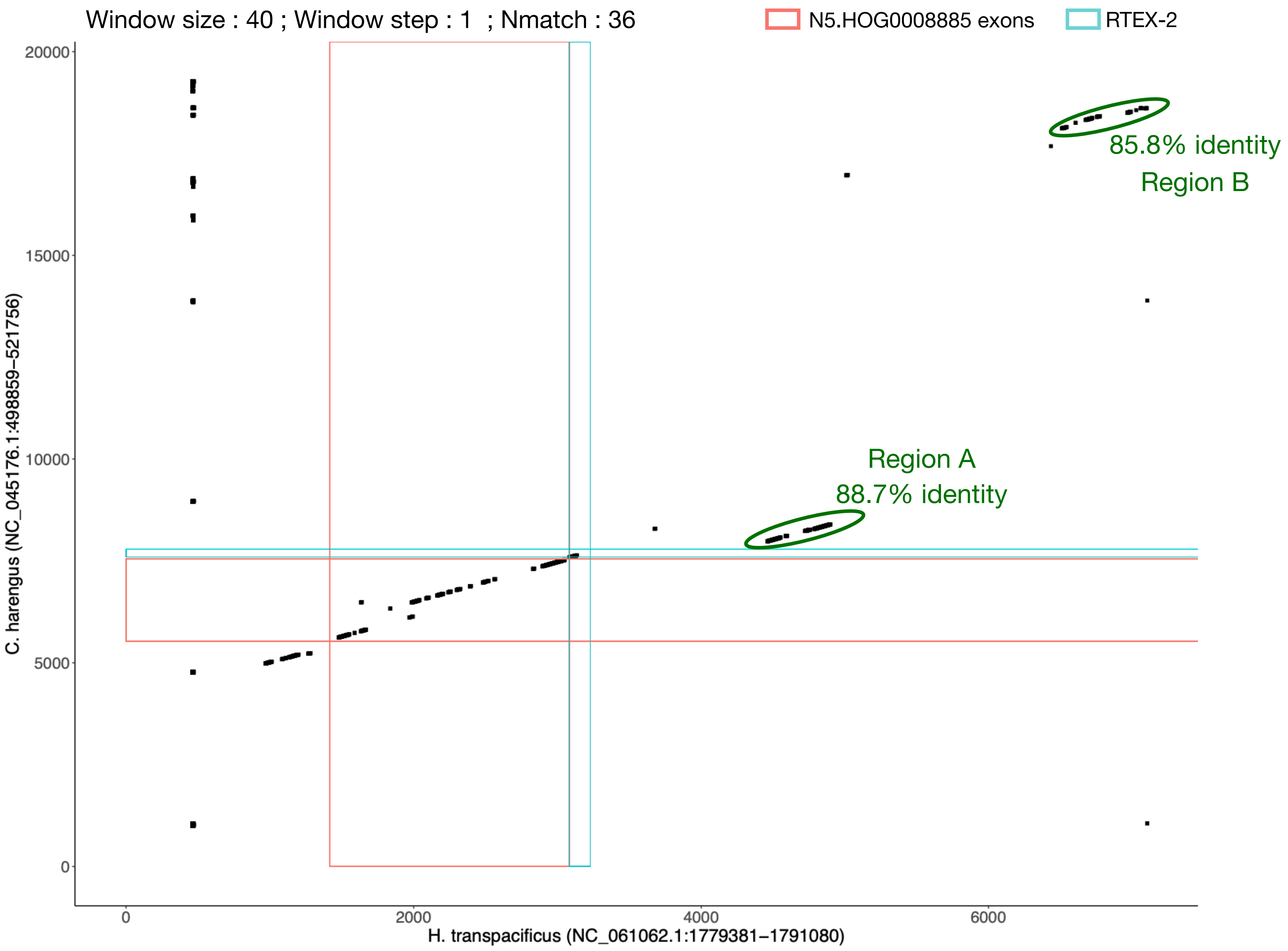

### N5.HOG0008885 Blastn of conserved regions against all genome assemblies

#### Region A (1/10)

| sseqid | pident | length | mismatch | gapopen | qstart | qend | sstart | send | evalue | bitscore |
|---|---|---|---|---|---|---|---|---|---|---|
| Hypomesus_transpacificus-NC_061062.1 | 100.000 | 601 | 0 | 0 | 1 | 601 | 1783780 | 1784380 | 0.0 | 1110 |
| Clupea_harengus-NC_045163.1 | 92.445 | 503 | 29 | 5 | 41 | 536 | 23572368 | 23571868 | 0.0 | 710 |
| Clupea_harengus-NC_045163.1 | 89.200 | 500 | 48 | 4 | 43 | 536 | 8556386 | 8556885 | 3.80e-173 | 619 |
| Clupea_harengus-NC_045163.1 | 89.022 | 501 | 43 | 6 | 48 | 536 | 10392746 | 10392246 | 2.29e-170 | 610 |
| Clupea_harengus-NC_045163.1 | 89.069 | 494 | 45 | 5 | 52 | 536 | 6990178 | 6989685 | 1.07e-168 | 604 |
| Clupea_harengus-NC_045163.1 | 88.732 | 497 | 44 | 6 | 52 | 536 | 6516134 | 6515638 | 1.78e-166 | 597 |
| Clupea_harengus-NC_045163.1 | 88.732 | 497 | 44 | 7 | 52 | 536 | 9068555 | 9068059 | 1.78e-166 | 597 |
| Clupea_harengus-NC_045163.1 | 88.086 | 512 | 43 | 8 | 43 | 536 | 7786544 | 7787055 | 8.29e-165 | 592 |
| Clupea_harengus-NC_045163.1 | 87.843 | 510 | 46 | 8 | 43 | 536 | 22151718 | 22152227 | 1.39e-162 | 584 |
| Clupea_harengus-NC_045163.1 | 87.695 | 512 | 45 | 11 | 43 | 536 | 10328661 | 10329172 | 1.79e-161 | 580 |
| Clupea_harengus-NC_045163.1 | 87.379 | 515 | 44 | 12 | 43 | 536 | 3743129 | 3743643 | 1.08e-158 | 571 |
| Clupea_harengus-NC_045163.1 | 87.426 | 509 | 49 | 7 | 43 | 536 | 12843474 | 12843982 | 1.08e-158 | 571 |
| Clupea_harengus-NC_045163.1 | 87.078 | 503 | 39 | 11 | 52 | 536 | 171444 | 170950 | 6.55e-151 | 545 |
| Clupea_harengus-NC_045163.1 | 86.248 | 509 | 55 | 8 | 43 | 536 | 6970709 | 6971217 | 1.10e-148 | 538 |
| Clupea_harengus-NC_045163.1 | 86.680 | 503 | 46 | 7 | 43 | 536 | 8312401 | 8312891 | 1.10e-148 | 538 |
| Clupea_harengus-NC_045163.1 | 86.707 | 331 | 38 | 3 | 43 | 367 | 10329466 | 10329796 | 7.08e-96 | 363 |
| Clupea_harengus-NC_045163.1 | 85.836 | 353 | 28 | 14 | 43 | 386 | 7126364 | 7126703 | 1.18e-93 | 355 |
| Clupea_harengus-NC_045163.1 | 85.538 | 325 | 38 | 5 | 43 | 358 | 8555877 | 8556201 | 2.00e-86 | 331 |
| Clupea_harengus-NC_045163.1 | 85.849 | 318 | 31 | 9 | 129 | 432 | 6311337 | 6311020 | 9.29e-85 | 326 |
| Clupea_harengus-NC_045163.1 | 95.556 | 180 | 8 | 0 | 357 | 536 | 8557929 | 8558108 | 1.22e-73 | 289 |
| Clupea_harengus-NC_045163.1 | 93.985 | 133 | 8 | 0 | 404 | 536 | 10329794 | 10329926 | 1.63e-47 | 202 |
| Synaphobranchus_kaupii-CM056379.1 | 91.865 | 504 | 31 | 6 | 43 | 536 | 118502789 | 118502286 | 0.0 | 695 |
| Synaphobranchus_kaupii-CM056379.1 | 87.810 | 525 | 27 | 15 | 43 | 536 | 8962172 | 8961654 | 1.79e-161 | 580 |
| Synaphobranchus_kaupii-CM056379.1 | 90.583 | 446 | 29 | 8 | 43 | 477 | 36536544 | 36536101 | 6.46e-161 | 579 |
| Synaphobranchus_kaupii-CM056379.1 | 90.993 | 433 | 31 | 5 | 112 | 536 | 111868698 | 111868266 | 2.32e-160 | 577 |
| Synaphobranchus_kaupii-CM056379.1 | 84.120 | 466 | 57 | 7 | 52 | 515 | 11434819 | 11434369 | 1.48e-117 | 435 |
| Synaphobranchus_kaupii-CM056379.1 | 87.989 | 358 | 33 | 7 | 43 | 390 | 82479657 | 82479300 | 1.93e-111 | 414 |
| Synaphobranchus_kaupii-CM056379.1 | 89.524 | 315 | 24 | 5 | 43 | 350 | 43908718 | 43908406 | 3.25e-104 | 390 |
| Synaphobranchus_kaupii-CM056379.1 | 89.423 | 312 | 28 | 3 | 43 | 350 | 6934121 | 6933811 | 1.17e-103 | 388 |
| Synaphobranchus_kaupii-CM056379.1 | 89.423 | 312 | 28 | 3 | 43 | 350 | 42696786 | 42696476 | 1.17e-103 | 388 |
| Synaphobranchus_kaupii-CM056379.1 | 89.423 | 312 | 28 | 3 | 43 | 350 | 54868900 | 54869210 | 1.17e-103 | 388 |
| Synaphobranchus_kaupii-CM056379.1 | 89.423 | 312 | 28 | 3 | 43 | 350 | 106106219 | 106106529 | 1.17e-103 | 388 |
| Synaphobranchus_kaupii-CM056379.1 | 88.959 | 317 | 29 | 4 | 38 | 350 | 8271009 | 8270695 | 4.20e-103 | 387 |
| Synaphobranchus_kaupii-CM056379.1 | 89.836 | 305 | 26 | 3 | 38 | 338 | 8489504 | 8489807 | 4.20e-103 | 387 |
| Synaphobranchus_kaupii-CM056379.1 | 90.033 | 301 | 26 | 2 | 43 | 339 | 46254599 | 46254299 | 4.20e-103 | 387 |
| Synaphobranchus_kaupii-CM056379.1 | 90.000 | 300 | 26 | 2 | 43 | 338 | 46398408 | 46398707 | 1.51e-102 | 385 |
| Synaphobranchus_kaupii-CM056379.1 | 89.803 | 304 | 25 | 3 | 43 | 340 | 79998037 | 79998340 | 1.51e-102 | 385 |
| Synaphobranchus_kaupii-CM056379.1 | 88.889 | 315 | 29 | 4 | 43 | 353 | 17443248 | 17442936 | 5.43e-102 | 383 |
| Synaphobranchus_kaupii-CM056379.1 | 89.103 | 312 | 29 | 3 | 43 | 350 | 82476142 | 82476452 | 5.43e-102 | 383 |
| Synaphobranchus_kaupii-CM056379.1 | 89.103 | 312 | 29 | 3 | 43 | 350 | 95129870 | 95129560 | 5.43e-102 | 383 |
| Synaphobranchus_kaupii-CM056379.1 | 90.000 | 300 | 24 | 3 | 43 | 340 | 96161143 | 96160848 | 5.43e-102 | 383 |
| Synaphobranchus_kaupii-CM056379.1 | 89.701 | 301 | 27 | 2 | 43 | 339 | 7904718 | 7904418 | 1.95e-101 | 381 |
| Synaphobranchus_kaupii-CM056379.1 | 88.644 | 317 | 30 | 4 | 38 | 350 | 82474761 | 82474447 | 1.95e-101 | 381 |
| Synaphobranchus_kaupii-CM056379.1 | 88.571 | 315 | 30 | 4 | 43 | 353 | 64327941 | 64328253 | 2.53e-100 | 377 |
| Synaphobranchus_kaupii-CM056379.1 | 88.462 | 312 | 31 | 3 | 43 | 350 | 8959577 | 8959887 | 1.18e-98 | 372 |
| Synaphobranchus_kaupii-CM056379.1 | 87.821 | 312 | 27 | 7 | 43 | 350 | 8812717 | 8813021 | 1.18e-93 | 355 |
| Synaphobranchus_kaupii-CM056379.1 | 87.821 | 312 | 27 | 7 | 43 | 350 | 8815492 | 8815796 | 1.18e-93 | 355 |
| Synaphobranchus_kaupii-CM056379.1 | 87.821 | 312 | 27 | 7 | 43 | 350 | 8818267 | 8818571 | 1.18e-93 | 355 |
| Synaphobranchus_kaupii-CM056379.1 | 87.821 | 312 | 27 | 7 | 43 | 350 | 8821042 | 8821346 | 1.18e-93 | 355 |
| Synaphobranchus_kaupii-CM056379.1 | 85.897 | 312 | 29 | 5 | 43 | 350 | 2872467 | 2872767 | 1.55e-82 | 318 |
| Synaphobranchus_kaupii-CM056379.1 | 83.974 | 312 | 25 | 4 | 43 | 350 | 8834287 | 8833997 | 9.49e-70 | 276 |
| Synaphobranchus_kaupii-CM056379.1 | 92.973 | 185 | 11 | 1 | 43 | 225 | 10437936 | 10438120 | 1.59e-67 | 268 |
| Synaphobranchus_kaupii-CM056379.1 | 86.772 | 189 | 21 | 2 | 48 | 234 | 76414847 | 76414661 | 3.51e-49 | 207 |
| Synaphobranchus_kaupii-CM056379.1 | 90.400 | 125 | 7 | 2 | 43 | 165 | 85616995 | 85616874 | 9.97e-35 | 159 |
| Synaphobranchus_kaupii-CM056378.1 | 91.552 | 509 | 32 | 7 | 38 | 536 | 43970180 | 43969673 | 0.0 | 691 |
| Synaphobranchus_kaupii-CM056378.1 | 88.762 | 525 | 28 | 12 | 43 | 536 | 95106340 | 95106864 | 1.77e-171 | 614 |
| Synaphobranchus_kaupii-CM056378.1 | 89.744 | 312 | 27 | 3 | 43 | 350 | 43721363 | 43721673 | 2.51e-105 | 394 |
| Synaphobranchus_kaupii-CM056378.1 | 89.274 | 317 | 28 | 5 | 38 | 350 | 70371837 | 70371523 | 9.03e-105 | 392 |
| Synaphobranchus_kaupii-CM056378.1 | 90.572 | 297 | 24 | 3 | 43 | 335 | 107270871 | 107270575 | 3.25e-104 | 390 |
| Synaphobranchus_kaupii-CM056378.1 | 89.206 | 315 | 30 | 3 | 38 | 350 | 117683895 | 117684207 | 3.25e-104 | 390 |
| Synaphobranchus_kaupii-CM056378.1 | 89.423 | 312 | 28 | 3 | 43 | 350 | 2922414 | 2922104 | 1.17e-103 | 388 |
| Synaphobranchus_kaupii-CM056378.1 | 89.423 | 312 | 28 | 3 | 43 | 350 | 6421908 | 6421598 | 1.17e-103 | 388 |
| Synaphobranchus_kaupii-CM056378.1 | 89.423 | 312 | 28 | 3 | 43 | 350 | 12044708 | 12044398 | 1.17e-103 | 388 |
| Synaphobranchus_kaupii-CM056378.1 | 89.423 | 312 | 28 | 3 | 43 | 350 | 26972227 | 26972537 | 1.17e-103 | 388 |
| Synaphobranchus_kaupii-CM056378.1 | 89.423 | 312 | 28 | 3 | 43 | 350 | 45924479 | 45924169 | 1.17e-103 | 388 |
| Synaphobranchus_kaupii-CM056378.1 | 89.423 | 312 | 28 | 3 | 43 | 350 | 89247987 | 89248297 | 1.17e-103 | 388 |
| Synaphobranchus_kaupii-CM056378.1 | 89.423 | 312 | 28 | 3 | 43 | 350 | 124134756 | 124135066 | 1.17e-103 | 388 |
| Synaphobranchus_kaupii-CM056378.1 | 90.033 | 301 | 26 | 2 | 43 | 339 | 2915316 | 2915616 | 4.20e-103 | 387 |
| Synaphobranchus_kaupii-CM056378.1 | 89.206 | 315 | 26 | 4 | 43 | 350 | 62823166 | 62823479 | 4.20e-103 | 387 |
| Synaphobranchus_kaupii-CM056378.1 | 89.206 | 315 | 26 | 4 | 43 | 350 | 69339287 | 69339600 | 4.20e-103 | 387 |
| Synaphobranchus_kaupii-CM056378.1 | 89.206 | 315 | 26 | 4 | 43 | 350 | 115941306 | 115941619 | 4.20e-103 | 387 |
| Synaphobranchus_kaupii-CM056378.1 | 89.103 | 312 | 29 | 3 | 43 | 350 | 4047737 | 4047427 | 5.43e-102 | 383 |
| Synaphobranchus_kaupii-CM056378.1 | 89.103 | 312 | 29 | 3 | 43 | 350 | 6428820 | 6428510 | 5.43e-102 | 383 |
| Synaphobranchus_kaupii-CM056378.1 | 89.103 | 312 | 29 | 3 | 43 | 350 | 26971544 | 26971854 | 5.43e-102 | 383 |
| Synaphobranchus_kaupii-CM056378.1 | 89.103 | 312 | 29 | 3 | 43 | 350 | 43036744 | 43036434 | 5.43e-102 | 383 |
| Synaphobranchus_kaupii-CM056378.1 | 89.103 | 312 | 29 | 3 | 43 | 350 | 43713414 | 43713104 | 5.43e-102 | 383 |
| Synaphobranchus_kaupii-CM056378.1 | 89.542 | 306 | 27 | 3 | 38 | 339 | 69962008 | 69962312 | 5.43e-102 | 383 |
| Synaphobranchus_kaupii-CM056378.1 | 89.103 | 312 | 29 | 3 | 43 | 350 | 107272238 | 107271928 | 5.43e-102 | 383 |
| Synaphobranchus_kaupii-CM056378.1 | 89.735 | 302 | 27 | 2 | 43 | 340 | 112005935 | 112005634 | 5.43e-102 | 383 |
| Synaphobranchus_kaupii-CM056378.1 | 89.103 | 312 | 29 | 3 | 43 | 350 | 125800852 | 125801162 | 5.43e-102 | 383 |
| Synaphobranchus_kaupii-CM056378.1 | 88.644 | 317 | 30 | 4 | 38 | 350 | 25856844 | 25856530 | 1.95e-101 | 381 |
| Synaphobranchus_kaupii-CM056378.1 | 88.889 | 315 | 28 | 5 | 43 | 353 | 43727337 | 43727026 | 1.95e-101 | 381 |
| Synaphobranchus_kaupii-CM056378.1 | 88.889 | 315 | 27 | 4 | 43 | 350 | 62821288 | 62821601 | 1.95e-101 | 381 |
| Synaphobranchus_kaupii-CM056378.1 | 88.924 | 316 | 26 | 4 | 43 | 350 | 99830796 | 99831110 | 1.95e-101 | 381 |
| Synaphobranchus_kaupii-CM056378.1 | 88.818 | 313 | 29 | 4 | 43 | 350 | 12852421 | 12852110 | 7.03e-101 | 379 |
| Synaphobranchus_kaupii-CM056378.1 | 88.782 | 312 | 30 | 3 | 43 | 350 | 5079217 | 5079527 | 2.53e-100 | 377 |
| Synaphobranchus_kaupii-CM056378.1 | 88.818 | 313 | 28 | 5 | 43 | 350 | 43719638 | 43719328 | 2.53e-100 | 377 |
| Synaphobranchus_kaupii-CM056378.1 | 88.782 | 312 | 30 | 3 | 43 | 350 | 62825044 | 62825354 | 2.53e-100 | 377 |
| Synaphobranchus_kaupii-CM056378.1 | 88.782 | 312 | 30 | 3 | 43 | 350 | 95577127 | 95576817 | 2.53e-100 | 377 |
| Synaphobranchus_kaupii-CM056378.1 | 88.782 | 312 | 30 | 3 | 43 | 350 | 95582380 | 95582690 | 2.53e-100 | 377 |
| Synaphobranchus_kaupii-CM056378.1 | 89.369 | 301 | 28 | 2 | 43 | 339 | 69396835 | 69396535 | 9.09e-100 | 375 |
| Synaphobranchus_kaupii-CM056378.1 | 88.462 | 312 | 31 | 3 | 43 | 350 | 14728987 | 14728677 | 1.18e-98 | 372 |
| Synaphobranchus_kaupii-CM056378.1 | 88.462 | 312 | 31 | 3 | 43 | 350 | 46779453 | 46779143 | 1.18e-98 | 372 |
| Synaphobranchus_kaupii-CM056378.1 | 88.462 | 312 | 31 | 3 | 43 | 350 | 99824966 | 99825276 | 1.18e-98 | 372 |
| Synaphobranchus_kaupii-CM056378.1 | 88.704 | 301 | 30 | 2 | 43 | 339 | 124130360 | 124130060 | 1.97e-96 | 364 |
| Synaphobranchus_kaupii-CM056386.1 | 91.667 | 504 | 32 | 6 | 43 | 536 | 2552647 | 2553150 | 0.0 | 689 |
| Synaphobranchus_kaupii-CM056386.1 | 90.945 | 508 | 32 | 8 | 43 | 536 | 33549245 | 33549752 | 0.0 | 671 |

### N5.HOG0008885 Blastn of conserved regions against all genome assemblies

#### Region A (2/10)

| sseqid | pident | length | mismatch | gapopen | qstart | qend | sstart | send | evalue | bitscore |
|---|---|---|---|---|---|---|---|---|---|---|
| Synaphobranchus_kaupii-CM056386.1 | 88.952 | 525 | 27 | 20 | 43 | 536 | 21670621 | 21670097 | 3.80e-173 | 619 |
| Synaphobranchus_kaupii-CM056386.1 | 88.247 | 519 | 32 | 14 | 43 | 536 | 30575942 | 30575428 | 2.31e-165 | 593 |
| Synaphobranchus_kaupii-CM056386.1 | 87.554 | 466 | 25 | 10 | 43 | 477 | 9925263 | 9925726 | 8.59e-140 | 508 |
| Synaphobranchus_kaupii-CM056386.1 | 89.644 | 309 | 28 | 2 | 43 | 347 | 23917303 | 23916995 | 3.25e-104 | 390 |
| Synaphobranchus_kaupii-CM056386.1 | 88.959 | 317 | 29 | 4 | 38 | 350 | 9957150 | 9956836 | 4.20e-103 | 387 |
| Synaphobranchus_kaupii-CM056386.1 | 89.597 | 298 | 26 | 4 | 38 | 335 | 42329445 | 42329737 | 3.27e-99 | 374 |
| Synaphobranchus_kaupii-CM056386.1 | 88.462 | 312 | 31 | 3 | 43 | 350 | 23913800 | 23914110 | 1.18e-98 | 372 |
| Synaphobranchus_kaupii-CM056386.1 | 89.492 | 295 | 25 | 4 | 43 | 335 | 10043251 | 10043541 | 1.52e-97 | 368 |
| Synaphobranchus_kaupii-CM056386.1 | 88.816 | 304 | 23 | 10 | 43 | 340 | 33959789 | 33959491 | 7.08e-96 | 363 |
| Synaphobranchus_kaupii-CM056386.1 | 92.627 | 217 | 14 | 1 | 43 | 257 | 16181564 | 16181348 | 2.60e-80 | 311 |
| Synaphobranchus_kaupii-CM056386.1 | 87.192 | 203 | 23 | 2 | 150 | 350 | 21668222 | 21668423 | 2.69e-55 | 228 |
| Synaphobranchus_kaupii-CM056386.1 | 85.235 | 149 | 19 | 2 | 204 | 350 | 21221270 | 21221417 | 6.00e-32 | 150 |
| Synaphobranchus_kaupii-CM056384.1 | 91.667 | 504 | 32 | 6 | 43 | 536 | 5407141 | 5407644 | 0.0 | 689 |
| Synaphobranchus_kaupii-CM056384.1 | 88.952 | 525 | 27 | 14 | 43 | 536 | 10851306 | 10851830 | 3.80e-173 | 619 |
| Synaphobranchus_kaupii-CM056384.1 | 89.423 | 312 | 28 | 3 | 43 | 350 | 67177811 | 67178121 | 1.17e-103 | 388 |
| Synaphobranchus_kaupii-CM056384.1 | 89.032 | 310 | 25 | 4 | 43 | 345 | 59591497 | 59591804 | 9.09e-100 | 375 |
| Synaphobranchus_kaupii-CM056380.1 | 91.667 | 504 | 32 | 6 | 43 | 536 | 59402517 | 59403020 | 0.0 | 689 |
| Synaphobranchus_kaupii-CM056380.1 | 87.833 | 526 | 29 | 16 | 43 | 536 | 50963978 | 50963456 | 1.39e-162 | 584 |
| Synaphobranchus_kaupii-CM056380.1 | 89.524 | 315 | 25 | 4 | 43 | 350 | 55939203 | 55938890 | 9.03e-105 | 392 |
| Synaphobranchus_kaupii-CM056380.1 | 89.423 | 312 | 28 | 3 | 43 | 350 | 16752645 | 16752335 | 1.17e-103 | 388 |
| Synaphobranchus_kaupii-CM056380.1 | 89.423 | 312 | 28 | 3 | 43 | 350 | 21387838 | 21387528 | 1.17e-103 | 388 |
| Synaphobranchus_kaupii-CM056380.1 | 89.423 | 312 | 28 | 3 | 43 | 350 | 47961734 | 47962044 | 1.17e-103 | 388 |
| Synaphobranchus_kaupii-CM056380.1 | 89.423 | 312 | 28 | 3 | 43 | 350 | 64661178 | 64660868 | 1.17e-103 | 388 |
| Synaphobranchus_kaupii-CM056380.1 | 89.423 | 312 | 28 | 3 | 43 | 350 | 112818790 | 112819100 | 1.17e-103 | 388 |
| Synaphobranchus_kaupii-CM056380.1 | 90.333 | 300 | 22 | 3 | 43 | 335 | 374091 | 373792 | 4.20e-103 | 387 |
| Synaphobranchus_kaupii-CM056380.1 | 89.206 | 315 | 26 | 4 | 43 | 350 | 84899359 | 84899046 | 4.20e-103 | 387 |
| Synaphobranchus_kaupii-CM056380.1 | 89.836 | 305 | 24 | 4 | 38 | 340 | 60520115 | 60520414 | 1.51e-102 | 385 |
| Synaphobranchus_kaupii-CM056380.1 | 89.103 | 312 | 29 | 3 | 43 | 350 | 49644270 | 49643960 | 5.43e-102 | 383 |
| Synaphobranchus_kaupii-CM056380.1 | 89.103 | 312 | 29 | 3 | 43 | 350 | 60529352 | 60529042 | 5.43e-102 | 383 |
| Synaphobranchus_kaupii-CM056380.1 | 89.803 | 304 | 24 | 3 | 43 | 339 | 110956904 | 110956601 | 5.43e-102 | 383 |
| Synaphobranchus_kaupii-CM056380.1 | 89.103 | 312 | 29 | 3 | 43 | 350 | 110972337 | 110972647 | 5.43e-102 | 383 |
| Synaphobranchus_kaupii-CM056380.1 | 89.701 | 301 | 27 | 2 | 43 | 339 | 32919445 | 32919145 | 1.95e-101 | 381 |
| Synaphobranchus_kaupii-CM056380.1 | 89.701 | 301 | 27 | 3 | 43 | 339 | 51573397 | 51573697 | 1.95e-101 | 381 |
| Synaphobranchus_kaupii-CM056380.1 | 89.701 | 301 | 27 | 2 | 43 | 339 | 51575764 | 51575464 | 1.95e-101 | 381 |
| Synaphobranchus_kaupii-CM056380.1 | 88.644 | 317 | 28 | 6 | 38 | 350 | 114884553 | 114884241 | 7.03e-101 | 379 |
| Synaphobranchus_kaupii-CM056380.1 | 88.818 | 313 | 29 | 4 | 43 | 350 | 115182763 | 115182452 | 7.03e-101 | 379 |
| Synaphobranchus_kaupii-CM056380.1 | 88.782 | 312 | 30 | 3 | 43 | 350 | 47967644 | 47967334 | 2.53e-100 | 377 |
| Synaphobranchus_kaupii-CM056380.1 | 88.782 | 312 | 30 | 3 | 43 | 350 | 66492592 | 66492902 | 2.53e-100 | 377 |
| Synaphobranchus_kaupii-CM056380.1 | 88.782 | 312 | 30 | 3 | 43 | 350 | 104067606 | 104067916 | 2.53e-100 | 377 |
| Synaphobranchus_kaupii-CM056380.1 | 89.831 | 295 | 24 | 4 | 43 | 335 | 101051164 | 101051454 | 3.27e-99 | 374 |
| Synaphobranchus_kaupii-CM056380.1 | 89.369 | 301 | 23 | 5 | 43 | 339 | 36789681 | 36789386 | 4.23e-98 | 370 |
| Synaphobranchus_kaupii-CM056380.1 | 95.000 | 220 | 11 | 0 | 317 | 536 | 36789366 | 36789147 | 7.13e-91 | 346 |
| Synaphobranchus_kaupii-CM056380.1 | 83.912 | 317 | 24 | 4 | 43 | 352 | 110969430 | 110969134 | 2.64e-70 | 278 |
| Synaphobranchus_kaupii-CM056380.1 | 83.654 | 312 | 26 | 4 | 43 | 350 | 51572234 | 51571944 | 4.41e-68 | 270 |
| Synaphobranchus_kaupii-CM056380.1 | 83.175 | 315 | 27 | 5 | 43 | 353 | 32900188 | 32899896 | 2.05e-66 | 265 |
| Coryphaenoides_rupestris-PIJE02000202.1 | 91.683 | 505 | 31 | 6 | 43 | 536 | 509108 | 508604 | 0.0 | 689 |
| Synaphobranchus_kaupii-CM056385.1 | 91.617 | 501 | 35 | 5 | 38 | 536 | 47140097 | 47139602 | 0.0 | 686 |
| Synaphobranchus_kaupii-CM056385.1 | 89.744 | 312 | 27 | 3 | 43 | 350 | 58698256 | 58698566 | 2.51e-105 | 394 |
| Synaphobranchus_kaupii-CM056385.1 | 89.744 | 312 | 27 | 3 | 43 | 350 | 58946125 | 58945815 | 2.51e-105 | 394 |
| Synaphobranchus_kaupii-CM056385.1 | 90.365 | 301 | 25 | 2 | 43 | 339 | 49520690 | 49520990 | 9.03e-105 | 392 |
| Synaphobranchus_kaupii-CM056385.1 | 90.066 | 302 | 26 | 2 | 43 | 340 | 49965210 | 49964909 | 1.17e-103 | 388 |
| Synaphobranchus_kaupii-CM056385.1 | 89.423 | 312 | 28 | 3 | 43 | 350 | 50069768 | 50070078 | 1.17e-103 | 388 |
| Synaphobranchus_kaupii-CM056385.1 | 88.959 | 317 | 29 | 4 | 38 | 350 | 34394782 | 34395096 | 4.20e-103 | 387 |
| Synaphobranchus_kaupii-CM056385.1 | 89.137 | 313 | 28 | 4 | 43 | 350 | 16861536 | 16861847 | 1.51e-102 | 385 |
| Synaphobranchus_kaupii-CM056385.1 | 88.994 | 318 | 26 | 5 | 43 | 352 | 28324039 | 28323723 | 1.51e-102 | 385 |
| Synaphobranchus_kaupii-CM056385.1 | 88.994 | 318 | 26 | 5 | 43 | 352 | 32975051 | 32975367 | 1.51e-102 | 385 |
| Synaphobranchus_kaupii-CM056385.1 | 89.542 | 306 | 27 | 3 | 38 | 339 | 3088814 | 3089118 | 5.43e-102 | 383 |
| Synaphobranchus_kaupii-CM056385.1 | 89.103 | 312 | 29 | 3 | 43 | 350 | 43483131 | 43482821 | 5.43e-102 | 383 |
| Synaphobranchus_kaupii-CM056385.1 | 89.103 | 312 | 29 | 3 | 43 | 350 | 46853664 | 46853974 | 5.43e-102 | 383 |
| Synaphobranchus_kaupii-CM056385.1 | 89.735 | 302 | 25 | 4 | 43 | 339 | 27943626 | 27943926 | 1.95e-101 | 381 |
| Synaphobranchus_kaupii-CM056385.1 | 89.508 | 305 | 25 | 3 | 43 | 340 | 58703554 | 58703858 | 7.03e-101 | 379 |
| Synaphobranchus_kaupii-CM056385.1 | 88.818 | 313 | 28 | 5 | 43 | 350 | 13369315 | 13369625 | 2.53e-100 | 377 |
| Synaphobranchus_kaupii-CM056385.1 | 88.782 | 312 | 30 | 3 | 43 | 350 | 27950024 | 27950334 | 2.53e-100 | 377 |
| Synaphobranchus_kaupii-CM056385.1 | 88.782 | 312 | 30 | 3 | 43 | 350 | 28316248 | 28316558 | 2.53e-100 | 377 |
| Synaphobranchus_kaupii-CM056385.1 | 89.369 | 301 | 28 | 2 | 43 | 339 | 28317785 | 28317485 | 9.09e-100 | 375 |
| Synaphobranchus_kaupii-CM056385.1 | 88.571 | 315 | 26 | 5 | 43 | 347 | 58955456 | 58955770 | 3.27e-99 | 374 |
| Synaphobranchus_kaupii-CM056385.1 | 88.462 | 312 | 31 | 3 | 43 | 350 | 49525235 | 49524925 | 1.18e-98 | 372 |
| Synaphobranchus_kaupii-CM056385.1 | 87.697 | 317 | 29 | 4 | 43 | 350 | 36361719 | 36361404 | 2.55e-95 | 361 |
| Synaphobranchus_kaupii-CM056385.1 | 89.273 | 289 | 23 | 6 | 43 | 329 | 55993719 | 55993437 | 1.18e-93 | 355 |
| Synaphobranchus_kaupii-CM056385.1 | 87.227 | 321 | 28 | 8 | 43 | 356 | 56000409 | 56000723 | 4.26e-93 | 353 |
| Synaphobranchus_kaupii-CM056385.1 | 85.276 | 163 | 22 | 2 | 347 | 508 | 26602911 | 26602750 | 5.96e-37 | 167 |
| Synaphobranchus_kaupii-CM056385.1 | 97.561 | 82 | 2 | 0 | 396 | 477 | 55993438 | 55993357 | 3.61e-29 | 141 |
| Synaphobranchus_kaupii- | 91.468 | 504 | 33 | 6 | 43 | 536 | 533674 | 533171 | 0.0 | 684 |
| Synaphobranchus_kaupii- | 89.103 | 312 | 29 | 3 | 43 | 350 | 2459235 | 2459545 | 5.43e-102 | 383 |
| Synaphobranchus_kaupii- | 89.701 | 301 | 27 | 3 | 43 | 339 | 532061 | 531761 | 1.95e-101 | 381 |
| Clupea_harengus-NC_045175.1 | 91.182 | 499 | 39 | 2 | 43 | 536 | 11408026 | 11408524 | 0.0 | 673 |
| Clupea_harengus-NC_045175.1 | 89.938 | 487 | 47 | 2 | 52 | 536 | 7152889 | 7152403 | 2.27e-175 | 627 |
| Clupea_harengus-NC_045175.1 | 89.066 | 503 | 46 | 6 | 43 | 536 | 7135381 | 7135883 | 4.92e-172 | 616 |
| Clupea_harengus-NC_045175.1 | 89.336 | 497 | 41 | 6 | 52 | 536 | 4812949 | 4812453 | 1.77e-171 | 614 |
| Clupea_harengus-NC_045175.1 | 88.735 | 506 | 45 | 6 | 43 | 536 | 5461866 | 5462371 | 8.23e-170 | 608 |
| Clupea_harengus-NC_045175.1 | 88.735 | 506 | 45 | 6 | 43 | 536 | 9279907 | 9280412 | 8.23e-170 | 608 |
| Clupea_harengus-NC_045175.1 | 88.735 | 506 | 44 | 7 | 43 | 536 | 7066682 | 7067186 | 2.96e-169 | 606 |
| Clupea_harengus-NC_045175.1 | 88.477 | 512 | 33 | 11 | 43 | 536 | 10493133 | 10493636 | 6.41e-166 | 595 |
| Clupea_harengus-NC_045175.1 | 88.664 | 494 | 44 | 7 | 52 | 536 | 9668180 | 9667690 | 8.29e-165 | 592 |
| Clupea_harengus-NC_045175.1 | 88.469 | 503 | 39 | 13 | 52 | 536 | 17737575 | 17737074 | 2.98e-164 | 590 |
| Clupea_harengus-NC_045175.1 | 88.333 | 480 | 41 | 9 | 52 | 516 | 7066018 | 7066497 | 6.50e-156 | 562 |
| Clupea_harengus-NC_045175.1 | 86.796 | 515 | 47 | 12 | 43 | 536 | 9669100 | 9669614 | 1.09e-153 | 555 |
| Clupea_harengus-NC_045175.1 | 83.083 | 532 | 50 | 21 | 38 | 536 | 9278491 | 9279015 | 1.90e-121 | 448 |
| Clupea_harengus-NC_045175.1 | 83.936 | 498 | 42 | 11 | 43 | 536 | 3891454 | 3891917 | 8.84e-120 | 442 |
| Clupea_harengus-NC_045175.1 | 83.846 | 390 | 36 | 14 | 147 | 536 | 11159433 | 11159071 | 7.13e-91 | 346 |
| Clupea_harengus-NC_045175.1 | 93.805 | 226 | 14 | 0 | 311 | 536 | 10544980 | 10544755 | 3.32e-89 | 340 |
| Clupea_harengus-NC_045175.1 | 83.249 | 394 | 32 | 17 | 147 | 537 | 11107579 | 11107941 | 2.00e-86 | 331 |
| Clupea_harengus-NC_045175.1 | 86.219 | 283 | 34 | 4 | 43 | 321 | 10545900 | 10545619 | 1.56e-77 | 302 |
| Clupea_harengus-NC_045175.1 | 93.103 | 145 | 9 | 1 | 43 | 186 | 15925904 | 15926048 | 2.71e-50 | 211 |
| Clupea_harengus-NC_045175.1 | 91.200 | 125 | 10 | 1 | 43 | 166 | 11158111 | 11158235 | 1.66e-37 | 169 |
| Coryphaenoides_rupestris-PIJE02006124.1 | 91.700 | 494 | 28 | 6 | 43 | 536 | 13176 | 12696 | 0.0 | 673 |
| Clupea_harengus-NC_045154.1 | 91.111 | 495 | 35 | 7 | 43 | 536 | 15154514 | 15154028 | 0.0 | 662 |
| Clupea_harengus-NC_045154.1 | 89.980 | 499 | 38 | 6 | 43 | 536 | 990397 | 990888 | 1.36e-177 | 634 |
| Clupea_harengus-NC_045154.1 | 89.271 | 494 | 44 | 5 | 52 | 536 | 21262099 | 21261606 | 2.29e-170 | 610 |
| Clupea_harengus-NC_045154.1 | 88.664 | 494 | 42 | 11 | 52 | 536 | 15165841 | 15165353 | 2.98e-164 | 590 |
| Clupea_harengus-NC_045154.1 | 88.016 | 509 | 46 | 7 | 43 | 536 | 9572965 | 9573473 | 1.07e-163 | 588 |

### N5.HOG0008885 Blastn of conserved regions against all genome assemblies

#### Region A (3/10)

| sseqid | pident | length | mismatch | gapopen | qstart | qend | sstart | send | evalue | bitscore |
|---|---|---|---|---|---|---|---|---|---|---|
| Clupea_harengus-NC_045154.1 | 88.400 | 500 | 43 | 7 | 52 | 536 | 28107431 | 28106932 | 1.07e-163 | 588 |
| Clupea_harengus-NC_045154.1 | 87.945 | 506 | 49 | 6 | 43 | 536 | 20015725 | 20016230 | 3.86e-163 | 586 |
| Clupea_harengus-NC_045154.1 | 87.305 | 512 | 47 | 11 | 43 | 536 | 17352262 | 17352773 | 3.89e-158 | 569 |
| Clupea_harengus-NC_045154.1 | 87.402 | 508 | 47 | 7 | 43 | 533 | 15163348 | 15163855 | 1.40e-157 | 568 |
| Clupea_harengus-NC_045154.1 | 86.756 | 521 | 42 | 21 | 43 | 536 | 29786942 | 29787462 | 1.09e-153 | 555 |
| Clupea_harengus-NC_045154.1 | 86.561 | 506 | 44 | 9 | 41 | 536 | 12963564 | 12963073 | 3.94e-148 | 536 |
| Clupea_harengus-NC_045154.1 | 86.263 | 495 | 54 | 9 | 43 | 536 | 17351424 | 17351905 | 8.53e-145 | 525 |
| Clupea_harengus-NC_045154.1 | 85.472 | 530 | 41 | 24 | 43 | 536 | 24863122 | 24863651 | 3.97e-143 | 520 |
| Clupea_harengus-NC_045154.1 | 90.964 | 166 | 12 | 2 | 43 | 207 | 16567168 | 16567331 | 4.51e-53 | 220 |
| Synaphobranchus_kaupii-CM056396.1 | 90.643 | 513 | 26 | 12 | 43 | 536 | 23717984 | 23717475 | 0.0 | 662 |
| Synaphobranchus_kaupii-CM056396.1 | 89.524 | 315 | 25 | 4 | 43 | 350 | 392557 | 392244 | 9.03e-105 | 392 |
| Synaphobranchus_kaupii-CM056396.1 | 89.423 | 312 | 28 | 3 | 43 | 350 | 493584 | 493274 | 1.17e-103 | 388 |
| Synaphobranchus_kaupii-CM056396.1 | 88.959 | 317 | 29 | 4 | 38 | 350 | 1264113 | 1263799 | 4.20e-103 | 387 |
| Synaphobranchus_kaupii-CM056396.1 | 89.836 | 305 | 24 | 4 | 43 | 340 | 1151445 | 1151749 | 1.51e-102 | 385 |
| Synaphobranchus_kaupii-CM056396.1 | 89.137 | 313 | 28 | 4 | 43 | 350 | 29008721 | 29008410 | 1.51e-102 | 385 |
| Synaphobranchus_kaupii-CM056396.1 | 89.701 | 301 | 27 | 2 | 43 | 339 | 22080139 | 22079839 | 1.95e-101 | 381 |
| Synaphobranchus_kaupii-CM056396.1 | 88.438 | 320 | 30 | 5 | 43 | 356 | 29009981 | 29009663 | 7.03e-101 | 379 |
| Clupea_harengus-NC_045164.1 | 90.818 | 501 | 35 | 4 | 43 | 536 | 28641043 | 28640547 | 0.0 | 660 |
| Clupea_harengus-NC_045164.1 | 88.933 | 506 | 44 | 6 | 43 | 536 | 14892924 | 14893429 | 1.77e-171 | 614 |
| Clupea_harengus-NC_045164.1 | 88.735 | 506 | 45 | 6 | 43 | 536 | 7568628 | 7569133 | 8.23e-170 | 608 |
| Clupea_harengus-NC_045164.1 | 88.623 | 501 | 49 | 6 | 43 | 536 | 8559688 | 8560187 | 3.83e-168 | 603 |
| Clupea_harengus-NC_045164.1 | 88.600 | 500 | 51 | 4 | 43 | 536 | 17861794 | 17862293 | 3.83e-168 | 603 |
| Clupea_harengus-NC_045164.1 | 88.538 | 506 | 45 | 8 | 43 | 536 | 3682928 | 3683432 | 1.38e-167 | 601 |
| Clupea_harengus-NC_045164.1 | 88.409 | 509 | 44 | 7 | 43 | 536 | 19553641 | 19554149 | 4.96e-167 | 599 |
| Clupea_harengus-NC_045164.1 | 88.732 | 497 | 44 | 6 | 52 | 536 | 4679841 | 4679345 | 1.78e-166 | 597 |
| Clupea_harengus-NC_045164.1 | 88.400 | 500 | 43 | 7 | 52 | 536 | 22647245 | 22646746 | 1.07e-163 | 588 |
| Clupea_harengus-NC_045164.1 | 87.549 | 506 | 42 | 12 | 52 | 536 | 8565404 | 8564899 | 5.03e-157 | 566 |
| Clupea_harengus-NC_045164.1 | 87.084 | 511 | 40 | 19 | 52 | 536 | 1736621 | 1736111 | 1.09e-153 | 555 |
| Clupea_harengus-NC_045164.1 | 90.526 | 285 | 23 | 3 | 43 | 327 | 16838490 | 16838210 | 3.27e-99 | 374 |
| Clupea_harengus-NC_045160.1 | 90.354 | 508 | 36 | 7 | 41 | 536 | 30193594 | 30193088 | 0.0 | 654 |
| Clupea_harengus-NC_045160.1 | 90.297 | 505 | 38 | 7 | 43 | 536 | 25913664 | 25913160 | 0.0 | 651 |
| Clupea_harengus-NC_045160.1 | 90.631 | 491 | 26 | 7 | 46 | 536 | 7941280 | 7940810 | 1.36e-177 | 634 |
| Clupea_harengus-NC_045160.1 | 88.933 | 506 | 44 | 6 | 43 | 536 | 20762310 | 20762815 | 1.77e-171 | 614 |
| Clupea_harengus-NC_045160.1 | 88.802 | 509 | 41 | 11 | 41 | 536 | 20469670 | 20469165 | 2.29e-170 | 610 |
| Clupea_harengus-NC_045160.1 | 89.184 | 490 | 46 | 5 | 54 | 537 | 17661748 | 17661260 | 1.07e-168 | 604 |
| Clupea_harengus-NC_045160.1 | 88.538 | 506 | 46 | 6 | 43 | 536 | 20760159 | 20760664 | 3.83e-168 | 603 |
| Clupea_harengus-NC_045160.1 | 88.732 | 497 | 44 | 6 | 52 | 536 | 20279415 | 20278919 | 1.78e-166 | 597 |
| Clupea_harengus-NC_045160.1 | 88.039 | 510 | 46 | 8 | 43 | 537 | 18783087 | 18782578 | 2.98e-164 | 590 |
| Clupea_harengus-NC_045160.1 | 88.072 | 503 | 42 | 11 | 52 | 536 | 10805058 | 10804556 | 1.79e-161 | 580 |
| Clupea_harengus-NC_045160.1 | 87.574 | 507 | 50 | 8 | 43 | 536 | 365244 | 364738 | 8.35e-160 | 575 |
| Clupea_harengus-NC_045160.1 | 87.835 | 485 | 41 | 8 | 52 | 518 | 10803313 | 10802829 | 3.91e-153 | 553 |
| Clupea_harengus-NC_045160.1 | 83.202 | 506 | 47 | 9 | 45 | 536 | 27523715 | 27524196 | 6.88e-116 | 429 |
| Clupea_harengus-NC_045160.1 | 83.004 | 506 | 48 | 9 | 45 | 536 | 27515802 | 27516283 | 3.20e-114 | 424 |
| Clupea_harengus-NC_045160.1 | 83.940 | 467 | 42 | 6 | 52 | 507 | 27577717 | 27577273 | 5.36e-112 | 416 |
| Clupea_harengus-NC_045160.1 | 87.261 | 314 | 24 | 10 | 238 | 536 | 17743097 | 17743409 | 2.56e-90 | 344 |
| Clupea_harengus-NC_045160.1 | 85.304 | 313 | 40 | 5 | 41 | 350 | 7532769 | 7532460 | 1.55e-82 | 318 |
| Clupea_harengus-NC_045160.1 | 94.536 | 183 | 10 | 0 | 43 | 225 | 363206 | 363388 | 5.67e-72 | 283 |
| Clupea_harengus-NC_045160.1 | 89.151 | 212 | 17 | 4 | 326 | 536 | 27606980 | 27606774 | 9.55e-65 | 259 |
| Clupea_harengus-NC_045160.1 | 90.860 | 186 | 13 | 3 | 43 | 227 | 27443862 | 27443680 | 7.44e-61 | 246 |
| Clupea_harengus-NC_045160.1 | 94.040 | 151 | 9 | 0 | 43 | 193 | 27607131 | 27606981 | 7.49e-56 | 230 |
| Clupea_harengus-NC_045160.1 | 93.284 | 134 | 7 | 2 | 45 | 176 | 27445182 | 27445049 | 7.60e-46 | 196 |
| Clupea_harengus-NC_045173.1 | 90.909 | 495 | 32 | 7 | 43 | 536 | 10011698 | 10011216 | 0.0 | 652 |
| Clupea_harengus-NC_045173.1 | 89.336 | 497 | 41 | 6 | 52 | 536 | 21090032 | 21089536 | 1.77e-171 | 614 |
| Clupea_harengus-NC_045173.1 | 88.538 | 506 | 46 | 6 | 43 | 536 | 17481585 | 17482090 | 3.83e-168 | 603 |
| Clupea_harengus-NC_045173.1 | 88.538 | 506 | 46 | 6 | 43 | 536 | 19795967 | 19796472 | 3.83e-168 | 603 |
| Clupea_harengus-NC_045173.1 | 88.800 | 500 | 41 | 8 | 52 | 536 | 7610560 | 7610061 | 4.96e-167 | 599 |
| Clupea_harengus-NC_045173.1 | 88.800 | 500 | 41 | 8 | 52 | 536 | 13020858 | 13020359 | 4.96e-167 | 599 |
| Clupea_harengus-NC_045173.1 | 88.363 | 507 | 47 | 7 | 41 | 536 | 18597546 | 18597041 | 4.96e-167 | 599 |
| Clupea_harengus-NC_045173.1 | 88.340 | 506 | 47 | 7 | 43 | 536 | 22774702 | 22775207 | 1.78e-166 | 597 |
| Clupea_harengus-NC_045173.1 | 88.212 | 509 | 45 | 7 | 43 | 536 | 11151749 | 11152257 | 2.31e-165 | 593 |
| Clupea_harengus-NC_045173.1 | 88.200 | 500 | 44 | 9 | 52 | 536 | 1048441 | 1047942 | 4.99e-162 | 582 |
| Clupea_harengus-NC_045173.1 | 87.549 | 514 | 44 | 12 | 43 | 536 | 7416003 | 7415490 | 2.32e-160 | 577 |
| Clupea_harengus-NC_045173.1 | 84.877 | 529 | 45 | 23 | 43 | 536 | 7416393 | 7416921 | 1.44e-137 | 501 |
| Clupea_harengus-NC_045173.1 | 89.752 | 322 | 26 | 4 | 43 | 362 | 2753222 | 2753538 | 1.16e-108 | 405 |
| Clupea_harengus-NC_045173.1 | 88.785 | 321 | 25 | 5 | 227 | 536 | 19826430 | 19826750 | 5.43e-102 | 383 |
| Clupea_harengus-NC_045173.1 | 90.217 | 276 | 13 | 10 | 108 | 377 | 8488419 | 8488686 | 1.98e-91 | 348 |
| Clupea_harengus-NC_045173.1 | 85.061 | 328 | 37 | 8 | 43 | 358 | 6911049 | 6911376 | 3.34e-84 | 324 |
| Clupea_harengus-NC_045173.1 | 95.882 | 170 | 7 | 0 | 367 | 536 | 18674723 | 18674554 | 9.49e-70 | 276 |
| Clupea_harengus-NC_045173.1 | 95.420 | 131 | 6 | 0 | 406 | 536 | 8488684 | 8488814 | 9.76e-50 | 209 |
| Clupea_harengus-NC_045173.1 | 93.985 | 133 | 8 | 0 | 404 | 536 | 6912909 | 6913041 | 1.63e-47 | 202 |
| Clupea_harengus-NC_045162.1 | 90.361 | 498 | 44 | 2 | 43 | 536 | 18437938 | 18438435 | 0.0 | 651 |
| Clupea_harengus-NC_045162.1 | 90.020 | 491 | 43 | 4 | 52 | 536 | 17031175 | 17030685 | 1.76e-176 | 630 |
| Clupea_harengus-NC_045162.1 | 89.251 | 521 | 30 | 9 | 41 | 536 | 4593890 | 4593371 | 6.32e-176 | 628 |
| Clupea_harengus-NC_045162.1 | 89.400 | 500 | 47 | 4 | 43 | 536 | 15417867 | 15418366 | 8.18e-175 | 625 |
| Clupea_harengus-NC_045162.1 | 89.676 | 494 | 42 | 5 | 52 | 536 | 4519976 | 4519483 | 1.06e-173 | 621 |
| Clupea_harengus-NC_045162.1 | 89.676 | 494 | 42 | 5 | 52 | 536 | 4555533 | 4555040 | 1.06e-173 | 621 |
| Clupea_harengus-NC_045162.1 | 89.613 | 491 | 38 | 6 | 52 | 536 | 15407955 | 15407472 | 6.36e-171 | 612 |
| Clupea_harengus-NC_045162.1 | 88.016 | 509 | 46 | 7 | 43 | 536 | 11427599 | 11428107 | 1.07e-163 | 588 |
| Clupea_harengus-NC_045162.1 | 88.016 | 509 | 46 | 7 | 43 | 536 | 24273214 | 24273722 | 1.07e-163 | 588 |
| Clupea_harengus-NC_045162.1 | 88.400 | 500 | 43 | 7 | 52 | 536 | 27532757 | 27532258 | 1.07e-163 | 588 |
| Clupea_harengus-NC_045162.1 | 87.945 | 506 | 40 | 12 | 52 | 536 | 10582461 | 10581956 | 2.32e-160 | 577 |
| Clupea_harengus-NC_045162.1 | 86.519 | 497 | 62 | 3 | 43 | 536 | 26454864 | 26455358 | 8.47e-150 | 542 |
| Clupea_harengus-NC_045162.1 | 86.187 | 514 | 50 | 11 | 43 | 536 | 10239494 | 10238982 | 3.94e-148 | 536 |
| Clupea_harengus-NC_045162.1 | 85.910 | 511 | 55 | 7 | 43 | 536 | 11778152 | 11777642 | 6.59e-146 | 529 |
| Clupea_harengus-NC_045162.1 | 85.825 | 515 | 44 | 10 | 38 | 540 | 27530717 | 27531214 | 3.97e-143 | 520 |
| Clupea_harengus-NC_045162.1 | 85.630 | 508 | 56 | 15 | 43 | 536 | 4512514 | 4512010 | 1.43e-142 | 518 |
| Clupea_harengus-NC_045162.1 | 83.149 | 362 | 38 | 12 | 43 | 392 | 29834172 | 29834522 | 9.35e-80 | 309 |
| Clupea_harengus-NC_045162.1 | 93.865 | 163 | 10 | 0 | 374 | 536 | 2025210 | 2025048 | 7.44e-61 | 246 |
| Clupea_harengus-NC_045162.1 | 93.750 | 160 | 8 | 2 | 43 | 201 | 2025373 | 2025215 | 1.24e-58 | 239 |
| Clupea_harengus-NC_045167.1 | 90.239 | 502 | 41 | 5 | 43 | 536 | 12736807 | 12737308 | 0.0 | 649 |
| Clupea_harengus-NC_045167.1 | 89.336 | 497 | 41 | 6 | 52 | 536 | 3342055 | 3341559 | 1.77e-171 | 614 |
| Clupea_harengus-NC_045167.1 | 88.713 | 505 | 50 | 5 | 38 | 536 | 1631292 | 1631795 | 2.29e-170 | 610 |
| Clupea_harengus-NC_045167.1 | 88.538 | 506 | 50 | 6 | 38 | 536 | 1608062 | 1608566 | 2.96e-169 | 606 |
| Clupea_harengus-NC_045167.1 | 89.069 | 494 | 45 | 5 | 52 | 536 | 17087517 | 17087024 | 1.07e-168 | 604 |
| Clupea_harengus-NC_045167.1 | 88.538 | 506 | 46 | 6 | 43 | 536 | 12481885 | 12482390 | 3.83e-168 | 603 |
| Clupea_harengus-NC_045167.1 | 88.732 | 497 | 44 | 6 | 52 | 536 | 16582610 | 16582114 | 1.78e-166 | 597 |
| Clupea_harengus-NC_045167.1 | 89.375 | 480 | 36 | 5 | 63 | 536 | 6682225 | 6682695 | 2.98e-164 | 590 |

### N5.HOG0008885 Blastn of conserved regions against all genome assemblies

#### Region A (4/10)

| sseqid | pident | length | mismatch | gapopen | qstart | qend | sstart | send | evalue | bitscore |
|---|---|---|---|---|---|---|---|---|---|---|
| Clupea harengus-NC 045167.1 | 88.072 | 503 | 43 | 8 | 52 | 537 | 12483930 | 12483428 | 1.79e-161 | 580 |
| Clupea harengus-NC 045167.1 | 87.598 | 508 | 45 | 11 | 43 | 532 | 14800860 | 14801367 | 3.00e-159 | 573 |
| Clupea harengus-NC 045167.1 | 87.033 | 509 | 49 | 11 | 43 | 536 | 24540991 | 24541497 | 8.41e-155 | 558 |
| Clupea harengus-NC 045167.1 | 86.117 | 497 | 44 | 12 | 43 | 536 | 8466443 | 8466917 | 6.64e-141 | 512 |
| Clupea harengus-NC 045167.1 | 86.765 | 476 | 39 | 17 | 52 | 503 | 14800172 | 14799697 | 8.59e-140 | 508 |
| Clupea harengus-NC 045167.1 | 91.154 | 260 | 15 | 4 | 285 | 536 | 9060875 | 9060616 | 7.13e-91 | 346 |
| Clupea harengus-NC 045167.1 | 95.000 | 180 | 9 | 0 | 357 | 536 | 16581068 | 16580889 | 5.67e-72 | 283 |
| Clupea harengus-NC 045167.1 | 90.783 | 217 | 11 | 1 | 320 | 536 | 6010274 | 6010481 | 2.04e-71 | 281 |
| Clupea harengus-NC 045167.1 | 92.424 | 198 | 12 | 3 | 43 | 239 | 6010081 | 6010276 | 7.33e-71 | 279 |
| Clupea harengus-NC 045167.1 | 90.426 | 188 | 17 | 1 | 52 | 238 | 9061061 | 9060874 | 7.44e-61 | 246 |
| Clupea harengus-NC 045167.1 | 94.891 | 137 | 7 | 0 | 400 | 536 | 3811003 | 3811139 | 2.10e-51 | 215 |
| Clupea harengus-NC 045167.1 | 90.000 | 100 | 8 | 2 | 41 | 139 | 9381822 | 9381724 | 2.81e-25 | 128 |
| Clupea harengus-NC 045166.1 | 90.890 | 483 | 40 | 3 | 43 | 521 | 14514288 | 14514770 | 0.0 | 645 |
| Clupea harengus-NC 045166.1 | 89.400 | 500 | 47 | 4 | 43 | 536 | 9593990 | 9594489 | 8.18e-175 | 625 |
| Clupea harengus-NC 045166.1 | 89.130 | 506 | 43 | 6 | 43 | 536 | 14269265 | 14268760 | 3.80e-173 | 619 |
| Clupea harengus-NC 045166.1 | 89.336 | 497 | 41 | 6 | 52 | 536 | 14513605 | 14513109 | 1.77e-171 | 614 |
| Clupea harengus-NC 045166.1 | 89.271 | 494 | 44 | 5 | 52 | 536 | 2162291 | 2161798 | 2.29e-170 | 610 |
| Clupea harengus-NC 045166.1 | 89.243 | 502 | 32 | 9 | 41 | 536 | 17925597 | 17925112 | 8.23e-170 | 608 |
| Clupea harengus-NC 045166.1 | 88.538 | 506 | 46 | 6 | 43 | 536 | 10840846 | 10841351 | 3.83e-168 | 603 |
| Clupea harengus-NC 045166.1 | 88.538 | 506 | 46 | 6 | 43 | 536 | 10872940 | 10873445 | 3.83e-168 | 603 |
| Clupea harengus-NC 045166.1 | 88.934 | 497 | 43 | 6 | 52 | 536 | 12407006 | 12406510 | 3.83e-168 | 603 |
| Clupea harengus-NC 045166.1 | 88.538 | 506 | 46 | 6 | 43 | 536 | 13548785 | 13549290 | 3.83e-168 | 603 |
| Clupea harengus-NC 045166.1 | 88.934 | 497 | 43 | 6 | 52 | 536 | 14512401 | 14511905 | 3.83e-168 | 603 |
| Clupea harengus-NC 045166.1 | 88.492 | 504 | 48 | 6 | 43 | 536 | 11366062 | 11366565 | 1.38e-167 | 601 |
| Clupea harengus-NC 045166.1 | 88.800 | 500 | 41 | 7 | 52 | 536 | 14510170 | 14509671 | 4.96e-167 | 599 |
| Clupea harengus-NC 045166.1 | 88.212 | 509 | 45 | 7 | 43 | 536 | 15826320 | 15826828 | 2.31e-165 | 593 |
| Clupea harengus-NC 045166.1 | 88.086 | 512 | 43 | 10 | 43 | 536 | 20001155 | 20001666 | 8.29e-165 | 592 |
| Clupea harengus-NC 045166.1 | 88.000 | 500 | 45 | 7 | 52 | 536 | 4323971 | 4323472 | 2.32e-160 | 577 |
| Clupea harengus-NC 045166.1 | 87.072 | 526 | 40 | 21 | 38 | 536 | 16186413 | 16186937 | 3.89e-158 | 569 |
| Clupea harengus-NC 045166.1 | 87.576 | 491 | 41 | 12 | 52 | 536 | 11825747 | 11825271 | 1.41e-152 | 551 |
| Clupea harengus-NC 045166.1 | 89.051 | 411 | 36 | 5 | 43 | 444 | 22145288 | 22145698 | 1.44e-137 | 501 |
| Clupea harengus-NC 045166.1 | 85.714 | 497 | 41 | 8 | 52 | 536 | 7507072 | 7506594 | 1.86e-136 | 497 |
| Clupea harengus-NC 045166.1 | 84.813 | 507 | 41 | 18 | 41 | 536 | 20400040 | 20399559 | 2.42e-130 | 477 |
| Clupea harengus-NC 045166.1 | 94.470 | 217 | 12 | 0 | 320 | 536 | 13693286 | 13693502 | 1.54e-87 | 335 |
| Clupea harengus-NC 045166.1 | 95.556 | 180 | 8 | 0 | 357 | 536 | 1966413 | 1966234 | 1.22e-73 | 289 |
| Clupea harengus-NC 045166.1 | 95.556 | 180 | 8 | 0 | 357 | 536 | 12407200 | 12407379 | 1.22e-73 | 289 |
| Clupea harengus-NC 045166.1 | 90.173 | 173 | 16 | 1 | 43 | 214 | 20287146 | 20286974 | 3.48e-54 | 224 |
| Clupea harengus-NC 045166.1 | 89.286 | 168 | 16 | 1 | 43 | 210 | 13693103 | 13693268 | 9.76e-50 | 209 |
| Clupea harengus-NC 045166.1 | 91.379 | 116 | 5 | 2 | 424 | 536 | 10276923 | 10276810 | 4.64e-33 | 154 |
| Clupea harengus-NC 045166.1 | 97.297 | 74 | 2 | 0 | 463 | 536 | 22145689 | 22145762 | 1.01e-24 | 126 |
| Coryphaenoides rupestris-PIJE02000572.1 | 90.891 | 494 | 26 | 9 | 43 | 536 | 235489 | 235015 | 0.0 | 645 |
| Clupea harengus-NC 045153.1 | 90.524 | 496 | 33 | 10 | 43 | 536 | 3105406 | 3105889 | 2.26e-180 | 643 |
| Clupea harengus-NC 045153.1 | 90.220 | 501 | 34 | 8 | 38 | 536 | 3129583 | 3130070 | 2.92e-179 | 640 |
| Clupea harengus-NC 045153.1 | 89.600 | 500 | 46 | 4 | 43 | 536 | 6216749 | 6217248 | 1.76e-176 | 630 |
| Clupea harengus-NC 045153.1 | 89.336 | 497 | 41 | 6 | 52 | 536 | 14268193 | 14267697 | 1.77e-171 | 614 |
| Clupea harengus-NC 045153.1 | 88.933 | 506 | 41 | 6 | 43 | 536 | 13955304 | 13955806 | 2.29e-170 | 610 |
| Clupea harengus-NC 045153.1 | 88.934 | 497 | 43 | 6 | 52 | 536 | 15394945 | 15394449 | 3.83e-168 | 603 |
| Clupea harengus-NC 045153.1 | 88.934 | 497 | 43 | 7 | 52 | 536 | 31817410 | 31816914 | 3.83e-168 | 603 |
| Clupea harengus-NC 045153.1 | 88.492 | 504 | 46 | 6 | 45 | 536 | 6175526 | 6176029 | 4.96e-167 | 599 |
| Clupea harengus-NC 045153.1 | 88.363 | 507 | 46 | 7 | 43 | 536 | 27183138 | 27183644 | 1.78e-166 | 597 |
| Clupea harengus-NC 045153.1 | 88.400 | 500 | 43 | 7 | 52 | 536 | 20557595 | 20557096 | 1.07e-163 | 588 |
| Clupea harengus-NC 045153.1 | 87.891 | 512 | 44 | 11 | 43 | 536 | 31198756 | 31199267 | 3.86e-163 | 586 |
| Clupea harengus-NC 045153.1 | 87.623 | 509 | 48 | 7 | 43 | 536 | 15519059 | 15519567 | 2.32e-160 | 577 |
| Clupea harengus-NC 045153.1 | 88.306 | 496 | 32 | 14 | 43 | 536 | 3184709 | 3185180 | 1.08e-158 | 571 |
| Clupea harengus-NC 045153.1 | 83.806 | 494 | 70 | 9 | 52 | 536 | 15176162 | 15175670 | 2.44e-125 | 460 |
| Clupea harengus-NC 045153.1 | 90.087 | 343 | 29 | 4 | 196 | 536 | 2994997 | 2994658 | 3.18e-119 | 440 |
| Clupea harengus-NC 045153.1 | 88.732 | 355 | 30 | 7 | 43 | 394 | 10664517 | 10664170 | 8.90e-115 | 425 |
| Clupea harengus-NC 045153.1 | 87.966 | 349 | 29 | 6 | 43 | 379 | 30037968 | 30037621 | 5.39e-107 | 399 |
| Clupea harengus-NC 045153.1 | 87.719 | 342 | 28 | 7 | 43 | 382 | 3137937 | 3138266 | 4.20e-103 | 387 |
| Clupea harengus-NC 045153.1 | 85.165 | 364 | 26 | 18 | 38 | 386 | 6112964 | 6113314 | 1.98e-91 | 348 |
| Clupea harengus-NC 045153.1 | 84.795 | 342 | 37 | 6 | 198 | 536 | 30037621 | 30037292 | 7.18e-86 | 329 |
| Clupea harengus-NC 045153.1 | 95.570 | 158 | 7 | 0 | 379 | 536 | 3140154 | 3140311 | 4.44e-63 | 254 |
| Clupea harengus-NC 045153.1 | 91.954 | 174 | 9 | 2 | 43 | 216 | 4165233 | 4165401 | 1.24e-58 | 239 |
| Clupea harengus-NC 045153.1 | 92.661 | 109 | 7 | 1 | 43 | 150 | 6112208 | 6112316 | 1.29e-33 | 156 |
| Clupea harengus-NC 045171.1 | 89.738 | 497 | 48 | 2 | 43 | 536 | 26359413 | 26358917 | 4.89e-177 | 632 |
| Clupea harengus-NC 045171.1 | 89.676 | 494 | 42 | 5 | 52 | 536 | 19142169 | 19141676 | 1.06e-173 | 621 |
| Clupea harengus-NC 045171.1 | 89.336 | 497 | 41 | 7 | 52 | 536 | 14046901 | 14046405 | 1.77e-171 | 614 |
| Clupea harengus-NC 045171.1 | 89.388 | 490 | 42 | 5 | 49 | 535 | 10689682 | 10690164 | 8.23e-170 | 608 |
| Clupea harengus-NC 045171.1 | 88.016 | 509 | 46 | 9 | 43 | 536 | 19943577 | 19944085 | 1.07e-163 | 588 |
| Clupea harengus-NC 045171.1 | 88.016 | 509 | 46 | 7 | 43 | 536 | 20042709 | 20043217 | 1.07e-163 | 588 |
| Clupea harengus-NC 045171.1 | 87.873 | 503 | 43 | 11 | 52 | 536 | 10189778 | 10189276 | 8.35e-160 | 575 |
| Clupea harengus-NC 045171.1 | 88.795 | 473 | 47 | 4 | 48 | 514 | 19945386 | 19944914 | 8.35e-160 | 575 |
| Clupea harengus-NC 045171.1 | 87.426 | 509 | 40 | 19 | 52 | 536 | 14034945 | 14034437 | 1.81e-156 | 564 |
| Clupea harengus-NC 045171.1 | 87.160 | 514 | 45 | 13 | 43 | 536 | 21894533 | 21895045 | 1.81e-156 | 564 |
| Clupea harengus-NC 045171.1 | 87.352 | 506 | 43 | 13 | 52 | 536 | 13114771 | 13114266 | 2.34e-155 | 560 |
| Clupea harengus-NC 045171.1 | 87.200 | 500 | 49 | 5 | 52 | 536 | 14047630 | 14047131 | 1.09e-153 | 555 |
| Clupea harengus-NC 045171.1 | 90.164 | 427 | 39 | 2 | 113 | 536 | 19945760 | 19946186 | 3.91e-153 | 553 |
| Clupea harengus-NC 045171.1 | 86.260 | 524 | 42 | 21 | 43 | 536 | 13389051 | 13389574 | 8.47e-150 | 542 |
| Clupea harengus-NC 045171.1 | 83.539 | 486 | 57 | 13 | 52 | 536 | 19044559 | 19044096 | 5.32e-117 | 433 |
| Clupea harengus-NC 045171.1 | 83.539 | 486 | 57 | 13 | 52 | 536 | 19069043 | 19068580 | 5.32e-117 | 433 |
| Clupea harengus-NC 045171.1 | 88.535 | 157 | 13 | 3 | 44 | 199 | 25839391 | 25839239 | 1.64e-42 | 185 |
| Clupea harengus-NC 045171.1 | 87.248 | 149 | 18 | 1 | 43 | 190 | 775581 | 775729 | 1.66e-37 | 169 |
| Clupea harengus-NC 045171.1 | 93.814 | 97 | 6 | 0 | 43 | 139 | 7209532 | 7209436 | 7.76e-31 | 147 |
| Clupea harengus-NC 045171.1 | 95.000 | 80 | 4 | 0 | 189 | 268 | 7209439 | 7209360 | 1.01e-24 | 126 |
| Clupea harengus-NW 024879570.1 | 90.020 | 491 | 43 | 4 | 52 | 536 | 46372 | 45882 | 1.76e-176 | 630 |
| Clupea harengus-NC 045169.1 | 89.600 | 500 | 46 | 4 | 43 | 536 | 13224562 | 13224063 | 1.76e-176 | 630 |
| Clupea harengus-NC 045169.1 | 89.130 | 506 | 42 | 7 | 43 | 536 | 7149321 | 7149825 | 1.37e-172 | 617 |
| Clupea harengus-NC 045169.1 | 88.933 | 506 | 44 | 6 | 43 | 536 | 8050426 | 8050931 | 1.77e-171 | 614 |
| Clupea harengus-NC 045169.1 | 88.735 | 506 | 45 | 6 | 43 | 536 | 7110577 | 7111082 | 8.23e-170 | 608 |
| Clupea harengus-NC 045169.1 | 89.135 | 497 | 42 | 6 | 52 | 536 | 25305226 | 25304730 | 8.23e-170 | 608 |
| Clupea harengus-NC 045169.1 | 88.800 | 500 | 41 | 7 | 52 | 536 | 24962965 | 24962466 | 4.96e-167 | 599 |
| Clupea harengus-NC 045169.1 | 88.212 | 509 | 45 | 7 | 43 | 536 | 23889247 | 23889755 | 2.31e-165 | 593 |
| Clupea harengus-NC 045169.1 | 88.531 | 497 | 45 | 6 | 52 | 536 | 13502747 | 13502251 | 8.29e-165 | 592 |
| Clupea harengus-NC 045169.1 | 87.891 | 512 | 44 | 10 | 43 | 536 | 5690628 | 5691139 | 3.86e-163 | 586 |
| Clupea harengus-NC 045169.1 | 88.200 | 500 | 44 | 7 | 52 | 536 | 24746625 | 24746126 | 4.99e-162 | 582 |
| Clupea harengus-NC 045169.1 | 88.048 | 502 | 42 | 11 | 52 | 536 | 13226450 | 13225950 | 6.46e-161 | 579 |
| Clupea harengus-NC 045169.1 | 87.623 | 509 | 48 | 10 | 43 | 536 | 8368401 | 8368909 | 2.32e-160 | 577 |
| Clupea harengus-NC 045169.1 | 87.259 | 518 | 42 | 19 | 43 | 536 | 25459612 | 25459095 | 3.89e-158 | 569 |
| Clupea harengus-NC 045169.1 | 86.719 | 512 | 50 | 11 | 43 | 536 | 13222037 | 13222548 | 3.91e-153 | 553 |
| Clupea harengus-NC 045169.1 | 86.260 | 524 | 42 | 22 | 43 | 536 | 26505644 | 26506167 | 8.47e-150 | 542 |
| Clupea harengus-NC 045169.1 | 87.273 | 275 | 31 | 4 | 43 | 314 | 23712780 | 23713053 | 2.60e-80 | 311 |

### N5.HOG0008885 Blastn of conserved regions against all genome assemblies

#### Region A (5/10)

| sseqid | pident | length | mismatch | gapopen | qstart | qend | sstart | send | evalue | bitscore |
|---|---|---|---|---|---|---|---|---|---|---|
| Clupea_harengus-NC_045169.1 | 86.909 | 275 | 32 | 4 | 43 | 314 | 23888271 | 23888544 | 1.21e-78 | 305 |
| Clupea_harengus-NC_045168.1 | 89.600 | 500 | 46 | 3 | 43 | 536 | 26369104 | 26369603 | 1.76e-176 | 630 |
| Clupea_harengus-NC_045168.1 | 88.735 | 506 | 45 | 6 | 43 | 536 | 17305372 | 17305877 | 8.23e-170 | 608 |
| Clupea_harengus-NC_045168.1 | 88.340 | 506 | 47 | 6 | 43 | 536 | 14346382 | 14346887 | 1.78e-166 | 597 |
| Clupea_harengus-NC_045168.1 | 88.212 | 509 | 45 | 7 | 43 | 536 | 13099934 | 13100442 | 2.31e-165 | 593 |
| Clupea_harengus-NC_045168.1 | 88.166 | 507 | 47 | 7 | 43 | 536 | 3104252 | 3103746 | 8.29e-165 | 592 |
| Clupea_harengus-NC_045168.1 | 88.039 | 510 | 45 | 9 | 43 | 536 | 24389 | 24898 | 2.98e-164 | 590 |
| Clupea_harengus-NC_045168.1 | 87.549 | 514 | 44 | 12 | 43 | 536 | 12456202 | 12455689 | 2.32e-160 | 577 |
| Clupea_harengus-NC_045168.1 | 86.260 | 524 | 41 | 17 | 43 | 536 | 3105366 | 3105888 | 3.05e-149 | 540 |
| Clupea_harengus-NC_045168.1 | 87.368 | 285 | 23 | 9 | 113 | 394 | 19851491 | 19851217 | 2.01e-81 | 315 |
| Clupea_harengus-NC_045156.1 | 89.738 | 497 | 45 | 5 | 43 | 536 | 29147365 | 29146872 | 1.76e-176 | 630 |
| Clupea_harengus-NC_045156.1 | 89.817 | 491 | 44 | 4 | 52 | 536 | 8606537 | 8606047 | 8.18e-175 | 625 |
| Clupea_harengus-NC_045156.1 | 89.066 | 503 | 45 | 6 | 43 | 536 | 21268696 | 21269197 | 4.92e-172 | 616 |
| Clupea_harengus-NC_045156.1 | 88.933 | 506 | 44 | 6 | 43 | 536 | 10385127 | 10385632 | 1.77e-171 | 614 |
| Clupea_harengus-NC_045156.1 | 88.735 | 506 | 45 | 6 | 43 | 536 | 20636205 | 20636710 | 8.23e-170 | 608 |
| Clupea_harengus-NC_045156.1 | 88.934 | 497 | 43 | 6 | 52 | 536 | 30663879 | 30663383 | 3.83e-168 | 603 |
| Clupea_harengus-NC_045156.1 | 87.500 | 512 | 46 | 11 | 43 | 536 | 5801661 | 5802172 | 8.35e-160 | 575 |
| Clupea_harengus-NC_045156.1 | 87.747 | 506 | 41 | 12 | 52 | 536 | 7928596 | 7928091 | 1.08e-158 | 571 |
| Clupea_harengus-NC_045156.1 | 87.800 | 500 | 46 | 7 | 52 | 536 | 10373163 | 10372664 | 1.08e-158 | 571 |
| Clupea_harengus-NC_045156.1 | 87.747 | 506 | 41 | 12 | 52 | 536 | 16625321 | 16624816 | 1.08e-158 | 571 |
| Clupea_harengus-NC_045156.1 | 87.549 | 506 | 42 | 12 | 52 | 536 | 26414743 | 26414238 | 5.03e-157 | 566 |
| Clupea_harengus-NC_045156.1 | 86.796 | 515 | 47 | 12 | 43 | 536 | 14694769 | 14695283 | 1.09e-153 | 555 |
| Clupea_harengus-NC_045156.1 | 86.680 | 518 | 45 | 19 | 43 | 536 | 13019067 | 13019584 | 3.91e-153 | 553 |
| Clupea_harengus-NC_045156.1 | 86.573 | 499 | 46 | 10 | 43 | 529 | 10367828 | 10367339 | 1.83e-146 | 531 |
| Clupea_harengus-NC_045156.1 | 86.166 | 506 | 46 | 9 | 43 | 536 | 14693917 | 14694410 | 8.53e-145 | 525 |
| Clupea_harengus-NC_045156.1 | 86.058 | 416 | 40 | 8 | 43 | 440 | 7921397 | 7921812 | 1.91e-116 | 431 |
| Clupea_harengus-NC_045156.1 | 93.088 | 217 | 13 | 2 | 320 | 536 | 25694758 | 25694544 | 5.59e-82 | 316 |
| Clupea_harengus-NC_045156.1 | 79.482 | 502 | 60 | 20 | 38 | 536 | 29130948 | 29131409 | 5.59e-82 | 316 |
| Clupea_harengus-NC_045156.1 | 92.929 | 198 | 12 | 2 | 43 | 239 | 25694952 | 25694756 | 4.38e-73 | 287 |
| Clupea_harengus-NC_045156.1 | 88.312 | 231 | 12 | 3 | 309 | 536 | 7923134 | 7922916 | 7.38e-66 | 263 |
| Clupea_harengus-NC_045156.1 | 89.189 | 148 | 16 | 0 | 154 | 301 | 21267769 | 21267916 | 1.64e-42 | 185 |
| Clupea_harengus-NC_045156.1 | 89.256 | 121 | 11 | 1 | 43 | 161 | 21164127 | 21164247 | 6.00e-32 | 150 |
| Clupea_harengus-NC_045156.1 | 88.430 | 121 | 12 | 1 | 43 | 161 | 21267132 | 21267252 | 2.79e-30 | 145 |
| Clupea_harengus-NC_045172.1 | 89.817 | 491 | 44 | 4 | 52 | 536 | 11082556 | 11082066 | 8.18e-175 | 625 |
| Clupea_harengus-NC_045172.1 | 89.336 | 497 | 41 | 6 | 52 | 536 | 18204658 | 18204162 | 1.77e-171 | 614 |
| Clupea_harengus-NC_045172.1 | 88.627 | 510 | 42 | 8 | 43 | 536 | 16438903 | 16438394 | 2.96e-169 | 606 |
| Clupea_harengus-NC_045172.1 | 88.409 | 509 | 44 | 7 | 43 | 536 | 18202459 | 18202967 | 4.96e-167 | 599 |
| Clupea_harengus-NC_045172.1 | 88.531 | 497 | 44 | 7 | 52 | 536 | 9053948 | 9053453 | 2.98e-164 | 590 |
| Clupea_harengus-NC_045172.1 | 88.016 | 509 | 46 | 7 | 43 | 536 | 19893992 | 19894500 | 1.07e-163 | 588 |
| Clupea_harengus-NC_045172.1 | 87.623 | 509 | 43 | 12 | 48 | 536 | 19923410 | 19922902 | 3.00e-159 | 573 |
| Clupea_harengus-NC_045172.1 | 86.640 | 509 | 41 | 14 | 48 | 536 | 19892972 | 19893473 | 1.10e-148 | 538 |
| Clupea_harengus-NC_045152.1 | 89.400 | 500 | 47 | 4 | 43 | 536 | 32362636 | 32363135 | 8.18e-175 | 625 |
| Clupea_harengus-NC_045152.1 | 89.655 | 493 | 43 | 6 | 52 | 536 | 20260213 | 20259721 | 1.06e-173 | 621 |
| Clupea_harengus-NC_045152.1 | 89.613 | 491 | 45 | 4 | 52 | 536 | 3493191 | 3492701 | 3.80e-173 | 619 |
| Clupea_harengus-NC_045152.1 | 89.613 | 491 | 45 | 4 | 52 | 536 | 5932303 | 5931813 | 3.80e-173 | 619 |
| Clupea_harengus-NC_045152.1 | 89.087 | 504 | 45 | 6 | 43 | 536 | 12572613 | 12573116 | 1.37e-172 | 617 |
| Clupea_harengus-NC_045152.1 | 89.271 | 494 | 44 | 5 | 52 | 536 | 13652362 | 13651869 | 2.29e-170 | 610 |
| Clupea_harengus-NC_045152.1 | 89.135 | 497 | 42 | 6 | 52 | 536 | 31821615 | 31821119 | 8.23e-170 | 608 |
| Clupea_harengus-NC_045152.1 | 88.934 | 497 | 42 | 10 | 52 | 536 | 17157788 | 17157293 | 1.38e-167 | 601 |
| Clupea_harengus-NC_045152.1 | 88.386 | 508 | 45 | 8 | 43 | 536 | 17617852 | 17618359 | 4.96e-167 | 599 |
| Clupea_harengus-NC_045152.1 | 88.340 | 506 | 47 | 6 | 43 | 536 | 13647402 | 13647907 | 1.78e-166 | 597 |
| Clupea_harengus-NC_045152.1 | 88.732 | 497 | 44 | 6 | 52 | 536 | 20180869 | 20180373 | 1.78e-166 | 597 |
| Clupea_harengus-NC_045152.1 | 88.212 | 509 | 45 | 9 | 43 | 536 | 20174630 | 20175138 | 2.31e-165 | 593 |
| Clupea_harengus-NC_045152.1 | 88.400 | 500 | 43 | 7 | 52 | 536 | 5943009 | 5942510 | 1.07e-163 | 588 |
| Clupea_harengus-NC_045152.1 | 87.945 | 506 | 48 | 7 | 43 | 536 | 16941312 | 16940808 | 1.39e-162 | 584 |
| Clupea_harengus-NC_045152.1 | 87.695 | 512 | 45 | 11 | 43 | 536 | 10184629 | 10185140 | 1.79e-161 | 580 |
| Clupea_harengus-NC_045152.1 | 88.081 | 495 | 49 | 5 | 43 | 536 | 5937114 | 5937599 | 6.46e-161 | 579 |
| Clupea_harengus-NC_045152.1 | 87.623 | 509 | 48 | 4 | 43 | 536 | 22362326 | 22362834 | 2.32e-160 | 577 |
| Clupea_harengus-NC_045152.1 | 87.305 | 512 | 47 | 11 | 43 | 536 | 17021419 | 17021930 | 3.89e-158 | 569 |
| Clupea_harengus-NC_045152.1 | 86.492 | 533 | 43 | 13 | 32 | 536 | 28209603 | 28209072 | 8.41e-155 | 558 |
| Clupea_harengus-NC_045152.1 | 87.273 | 495 | 46 | 9 | 48 | 536 | 14861790 | 14861307 | 5.06e-152 | 549 |
| Clupea_harengus-NC_045152.1 | 84.990 | 513 | 47 | 17 | 43 | 536 | 5938348 | 5938849 | 2.41e-135 | 494 |
| Clupea_harengus-NC_045152.1 | 83.752 | 517 | 47 | 16 | 43 | 536 | 17149670 | 17150172 | 1.14e-123 | 455 |
| Clupea_harengus-NC_045152.1 | 82.528 | 538 | 50 | 21 | 43 | 536 | 13475850 | 13476387 | 5.32e-117 | 433 |
| Clupea_harengus-NC_045152.1 | 82.874 | 508 | 41 | 19 | 52 | 536 | 17159157 | 17158673 | 1.93e-111 | 414 |
| Clupea_harengus-NC_045152.1 | 82.716 | 486 | 55 | 17 | 52 | 536 | 21689375 | 21688918 | 1.16e-108 | 405 |
| Coryphaenoides_rupestris-PIJE02000106.1 | 89.237 | 511 | 38 | 7 | 43 | 536 | 130825 | 130315 | 2.94e-174 | 623 |
| Coryphaenoides_rupestris-PIJE02000106.1 | 87.263 | 369 | 30 | 7 | 43 | 394 | 131000 | 131368 | 1.16e-108 | 405 |
| Coryphaenoides_rupestris-PIJE02000106.1 | 85.836 | 353 | 34 | 8 | 43 | 394 | 116726 | 116389 | 2.55e-95 | 361 |
| Clupea_harengus-NC_045177.1 | 89.264 | 503 | 45 | 5 | 43 | 536 | 12262981 | 12263483 | 1.06e-173 | 621 |
| Clupea_harengus-NC_045177.1 | 86.811 | 508 | 53 | 5 | 43 | 536 | 1292421 | 1292928 | 1.09e-153 | 555 |
| Clupea_harengus-NC_045177.1 | 87.734 | 481 | 44 | 9 | 43 | 508 | 858084 | 858564 | 1.82e-151 | 547 |
| Clupea_harengus-NC_045177.1 | 86.497 | 511 | 52 | 7 | 43 | 536 | 1288731 | 1289241 | 6.55e-151 | 545 |
| Clupea_harengus-NC_045177.1 | 89.254 | 335 | 30 | 4 | 63 | 394 | 11901245 | 11900914 | 1.93e-111 | 414 |
| Clupea_harengus-NC_045177.1 | 97.015 | 67 | 2 | 0 | 470 | 536 | 11900902 | 11900836 | 7.87e-21 | 113 |
| Clupea_harengus-NC_045165.1 | 90.566 | 477 | 34 | 7 | 61 | 536 | 19471403 | 19470937 | 1.06e-173 | 621 |
| Clupea_harengus-NC_045165.1 | 88.716 | 514 | 42 | 8 | 38 | 536 | 778775 | 779287 | 1.77e-171 | 614 |
| Clupea_harengus-NC_045165.1 | 88.800 | 500 | 41 | 8 | 52 | 536 | 11796188 | 11795689 | 4.96e-167 | 599 |
| Clupea_harengus-NC_045165.1 | 88.469 | 503 | 49 | 5 | 43 | 536 | 28490247 | 28490749 | 4.96e-167 | 599 |
| Clupea_harengus-NC_045165.1 | 88.212 | 509 | 45 | 7 | 43 | 536 | 24428012 | 24428520 | 2.31e-165 | 593 |
| Clupea_harengus-NC_045165.1 | 88.571 | 280 | 29 | 2 | 43 | 319 | 1544306 | 1544585 | 4.29e-88 | 337 |
| Clupea_harengus-NC_045165.1 | 96.111 | 180 | 7 | 0 | 357 | 536 | 11794641 | 11794462 | 2.62e-75 | 294 |
| Clupea_harengus-NC_045170.1 | 89.613 | 491 | 45 | 4 | 52 | 536 | 20447853 | 20447363 | 3.80e-173 | 619 |
| Clupea_harengus-NC_045170.1 | 89.474 | 494 | 43 | 5 | 52 | 536 | 2007083 | 2006590 | 4.92e-172 | 616 |
| Clupea_harengus-NC_045170.1 | 89.069 | 494 | 45 | 5 | 52 | 536 | 18177048 | 18176555 | 1.07e-168 | 604 |
| Clupea_harengus-NC_045170.1 | 88.212 | 509 | 45 | 7 | 43 | 536 | 14502401 | 14502909 | 2.31e-165 | 593 |
| Clupea_harengus-NC_045170.1 | 88.423 | 501 | 42 | 8 | 52 | 536 | 20448719 | 20448219 | 2.98e-164 | 590 |
| Clupea_harengus-NC_045170.1 | 87.819 | 509 | 47 | 7 | 43 | 536 | 649524 | 650032 | 4.99e-162 | 582 |
| Clupea_harengus-NC_045170.1 | 87.873 | 503 | 43 | 11 | 52 | 536 | 18176187 | 18175685 | 8.35e-160 | 575 |
| Clupea_harengus-NC_045170.1 | 86.561 | 506 | 46 | 16 | 43 | 536 | 26417233 | 26417728 | 1.10e-148 | 538 |
| Clupea_harengus-NC_045170.1 | 82.828 | 495 | 59 | 15 | 43 | 536 | 23296920 | 23297389 | 4.14e-113 | 420 |
| Clupea_harengus-NC_045170.1 | 93.939 | 231 | 13 | 1 | 306 | 536 | 2155292 | 2155063 | 1.98e-91 | 348 |
| Clupea_harengus-NC_045170.1 | 93.939 | 231 | 13 | 1 | 306 | 536 | 2178357 | 2178586 | 1.98e-91 | 348 |
| Clupea_harengus-NC_045159.1 | 89.613 | 491 | 45 | 4 | 52 | 536 | 20277754 | 20277264 | 3.80e-173 | 619 |
| Clupea_harengus-NC_045159.1 | 89.474 | 494 | 43 | 5 | 52 | 536 | 9122778 | 9122285 | 4.92e-172 | 616 |
| Clupea_harengus-NC_045159.1 | 89.336 | 497 | 41 | 7 | 52 | 536 | 2396766 | 2396270 | 1.77e-171 | 614 |
| Clupea_harengus-NC_045159.1 | 88.867 | 503 | 47 | 4 | 43 | 536 | 2031425 | 2031927 | 2.29e-170 | 610 |

### N5.HOG0008885 Blastn of conserved regions against all genome assemblies

#### Region A (6/10)

| sseqid | pident | length | mismatch | gapopen | qstart | qend | sstart | send | evalue | bitscore |
|---|---|---|---|---|---|---|---|---|---|---|
| Clupea_harengus-NC_045159.1 | 88.538 | 506 | 46 | 6 | 43 | 536 | 966081 | 966586 | 3.83e-168 | 603 |
| Clupea_harengus-NC_045159.1 | 88.538 | 506 | 46 | 6 | 43 | 536 | 5764983 | 5765488 | 3.83e-168 | 603 |
| Clupea_harengus-NC_045159.1 | 88.583 | 508 | 43 | 9 | 43 | 536 | 8755289 | 8755795 | 3.83e-168 | 603 |
| Clupea_harengus-NC_045159.1 | 88.600 | 500 | 47 | 8 | 43 | 536 | 2393366 | 2393861 | 4.96e-167 | 599 |
| Clupea_harengus-NC_045159.1 | 88.400 | 500 | 43 | 7 | 52 | 536 | 13898324 | 13897825 | 1.07e-163 | 588 |
| Clupea_harengus-NC_045159.1 | 87.767 | 515 | 42 | 12 | 43 | 536 | 12366228 | 12366742 | 4.99e-162 | 582 |
| Clupea_harengus-NC_045159.1 | 88.200 | 500 | 44 | 7 | 52 | 536 | 18896036 | 18895537 | 4.99e-162 | 582 |
| Clupea_harengus-NC_045159.1 | 87.726 | 497 | 43 | 9 | 52 | 536 | 1583548 | 1583058 | 1.81e-156 | 564 |
| Clupea_harengus-NC_045159.1 | 86.914 | 512 | 49 | 11 | 43 | 536 | 12368428 | 12368939 | 8.41e-155 | 558 |
| Clupea_harengus-NC_045159.1 | 87.352 | 506 | 40 | 15 | 52 | 536 | 27139682 | 27139180 | 8.41e-155 | 558 |
| Clupea_harengus-NC_045159.1 | 87.247 | 494 | 42 | 8 | 52 | 536 | 13881348 | 13880867 | 2.35e-150 | 544 |
| Clupea_harengus-NC_045159.1 | 85.798 | 514 | 51 | 9 | 41 | 536 | 15199018 | 15198509 | 8.53e-145 | 525 |
| Clupea_harengus-NC_045159.1 | 85.600 | 500 | 52 | 7 | 41 | 536 | 5318290 | 5317807 | 3.09e-139 | 507 |
| Clupea_harengus-NC_045159.1 | 85.352 | 512 | 44 | 18 | 43 | 536 | 21981996 | 21981498 | 1.44e-137 | 501 |
| Clupea_harengus-NC_045159.1 | 85.484 | 496 | 53 | 7 | 43 | 536 | 5329399 | 5329877 | 5.17e-137 | 499 |
| Clupea_harengus-NC_045159.1 | 90.451 | 377 | 31 | 3 | 165 | 536 | 5900159 | 5900535 | 8.65e-135 | 492 |
| Clupea_harengus-NC_045159.1 | 84.783 | 506 | 42 | 11 | 43 | 536 | 9133793 | 9134275 | 8.71e-130 | 475 |
| Clupea_harengus-NC_045159.1 | 88.385 | 353 | 30 | 6 | 41 | 393 | 9127310 | 9126969 | 1.93e-111 | 414 |
| Clupea_harengus-NC_045159.1 | 82.688 | 491 | 45 | 15 | 43 | 532 | 5269408 | 5268957 | 5.39e-107 | 399 |
| Clupea_harengus-NC_045159.1 | 95.455 | 88 | 4 | 0 | 449 | 536 | 9123063 | 9123150 | 3.61e-29 | 141 |
| Clupea_harengus-NC_045157.1 | 89.200 | 500 | 48 | 4 | 43 | 536 | 20216522 | 20217021 | 3.80e-173 | 619 |
| Clupea_harengus-NC_045157.1 | 88.933 | 506 | 44 | 6 | 43 | 536 | 18675161 | 18675666 | 1.77e-171 | 614 |
| Clupea_harengus-NC_045157.1 | 89.135 | 497 | 42 | 6 | 52 | 536 | 2889975 | 2889479 | 8.23e-170 | 608 |
| Clupea_harengus-NC_045157.1 | 88.538 | 506 | 46 | 6 | 43 | 536 | 2117708 | 2118213 | 3.83e-168 | 603 |
| Clupea_harengus-NC_045157.1 | 88.409 | 509 | 44 | 7 | 43 | 536 | 11606819 | 11607327 | 4.96e-167 | 599 |
| Clupea_harengus-NC_045157.1 | 88.600 | 500 | 42 | 7 | 52 | 536 | 11958814 | 11958315 | 2.31e-165 | 593 |
| Clupea_harengus-NC_045157.1 | 88.363 | 507 | 41 | 11 | 48 | 536 | 14628381 | 14627875 | 2.31e-165 | 593 |
| Clupea_harengus-NC_045157.1 | 88.016 | 509 | 46 | 8 | 43 | 536 | 8784102 | 8784610 | 1.07e-163 | 588 |
| Clupea_harengus-NC_045157.1 | 88.247 | 502 | 42 | 10 | 52 | 536 | 7548317 | 7547816 | 1.39e-162 | 584 |
| Clupea_harengus-NC_045157.1 | 87.873 | 503 | 43 | 11 | 52 | 536 | 21457478 | 21456976 | 8.35e-160 | 575 |
| Clupea_harengus-NC_045157.1 | 85.988 | 521 | 37 | 24 | 52 | 536 | 8786289 | 8785769 | 8.53e-145 | 525 |
| Clupea_harengus-NC_045157.1 | 84.879 | 496 | 27 | 13 | 45 | 536 | 4373319 | 4372868 | 3.16e-124 | 457 |
| Clupea_harengus-NC_045157.1 | 83.463 | 514 | 49 | 16 | 43 | 536 | 19539720 | 19540217 | 6.83e-121 | 446 |
| Clupea_harengus-NC_045157.1 | 85.676 | 377 | 39 | 9 | 43 | 404 | 13385537 | 13385913 | 5.43e-102 | 383 |
| Clupea_harengus-NC_045157.1 | 96.226 | 106 | 4 | 0 | 431 | 536 | 13385900 | 13386005 | 3.56e-39 | 174 |
| Clupea_harengus-NW_024880674.1 | 89.087 | 504 | 45 | 5 | 43 | 536 | 1208 | 1711 | 1.37e-172 | 617 |
| Clupea_harengus-NW_024879809.1 | 88.976 | 508 | 44 | 6 | 41 | 536 | 15625 | 15118 | 1.37e-172 | 617 |
| Clupea_harengus-NW_024879809.1 | 88.800 | 500 | 41 | 7 | 52 | 536 | 20832 | 20333 | 4.96e-167 | 599 |
| Clupea_harengus-NW_024879809.1 | 86.200 | 500 | 48 | 10 | 38 | 536 | 13563 | 14042 | 1.10e-143 | 521 |
| Clupea_harengus-NC_045161.1 | 89.066 | 503 | 46 | 5 | 43 | 536 | 14934477 | 14934979 | 4.92e-172 | 616 |
| Clupea_harengus-NC_045161.1 | 89.336 | 497 | 41 | 6 | 52 | 536 | 14931175 | 14930679 | 1.77e-171 | 614 |
| Clupea_harengus-NC_045161.1 | 89.000 | 500 | 40 | 7 | 52 | 536 | 17223265 | 17222766 | 1.07e-168 | 604 |
| Clupea_harengus-NC_045161.1 | 88.668 | 503 | 48 | 6 | 43 | 536 | 22464230 | 22464732 | 1.07e-168 | 604 |
| Clupea_harengus-NC_045161.1 | 88.600 | 500 | 42 | 7 | 52 | 536 | 25596803 | 25596304 | 2.31e-165 | 593 |
| Clupea_harengus-NC_045161.1 | 87.674 | 503 | 44 | 11 | 52 | 536 | 14954011 | 14953509 | 3.89e-158 | 569 |
| Clupea_harengus-NC_045161.1 | 87.860 | 486 | 43 | 9 | 52 | 536 | 20827768 | 20827298 | 3.02e-154 | 556 |
| Clupea_harengus-NC_045161.1 | 86.486 | 518 | 37 | 23 | 52 | 536 | 23357142 | 23356625 | 1.10e-148 | 538 |
| Clupea_harengus-NC_045161.1 | 86.561 | 506 | 44 | 9 | 43 | 536 | 17515556 | 17516049 | 3.94e-148 | 536 |
| Clupea_harengus-NC_045161.1 | 87.703 | 431 | 34 | 10 | 43 | 456 | 14945179 | 14945607 | 1.45e-132 | 484 |
| Clupea_harengus-NC_045161.1 | 94.258 | 209 | 9 | 3 | 329 | 536 | 11018034 | 11018240 | 5.59e-82 | 316 |
| Clupea_harengus-NC_045161.1 | 92.308 | 117 | 8 | 1 | 43 | 158 | 21327015 | 21326899 | 2.14e-36 | 165 |
| Clupea_harengus-NC_045161.1 | 94.382 | 89 | 5 | 0 | 448 | 536 | 14930455 | 14930367 | 4.67e-28 | 137 |
| Clupea_harengus-NW_024879599.1 | 88.933 | 506 | 44 | 6 | 43 | 536 | 41893 | 42398 | 1.77e-171 | 614 |
| Synaphobranchus_kaupii-CM056393.1 | 88.762 | 525 | 28 | 14 | 43 | 536 | 32110974 | 32110450 | 1.77e-171 | 614 |
| Synaphobranchus_kaupii-CM056393.1 | 89.274 | 317 | 28 | 4 | 38 | 350 | 6245601 | 6245915 | 9.03e-105 | 392 |
| Synaphobranchus_kaupii-CM056393.1 | 89.423 | 312 | 28 | 3 | 43 | 350 | 12839239 | 12839549 | 1.17e-103 | 388 |
| Synaphobranchus_kaupii-CM056393.1 | 90.333 | 300 | 23 | 4 | 43 | 340 | 22875105 | 22874810 | 1.17e-103 | 388 |
| Synaphobranchus_kaupii-CM056393.1 | 89.423 | 312 | 28 | 3 | 43 | 350 | 28457833 | 28457523 | 1.17e-103 | 388 |
| Synaphobranchus_kaupii-CM056393.1 | 89.103 | 312 | 29 | 3 | 43 | 350 | 32104846 | 32104536 | 5.43e-102 | 383 |
| Synaphobranchus_kaupii-CM056393.1 | 89.369 | 301 | 28 | 2 | 43 | 339 | 6247793 | 6248093 | 9.09e-100 | 375 |
| Synaphobranchus_kaupii-CM056393.1 | 88.013 | 317 | 32 | 4 | 38 | 350 | 7099356 | 7099042 | 4.23e-98 | 370 |
| Coryphaenoides_rupestris-PIJE02018372.1 | 89.062 | 512 | 33 | 12 | 43 | 536 | 2128 | 1622 | 1.77e-171 | 614 |
| Coryphaenoides_rupestris-PIJE02000809.1 | 89.286 | 504 | 33 | 12 | 43 | 536 | 199233 | 198741 | 6.36e-171 | 612 |
| Coryphaenoides_rupestris-PIJE02000809.1 | 87.354 | 514 | 39 | 13 | 43 | 536 | 184889 | 184382 | 5.03e-157 | 566 |
| Clupea_harengus-NW_024879816.1 | 88.735 | 506 | 45 | 6 | 43 | 536 | 12122 | 12627 | 8.23e-170 | 608 |
| Clupea_harengus-NC_045174.1 | 88.735 | 506 | 45 | 6 | 43 | 536 | 12636750 | 12637255 | 8.23e-170 | 608 |
| Clupea_harengus-NC_045174.1 | 88.690 | 504 | 47 | 6 | 43 | 537 | 12635537 | 12636039 | 2.96e-169 | 606 |
| Clupea_harengus-NC_045174.1 | 88.538 | 506 | 46 | 7 | 43 | 536 | 12424402 | 12424907 | 3.83e-168 | 603 |
| Clupea_harengus-NC_045174.1 | 88.538 | 506 | 46 | 6 | 43 | 536 | 14722229 | 14722734 | 3.83e-168 | 603 |
| Clupea_harengus-NC_045174.1 | 88.431 | 510 | 43 | 12 | 43 | 536 | 19544012 | 19544521 | 1.38e-167 | 601 |
| Clupea_harengus-NC_045174.1 | 88.800 | 500 | 41 | 7 | 52 | 536 | 12630707 | 12630208 | 4.96e-167 | 599 |
| Clupea_harengus-NC_045174.1 | 88.485 | 495 | 46 | 6 | 43 | 536 | 10615432 | 10615916 | 1.07e-163 | 588 |
| Clupea_harengus-NC_045174.1 | 88.400 | 500 | 43 | 7 | 52 | 536 | 12628672 | 12628173 | 1.07e-163 | 588 |
| Clupea_harengus-NC_045174.1 | 88.000 | 500 | 45 | 7 | 52 | 536 | 3465126 | 3464627 | 2.32e-160 | 577 |
| Clupea_harengus-NC_045174.1 | 85.907 | 518 | 49 | 12 | 43 | 536 | 9557432 | 9557949 | 1.83e-146 | 531 |
| Clupea_harengus-NC_045174.1 | 86.807 | 379 | 38 | 6 | 43 | 409 | 17668389 | 17668767 | 6.93e-111 | 412 |
| Clupea_harengus-NC_045174.1 | 91.209 | 273 | 22 | 1 | 266 | 536 | 272343 | 272071 | 4.23e-98 | 370 |
| Clupea_harengus-NC_045174.1 | 91.078 | 269 | 15 | 7 | 272 | 536 | 25138300 | 25138563 | 1.18e-93 | 355 |
| Clupea_harengus-NC_045174.1 | 94.248 | 226 | 13 | 0 | 43 | 268 | 21750920 | 21750695 | 7.13e-91 | 346 |
| Clupea_harengus-NC_045174.1 | 82.778 | 360 | 42 | 8 | 43 | 398 | 25109430 | 25109773 | 4.35e-78 | 303 |
| Clupea_harengus-NC_045174.1 | 91.284 | 218 | 18 | 1 | 43 | 259 | 276625 | 276408 | 7.28e-76 | 296 |
| Clupea_harengus-NC_045174.1 | 82.405 | 341 | 28 | 17 | 55 | 394 | 284555 | 284864 | 1.59e-67 | 268 |
| Clupea_harengus-NC_045174.1 | 88.750 | 160 | 16 | 1 | 43 | 202 | 25138132 | 25138289 | 2.73e-45 | 195 |
| Clupea_harengus-NC_045174.1 | 95.294 | 85 | 4 | 0 | 452 | 536 | 17668768 | 17668852 | 1.68e-27 | 135 |
| Clupea_harengus-NC_045174.1 | 95.349 | 86 | 3 | 1 | 43 | 127 | 21565217 | 21565302 | 1.68e-27 | 135 |
| Clupea_harengus-NC_045174.1 | 95.349 | 86 | 3 | 1 | 43 | 127 | 21587254 | 21587339 | 1.68e-27 | 135 |
| Clupea_harengus-NC_045174.1 | 94.186 | 86 | 4 | 1 | 43 | 127 | 21598879 | 21598964 | 7.81e-26 | 130 |
| Clupea_harengus-NC_045155.1 | 88.735 | 506 | 45 | 7 | 43 | 536 | 11248416 | 11247911 | 8.23e-170 | 608 |
| Clupea_harengus-NC_045155.1 | 88.605 | 509 | 43 | 7 | 43 | 536 | 20880358 | 20880866 | 1.07e-168 | 604 |
| Clupea_harengus-NC_045155.1 | 89.002 | 491 | 48 | 5 | 52 | 536 | 23610606 | 23610116 | 3.83e-168 | 603 |
| Clupea_harengus-NC_045155.1 | 88.956 | 498 | 41 | 8 | 52 | 536 | 23950978 | 23950482 | 3.83e-168 | 603 |
| Clupea_harengus-NC_045155.1 | 88.554 | 498 | 44 | 7 | 52 | 536 | 7742132 | 7741635 | 8.29e-165 | 592 |
| Clupea_harengus-NC_045155.1 | 88.142 | 506 | 48 | 6 | 43 | 536 | 12027160 | 12027665 | 8.29e-165 | 592 |
| Clupea_harengus-NC_045155.1 | 88.086 | 512 | 43 | 11 | 43 | 536 | 31466335 | 31466846 | 8.29e-165 | 592 |
| Clupea_harengus-NC_045155.1 | 88.400 | 500 | 43 | 8 | 52 | 536 | 16090126 | 16089627 | 1.07e-163 | 588 |
| Clupea_harengus-NC_045155.1 | 88.016 | 509 | 46 | 8 | 43 | 536 | 29748472 | 29748980 | 1.07e-163 | 588 |
| Clupea_harengus-NC_045155.1 | 88.200 | 500 | 44 | 7 | 52 | 536 | 29865864 | 29865365 | 4.99e-162 | 582 |
| Clupea_harengus-NC_045155.1 | 87.230 | 509 | 41 | 19 | 52 | 536 | 1428094 | 1427586 | 8.41e-155 | 558 |
| Clupea_harengus-NC_045155.1 | 87.746 | 457 | 35 | 10 | 43 | 499 | 11221839 | 11222274 | 1.85e-141 | 514 |
| Clupea_harengus-NC_045155.1 | 83.984 | 512 | 46 | 14 | 38 | 536 | 5900153 | 5900641 | 8.77e-125 | 459 |
| Clupea_harengus-NC_045155.1 | 86.970 | 330 | 30 | 7 | 43 | 361 | 31720923 | 31721250 | 9.16e-95 | 359 |
| Clupea_harengus-NC_045155.1 | 95.000 | 180 | 9 | 0 | 357 | 536 | 16630349 | 16630170 | 5.67e-72 | 283 |

### N5.HOG0008885 Blastn of conserved regions against all genome assemblies

#### Region A (7/10)

| sseqid | pident | length | mismatch | gapopen | qstart | qend | sstart | send | evalue | bitscore |
|---|---|---|---|---|---|---|---|---|---|---|
| Clupea_harengus-NC_045155.1 | 86.585 | 246 | 27 | 6 | 293 | 536 | 7822132 | 7821891 | 5.71e-67 | 267 |
| Clupea_harengus-NC_045155.1 | 93.706 | 143 | 8 | 1 | 52 | 193 | 4918593 | 4918451 | 7.54e-51 | 213 |
| Clupea_harengus-NW_024881132.1 | 88.668 | 503 | 48 | 5 | 43 | 536 | 181 | 683 | 1.07e-168 | 604 |
| Clupea_harengus-NW_024880687.1 | 89.069 | 494 | 45 | 5 | 52 | 536 | 6634 | 6141 | 1.07e-168 | 604 |
| Clupea_harengus-NW_024880687.1 | 86.117 | 497 | 48 | 8 | 41 | 536 | 3284 | 2808 | 5.13e-142 | 516 |
| Clupea_harengus-NW_024880600.1 | 89.069 | 494 | 45 | 5 | 52 | 536 | 9313 | 8820 | 1.07e-168 | 604 |
| Clupea_harengus-NW_024880517.1 | 89.069 | 494 | 45 | 5 | 52 | 536 | 19356 | 18863 | 1.07e-168 | 604 |
| Clupea_harengus-NW_024879697.1 | 89.069 | 494 | 45 | 5 | 52 | 536 | 16095 | 15602 | 1.07e-168 | 604 |
| Clupea_harengus-NW_024880578.1 | 88.538 | 506 | 46 | 7 | 43 | 536 | 611 | 106 | 3.83e-168 | 603 |
| Clupea_harengus-NW_024880578.1 | 88.934 | 497 | 43 | 6 | 52 | 536 | 2587 | 2091 | 3.83e-168 | 603 |
| Clupea_harengus-NW_024879662.1 | 88.538 | 506 | 46 | 7 | 43 | 536 | 62601 | 63106 | 3.83e-168 | 603 |
| Clupea_harengus-NW_024879662.1 | 88.732 | 497 | 44 | 7 | 52 | 536 | 65209 | 64713 | 1.78e-166 | 597 |
| Clupea_harengus-NW_024879662.1 | 88.732 | 497 | 44 | 6 | 52 | 536 | 75764 | 75268 | 1.78e-166 | 597 |
| Clupea_harengus-NW_024879642.1 | 88.538 | 506 | 46 | 6 | 43 | 536 | 77471 | 77976 | 3.83e-168 | 603 |
| Clupea_harengus-NW_024879642.1 | 88.340 | 506 | 47 | 6 | 43 | 536 | 76609 | 77114 | 1.78e-166 | 597 |
| Clupea_harengus-NW_024879642.1 | 95.146 | 103 | 5 | 0 | 434 | 536 | 76264 | 76162 | 7.70e-36 | 163 |
| Clupea_harengus-NW_024879615.1 | 88.538 | 506 | 46 | 6 | 43 | 536 | 59527 | 60032 | 3.83e-168 | 603 |
| Clupea_harengus-NW_024879615.1 | 88.538 | 506 | 46 | 6 | 43 | 536 | 61486 | 61991 | 3.83e-168 | 603 |
| Clupea_harengus-NW_024879615.1 | 88.340 | 506 | 47 | 6 | 43 | 536 | 57569 | 58074 | 1.78e-166 | 597 |
| Clupea_harengus-NW_024879615.1 | 88.577 | 499 | 45 | 6 | 43 | 529 | 63444 | 63942 | 6.41e-166 | 595 |
| Clupea_harengus-NW_024879615.1 | 90.238 | 420 | 40 | 1 | 43 | 462 | 67227 | 66809 | 1.82e-151 | 547 |
| Clupea_harengus-NW_024879595.1 | 88.538 | 506 | 46 | 6 | 43 | 536 | 9433 | 9938 | 3.83e-168 | 603 |
| Clupea_harengus-NW_024880834.1 | 88.409 | 509 | 44 | 9 | 43 | 536 | 2474 | 2982 | 4.96e-167 | 599 |
| Clupea_harengus-NW_024880663.1 | 88.409 | 509 | 44 | 7 | 43 | 536 | 6516 | 7024 | 4.96e-167 | 599 |
| Clupea_harengus-NW_024880506.1 | 88.800 | 500 | 41 | 7 | 52 | 536 | 10913 | 10414 | 4.96e-167 | 599 |
| Clupea_harengus-NW_024879565.1 | 88.031 | 518 | 46 | 10 | 34 | 536 | 110260 | 110776 | 4.96e-167 | 599 |
| Clupea_harengus-NC_045176.1 | 88.778 | 499 | 42 | 7 | 52 | 536 | 10599971 | 10599473 | 4.96e-167 | 599 |
| Clupea_harengus-NC_045176.1 | 88.531 | 497 | 49 | 4 | 43 | 536 | 7309864 | 7310355 | 6.41e-166 | 595 |
| Clupea_harengus-NC_045176.1 | 87.891 | 512 | 44 | 11 | 43 | 536 | 11458402 | 11458913 | 3.86e-163 | 586 |
| Clupea_harengus-NC_045176.1 | 87.819 | 509 | 47 | 8 | 43 | 536 | 10723284 | 10723792 | 4.99e-162 | 582 |
| Clupea_harengus-NC_045176.1 | 87.695 | 512 | 45 | 11 | 43 | 536 | 7705715 | 7706226 | 1.79e-161 | 580 |
| Clupea_harengus-NC_045176.1 | 85.913 | 504 | 34 | 8 | 38 | 536 | 506819 | 507290 | 4.00e-138 | 503 |
| Clupea_harengus-NC_045176.1 | 82.677 | 254 | 16 | 8 | 285 | 536 | 2063194 | 2063421 | 5.87e-47 | 200 |
| Clupea_harengus-NC_045176.1 | 96.970 | 99 | 3 | 0 | 438 | 536 | 2054867 | 2054965 | 5.96e-37 | 167 |
| Clupea_harengus-NC_045176.1 | 95.960 | 99 | 4 | 0 | 438 | 536 | 2069747 | 2069649 | 2.77e-35 | 161 |
| Clupea_harengus-NC_045176.1 | 93.684 | 95 | 5 | 1 | 443 | 536 | 2086340 | 2086246 | 3.61e-29 | 141 |
| Clupea_harengus-NC_045158.1 | 88.258 | 511 | 47 | 8 | 38 | 536 | 18861011 | 18861520 | 4.96e-167 | 599 |
| Clupea_harengus-NC_045158.1 | 88.212 | 509 | 45 | 7 | 43 | 536 | 9917273 | 9917781 | 2.31e-165 | 593 |
| Clupea_harengus-NC_045158.1 | 87.674 | 503 | 44 | 11 | 52 | 536 | 4065834 | 4065332 | 3.89e-158 | 569 |
| Clupea_harengus-NC_045158.1 | 87.426 | 509 | 40 | 19 | 52 | 536 | 18863247 | 18862739 | 1.81e-156 | 564 |
| Clupea_harengus-NC_045158.1 | 86.873 | 518 | 43 | 14 | 43 | 536 | 2932485 | 2933001 | 3.02e-154 | 556 |
| Clupea_harengus-NC_045158.1 | 87.381 | 420 | 40 | 7 | 43 | 450 | 10820409 | 10819991 | 4.05e-128 | 470 |
| Clupea_harengus-NC_045158.1 | 85.981 | 428 | 47 | 8 | 38 | 453 | 26015871 | 26016297 | 6.83e-121 | 446 |
| Clupea_harengus-NC_045158.1 | 90.000 | 270 | 19 | 6 | 272 | 536 | 10819602 | 10819336 | 9.22e-90 | 342 |
| Clupea_harengus-NC_045158.1 | 89.520 | 229 | 19 | 5 | 309 | 536 | 23206697 | 23206921 | 1.58e-72 | 285 |
| Clupea_harengus-NC_045158.1 | 93.789 | 161 | 10 | 0 | 376 | 536 | 1694315 | 1694155 | 9.62e-60 | 243 |
| Clupea_harengus-NC_045158.1 | 89.785 | 186 | 15 | 3 | 43 | 227 | 26001914 | 26002096 | 1.61e-57 | 235 |
| Clupea_harengus-NC_045158.1 | 85.057 | 174 | 11 | 7 | 97 | 268 | 23206265 | 23206425 | 7.70e-36 | 163 |
| Coryphaenoides_rupestris-PIJE02000183.1 | 91.383 | 441 | 32 | 4 | 43 | 477 | 46794 | 46354 | 4.96e-167 | 599 |
| Clupea_harengus-NW_024880879.1 | 88.732 | 497 | 44 | 6 | 52 | 536 | 4610 | 4114 | 1.78e-166 | 597 |
| Clupea_harengus-NW_024879608.1 | 88.340 | 506 | 47 | 7 | 43 | 536 | 139443 | 139948 | 1.78e-166 | 597 |
| Clupea_harengus-NW_024879608.1 | 88.212 | 509 | 45 | 7 | 43 | 536 | 134961 | 135469 | 2.31e-165 | 593 |
| Clupea_harengus-NW_024879608.1 | 88.401 | 319 | 23 | 6 | 232 | 536 | 136314 | 136632 | 1.18e-98 | 372 |
| Clupea_harengus-NW_024880046.1 | 88.212 | 509 | 45 | 7 | 43 | 536 | 4517 | 5025 | 2.31e-165 | 593 |
| Clupea_harengus-NW_024879629.1 | 88.600 | 500 | 42 | 7 | 52 | 536 | 97369 | 96870 | 2.31e-165 | 593 |
| Clupea_harengus-NW_024879601.1 | 88.189 | 508 | 46 | 8 | 43 | 536 | 33180 | 33687 | 2.31e-165 | 593 |
| Synaphobranchus_kaupii-CM056394.1 | 90.889 | 450 | 28 | 8 | 38 | 477 | 28241265 | 28241711 | 8.29e-165 | 592 |
| Synaphobranchus_kaupii-CM056394.1 | 89.423 | 312 | 28 | 3 | 43 | 350 | 20419466 | 20419776 | 1.17e-103 | 388 |
| Synaphobranchus_kaupii-CM056394.1 | 89.103 | 312 | 29 | 3 | 43 | 350 | 22547670 | 22547360 | 5.43e-102 | 383 |
| Synaphobranchus_kaupii-CM056394.1 | 88.782 | 312 | 30 | 3 | 43 | 350 | 22548332 | 22548022 | 2.53e-100 | 377 |
| Synaphobranchus_kaupii-CM056394.1 | 86.667 | 315 | 26 | 5 | 43 | 350 | 1783379 | 1783074 | 1.54e-87 | 335 |
| Coryphaenoides_rupestris-PIJE02000668.1 | 87.925 | 530 | 27 | 23 | 43 | 536 | 233078 | 233606 | 2.98e-164 | 590 |
| Clupea_harengus-NW_024880010.1 | 88.400 | 500 | 43 | 10 | 52 | 536 | 4125 | 3626 | 1.07e-163 | 588 |
| Clupea_harengus-NW_024879589.1 | 88.016 | 509 | 46 | 7 | 43 | 536 | 84724 | 85232 | 1.07e-163 | 588 |
| Clupea_harengus-NW_024879589.1 | 94.944 | 178 | 9 | 0 | 359 | 536 | 83998 | 83821 | 7.33e-71 | 279 |
| Clupea_harengus-NW_024879577.1 | 88.400 | 500 | 43 | 7 | 52 | 536 | 52618 | 52119 | 1.07e-163 | 588 |
| Coryphaenoides_rupestris-PIJE02002316.1 | 88.454 | 511 | 27 | 11 | 43 | 536 | 63665 | 64160 | 1.07e-163 | 588 |
| Coryphaenoides_rupestris-PIJE02001412.1 | 87.856 | 527 | 30 | 22 | 43 | 536 | 70874 | 71399 | 1.07e-163 | 588 |
| Clupea_harengus-NW_024880258.1 | 87.891 | 512 | 44 | 9 | 43 | 536 | 18148 | 18659 | 3.86e-163 | 586 |
| Clupea_harengus-NW_024879731.1 | 89.827 | 462 | 39 | 6 | 77 | 536 | 18963 | 19418 | 3.86e-163 | 586 |
| Coryphaenoides_rupestris-PIJE02001193.1 | 87.814 | 517 | 40 | 18 | 43 | 536 | 113197 | 112681 | 1.39e-162 | 584 |
| Coryphaenoides_rupestris-PIJE02000052.1 | 90.065 | 463 | 29 | 10 | 43 | 488 | 730580 | 730118 | 1.39e-162 | 584 |
| Clupea_harengus-NW_024880656.1 | 88.200 | 500 | 44 | 7 | 52 | 536 | 3342 | 2843 | 4.99e-162 | 582 |
| Clupea_harengus-NW_024879571.1 | 87.819 | 509 | 47 | 7 | 43 | 536 | 170827 | 171335 | 4.99e-162 | 582 |
| Clupea_harengus-NW_024879571.1 | 86.923 | 260 | 33 | 1 | 43 | 301 | 169599 | 169858 | 3.39e-74 | 291 |
| Clupea_harengus-NW_024879571.1 | 96.875 | 96 | 3 | 0 | 43 | 138 | 199584 | 199489 | 2.77e-35 | 161 |
| Synaphobranchus_kaupii-CM056381.1 | 90.466 | 451 | 29 | 9 | 38 | 477 | 70917845 | 70917398 | 4.99e-162 | 582 |
| Synaphobranchus_kaupii-CM056381.1 | 89.423 | 312 | 28 | 3 | 43 | 350 | 38484094 | 38483784 | 1.17e-103 | 388 |
| Synaphobranchus_kaupii-CM056381.1 | 89.423 | 312 | 28 | 3 | 43 | 350 | 80186319 | 80186629 | 1.17e-103 | 388 |
| Synaphobranchus_kaupii-CM056381.1 | 89.423 | 312 | 28 | 3 | 43 | 350 | 85287068 | 85286758 | 1.17e-103 | 388 |
| Synaphobranchus_kaupii-CM056381.1 | 89.103 | 312 | 29 | 3 | 43 | 350 | 59682860 | 59682550 | 5.43e-102 | 383 |
| Synaphobranchus_kaupii-CM056381.1 | 89.103 | 312 | 29 | 3 | 43 | 350 | 68842048 | 68841738 | 5.43e-102 | 383 |
| Synaphobranchus_kaupii-CM056381.1 | 89.369 | 301 | 28 | 2 | 43 | 339 | 35619458 | 35619158 | 9.09e-100 | 375 |
| Coryphaenoides_rupestris-PIJE02000459.1 | 88.063 | 511 | 35 | 14 | 43 | 534 | 34334 | 34837 | 4.99e-162 | 582 |
| Coryphaenoides_rupestris-PIJE02000459.1 | 87.755 | 392 | 28 | 9 | 43 | 414 | 29705 | 30096 | 3.18e-119 | 440 |
| Coryphaenoides_rupestris-PIJE02000007.1 | 88.431 | 510 | 23 | 29 | 43 | 536 | 1747586 | 1747097 | 4.99e-162 | 582 |
| Clupea_harengus-NW_024880019.1 | 87.695 | 512 | 45 | 11 | 43 | 536 | 32558 | 33069 | 1.79e-161 | 580 |
| Clupea_harengus-NW_024880618.1 | 87.897 | 504 | 46 | 10 | 43 | 536 | 15534 | 16032 | 6.46e-161 | 579 |
| Synaphobranchus_kaupii-CM056377.1 | 90.583 | 446 | 29 | 8 | 43 | 477 | 112348285 | 112347842 | 6.46e-161 | 579 |
| Synaphobranchus_kaupii-CM056377.1 | 86.433 | 457 | 26 | 14 | 43 | 477 | 129841345 | 129840903 | 1.46e-127 | 468 |
| Synaphobranchus_kaupii-CM056377.1 | 86.275 | 459 | 23 | 18 | 43 | 477 | 130019437 | 130018995 | 6.78e-126 | 462 |
| Synaphobranchus_kaupii-CM056377.1 | 89.423 | 312 | 28 | 3 | 43 | 350 | 75657486 | 75657176 | 1.17e-103 | 388 |
| Synaphobranchus_kaupii-CM056377.1 | 89.423 | 312 | 28 | 3 | 43 | 350 | 92422107 | 92422417 | 1.17e-103 | 388 |
| Synaphobranchus_kaupii-CM056377.1 | 89.423 | 312 | 28 | 3 | 43 | 350 | 116594486 | 116594796 | 1.17e-103 | 388 |
| Synaphobranchus_kaupii-CM056377.1 | 89.423 | 312 | 28 | 3 | 43 | 350 | 129005761 | 129005451 | 1.17e-103 | 388 |
| Synaphobranchus_kaupii-CM056377.1 | 89.423 | 312 | 28 | 3 | 43 | 350 | 129311842 | 129312152 | 1.17e-103 | 388 |

### N5.HOG0008885 Blastn of conserved regions against all genome assemblies

#### Region A (8/10)

| sseqid | pident | length | mismatch | gapopen | qstart | qend | sstart | send | evalue | bitscore |
|---|---|---|---|---|---|---|---|---|---|---|
| Synaphobranchus_kaupii-CM056377.1 | 89.423 | 312 | 28 | 3 | 43 | 350 | 130750092 | 130749782 | 1.17e-103 | 388 |
| Synaphobranchus_kaupii-CM056377.1 | 89.206 | 315 | 26 | 4 | 43 | 350 | 8720489 | 8720176 | 4.20e-103 | 387 |
| Synaphobranchus_kaupii-CM056377.1 | 90.033 | 301 | 26 | 2 | 43 | 339 | 122261860 | 122262160 | 4.20e-103 | 387 |
| Synaphobranchus_kaupii-CM056377.1 | 89.103 | 312 | 29 | 3 | 43 | 350 | 6101276 | 6100966 | 5.43e-102 | 383 |
| Synaphobranchus_kaupii-CM056377.1 | 89.967 | 299 | 26 | 2 | 43 | 337 | 50218781 | 50218483 | 5.43e-102 | 383 |
| Synaphobranchus_kaupii-CM056377.1 | 89.103 | 312 | 29 | 3 | 43 | 350 | 97215348 | 97215038 | 5.43e-102 | 383 |
| Synaphobranchus_kaupii-CM056377.1 | 89.701 | 301 | 27 | 2 | 43 | 339 | 116593900 | 116593600 | 1.95e-101 | 381 |
| Synaphobranchus_kaupii-CM056377.1 | 88.782 | 312 | 30 | 3 | 43 | 350 | 37504018 | 37503708 | 2.53e-100 | 377 |
| Synaphobranchus_kaupii-CM056377.1 | 88.782 | 312 | 30 | 3 | 43 | 350 | 129313412 | 129313102 | 2.53e-100 | 377 |
| Synaphobranchus_kaupii-CM056377.1 | 88.782 | 312 | 30 | 3 | 43 | 350 | 130010921 | 130011231 | 2.53e-100 | 377 |
| Synaphobranchus_kaupii-CM056377.1 | 88.141 | 312 | 32 | 3 | 43 | 350 | 19000065 | 18999755 | 5.47e-97 | 366 |
| Synaphobranchus_kaupii-CM056377.1 | 88.742 | 302 | 27 | 6 | 43 | 339 | 123330370 | 123330669 | 7.08e-96 | 363 |
| Synaphobranchus_kaupii-CM056377.1 | 87.500 | 312 | 28 | 7 | 43 | 350 | 130764822 | 130765126 | 5.51e-92 | 350 |
| Synaphobranchus_kaupii-CM056377.1 | 84.419 | 353 | 23 | 14 | 196 | 536 | 131738674 | 131739006 | 1.55e-82 | 318 |
| Synaphobranchus_kaupii-CM056377.1 | 82.372 | 312 | 26 | 5 | 43 | 350 | 24147309 | 24147023 | 2.67e-60 | 244 |
| Synaphobranchus_kaupii-CM056376.1 | 90.724 | 442 | 27 | 9 | 43 | 477 | 92791778 | 92792212 | 2.32e-160 | 577 |
| Synaphobranchus_kaupii-CM056376.1 | 89.968 | 309 | 27 | 2 | 43 | 347 | 127349520 | 127349212 | 6.98e-106 | 396 |
| Synaphobranchus_kaupii-CM056376.1 | 89.744 | 312 | 27 | 3 | 43 | 350 | 114805219 | 114805529 | 2.51e-105 | 394 |
| Synaphobranchus_kaupii-CM056376.1 | 89.423 | 312 | 28 | 3 | 43 | 350 | 47830005 | 47830315 | 1.17e-103 | 388 |
| Synaphobranchus_kaupii-CM056376.1 | 89.423 | 312 | 28 | 3 | 43 | 350 | 90265791 | 90266101 | 1.17e-103 | 388 |
| Synaphobranchus_kaupii-CM056376.1 | 89.423 | 312 | 28 | 3 | 43 | 350 | 125943096 | 125942786 | 1.17e-103 | 388 |
| Synaphobranchus_kaupii-CM056376.1 | 89.423 | 312 | 28 | 3 | 43 | 350 | 127349948 | 127350258 | 1.17e-103 | 388 |
| Synaphobranchus_kaupii-CM056376.1 | 89.423 | 312 | 28 | 3 | 43 | 350 | 141561631 | 141561321 | 1.17e-103 | 388 |
| Synaphobranchus_kaupii-CM056376.1 | 89.206 | 315 | 26 | 4 | 43 | 350 | 32315108 | 32315421 | 4.20e-103 | 387 |
| Synaphobranchus_kaupii-CM056376.1 | 90.033 | 301 | 26 | 2 | 43 | 339 | 96215277 | 96215577 | 4.20e-103 | 387 |
| Synaphobranchus_kaupii-CM056376.1 | 88.959 | 317 | 29 | 4 | 38 | 350 | 141560089 | 141560403 | 4.20e-103 | 387 |
| Synaphobranchus_kaupii-CM056376.1 | 89.137 | 313 | 29 | 3 | 43 | 351 | 109925261 | 109924950 | 1.51e-102 | 385 |
| Synaphobranchus_kaupii-CM056376.1 | 89.103 | 312 | 29 | 3 | 43 | 350 | 41597990 | 41597680 | 5.43e-102 | 383 |
| Synaphobranchus_kaupii-CM056376.1 | 89.103 | 312 | 29 | 3 | 43 | 350 | 85693182 | 85693492 | 5.43e-102 | 383 |
| Synaphobranchus_kaupii-CM056376.1 | 89.103 | 312 | 29 | 3 | 43 | 350 | 110508822 | 110508512 | 5.43e-102 | 383 |
| Synaphobranchus_kaupii-CM056376.1 | 89.542 | 306 | 27 | 3 | 38 | 339 | 124065588 | 124065284 | 5.43e-102 | 383 |
| Synaphobranchus_kaupii-CM056376.1 | 89.103 | 312 | 29 | 3 | 43 | 350 | 133770065 | 133770375 | 5.43e-102 | 383 |
| Synaphobranchus_kaupii-CM056376.1 | 89.103 | 312 | 29 | 3 | 43 | 350 | 133773714 | 133773404 | 5.43e-102 | 383 |
| Synaphobranchus_kaupii-CM056376.1 | 89.103 | 312 | 29 | 3 | 43 | 350 | 135535176 | 135535486 | 5.43e-102 | 383 |
| Synaphobranchus_kaupii-CM056376.1 | 89.103 | 312 | 29 | 3 | 43 | 350 | 141563186 | 141563496 | 5.43e-102 | 383 |
| Synaphobranchus_kaupii-CM056376.1 | 89.701 | 301 | 27 | 2 | 43 | 339 | 96897113 | 96896813 | 1.95e-101 | 381 |
| Synaphobranchus_kaupii-CM056376.1 | 89.701 | 301 | 27 | 2 | 43 | 339 | 135542843 | 135542543 | 1.95e-101 | 381 |
| Synaphobranchus_kaupii-CM056376.1 | 89.701 | 301 | 27 | 2 | 43 | 339 | 143011626 | 143011926 | 1.95e-101 | 381 |
| Synaphobranchus_kaupii-CM056376.1 | 88.782 | 312 | 29 | 4 | 43 | 350 | 74703994 | 74704303 | 2.53e-100 | 377 |
| Synaphobranchus_kaupii-CM056376.1 | 88.782 | 312 | 30 | 3 | 43 | 350 | 125135788 | 125135478 | 2.53e-100 | 377 |
| Synaphobranchus_kaupii-CM056376.1 | 88.462 | 312 | 31 | 3 | 43 | 350 | 125142209 | 125141899 | 1.18e-98 | 372 |
| Synaphobranchus_kaupii-CM056376.1 | 88.462 | 312 | 31 | 3 | 43 | 350 | 127355269 | 127354959 | 1.18e-98 | 372 |
| Synaphobranchus_kaupii-CM056376.1 | 88.254 | 315 | 29 | 4 | 43 | 350 | 133795004 | 133794691 | 4.23e-98 | 370 |
| Synaphobranchus_kaupii-CM056376.1 | 87.821 | 312 | 29 | 4 | 43 | 350 | 118313735 | 118313429 | 3.29e-94 | 357 |
| Synaphobranchus_kaupii-CM056376.1 | 84.365 | 307 | 24 | 3 | 43 | 345 | 96218587 | 96218301 | 7.33e-71 | 279 |
| Synaphobranchus_kaupii-CM056376.1 | 93.011 | 186 | 11 | 1 | 43 | 226 | 53081750 | 53081565 | 4.41e-68 | 270 |
| Synaphobranchus_kaupii-CM056399.1 | 90.337 | 445 | 32 | 6 | 43 | 477 | 3527507 | 3527950 | 3.00e-159 | 573 |
| Synaphobranchus_kaupii-CM056399.1 | 89.423 | 312 | 28 | 3 | 43 | 350 | 12562517 | 12562207 | 1.17e-103 | 388 |
| Synaphobranchus_kaupii-CM056399.1 | 89.103 | 312 | 29 | 3 | 43 | 350 | 376312 | 376002 | 5.43e-102 | 383 |
| Synaphobranchus_kaupii-CM056399.1 | 89.103 | 312 | 29 | 3 | 43 | 350 | 687671 | 687361 | 5.43e-102 | 383 |
| Synaphobranchus_kaupii-CM056399.1 | 89.103 | 312 | 29 | 3 | 43 | 350 | 2241783 | 2242093 | 5.43e-102 | 383 |
| Synaphobranchus_kaupii-CM056399.1 | 89.103 | 312 | 29 | 3 | 43 | 350 | 6212124 | 6211814 | 5.43e-102 | 383 |
| Synaphobranchus_kaupii-CM056399.1 | 89.103 | 312 | 29 | 3 | 43 | 350 | 8683151 | 8682841 | 5.43e-102 | 383 |
| Synaphobranchus_kaupii-CM056399.1 | 89.103 | 312 | 29 | 3 | 43 | 350 | 12648024 | 12647714 | 5.43e-102 | 383 |
| Synaphobranchus_kaupii-CM056399.1 | 89.701 | 301 | 27 | 2 | 43 | 339 | 6941108 | 6941408 | 1.95e-101 | 381 |
| Synaphobranchus_kaupii-CM056399.1 | 88.782 | 312 | 30 | 3 | 43 | 350 | 8638848 | 8638538 | 2.53e-100 | 377 |
| Synaphobranchus_kaupii-CM056399.1 | 88.782 | 312 | 28 | 4 | 43 | 350 | 12568281 | 12567973 | 9.09e-100 | 375 |
| Synaphobranchus_kaupii-CM056399.1 | 88.254 | 315 | 31 | 4 | 43 | 352 | 8636739 | 8637052 | 1.18e-98 | 372 |
| Synaphobranchus_kaupii-CM056399.1 | 88.852 | 305 | 28 | 4 | 43 | 345 | 12573825 | 12574125 | 4.23e-98 | 370 |
| Synaphobranchus_kaupii-CM056399.1 | 92.935 | 184 | 11 | 1 | 43 | 224 | 12648391 | 12648208 | 5.71e-67 | 267 |
| Clupea_harengus-NW_024879908.1 | 87.698 | 504 | 42 | 13 | 52 | 536 | 48854 | 48352 | 3.89e-158 | 569 |
| Clupea_harengus-NW_024879647.1 | 87.402 | 508 | 45 | 15 | 43 | 536 | 21464 | 21966 | 5.03e-157 | 566 |
| Clupea_harengus-NW_024879735.1 | 87.649 | 502 | 38 | 14 | 38 | 536 | 51486 | 51966 | 6.50e-156 | 562 |
| Clupea_harengus-NW_024879735.1 | 87.475 | 503 | 38 | 15 | 38 | 536 | 56240 | 56721 | 3.02e-154 | 556 |
| Clupea_harengus-NW_024879735.1 | 87.205 | 508 | 38 | 16 | 38 | 536 | 66515 | 67004 | 3.91e-153 | 553 |
| Clupea_harengus-NW_024879735.1 | 87.059 | 510 | 37 | 18 | 38 | 536 | 71272 | 71763 | 5.06e-152 | 549 |
| Clupea_harengus-NW_024879735.1 | 87.008 | 508 | 39 | 16 | 38 | 536 | 61376 | 61865 | 1.82e-151 | 547 |
| Clupea_harengus-NW_024879735.1 | 86.863 | 510 | 38 | 17 | 38 | 536 | 76389 | 76880 | 2.35e-150 | 544 |
| Clupea_harengus-NW_024879735.1 | 89.500 | 400 | 30 | 8 | 38 | 434 | 21242 | 21632 | 6.69e-136 | 496 |
| Clupea_harengus-NW_024879798.1 | 87.109 | 512 | 39 | 20 | 52 | 536 | 13190 | 12679 | 1.09e-153 | 555 |
| Clupea_harengus-NW_024880259.1 | 85.988 | 521 | 50 | 18 | 38 | 536 | 23490 | 24009 | 3.94e-148 | 536 |
| Synaphobranchus_kaupii-CM056382.1 | 88.789 | 446 | 28 | 10 | 43 | 477 | 23345544 | 23345110 | 2.37e-145 | 527 |
| Synaphobranchus_kaupii-CM056382.1 | 88.889 | 360 | 27 | 5 | 43 | 392 | 23347997 | 23347641 | 1.91e-116 | 431 |
| Synaphobranchus_kaupii-CM056382.1 | 89.744 | 312 | 27 | 3 | 43 | 350 | 55151188 | 55150878 | 2.51e-105 | 394 |
| Synaphobranchus_kaupii-CM056382.1 | 89.423 | 312 | 28 | 3 | 43 | 350 | 21921449 | 21921759 | 1.17e-103 | 388 |
| Synaphobranchus_kaupii-CM056382.1 | 89.423 | 312 | 28 | 3 | 43 | 350 | 21925416 | 21925106 | 1.17e-103 | 388 |
| Synaphobranchus_kaupii-CM056382.1 | 89.423 | 312 | 28 | 3 | 43 | 350 | 55985534 | 55985844 | 1.17e-103 | 388 |
| Synaphobranchus_kaupii-CM056382.1 | 89.103 | 312 | 29 | 3 | 43 | 350 | 23375657 | 23375967 | 5.43e-102 | 383 |
| Synaphobranchus_kaupii-CM056382.1 | 89.103 | 312 | 29 | 3 | 43 | 350 | 32455450 | 32455140 | 5.43e-102 | 383 |
| Synaphobranchus_kaupii-CM056382.1 | 89.103 | 312 | 29 | 3 | 43 | 350 | 44225938 | 44226248 | 5.43e-102 | 383 |
| Synaphobranchus_kaupii-CM056382.1 | 89.103 | 312 | 29 | 3 | 43 | 350 | 57547732 | 57547422 | 5.43e-102 | 383 |
| Synaphobranchus_kaupii-CM056382.1 | 89.103 | 312 | 29 | 3 | 43 | 350 | 73998974 | 73999284 | 5.43e-102 | 383 |
| Synaphobranchus_kaupii-CM056382.1 | 89.701 | 301 | 27 | 2 | 43 | 339 | 18672051 | 18672351 | 1.95e-101 | 381 |
| Synaphobranchus_kaupii-CM056382.1 | 89.369 | 301 | 28 | 2 | 43 | 339 | 41359303 | 41359003 | 9.09e-100 | 375 |
| Synaphobranchus_kaupii-CM056382.1 | 88.462 | 312 | 31 | 3 | 43 | 350 | 67920198 | 67919888 | 1.18e-98 | 372 |
| Synaphobranchus_kaupii-CM056382.1 | 81.513 | 357 | 49 | 8 | 45 | 397 | 15400811 | 15401154 | 2.64e-70 | 278 |
| Synaphobranchus_kaupii-CM056382.1 | 83.923 | 311 | 25 | 7 | 44 | 350 | 51980168 | 51979879 | 3.41e-69 | 274 |
| Synaphobranchus_kaupii-CM056382.1 | 83.492 | 315 | 26 | 5 | 43 | 353 | 67918312 | 67918020 | 4.41e-68 | 270 |
| Synaphobranchus_kaupii-CM056382.1 | 82.692 | 312 | 24 | 8 | 43 | 350 | 51978668 | 51978953 | 5.75e-62 | 250 |
| Coryphaenoides_rupestris-PIJE02008971.1 | 88.692 | 451 | 26 | 8 | 96 | 536 | 7956 | 8391 | 2.37e-145 | 527 |
| Coryphaenoides_rupestris-PIJE02008971.1 | 93.229 | 192 | 13 | 0 | 43 | 234 | 192 | 1 | 5.67e-72 | 283 |
| Clupea_harengus-NW_024879597.1 | 85.855 | 509 | 45 | 8 | 43 | 536 | 113772 | 114268 | 5.13e-142 | 516 |
| Coryphaenoides_rupestris-PIJE02001954.1 | 87.832 | 452 | 22 | 15 | 43 | 488 | 68015 | 68439 | 5.17e-137 | 499 |
| Coryphaenoides_rupestris-PIJE02001954.1 | 87.224 | 407 | 25 | 10 | 43 | 434 | 62390 | 61996 | 1.14e-118 | 438 |
| Coryphaenoides_rupestris-PIJE02002217.1 | 88.670 | 406 | 32 | 7 | 43 | 434 | 7791 | 7386 | 5.21e-132 | 483 |
| Coryphaenoides_rupestris-PIJE02000179.1 | 88.889 | 405 | 27 | 8 | 127 | 513 | 347061 | 347465 | 5.21e-132 | 483 |
| Coryphaenoides_rupestris-PIJE02000271.1 | 88.191 | 398 | 31 | 9 | 43 | 434 | 372875 | 373262 | 2.44e-125 | 460 |
| Coryphaenoides_rupestris-PIJE02000271.1 | 87.469 | 399 | 31 | 11 | 43 | 434 | 361263 | 361649 | 8.84e-120 | 442 |
| Coryphaenoides_rupestris-PIJE02018184.1 | 87.940 | 398 | 32 | 9 | 43 | 434 | 966 | 1353 | 1.14e-123 | 455 |
| Coryphaenoides_rupestris-PIJE02007852.1 | 88.161 | 397 | 27 | 9 | 43 | 434 | 885 | 1266 | 1.14e-123 | 455 |
| Coryphaenoides_rupestris-PIJE02000083.1 | 87.940 | 398 | 32 | 9 | 43 | 434 | 7920 | 7533 | 1.14e-123 | 455 |
| Coryphaenoides_rupestris-PIJE02000504.1 | 88.342 | 386 | 33 | 7 | 157 | 534 | 182190 | 182571 | 4.08e-123 | 453 |
| Coryphaenoides_rupestris-PIJE02012768.1 | 87.688 | 398 | 33 | 9 | 43 | 434 | 1803 | 2190 | 5.28e-122 | 449 |

### N5.HOG0008885 Blastn of conserved regions against all genome assemblies

#### Region A (9/10)

| sseqid | pident | length | mismatch | gapopen | qstart | qend | sstart | send | evalue | bitscore |
|---|---|---|---|---|---|---|---|---|---|---|
| Clupea_harengus-NW_024880560.1 | 84.040 | 495 | 47 | 5 | 43 | 536 | 18634 | 18171 | 1.90e-121 | 448 |
| Coryphaenoides_rupestris-PIJE02001593.1 | 87.848 | 395 | 26 | 10 | 45 | 434 | 20977 | 20600 | 2.46e-120 | 444 |
| Coryphaenoides_rupestris-PIJE02000229.1 | 87.437 | 398 | 33 | 10 | 43 | 434 | 229022 | 228636 | 8.84e-120 | 442 |
| Coryphaenoides_rupestris-PIJE02000217.1 | 87.848 | 395 | 25 | 9 | 43 | 434 | 227095 | 226721 | 8.84e-120 | 442 |
| Coryphaenoides_rupestris-PIJE02000389.1 | 87.406 | 397 | 30 | 10 | 43 | 434 | 37569 | 37950 | 1.14e-118 | 438 |
| Coryphaenoides_rupestris-PIJE02000389.1 | 97.500 | 80 | 2 | 0 | 355 | 434 | 35422 | 35343 | 4.67e-28 | 137 |
| Coryphaenoides_rupestris-PIJE02000389.1 | 97.500 | 80 | 2 | 0 | 355 | 434 | 35763 | 35842 | 4.67e-28 | 137 |
| Coryphaenoides_rupestris-PIJE02000076.1 | 87.186 | 398 | 34 | 10 | 43 | 434 | 549201 | 548815 | 4.11e-118 | 436 |
| Coryphaenoides_rupestris-PIJE02001839.1 | 87.245 | 392 | 34 | 10 | 43 | 434 | 69613 | 69988 | 5.32e-117 | 433 |
| Coryphaenoides_rupestris-PIJE02000560.1 | 86.935 | 398 | 36 | 9 | 43 | 434 | 204122 | 203735 | 5.32e-117 | 433 |
| Coryphaenoides_rupestris-PIJE02000136.1 | 87.154 | 397 | 31 | 10 | 43 | 434 | 23438 | 23819 | 5.32e-117 | 433 |
| Coryphaenoides_rupestris-PIJE02002971.1 | 87.532 | 393 | 22 | 9 | 43 | 434 | 34594 | 34960 | 6.88e-116 | 429 |
| Coryphaenoides_rupestris-PIJE02000393.1 | 86.935 | 398 | 30 | 7 | 43 | 434 | 139256 | 139637 | 2.47e-115 | 427 |
| Coryphaenoides_rupestris-PIJE02000215.1 | 86.683 | 398 | 37 | 9 | 43 | 434 | 139183 | 139570 | 2.47e-115 | 427 |
| Coryphaenoides_rupestris-PIJE02000113.1 | 86.750 | 400 | 33 | 10 | 43 | 434 | 260696 | 260309 | 2.47e-115 | 427 |
| Coryphaenoides_rupestris-PIJE02002957.1 | 88.108 | 370 | 22 | 11 | 43 | 412 | 4738 | 5085 | 4.14e-113 | 420 |
| Coryphaenoides_rupestris-PIJE02000114.1 | 86.352 | 403 | 34 | 13 | 43 | 434 | 680077 | 680469 | 4.14e-113 | 420 |
| Coryphaenoides_rupestris-PIJE02006056.1 | 87.467 | 375 | 31 | 9 | 66 | 434 | 15528 | 15164 | 1.49e-112 | 418 |
| Coryphaenoides_rupestris-PIJE02006056.1 | 85.169 | 236 | 13 | 16 | 219 | 434 | 11039 | 11272 | 1.25e-53 | 222 |
| Coryphaenoides_rupestris-PIJE02000766.1 | 86.990 | 392 | 20 | 15 | 43 | 434 | 57400 | 57760 | 6.93e-111 | 412 |
| Coryphaenoides_rupestris-PIJE02000766.1 | 85.360 | 403 | 29 | 9 | 43 | 434 | 58788 | 59171 | 3.25e-104 | 390 |
| Coryphaenoides_rupestris-PIJE02013360.1 | 86.480 | 392 | 32 | 8 | 43 | 434 | 1322 | 1692 | 2.49e-110 | 411 |
| Coryphaenoides_rupestris-PIJE02001381.1 | 86.104 | 403 | 30 | 16 | 43 | 434 | 24475 | 24862 | 2.49e-110 | 411 |
| Coryphaenoides_rupestris-PIJE02001381.1 | 84.323 | 421 | 26 | 24 | 43 | 434 | 25295 | 25704 | 9.09e-100 | 375 |
| Coryphaenoides_rupestris-PIJE02001381.1 | 90.948 | 232 | 9 | 11 | 309 | 536 | 26858 | 27081 | 1.56e-77 | 302 |
| Coryphaenoides_rupestris-PIJE02000201.1 | 87.003 | 377 | 31 | 14 | 66 | 434 | 383081 | 382715 | 8.96e-110 | 409 |
| Coryphaenoides_rupestris-PIJE02001073.1 | 89.538 | 325 | 23 | 6 | 43 | 356 | 106352 | 106676 | 1.50e-107 | 401 |
| Synaphobranchus_kaupii- | 90.064 | 312 | 26 | 3 | 43 | 350 | 785796 | 785486 | 5.39e-107 | 399 |
| Synaphobranchus_kaupii- | 89.274 | 317 | 25 | 5 | 38 | 347 | 196133 | 195819 | 1.17e-103 | 388 |
| Synaphobranchus_kaupii- | 89.735 | 302 | 27 | 2 | 43 | 340 | 714976 | 714675 | 5.43e-102 | 383 |
| Synaphobranchus_kaupii- | 88.571 | 315 | 30 | 4 | 43 | 352 | 787186 | 787499 | 2.53e-100 | 377 |
| Synaphobranchus_kaupii- | 88.462 | 312 | 31 | 3 | 43 | 350 | 3763882 | 3763572 | 1.18e-98 | 372 |
| Synaphobranchus_kaupii- | 89.262 | 298 | 26 | 4 | 43 | 338 | 717440 | 717147 | 1.52e-97 | 368 |
| Synaphobranchus_kaupii- | 83.974 | 312 | 25 | 4 | 43 | 350 | 4263515 | 4263225 | 9.49e-70 | 276 |
| Synaphobranchus_kaupii- | 83.492 | 315 | 22 | 13 | 43 | 347 | 4941255 | 4940961 | 5.71e-67 | 267 |
| Synaphobranchus_kaupii- | 93.258 | 178 | 9 | 2 | 38 | 213 | 183713 | 183889 | 9.55e-65 | 259 |
| Coryphaenoides_rupestris-PIJE02000208.1 | 86.189 | 391 | 27 | 11 | 45 | 434 | 361368 | 361732 | 1.94e-106 | 398 |
| Coryphaenoides_rupestris-PIJE02021828.1 | 89.164 | 323 | 26 | 8 | 43 | 356 | 2705 | 2383 | 2.51e-105 | 394 |
| Synaphobranchus_kaupii-CM056395.1 | 89.524 | 315 | 25 | 4 | 43 | 350 | 22066344 | 22066031 | 9.03e-105 | 392 |
| Synaphobranchus_kaupii-CM056395.1 | 89.423 | 312 | 28 | 3 | 43 | 350 | 24571124 | 24571434 | 1.17e-103 | 388 |
| Synaphobranchus_kaupii-CM056395.1 | 89.206 | 315 | 26 | 4 | 43 | 350 | 20053120 | 20053433 | 4.20e-103 | 387 |
| Synaphobranchus_kaupii-CM056395.1 | 89.206 | 315 | 26 | 4 | 43 | 350 | 20069708 | 20070021 | 4.20e-103 | 387 |
| Synaphobranchus_kaupii-CM056395.1 | 89.206 | 315 | 26 | 4 | 43 | 350 | 20071697 | 20071384 | 4.20e-103 | 387 |
| Synaphobranchus_kaupii-CM056395.1 | 89.103 | 312 | 29 | 3 | 43 | 350 | 7232437 | 7232127 | 5.43e-102 | 383 |
| Synaphobranchus_kaupii-CM056395.1 | 88.889 | 315 | 29 | 4 | 43 | 352 | 24572659 | 24572346 | 5.43e-102 | 383 |
| Synaphobranchus_kaupii-CM056395.1 | 89.103 | 312 | 27 | 4 | 43 | 350 | 5826311 | 5826003 | 1.95e-101 | 381 |
| Synaphobranchus_kaupii-CM056395.1 | 88.571 | 315 | 30 | 4 | 43 | 352 | 20061558 | 20061871 | 2.53e-100 | 377 |
| Synaphobranchus_kaupii-CM056392.1 | 89.274 | 317 | 28 | 4 | 38 | 350 | 13600561 | 13600875 | 9.03e-105 | 392 |
| Synaphobranchus_kaupii-CM056392.1 | 89.206 | 315 | 28 | 4 | 43 | 353 | 18248149 | 18248461 | 1.17e-103 | 388 |
| Synaphobranchus_kaupii-CM056392.1 | 89.423 | 312 | 28 | 3 | 43 | 350 | 23141525 | 23141835 | 1.17e-103 | 388 |
| Synaphobranchus_kaupii-CM056392.1 | 89.206 | 315 | 26 | 4 | 43 | 350 | 13607622 | 13607935 | 4.20e-103 | 387 |
| Synaphobranchus_kaupii-CM056392.1 | 89.103 | 312 | 29 | 3 | 43 | 350 | 13607164 | 13606854 | 5.43e-102 | 383 |
| Synaphobranchus_kaupii-CM056392.1 | 88.782 | 312 | 30 | 3 | 43 | 350 | 32393059 | 32392749 | 2.53e-100 | 377 |
| Synaphobranchus_kaupii-CM056392.1 | 88.462 | 312 | 31 | 3 | 43 | 350 | 14472363 | 14472053 | 1.18e-98 | 372 |
| Synaphobranchus_kaupii-CM056392.1 | 94.483 | 145 | 6 | 1 | 43 | 185 | 2630965 | 2630821 | 1.25e-53 | 222 |
| Coryphaenoides_rupestris-PIJE02000390.1 | 87.187 | 359 | 28 | 10 | 43 | 394 | 174560 | 174907 | 9.03e-105 | 392 |
| Coryphaenoides_rupestris-PIJE02025593.1 | 88.754 | 329 | 24 | 6 | 43 | 360 | 2 | 328 | 3.25e-104 | 390 |
| Synaphobranchus_kaupii- | 89.423 | 312 | 28 | 3 | 43 | 350 | 224851 | 224541 | 1.17e-103 | 388 |
| Synaphobranchus_kaupii-CM056398.1 | 89.423 | 312 | 28 | 3 | 43 | 350 | 4629840 | 4630150 | 1.17e-103 | 388 |
| Synaphobranchus_kaupii-CM056398.1 | 89.172 | 314 | 29 | 3 | 38 | 347 | 4282700 | 4282388 | 4.20e-103 | 387 |
| Synaphobranchus_kaupii-CM056398.1 | 89.667 | 300 | 27 | 2 | 43 | 338 | 6796717 | 6797016 | 7.03e-101 | 379 |
| Synaphobranchus_kaupii-CM056398.1 | 88.235 | 221 | 24 | 1 | 121 | 339 | 6800762 | 6800542 | 7.38e-66 | 263 |
| Synaphobranchus_kaupii-CM056397.1 | 89.423 | 312 | 28 | 3 | 43 | 350 | 5024667 | 5024357 | 1.17e-103 | 388 |
| Synaphobranchus_kaupii-CM056390.1 | 89.423 | 312 | 28 | 3 | 43 | 350 | 586642 | 586332 | 1.17e-103 | 388 |
| Synaphobranchus_kaupii-CM056390.1 | 89.542 | 306 | 27 | 3 | 38 | 339 | 6052363 | 6052667 | 5.43e-102 | 383 |
| Synaphobranchus_kaupii-CM056390.1 | 89.542 | 306 | 27 | 3 | 38 | 339 | 6491788 | 6492092 | 5.43e-102 | 383 |
| Synaphobranchus_kaupii-CM056390.1 | 89.103 | 312 | 29 | 3 | 43 | 350 | 22161631 | 22161941 | 5.43e-102 | 383 |
| Synaphobranchus_kaupii-CM056390.1 | 89.701 | 301 | 27 | 2 | 43 | 339 | 4303973 | 4304273 | 1.95e-101 | 381 |
| Synaphobranchus_kaupii-CM056390.1 | 89.701 | 301 | 27 | 2 | 43 | 339 | 17059906 | 17059606 | 1.95e-101 | 381 |
| Synaphobranchus_kaupii-CM056390.1 | 88.291 | 316 | 28 | 6 | 43 | 350 | 3790161 | 3790475 | 4.23e-98 | 370 |
| Synaphobranchus_kaupii-CM056390.1 | 86.992 | 246 | 28 | 3 | 108 | 350 | 1385747 | 1385991 | 3.41e-69 | 274 |
| Synaphobranchus_kaupii-CM056389.1 | 89.423 | 312 | 28 | 3 | 43 | 350 | 734676 | 734986 | 1.17e-103 | 388 |
| Synaphobranchus_kaupii-CM056389.1 | 89.423 | 312 | 28 | 3 | 43 | 350 | 25058084 | 25058394 | 1.17e-103 | 388 |
| Synaphobranchus_kaupii-CM056389.1 | 88.889 | 315 | 27 | 4 | 43 | 350 | 21428224 | 21428537 | 1.95e-101 | 381 |
| Synaphobranchus_kaupii-CM056389.1 | 88.782 | 312 | 30 | 3 | 43 | 350 | 19506722 | 19507032 | 2.53e-100 | 377 |
| Synaphobranchus_kaupii-CM056387.1 | 89.423 | 312 | 28 | 3 | 43 | 350 | 836821 | 837131 | 1.17e-103 | 388 |
| Synaphobranchus_kaupii-CM056387.1 | 89.423 | 312 | 28 | 3 | 43 | 350 | 3741401 | 3741711 | 1.17e-103 | 388 |
| Synaphobranchus_kaupii-CM056387.1 | 89.423 | 312 | 28 | 3 | 43 | 350 | 5197381 | 5197691 | 1.17e-103 | 388 |
| Synaphobranchus_kaupii-CM056387.1 | 88.580 | 324 | 30 | 5 | 38 | 356 | 2517631 | 2517310 | 4.20e-103 | 387 |
| Synaphobranchus_kaupii-CM056387.1 | 89.103 | 312 | 29 | 3 | 43 | 350 | 2096465 | 2096155 | 5.43e-102 | 383 |
| Synaphobranchus_kaupii-CM056387.1 | 88.782 | 312 | 30 | 3 | 43 | 350 | 14545370 | 14545680 | 2.53e-100 | 377 |
| Synaphobranchus_kaupii-CM056387.1 | 91.912 | 136 | 11 | 0 | 338 | 473 | 2882742 | 2882607 | 3.53e-44 | 191 |
| Coryphaenoides_rupestris-PIJE02000579.1 | 86.486 | 370 | 31 | 11 | 43 | 404 | 73821 | 74179 | 1.17e-103 | 388 |
| Synaphobranchus_kaupii-CM056391.1 | 88.994 | 318 | 27 | 6 | 38 | 350 | 7757441 | 7757127 | 4.20e-103 | 387 |
| Synaphobranchus_kaupii-CM056391.1 | 88.782 | 312 | 30 | 3 | 43 | 350 | 5140598 | 5140288 | 2.53e-100 | 377 |
| Synaphobranchus_kaupii-CM056391.1 | 88.462 | 312 | 31 | 3 | 43 | 350 | 7768962 | 7768652 | 1.18e-98 | 372 |
| Synaphobranchus_kaupii-CM056391.1 | 88.462 | 312 | 31 | 3 | 43 | 350 | 27853695 | 27854005 | 1.18e-98 | 372 |
| Synaphobranchus_kaupii-CM056391.1 | 86.218 | 312 | 28 | 4 | 43 | 350 | 23386236 | 23386536 | 3.34e-84 | 324 |
| Synaphobranchus_kaupii-CM056391.1 | 82.692 | 312 | 29 | 4 | 43 | 350 | 23864529 | 23864239 | 4.44e-63 | 254 |
| Synaphobranchus_kaupii-CM056388.1 | 88.959 | 317 | 29 | 4 | 38 | 350 | 6421842 | 6421528 | 4.20e-103 | 387 |
| Coryphaenoides_rupestris-PIJE02000411.1 | 88.650 | 326 | 25 | 10 | 43 | 356 | 270606 | 270281 | 4.20e-103 | 387 |
| Coryphaenoides_rupestris-PIJE02000191.1 | 88.450 | 329 | 26 | 5 | 43 | 360 | 414629 | 414956 | 4.20e-103 | 387 |
| Coryphaenoides_rupestris-PIJE02000739.1 | 89.967 | 299 | 28 | 1 | 43 | 339 | 95874 | 96172 | 1.51e-102 | 385 |
| Coryphaenoides_rupestris-PIJE02000417.1 | 89.735 | 302 | 27 | 3 | 43 | 341 | 349345 | 349045 | 5.43e-102 | 383 |
| Coryphaenoides_rupestris-PIJE02024745.1 | 86.630 | 359 | 30 | 10 | 43 | 394 | 504 | 157 | 1.95e-101 | 381 |

### N5.HOG0008885 Blastn of conserved regions against all genome assemblies

#### Region A (10/10)

| sseqid | pident | length | mismatch | gapopen | qstart | qend | sstart | send | evalue | bitscore |
|---|---|---|---|---|---|---|---|---|---|---|
| Coryphaenoides_rupestris-PIJE02005585.1 | 86.630 | 359 | 30 | 10 | 43 | 394 | 13865 | 14212 | 1.95e-101 | 381 |
| Synaphobranchus_kaupii-CM056383.1 | 88.782 | 312 | 30 | 3 | 43 | 350 | 52635354 | 52635664 | 2.53e-100 | 377 |
| Synaphobranchus_kaupii-CM056383.1 | 88.782 | 312 | 30 | 3 | 43 | 350 | 55736834 | 55736524 | 2.53e-100 | 377 |
| Synaphobranchus_kaupii-CM056383.1 | 88.571 | 315 | 28 | 4 | 43 | 350 | 24327200 | 24326887 | 9.09e-100 | 375 |
| Synaphobranchus_kaupii-CM056383.1 | 89.369 | 301 | 25 | 5 | 43 | 340 | 10452331 | 10452627 | 1.18e-98 | 372 |
| Synaphobranchus_kaupii-CM056383.1 | 88.462 | 312 | 31 | 3 | 43 | 350 | 52647218 | 52647528 | 1.18e-98 | 372 |
| Synaphobranchus_kaupii-CM056383.1 | 89.492 | 295 | 25 | 4 | 43 | 335 | 52670240 | 52670530 | 1.52e-97 | 368 |
| Synaphobranchus_kaupii-CM056383.1 | 87.697 | 317 | 29 | 4 | 43 | 350 | 60663928 | 60663613 | 2.55e-95 | 361 |
| Coryphaenoides_rupestris-PIJE02000196.1 | 85.496 | 393 | 22 | 15 | 43 | 434 | 230402 | 230044 | 2.53e-100 | 377 |
| Coryphaenoides_rupestris-PIJE02011586.1 | 84.337 | 415 | 32 | 18 | 45 | 434 | 5154 | 4748 | 9.09e-100 | 375 |
| Coryphaenoides_rupestris-PIJE02002692.1 | 86.969 | 353 | 22 | 9 | 43 | 394 | 22058 | 22387 | 9.09e-100 | 375 |
| Coryphaenoides_rupestris-PIJE02002692.1 | 86.080 | 352 | 20 | 12 | 43 | 394 | 17673 | 17995 | 1.53e-92 | 351 |
| Coryphaenoides_rupestris-PIJE02001659.1 | 85.224 | 379 | 39 | 9 | 58 | 434 | 67845 | 68208 | 3.27e-99 | 374 |
| Coryphaenoides_rupestris-PIJE02000700.1 | 85.309 | 388 | 28 | 16 | 62 | 432 | 28437 | 28812 | 3.27e-99 | 374 |
| Coryphaenoides_rupestris-PIJE02001017.1 | 87.692 | 325 | 29 | 4 | 43 | 356 | 26995 | 26671 | 1.52e-97 | 368 |
| Clupea_harengus-NW_024880723.1 | 87.234 | 329 | 31 | 6 | 43 | 362 | 7392 | 7718 | 1.97e-96 | 364 |
| Coryphaenoides_rupestris-PIJE02019434.1 | 91.892 | 259 | 21 | 0 | 43 | 301 | 2999 | 3257 | 7.08e-96 | 363 |
| Coryphaenoides_rupestris-PIJE02004644.1 | 84.733 | 393 | 27 | 18 | 43 | 434 | 1297 | 1657 | 7.08e-96 | 363 |
| Coryphaenoides_rupestris-PIJE02001555.1 | 86.455 | 347 | 27 | 12 | 43 | 381 | 7729 | 8063 | 7.08e-96 | 363 |
| Coryphaenoides_rupestris-PIJE02000454.1 | 87.821 | 312 | 35 | 3 | 43 | 354 | 4281 | 4589 | 7.08e-96 | 363 |
| Coryphaenoides_rupestris-PIJE02000454.1 | 84.000 | 225 | 26 | 3 | 134 | 356 | 279134 | 278918 | 3.51e-49 | 207 |
| Coryphaenoides_rupestris-PIJE02010167.1 | 85.411 | 377 | 18 | 19 | 59 | 434 | 688 | 348 | 3.29e-94 | 357 |
| Clupea_harengus-NW_024880607.1 | 91.078 | 269 | 15 | 7 | 272 | 536 | 13752 | 14015 | 1.18e-93 | 355 |
| Clupea_harengus-NW_024880607.1 | 88.750 | 160 | 16 | 1 | 43 | 202 | 13584 | 13741 | 2.73e-45 | 195 |
| Coryphaenoides_rupestris-PIJE02000154.1 | 87.138 | 311 | 36 | 3 | 43 | 351 | 433140 | 432832 | 5.51e-92 | 350 |
| Coryphaenoides_rupestris-PIJE02000154.1 | 94.231 | 208 | 8 | 4 | 332 | 536 | 432691 | 432485 | 2.01e-81 | 315 |
| Coryphaenoides_rupestris-PIJE02002288.1 | 94.248 | 226 | 13 | 0 | 311 | 536 | 2559 | 2784 | 7.13e-91 | 346 |
| Coryphaenoides_rupestris-PIJE02002269.1 | 84.073 | 383 | 35 | 15 | 69 | 434 | 21653 | 21280 | 7.13e-91 | 346 |
| Coryphaenoides_rupestris-PIJE02001023.1 | 96.209 | 211 | 8 | 0 | 326 | 536 | 120796 | 120586 | 7.13e-91 | 346 |
| Coryphaenoides_rupestris-PIJE02000763.1 | 94.248 | 226 | 13 | 0 | 311 | 536 | 55363 | 55138 | 7.13e-91 | 346 |
| Coryphaenoides_rupestris-PIJE02000763.1 | 94.570 | 221 | 12 | 0 | 43 | 263 | 56415 | 56195 | 9.22e-90 | 342 |
| Coryphaenoides_rupestris-PIJE02002510.1 | 82.974 | 417 | 36 | 21 | 45 | 434 | 6614 | 6206 | 2.56e-90 | 344 |
| Clupea_harengus-NW_024880647.1 | 95.392 | 217 | 7 | 2 | 320 | 536 | 8938 | 8725 | 9.22e-90 | 342 |
| Coryphaenoides_rupestris-PIJE02000079.1 | 93.805 | 226 | 14 | 0 | 311 | 536 | 117414 | 117639 | 3.32e-89 | 340 |
| Coryphaenoides_rupestris-PIJE02000586.1 | 85.976 | 328 | 26 | 15 | 43 | 353 | 206723 | 207047 | 5.55e-87 | 333 |
| Coryphaenoides_rupestris-PIJE02000011.1 | 84.659 | 352 | 29 | 16 | 55 | 394 | 1090999 | 1090661 | 2.58e-85 | 327 |
| Coryphaenoides_rupestris-PIJE02000589.1 | 85.714 | 322 | 30 | 9 | 119 | 434 | 109919 | 110230 | 9.29e-85 | 326 |
| Coryphaenoides_rupestris-PIJE02000589.1 | 85.127 | 316 | 22 | 12 | 119 | 434 | 117481 | 117771 | 5.63e-77 | 300 |
| Coryphaenoides_rupestris-PIJE02000344.1 | 85.000 | 340 | 27 | 9 | 66 | 394 | 201788 | 202114 | 3.34e-84 | 324 |
| Coryphaenoides_rupestris-PIJE02000103.1 | 82.456 | 399 | 36 | 18 | 43 | 434 | 178640 | 179011 | 1.55e-82 | 318 |
| Coryphaenoides_rupestris-PIJE02000359.1 | 85.669 | 314 | 29 | 9 | 127 | 434 | 35557 | 35254 | 5.59e-82 | 316 |
| Coryphaenoides_rupestris-PIJE02000144.1 | 86.333 | 300 | 30 | 3 | 43 | 339 | 445229 | 444938 | 5.59e-82 | 316 |
| Coryphaenoides_rupestris-PIJE02004308.1 | 91.026 | 234 | 15 | 6 | 279 | 509 | 5174 | 4944 | 2.60e-80 | 311 |
| Clupea_harengus-NW_024881065.1 | 86.572 | 283 | 35 | 3 | 52 | 333 | 281 | 1 | 9.35e-80 | 309 |
| Clupea_harengus-NW_024881095.1 | 96.721 | 183 | 5 | 1 | 355 | 536 | 379 | 197 | 4.35e-78 | 303 |
| Clupea_harengus-NW_024879711.1 | 82.778 | 360 | 42 | 8 | 43 | 398 | 12253 | 11910 | 4.35e-78 | 303 |
| Coryphaenoides_rupestris-PIJE02000356.1 | 93.204 | 206 | 12 | 2 | 43 | 248 | 330126 | 329923 | 1.56e-77 | 302 |
| Coryphaenoides_rupestris-PIJE02000970.1 | 86.505 | 289 | 17 | 8 | 43 | 328 | 37248 | 36979 | 2.02e-76 | 298 |
| Coryphaenoides_rupestris-PIJE02000692.1 | 85.859 | 297 | 21 | 8 | 43 | 338 | 54517 | 54241 | 7.28e-76 | 296 |
| Coryphaenoides_rupestris-PIJE02002354.1 | 90.950 | 221 | 13 | 1 | 43 | 263 | 22606 | 22393 | 3.39e-74 | 291 |
| Coryphaenoides_rupestris-PIJE02000091.1 | 90.498 | 221 | 14 | 1 | 43 | 263 | 283741 | 283954 | 1.58e-72 | 285 |
| Clupea_harengus-NW_024879593.1 | 91.707 | 205 | 16 | 1 | 43 | 246 | 83020 | 83224 | 5.67e-72 | 283 |
| Coryphaenoides_rupestris-PIJE02003069.1 | 83.912 | 317 | 29 | 4 | 43 | 356 | 9031 | 9328 | 5.67e-72 | 283 |
| Coryphaenoides_rupestris-PIJE02003069.1 | 91.753 | 97 | 6 | 2 | 45 | 139 | 42006 | 42102 | 6.04e-27 | 134 |
| Coryphaenoides_rupestris-PIJE02010023.1 | 86.447 | 273 | 17 | 9 | 167 | 434 | 2840 | 2583 | 2.04e-71 | 281 |
| Clupea_harengus-NW_024880188.1 | 95.758 | 165 | 7 | 0 | 43 | 207 | 19276 | 19112 | 5.71e-67 | 267 |
| Clupea_harengus-NW_024880706.1 | 93.865 | 163 | 10 | 0 | 374 | 536 | 2870 | 3032 | 7.44e-61 | 246 |
| Clupea_harengus-NW_024880706.1 | 93.750 | 160 | 8 | 2 | 43 | 201 | 2707 | 2865 | 1.24e-58 | 239 |
| Clupea_harengus-NW_024879567.1 | 94.805 | 154 | 8 | 0 | 383 | 536 | 74630 | 74477 | 3.46e-59 | 241 |
| Clupea_harengus-NW_024879754.1 | 89.529 | 191 | 15 | 5 | 43 | 230 | 14833 | 15021 | 4.48e-58 | 237 |
| Coryphaenoides_rupestris-PIJE02000124.1 | 83.643 | 269 | 26 | 11 | 173 | 434 | 254925 | 254668 | 4.48e-58 | 237 |
| Coryphaenoides_rupestris-PIJE02001348.1 | 92.121 | 165 | 13 | 0 | 45 | 209 | 79252 | 79088 | 5.79e-57 | 233 |
| Coryphaenoides_rupestris-PIJE02003827.1 | 81.115 | 323 | 27 | 19 | 43 | 356 | 22153 | 22450 | 2.69e-55 | 228 |
| Coryphaenoides_rupestris-PIJE02003639.1 | 87.379 | 206 | 15 | 6 | 240 | 434 | 5280 | 5075 | 9.69e-55 | 226 |
| Coryphaenoides_rupestris-PIJE02017629.1 | 92.357 | 157 | 12 | 0 | 43 | 199 | 162 | 6 | 3.48e-54 | 224 |
| Clupea_harengus-NW_024880797.1 | 90.964 | 166 | 12 | 2 | 43 | 207 | 2411 | 2574 | 4.51e-53 | 220 |
| Coryphaenoides_rupestris-PIJE02000170.1 | 88.649 | 185 | 12 | 3 | 67 | 249 | 47353 | 47530 | 5.83e-52 | 217 |
| Coryphaenoides_rupestris-PIJE02000891.1 | 92.466 | 146 | 11 | 0 | 311 | 456 | 43990 | 43845 | 9.76e-50 | 209 |
| Coryphaenoides_rupestris-PIJE02044490.1 | 82.625 | 259 | 17 | 10 | 80 | 336 | 233 | 1 | 4.54e-48 | 204 |
| Coryphaenoides_rupestris-PIJE02000014.1 | 84.360 | 211 | 23 | 9 | 230 | 434 | 175936 | 176142 | 2.11e-46 | 198 |
| Coryphaenoides_rupestris-PIJE02000499.1 | 88.024 | 167 | 9 | 6 | 279 | 434 | 285738 | 285904 | 4.57e-43 | 187 |
| Coryphaenoides_rupestris-PIJE02044399.1 | 93.496 | 123 | 8 | 0 | 312 | 434 | 1 | 123 | 5.91e-42 | 183 |
| Hoplostethus_atlanticus- | 88.811 | 143 | 16 | 0 | 344 | 486 | 16711 | 16853 | 9.90e-40 | 176 |
| Clupea_harengus-NW_024879582.1 | 96.875 | 96 | 3 | 0 | 43 | 138 | 12224 | 12129 | 2.77e-35 | 161 |
| Clupea_harengus-NW_024879582.1 | 96.875 | 96 | 3 | 0 | 43 | 138 | 178213 | 178118 | 2.77e-35 | 161 |
| Coryphaenoides_rupestris-PIJE02000121.1 | 86.986 | 146 | 13 | 1 | 47 | 192 | 464850 | 464989 | 9.97e-35 | 159 |
| Coryphaenoides_rupestris-PIJE02000121.1 | 93.478 | 92 | 4 | 2 | 344 | 434 | 465021 | 465111 | 1.68e-27 | 135 |
| Hoplostethus_atlanticus- | 82.979 | 188 | 18 | 5 | 321 | 508 | 18450 | 18277 | 3.58e-34 | 158 |
| Clupea_harengus-NW_024879846.1 | 95.833 | 96 | 4 | 0 | 43 | 138 | 11490 | 11395 | 1.29e-33 | 156 |
| Hoplostethus_mediterraneus- | 83.648 | 159 | 25 | 1 | 357 | 515 | 15664 | 15821 | 2.16e-31 | 148 |
| Clupea_harengus-NW_024879834.1 | 93.814 | 97 | 5 | 1 | 43 | 138 | 36514 | 36418 | 2.79e-30 | 145 |
| Coryphaenoides_rupestris-PIJE02001024.1 | 95.000 | 80 | 4 | 0 | 355 | 434 | 68087 | 68166 | 1.01e-24 | 126 |
| Coregonus_clupeaformis-NW_025534906.1 | 79.775 | 178 | 23 | 10 | 303 | 473 | 64472 | 64643 | 6.08e-22 | 117 |
| Coryphaenoides_rupestris-PIJE02001778.1 | 93.506 | 77 | 5 | 0 | 43 | 119 | 2892 | 2968 | 2.19e-21 | 115 |

### N5.HOG0008885 Blastn of conserved regions against all genome assemblies

#### Region B

| sseqid | pident | length | mismatch | gapopen | qstart | qend | sstart | send | evalue | bitscore |
|---|---|---|---|---|---|---|---|---|---|---|
| Hypomesus_transpacificus-NC_061062.1 | 100.000 | 801 | 0 | 0 | 1 | 801 | 1785680 | 1786480 | 0.0 | 1480 |
| Clupea_harengus-NC_045176.1 | 88.444 | 225 | 16 | 4 | 294 | 515 | 517099 | 517316 | 9.99e-66 | 263 |
| Clupea_harengus-NC_045176.1 | 90.299 | 134 | 9 | 4 | 671 | 801 | 517363 | 517495 | 1.73e-38 | 172 |
| Clupea_harengus-NW_024879780.1 | 90.090 | 111 | 11 | 0 | 1 | 111 | 34172 | 34062 | 3.78e-30 | 145 |
| Clupea_harengus-NW_024879598.1 | 90.090 | 111 | 11 | 0 | 1 | 111 | 134625 | 134515 | 3.78e-30 | 145 |
| Clupea_harengus-NC_045168.1 | 91.346 | 104 | 9 | 0 | 1 | 104 | 12477663 | 12477766 | 1.36e-29 | 143 |
| Clupea_harengus-NW_024879731.1 | 86.154 | 130 | 17 | 1 | 624 | 752 | 516 | 645 | 1.76e-28 | 139 |
| Clupea_harengus-NW_024879595.1 | 86.885 | 122 | 15 | 1 | 624 | 744 | 44590 | 44711 | 2.27e-27 | 135 |
| Clupea_harengus-NW_024880355.1 | 89.320 | 103 | 11 | 0 | 1 | 103 | 21839 | 21941 | 1.06e-25 | 130 |
| Clupea_harengus-NC_045159.1 | 89.216 | 102 | 10 | 1 | 10 | 110 | 4361260 | 4361159 | 1.37e-24 | 126 |
| Clupea_harengus-NC_045159.1 | 87.736 | 106 | 13 | 0 | 10 | 115 | 4115859 | 4115754 | 4.92e-24 | 124 |
| Clupea_harengus-NC_045155.1 | 86.325 | 117 | 14 | 1 | 1 | 117 | 9964675 | 9964561 | 1.37e-24 | 126 |
| Clupea_harengus-NC_045166.1 | 87.037 | 108 | 14 | 0 | 1 | 108 | 16110188 | 16110081 | 1.77e-23 | 122 |
| Clupea_harengus-NC_045162.1 | 87.156 | 109 | 13 | 1 | 1 | 108 | 7566339 | 7566231 | 1.77e-23 | 122 |
| Clupea_harengus-NC_045165.1 | 86.239 | 109 | 12 | 2 | 1 | 106 | 4696074 | 4695966 | 2.96e-21 | 115 |
| Clupea_harengus-NC_045157.1 | 90.000 | 90 | 8 | 1 | 15 | 103 | 12746054 | 12746143 | 2.96e-21 | 115 |
| Clupea_harengus-NC_045156.1 | 86.667 | 105 | 13 | 1 | 10 | 113 | 6523649 | 6523545 | 2.96e-21 | 115 |

N5.HOG0008885

In *O. eperlanus*, the N5.HOG0008885 region does not exist

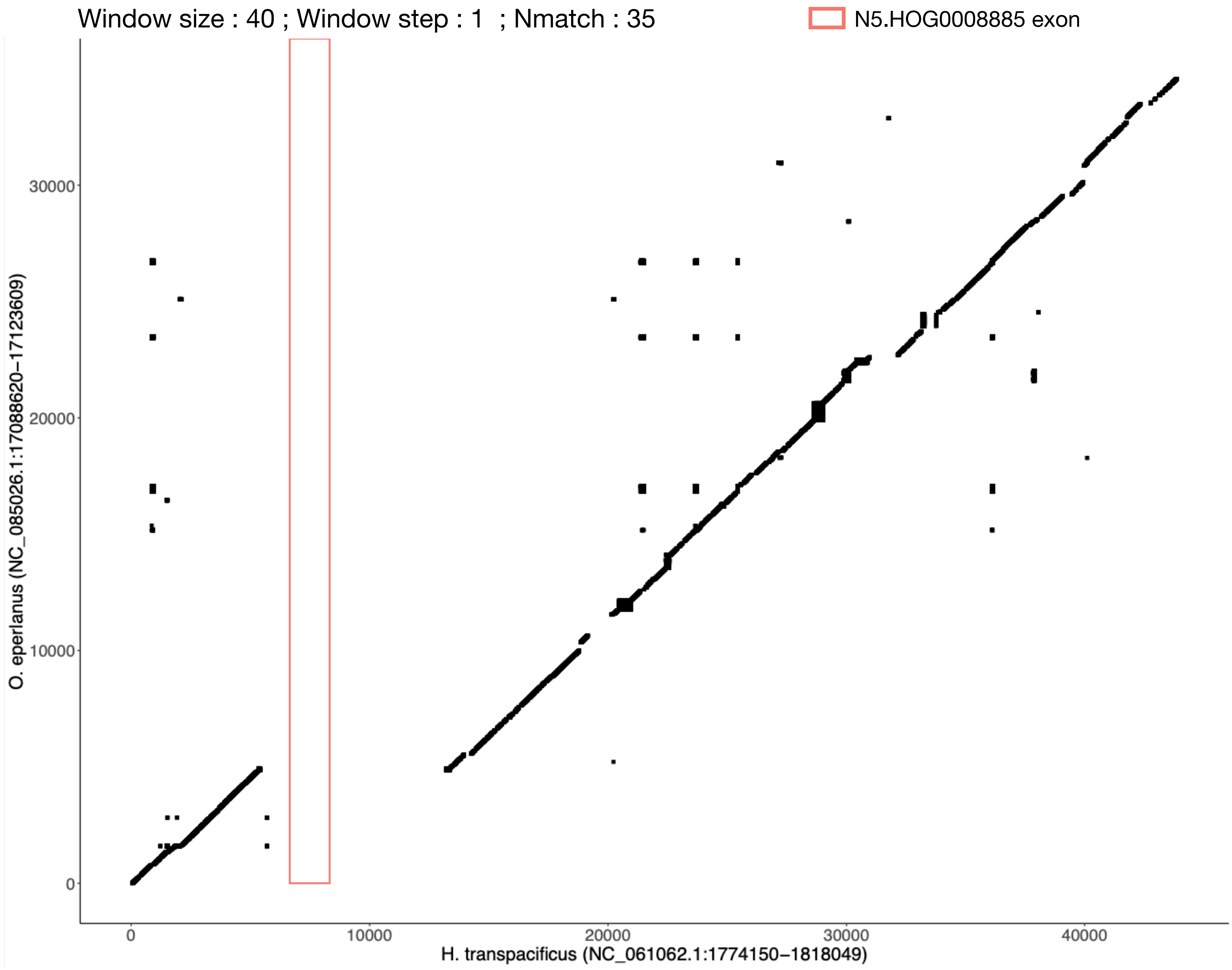

N5.HOG0008885

The gene is found in an alernative assembly of *H. transpacificus*

(CM038948.1-2566341-2564674 in GCA_021870715.1)

Alignment between the two gene versions : 99.8% similarity

```
  1 MGRTTILWFLLLLSLSVPSVSSIKTLNSIRELADNNIQFGKTFAGHGLKL     50
    ||||||||||||||||||||||||||||||||||||||||||||||||||
  1 MGRTTILWFLLLLSLSVPSVSSIKTLNSIRELADNNIQFGKTFAGHGLKL     50

 51 LYWLAHNISIDQNDVIDLGYINPSQGHYGFHYFGNRDQQGLYIFPQANNI    100
    ||||||||||||||||||||||||||||||||||||||||||||||||||
 51 LYWLAHNISIDQNDVIDLGYINPSQGHYGFHYFGNRDQQGLYIFPQANNI    100

101 HYYALGNLNGNIHPGSNTLPSYVTEDFTNTRWDTTHRNDERIIVQSYRYN    150
    ||||||||||||||||||||||||||||||||||||||||||||||||||
101 HYYALGNLNGNIHPGSNTLPSYVTEDFTNTRWDTTHRNDERIIVQSYRYN    150

151 PRRVRRVYASLHYDPDRTWEISPQLLREIRNIPTLGAFLRQVDYDFNSRN    200
    ||||||||||||||||||||||||||||||||||||||||||||||||||
151 PRRVRRVYASLHYDPDRTWEISPQLLREIRNIPTLGAFLRQVDYDFNSRN    200

201 NFYGKIFHCTQSHDLRRRRSLPECDTSEGLALDMKPVYSNANARITWSGI    250
    ||||||||||||||||||||||||||||||||||||||||||||||||||
201 NFYGKIFHCTQSHDLRRRRSLPECDTSEGLALDMKPVYSNANARITWSGI    250

251 PKTILTSNLFLNVYANMDSSTALESYDVGWQKSGSRDTSVLLHPGLQVRL    300
    |||||||||||||||||||||||||||||||||||||||||||||||:||
251 PKTILTSNLFLNVYANMDSSTALESYDVGWQKSGSRDTSVLLHPGLQIRL    300

301 MKKTGVWPWAQFTQIWAGPEFDEGNRKLPTDVPGYDTSLLLFVKQGKACV    350
    ||||||||||||||||||||||||||||||||||||||||||||||||||
301 MKKTGVWPWAQFTQIWAGPEFDEGNRKLPTDVPGYDTSLLLFVKQGKACV    350

351 RLYIKKSFTDWKNTFKYSWVGFYLTRDIGHSIYVAWQWATKFYENQNKHT    400
    |||||||||||||||||||||||||||||||.||||||||||||||||||
351 RLYIKKSFTDWKNTFKYSWVGFYLTRDIGHSSYVAWQWATKFYENQNKHT    400

401 DEYLAYEYDFGLDIHRAEQVRFFLEKAPSSVRAQAVPWGQEAVEKIQHNV    450
    ||||||||||||||||||||||||||||||||||||||||||||||||||
401 DEYLAYEYDFGLDIHRAEQVRFFLEKAPSSVRAQAVPWGQEAVEKIQHNV    450

451 RGYEARLQPIAIGGKVWARLYIKKSFTDWKDQFEYAWVGFYSSNSALSYS    500
    ||||||||||||||||||||||||||||||||||||||||||||||||||
451 RGYEARLQPIAIGGKVWARLYIKKSFTDWKDQFEYAWVGFYSSNSALSYS    500

501 YDTWQSVIKFNRNEDQDTEEYMAYDYNSNMDVRQGMETRFLMTKGYDQEK    550
    ||||||||||||||||||||||||||||||||||||||||||||||||||
501 YDTWQSVIKFNRNEDQDTEEYMAYDYNSNMDVRQGMETRFLMTKGYDQEK    550

551 CRARL    555
    |||||
551 CRARL    555
```

N5.HOG0008885

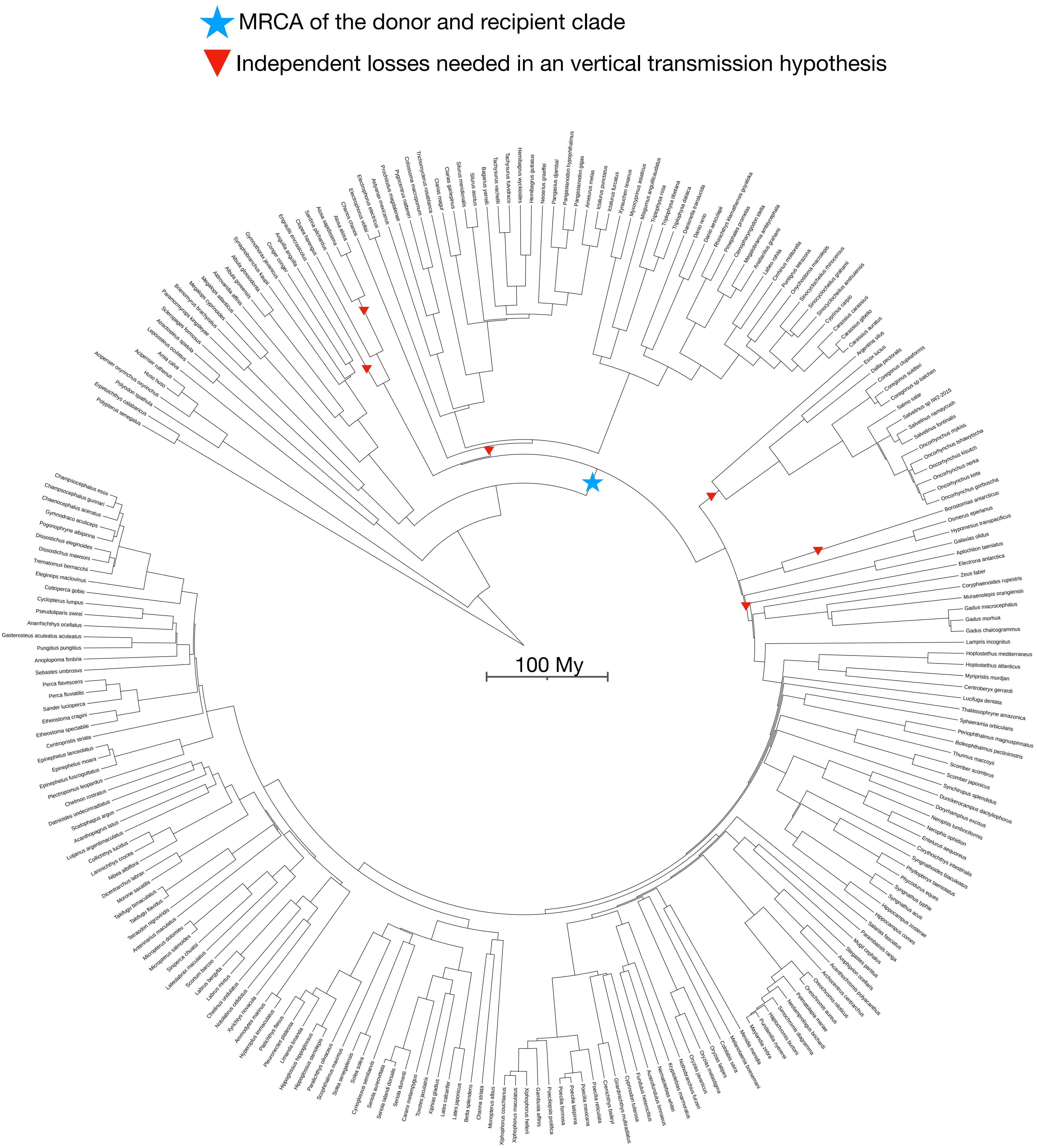

N5.HOG0004480

### N5.HOG0004480

Species tree

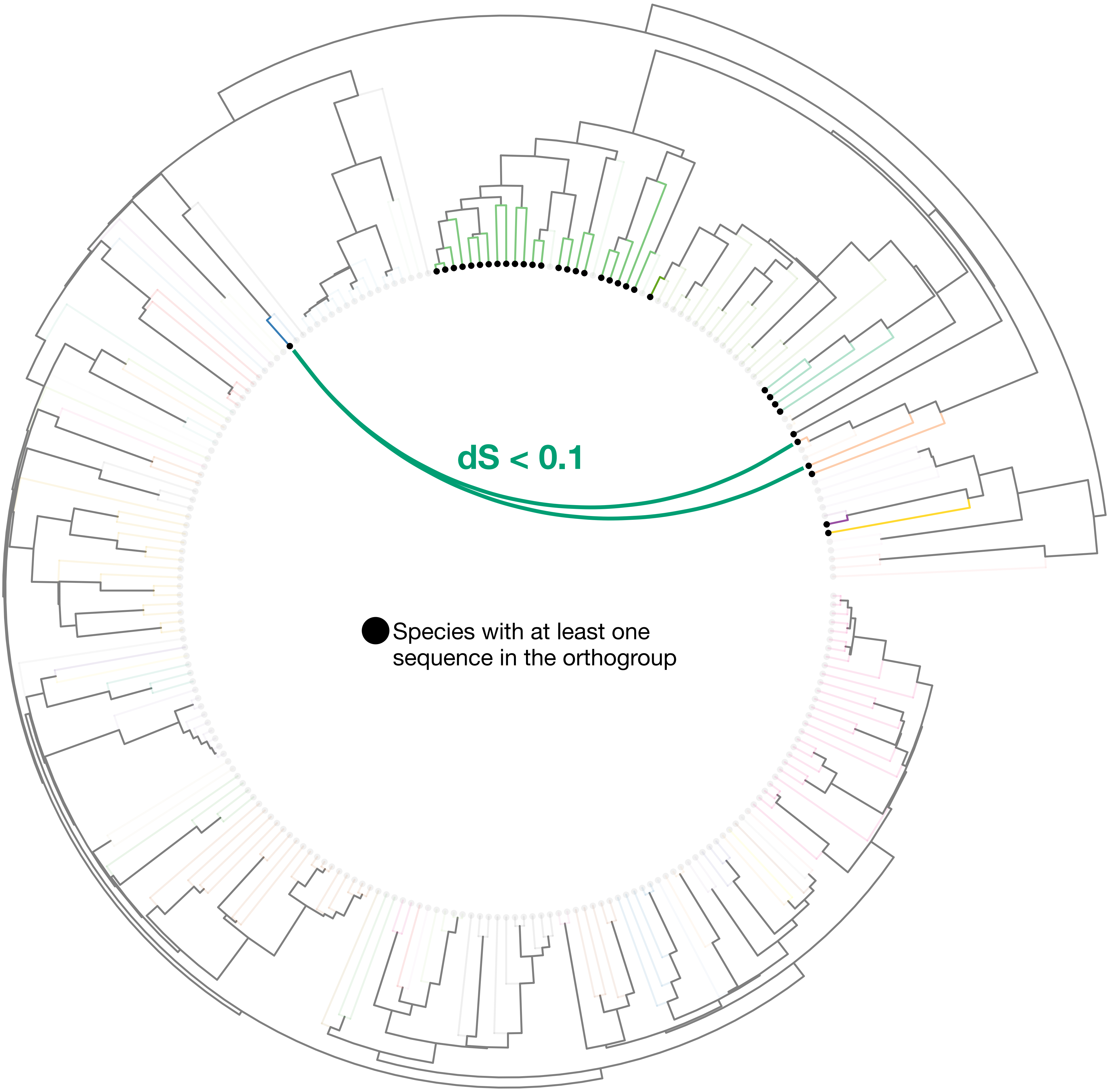

### N5.HOG0004480

Orthogroup phylogeny
and
protein alignment

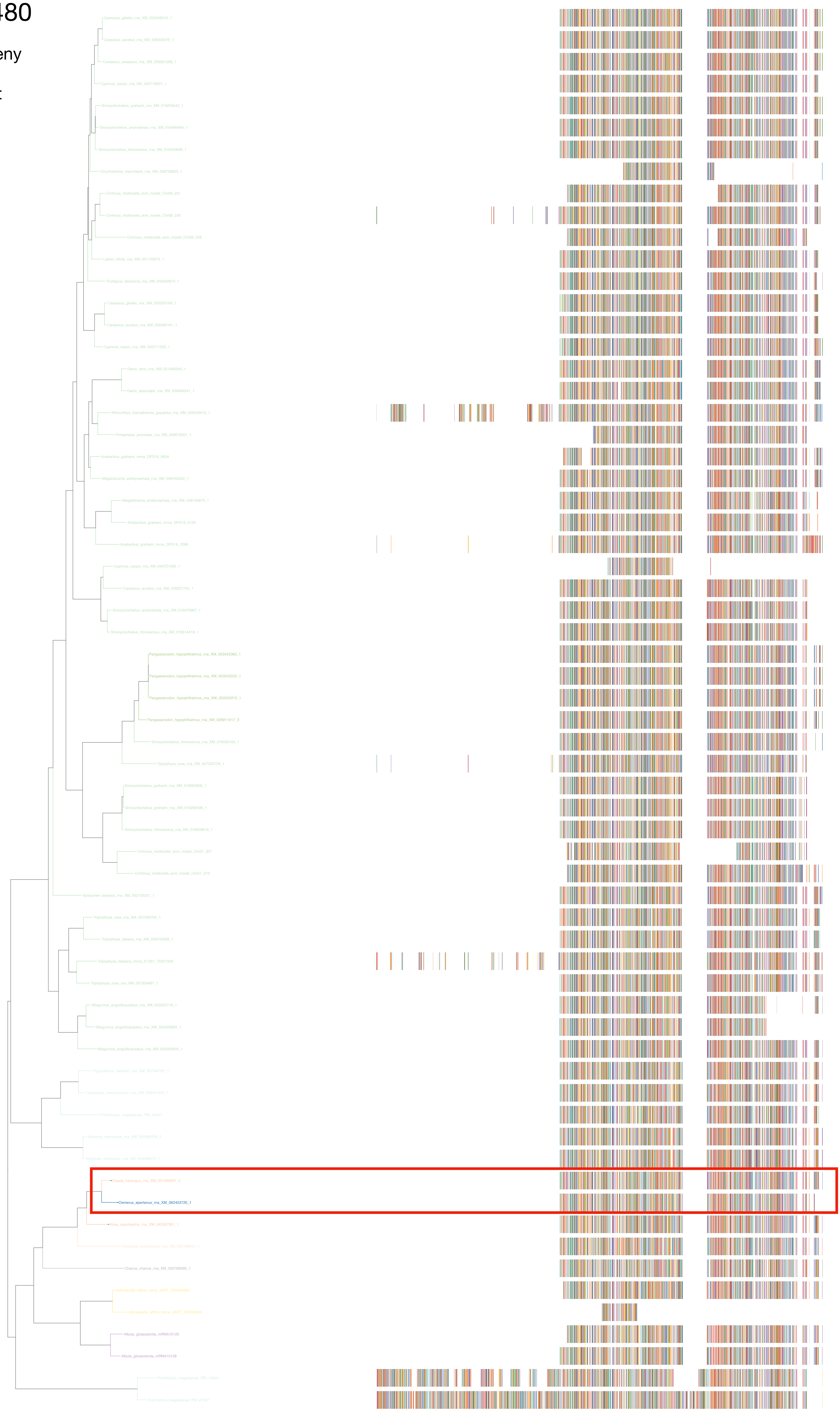

### N5.HOG0004480

Zoom on HGT clade with bootstrap values

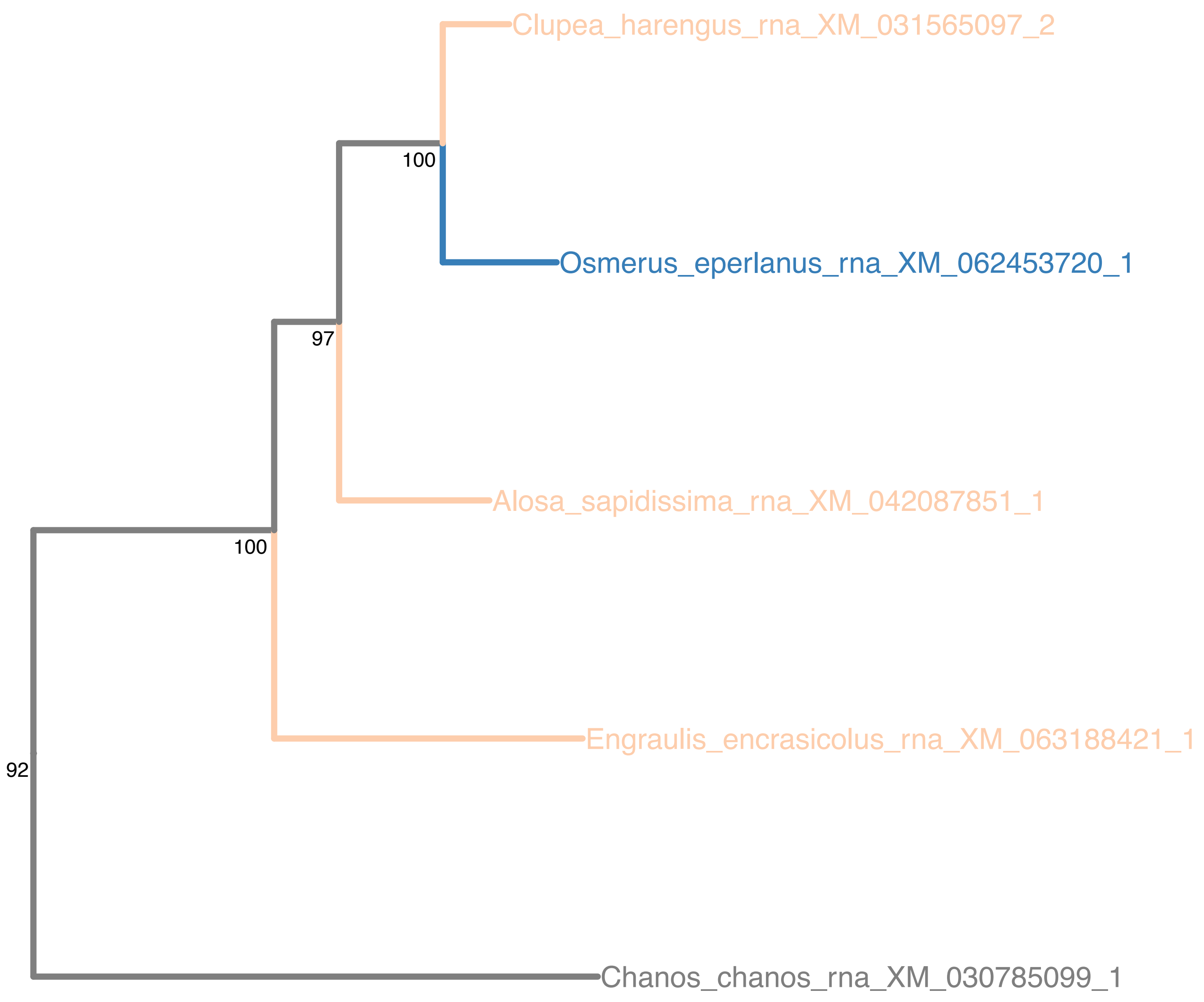

### N5.HOG0004480

Orthogroup phylogeny
and
trimmed
protein alignment

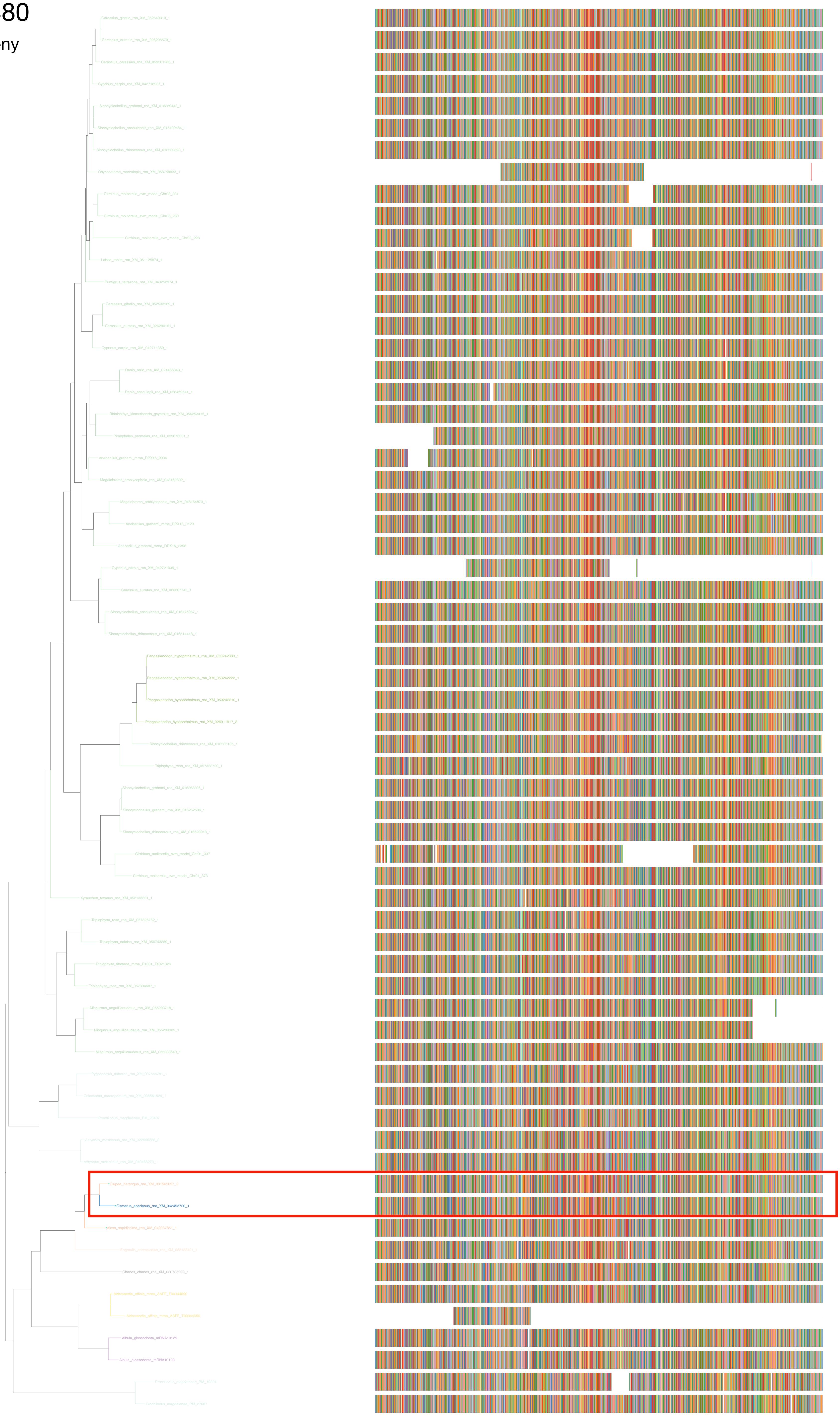

### N5.HOG0004480

« Best-match » phylogeny
and
protein alignment

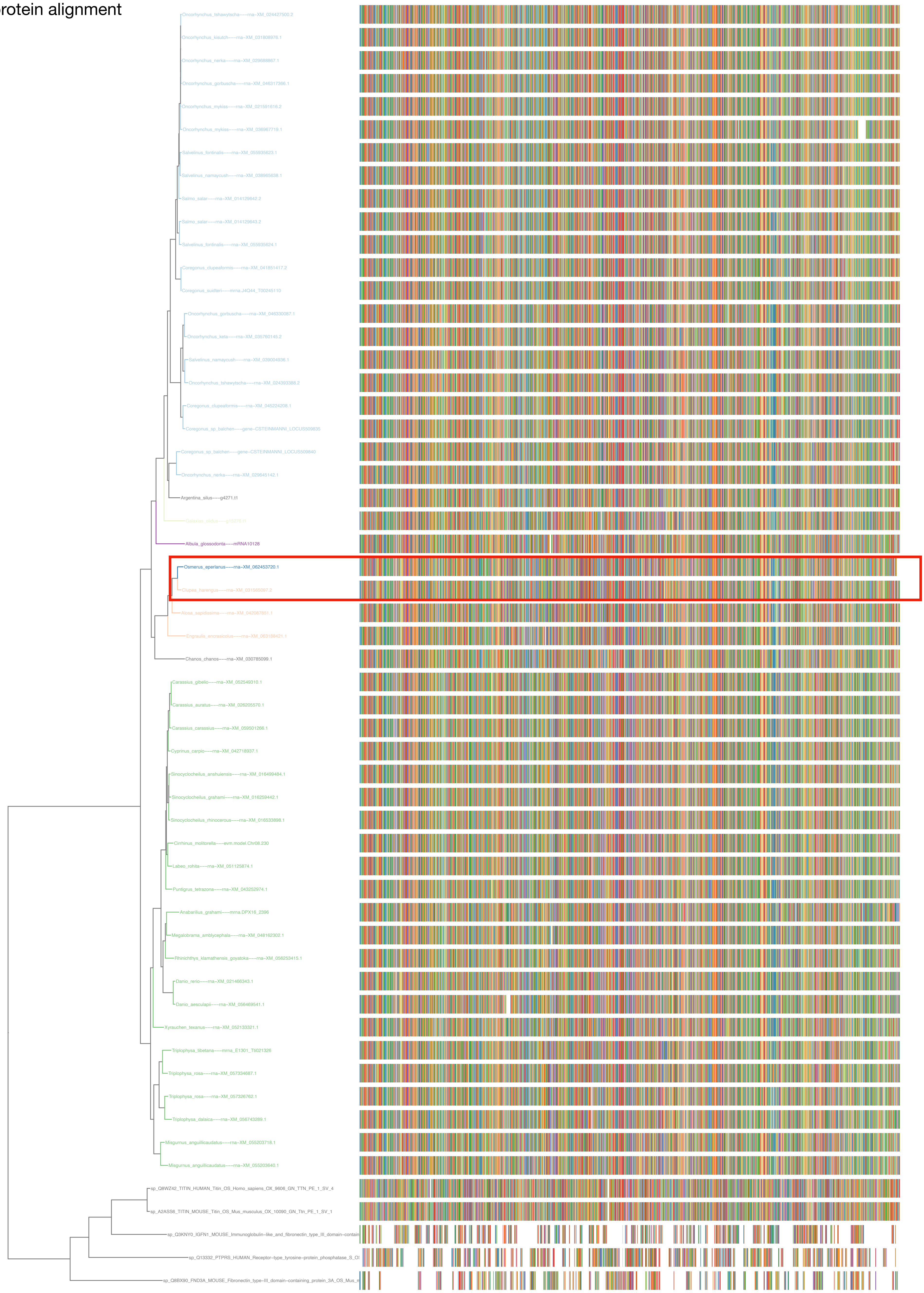

N5.HOG0004480

dS distribution

Inter-species dS distribution

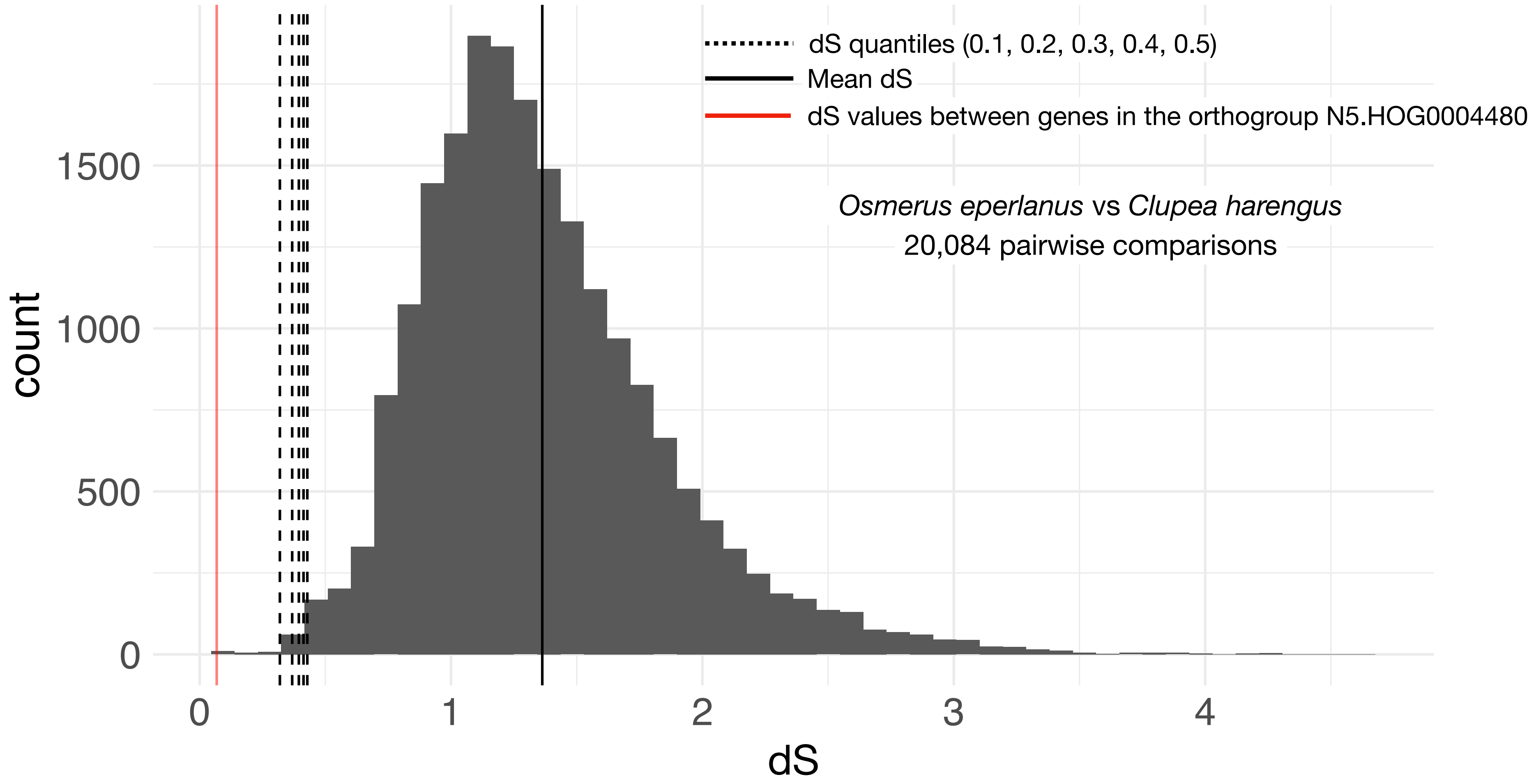


Intra-orthogroup dS

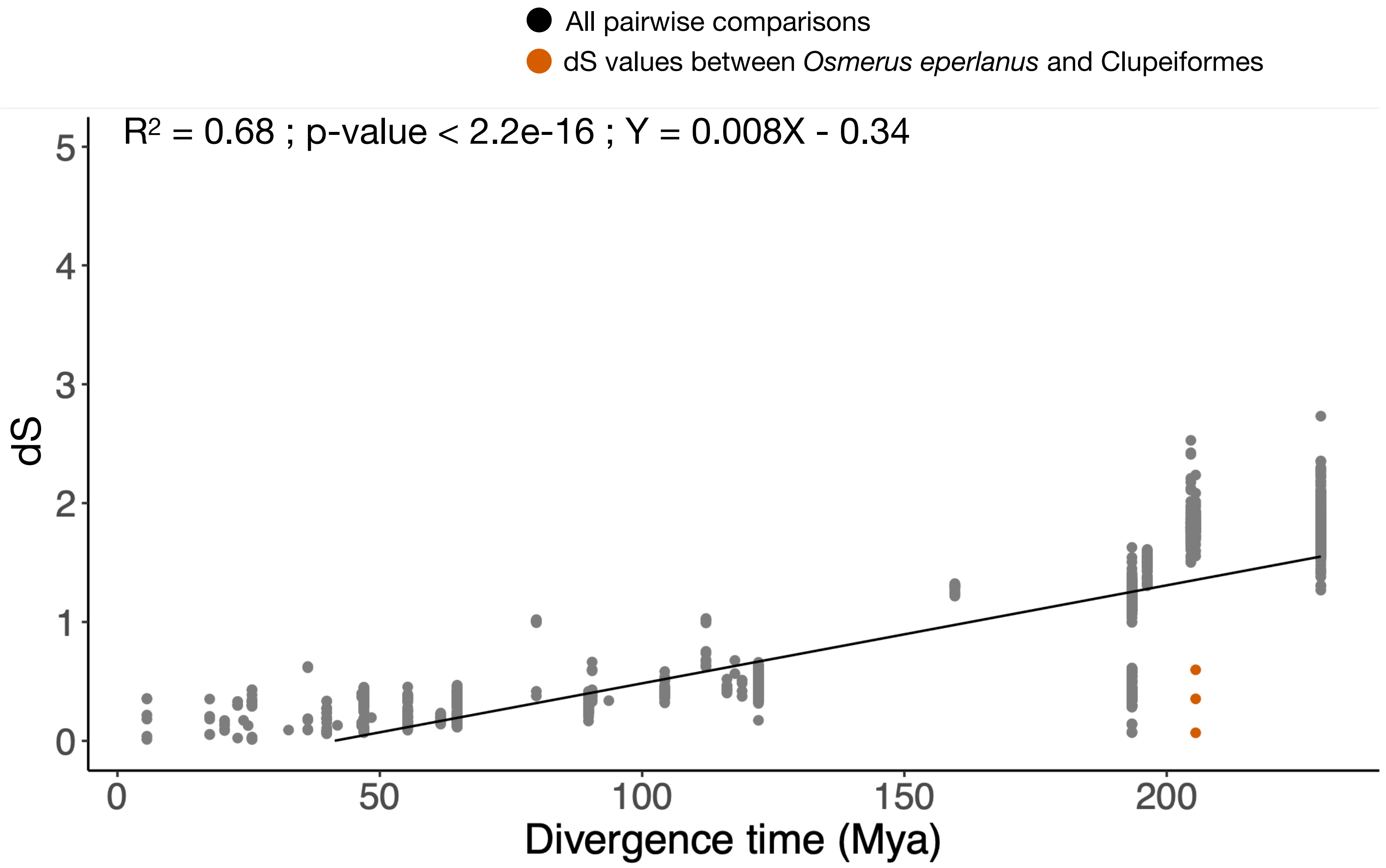

### N5.HOG0004480

#### Micro-synteny

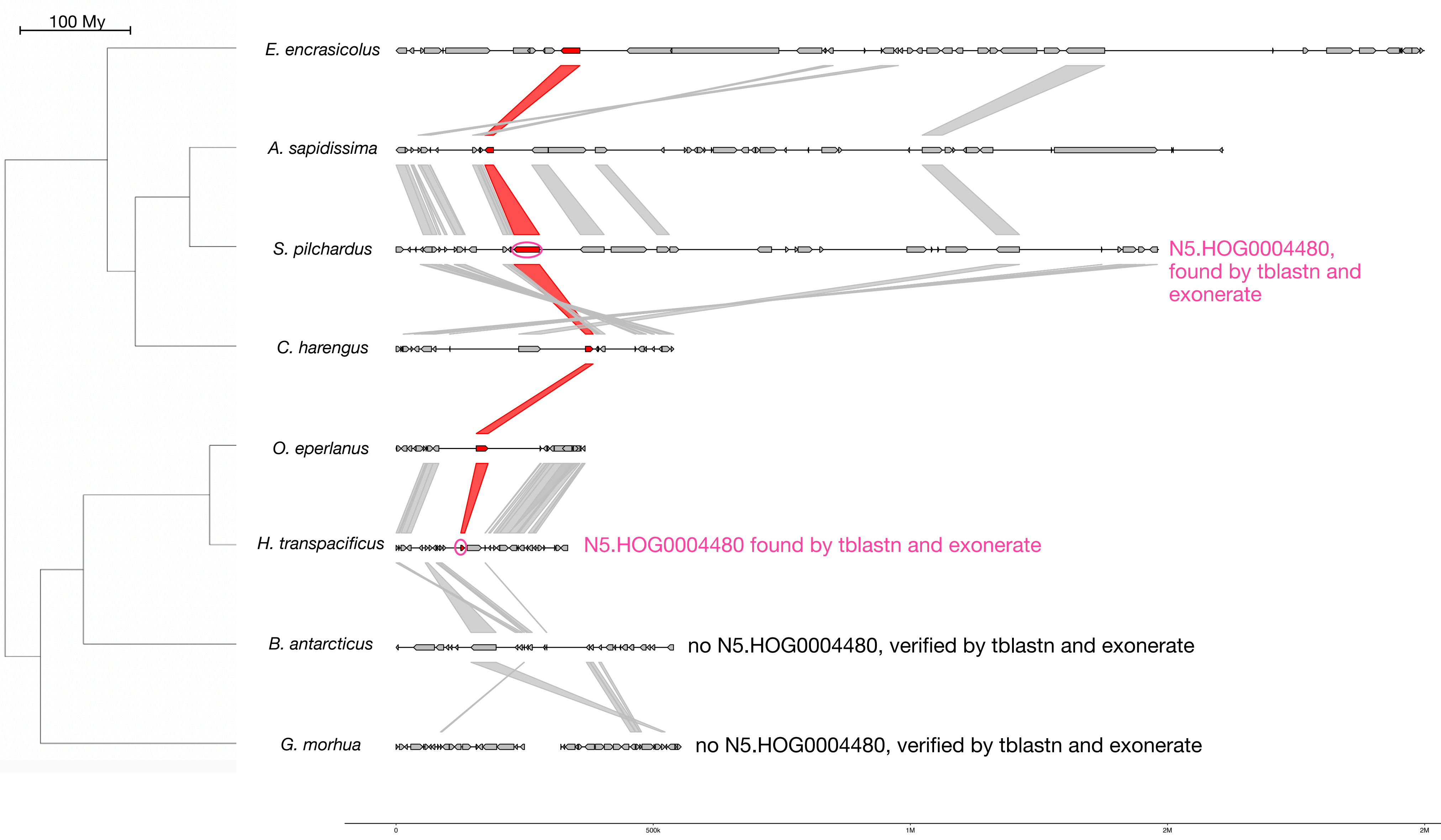


#### Zoom on *C. harengus* and *O. eperlanus*

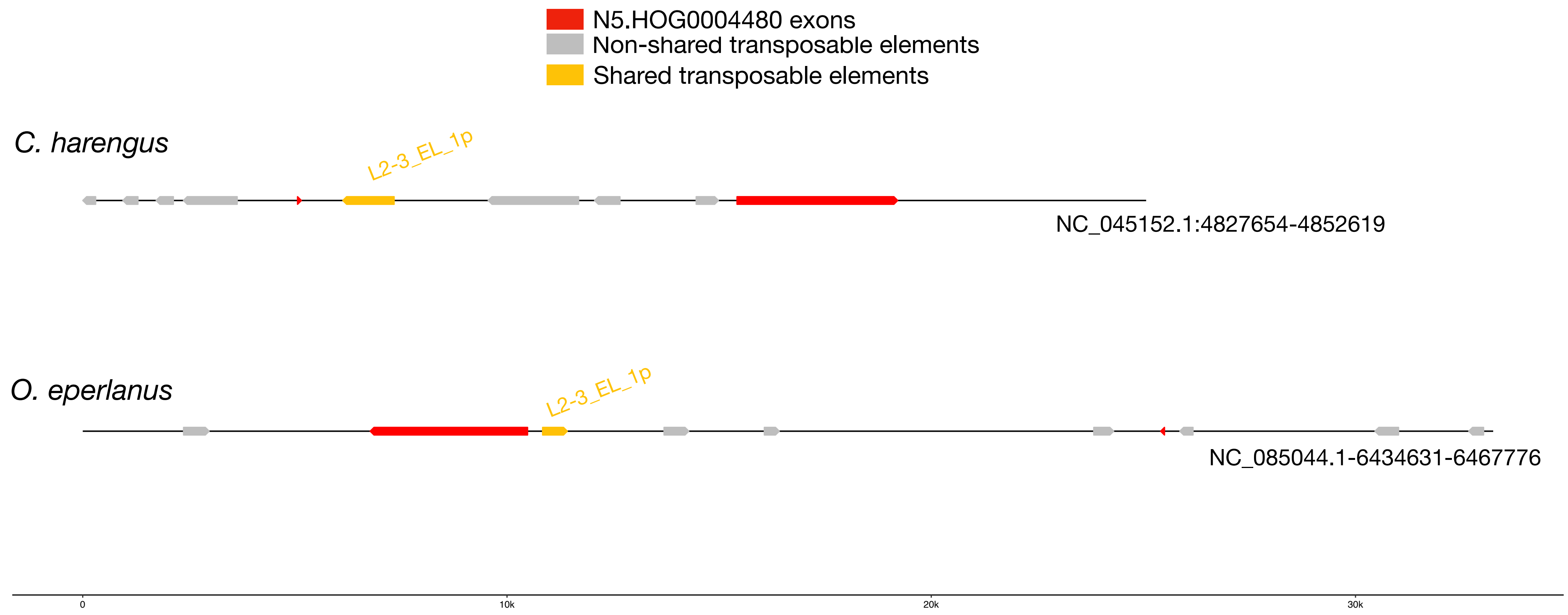


Both have the same TE in the intron but these TE shows no strong identity

## L2-3_EL_1p :

| | |
|---|---|
| *E. encrasicolus* | *10 copies scattered in the genome* |
| *A. sapidissima* | *252 copies scattered in the genome* |
| *S. pilchardus* | *85 copies scattered in the genome* |
| *C. harengus* | *42 copies scattered in the genome* |
| *H. transpacificus* | *25 copies scattered in the genome* |
| *O. eperlanus* | *33 copies scattered in the genome* |
| *B. antarcticus* | *172 copies scattered in the genome* |
| *G. morhua* | *28 copies scattered in the genome* |

### N5.HOG0004480

N5.HOG0004480 is present in H. transpacificus and S. pilchardus.
Full length sequence extracted by Exonerate

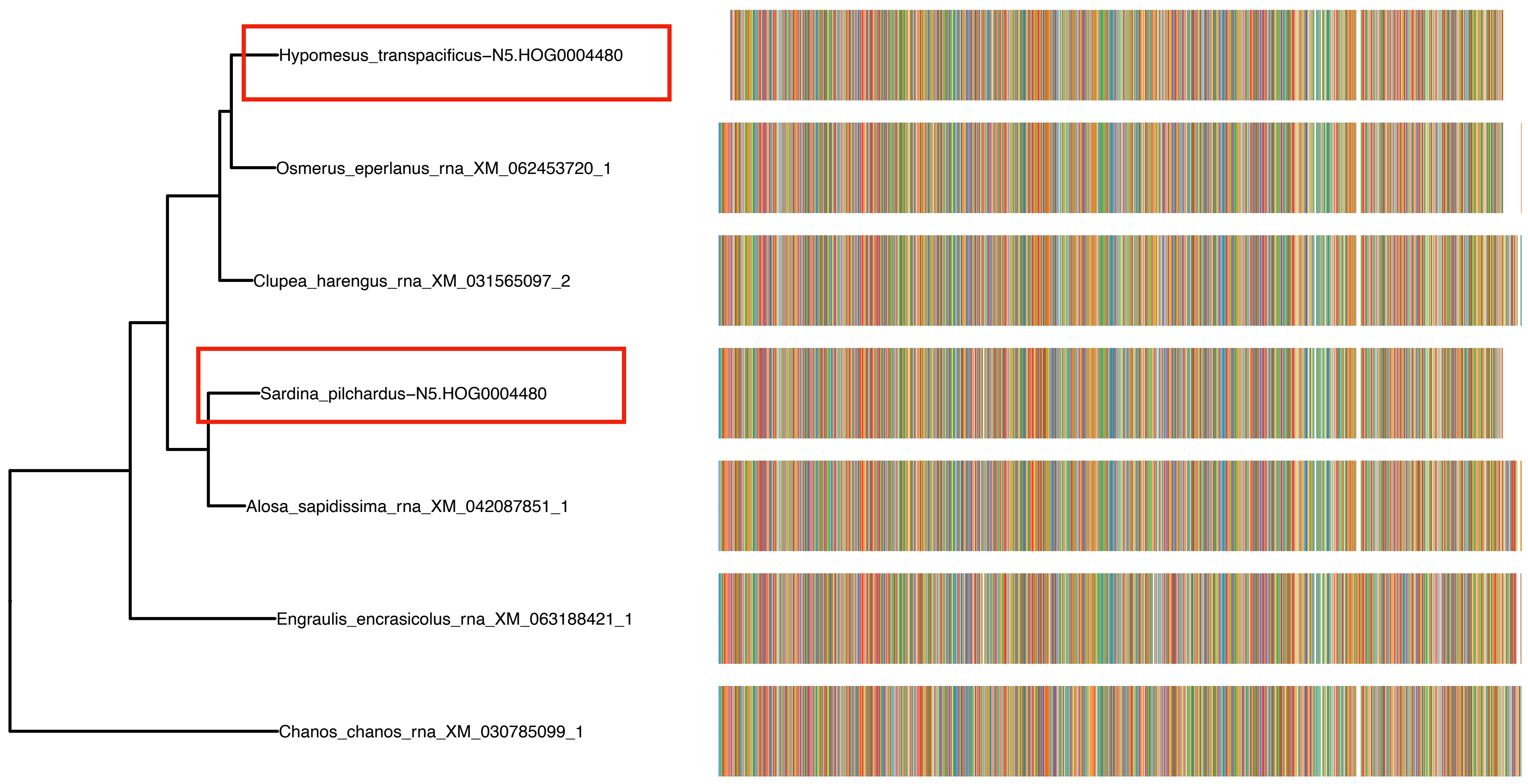

N5.HOG0004480

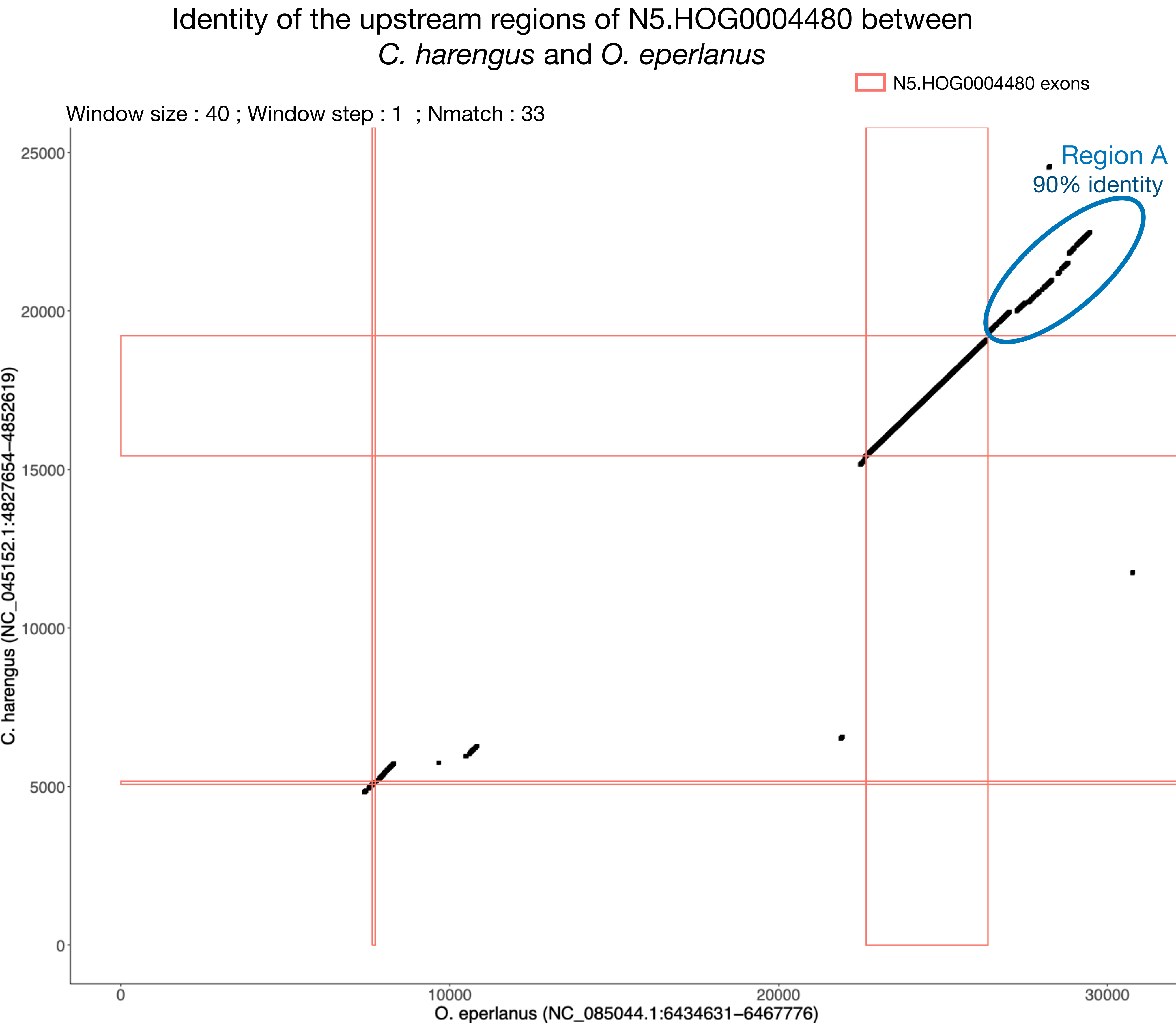

### N5.HOG0004480 Blastn of conserved regions against all genome assemblies

#### Region A

| sseqid | pident | length | mismatch | gapopen | qstart | qend | sstart | send | evalue | bitscore |
|---|---|---|---|---|---|---|---|---|---|---|
| Osmerus_eperlanus-NC_085044.1 | 100.000 | 3530 | 0 | 0 | 1 | 3530 | 6441406 | 6437877 | 0.0 | 6519 |
| Osmerus_eperlanus-NC_085044.1 | 95.187 | 187 | 6 | 1 | 688 | 871 | 611628 | 611442 | 5.81e-74 | 292 |
| Osmerus_eperlanus-NC_085044.1 | 94.652 | 187 | 7 | 1 | 688 | 871 | 7472004 | 7471818 | 2.70e-72 | 287 |
| Osmerus_eperlanus-NC_085044.1 | 94.118 | 187 | 8 | 1 | 688 | 871 | 3832120 | 3832306 | 1.26e-70 | 281 |
| Osmerus_eperlanus-NC_085044.1 | 94.054 | 185 | 8 | 1 | 688 | 869 | 7052671 | 7052855 | 1.63e-69 | 278 |
| Osmerus_eperlanus-NC_085044.1 | 92.746 | 193 | 11 | 1 | 682 | 871 | 3086388 | 3086580 | 5.85e-69 | 276 |
| Osmerus_eperlanus-NC_085044.1 | 92.308 | 195 | 12 | 1 | 681 | 872 | 2113911 | 2114105 | 2.11e-68 | 274 |
| Osmerus_eperlanus-NC_085044.1 | 91.584 | 202 | 11 | 4 | 680 | 875 | 4087083 | 4086882 | 2.11e-68 | 274 |
| Osmerus_eperlanus-NC_085044.1 | 92.708 | 192 | 11 | 1 | 681 | 869 | 5206336 | 5206527 | 2.11e-68 | 274 |
| Osmerus_eperlanus-NC_085044.1 | 92.670 | 191 | 10 | 2 | 688 | 875 | 1252408 | 1252219 | 7.57e-68 | 272 |
| Hypomesus_transpacificus-NC_061079.1 | 90.591 | 1201 | 83 | 15 | 2349 | 3530 | 3812098 | 3813287 | 0.0 | 1565 |
| Hypomesus_transpacificus-NC_061079.1 | 86.287 | 1371 | 84 | 43 | 873 | 2211 | 3810136 | 3811434 | 0.0 | 1395 |
| Hypomesus_transpacificus-NC_061079.1 | 94.340 | 689 | 39 | 0 | 1 | 689 | 3809450 | 3810138 | 0.0 | 1057 |
| Hypomesus_transpacificus-NC_061079.1 | 92.708 | 192 | 9 | 4 | 682 | 868 | 2881790 | 2881981 | 7.57e-68 | 272 |
| Hypomesus_transpacificus-NC_061079.1 | 92.973 | 185 | 10 | 1 | 688 | 869 | 1076065 | 1075881 | 3.52e-66 | 267 |
| Hypomesus_transpacificus-NC_061079.1 | 92.513 | 187 | 10 | 2 | 688 | 871 | 5074365 | 5074180 | 1.27e-65 | 265 |
| Hypomesus_transpacificus-NC_061079.1 | 92.473 | 186 | 10 | 2 | 688 | 870 | 3115380 | 3115196 | 4.56e-65 | 263 |
| Hypomesus_transpacificus-NC_061079.1 | 91.667 | 192 | 11 | 3 | 681 | 869 | 1561948 | 1562137 | 1.64e-64 | 261 |
| Hypomesus_transpacificus-NC_061079.1 | 91.935 | 186 | 12 | 1 | 688 | 870 | 4898898 | 4899083 | 2.12e-63 | 257 |
| Hypomesus_transpacificus-NC_061079.1 | 89.604 | 202 | 15 | 4 | 679 | 875 | 98262 | 98462 | 9.86e-62 | 252 |
| Clupea_harengus-NC_045152.1 | 91.966 | 697 | 41 | 7 | 1 | 689 | 4846968 | 4847657 | 0.0 | 963 |
| Clupea_harengus-NC_045152.1 | 87.866 | 717 | 50 | 15 | 2460 | 3141 | 4849468 | 4850182 | 0.0 | 808 |
| Clupea_harengus-NC_045152.1 | 86.269 | 772 | 52 | 20 | 1221 | 1972 | 4847923 | 4848660 | 0.0 | 789 |
| Clupea_harengus-NC_045152.1 | 91.419 | 303 | 23 | 3 | 873 | 1173 | 4847655 | 4847956 | 4.28e-110 | 412 |
| Clupea_harengus-NC_045152.1 | 88.333 | 240 | 18 | 6 | 2228 | 2467 | 4848980 | 4849209 | 4.53e-70 | 279 |
| Clupea_harengus-NC_045152.1 | 82.288 | 271 | 31 | 11 | 3149 | 3412 | 22084757 | 22085017 | 1.00e-51 | 219 |
| Clupea_harengus-NC_045152.1 | 95.294 | 85 | 2 | 1 | 2129 | 2211 | 4848834 | 4848918 | 3.73e-26 | 134 |
| Clupea_harengus-NW_024880352.1 | 88.542 | 384 | 27 | 9 | 3142 | 3519 | 21790 | 22162 | 3.26e-121 | 449 |
| Clupea_harengus-NC_045153.1 | 86.260 | 393 | 41 | 6 | 3141 | 3530 | 4979591 | 4979209 | 1.19e-110 | 414 |
| Clupea_harengus-NC_045153.1 | 85.459 | 392 | 45 | 5 | 3141 | 3530 | 5026178 | 5025797 | 1.20e-105 | 398 |
| Clupea_harengus-NC_045153.1 | 75.070 | 357 | 70 | 16 | 3142 | 3489 | 11665397 | 11665743 | 1.33e-30 | 148 |
| Clupea_harengus-NC_045160.1 | 86.375 | 389 | 25 | 6 | 3142 | 3530 | 9080502 | 9080862 | 3.33e-106 | 399 |
| Clupea_harengus-NC_045160.1 | 76.149 | 348 | 60 | 20 | 3141 | 3477 | 9061679 | 9062014 | 1.71e-34 | 161 |
| Clupea_harengus-NC_045156.1 | 85.128 | 390 | 39 | 9 | 3133 | 3519 | 13123277 | 13123650 | 1.21e-100 | 381 |
| Clupea_harengus-NC_045168.1 | 84.439 | 392 | 49 | 5 | 3142 | 3530 | 16642471 | 16642853 | 5.61e-99 | 375 |
| Clupea_harengus-NC_045168.1 | 77.099 | 393 | 57 | 25 | 3142 | 3527 | 17607696 | 17607330 | 4.69e-45 | 196 |
| Clupea_harengus-NC_045166.1 | 83.210 | 405 | 53 | 12 | 3130 | 3527 | 10987541 | 10987145 | 2.03e-93 | 357 |
| Clupea_harengus-NC_045166.1 | 82.908 | 392 | 53 | 7 | 3142 | 3530 | 27243063 | 27243443 | 2.05e-88 | 340 |
| Clupea_harengus-NC_045157.1 | 83.503 | 394 | 51 | 7 | 3139 | 3527 | 31075142 | 31075526 | 7.31e-93 | 355 |
| Clupea_harengus-NC_045157.1 | 79.444 | 360 | 47 | 12 | 3142 | 3498 | 23384059 | 23384394 | 4.62e-55 | 230 |
| Clupea_harengus-NC_045170.1 | 85.593 | 236 | 20 | 7 | 3138 | 3369 | 13389790 | 13389565 | 9.93e-57 | 235 |

N5.HOG0004480

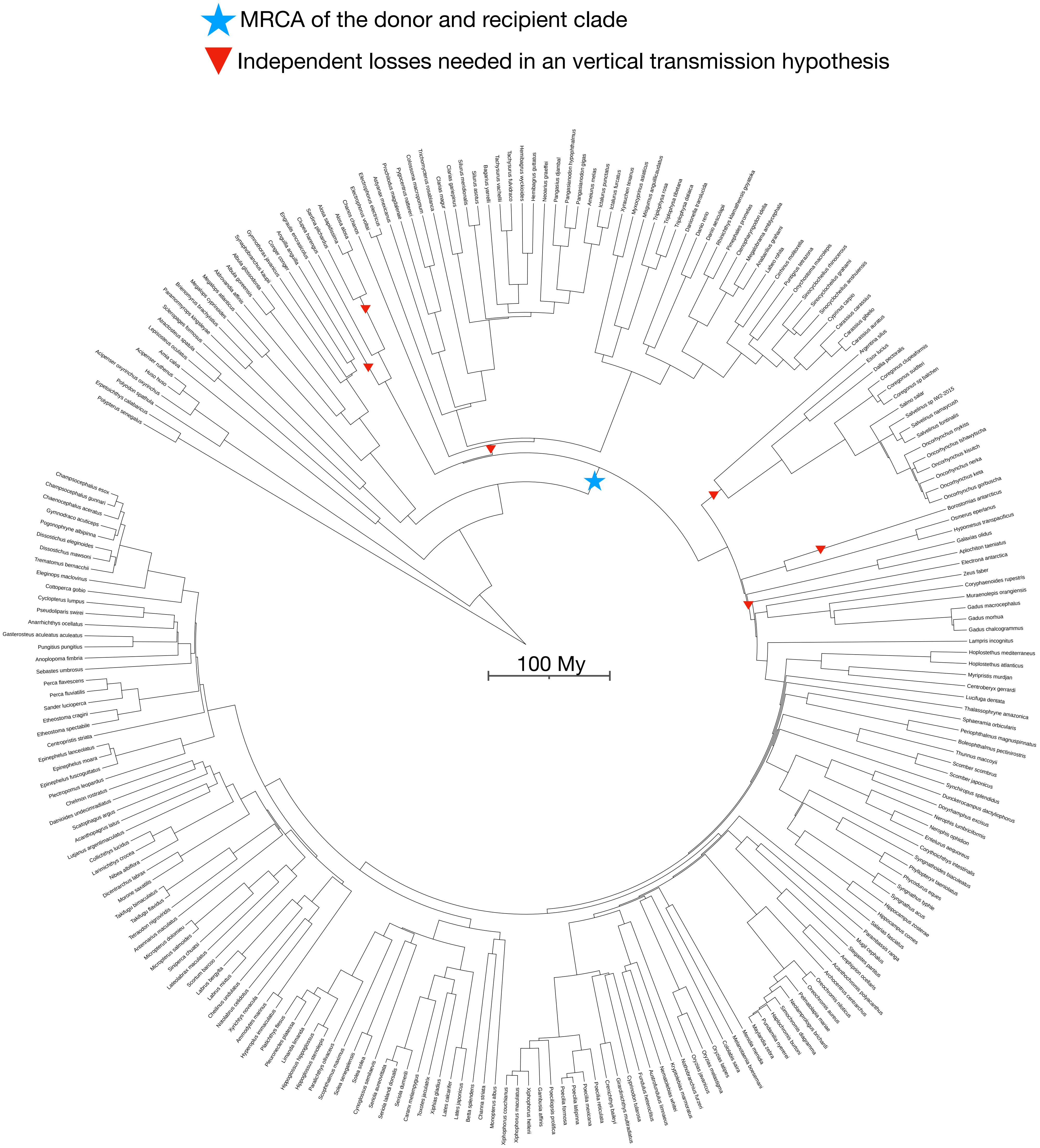

N5.HOG0028899

N5.HOG0028899

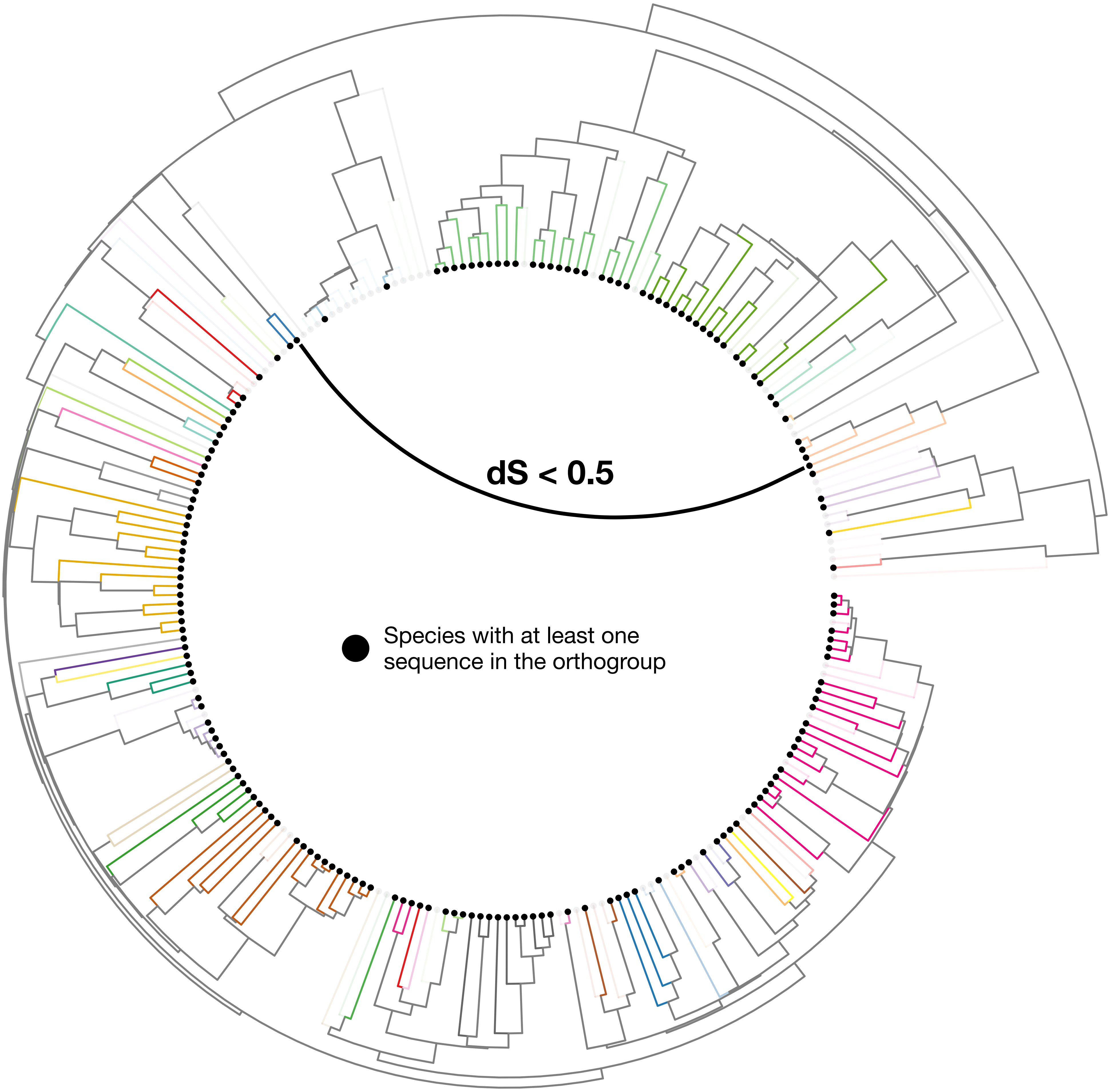

### N5.HOG0028899

Orthogroup phylogeny and protein alignment

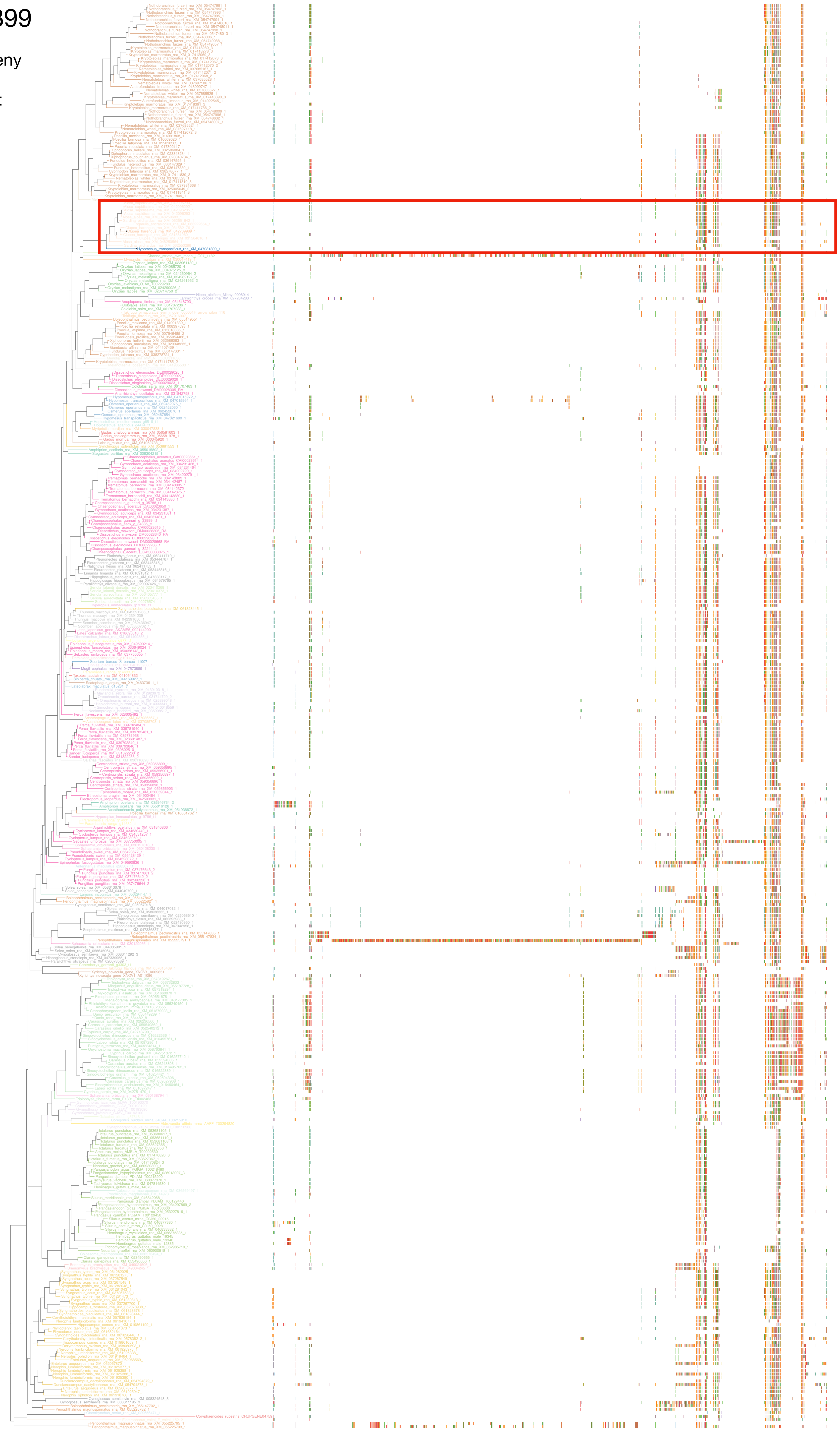

### N5.HOG0028899

Zoom on HGT clade with bootstrap values

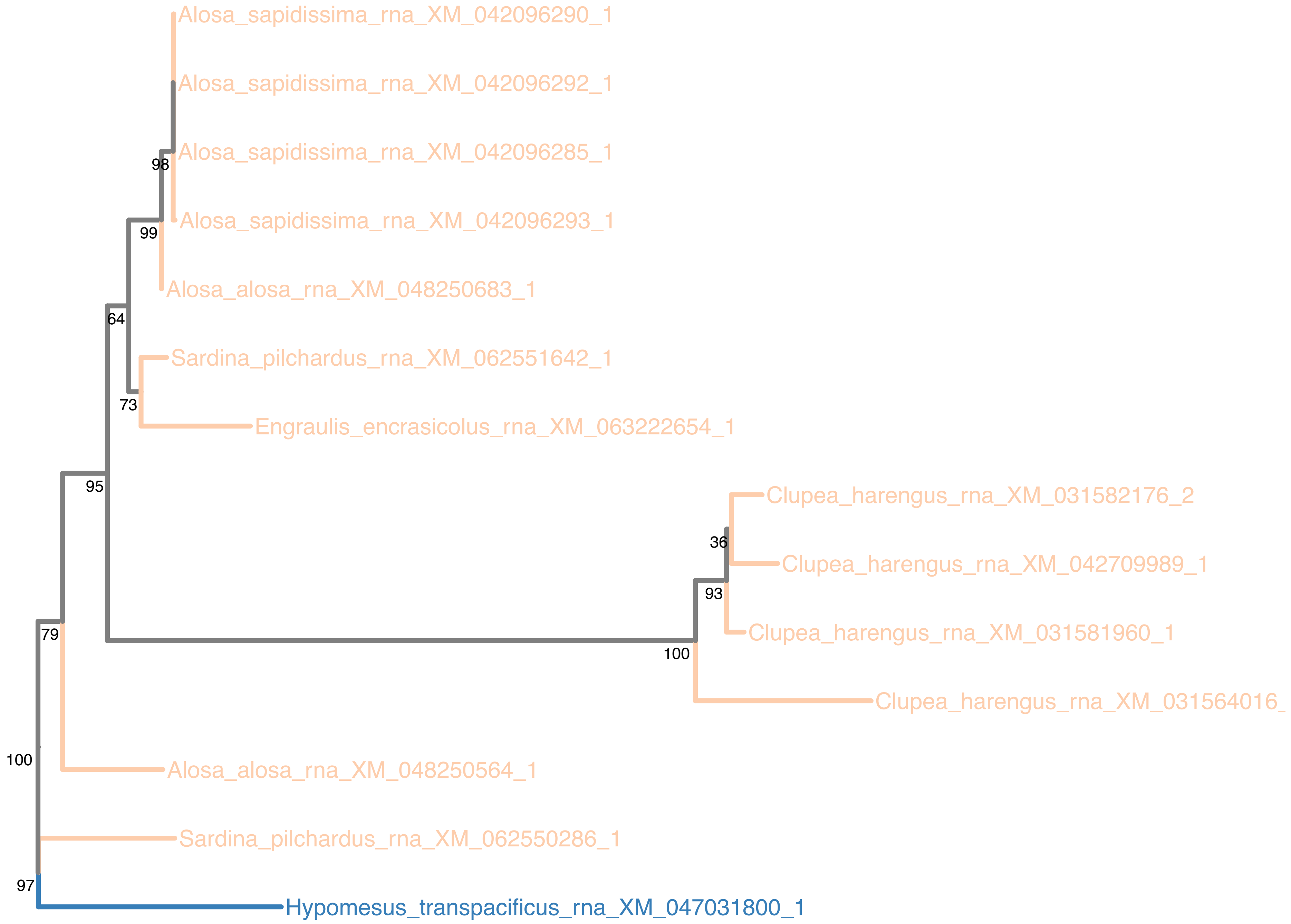

### N5.HOG0028899

Orthogroup phylogeny and trimmed protein alignment

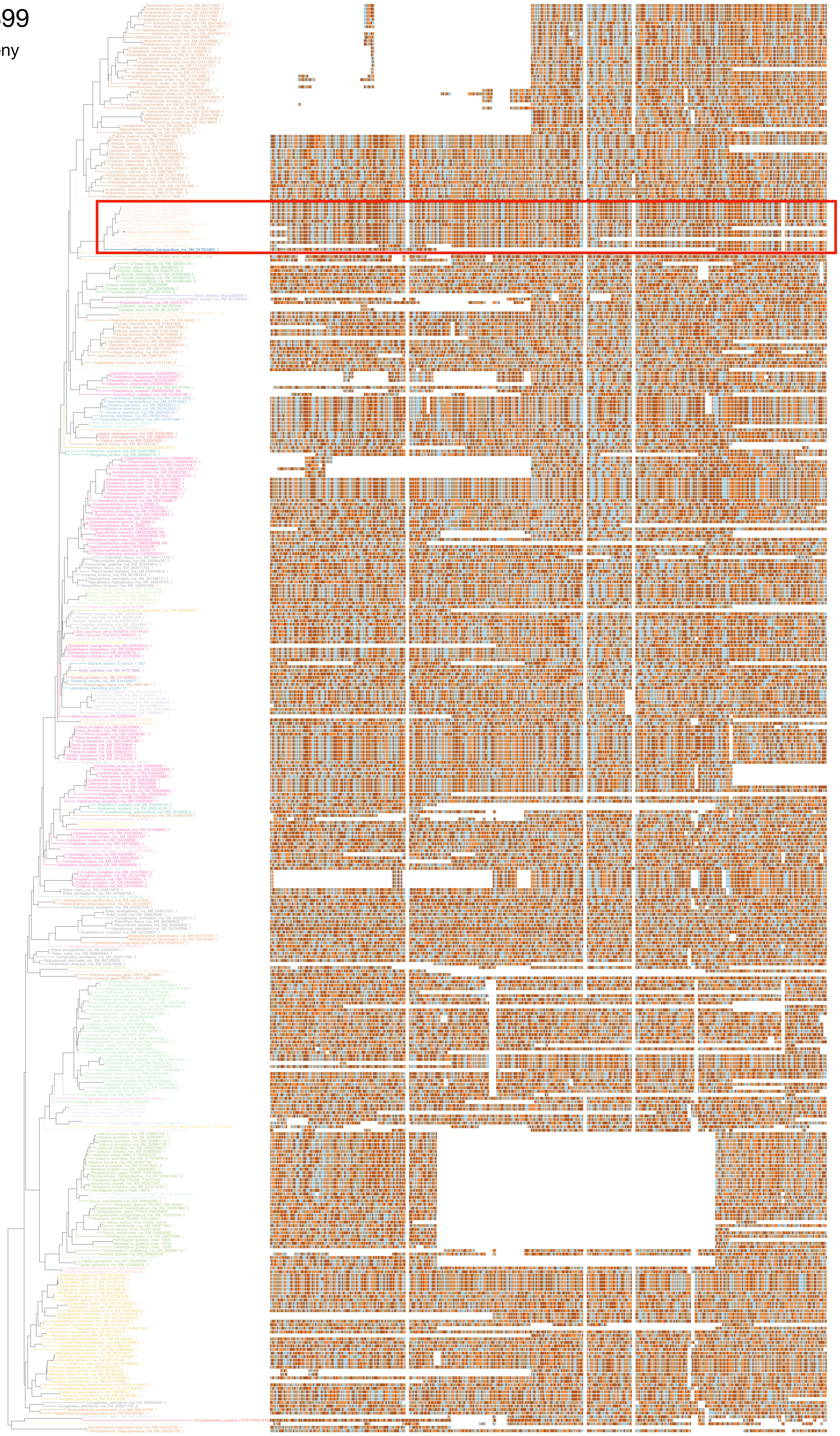

### N5.HOG0028899

« Best-match » phylogeny
and
protein alignment

50 best match against ray-finned fishes protein databases
10 best match against non-fish proteins from UniProt

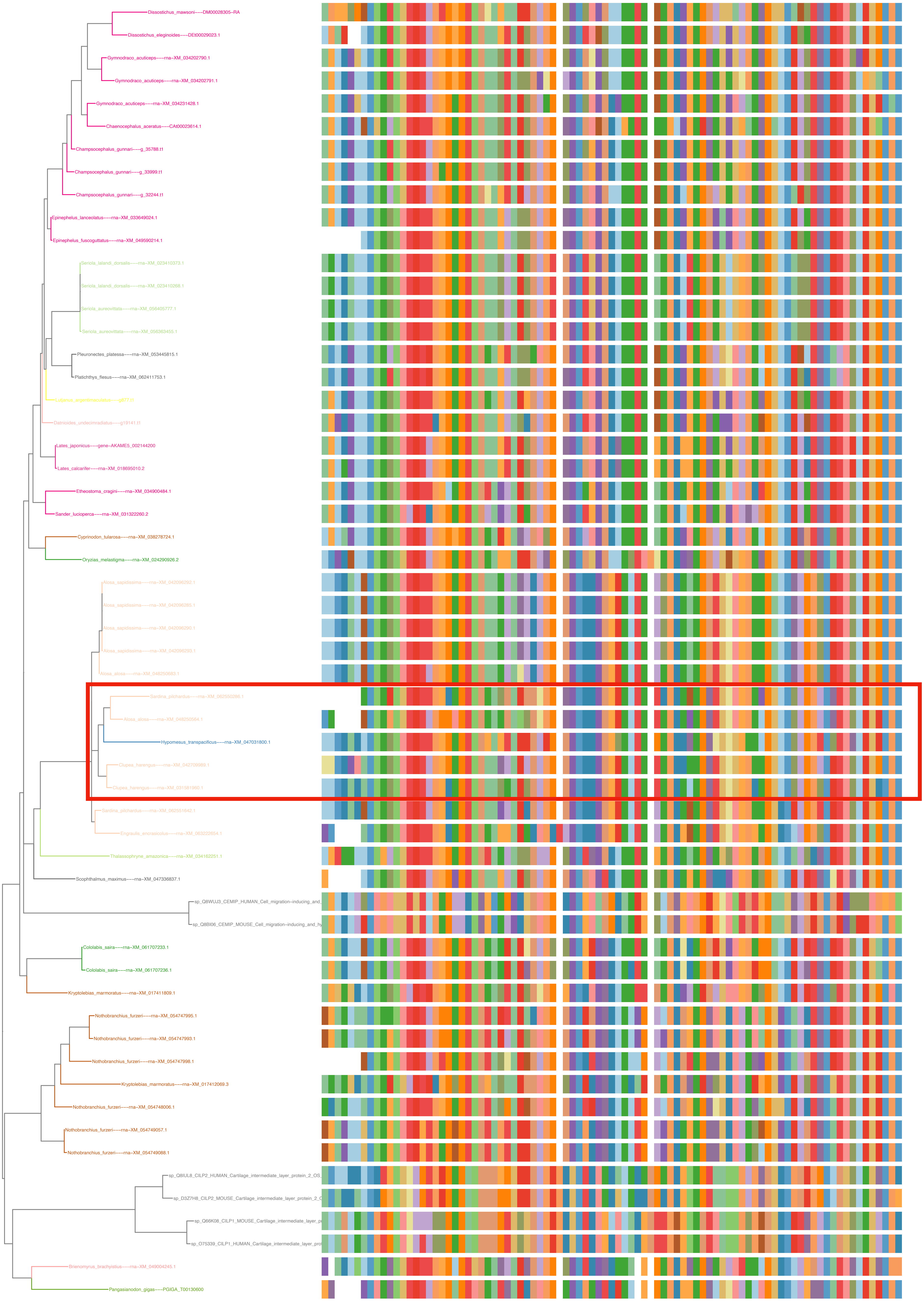

### N5.HOG0028899

dS distribution

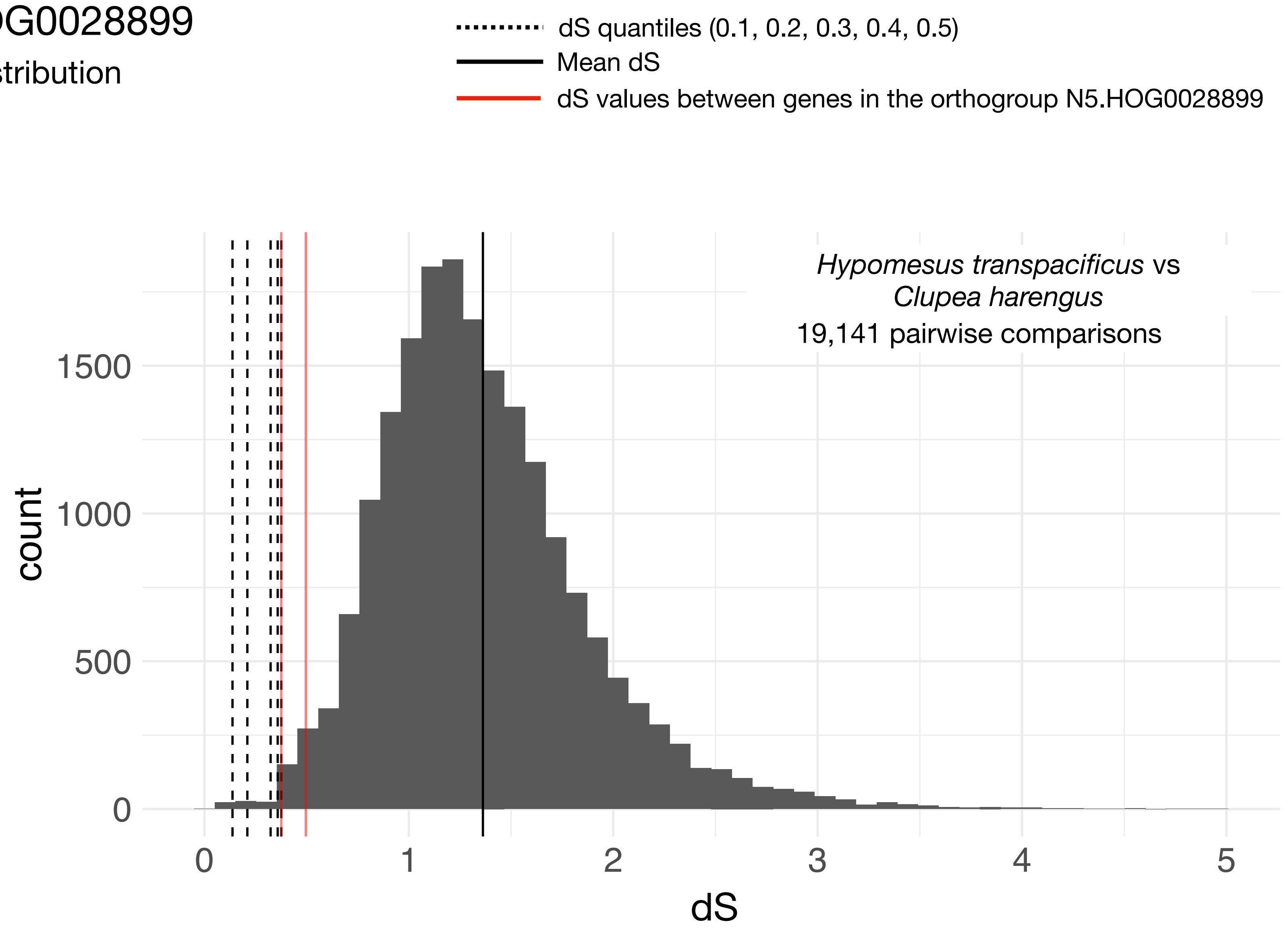


Intra-orthogroup dS

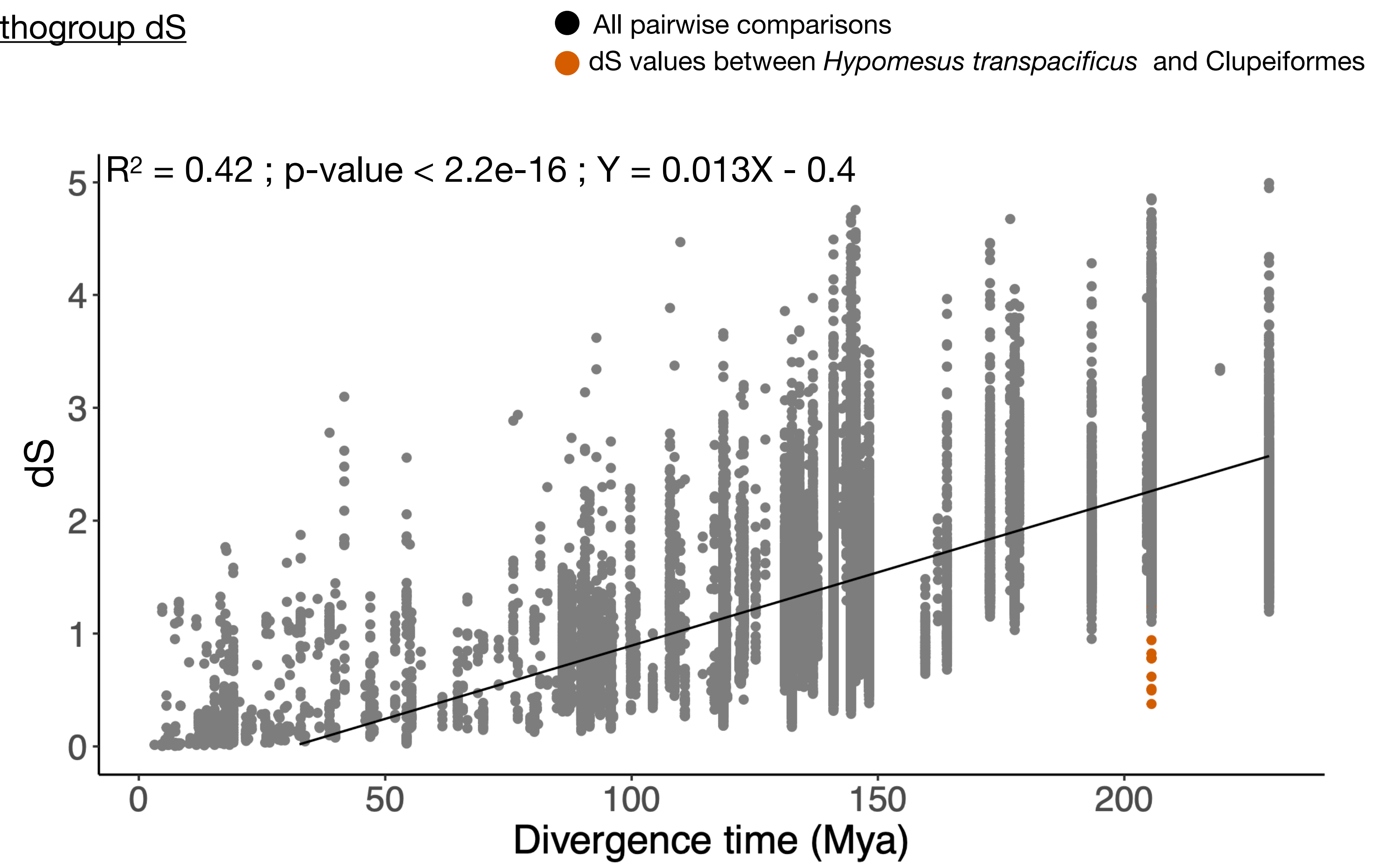

### N5.HOG0028899

The gene found in *H. transpacificus* is much shorter than the version of the gene found in *C. harengus*. Only 2 exons (on 4) remains intact in *H. transpacificus*

```
  138 : LeuCysTrpThrLysTrpTyrAspArgAspAsnProSerGlyThrGlyAspTrpGluLeu :   157
        :!!|||||||||||||||||||||||||||:!!||||||||||||||||||||||||||! !
        ValCysTrpThrLysTrpTyrAspArgAspAspProSerGlyThrGlyAspTrpGluSer
30088 : GTGTGCTGGACAAAGTGGTACGATCGTGATGATCCCTCCGGTACCGGTGACTGGGAGTCG : 30031

  158 : LeuGlnAsnLeuArgAsnGluAsnProGlyGluIleCysAlaAspProIleAlaIleGlu :   177
        |||..!:!!||||||!..||||||! !|||||||||||||||   |||||||||||||||
        LeuSerAspLeuArgArgGluAsnArgGlyGluIleCysAlaArgProIleAlaIleGlu
30030 : CTGTCGGACCTGCGGAGAGAGAACCGTGGTGAGATCTGTGCCAGACCAATCGCCATTGAG : 29971

  178 : SerArgThrValAspThrAspThrProAlaAlaThrThrGlyGlnAspPheLeu{Hi}   :   196
        |||!:!|||||||||||||!  ||||||||||||  !||||||||| !!|||||||{||}
        SerLysThrValAspThrGlyThrProAlaAlaGlnThrGlyGlnHisPheLeu{Hi}++
29970 : AGCAAGACTGTGGACACAGGGACTCCTGCCGCCCAGACTGGCCAACACTTCCTC{CA}gt : 29912

  197 : >>>> Target Intron 1 >>>>  {s}PheSerProThrThrGlyPheValCysLys :   206
                 345 bp            {|}|||||| !!||||..!|||||||||||||||
                                 ++{s}PheSerThrThrValGlyPheValCysLys
29911 : .........................ag{C}TTCTCTACGACAGTTGGGTTTGTATGCAAG : 29539

  207 : AsnGlyProAsnGlnTyrCysArgAspTyrLysValArgPheGlyCysProCysLys :   225
        |||||||||:!!||||||||||!:!||||||:!!|||||||||| !!||||||||||::!
        AsnGlyProAspGlnTyrCysLysAspTyrGlnValArgPheArgCysProCysArg
29538 : AACGGCCCCGACCAATACTGCAAGGACTACCAAGTCCGGTTTCGGTGTCCTTGTCGG : 29480
```

(*H. transpacificus* annotated gene have 4 exons but two are very small and not homologous to the Clupea gene version)

These 2 exons are also retrieved in *O. eperlanus* but it is not annotated by RefSeq

Window size : 40 ; Window step : 1 ; Nmatch : 27

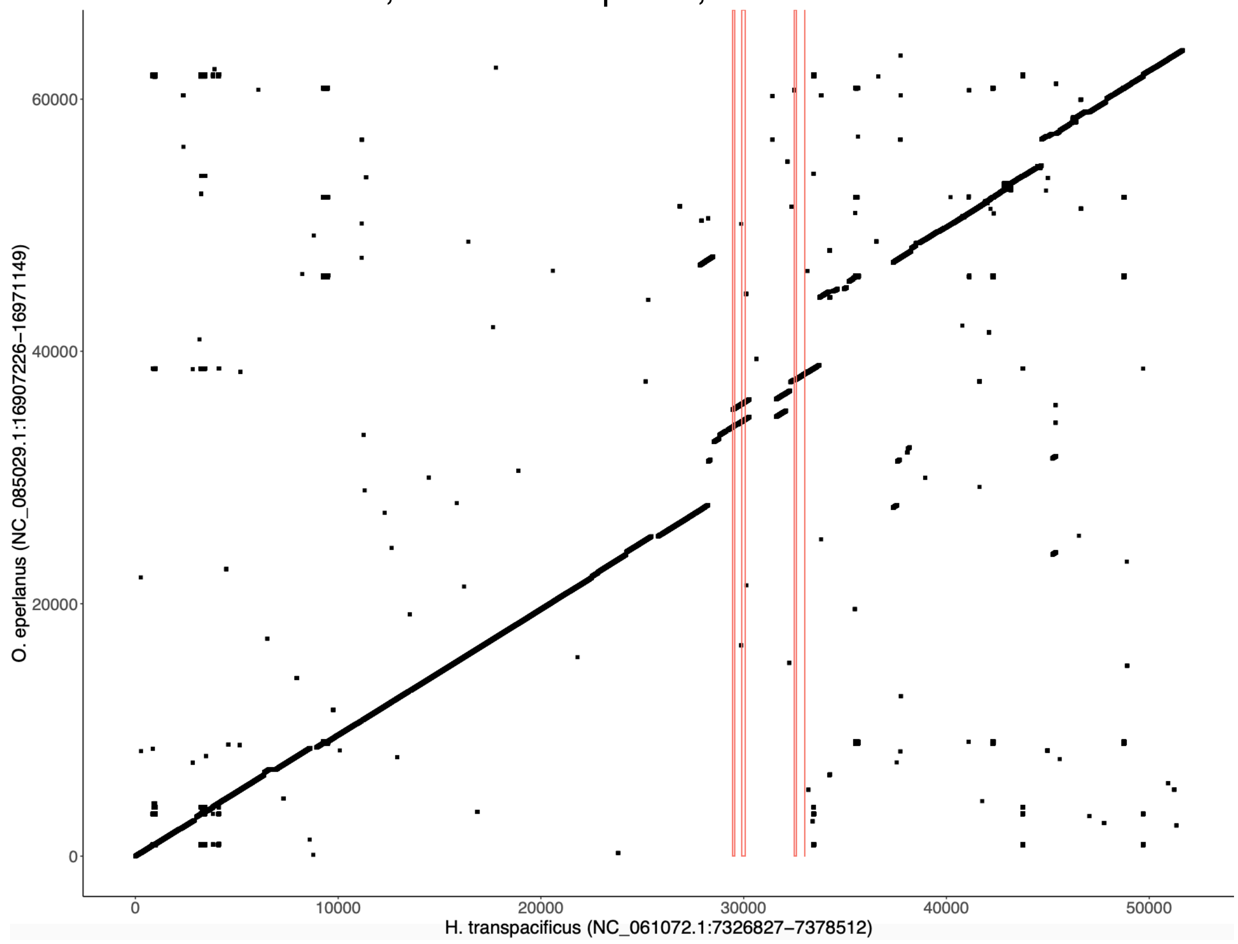


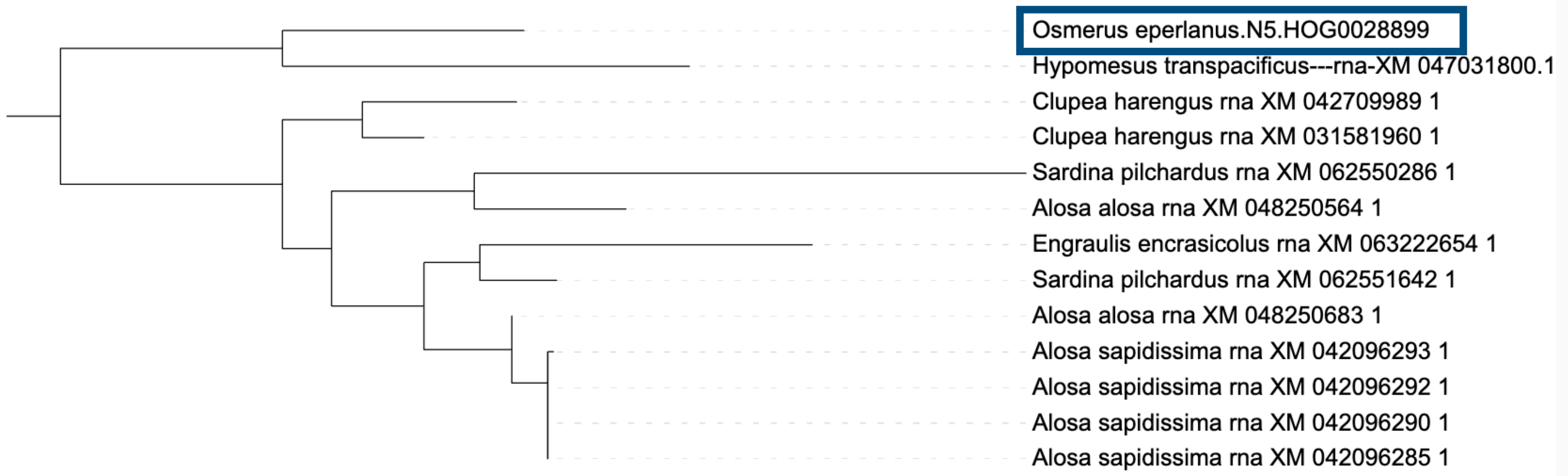

### N5.HOG0028899

#### Micro-synteny

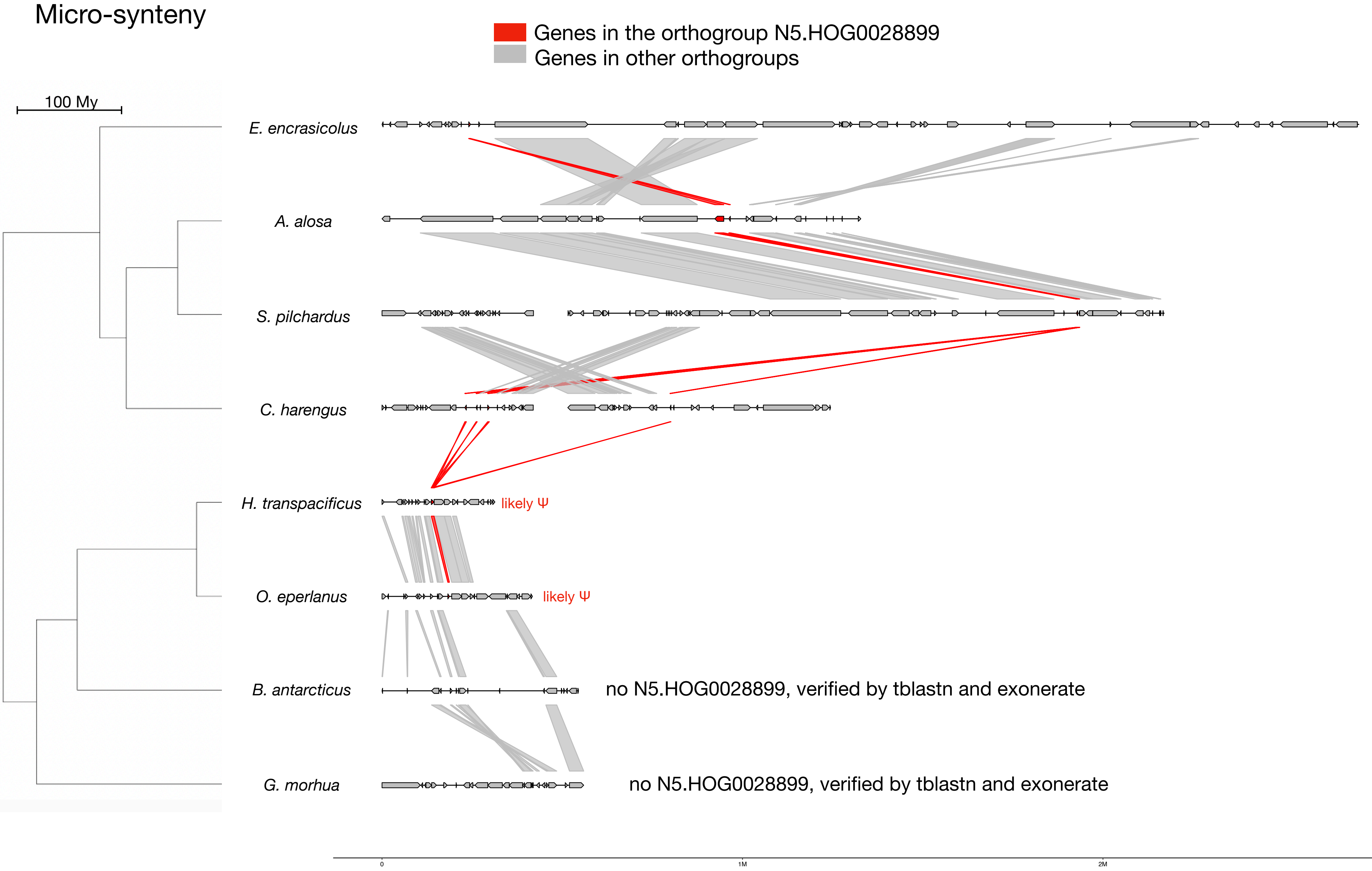


#### Zoom on *C. harengus* and *H. transpacificus*

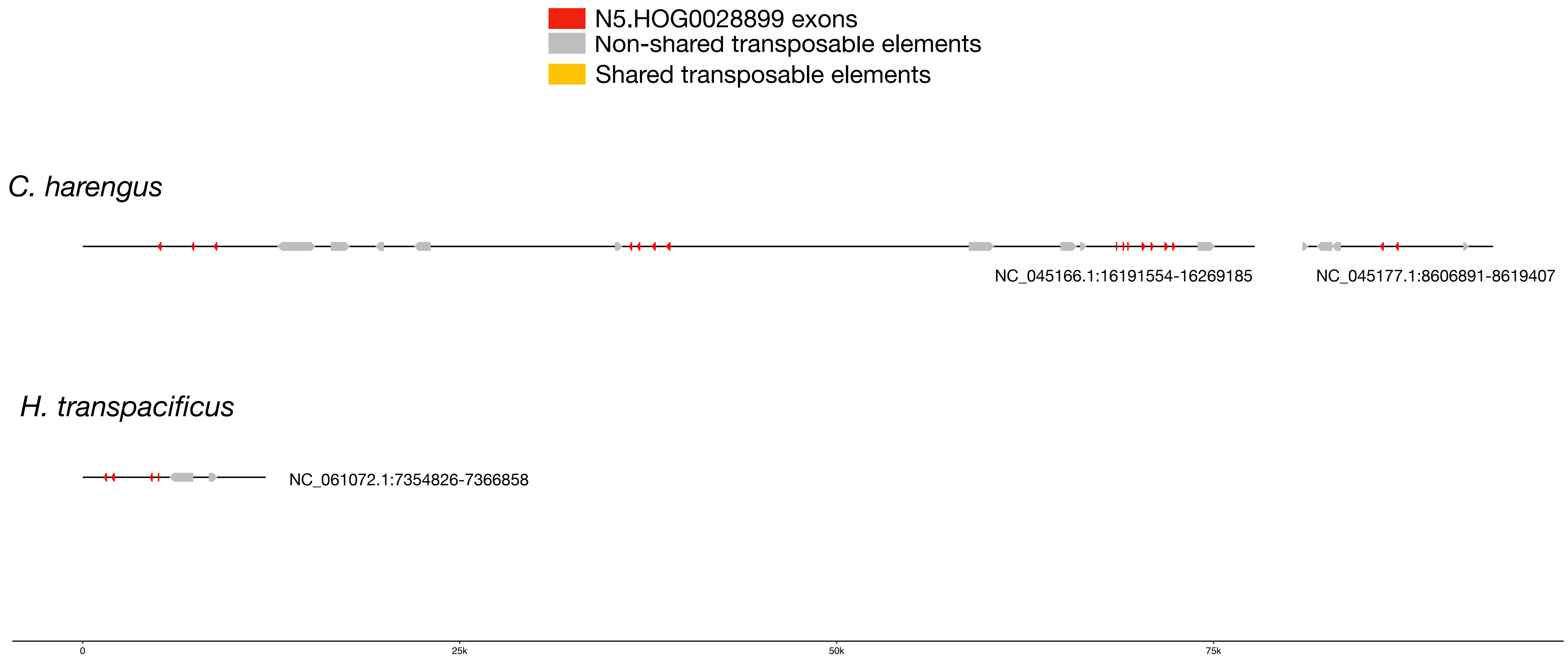

### N5.HOG0028899

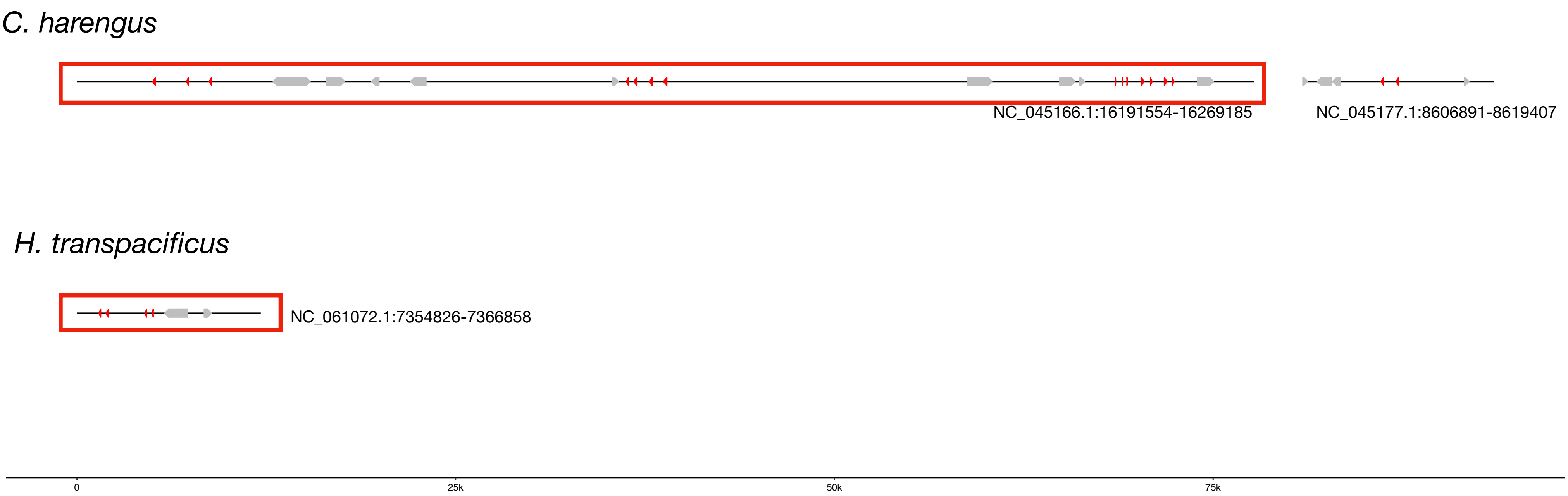


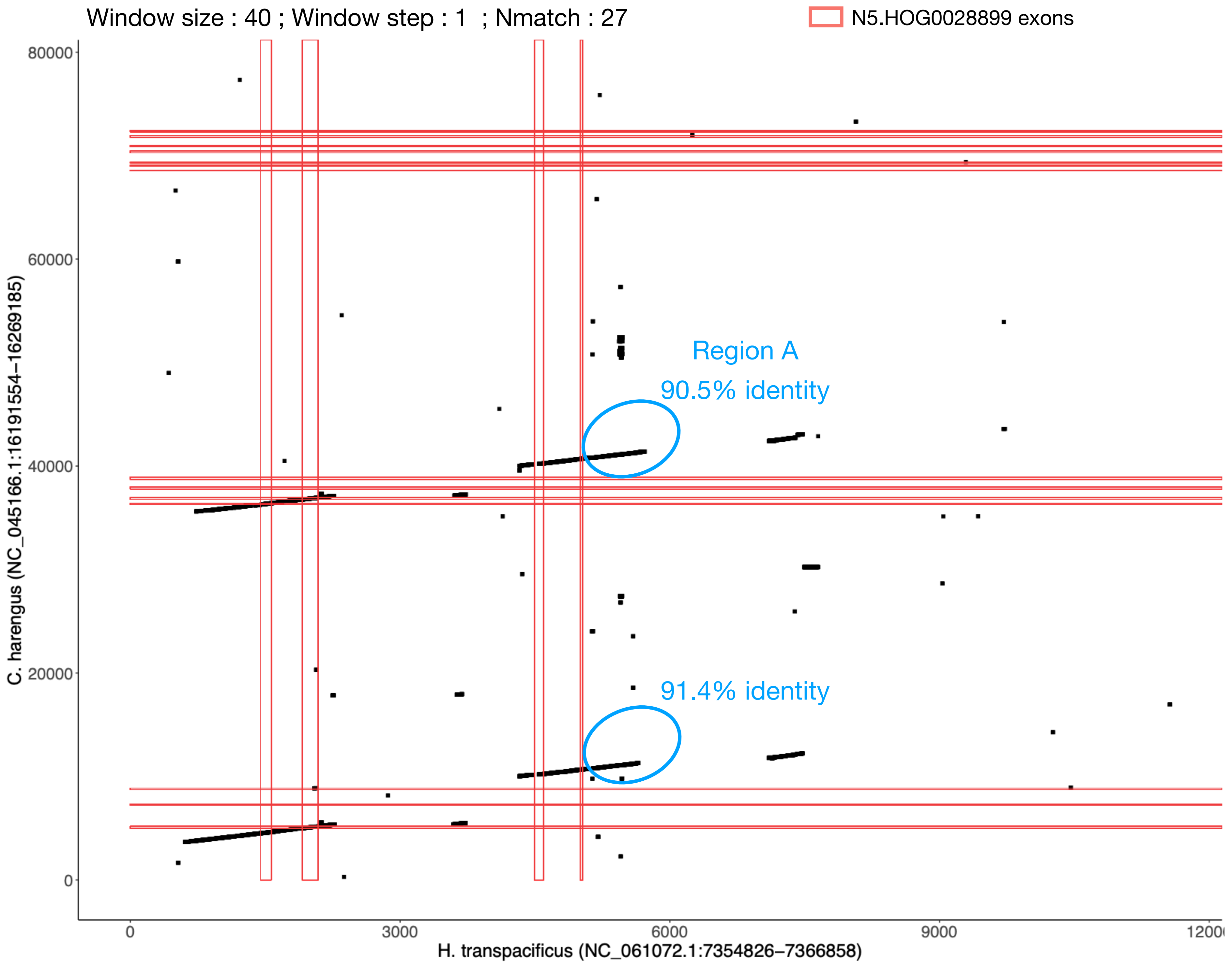

### N5.HOG0028899 Blastn of conserved regions against all genome assemblies

#### Region A

| sseqid | pident | length | mismatch | gapopen | qstart | qend | sstart | send | evalue | bitscore |
|---|---|---|---|---|---|---|---|---|---|---|
| Hypomesus_transpacificus-NC_061072.1 | 100.000 | 751 | 0 | 0 | 1 | 751 | 7359875 | 7360625 | 0.0 | 1387 |
| Osmerus_eperlanus-NC_085029.1 | 87.092 | 705 | 47 | 18 | 1 | 695 | 16932900 | 16932230 | 0.0 | 758 |
| Clupea_harengus-NC_045166.1 | 85.896 | 709 | 51 | 22 | 1 | 692 | 16280330 | 16281006 | 0.0 | 710 |
| Clupea_harengus-NC_045166.1 | 85.237 | 718 | 51 | 13 | 1 | 692 | 16232286 | 16232974 | 0.0 | 688 |
| Clupea_harengus-NC_045166.1 | 86.557 | 610 | 40 | 9 | 1 | 601 | 16202257 | 16202833 | 1.72e-177 | 634 |
| Clupea_harengus-NC_045166.1 | 82.396 | 409 | 28 | 22 | 1 | 393 | 16260078 | 16259698 | 7.07e-82 | 316 |
| Clupea_harengus-NC_045166.1 | 84.932 | 292 | 28 | 4 | 420 | 695 | 16254520 | 16254229 | 2.58e-71 | 281 |
| Clupea_harengus-NC_045177.1 | 81.431 | 587 | 67 | 14 | 117 | 692 | 8599837 | 8600392 | 1.12e-119 | 442 |
| Clupea_harengus-NC_045177.1 | 81.090 | 587 | 63 | 16 | 117 | 692 | 8663660 | 8664209 | 1.13e-114 | 425 |
| Clupea_harengus-NC_045177.1 | 81.316 | 380 | 36 | 17 | 221 | 596 | 8587481 | 8587829 | 1.20e-69 | 276 |

### N5.HOG0028899

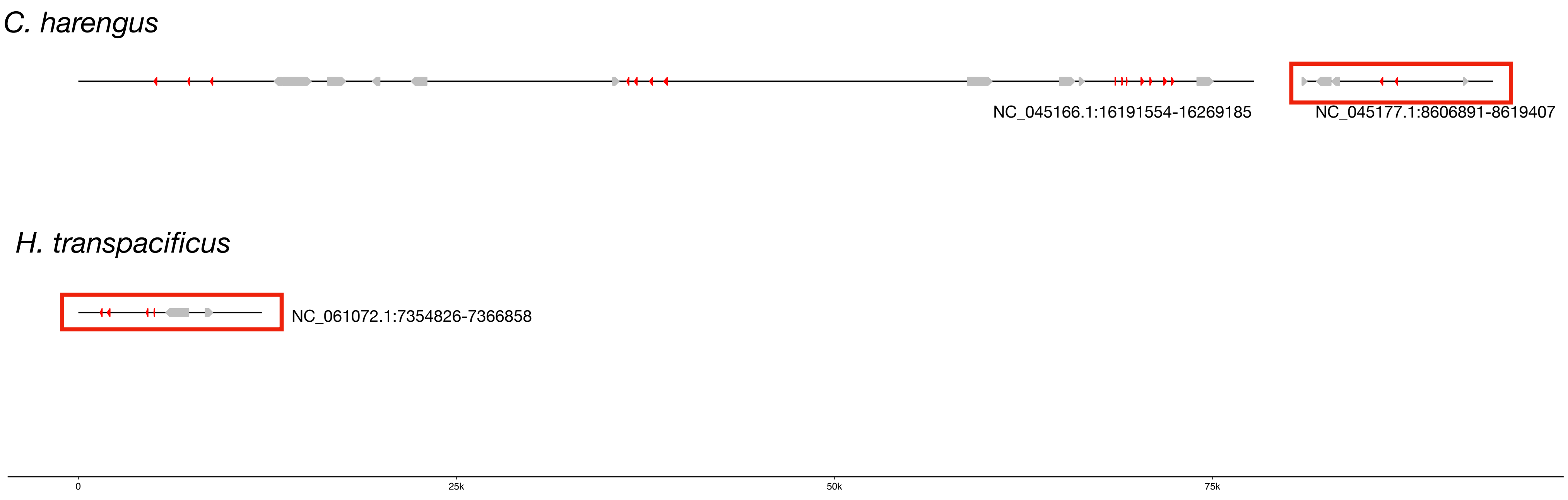


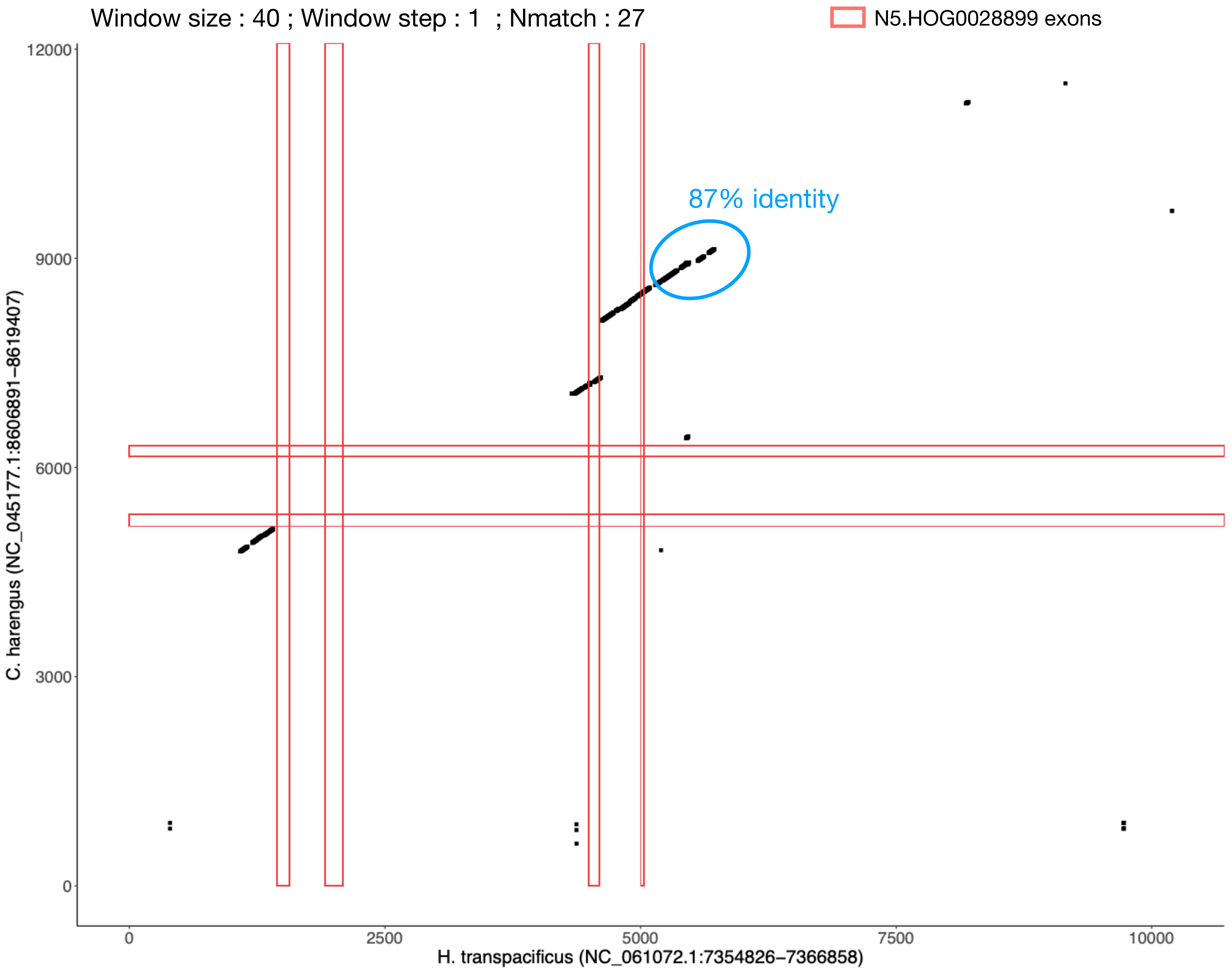

N5.HOG0028899

### Region1

#### Phylogeny and alignment of this non-coding conserved region: (between 87 and 91.4% identity)

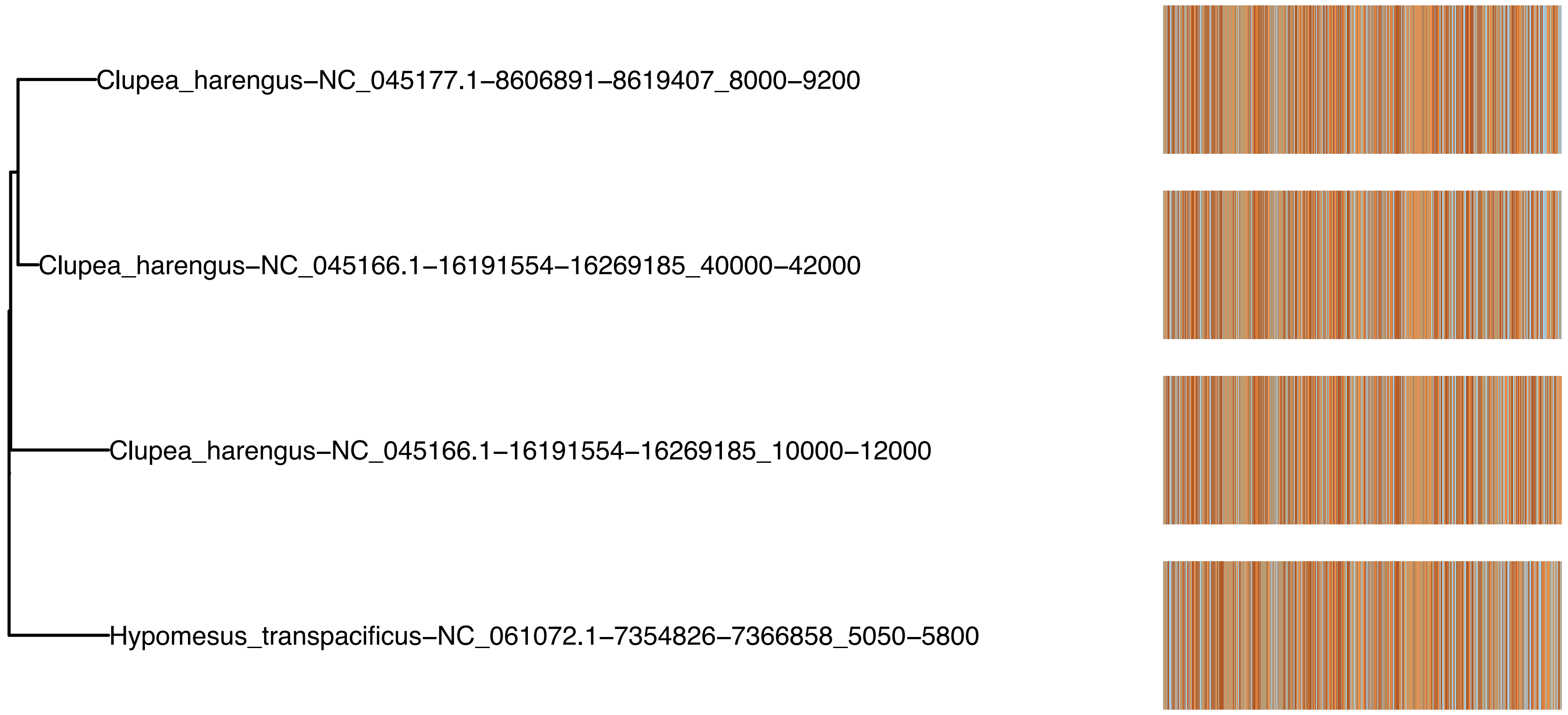


#### Blastn again all NCBI RefSeq genomes : Only match again Osmeriformes (*H. transpacificus* and *Osmerus spp.)* and *Clupea harengus*, next to N5.HOG0028899

| | |
|---|---|
| **Job Title** | **NC_061072.1-7354826-7366858:5050-5800** |
| **RID** | E55A7P82013 *Search expires on 09-13 15:16 pm* Download All |
| **Program** | BLASTN Citation |
| **Database** | refseq_reference_genomes (204 databases) See details |
| **Query ID** | lcl\|Query_3512079 |
| **Description** | NC_061072.1-7354826-7366858:5050-5800 |
| **Molecule type** | dna |
| **Query Length** | 751 |
| **Other reports** | Distance tree of results MSA viewer |

**Filter Results**

**Organism** *only top 20 will appear* exclude

Type common name, binomial, taxid or group name

+ Add organism

**Percent Identity** to **E value** to **Query Coverage** to

Filter Reset

**Descriptions** | Graphic Summary | Alignments | Taxonomy

**Sequences producing significant alignments** Download Select columns Show 100

select all *78 sequences selected* GenBank Graphics Distance tree of results MSA Viewer

| Description | Scientific Name | Max Score | Total Score | Query Cover | E value | Per. Ident | Acc. Len | Accession |
|---|---|---|---|---|---|---|---|---|
| Hypomesus transpacificus isolate Combined female chromosome 13, fHypTra1 | Hypomesus transpacificus | 1387 | 1387 | 100% | 0.0 | 100.00% | 14850352 | NC_061072.1 |
| Osmerus mordax isolate fOsmMor3 chromosome 10, fOsmMor3.pri | Osmerus mordax | 889 | 889 | 92% | 0.0 | 89.82% | 19003928 | NC_090059.1 |
| Osmerus eperlanus chromosome 12, fOsmEpe2.1 | Osmerus eperlanus | 758 | 758 | 92% | 0.0 | 87.09% | 18633437 | NC_085029.1 |
| Clupea harengus chromosome 15, Ch_v2.0.2 | Clupea harengus | 710 | 3606 | 98% | 0.0 | 85.90% | 28713521 | NC_045166.1 |
| Clupea harengus chromosome 26, Ch_v2.0.2 | Clupea harengus | 442 | 2042 | 82% | 1e-119 | 81.43% | 12443209 | NC_045177.1 |
| Clupea harengus unplaced genomic scaffold, Ch_v2.0.2 | Clupea harengus | 82.4 | 82.4 | 6% | 2e-11 | 97.87% | 76004 | NW_024879754.1 |

N5.HOG0028899

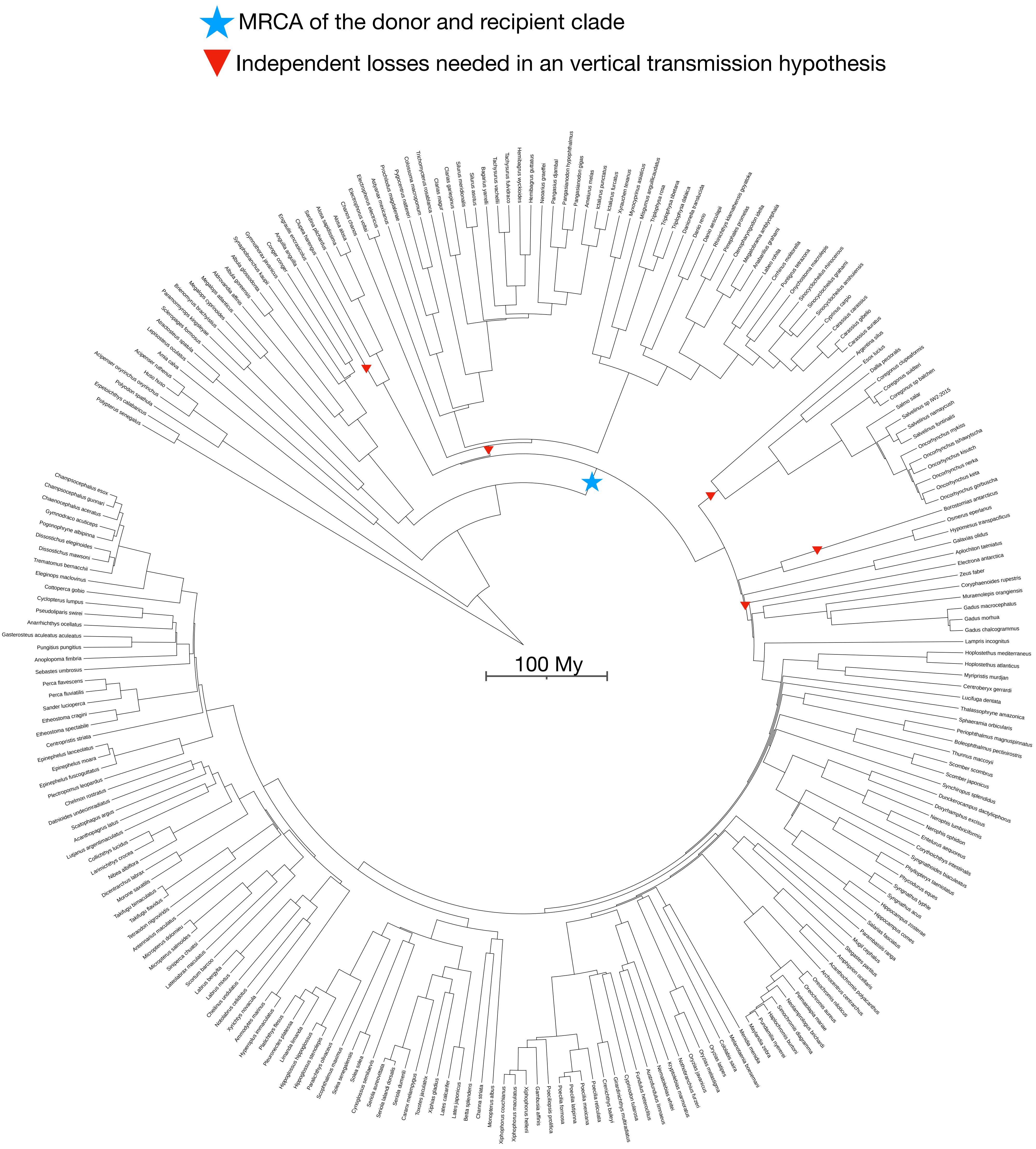

N5.HOG0004450

N5.HOG0004450

Species tree

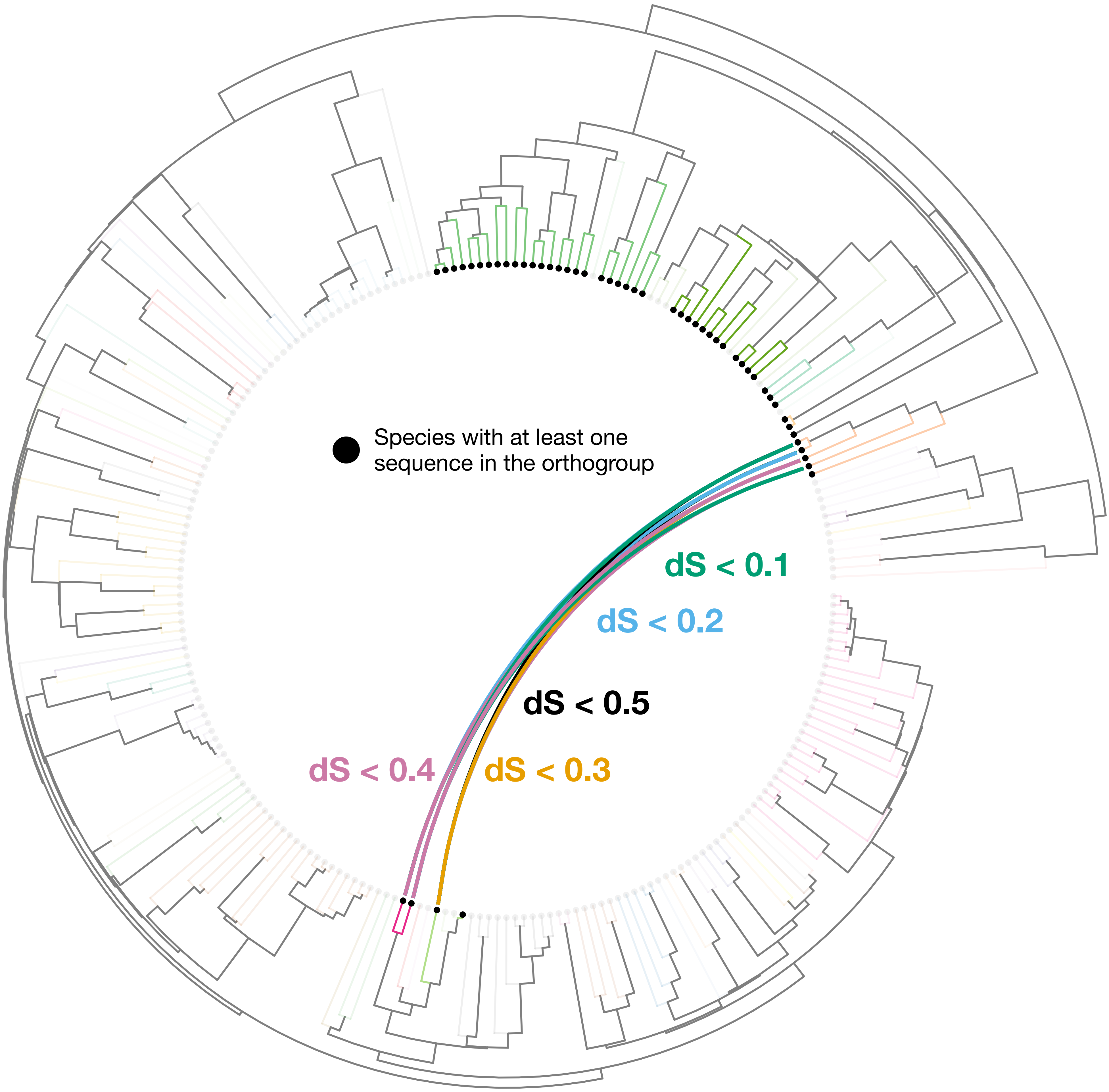

### N5.HOG0004450

Orthogroup phylogeny
and
protein alignment

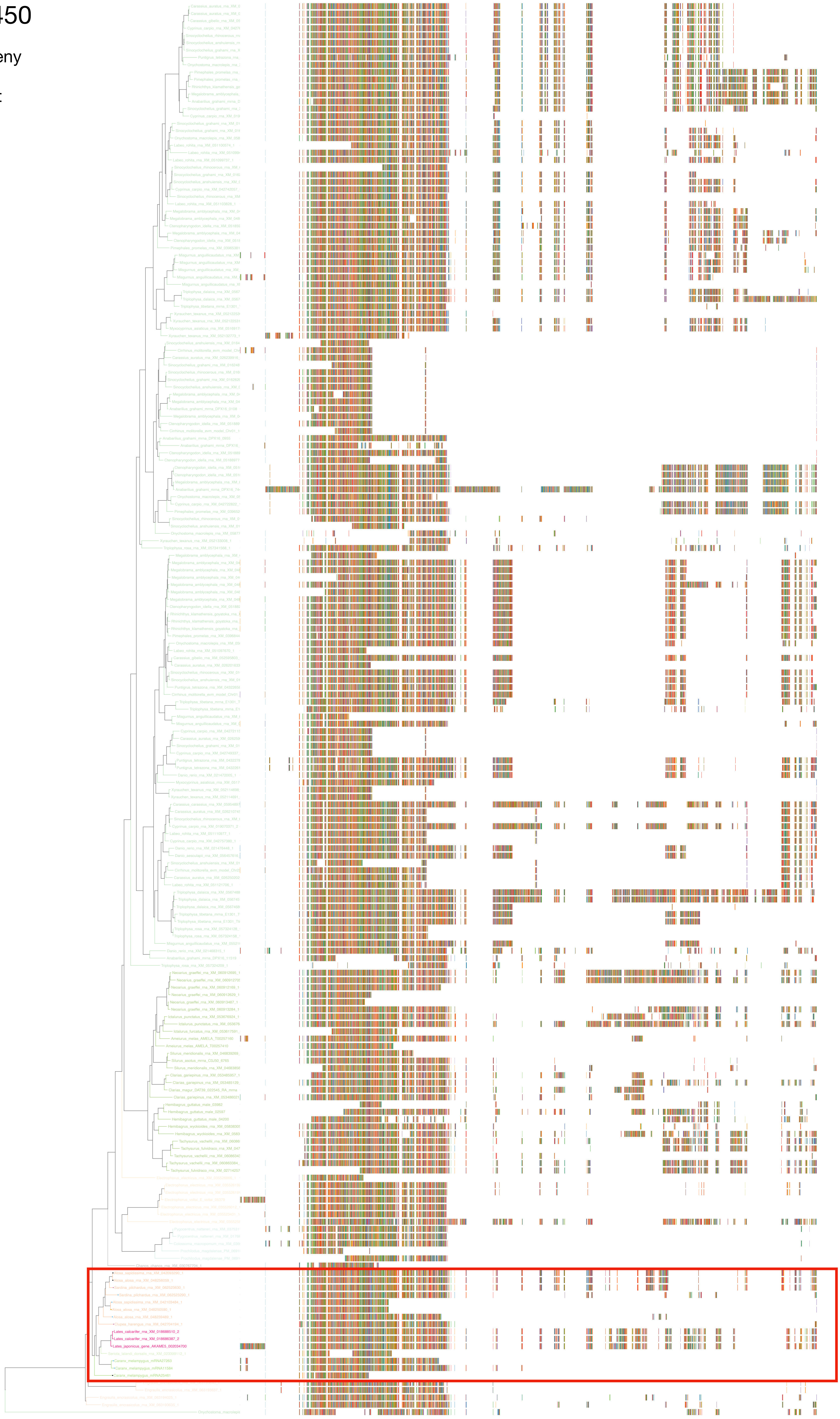

### N5.HOG0004450

Zoom on HGT clade with bootstrap values

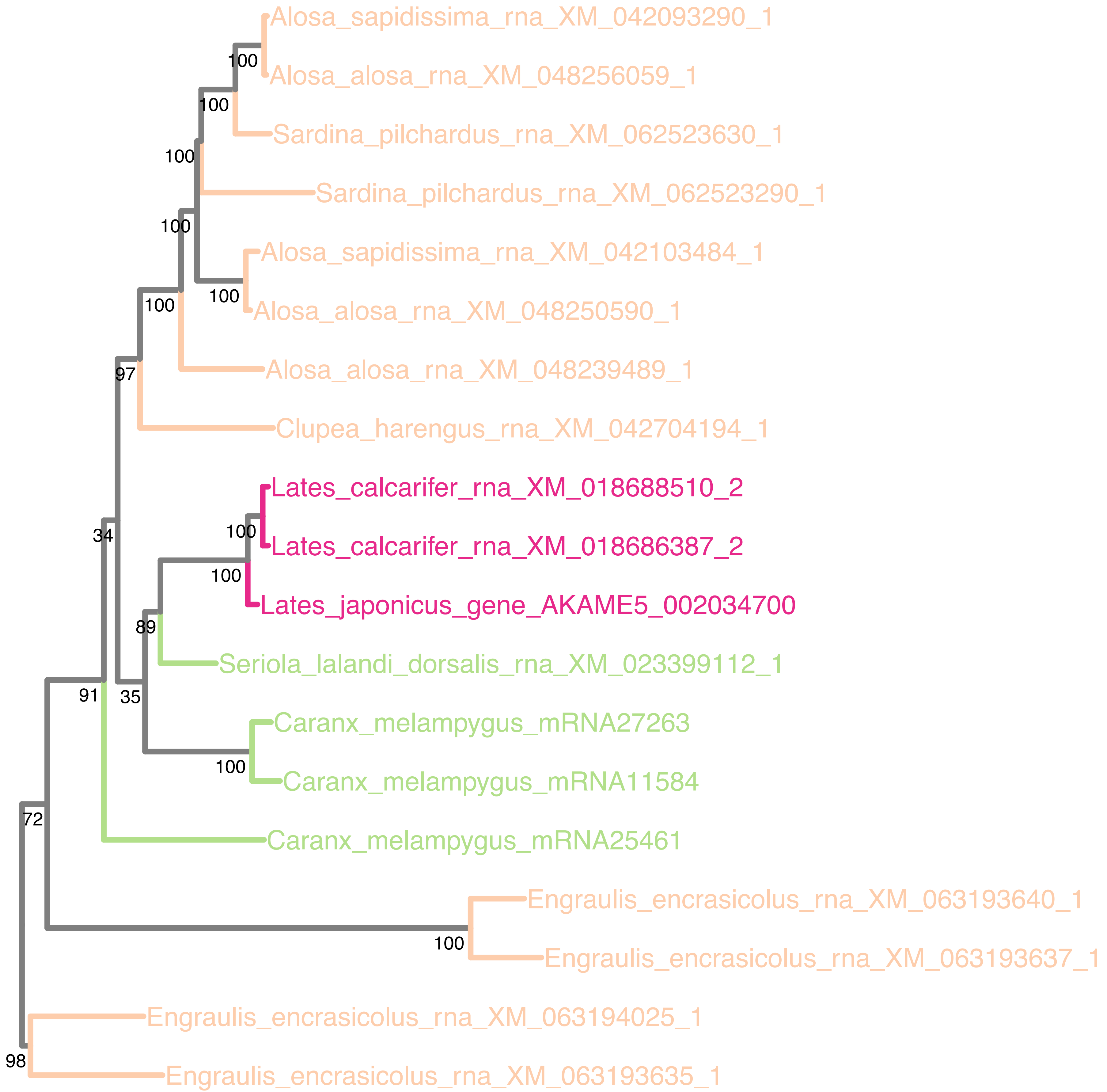

### N5.HOG0004450

Orthogroup phylogeny
and
trimmed
protein alignment

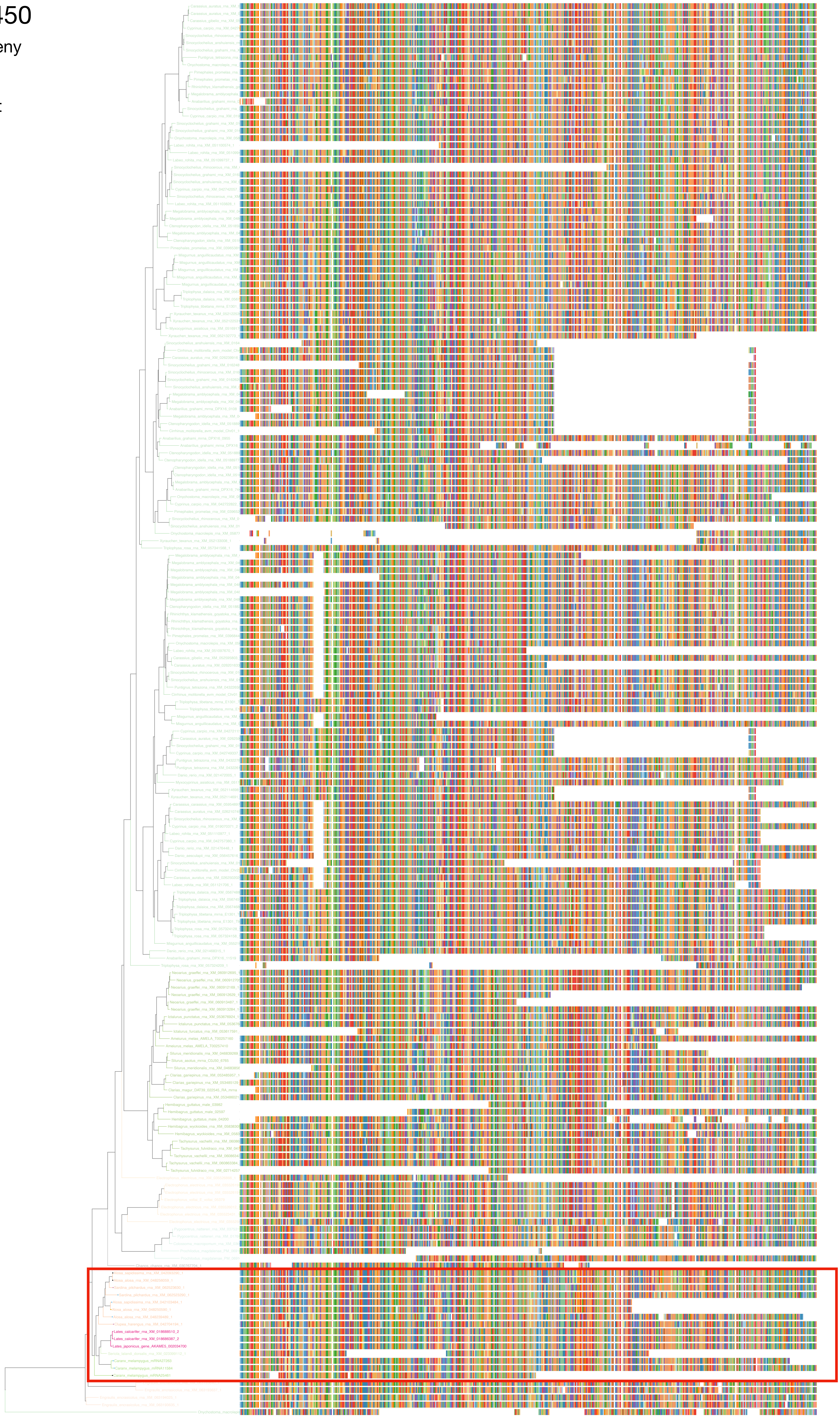

### N5.HOG0004450

« Best-match » phylogeny
and
protein alignment

50 best match against ray-finned fishes protein databases
10 best match against non-fish proteins from UniProt

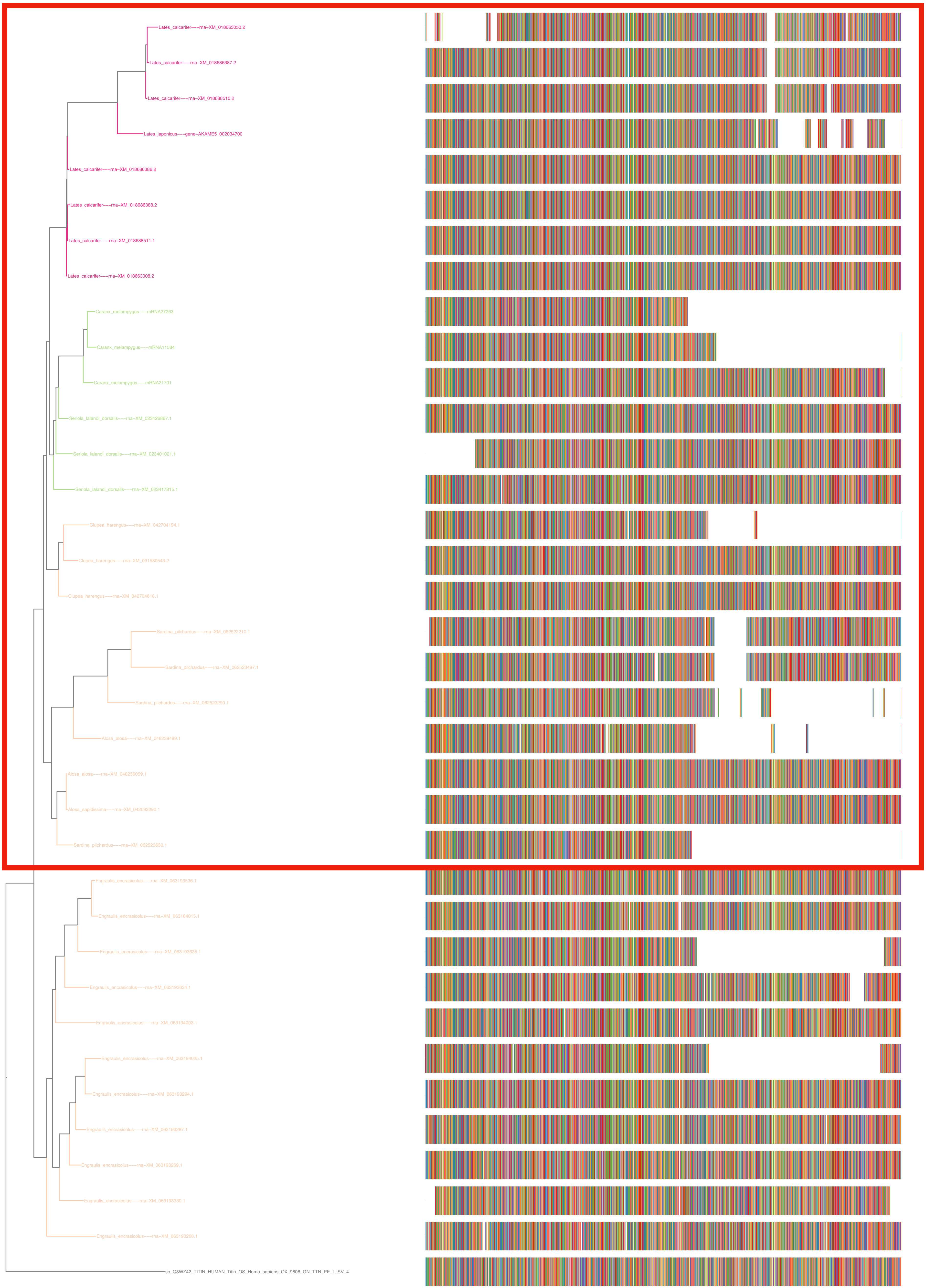

### N5.HOG0004450

#### dS distribution

Inter-species dS distribution

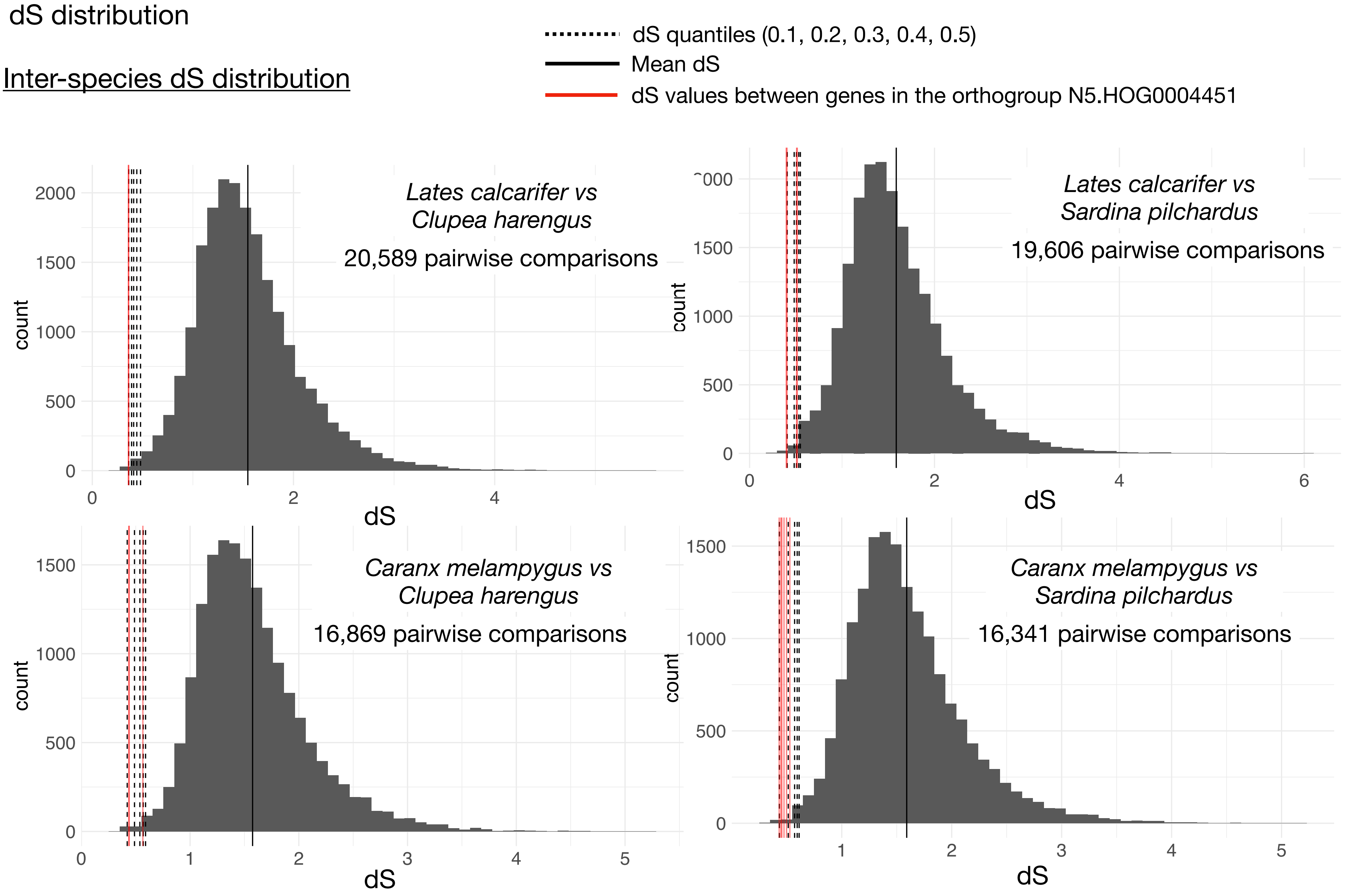


Intra-orthogroup dS

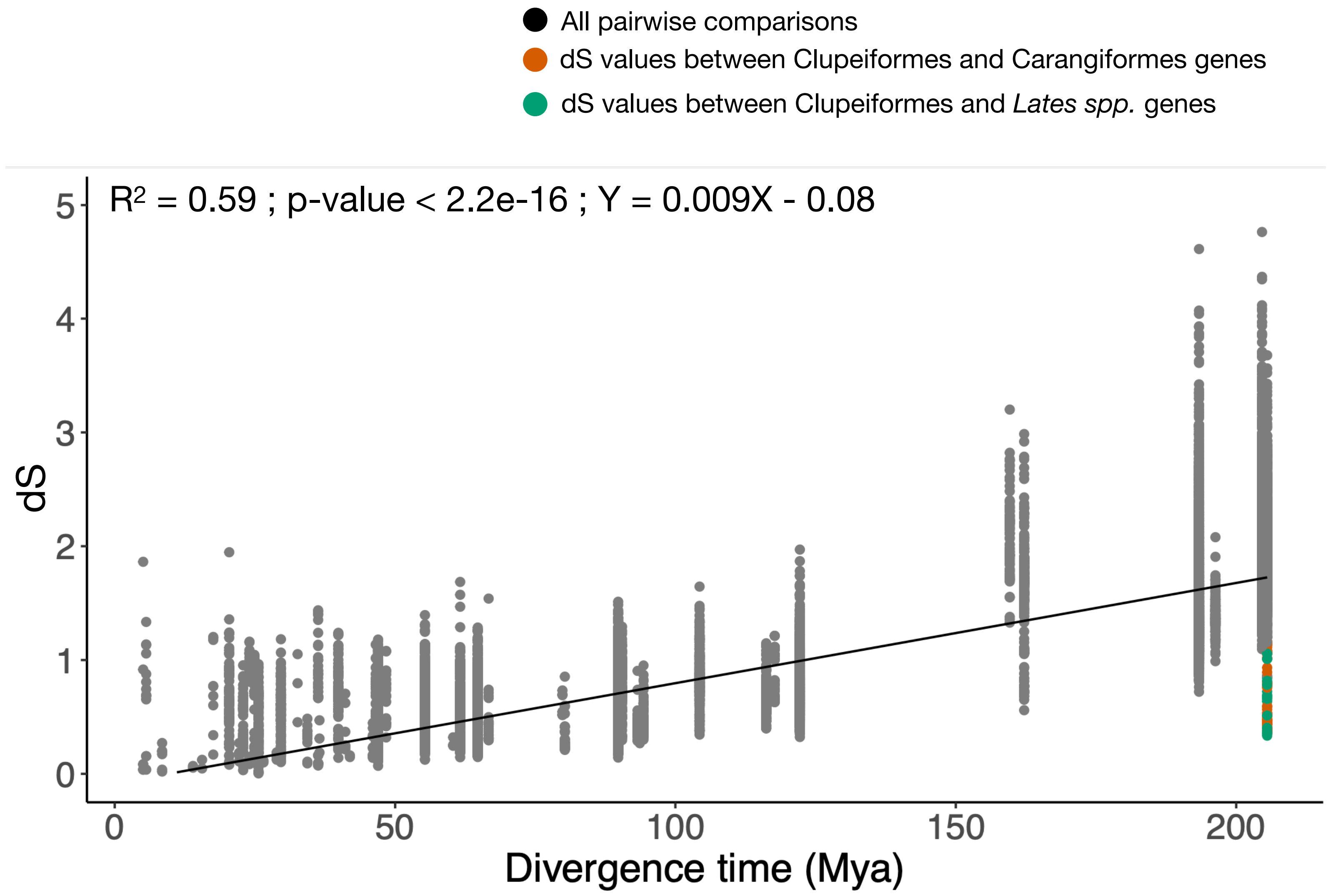

### N5.HOG0004450

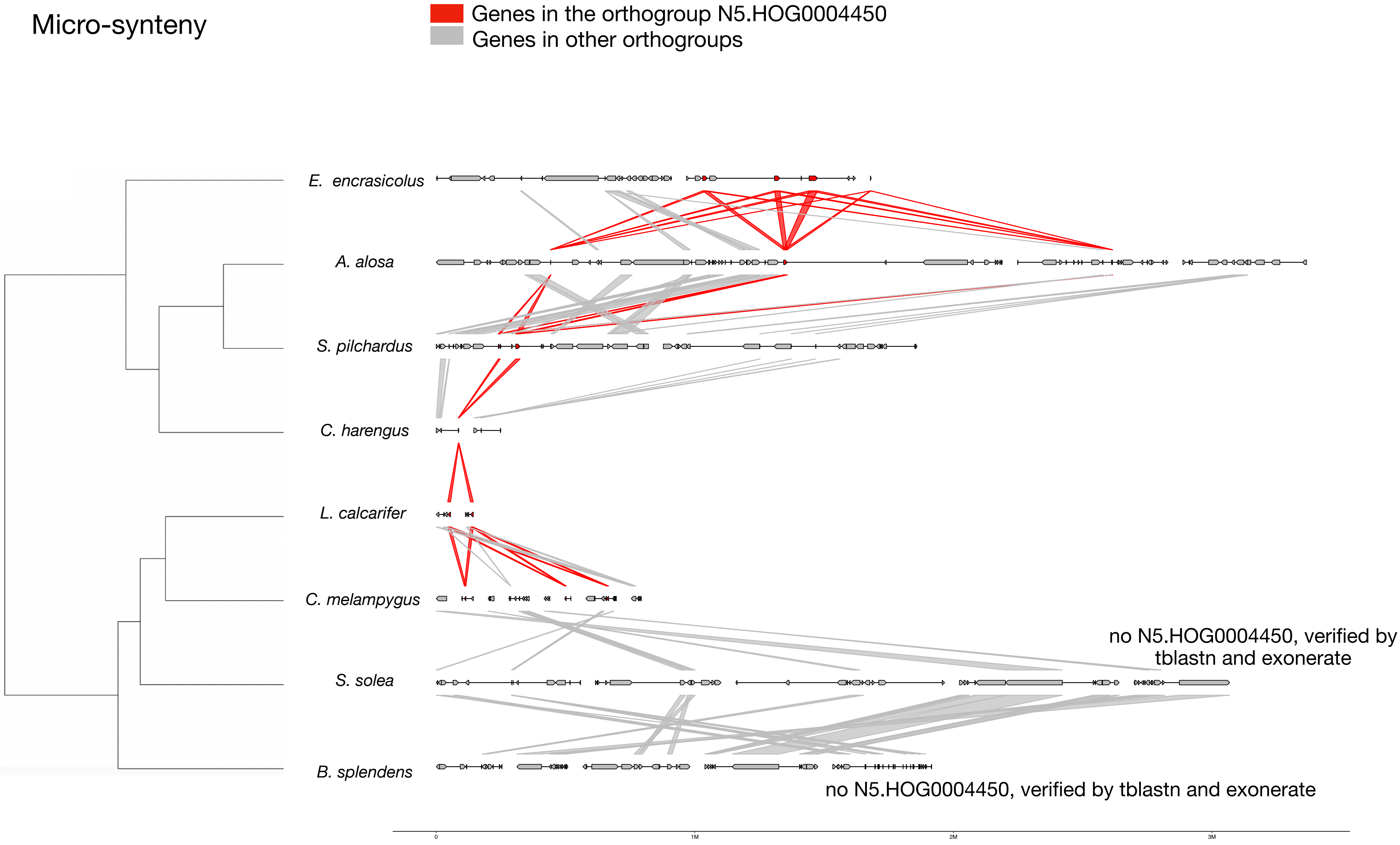


#### Zoom on *S. pilchardus* and *C. melampygus*

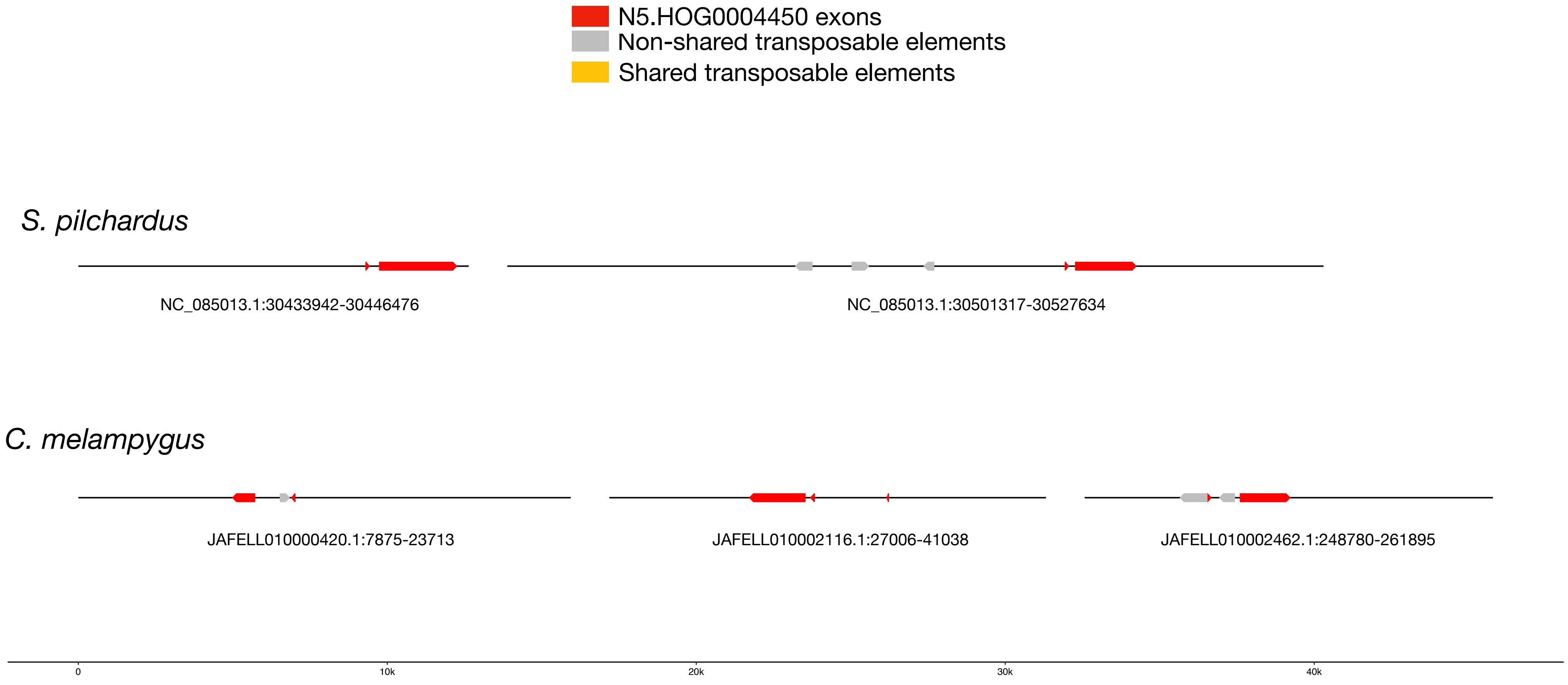

### N5.HOG0004450

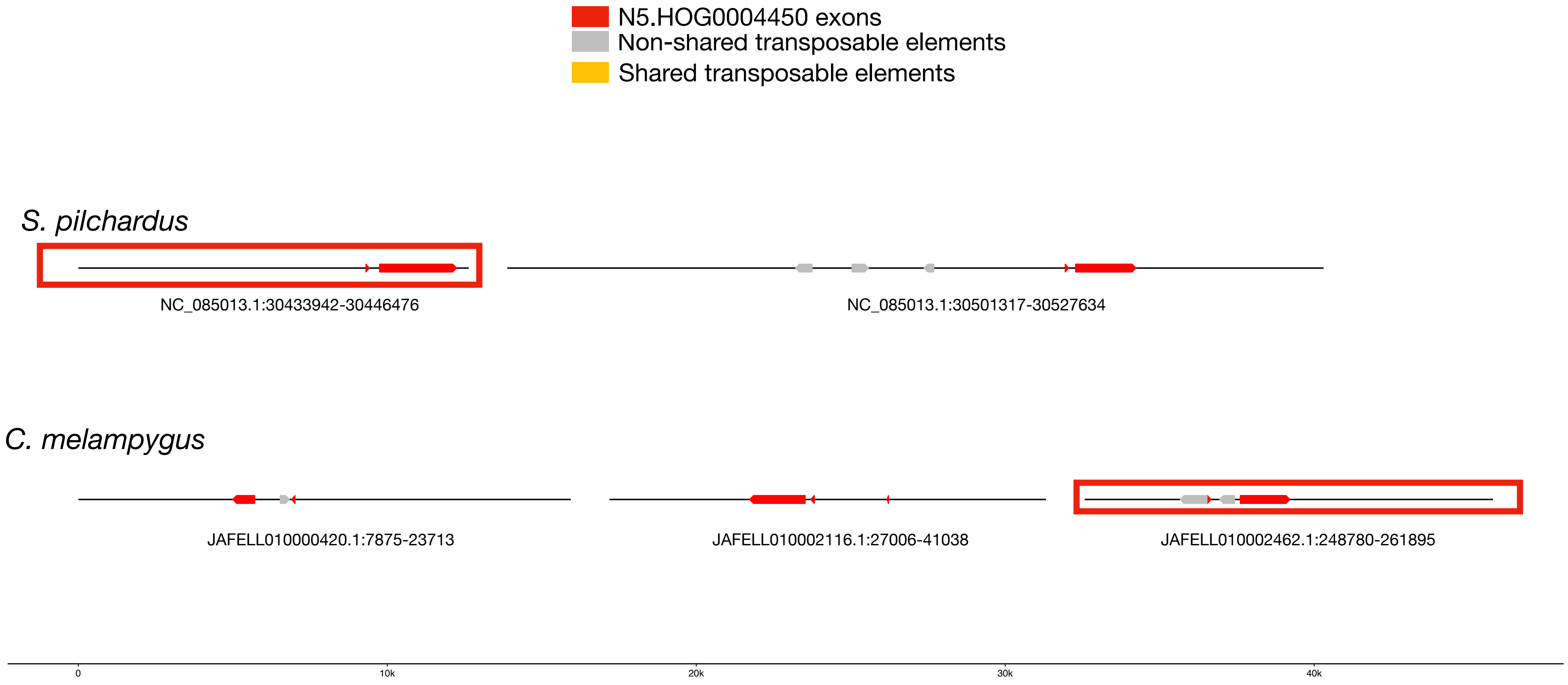


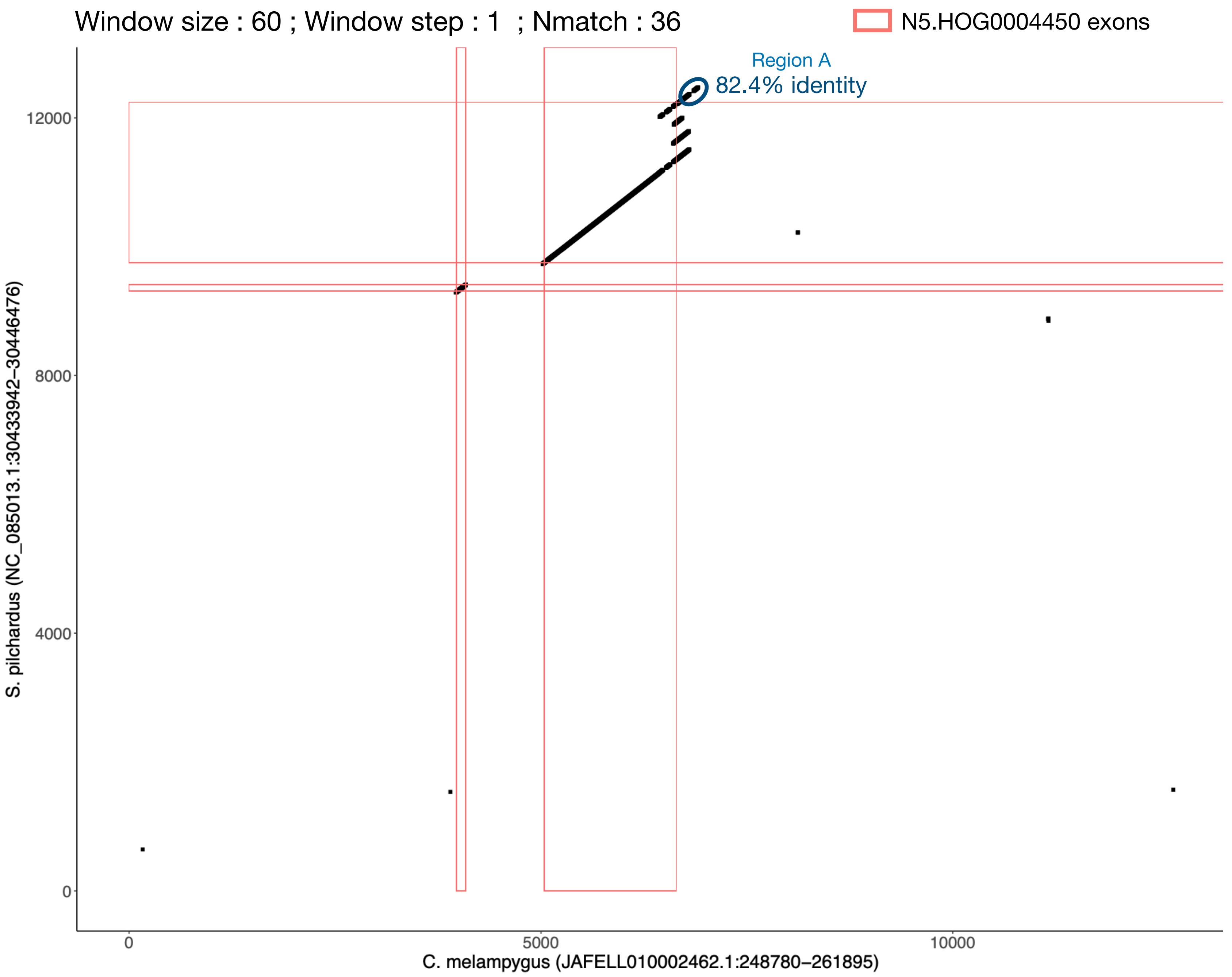

### N5.HOG0004450 Blastn of conserved regions against all genome assemblies

#### Region A

| sseqid | pident | length | mismatch | gapopen | qstart | qend | sstart | send | evalue | bitscore |
|---|---|---|---|---|---|---|---|---|---|---|
| Caranx_melampygus-JAFELL010002462.1 | 100.000 | 360 | 0 | 0 | 1 | 360 | 255420 | 255779 | 0.0 | 665 |
| Caranx_melampygus-JAFELL010000554.1 | 97.759 | 357 | 6 | 2 | 1 | 357 | 206965 | 207319 | 1.02e-171 | 614 |
| Caranx_melampygus-JAFELL010002120.1 | 97.479 | 357 | 8 | 1 | 1 | 357 | 31739 | 31384 | 4.74e-170 | 608 |
| Caranx_melampygus-JAFELL010002120.1 | 93.056 | 360 | 14 | 9 | 1 | 357 | 1419 | 1068 | 2.96e-142 | 516 |
| Caranx_melampygus-JAFELL010003188.1 | 96.703 | 364 | 7 | 3 | 1 | 360 | 32266 | 32628 | 7.94e-168 | 601 |
| Caranx_melampygus-JAFELL010003188.1 | 98.208 | 279 | 5 | 0 | 79 | 357 | 15919 | 16197 | 6.45e-134 | 488 |
| Caranx_melampygus-JAFELL010002129.1 | 96.639 | 357 | 8 | 4 | 1 | 357 | 2339 | 1987 | 1.72e-164 | 590 |
| Seriola_lalandi_dorsalis-NW_019525255.1 | 85.593 | 354 | 47 | 3 | 2 | 352 | 1828 | 1476 | 8.76e-98 | 368 |
| Seriola_aureovittata-NC_079375.1 | 85.352 | 355 | 47 | 4 | 2 | 352 | 25420596 | 25420949 | 4.08e-96 | 363 |
| Seriola_lalandi_dorsalis-NW_019523873.1 | 84.225 | 355 | 51 | 4 | 2 | 352 | 4010 | 3657 | 1.91e-89 | 340 |
| Seriola_dumerili-NW_019174519.1 | 83.989 | 356 | 50 | 6 | 2 | 352 | 18475 | 18828 | 8.89e-88 | 335 |
| Seriola_dumerili-NW_019174519.1 | 83.989 | 356 | 50 | 6 | 2 | 352 | 41931 | 41578 | 8.89e-88 | 335 |
| Clupea_harengus-NW_024879688.1 | 80.845 | 355 | 63 | 4 | 2 | 352 | 84591 | 84944 | 1.96e-69 | 274 |
| Clupea_harengus-NC_045165.1 | 80.337 | 356 | 63 | 6 | 2 | 352 | 1167025 | 1166672 | 4.25e-66 | 263 |

N5.HOG0004450

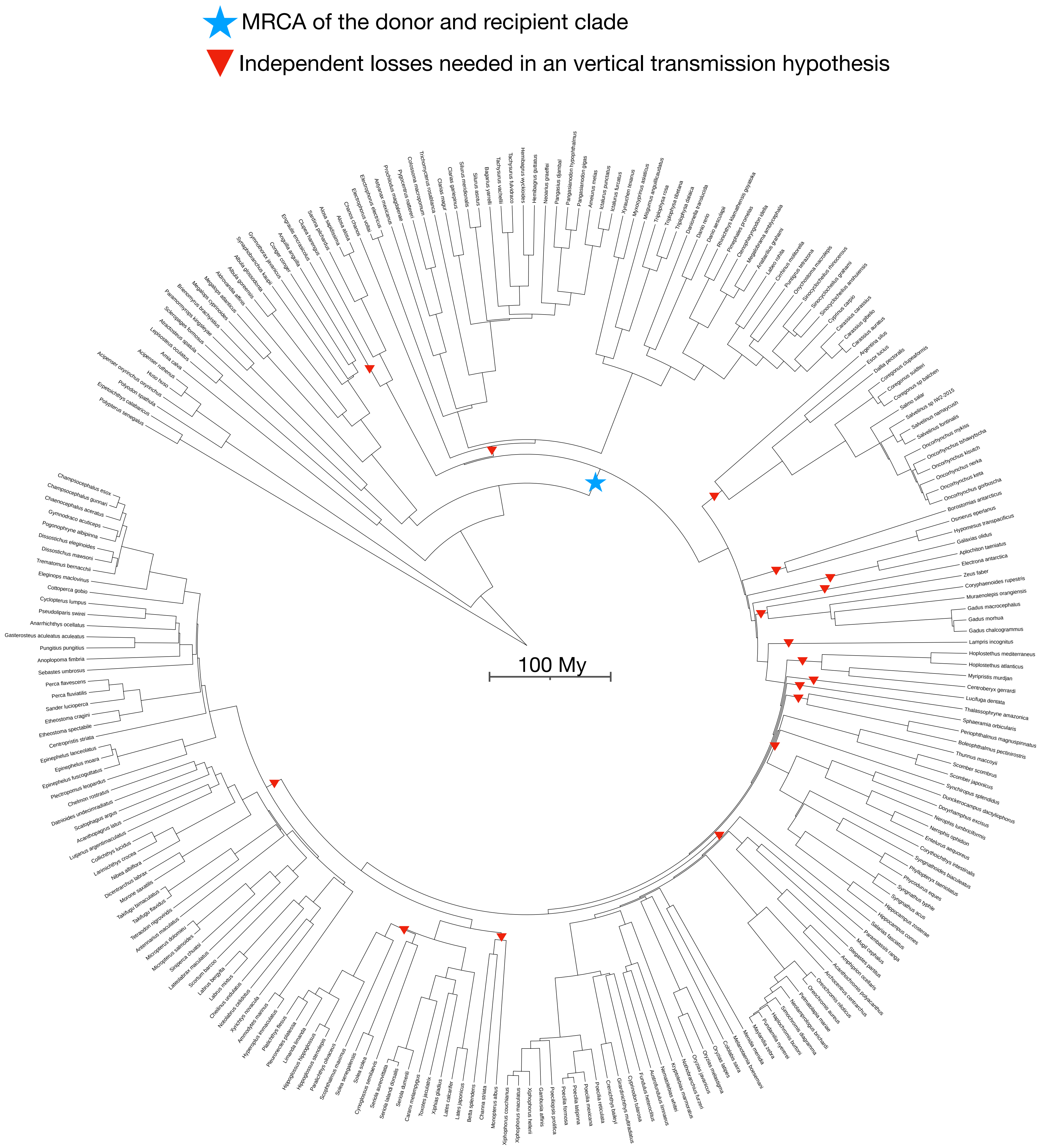

N5.HOG0029633 and
N5.HOG0046344

N5.HOG0029633

Species tree

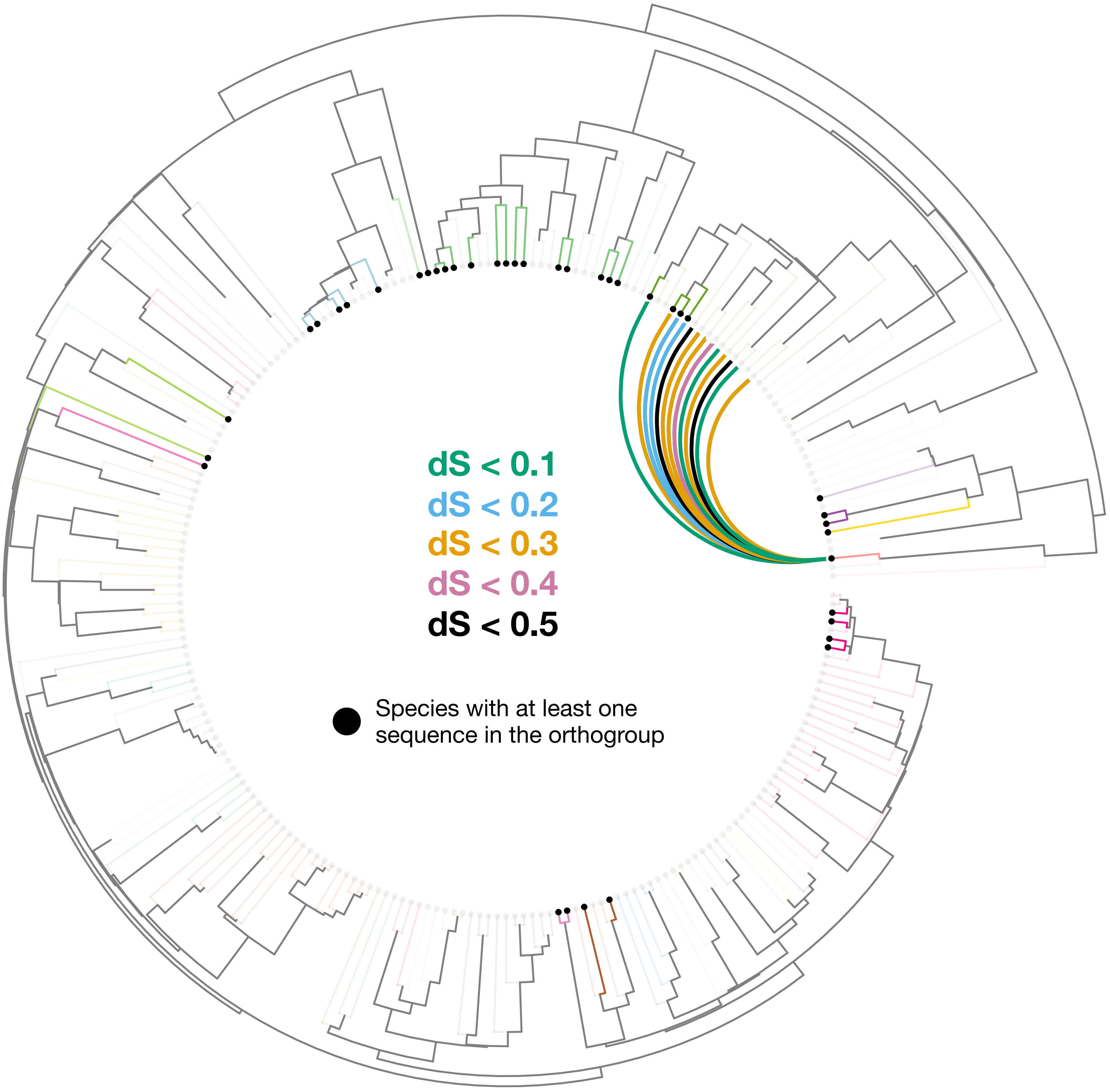

### N5.HOG0029633

Orthogroup phylogeny
and
protein alignment

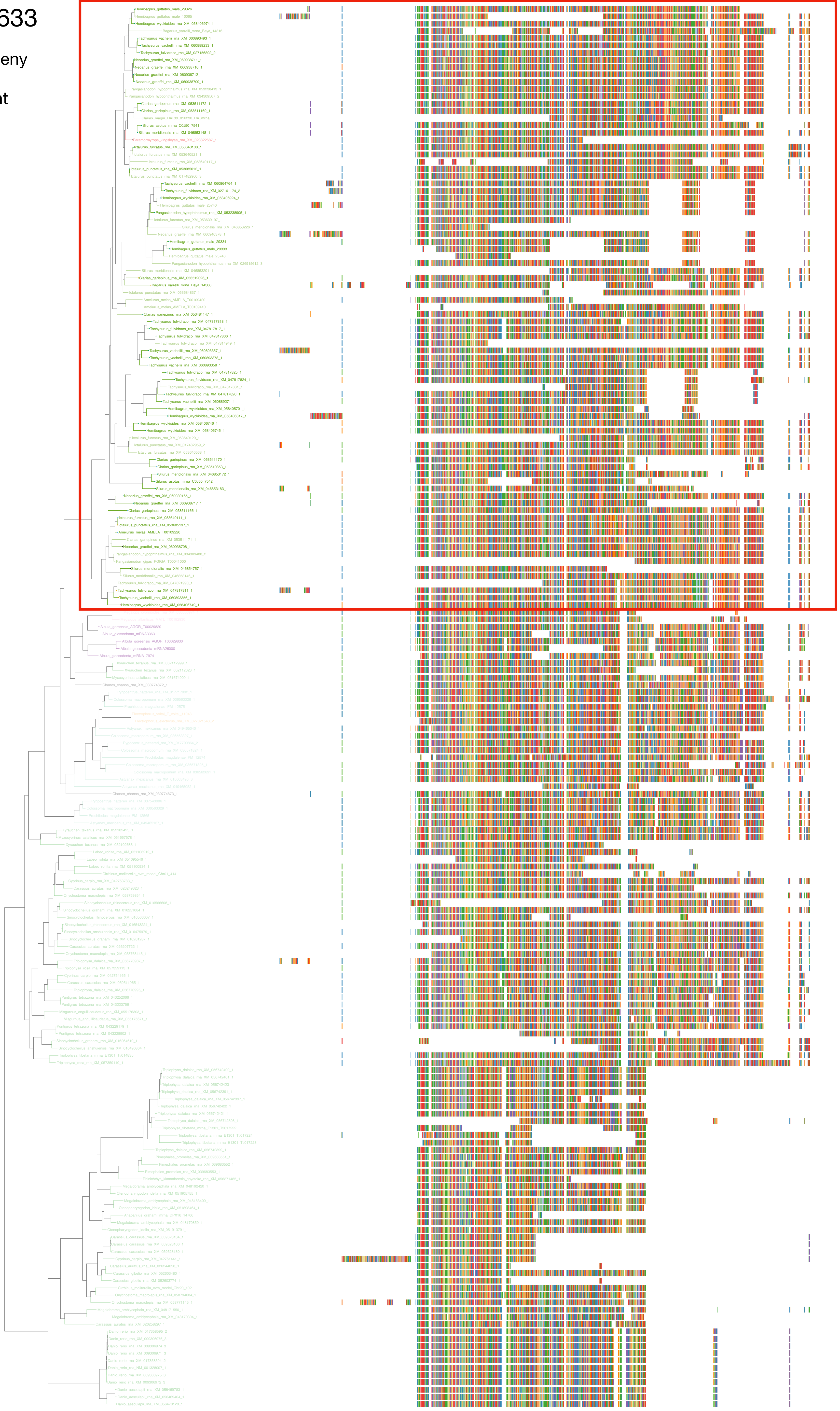

### N5.HOG0029633

Zoom on HGT clade with bootstrap values

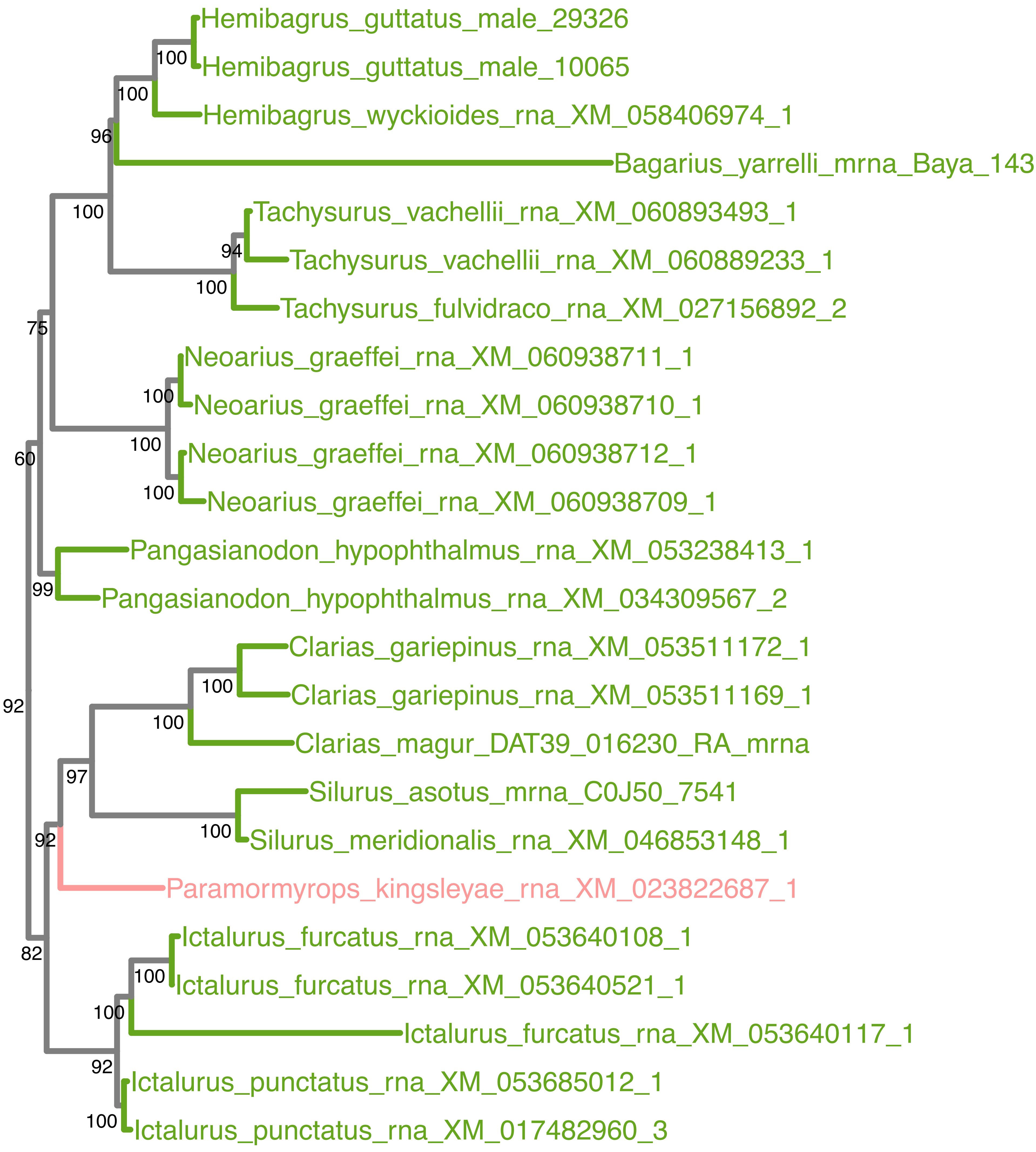

### N5.HOG0029633

Orthogroup phylogeny
and
trimmed
protein alignment

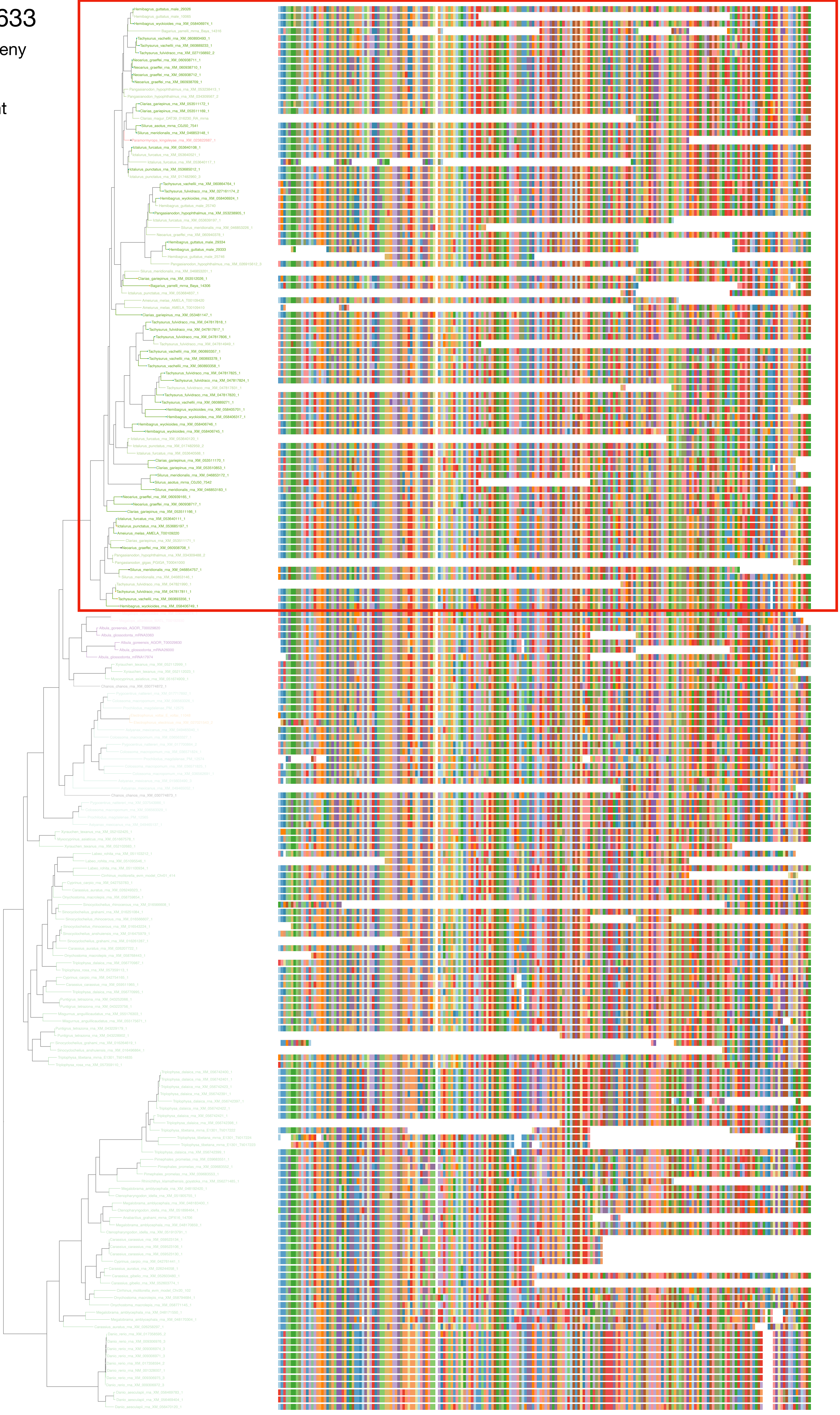

### N5.HOG0029633

« Best-match » phylogeny
and
protein alignment

50 best match against ray-finned fishes protein databases
10 best match against non-fish proteins from UniProt

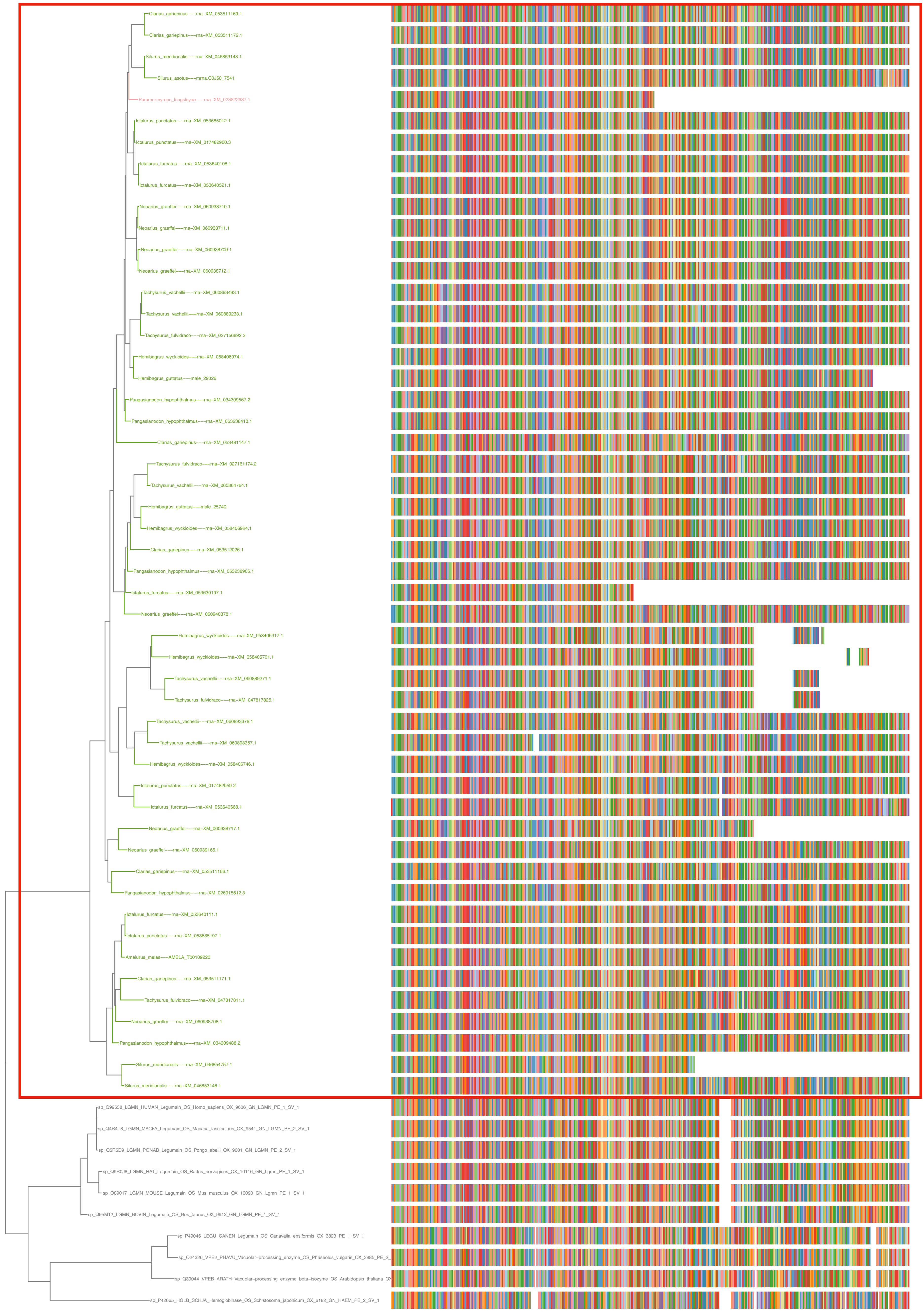

N5.HOG0029633

The sequence is not correctly annotated on the genome of *Paramormyrops kingsleyae*. The full length sequence was retrieved with Exonerate. The gene can also be retrieved entirely on an other genome assembly of *Paramormyrops kingsleyae* + on the genome of the closely related *Paramormyrops hopkinsi*

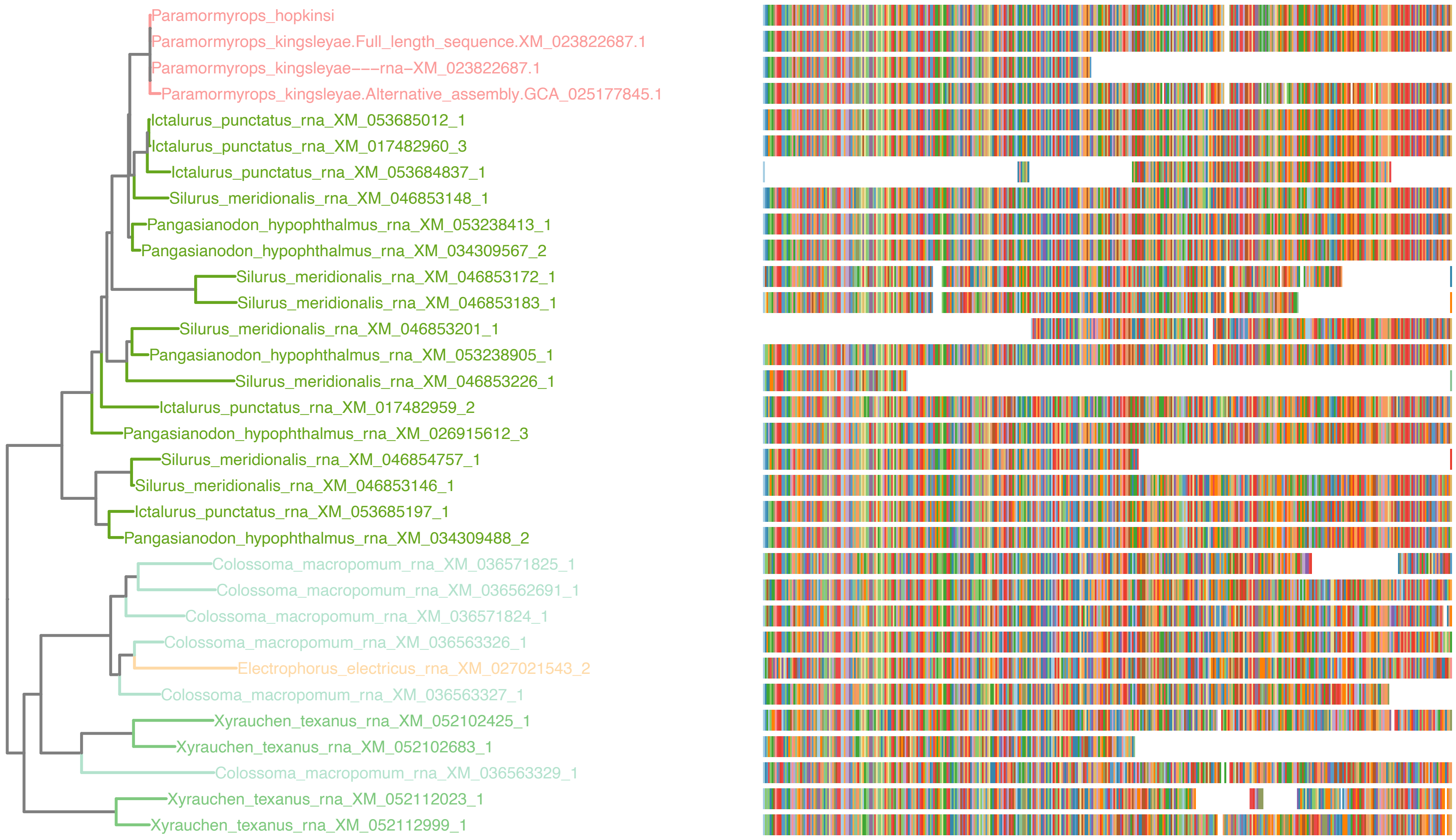

### N5.HOG0029633

#### dS distribution

Inter-species dS distribution

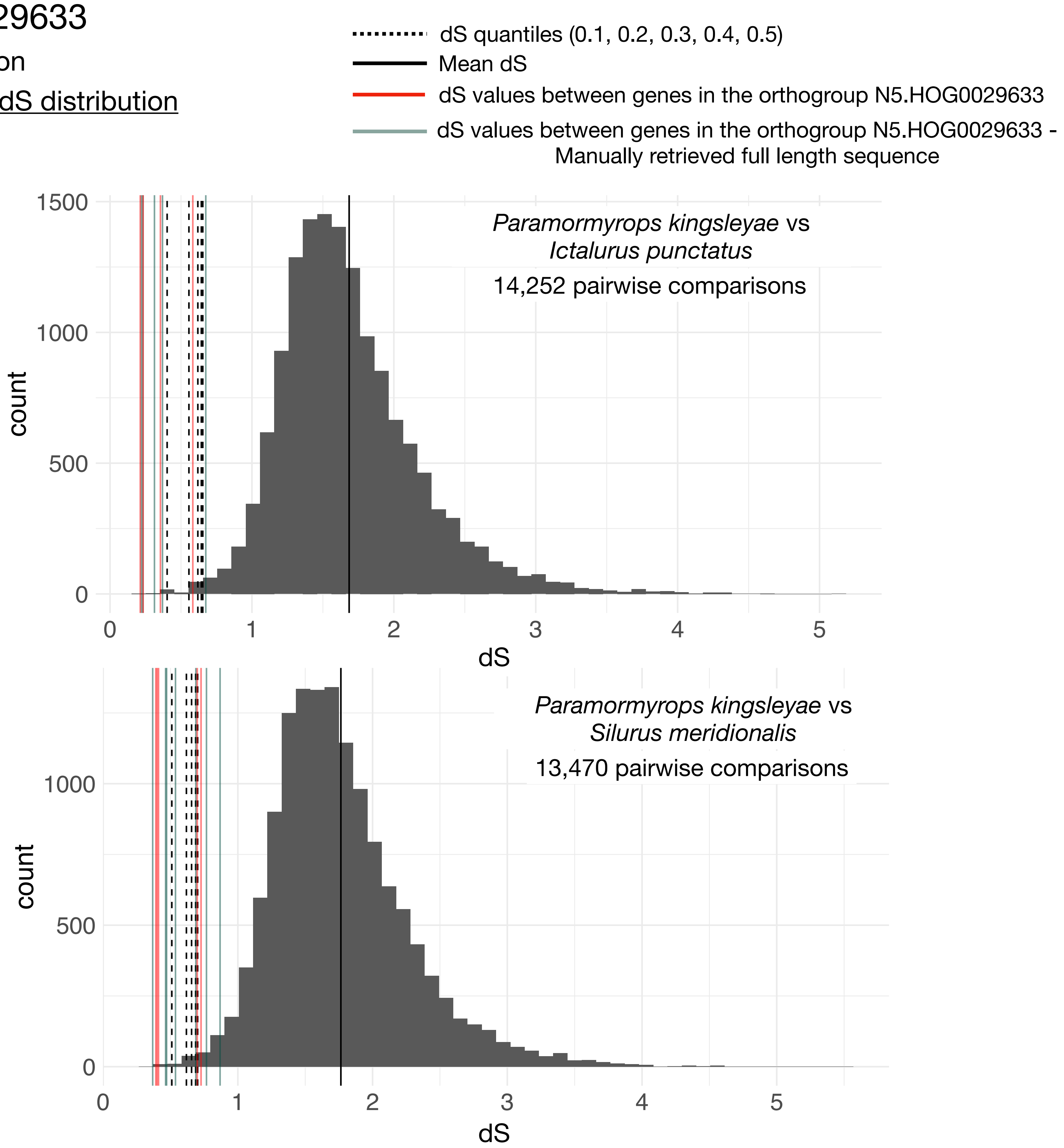


Intra-orthogroup dS

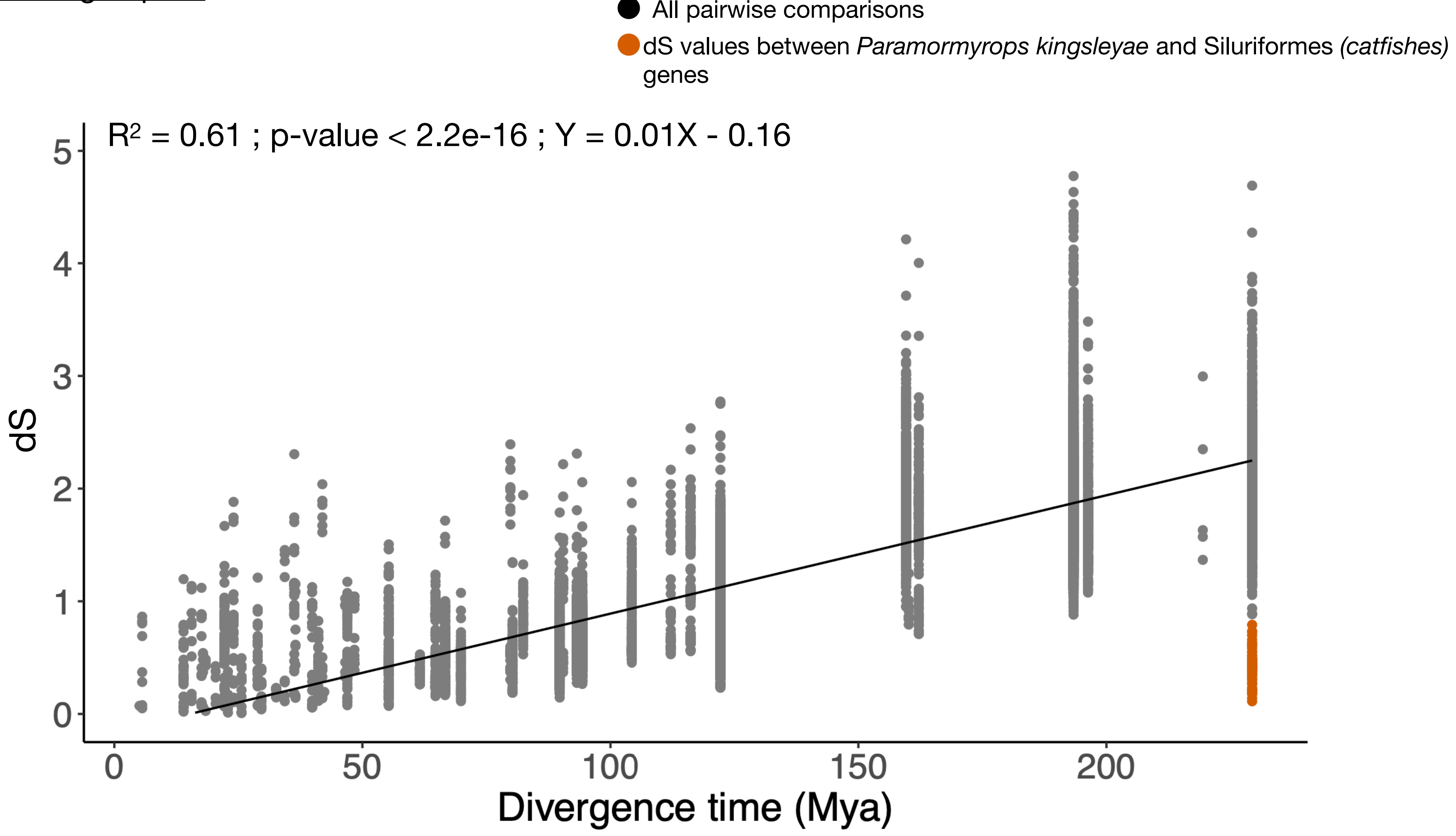

### N5.HOG0029633

#### Micro-synteny

- Genes in the orthogroup N5.HOG0029633
- Genes in other orthogroups
- Genes in the orthogroup N5.HOG0046344

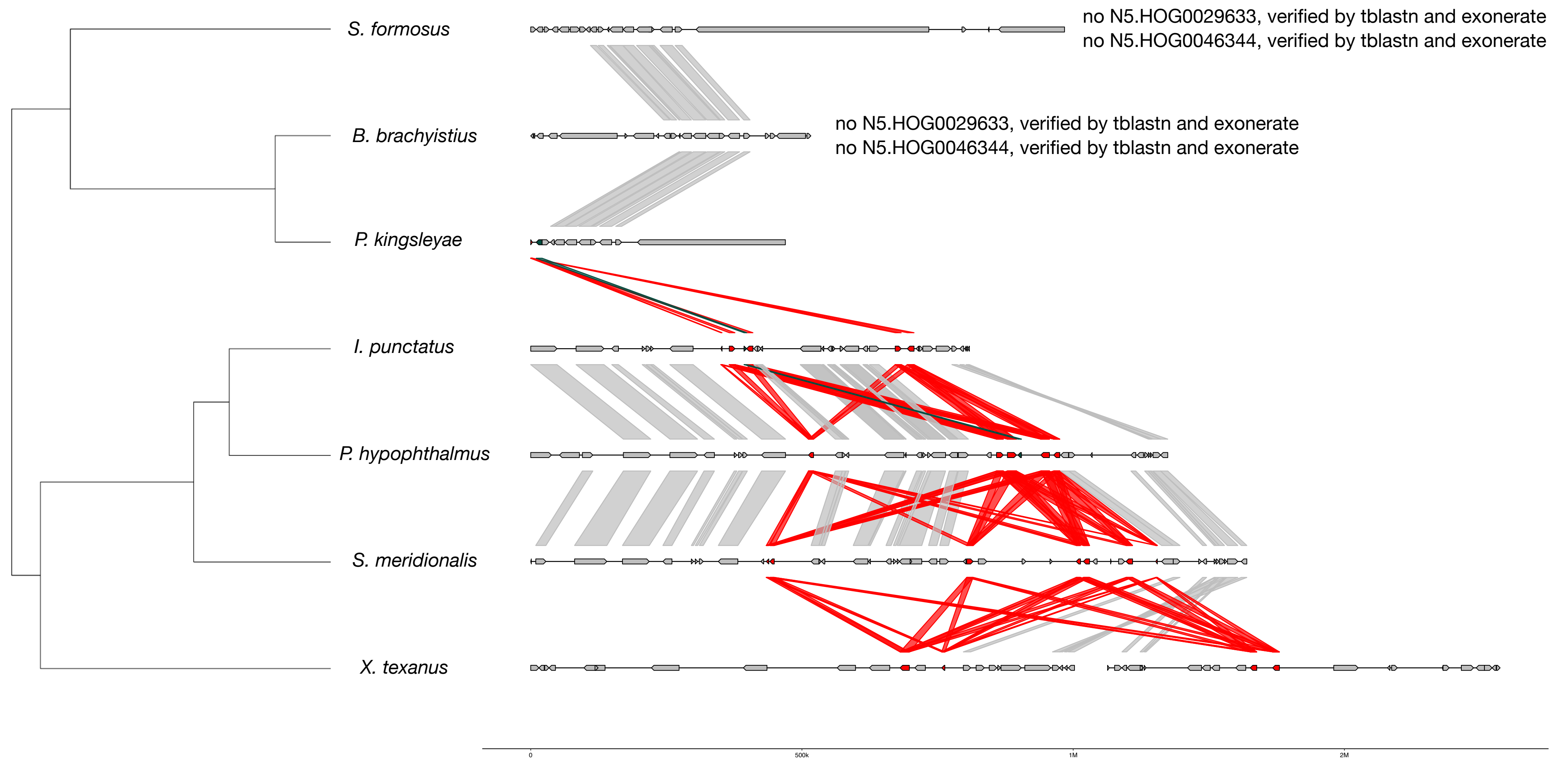


#### Zoom on *P. kingsleyae* and *I. punctatus*

- N5.HOG0029633 exons
- N5.HOG0046344 exons
- Non-shared transposable elements
- Shared transposable elements

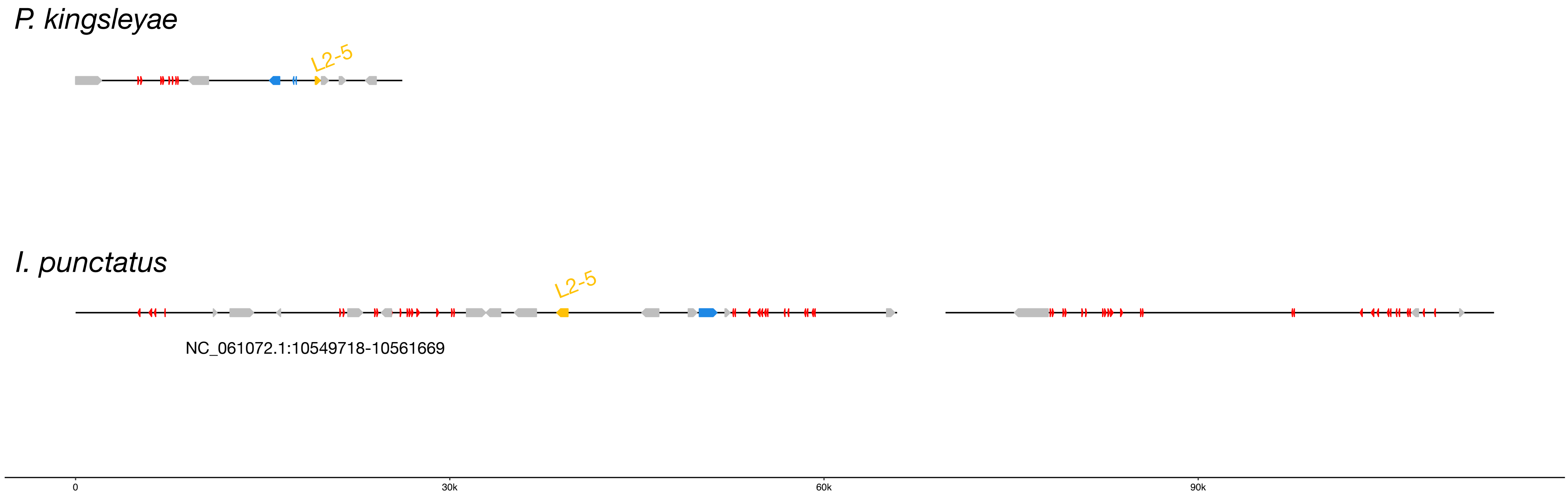


## L2-5 :

| | |
|---|---|
| *S. formosus* | *1241 copies scattered in the genome* |
| *B. brachyistius* | *492 copies scattered in the genome* |
| *P. kingsleyae* | *196 copies scattered in the genome* |
| *I. punctatus* | *1062 copies scattered in the genome* |
| *P. hypophthalmus* | *1426 copies scattered in the genome* |
| *S. meridionalis* | *2236 copies scattered in the genome* |
| *X. texanus* | *2672 copies scattered in the genome* |

The gene N5.HOG0046344 is probably a co-transfered gene. The gene phylogeny is similar to the one of N5.HOG0029633 , with *P. kingsleyae* clustering with catfishes and low dS values are also retrieved (just above the quantile 0.05)

N5.HOG0046344 gene tree:

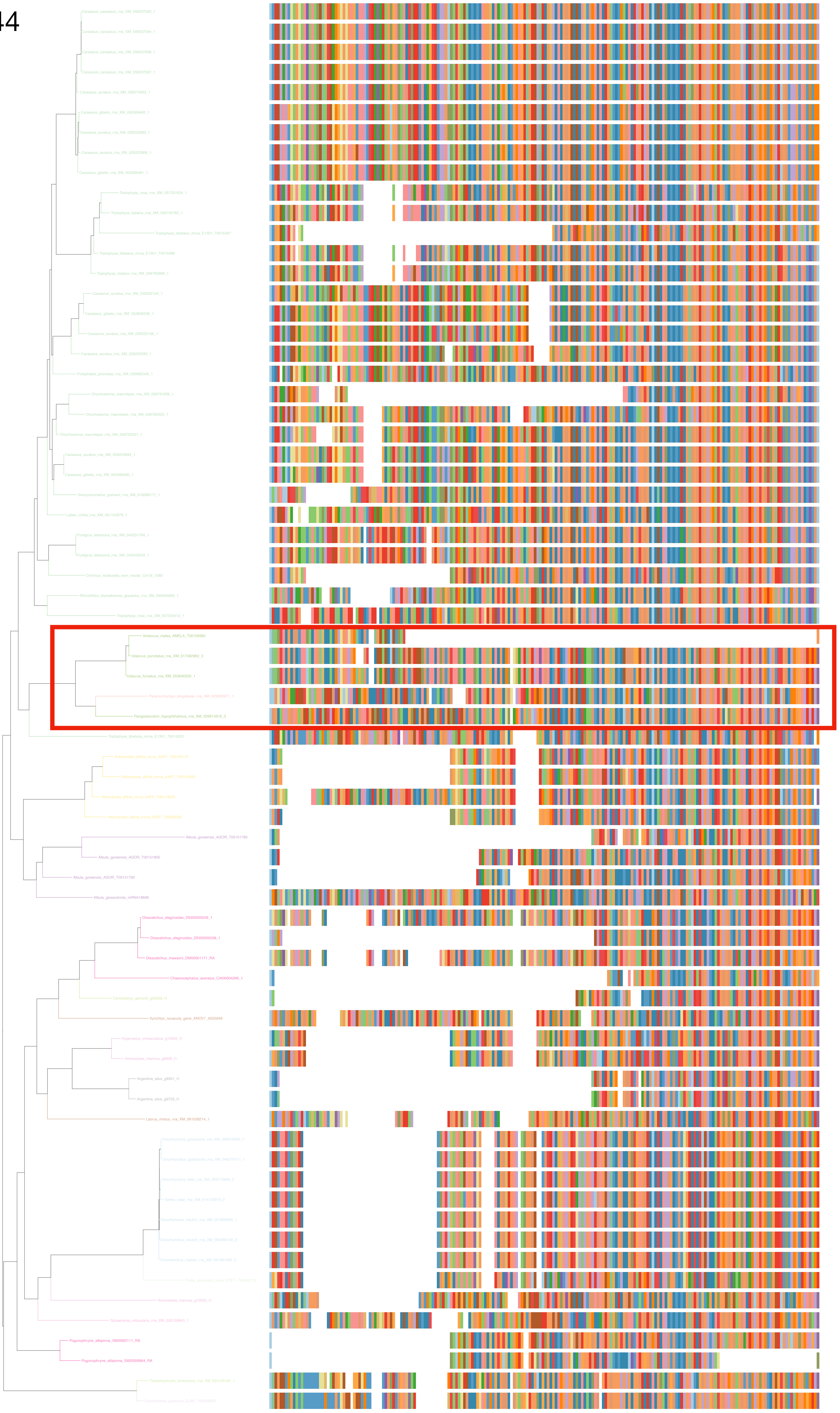

N5.HOG0046344

Zoom on HGT clade with bootstrap values

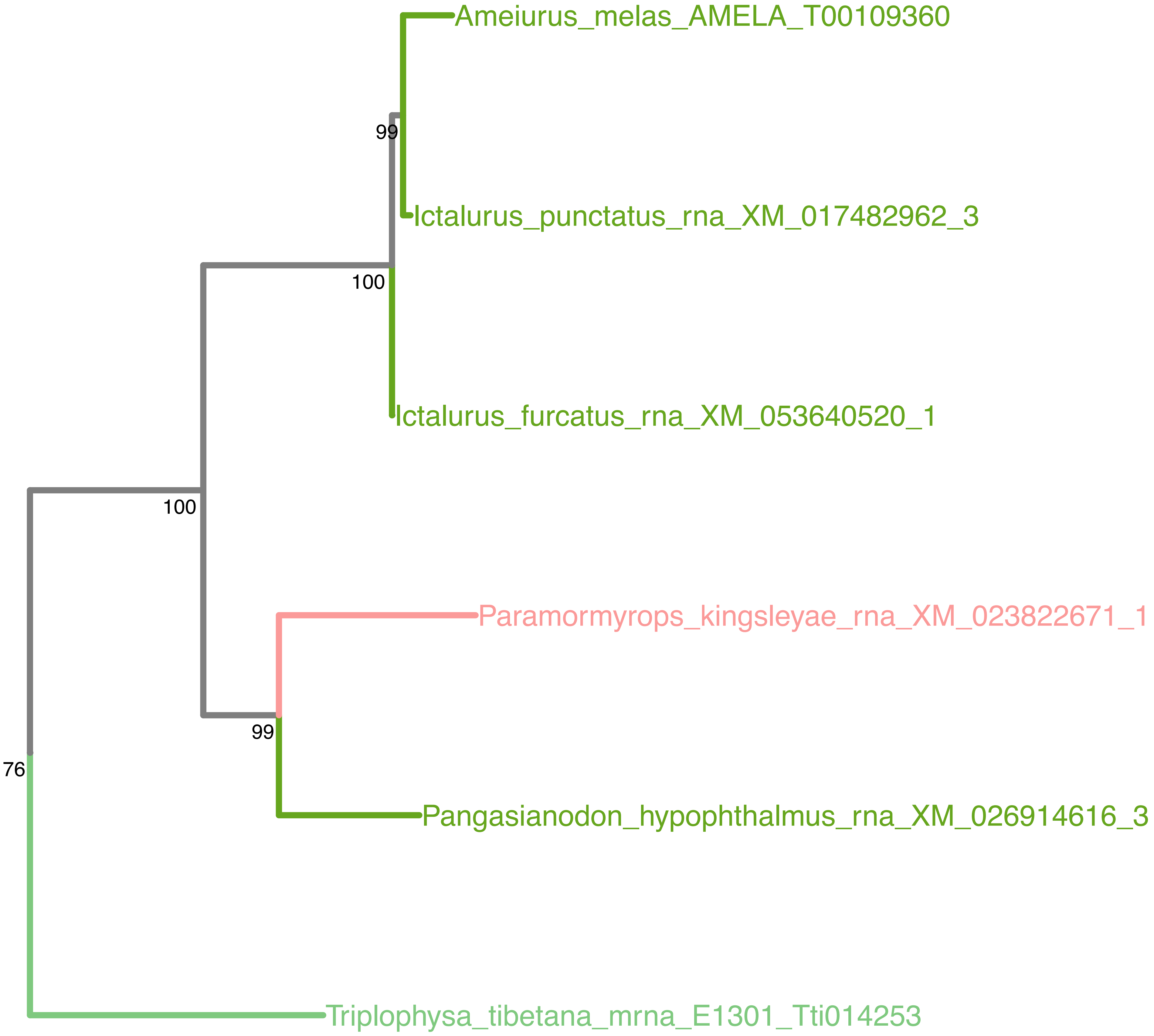

N5.HOG0046344

N5.HOG0046344 can also be retrieved entirely on an other genome assembly of *Paramormyrops kingsleyae* + on the genome of the closely related *Paramormyrops hopkinsi* (and are also located next to N5.HOG0029633)

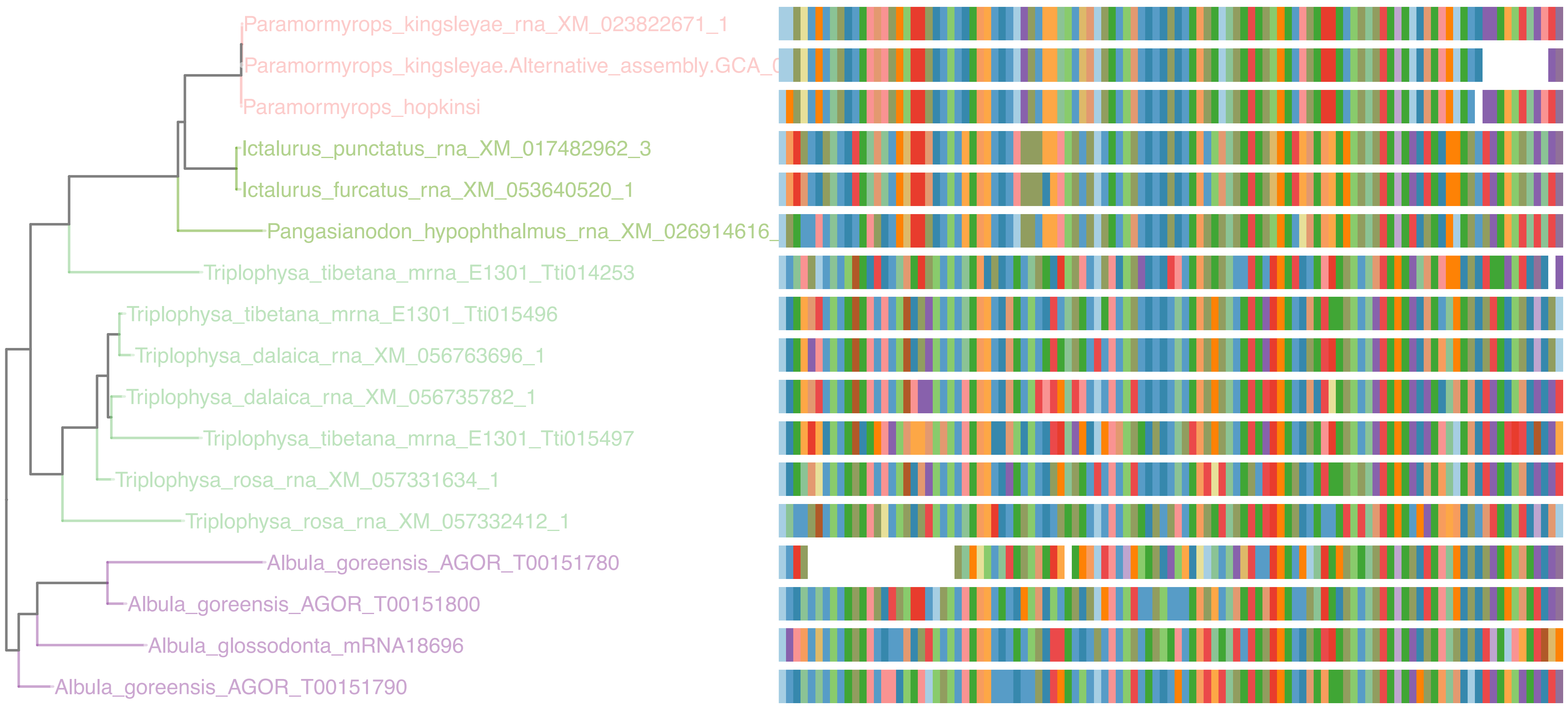

N5.HOG0046344

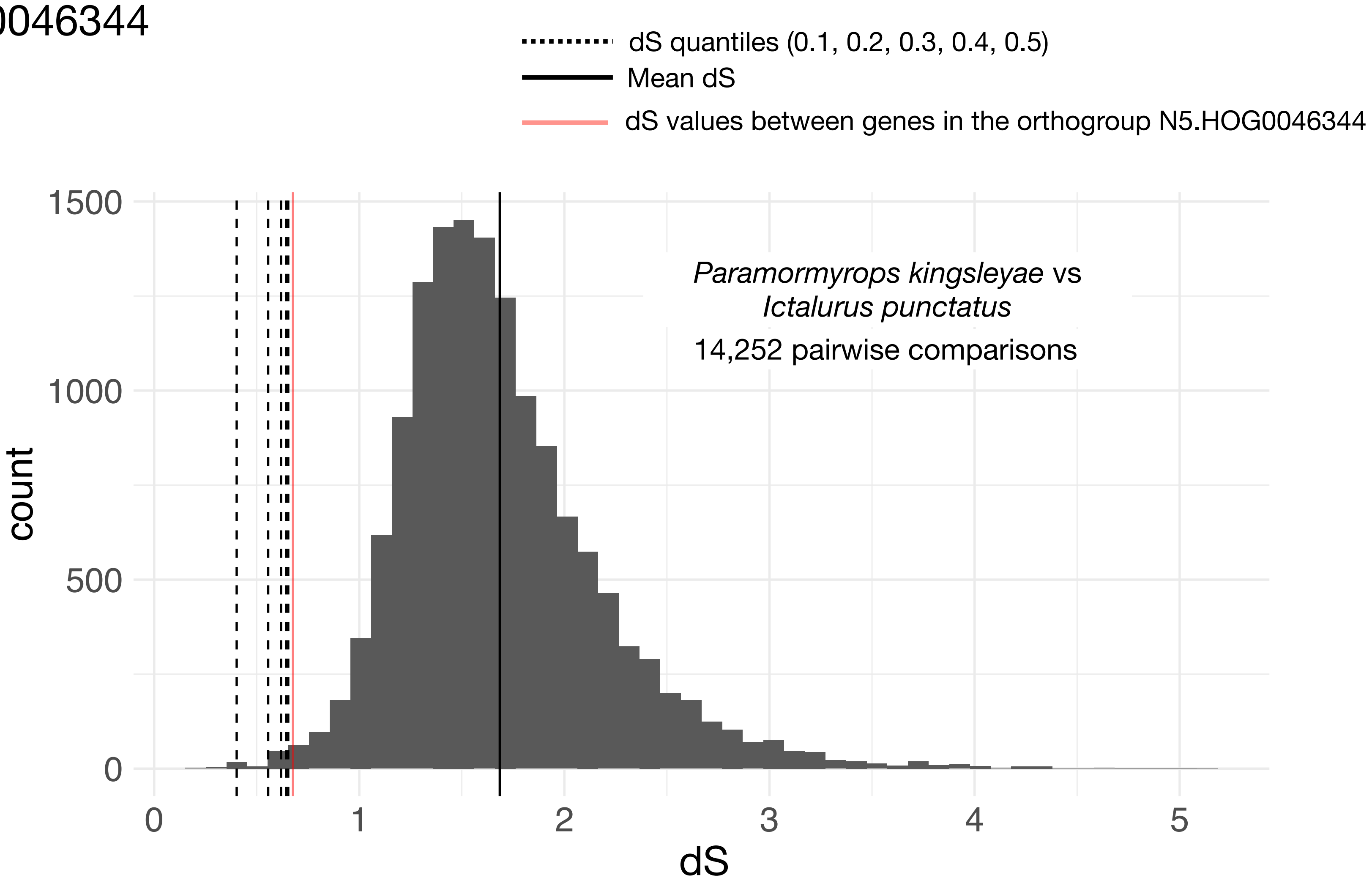

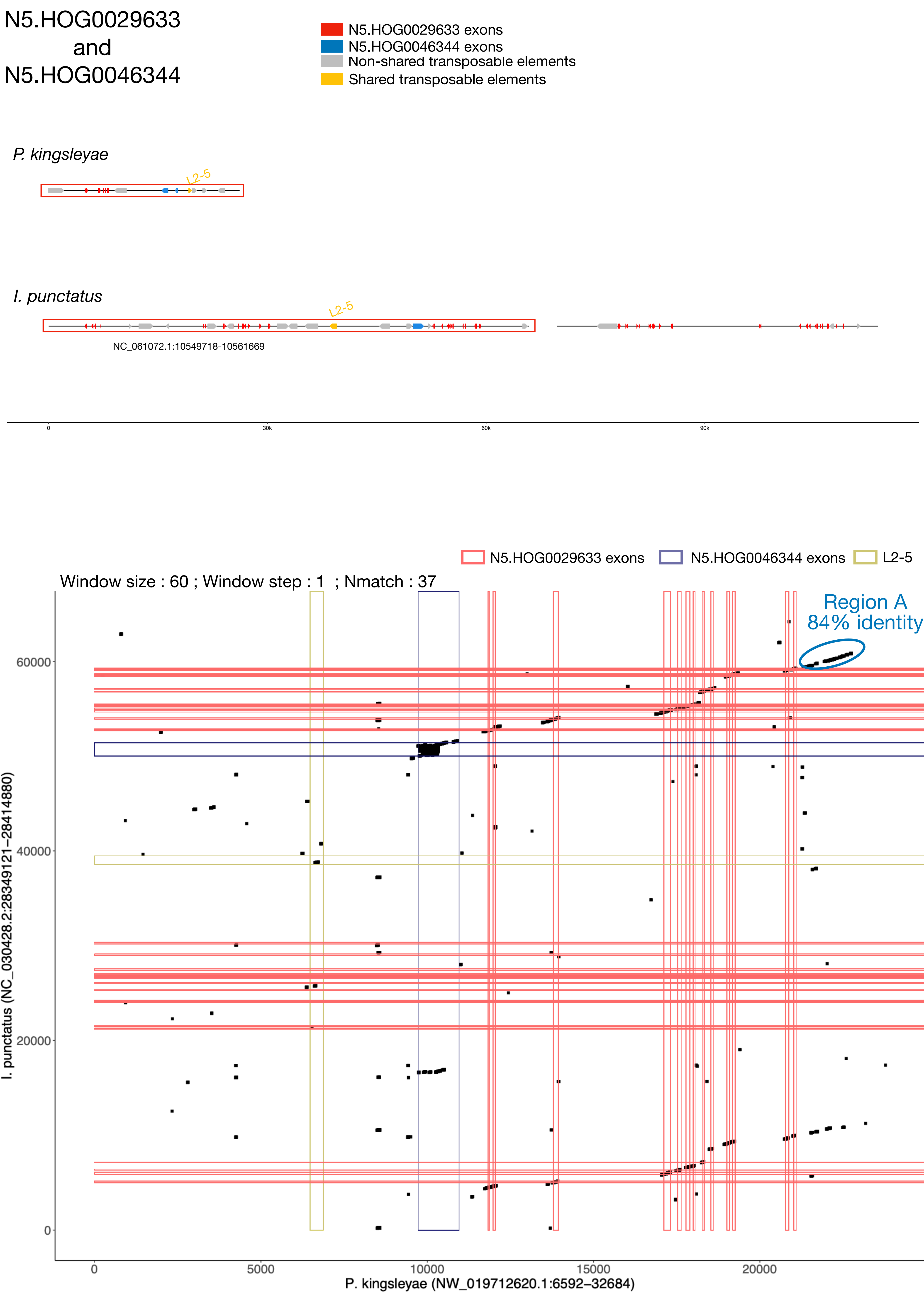
N5.HOG0029633
and
N5.HOG0046344
N5.HOG0029633 exons
N5.HOG0046344 exons
Non-shared transposable elements
Shared transposable elements
P. kingsleyae
L2-5
I. punctatus
L2-5
NC_061072.1:10549718-10561669
0
30k
60k
90k
N5.HOG0029633 exons
N5.HOG0046344 exons
L2-5
Window size : 60 ; Window step : 1 ; Nmatch : 37
Region A
84% identity
I. punctatus (NC_030428.2:28349121–28414880)
P. kingsleyae (NW_019712620.1:6592–32684)
0
20000
40000
60000
0
5000
10000
15000
20000

### N5.HOG0029633 and N5.HOG0046344

Blastn of conserved regions against all genome assemblies

#### Region A

| sseqid | pident | length | mismatch | gapopen | qstart | qend | sstart | send | evalue | bitscore |
|---|---|---|---|---|---|---|---|---|---|---|
| Paramormyrops_kingsleyae-NW_019712620.1 | 100.000 | 1708 | 0 | 0 | 1 | 1708 | 11592 | 9885 | 0.0 | 3155 |
| Pangasianodon_hypophthalmus-NC_069721.1 | 80.510 | 862 | 109 | 35 | 878 | 1697 | 25091173 | 25090329 | 8.75e-169 | 606 |
| Pangasianodon_hypophthalmus-NC_069721.1 | 78.491 | 623 | 87 | 29 | 610 | 1192 | 25072686 | 25072071 | 5.81e-96 | 364 |
| Pangasianodon_gigas-CM040465.1 | 79.977 | 864 | 113 | 35 | 878 | 1697 | 24716897 | 24716050 | 1.47e-161 | 582 |
| Pangasianodon_gigas-CM040465.1 | 78.528 | 652 | 91 | 30 | 587 | 1197 | 24697421 | 24696778 | 1.60e-101 | 383 |
| Pangasius_djambal-CM040986.1 | 78.196 | 876 | 103 | 44 | 878 | 1697 | 24587433 | 24586590 | 1.99e-130 | 479 |
| Pangasius_djambal-CM040986.1 | 77.518 | 854 | 125 | 39 | 899 | 1697 | 24567193 | 24566352 | 4.34e-122 | 451 |
| Neoarius_graeffei-NC_083582.1 | 81.409 | 511 | 71 | 13 | 981 | 1479 | 4443801 | 4444299 | 2.06e-105 | 396 |
| Neoarius_graeffei-NC_083582.1 | 81.202 | 516 | 72 | 13 | 981 | 1485 | 3973940 | 3973439 | 2.67e-104 | 392 |
| Neoarius_graeffei-NC_083582.1 | 80.943 | 509 | 77 | 12 | 981 | 1479 | 4248365 | 4248863 | 4.46e-102 | 385 |
| Neoarius_graeffei-NC_083582.1 | 80.980 | 510 | 72 | 13 | 981 | 1479 | 4507979 | 4508474 | 5.77e-101 | 381 |
| Hemibagrus_wyckioides-NC_080722.1 | 84.426 | 244 | 30 | 6 | 1002 | 1244 | 1695812 | 1696048 | 1.71e-56 | 233 |
| Silurus_meridionalis-NC_060890.1 | 75.103 | 486 | 86 | 24 | 1009 | 1473 | 26728224 | 26727753 | 8.07e-45 | 195 |
| Tachysurus_vachellii-NC_083461.1 | 86.364 | 154 | 20 | 1 | 1025 | 1177 | 2028142 | 2028295 | 1.76e-36 | 167 |
| Tachysurus_vachellii-NC_083461.1 | 86.364 | 154 | 20 | 1 | 1025 | 1177 | 2113873 | 2114026 | 1.76e-36 | 167 |
| Pangasius_djambal-CM040975.1 | 87.500 | 104 | 9 | 2 | 476 | 577 | 33099338 | 33099237 | 1.80e-21 | 117 |

N5.HOG0029633
and
N5.HOG0046344

MRCA of the donor and recipient clade

Independent losses needed in an vertical transmission hypothesis

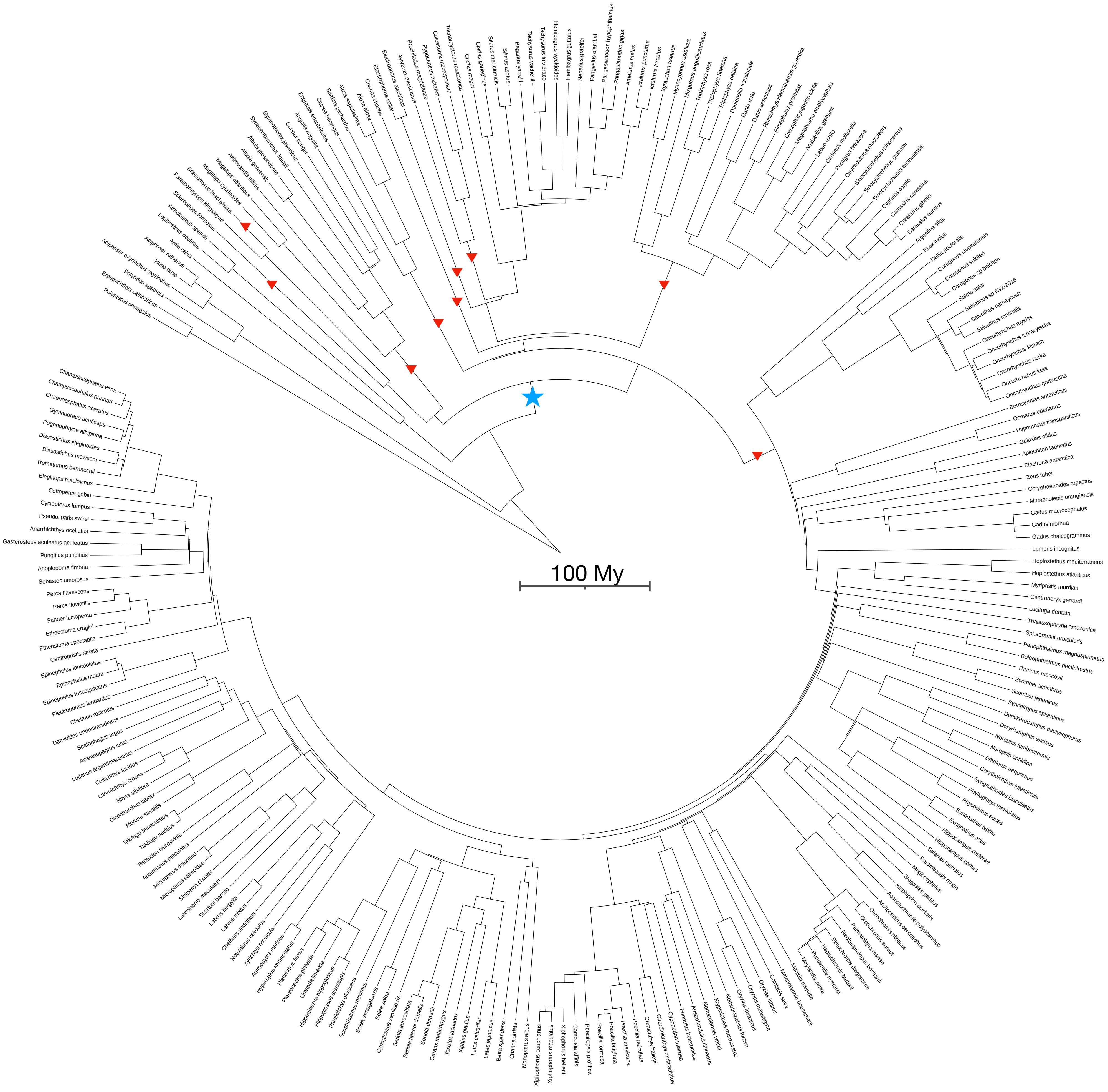

N5.HOG0010622

### N5.HOG0010622

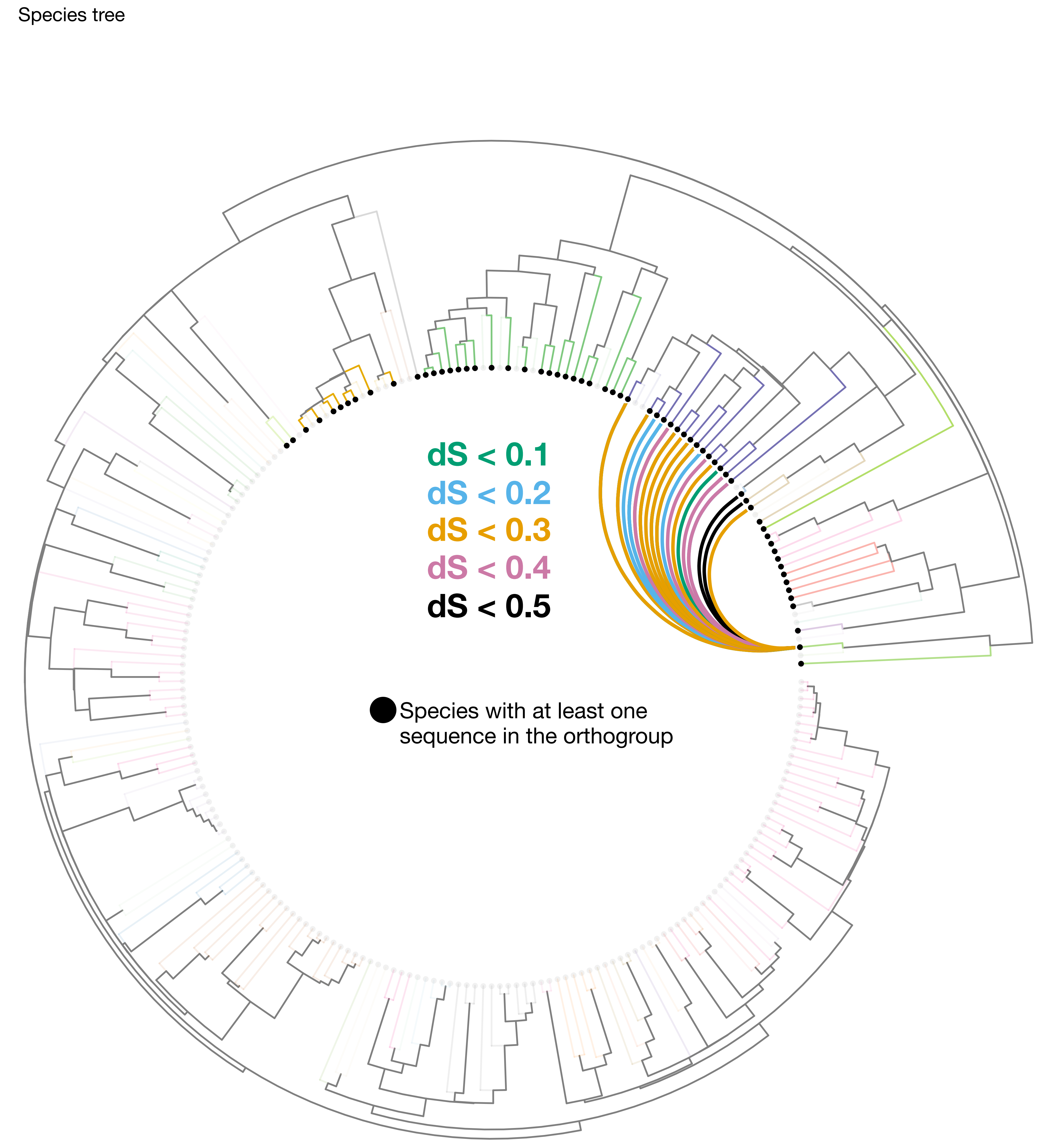

### N5.HOG0010622

Orthogroup phylogeny
and
protein alignment

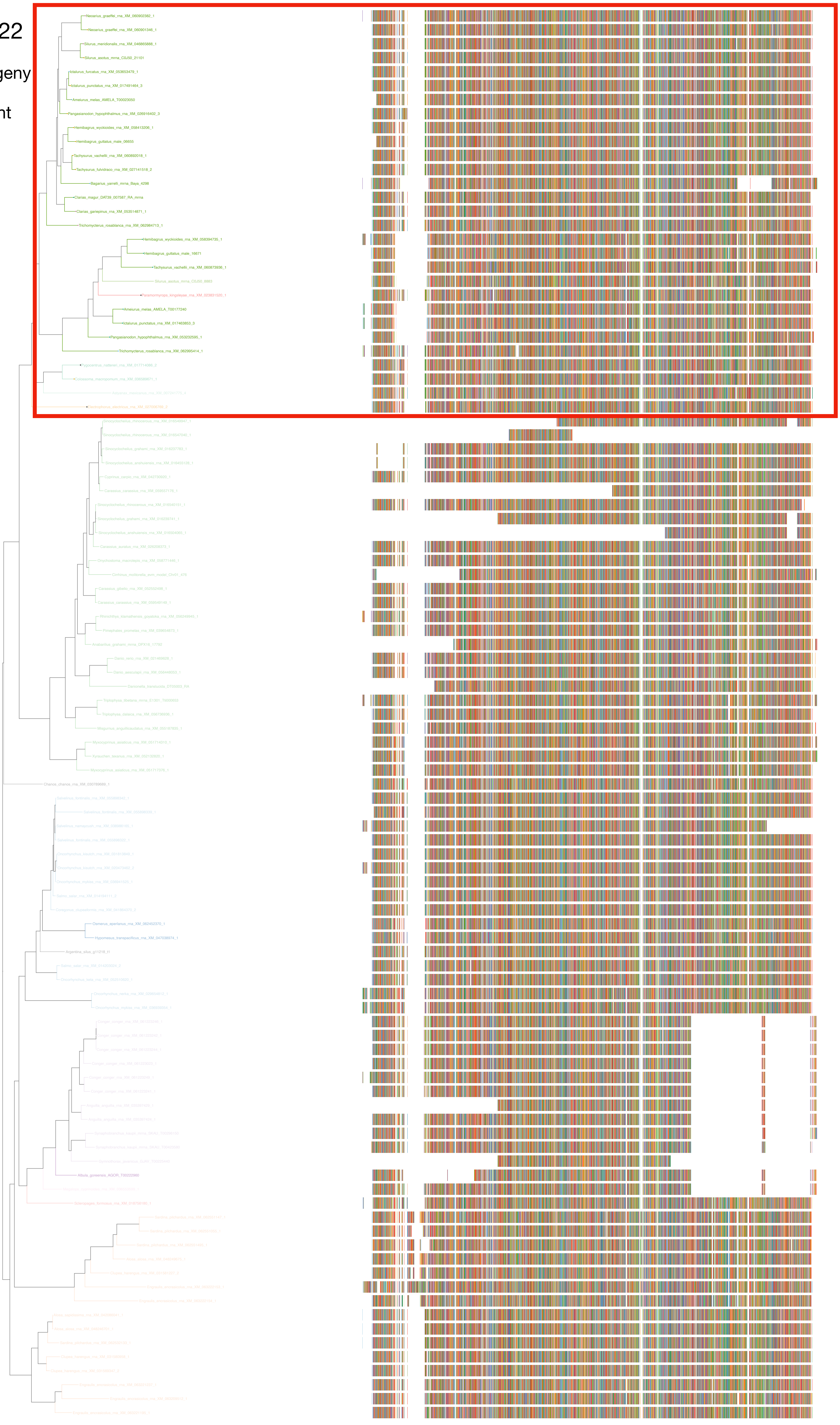

N5.HOG0010622

Zoom on HGT clade with bootstrap values

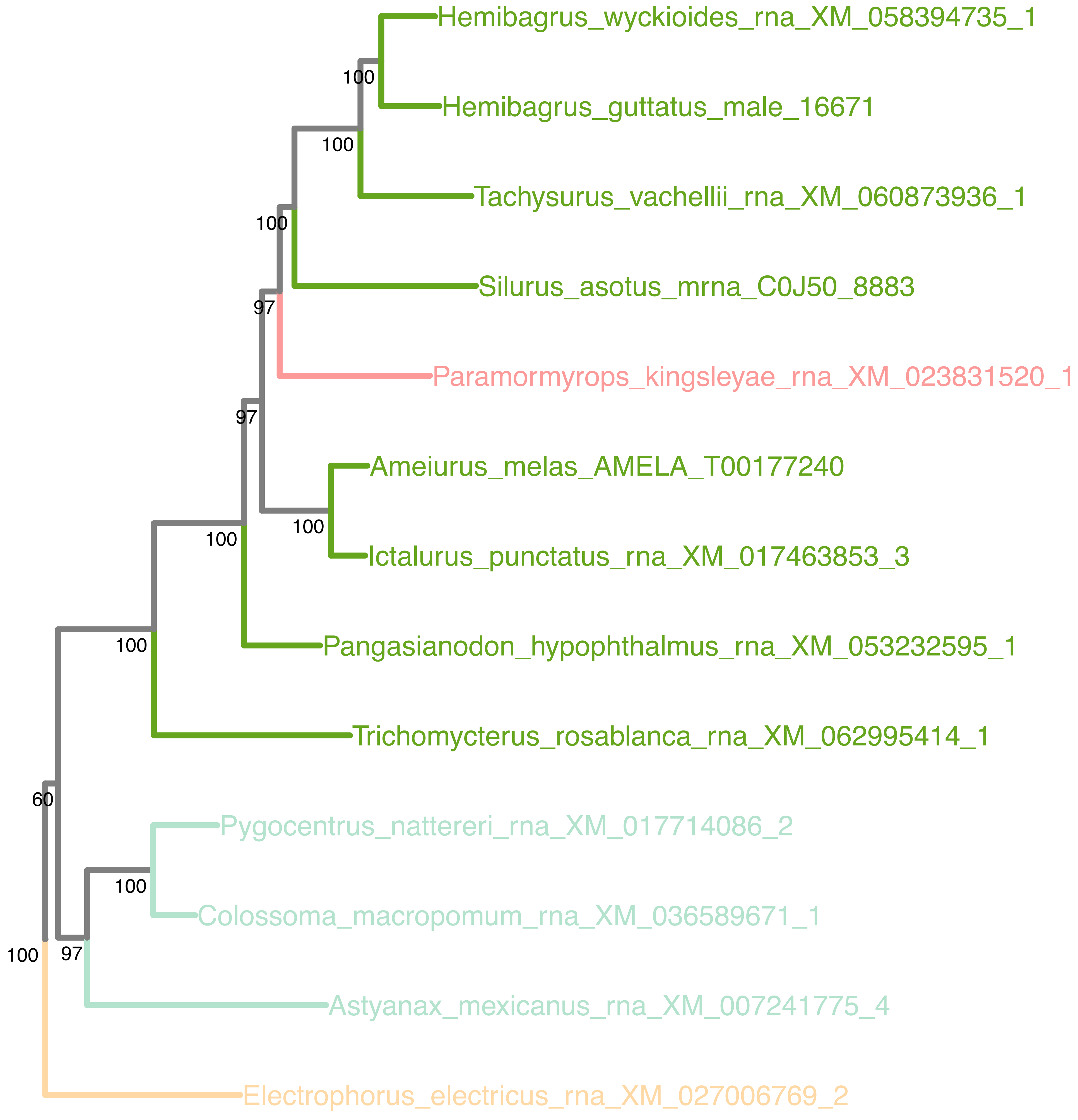

### N5.HOG0010622

Orthogroup phylogeny
and
trimmed
protein alignment

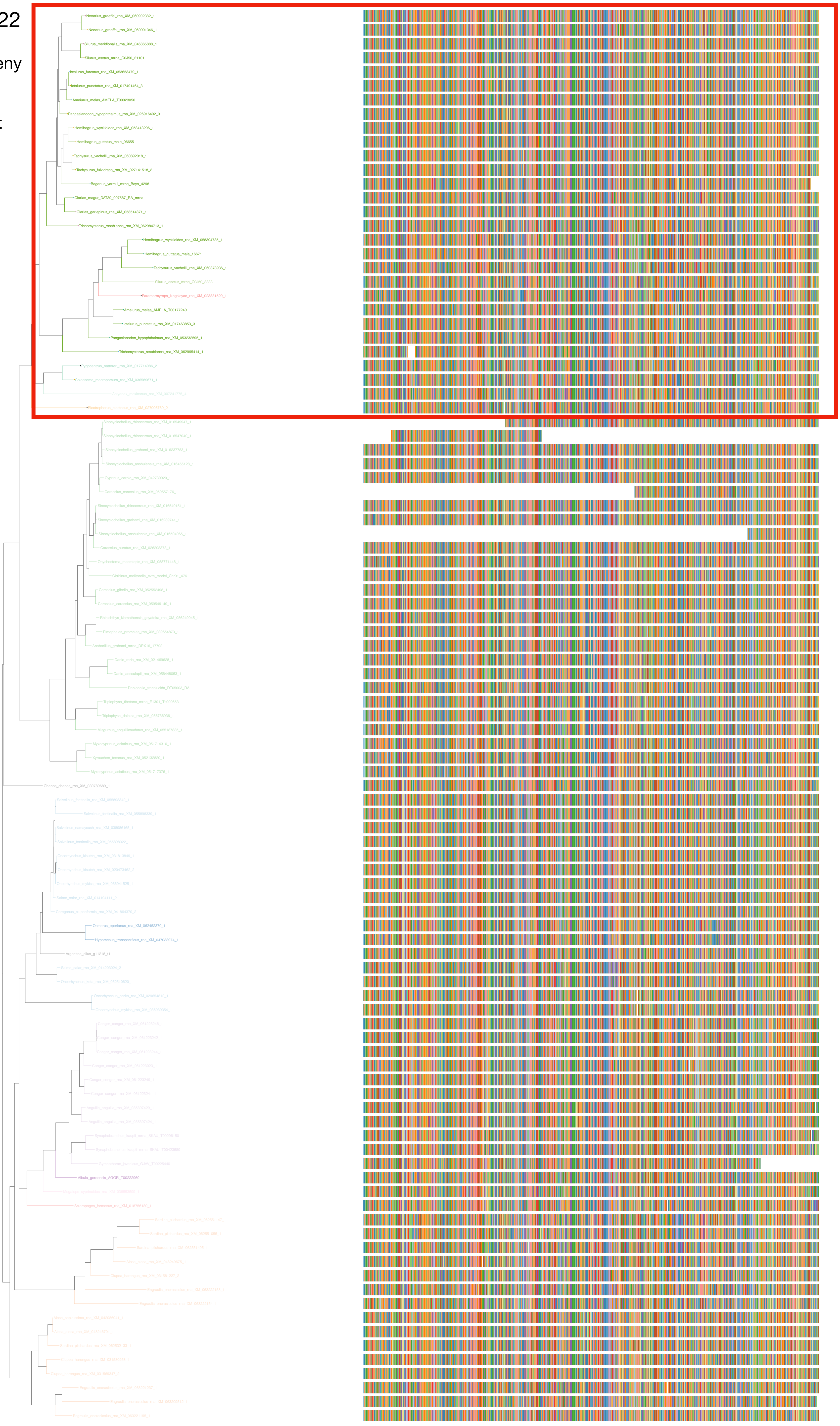

### N5.HOG0010622

« Best-match » phylogeny
and
protein alignment

50 best match against ray-finned fishes protein databases
10 best match against non-fish proteins from UniProt

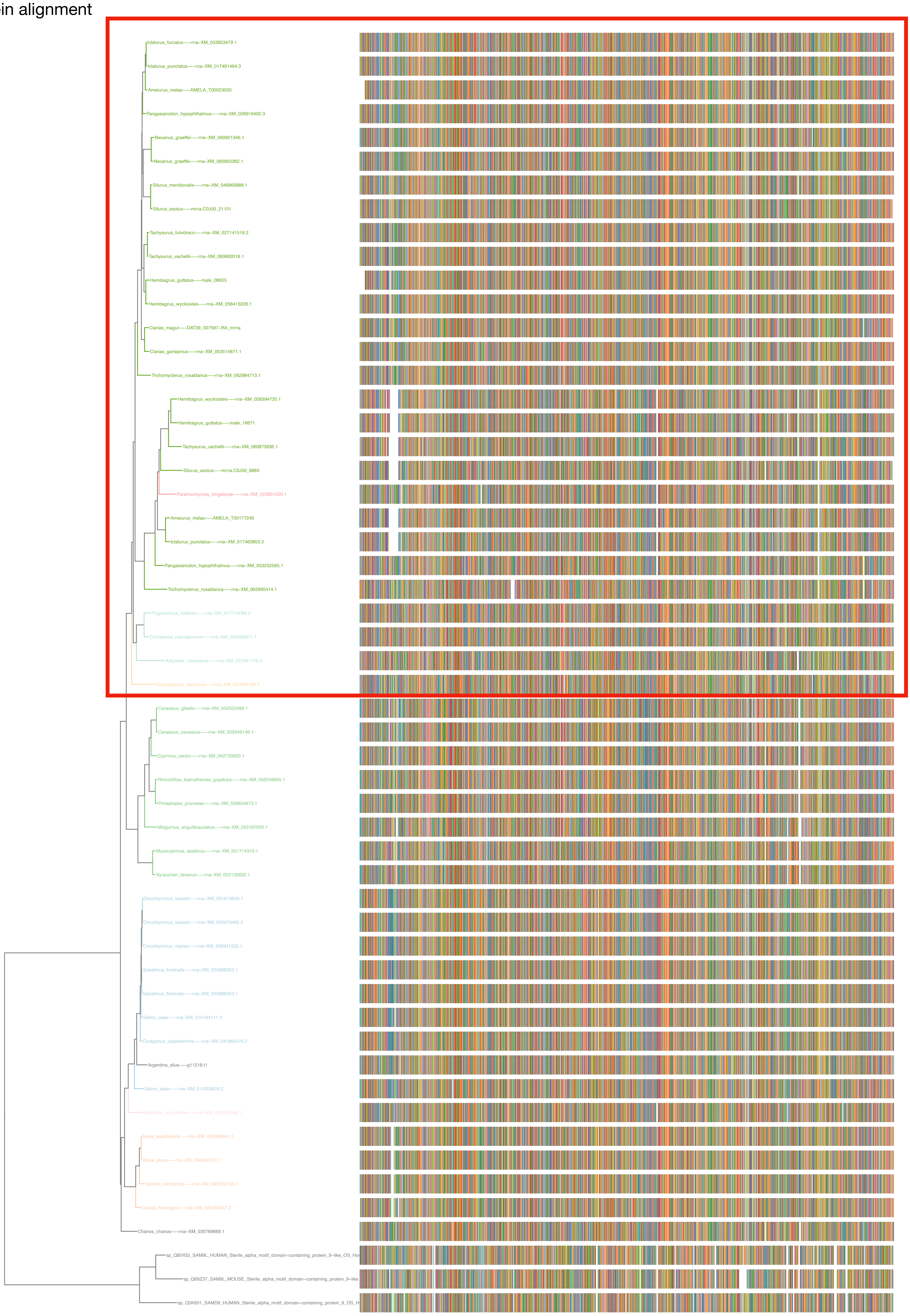

### N5.HOG0010622

dS distribution

Inter-species dS distribution

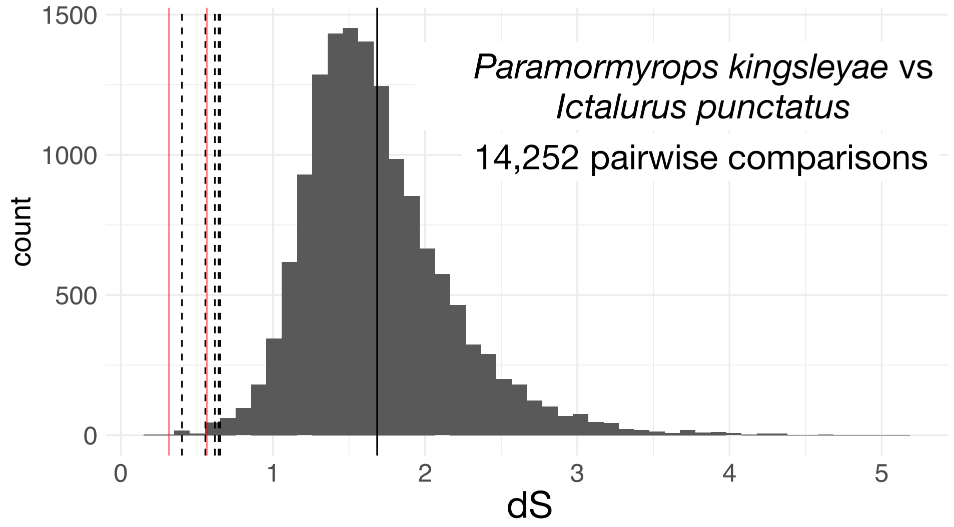


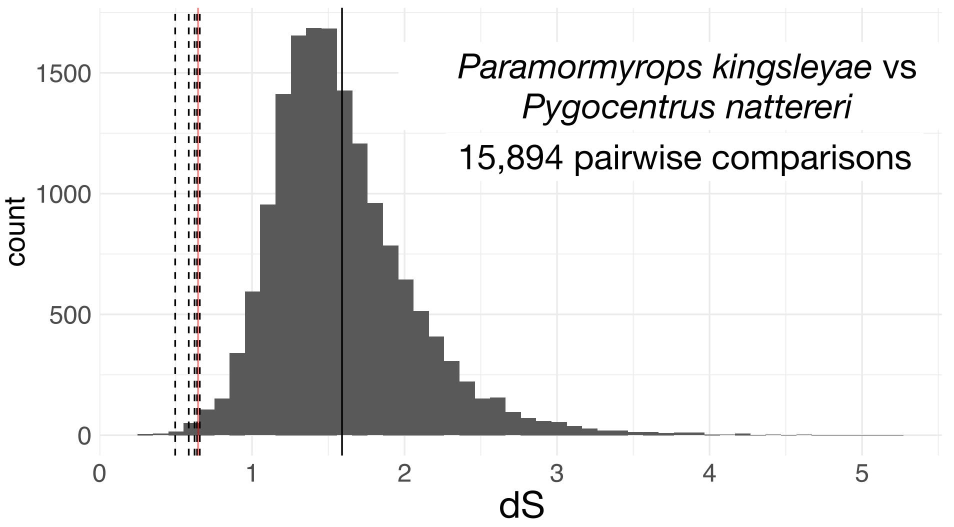


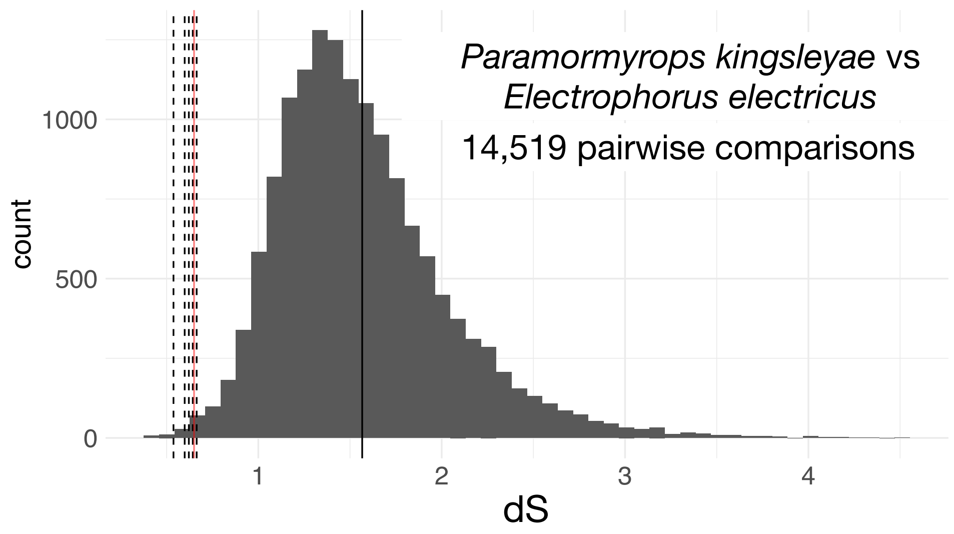


Intra-orthogroup dS

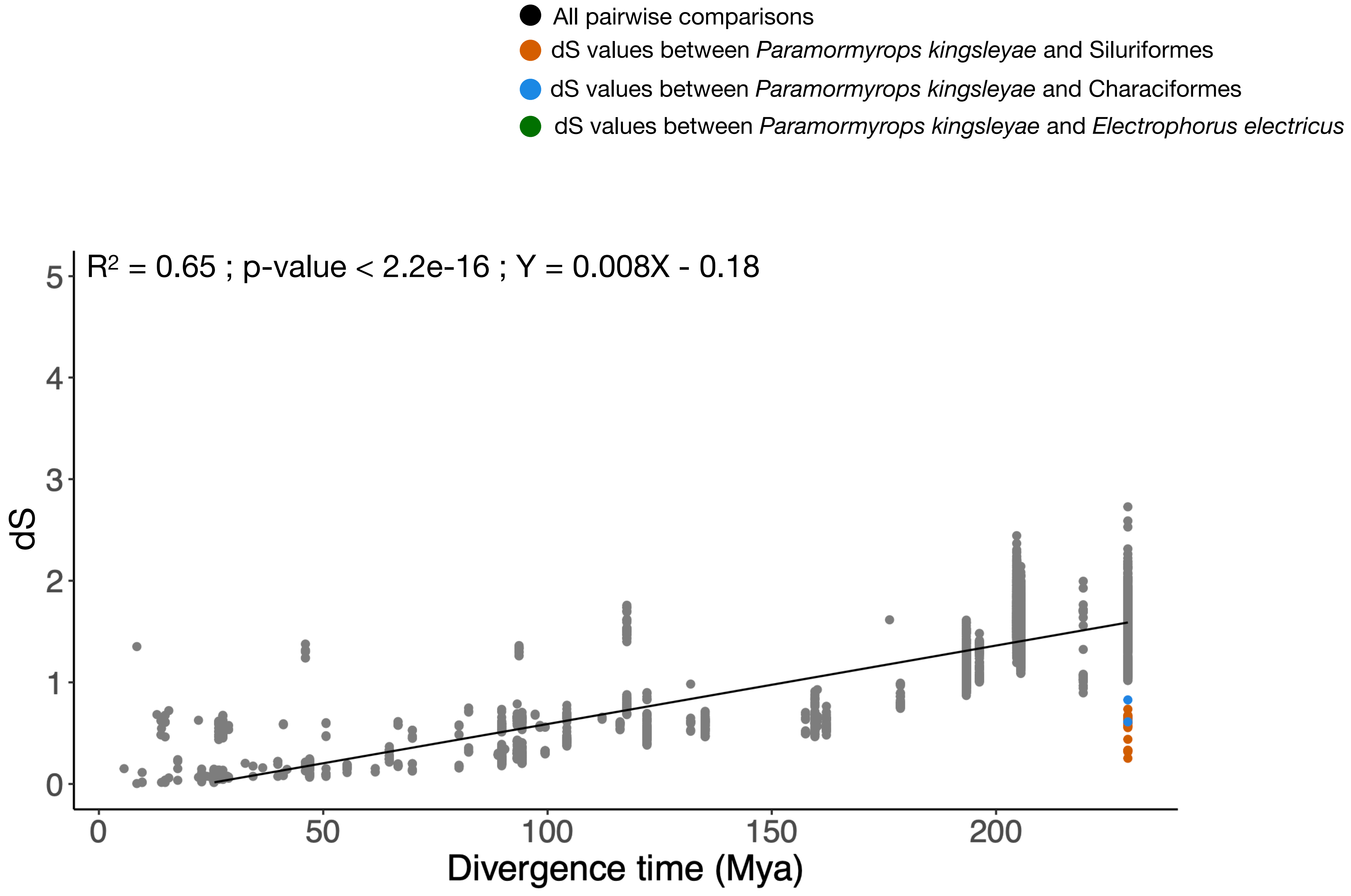

N5.HOG0010622

The gene can also be retrieved entirely on an other genome assembly of *Paramormyrops kingsleyae* + on the genome of the closely related *Paramormyrops hopkinsi*

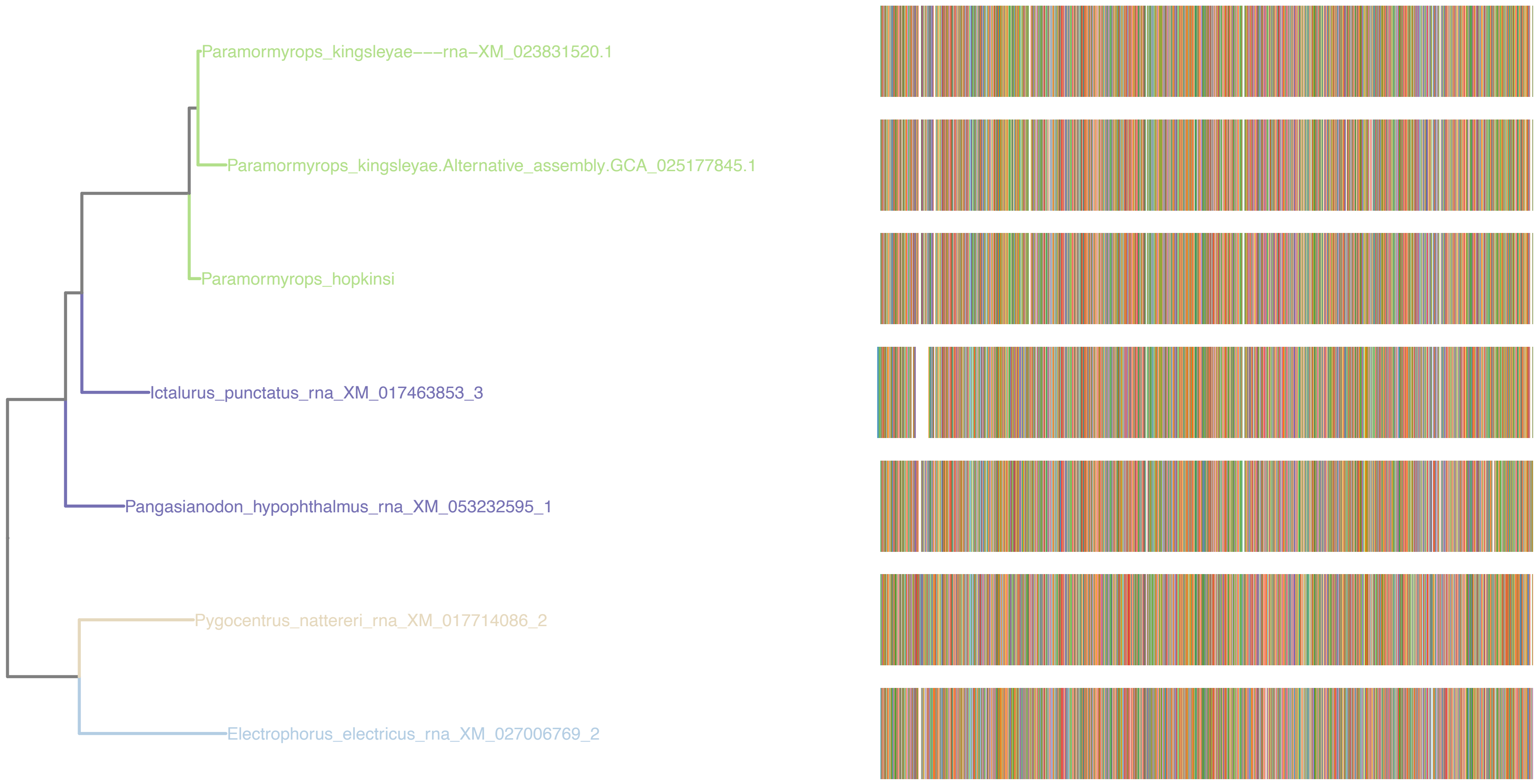

### N5.HOG0010622

#### Micro-synteny

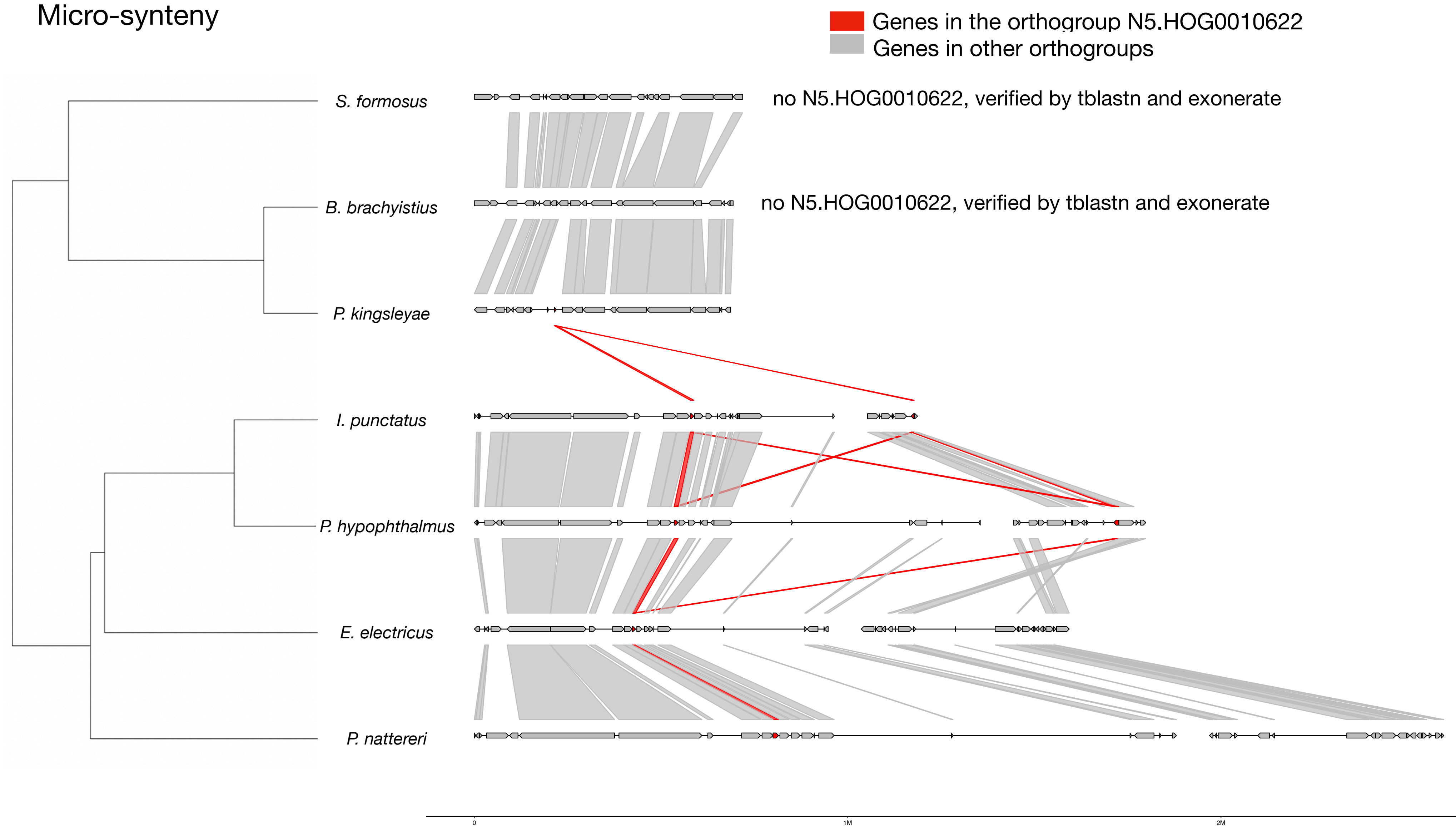


#### <u>Zoom on *P. kingsleyae* and *I. punctatus*</u>

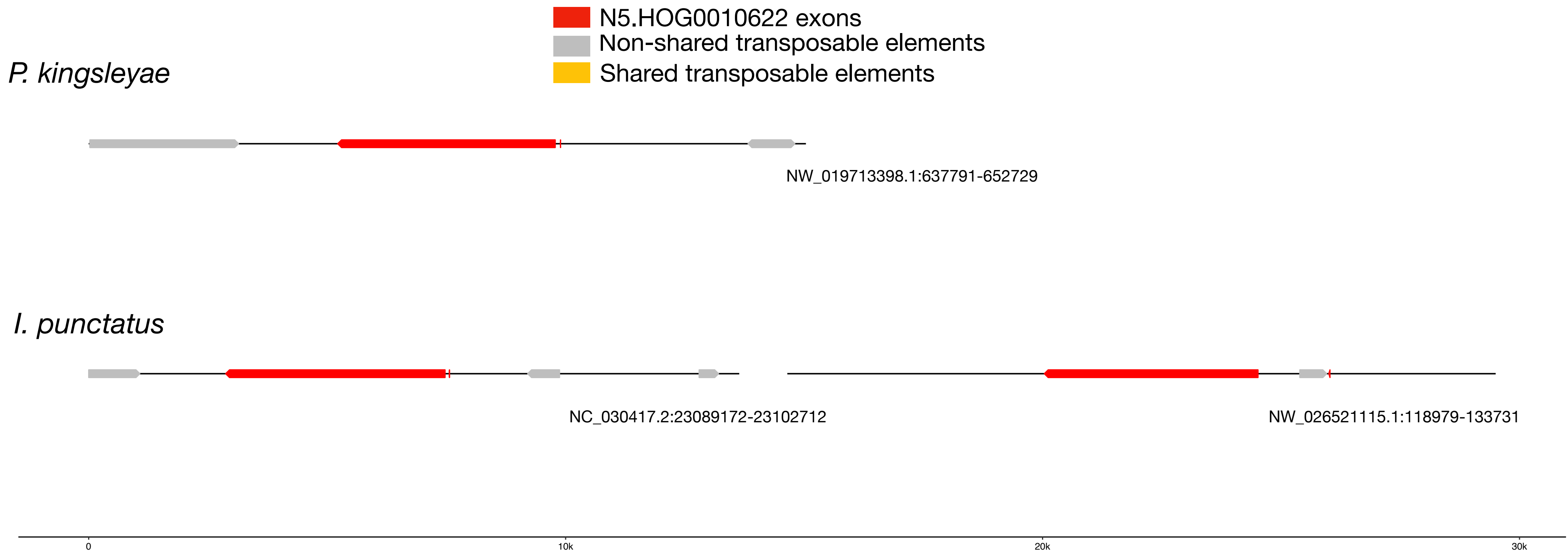

### N5.HOG0010622

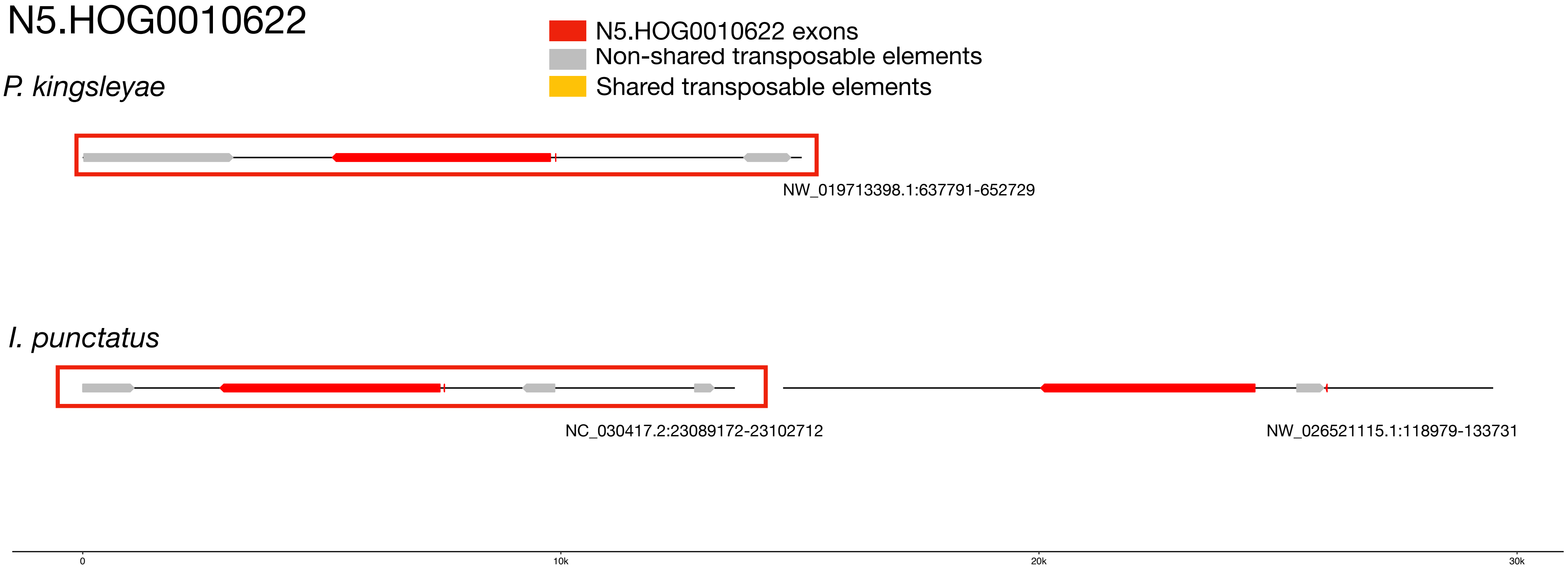


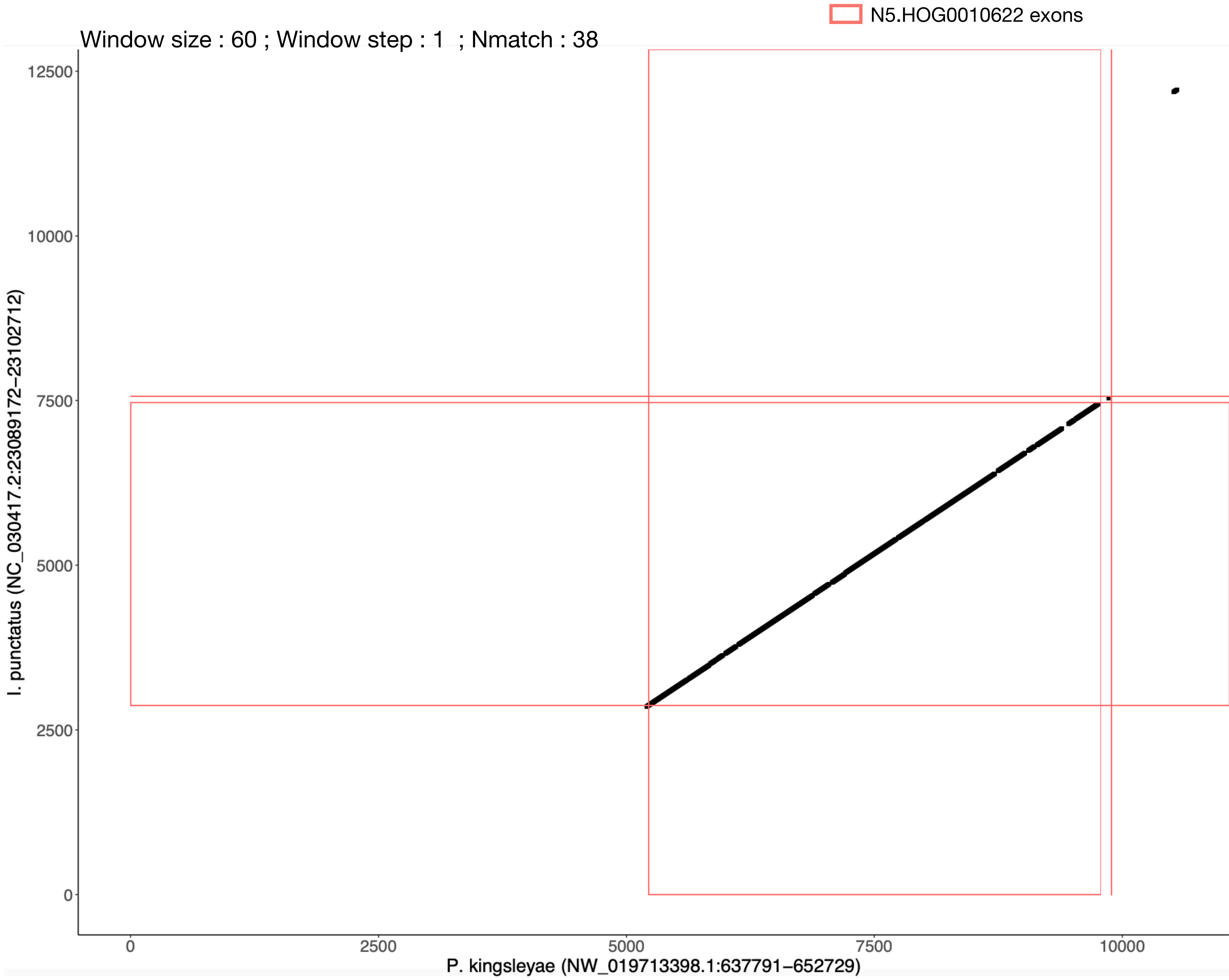

### N5.HOG0010622

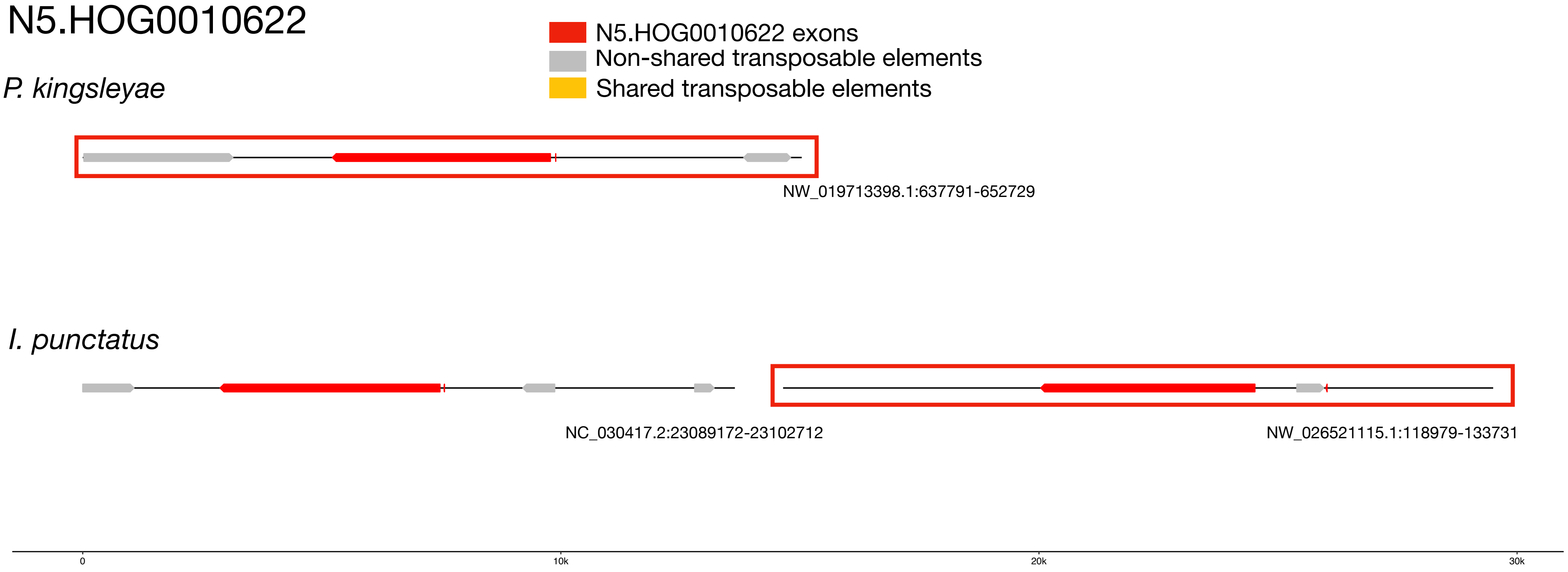


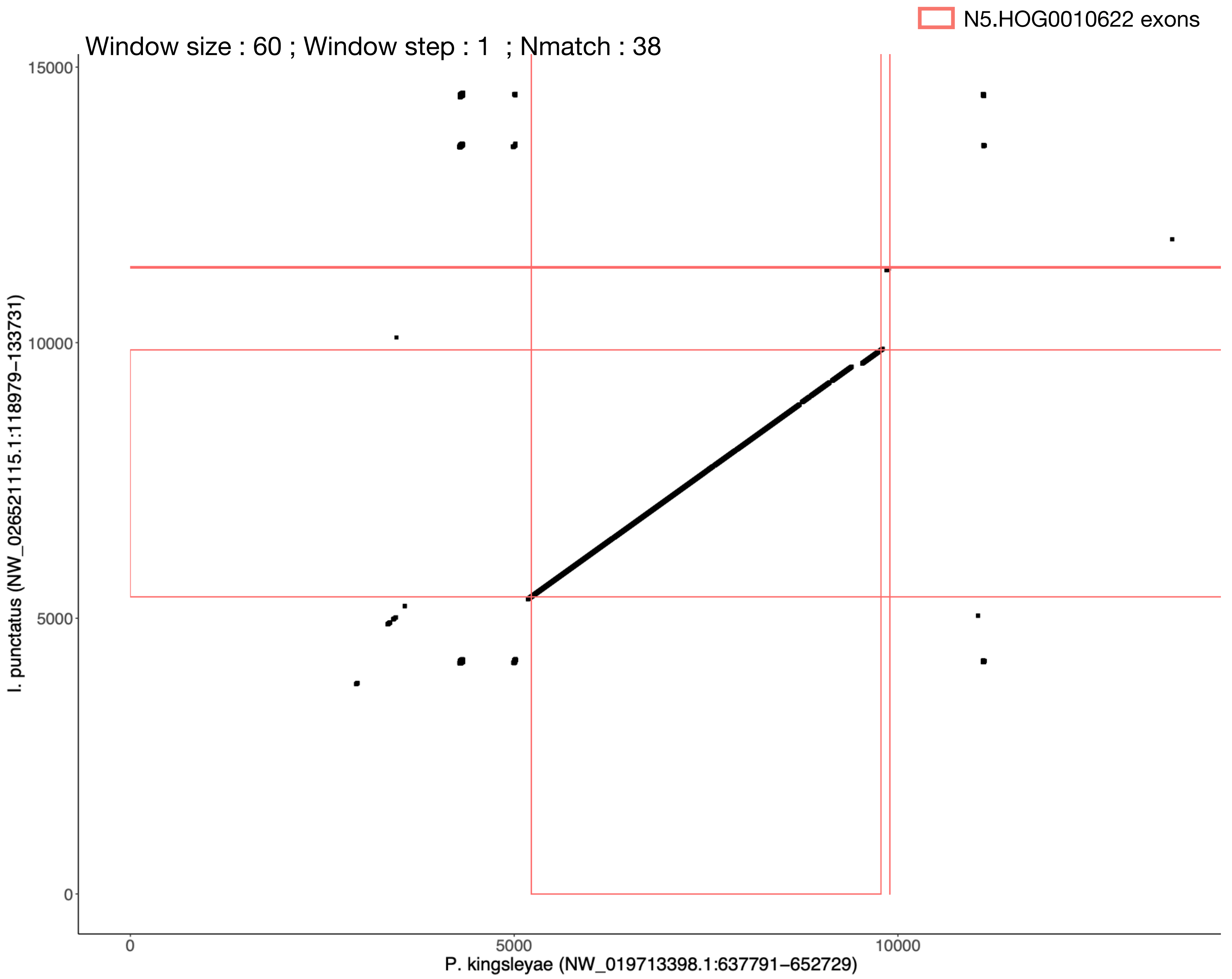

N5.HOG0010622

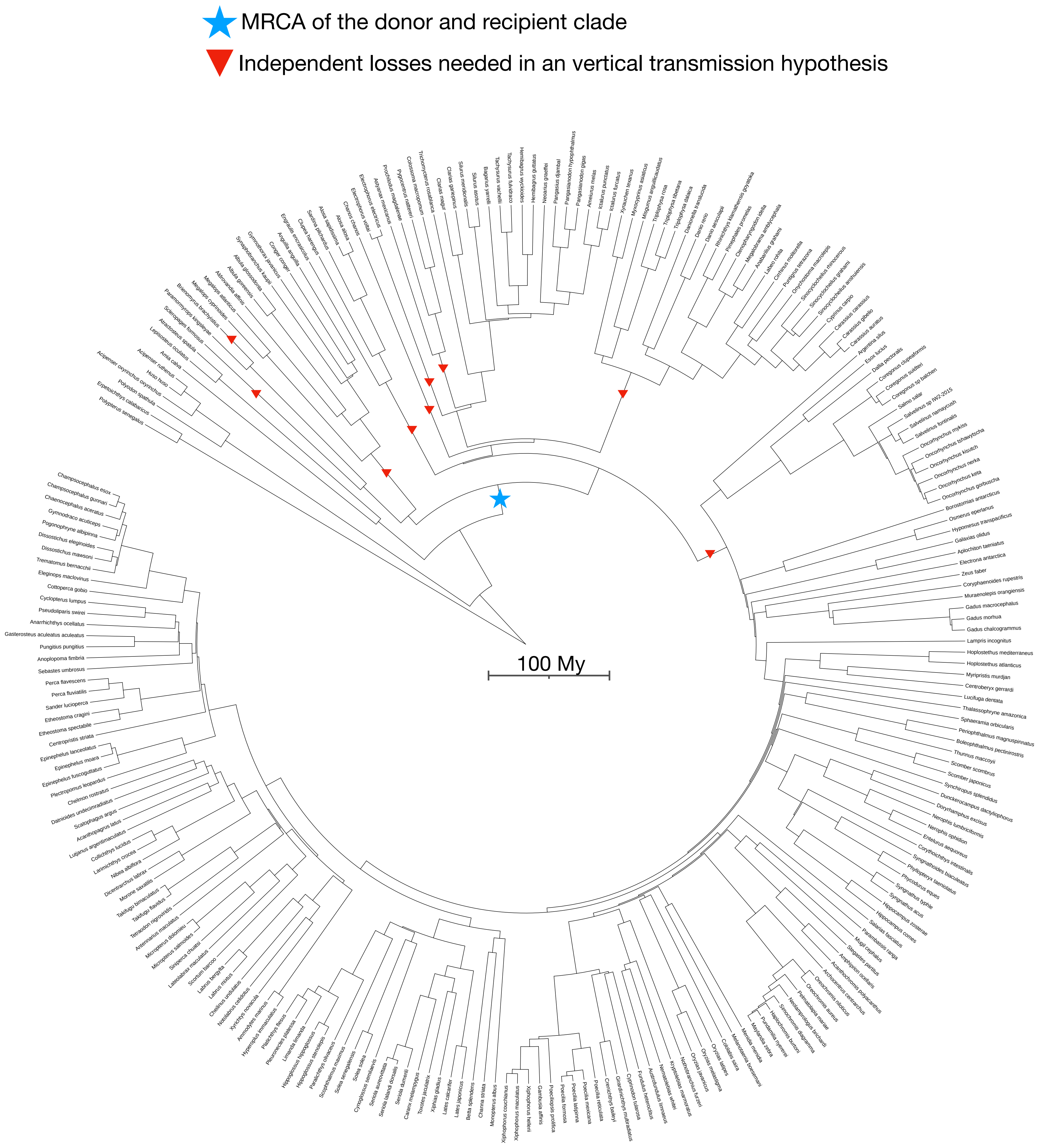

N5.HOG0013500

N5.HOG0013500

Species tree

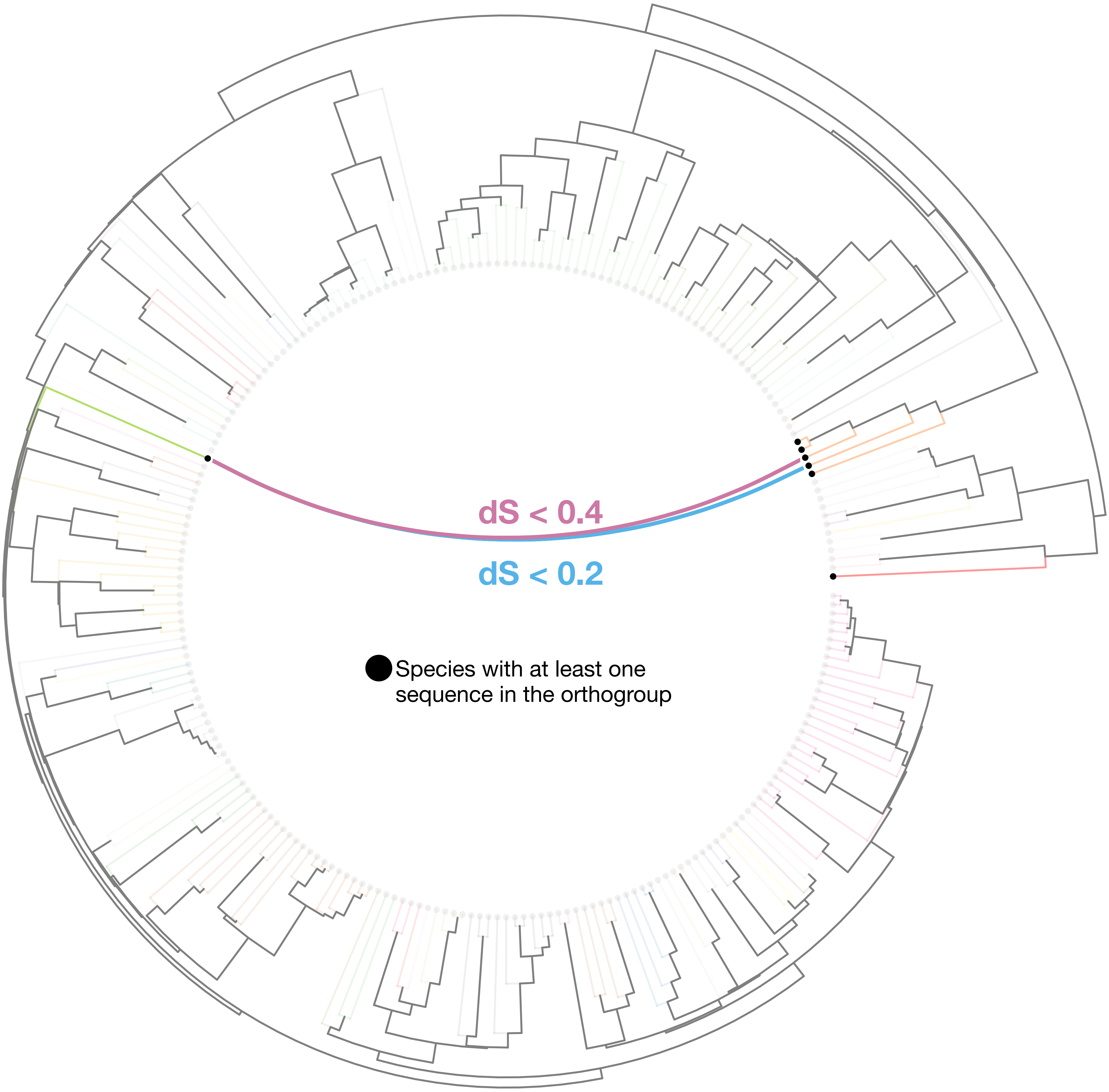

### N5.HOG0013500

Orthogroup phylogeny
and
protein alignment

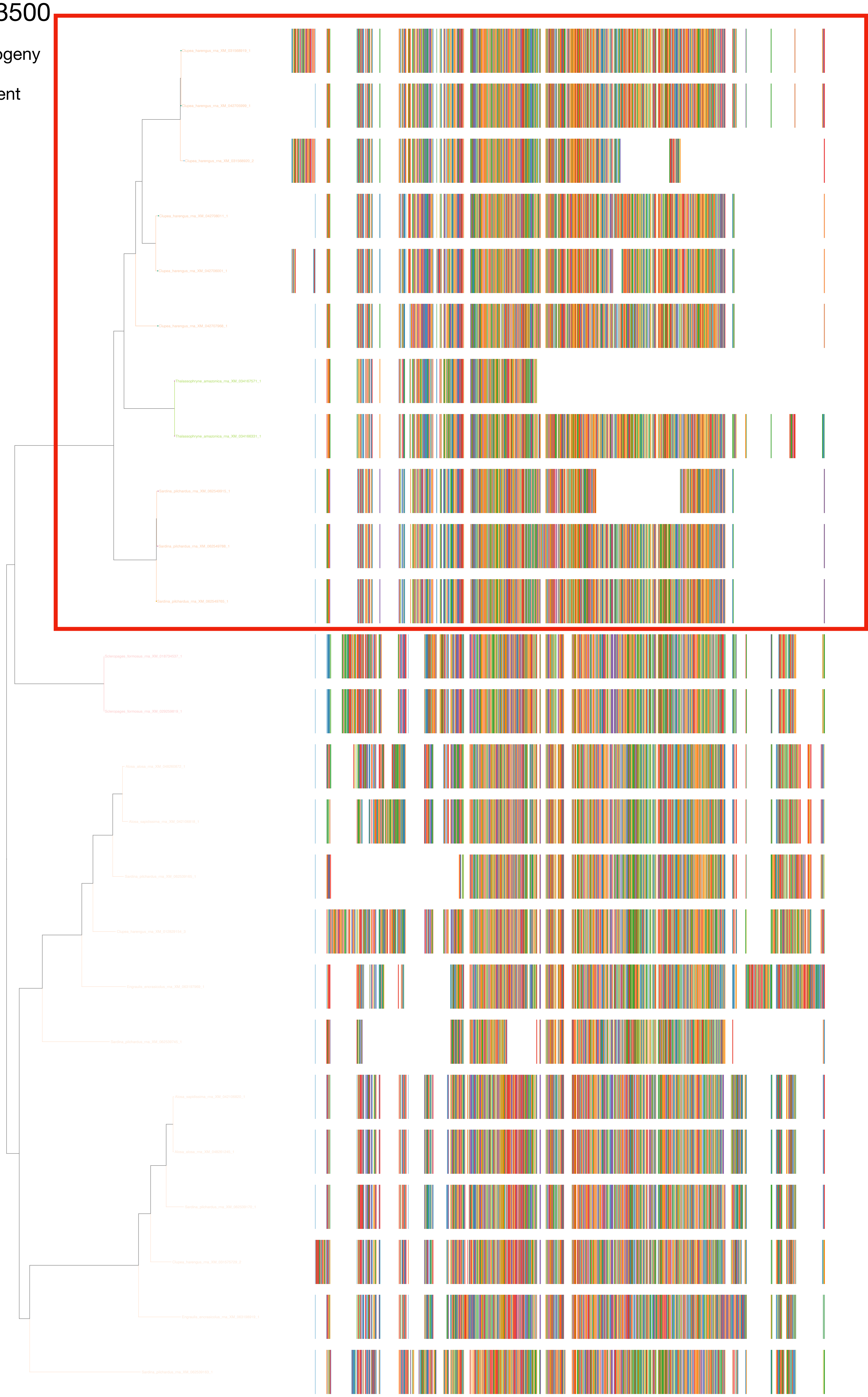

### N5.HOG0013500

Zoom on HGT clade with bootstrap values

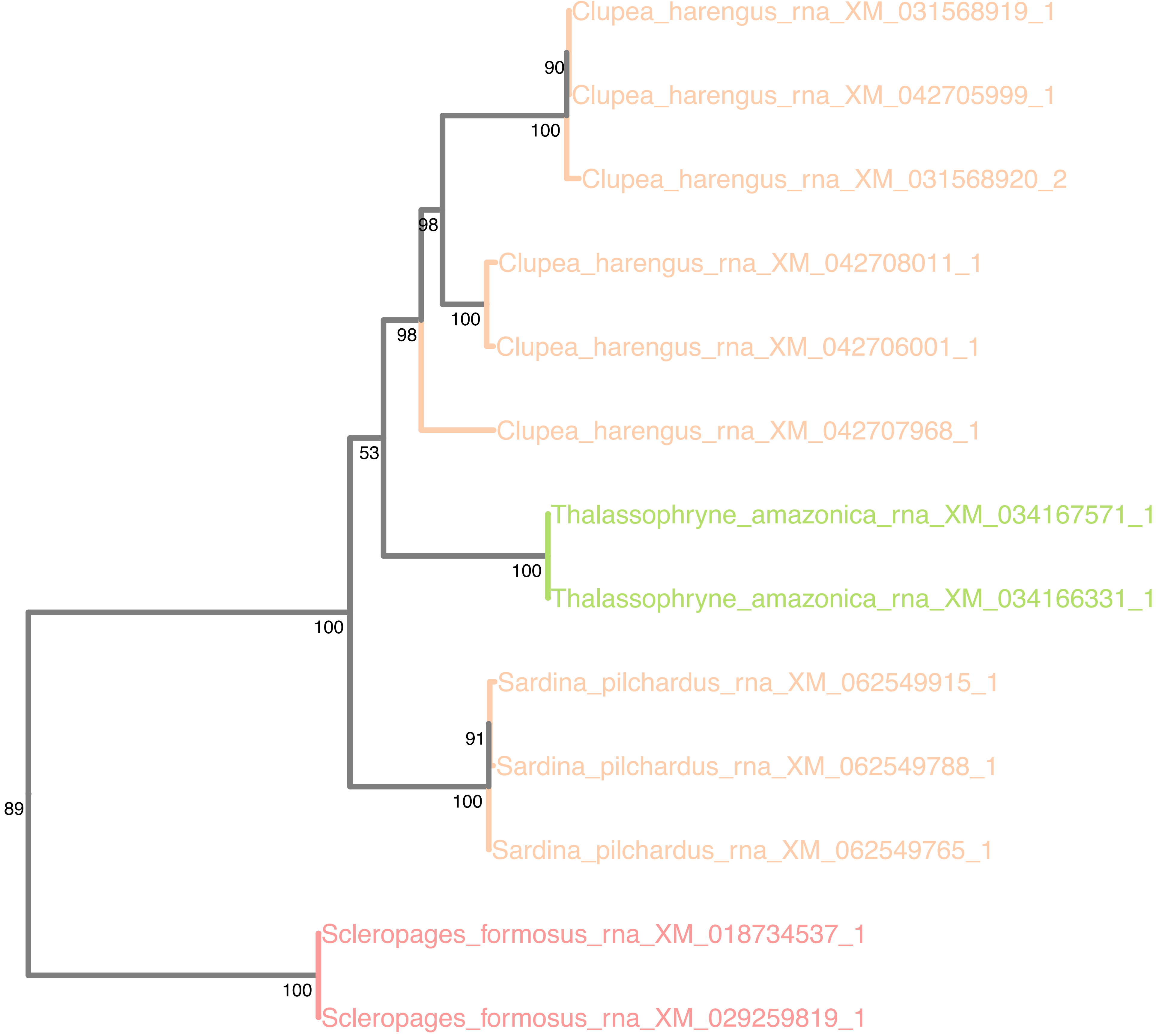

### N5.HOG0013500

Orthogroup phylogeny
and
trimmed
protein alignment

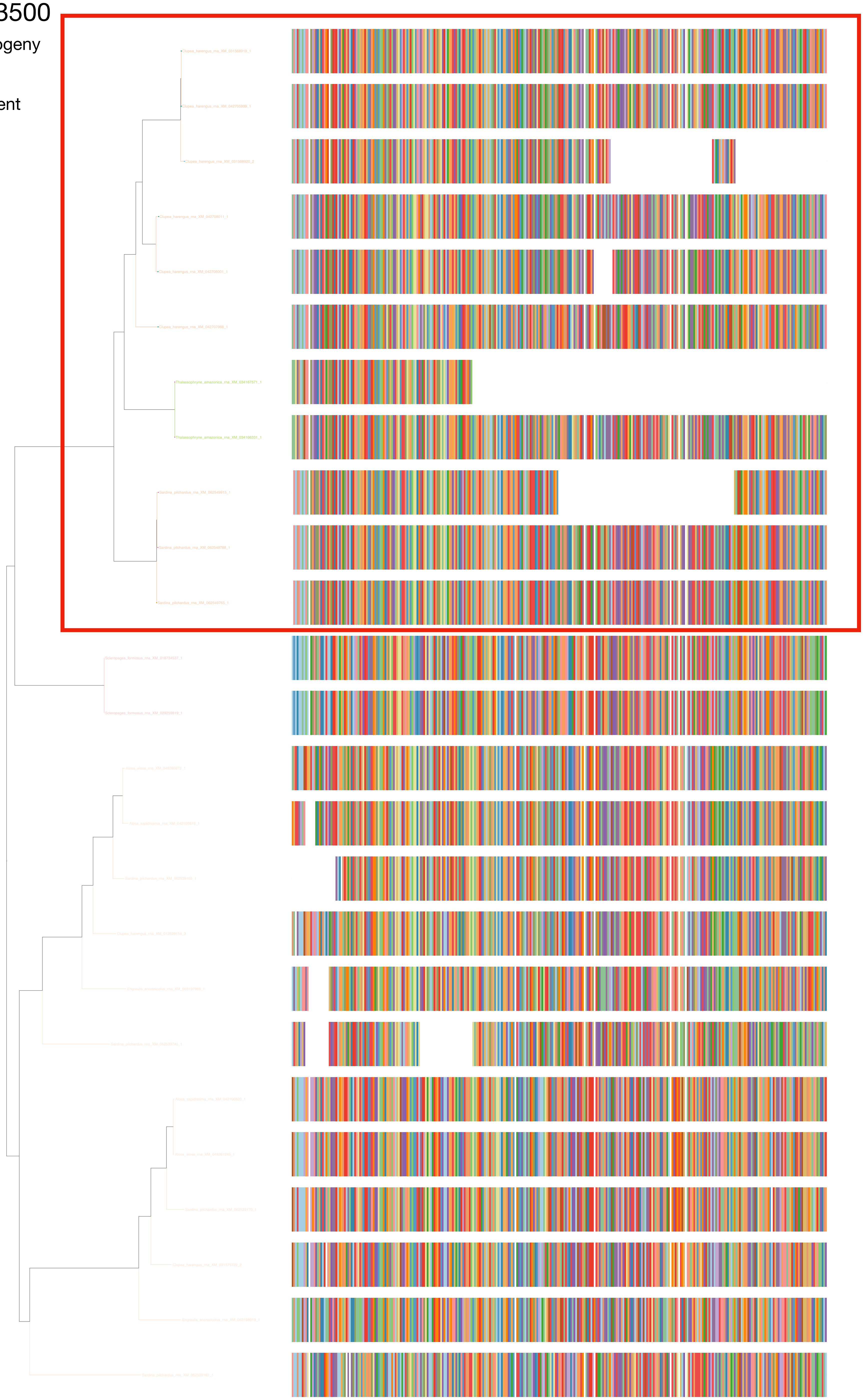

### N5.HOG0013500

« Best-match » phylogeny
and
protein alignment

50 best match against ray-finned fishes protein databases
10 best match against non-fish proteins from UniProt

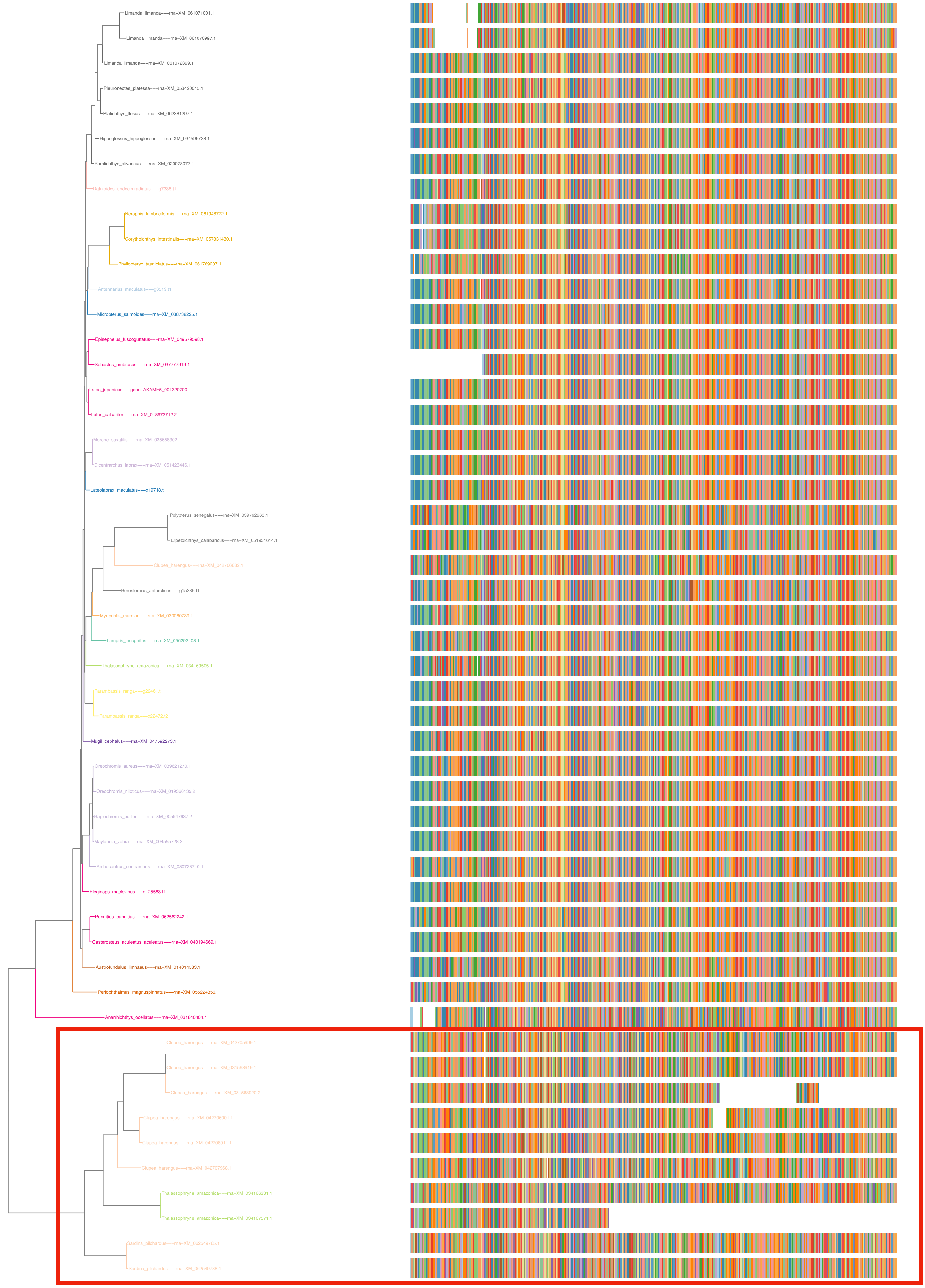

One of the two *Thalassophryne amazonica* copy is truncated in the annotation (XM_034167571)

By using Exonerate, we could find the full length sequence. This copy contains multiple frameshifts (# in the alignment below, with the full length and complete copy (XM_034166331) as query ). Thus, it is a pseudogene

```
       Query: Thalassophryne_amazonica---rna-XM_034166331.1
      Target: NC_047105.1:54418817-54442016 [revcomp]
       Model: protein2genome:local
   Raw score: 2019
 Query range: 0 -> 428
Target range: 12279 -> 9256

    1 : MetArgPheTyrLeuSerValValPheThrIleLeuValMetAlaAlaAspHisThrArg :    20
        ||||||||||||||||||||||||||||||||||||||||||||||||||||||||||||
        MetArgPheTyrLeuSerValValPheThrIleLeuValMetAlaAlaAspHisThrArg
12279 : ATGAGGTTCTACCTGTCTGTTGTTTTCACCATTCTGGTCATGGCTGCGGACCACACCCGT : 12222

   21 : AspCysGluProLeuThrGluThrValArgLeuProSerValLysValSerLysGluIle :    40
        ||||||||||||||||||||||||||||||||||||||||||||||||||||||||||||
        AspCysGluProLeuThrGluThrValArgLeuProSerValLysValSerLysGluIle
12221 : GATTGTGAACCACTTACCGAAACAGTGAGACTACCCTCAGTCAAGGTTTCCAAGGAAATC : 12162

   41 : LysGluThrGlnIle{S}  >>>> Target Intron 1 >>>>  {er}ThrValAla :    49
        |||||||||||||||{|}           1742 bp           {||}|||||||||
        LysGluThrGlnIle{S}++                         ++{er}ThrValAla
12161 : AAAGAAACTCAGATC{T}gt.........................ag{CG}ACCGTTGCT : 10393

   50 : GluGluProAlaProSerAlaGlnIleAlaPheHisLysAlaLeuIleAspGlySerThr :    69
        ||||||||||||||||||||||||||||||||||||||||||||||||||||||||||||
        GluGluProAlaProSerAlaGlnIleAlaPheHisLysAlaLeuIleAspGlySerThr
10392 : GAAGAGCCTGCTCCATCAgctcagattgcatttcacaaggCATTGATTGATGGCAGCACC : 10333

   70 : SerArgSerAspSerHisValProProTyrThrPheAspAspAsnProAsnLeuGluTrp :    89
        ||||||||||||||||||||||||||||||||||||||||||||||||||||||||||||
        SerArgSerAspSerHisValProProTyrThrPheAspAspAsnProAsnLeuGluTrp
10332 : TCGAGAAGTGATTCTCACGTCCCTCCATACACCTTTGATGACAACCCCAACCTTGAGTGG : 10273

   90 : GlnLeuPheAsnGlySerLeuProAsnGlyAlaValGlyIleTrpAsnSerTyrThrAsn :   109
        ||||||||||||||||||||||||||||||||||||||||||||||||||||||||||||
        GlnLeuPheAsnGlySerLeuProAsnGlyAlaValGlyIleTrpAsnSerTyrThrAsn
10272 : CAGTTGTTTAATGGGTCTCTTCCCAACGGAGCAGTGGGCATCTGGAACAGTTACACAAAT : 10213

  110 : ArgTyrAspTyrValCysArgAlaTyrAspGlyCysGluSerGlyPheTyrSerSerAsp :   129
        ||||||||||||||||||||||||||||||||||||||||||||||||||||||||||||
        ArgTyrAspTyrValCysArgAlaTyrAspGlyCysGluSerGlyPheTyrSerSerAsp
10212 : CGCTATGACTATGTGTGCAGAGCATATGATGGTTGTGAGAGCGGATTCTACAGCAGTGAC : 10153

  130 : PheGlyLysCysLeuPheProSerTyrProLeuLeuLeuIleThrGlnSerPheTyrIle :   149
        ||||||||||||||||||||||||||||||||||||||||||||||||||||||||||||
        PheGlyLysCysLeuPheProSerTyrProLeuLeuLeuIleThrGlnSerPheTyrIle
10152 : TTTGGTAAATGCTTGTTCCCAAGTTACCCTTTGTTGTTGATTACTCAGAGTTTTTACATC : 10093

  150 : LeuValAsnLysAspAspPheValPheLeuGluTrpLysTrpGlySerGlyGlySerVal :   169
        ||||||||||||||||||||||||||||||||||||||||||||||||||||||||||||
        LeuValAsnLysAspAspPheValPheLeuGluTrpLysTrpGlySerGlyGlySerVal
10092 : CTCGTAAACAAAGACGACTTTGTATTTCTGGAGTGGAAATGGGGATCAGGTGGGTCTGTT : 10033

  170 : ProGlnAsnSerIleArgThrCys-PheLysProGlnPheTyrValGlyLysAsnLysTy :   189
        ||||||||||||||||||||||||#|||||||||||||||||||||||||||||||||||
        ProGlnAsnSerIleArgThrCys#PheLysProGlnPheTyrValGlyLysAsnLysTy
10032 : CCCCAAAACTCAATCAGGACATGTTTTTAAGCCACAGTTTTATGTTGGAAAAAATAAGTA :  9972

  190 : rGlyLeuGlyLeuLeuSerSerGlyAspSerPheTyrLeuProTrpValGluGluTyrSe :   209
        ||||||||||||||||||||||||||||||||||||||||||||||||||||||||||||
        rGlyLeuGlyLeuLeuSerSerGlyAspSerPheTyrLeuProTrpValGluGluTyrSe
 9971 : TGGTTTAGGGTTGCTTTCCTCTGGAGATTCATTCTATTTGCCATGGGTTGAAGAATATTC :  9912

  210 : rAsnGlyThrIleTyrGlyTyrSerThrTrpTyrArgSerSerTyrGlnAlaLeuThr-- :   229
        ||||||||||||||||||||||||||||||||||||||||||||||||||||||||||##
        rAsnGlyThrIleTyrGlyTyrSerThrTrpTyrArgSerSerTyrGlnAlaLeuThr##
 9911 : CAATGGTACAATCTATGGGTATAGTACATGGTATAGAAGCTCTTACCAGGCTTTGACAGT :  9852

  230 : ValAsnThrAspLysTyrLysGlnGluIleLysAspValLysTyrPheThrAspGlnAla :   248
           |||||||||||||||||||||||||||||||||||||||||||||||||||||||||
        ---AsnThrAspLysTyrLysGlnGluIleLysAspValLysTyrPheThrAspGlnAla
 9851 : ---AATACAGACAAATATAAGCAGGAAATCAAAGATGTAAAGTATTTCACGGATCAAGCT :  9796

  249 : LysIleIleGlyThrProProPheGlyIleValLysAsnThrLeuSerAsnArgGluCys :   268
        ||||||||||||||||||||||||||||||||||||||||||||||||||||||||||||
        LysIleIleGlyThrProProPheGlyIleValLysAsnThrLeuSerAsnArgGluCys
 9795 : AAGATCATAGGCACTCCCCCATTCGGTATTGTCAAAAACACTTTAAGCAACAGAGAATGC :  9736

  269 : HisProValThrLeuThrThrThrLeuSerThrSerGlnThrLysThrSerAsnTrpGln :   288
        ||||||||||||||||||||||||||||||||||||||||||||||||||||||||||||
        HisProValThrLeuThrThrThrLeuSerThrSerGlnThrLysThrSerAsnTrpGln
 9735 : CATCCTGTAACTTTGACGACCACATTGTCTACATctcaaacaaaaacaagcaattGGCAA :  9676

  289 : PheSerTyrSerMetThrLeuSerIleSerThrThrValSerAlaSerIleProAspIle :   308
        ||||||||||||||||||||||||||||||||||||||||||||||||||||||||||||
        PheSerTyrSerMetThrLeuSerIleSerThrThrValSerAlaSerIleProAspIle
 9675 : TTCTCTTACTCCATGACCCTTTCCATCAGCACCACTGTATCTGCTAGCATCCCTGACATC :  9616

  309 : ValAspPheSerValThrIleGlyValGluGln--ThrPheThrValThrLysGluIleS :   328
        |||||||||||||||||||||||||||||||||##   ||||||||||||||||||||||
        ValAspPheSerValThrIleGlyValGluGln##---PheThrValThrLysGluIleS
 9615 : GTTGACTTTAGTGTCACCATTGGTGTAGAGCAGAC---TTTACAGTAACAAAGGAAATAT :  9557

  329 : erLeuSerGlnThrGlnThrThrSerLeuValValGluValThrValProProAsnMetA :   348
        ||||||||||||||||||||||||||||||||||||||||||||||||||||||||||||
        erLeuSerGlnThrGlnThrThrSerLeuValValGluValThrValProProAsnMetA
 9556 : CCCTGTCTCAAACTCAAACTACTAGCCTTGTGGTTGAAGTCACTGTGCCCCCAAACATGA :  9497

  349 : rgCysThrIleGluMetGluGlyLysLysPheThr--SerAsnIleProPheGlnAlaAr :   367
        |||||||||||||||||||||||||||||||||||##   ||||||||||||||||||||
        rgCysThrIleGluMetGluGlyLysLysPheThr##---AsnIleProPheGlnAlaAr
 9496 : GGTGCACTATTGAGATGGAAGGCAAGAAATTCACATC---AATATACCTTTTCAGGCTCG :  9441

  368 : gLeuGlyArgThrTyrSer--AsnGlyAspThrLysTrpThrThrIleThrGlyIleTyr :   386
        |||||||||||||||||||##   ||||||||||||||||||||||||||||||||||||
        gLeuGlyArgThrTyrSer##---GlyAspThrLysTrpThrThrIleThrGlyIleTyr
 9440 : CCTTGGCCGTACCTACAGCAT---GGTGACACAAAGTGGACCACCATAACCGGCATATAT :  9385

  387 : AspGlyLeuGlnValSerAspPheGlnAlaThrAlaLysThrCysGluProValAspGlu :   406
        ||||||||||||||||||||||||||||||||||||||||||||||||||||||||||||
        AspGlyLeuGlnValSerAspPheGlnAlaThrAlaLysThrCysGluProValAspGlu
 9384 : GATGGGCTTCAGGTGTCTGATTTTCAGGCTACTGCAAAGACTTGTGAACCTGTGGATGAA :  9325

  407 : LeuValSerCysGlyAsnAspIleTyrValSerAspLeuSerSerLeuHisGlnGlnSer :   426
        ||||||||||||||||||||||||||||||||||||||||||||||||||||||||||||
        LeuValSerCysGlyAsnAspIleTyrValSerAspLeuSerSerLeuHisGlnGlnSer
 9324 : CTTGTGTCATGTGGGAATGATATCTATGTTTCTGATCTATCCTCCTTACATCAACAATCC :  9265

  427 : ProVal :   428
        ||||||
        ProVal
 9264 : CCTGTT :  9257
```

### N5.HOG0013500

dS distribution

Inter-species dS distribution

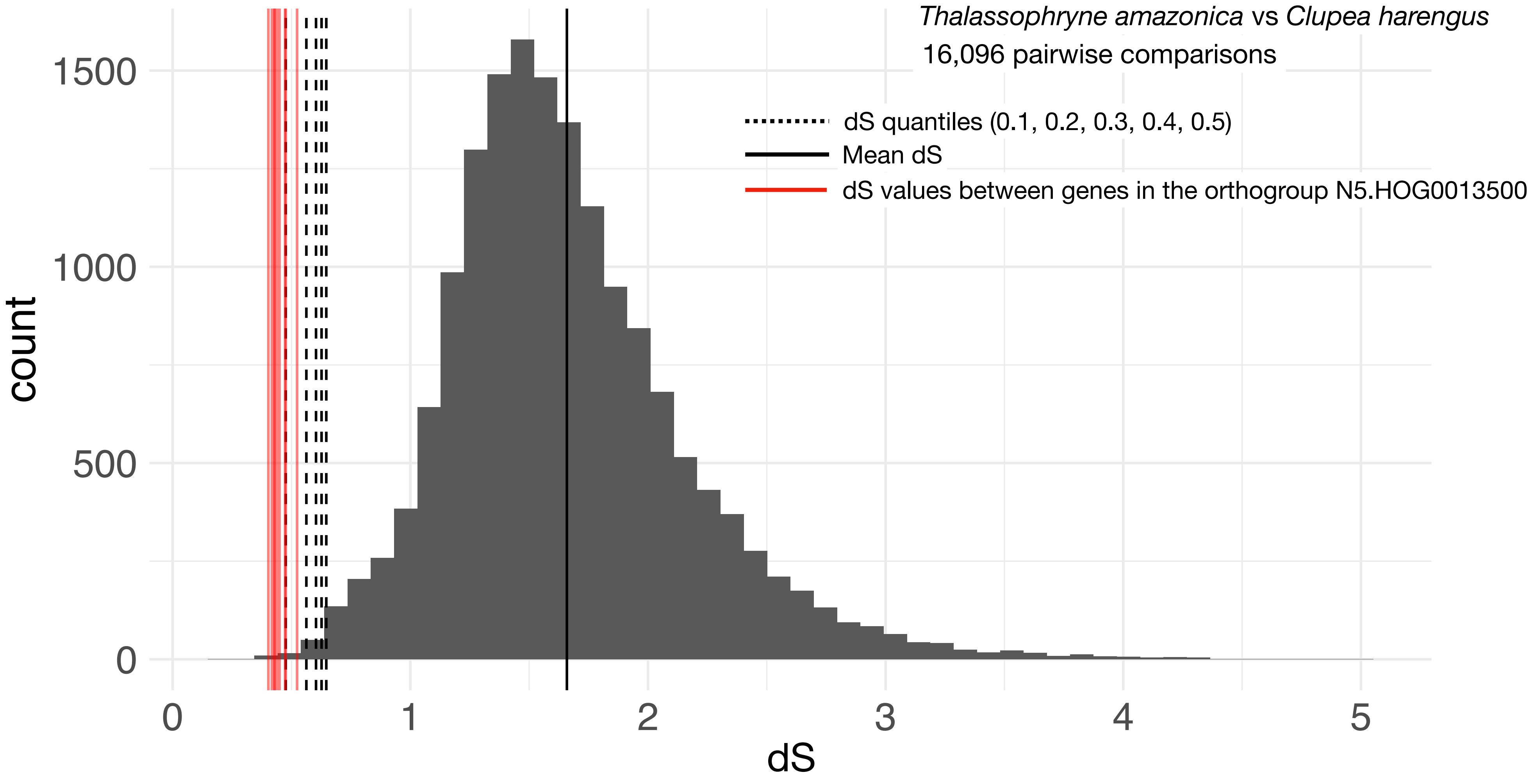


Intra-orthogroup dS

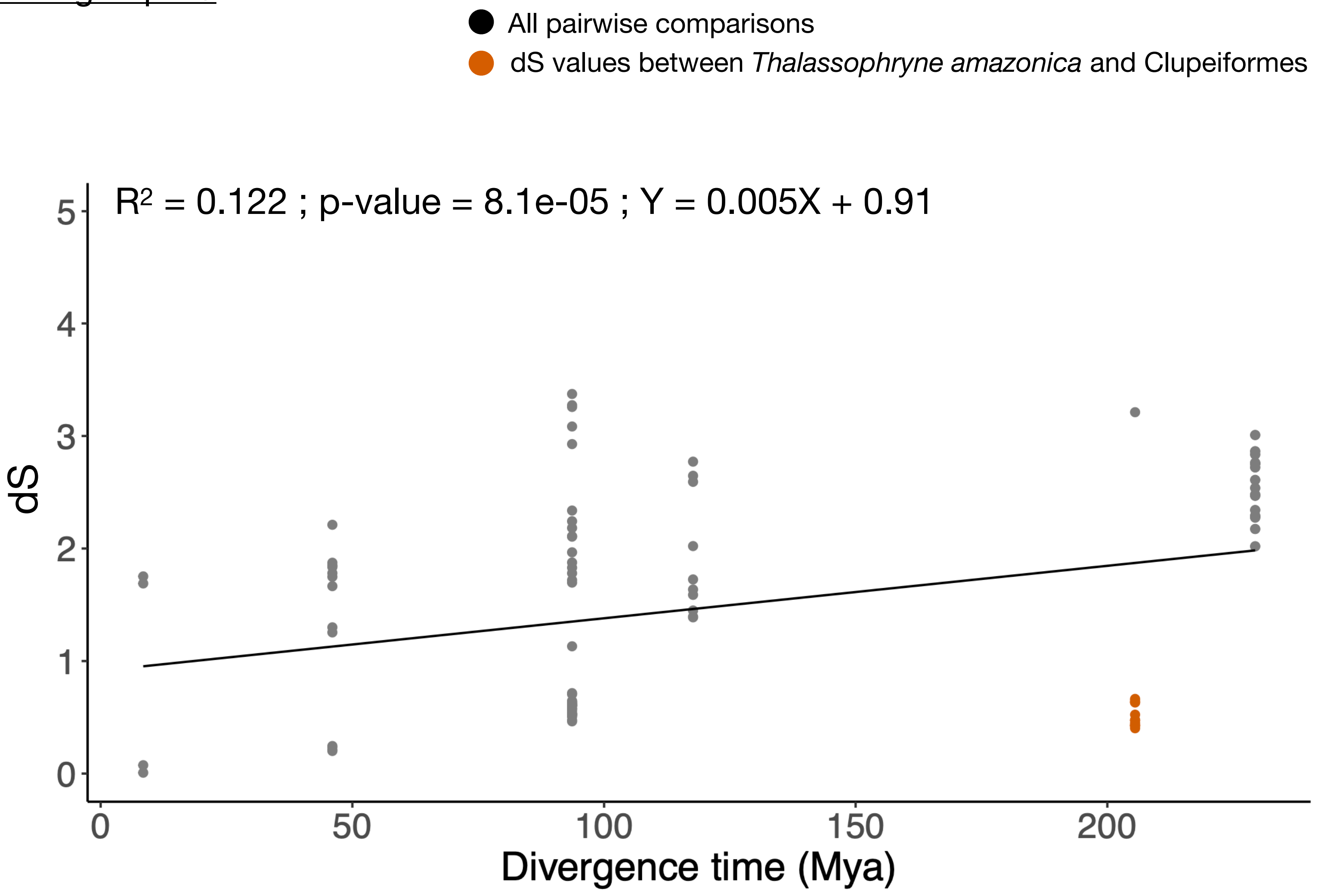

### N5.HOG0013500

#### Micro-synteny

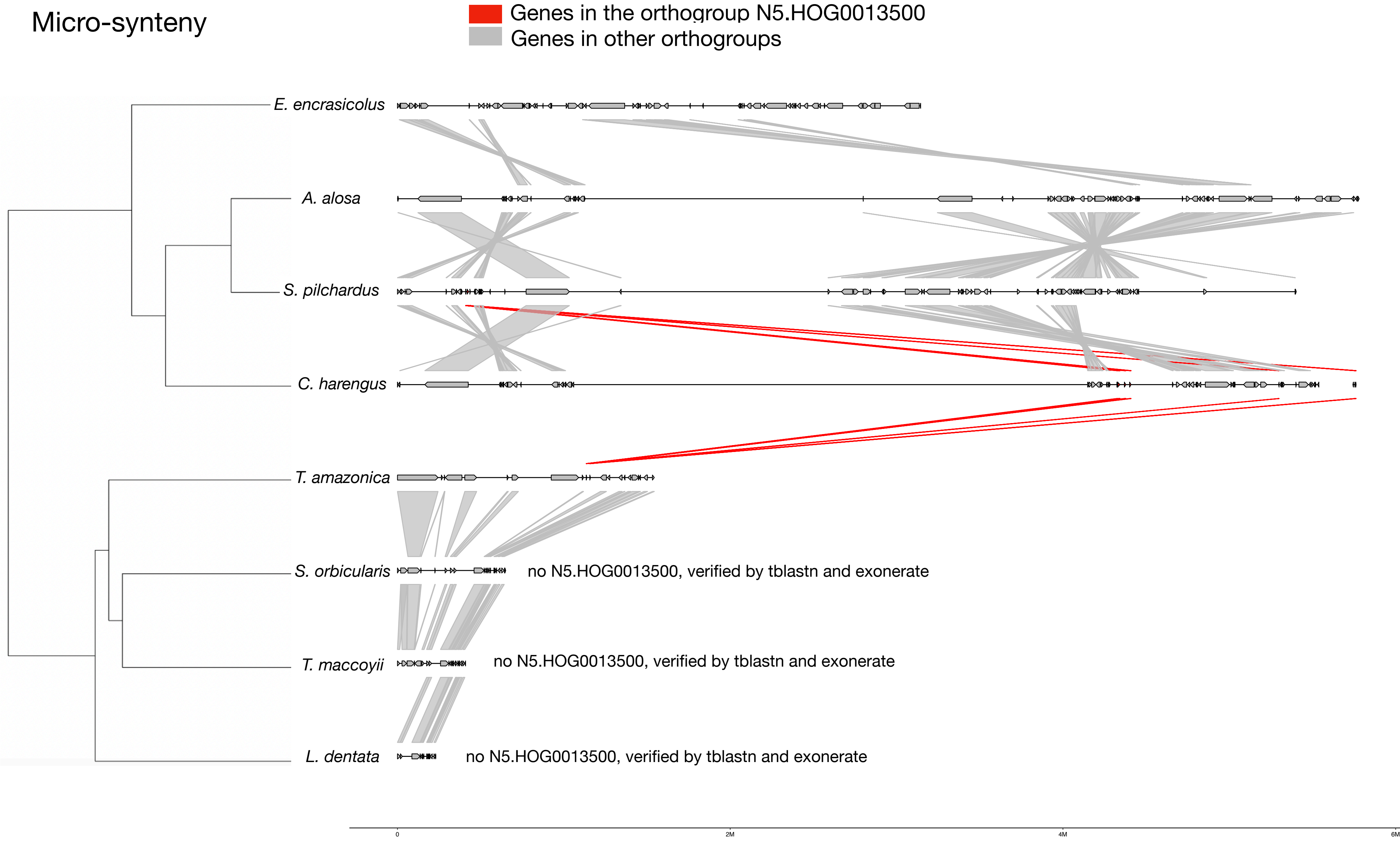


#### <u>Zoom on *T. amazonica* and *C. harengus*</u>

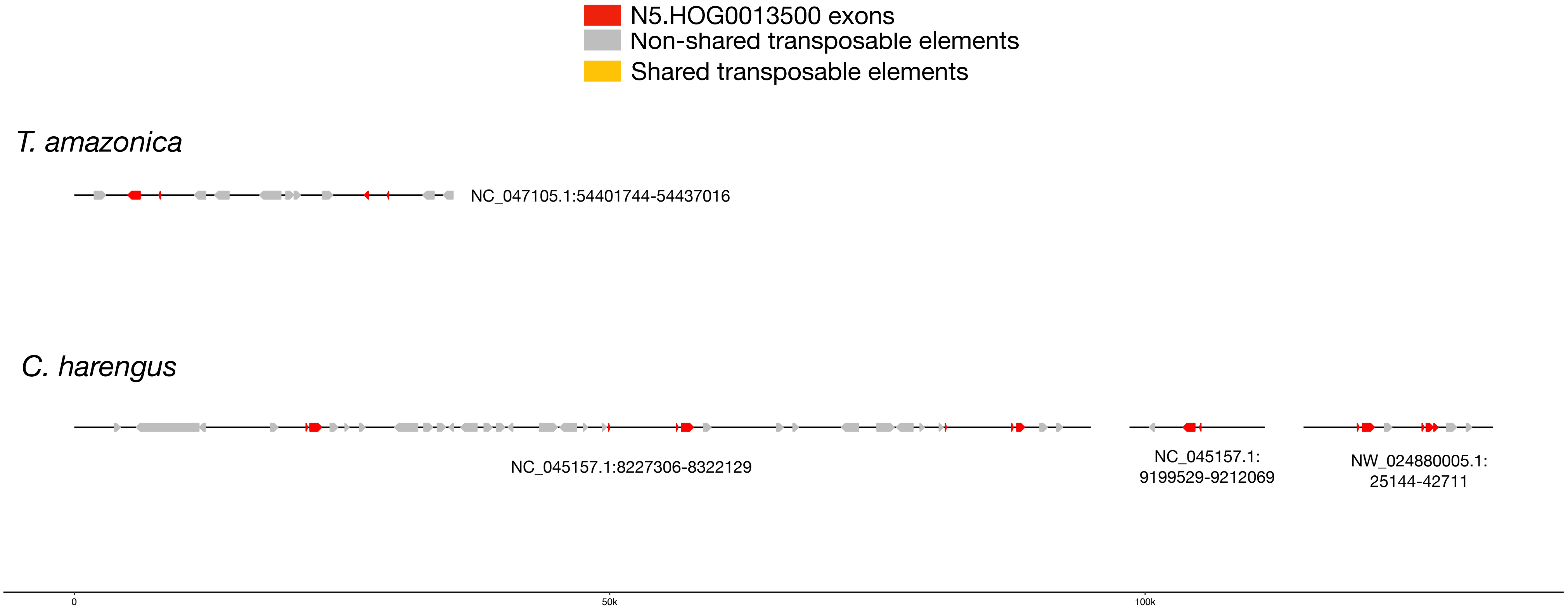

### N5.HOG0013500

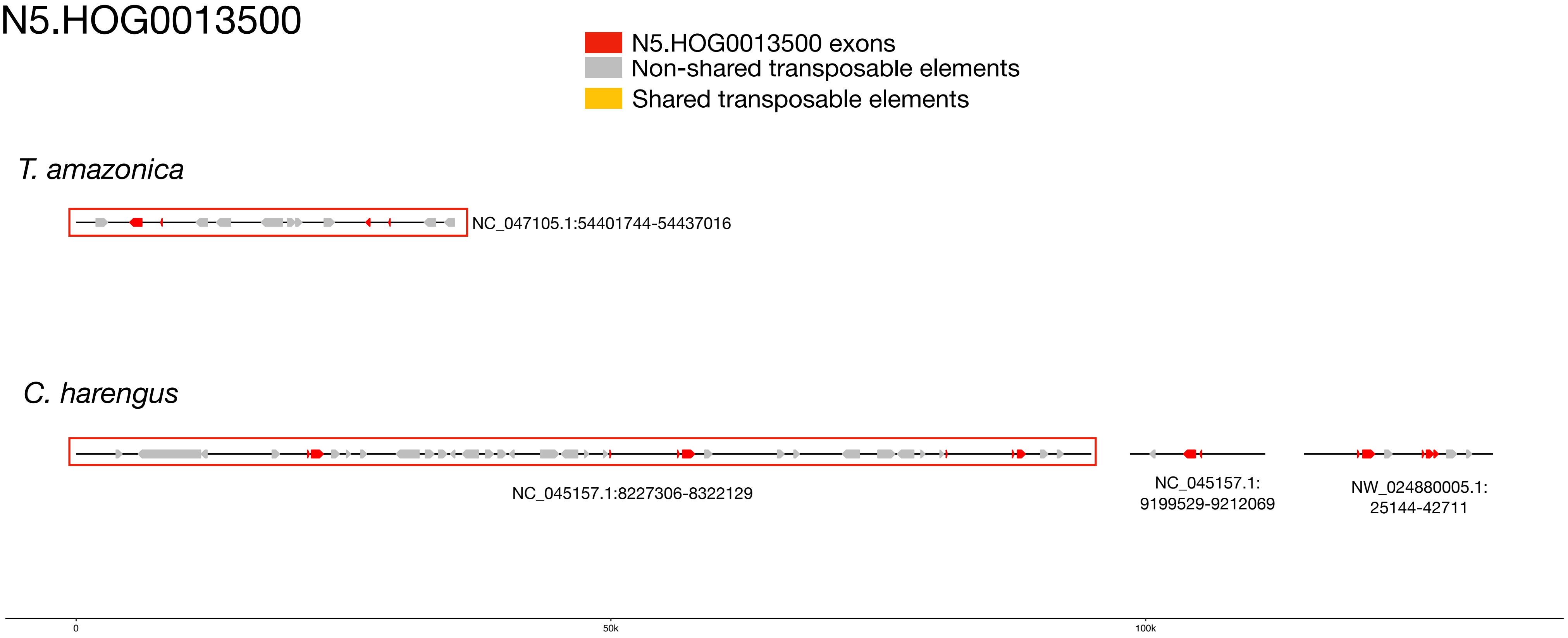


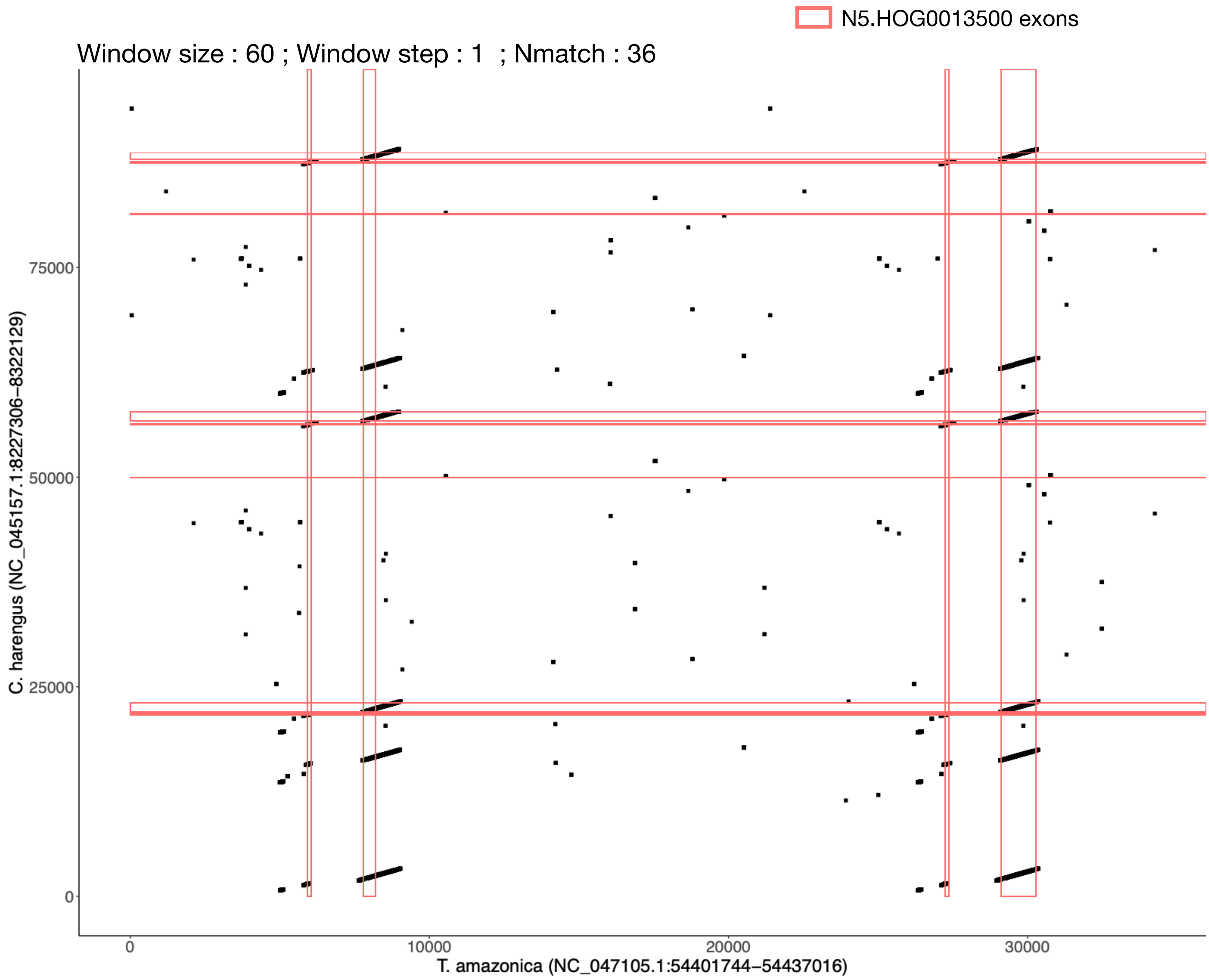

### N5.HOG0013500

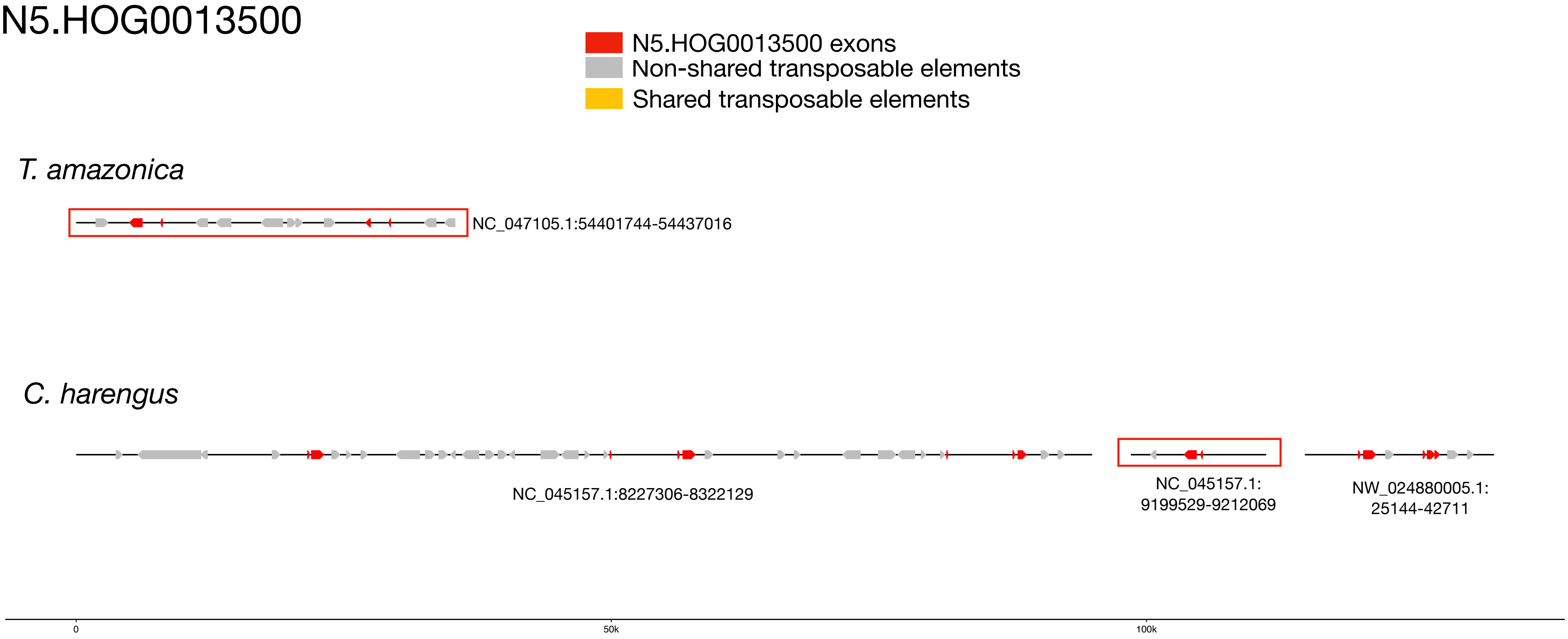


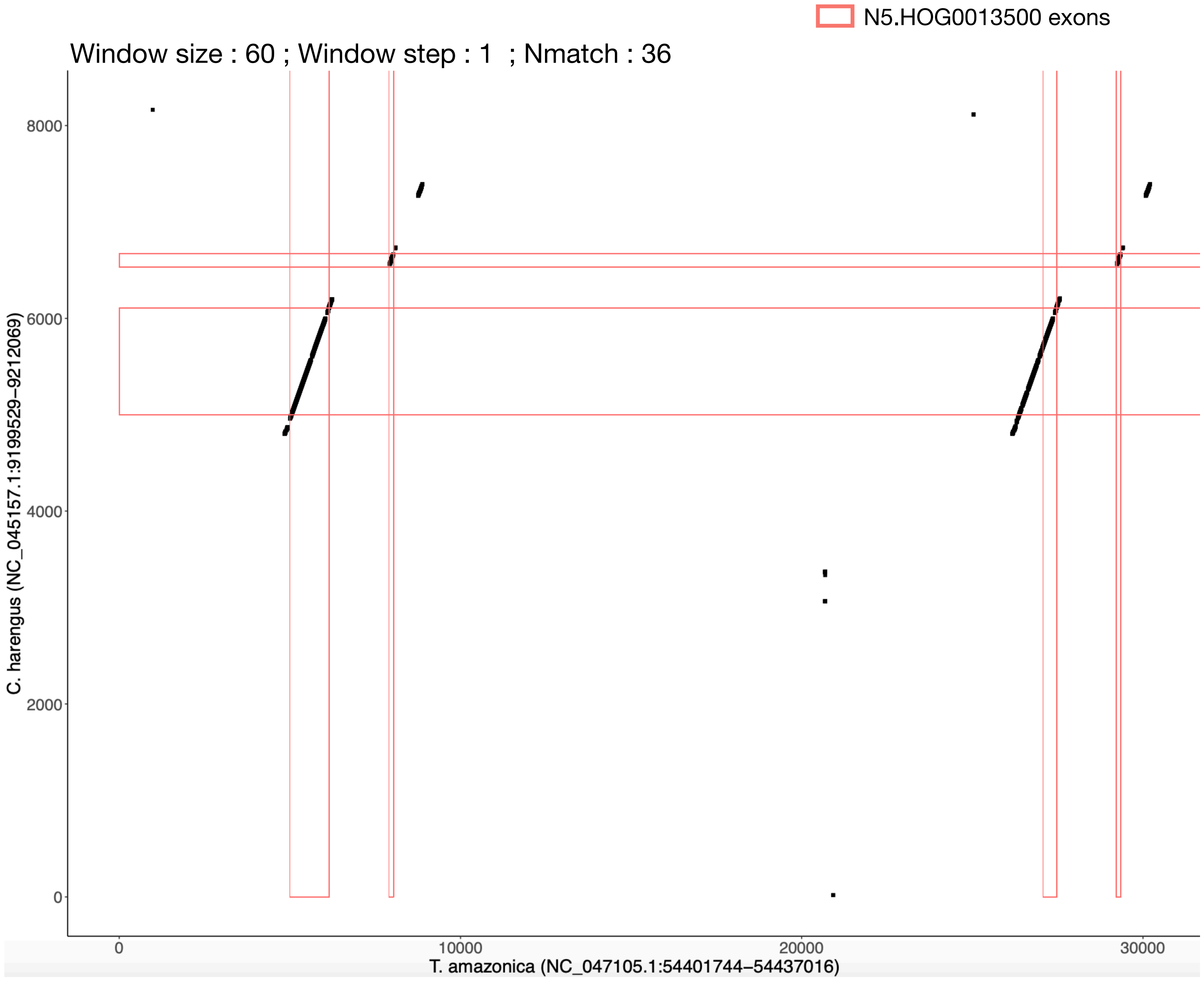

### N5.HOG0013500

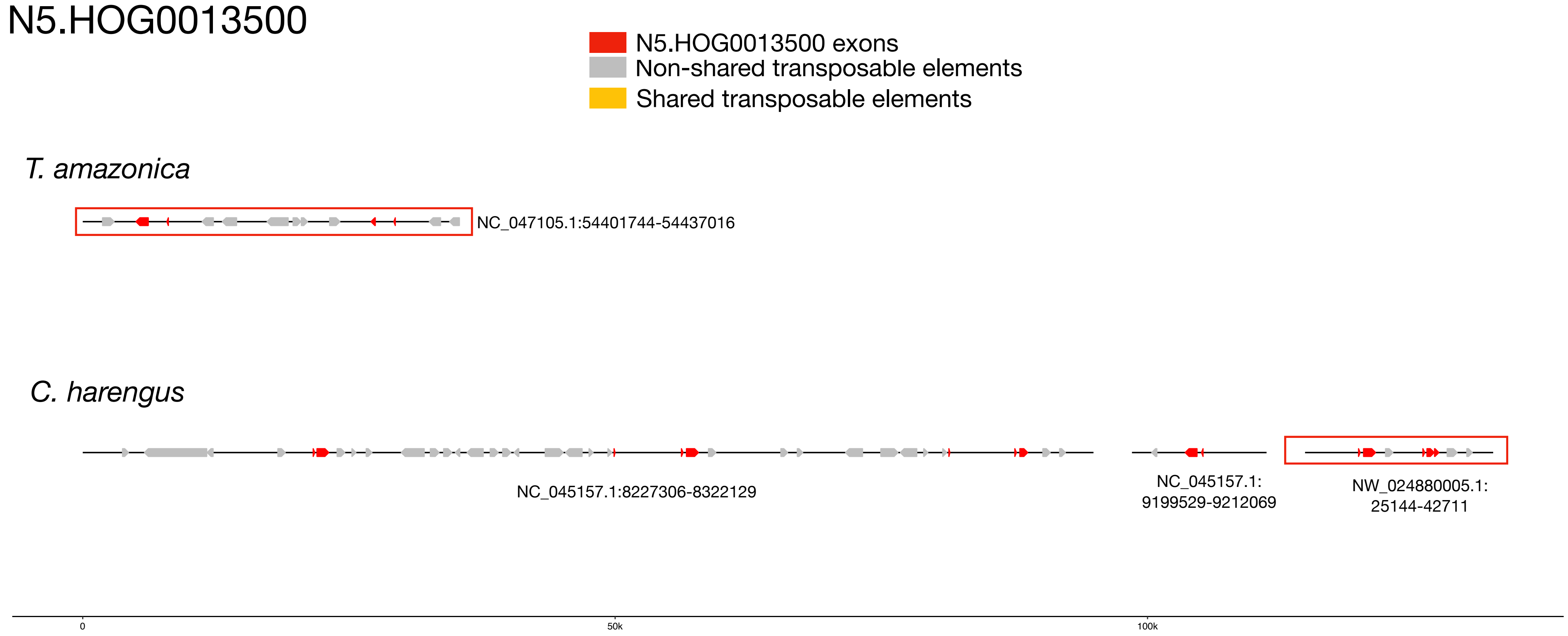


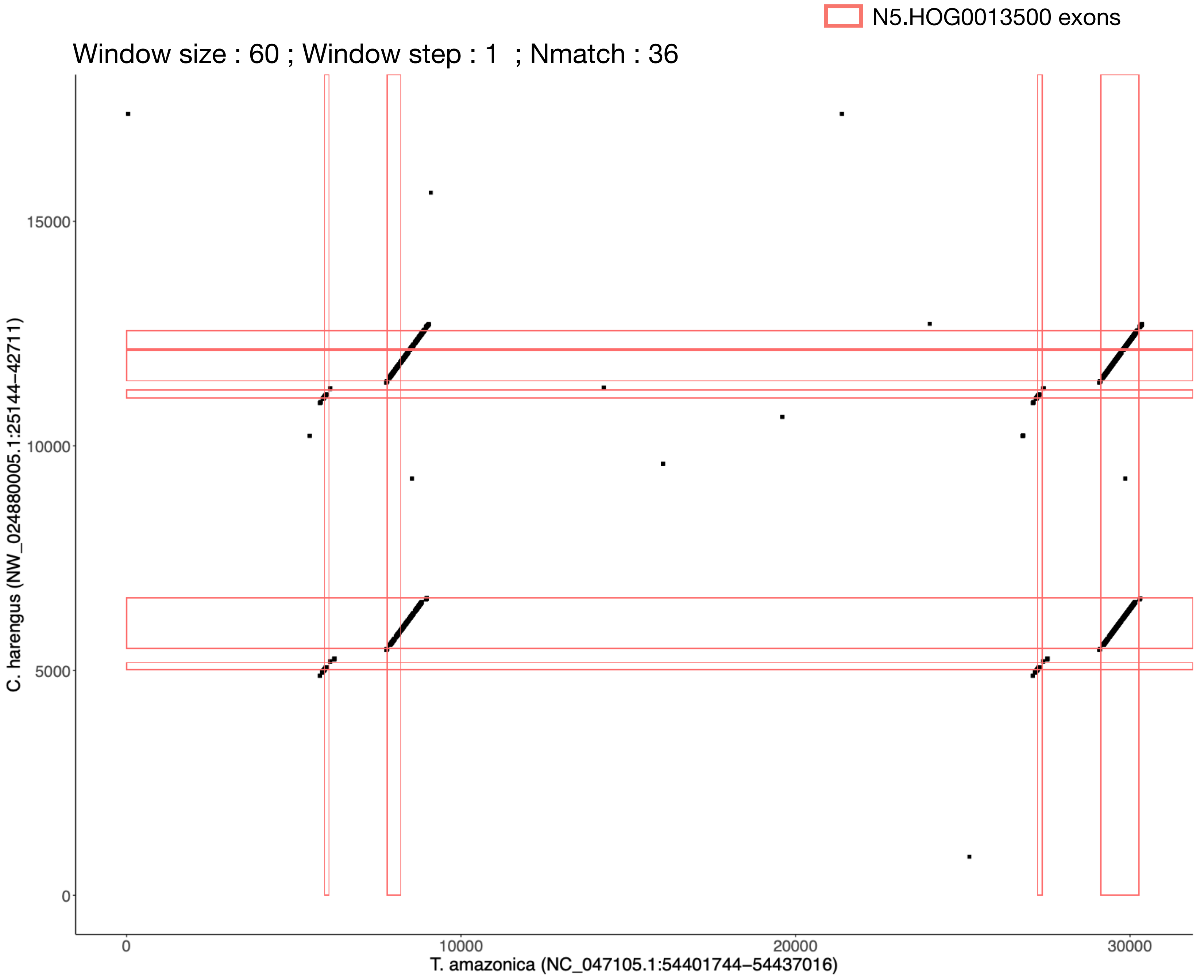

N5.HOG0013500

The gene was search with tblastn and exonerate on the two other non-annotated Batrachoididae genome assemblies (*Opsanus beta* ; *Chatrabus_melanurus*)

One complete version of the gene retrieved in *Opsanus beta (92.1%* DNA identity with the Thalassophryne amazonica complete copy) and one pseudogene retrieved in *Chatrabus melanurus (*77.4% DNA identity with the *Thalassophryne amazonica* complete copy)

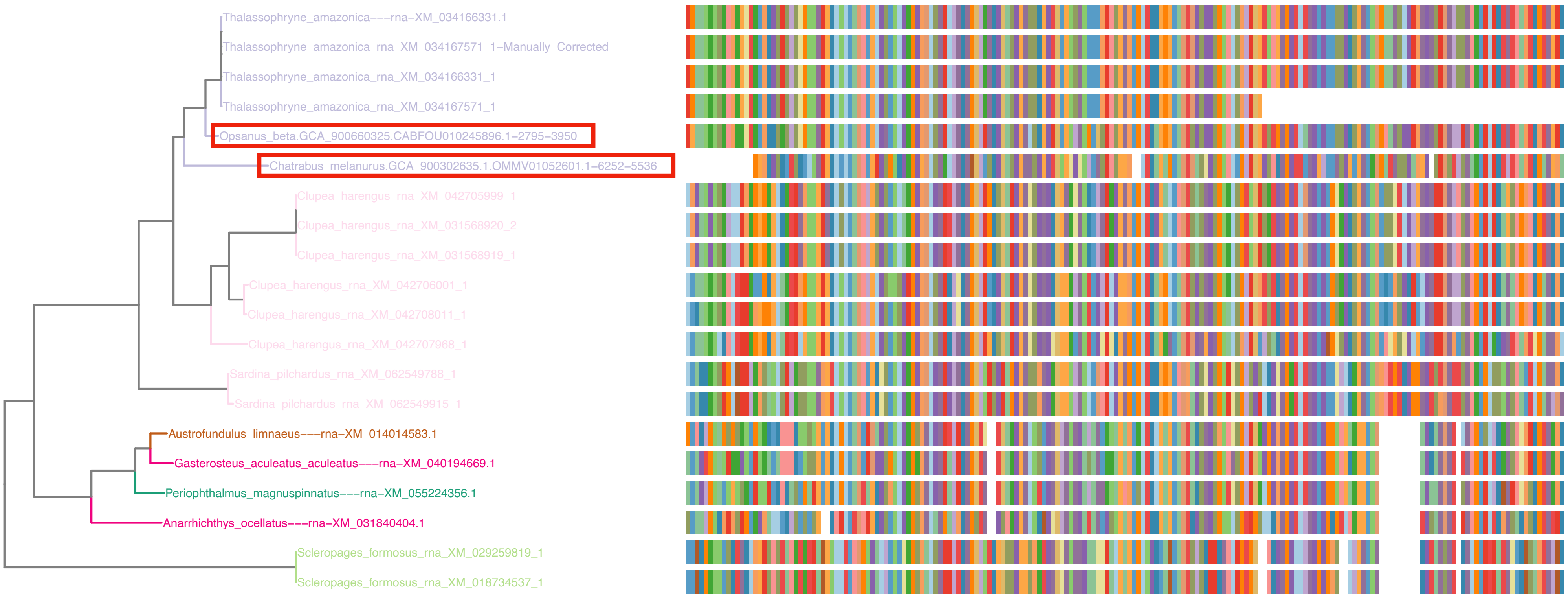


*Opsanus beta* gene alo found in the TSA database

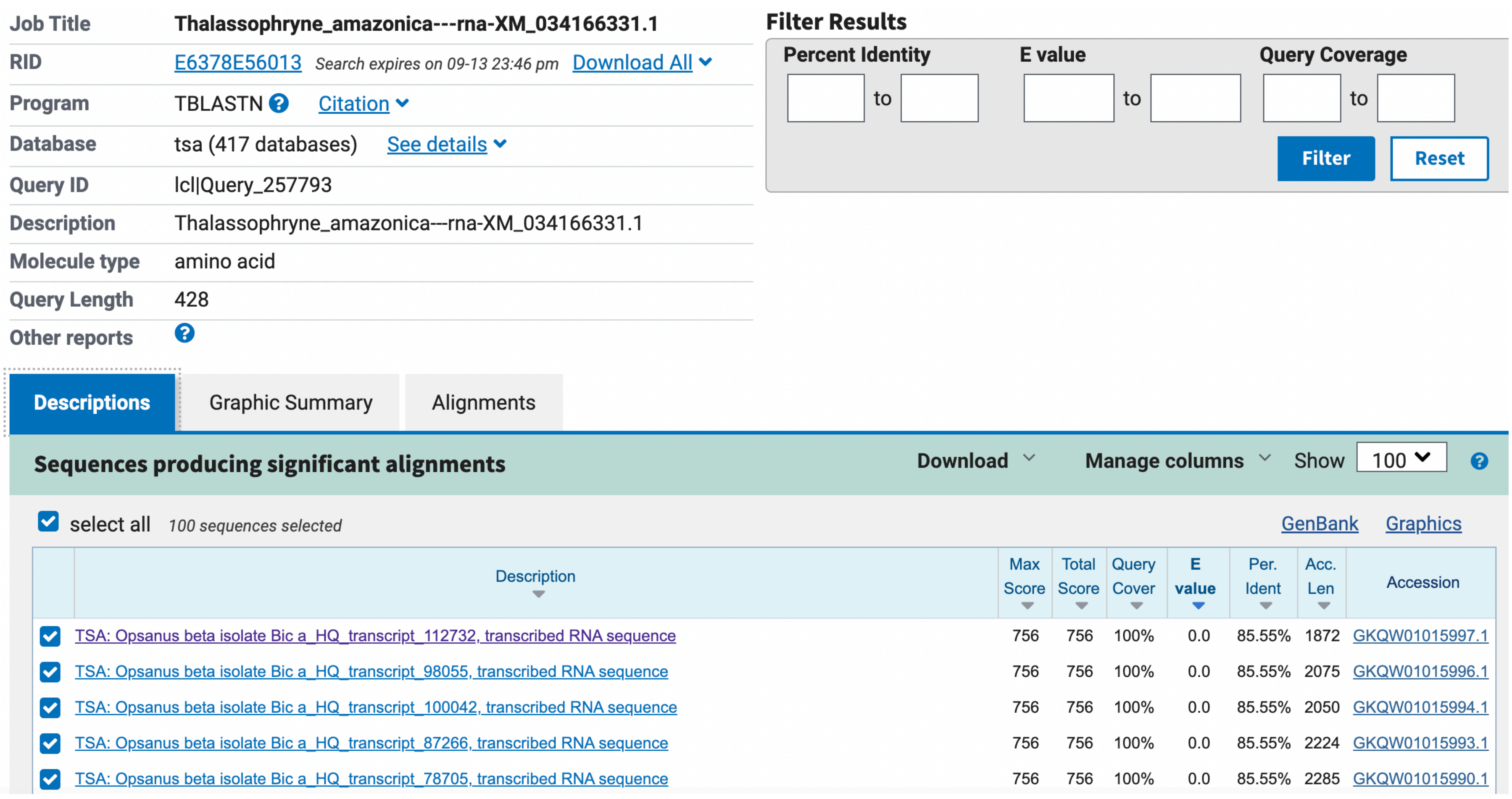

**Job Title** Thalassophryne_amazonica---rna-XM_034166331.1
**RID** E6378E56013 *Search expires on 09-13 23:46 pm* Download All
**Program** TBLASTN Citation
**Database** tsa (417 databases) See details
**Query ID** lcl|Query_257793
**Description** Thalassophryne_amazonica---rna-XM_034166331.1
**Molecule type** amino acid
**Query Length** 428
**Other reports**

**Filter Results**
Percent Identity ___ to ___ E value ___ to ___ Query Coverage ___ to ___
Filter Reset

Descriptions | Graphic Summary | Alignments

**Sequences producing significant alignments** Download Manage columns Show 100

select all *100 sequences selected* GenBank Graphics

| Description | Max Score | Total Score | Query Cover | E value | Per. Ident | Acc. Len | Accession |
|---|---|---|---|---|---|---|---|
| TSA: Opsanus beta isolate Bic a_HQ_transcript_112732, transcribed RNA sequence | 756 | 756 | 100% | 0.0 | 85.55% | 1872 | GKQW01015997.1 |
| TSA: Opsanus beta isolate Bic a_HQ_transcript_98055, transcribed RNA sequence | 756 | 756 | 100% | 0.0 | 85.55% | 2075 | GKQW01015996.1 |
| TSA: Opsanus beta isolate Bic a_HQ_transcript_100042, transcribed RNA sequence | 756 | 756 | 100% | 0.0 | 85.55% | 2050 | GKQW01015994.1 |
| TSA: Opsanus beta isolate Bic a_HQ_transcript_87266, transcribed RNA sequence | 756 | 756 | 100% | 0.0 | 85.55% | 2224 | GKQW01015993.1 |
| TSA: Opsanus beta isolate Bic a_HQ_transcript_78705, transcribed RNA sequence | 756 | 756 | 100% | 0.0 | 85.55% | 2285 | GKQW01015990.1 |

### N5.HOG0013500

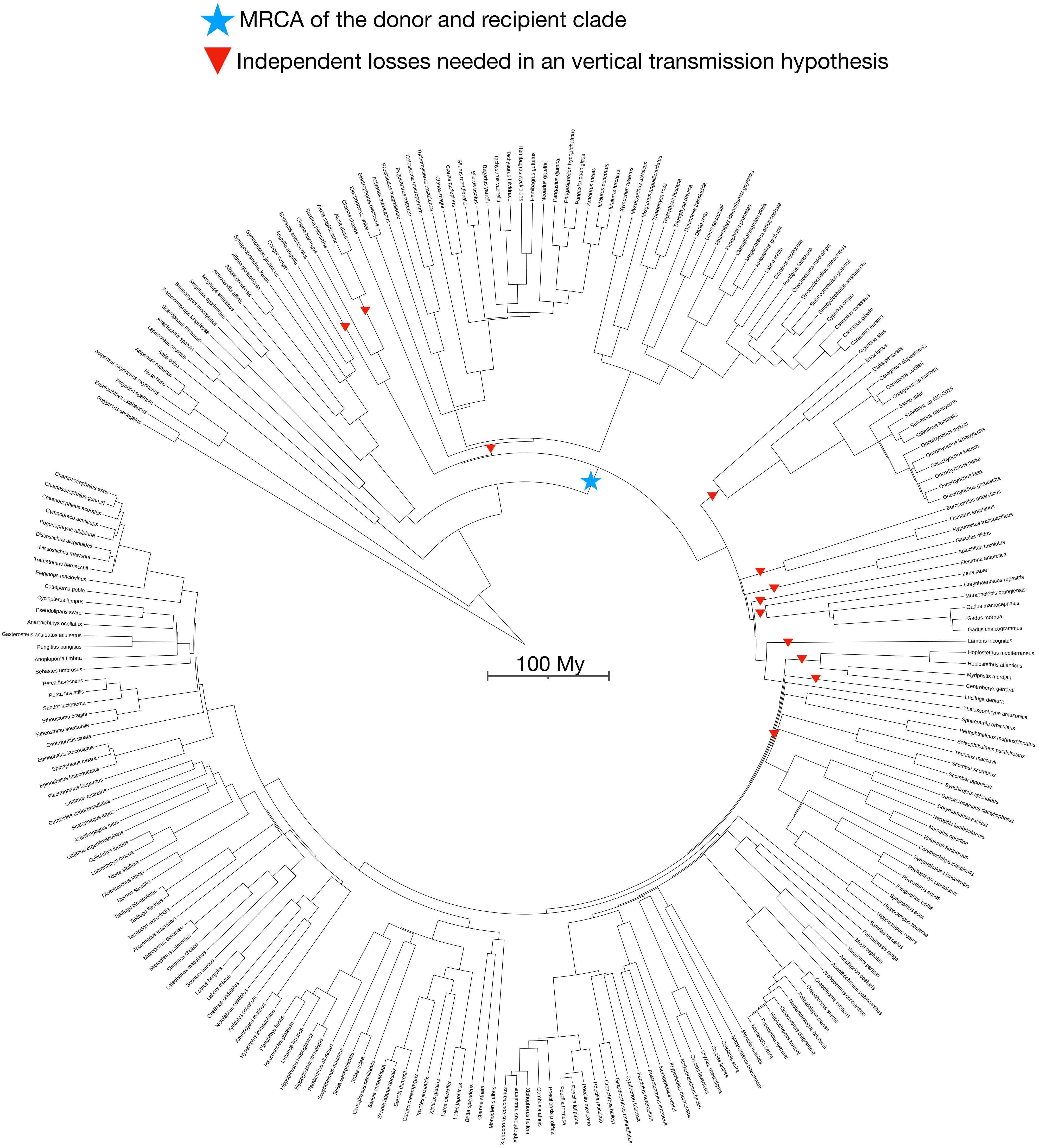

N5.HOG0019032

### N5.HOG0019032

Species tree

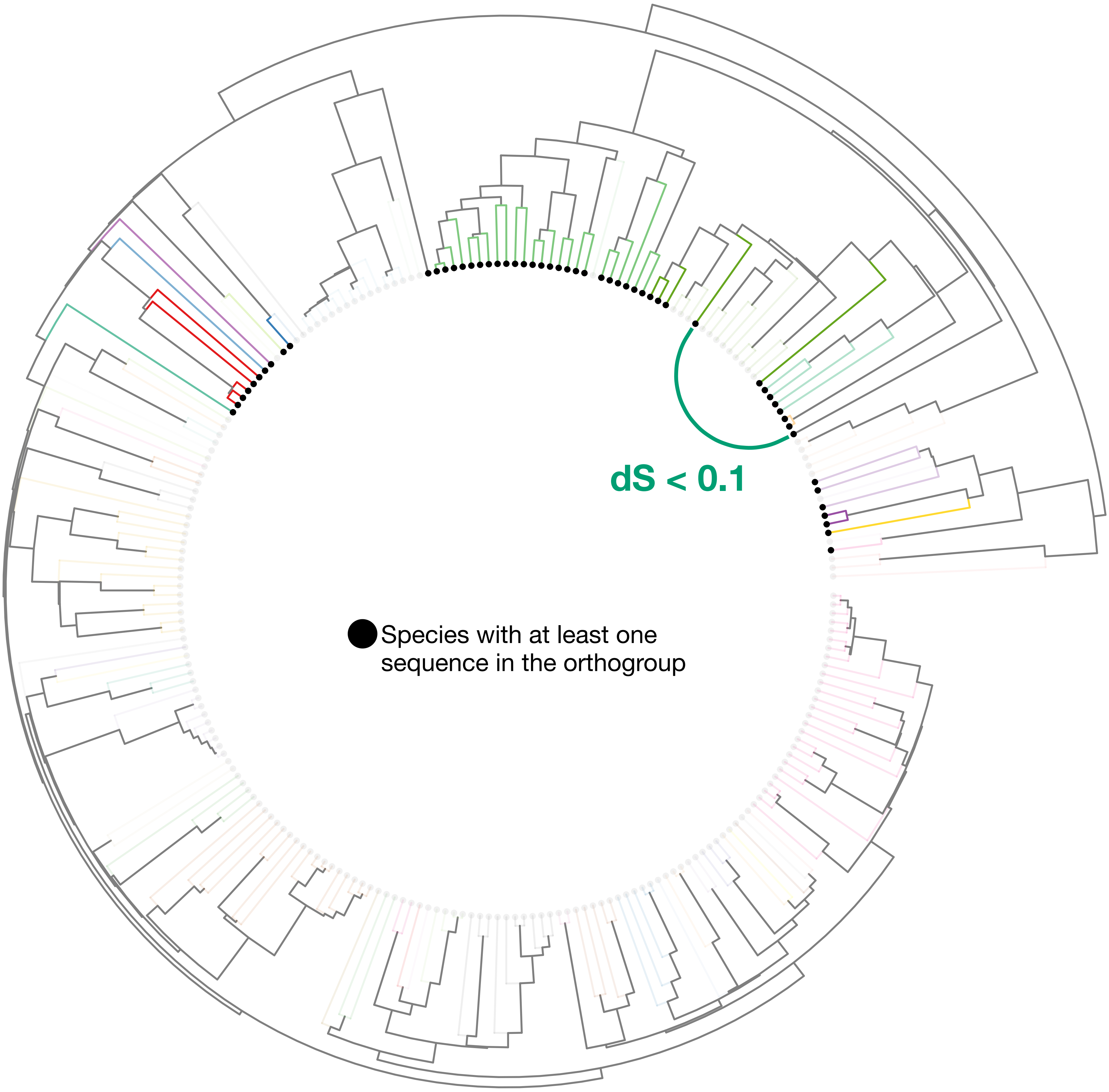

### N5.HOG0019032

Orthogroup phylogeny and protein alignment

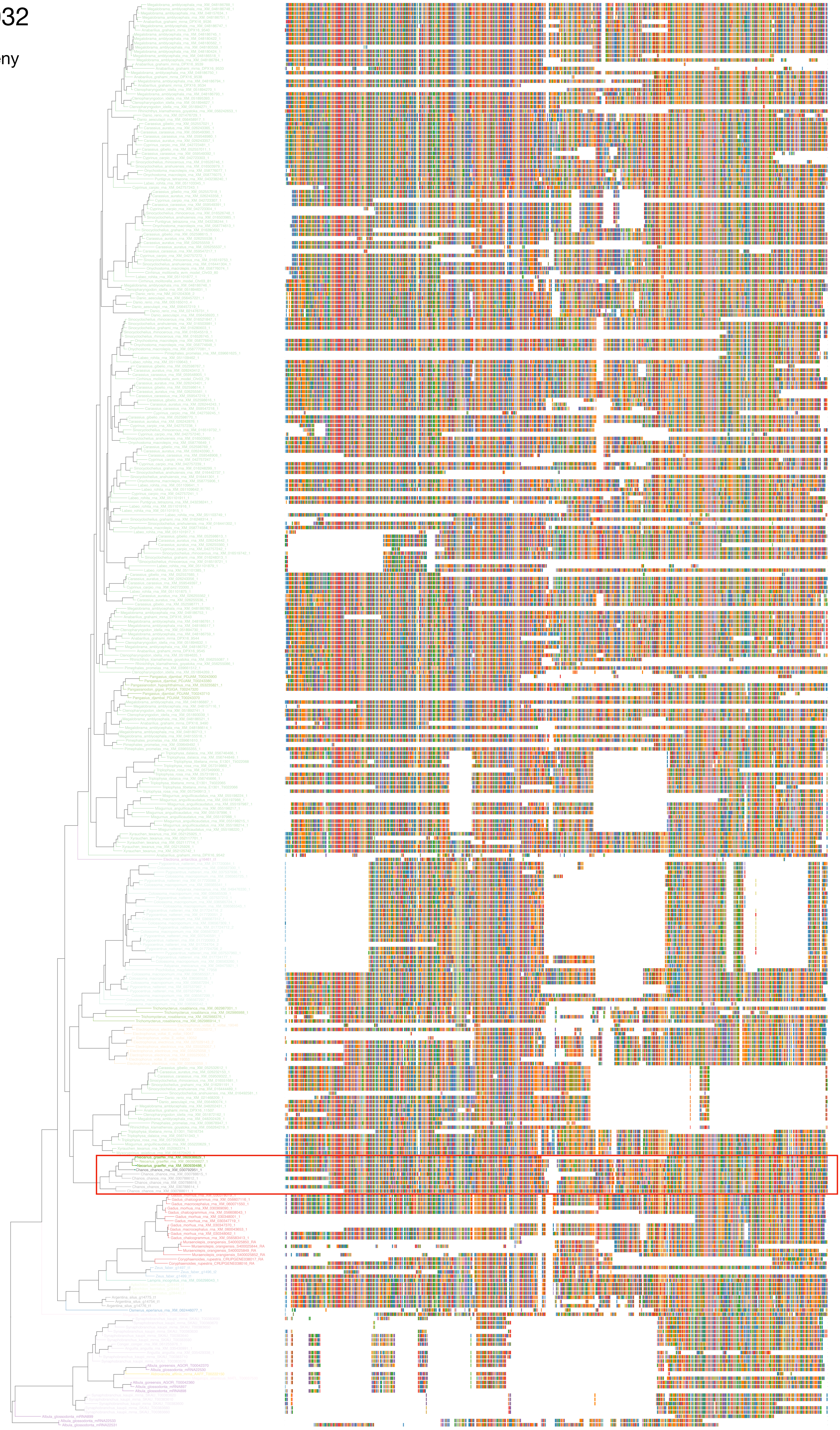

### N5.HOG0019032

Zoom on HGT clade with bootstrap values

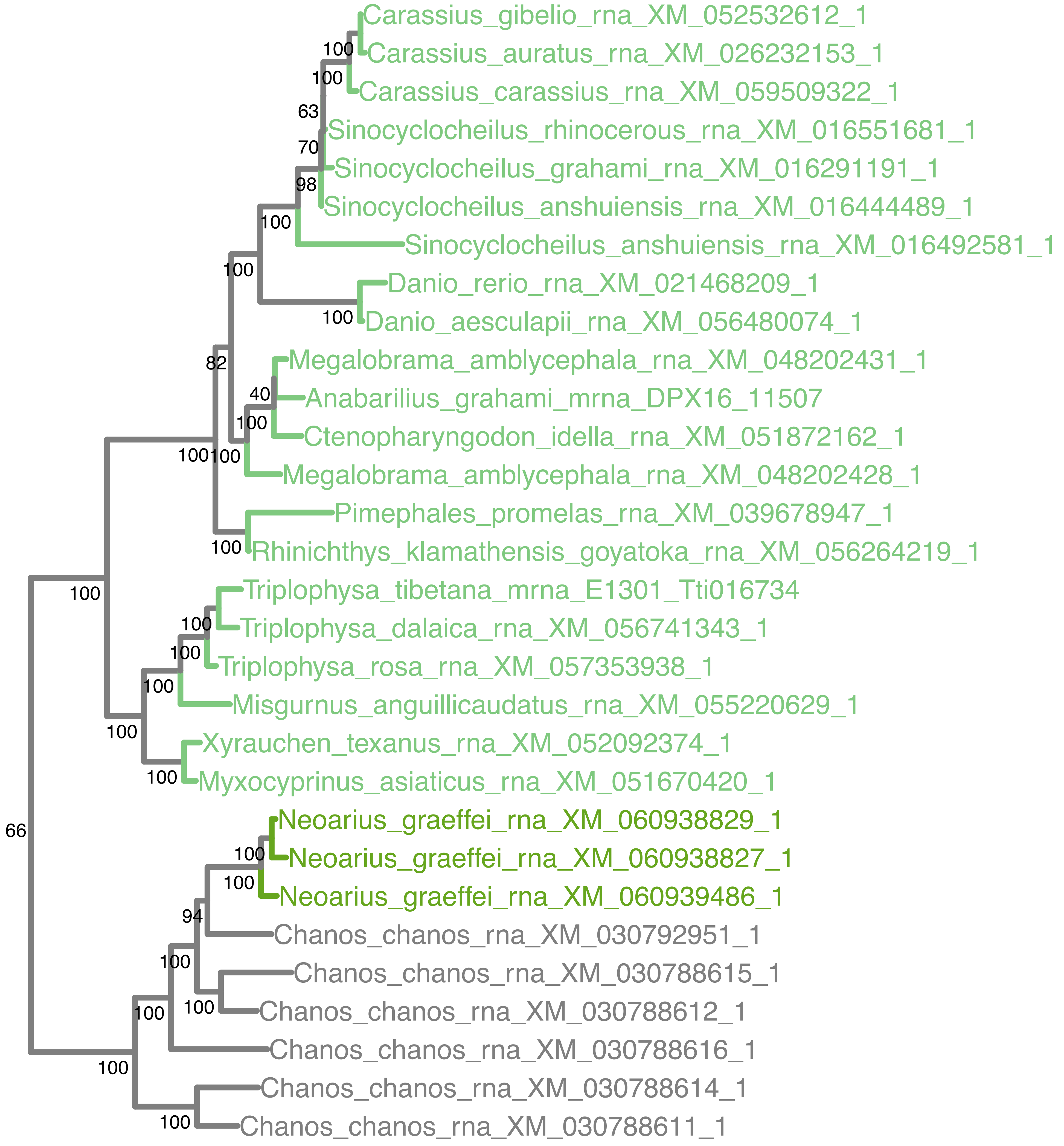

### N5.HOG0019032

Orthogroup phylogeny
and
trimmed
protein alignment

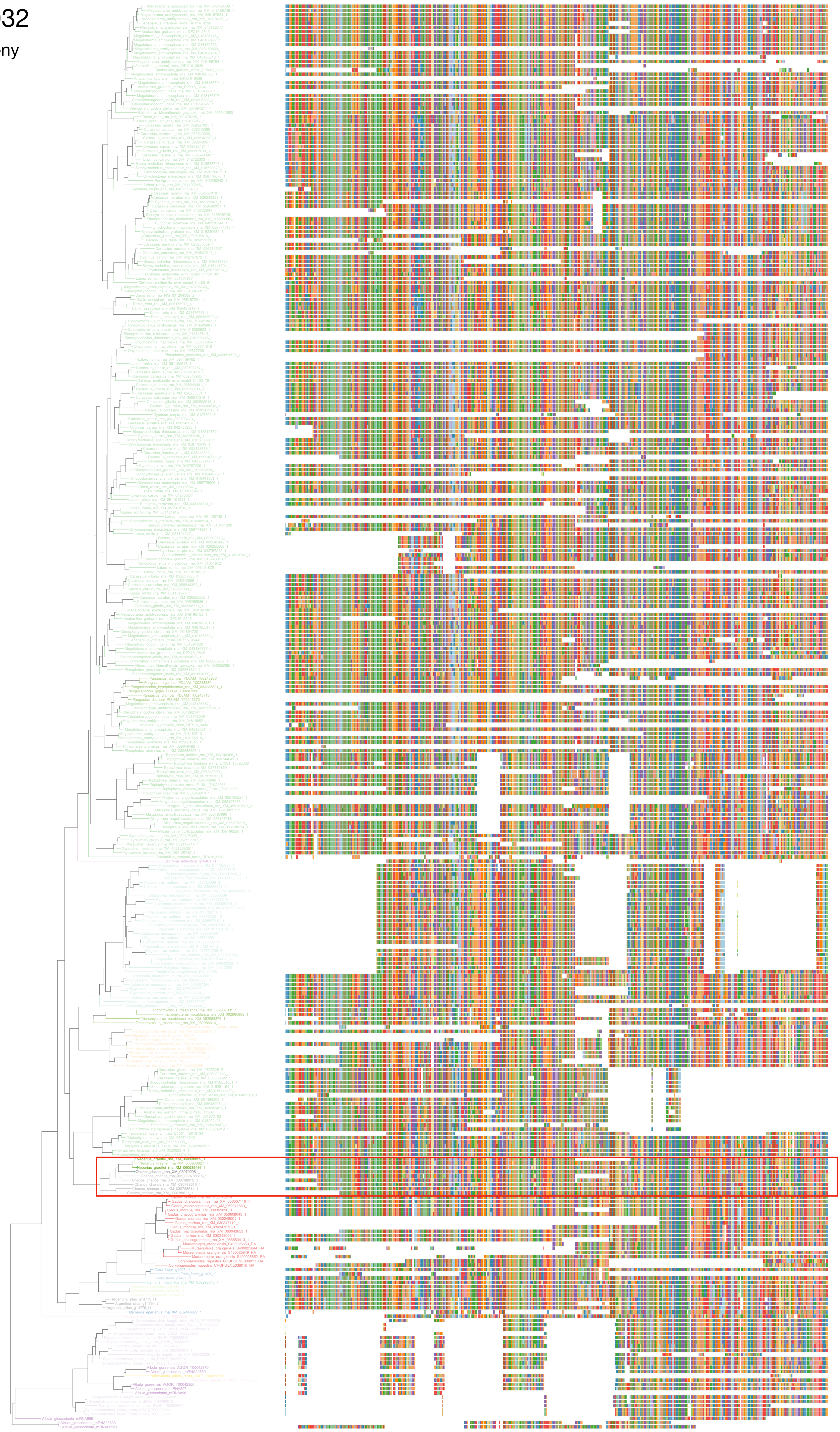

### N5.HOG0019032

« Best-match » phylogeny
and
protein alignment

50 best match against ray-finned fishes protein databases
10 best match against non-fish proteins from UniProt

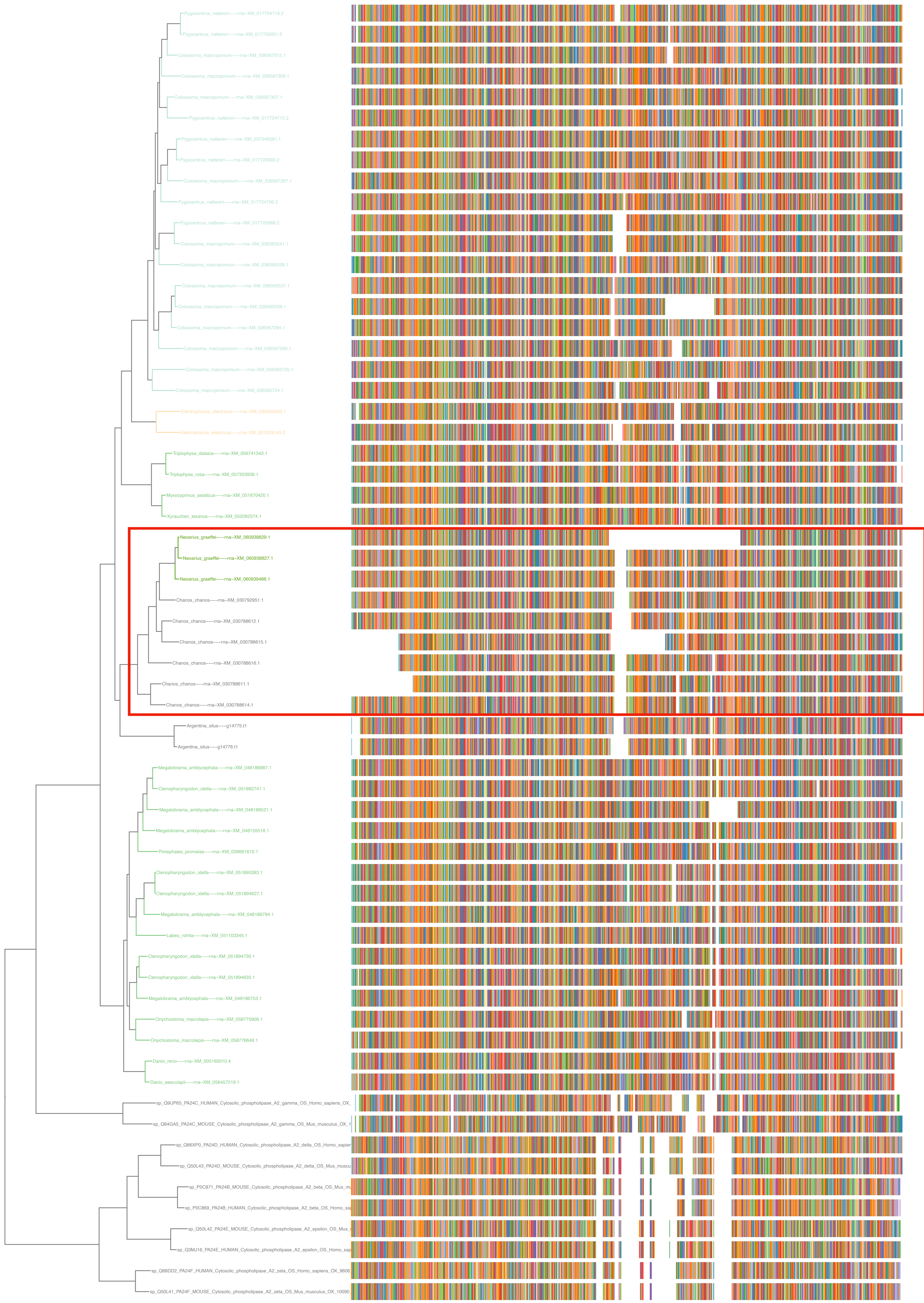

N5.HOG0019032

dS distribution

Inter-species dS distribution

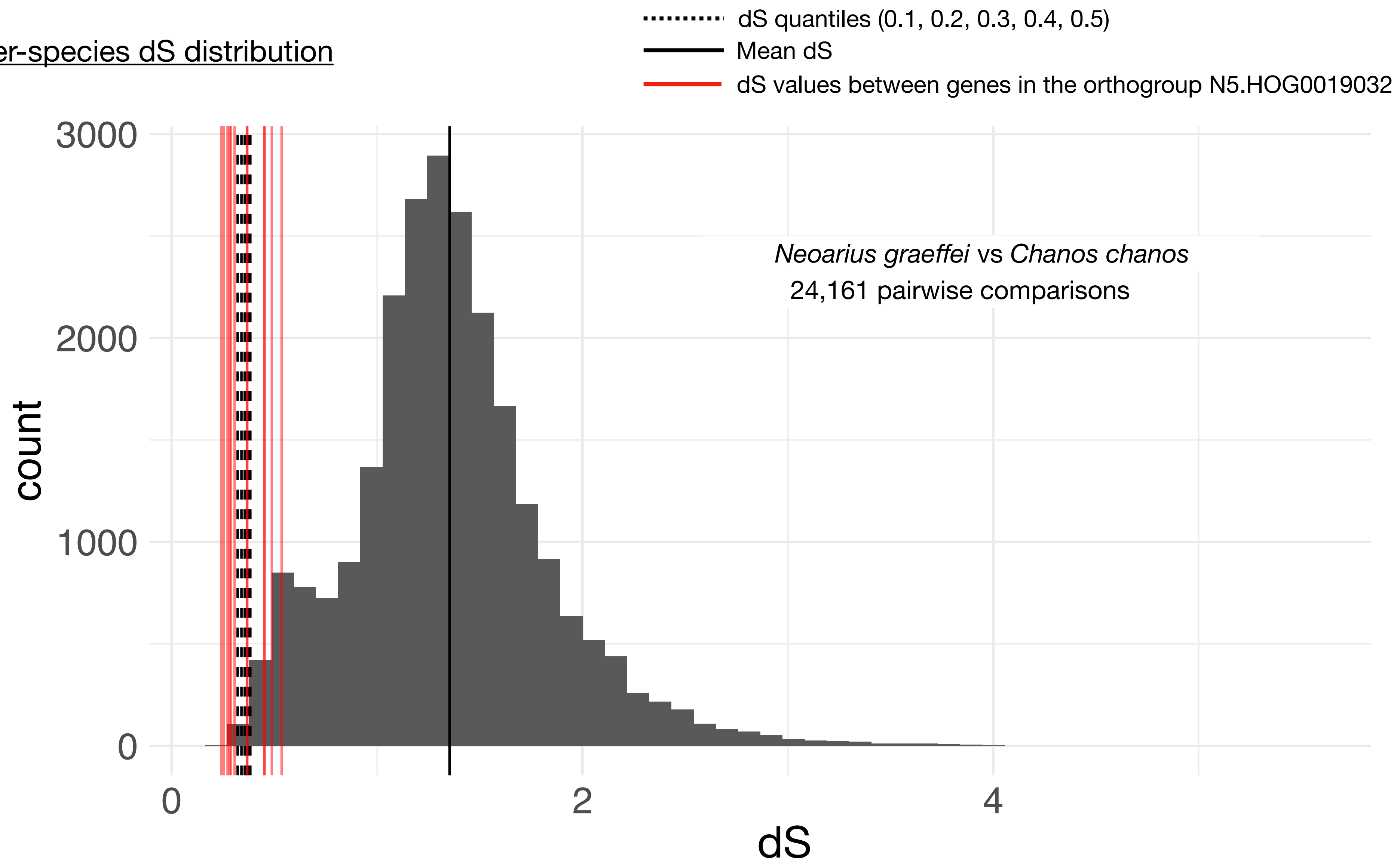


Intra-orthogroup dS

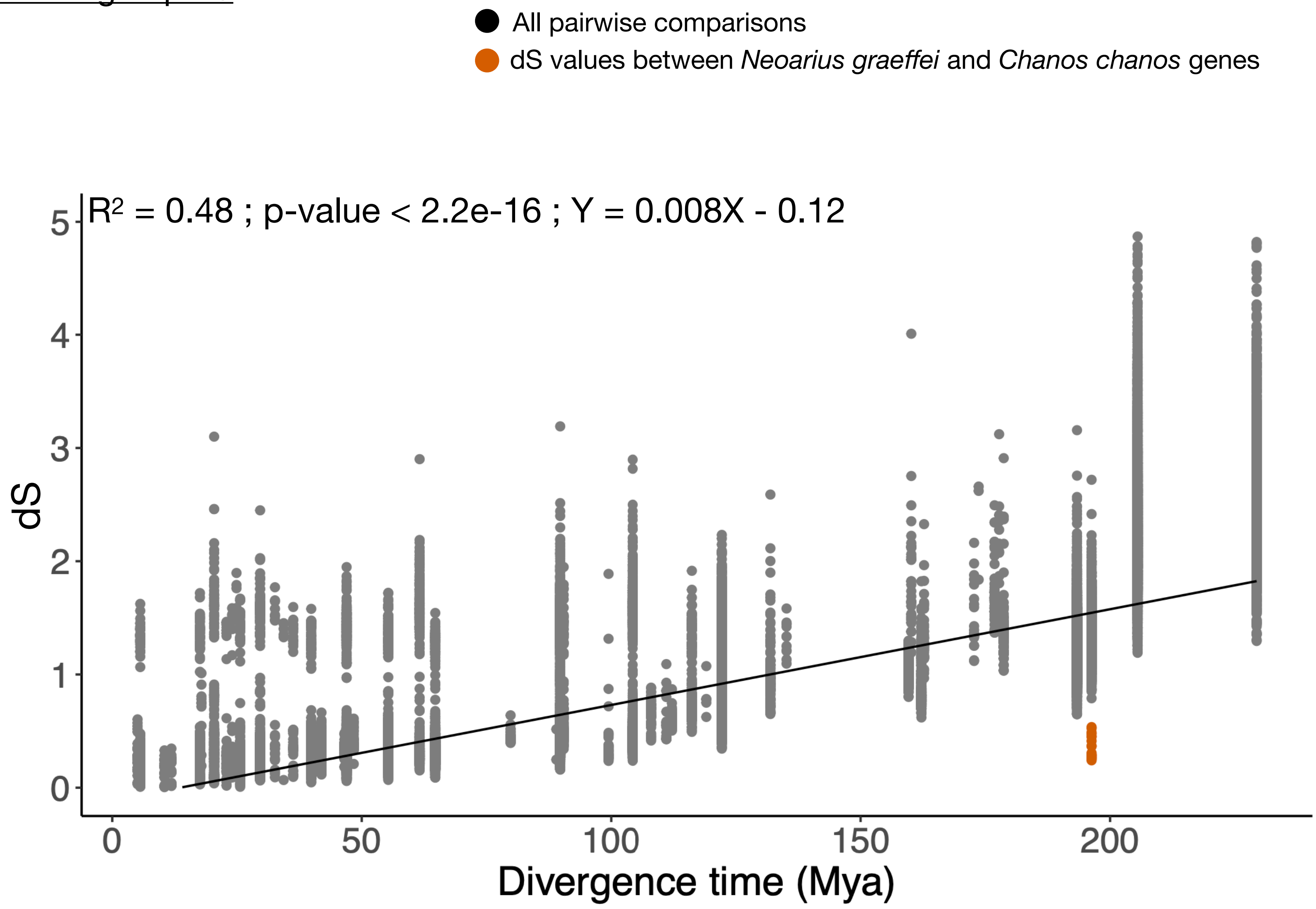

### N5.HOG0019032

#### Micro-synteny

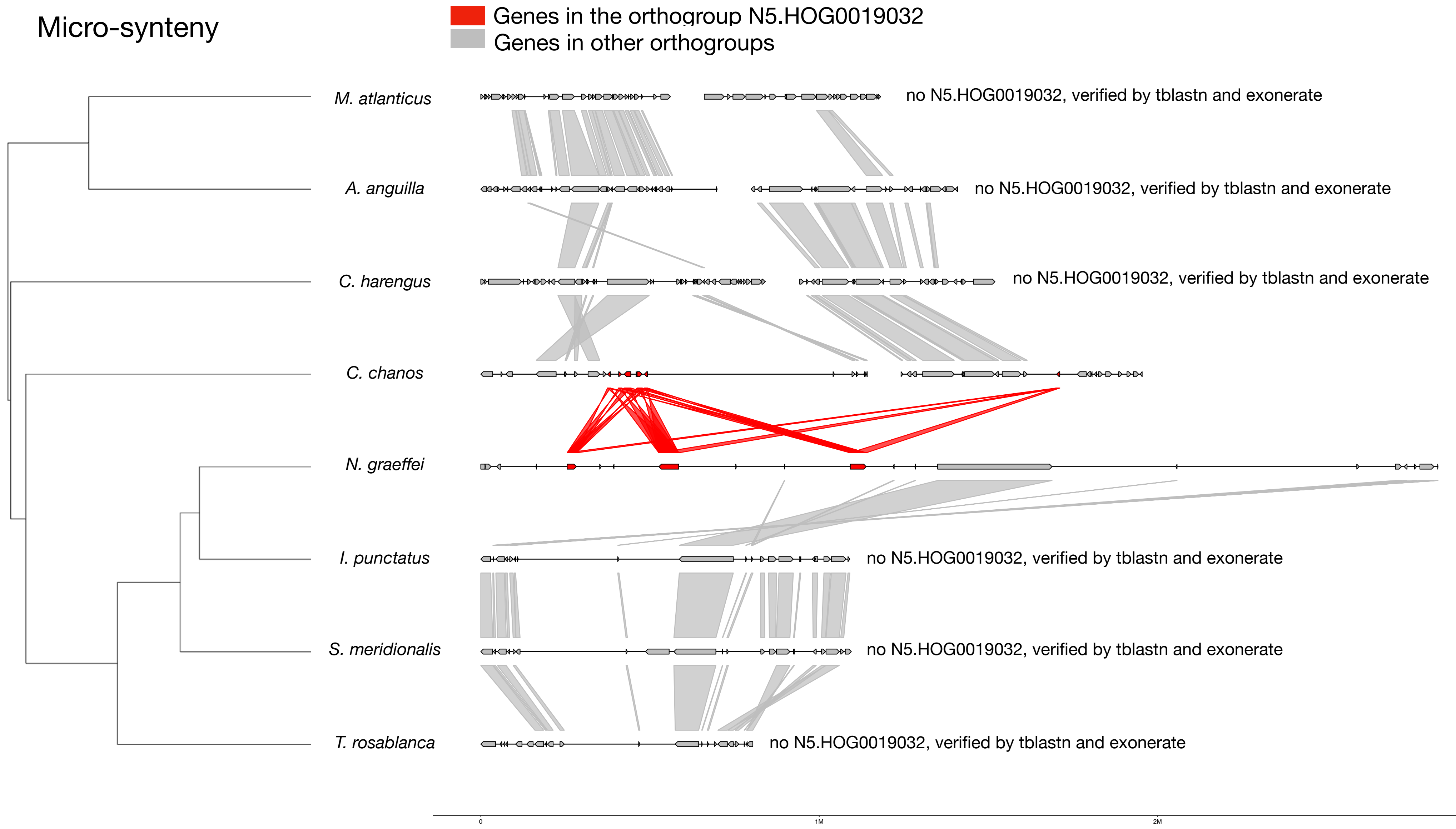


#### Zoom on *N. graeffei* and *C. chanos*

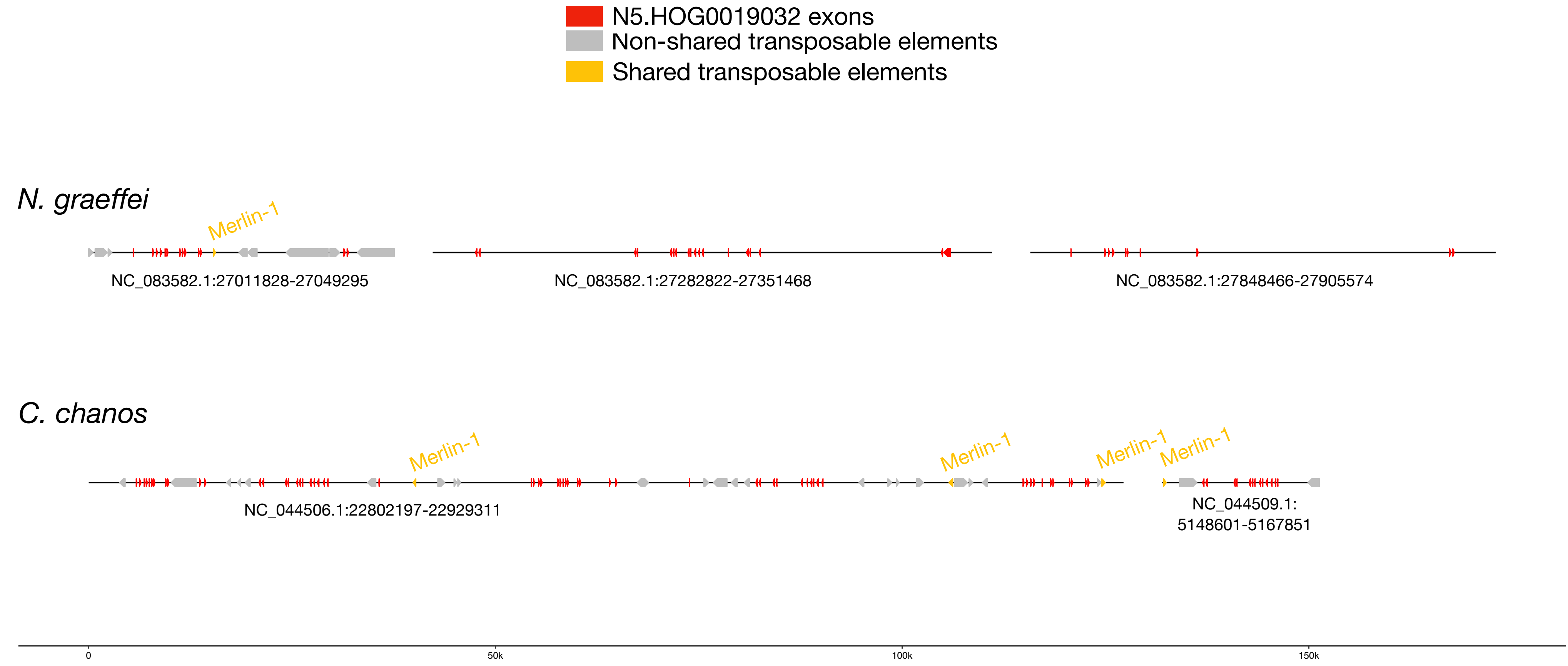


#### Merlin-1:

| | |
|---|---|
| *M. atlanticus* | *166 copies scattered in the genome* |
| *A. anguilla* | *764 copies scattered in the genome* |
| *C. harengus* | *204 copies scattered in the genome* |
| *C. chanos* | *895 copies scattered in the genome* |
| *N. graeffei* | *195,501 copies scattered in the genome* |
| *I. punctatus* | *64 copies scattered in the genome* |
| *S. meridionalis* | *2 copies scattered in the genome* |
| *T. rosablanca* | *27,398 copies scattered in the genome* |

### N5.HOG0019032

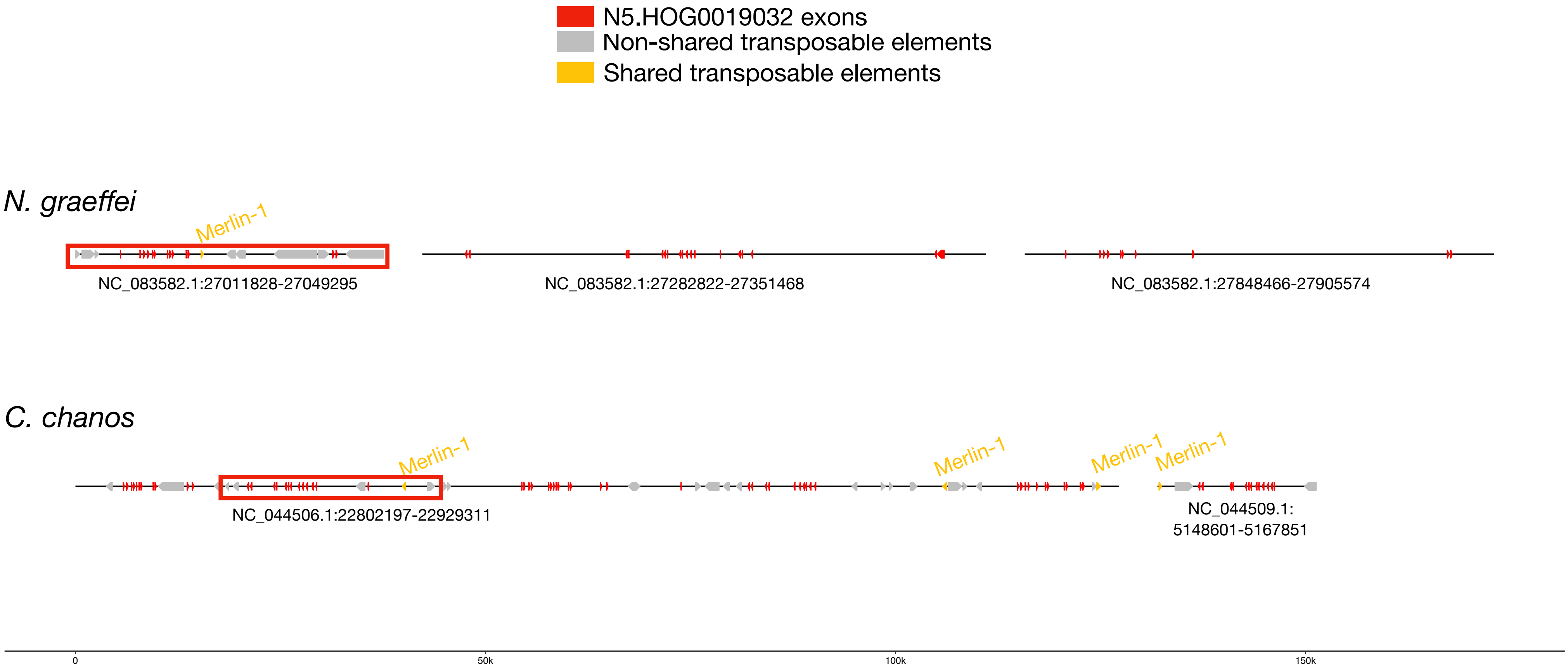


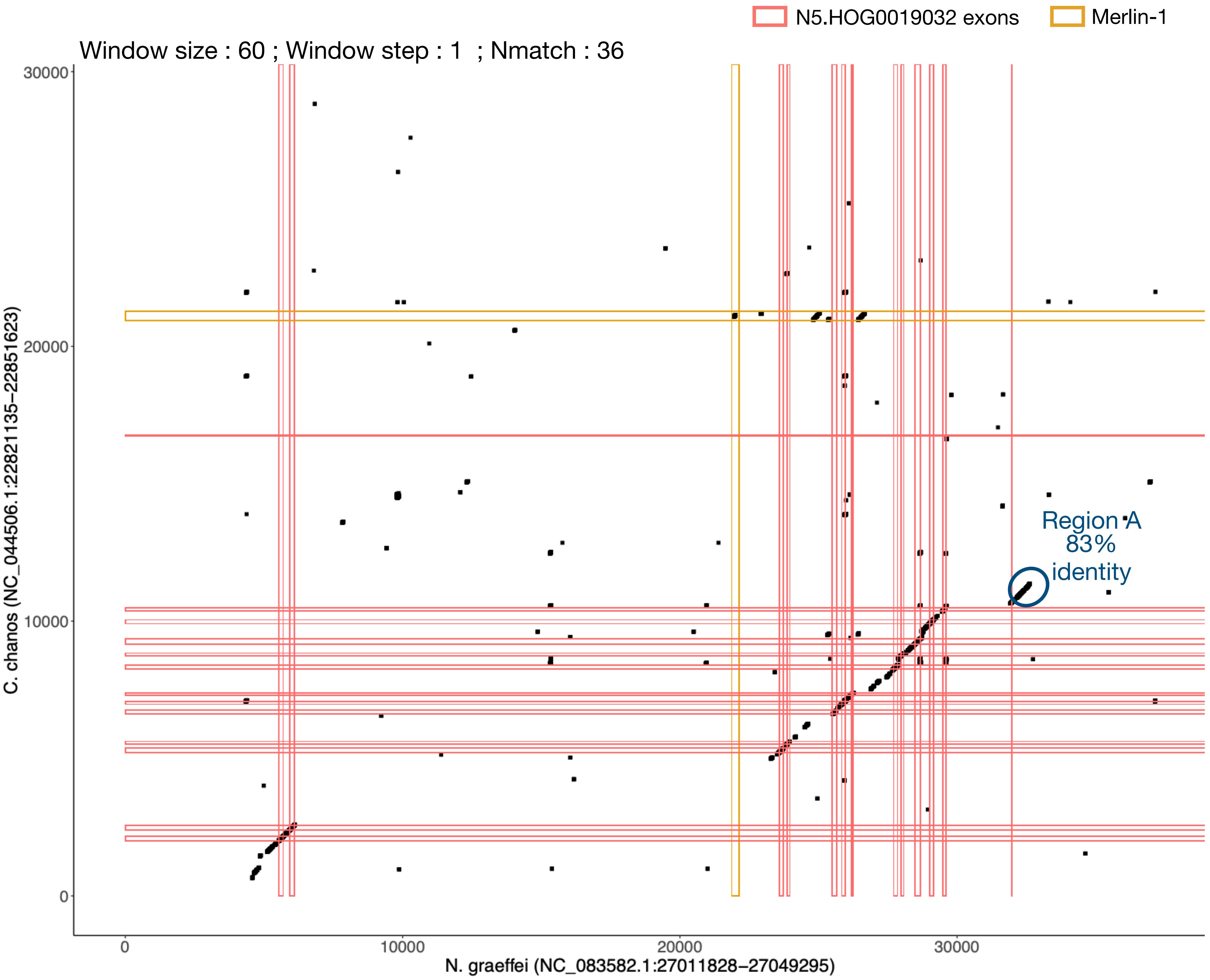

### N5.HOG0019032 Blastn of conserved regions against all genome assemblies

#### Region A

| sseqid | pident | length | mismatch | gapopen | qstart | qend | sstart | send | evalue | bitscore |
|---|---|---|---|---|---|---|---|---|---|---|
| Neoarius_graeffei-NC_083582.1 | 100.000 | 711 | 0 | 0 | 1 | 711 | 27017306 | 27016596 | 0.0 | 1314 |
| Neoarius_graeffei-NC_083582.1 | 97.001 | 667 | 17 | 1 | 1 | 667 | 5202069 | 5201406 | 0.0 | 1118 |
| Neoarius_graeffei-NC_083582.1 | 97.101 | 552 | 10 | 3 | 1 | 552 | 27319191 | 27319736 | 0.0 | 926 |
| Neoarius_graeffei-NC_083582.1 | 96.022 | 553 | 15 | 3 | 1 | 552 | 27853462 | 27852916 | 0.0 | 893 |
| Neoarius_graeffei-NC_083582.1 | 92.516 | 628 | 29 | 7 | 1 | 623 | 27639400 | 27640014 | 0.0 | 883 |
| Neoarius_graeffei-NC_083582.1 | 92.871 | 533 | 26 | 4 | 1 | 523 | 27229552 | 27230082 | 0.0 | 763 |
| Neoarius_graeffei-NC_083582.1 | 93.651 | 504 | 9 | 2 | 208 | 711 | 27147231 | 27146751 | 0.0 | 732 |
| Neoarius_graeffei-NC_083582.1 | 92.262 | 504 | 12 | 3 | 208 | 711 | 27577759 | 27577283 | 0.0 | 689 |
| Neoarius_graeffei-NC_083582.1 | 99.052 | 211 | 2 | 0 | 1 | 211 | 27587109 | 27586899 | 8.39e-101 | 379 |
| Neoarius_graeffei-NC_083582.1 | 95.783 | 166 | 2 | 1 | 551 | 711 | 27335051 | 27335216 | 8.82e-66 | 263 |
| Neoarius_graeffei-NC_083582.1 | 95.783 | 166 | 2 | 1 | 551 | 711 | 27830113 | 27829948 | 8.82e-66 | 263 |
| Neoarius_graeffei-NC_083582.1 | 99.099 | 111 | 1 | 0 | 1 | 111 | 27164724 | 27164614 | 7.01e-47 | 200 |
| Neoarius_graeffei-NC_083582.1 | 100.000 | 97 | 0 | 0 | 102 | 198 | 27162264 | 27162168 | 9.14e-41 | 180 |
| Chanos_chanos-NC_044509.1 | 80.068 | 587 | 95 | 14 | 1 | 576 | 5162855 | 5163430 | 6.40e-112 | 416 |
| Chanos_chanos-NC_044506.1 | 78.971 | 447 | 78 | 10 | 197 | 630 | 22832019 | 22832462 | 4.05e-74 | 291 |

N5.HOG0019032 can be retrieved on the genome assemblies of other *Neoarius* species (*N. berneyi*, *N. pectoralis* and *N. leptaspis*). It can also be retrieved in an alternative assembly of *Chanos chanos* (GCA_018691325.1).

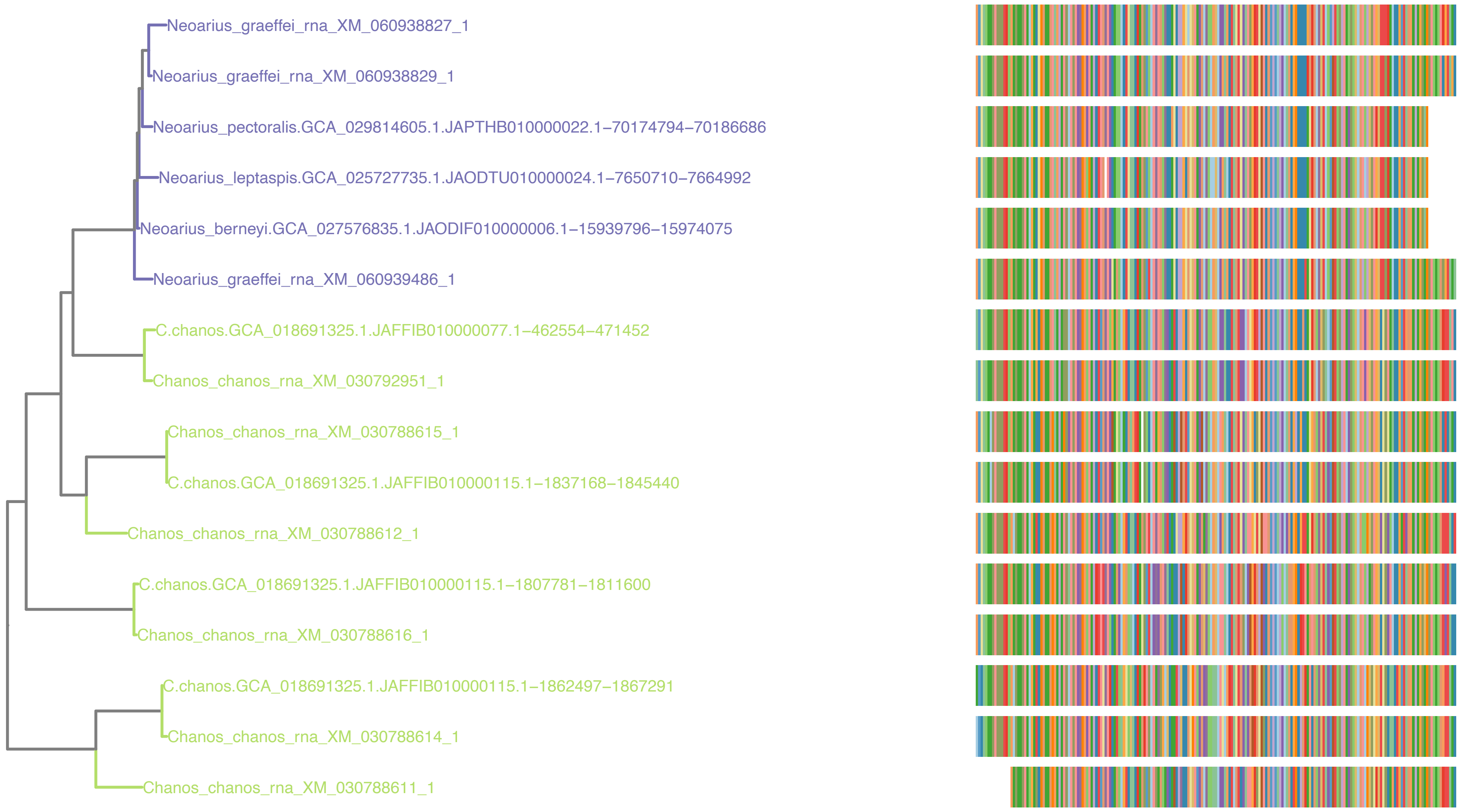

N5.HOG0019032

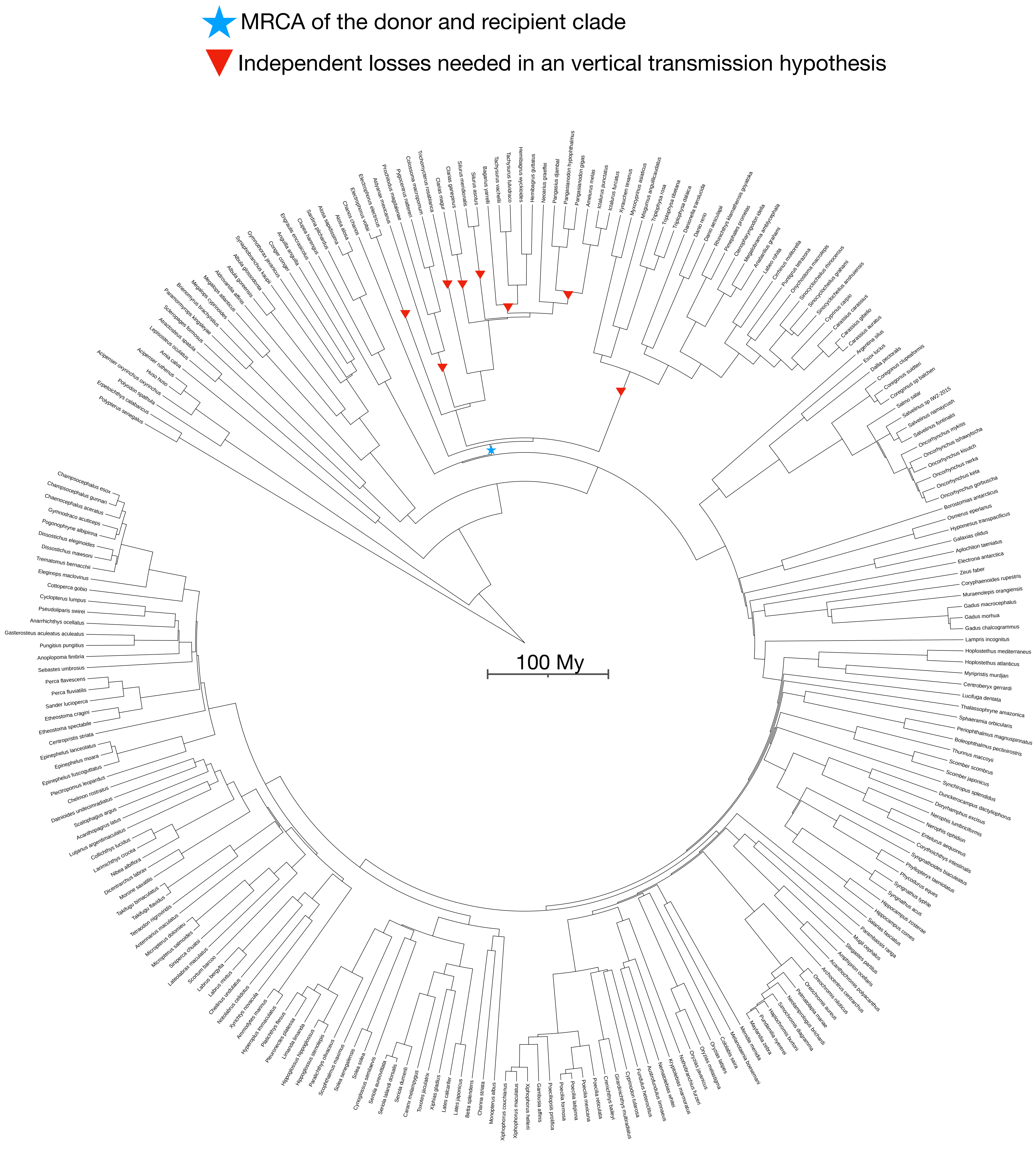

N5.HOG0004451

### N5.HOG0004451

Species tree

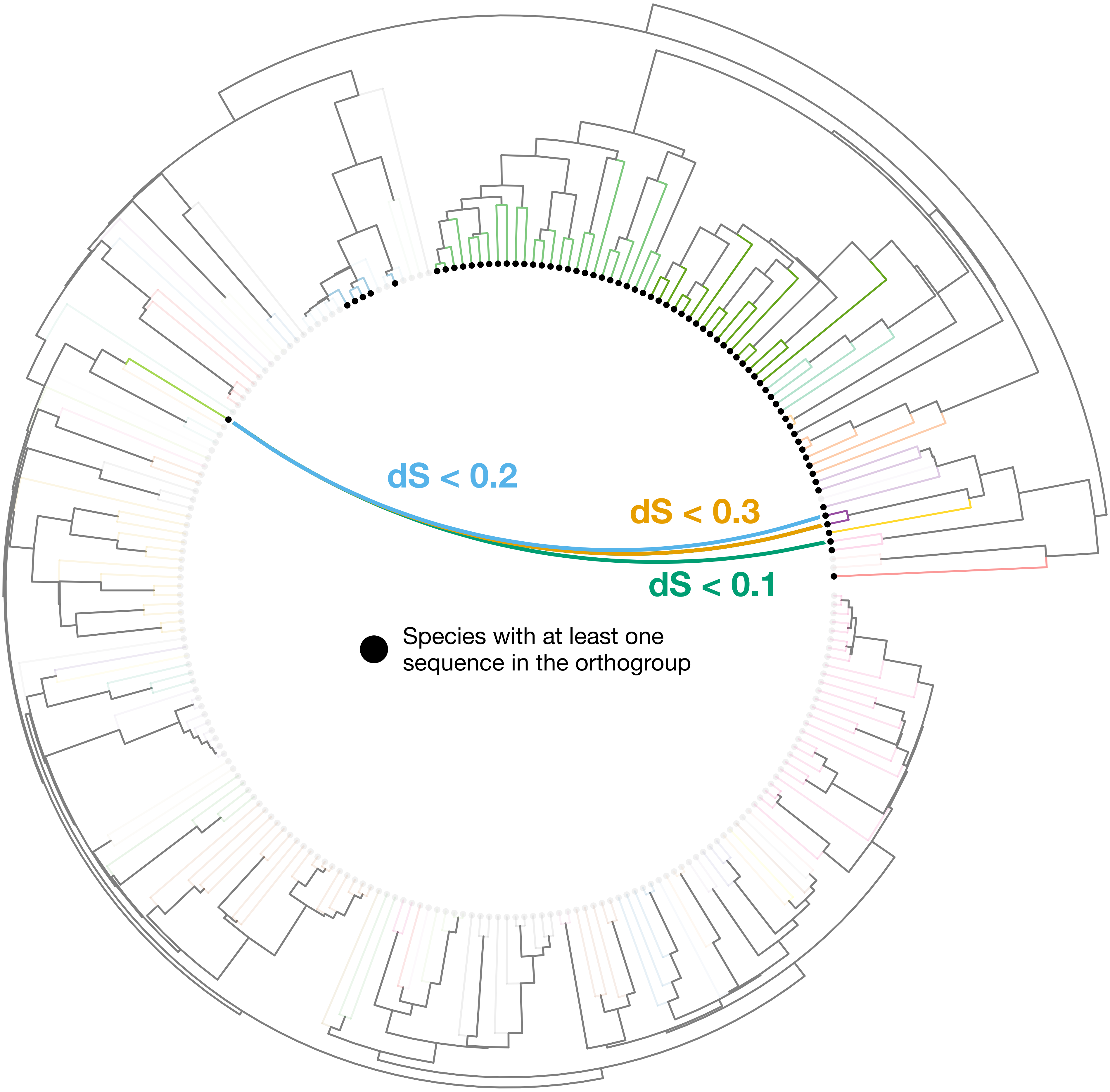

### N5.HOG0004451

Orthogroup phylogeny
and
protein alignment

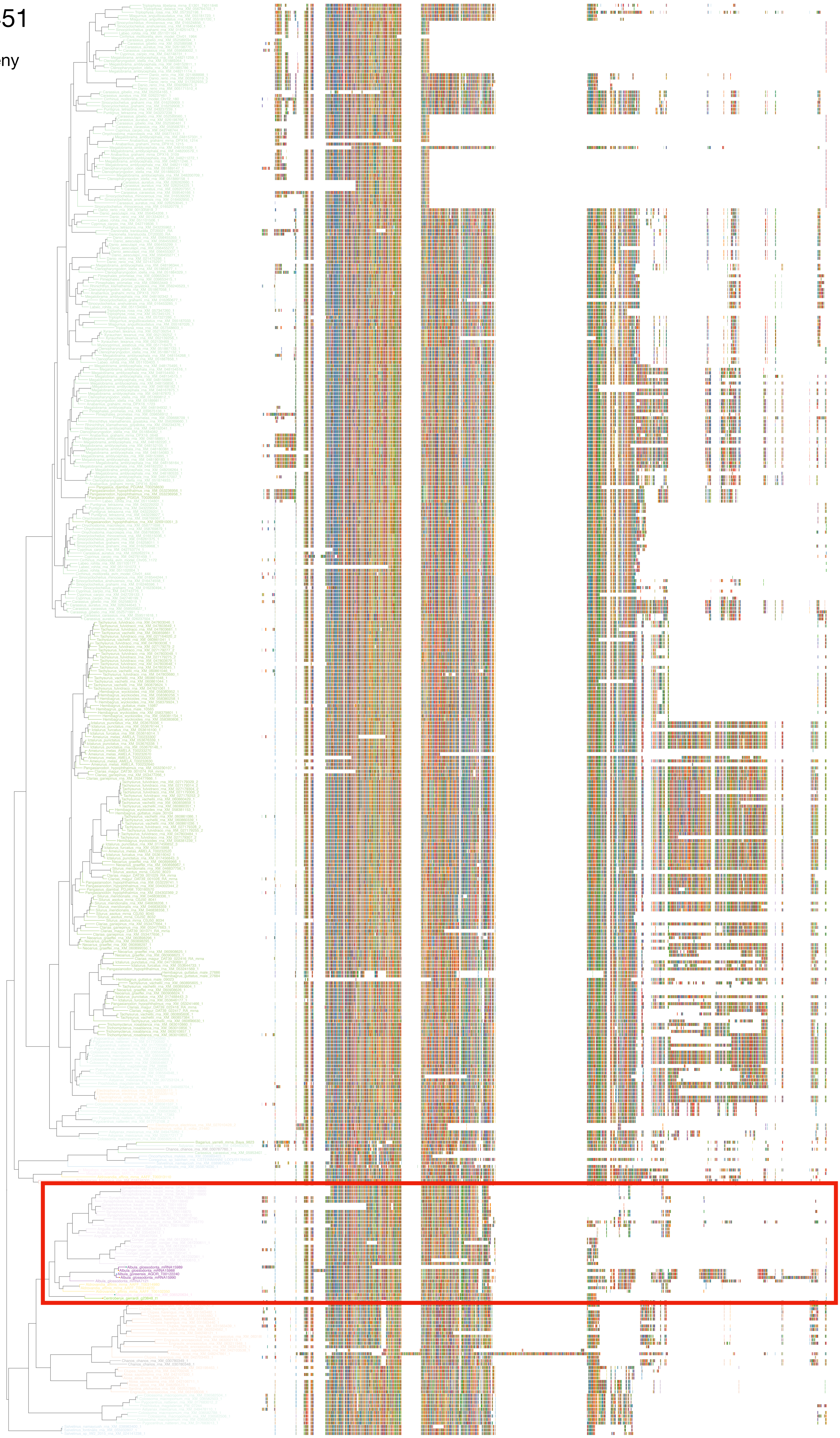

### N5.HOG0004451

Zoom on HGT clade with bootstrap values

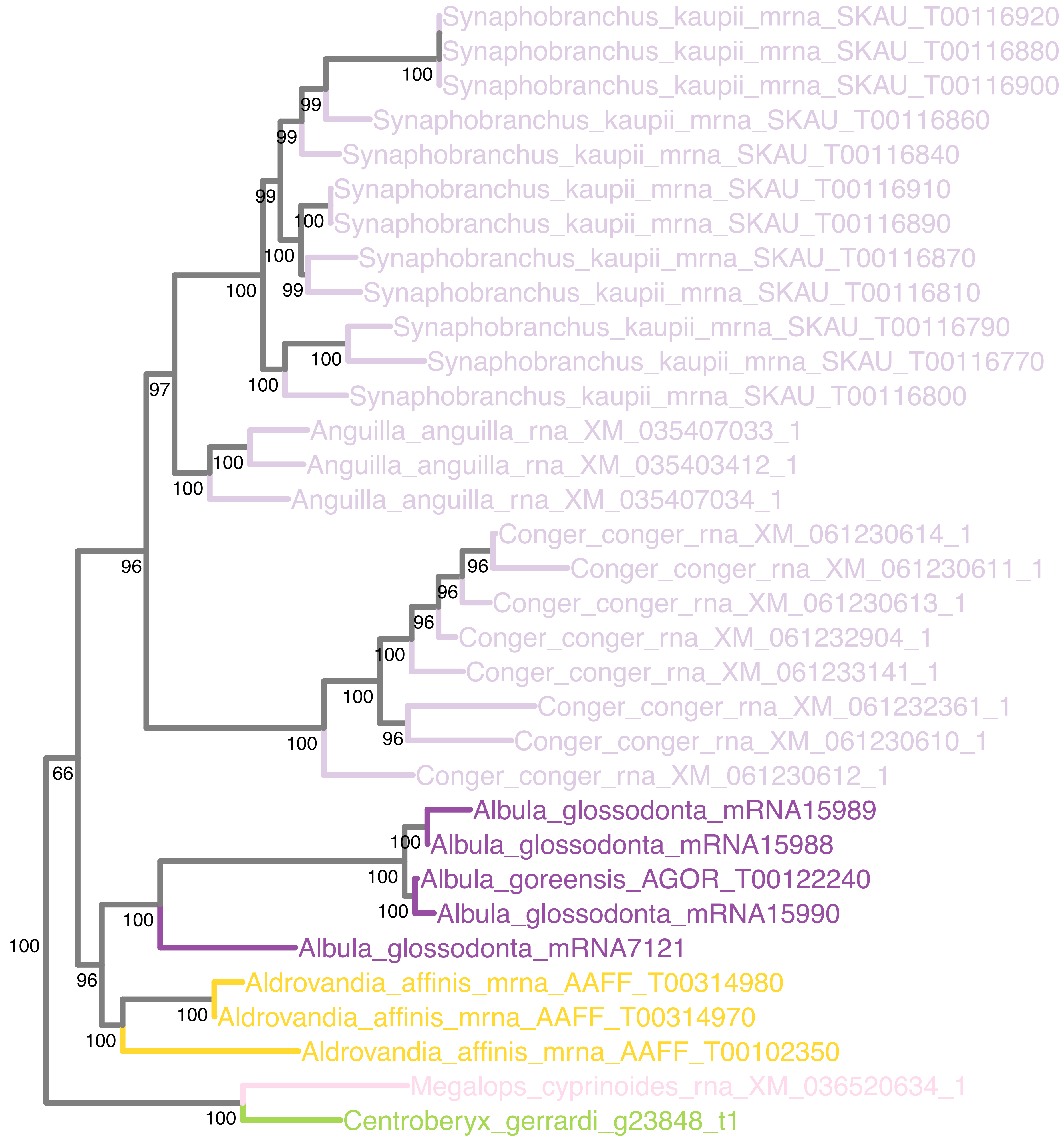

### N5.HOG0004451

Orthogroup phylogeny
and
trimmed
protein alignment

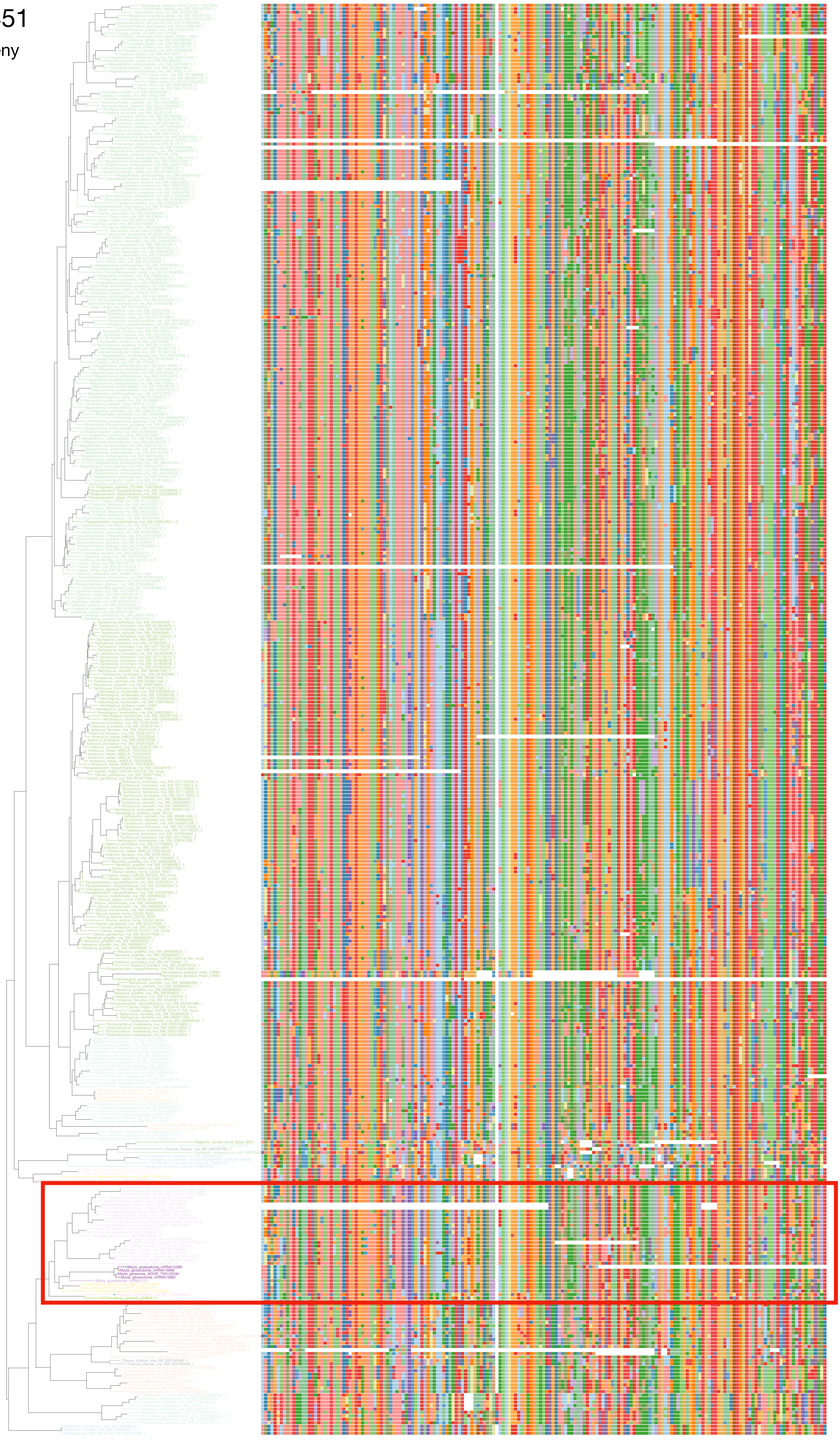

### N5.HOG0004451

« Best-match » phylogeny
and
protein alignment

50 best match against ray-finned fishes protein databases
10 best match against non-fish proteins from UniProt

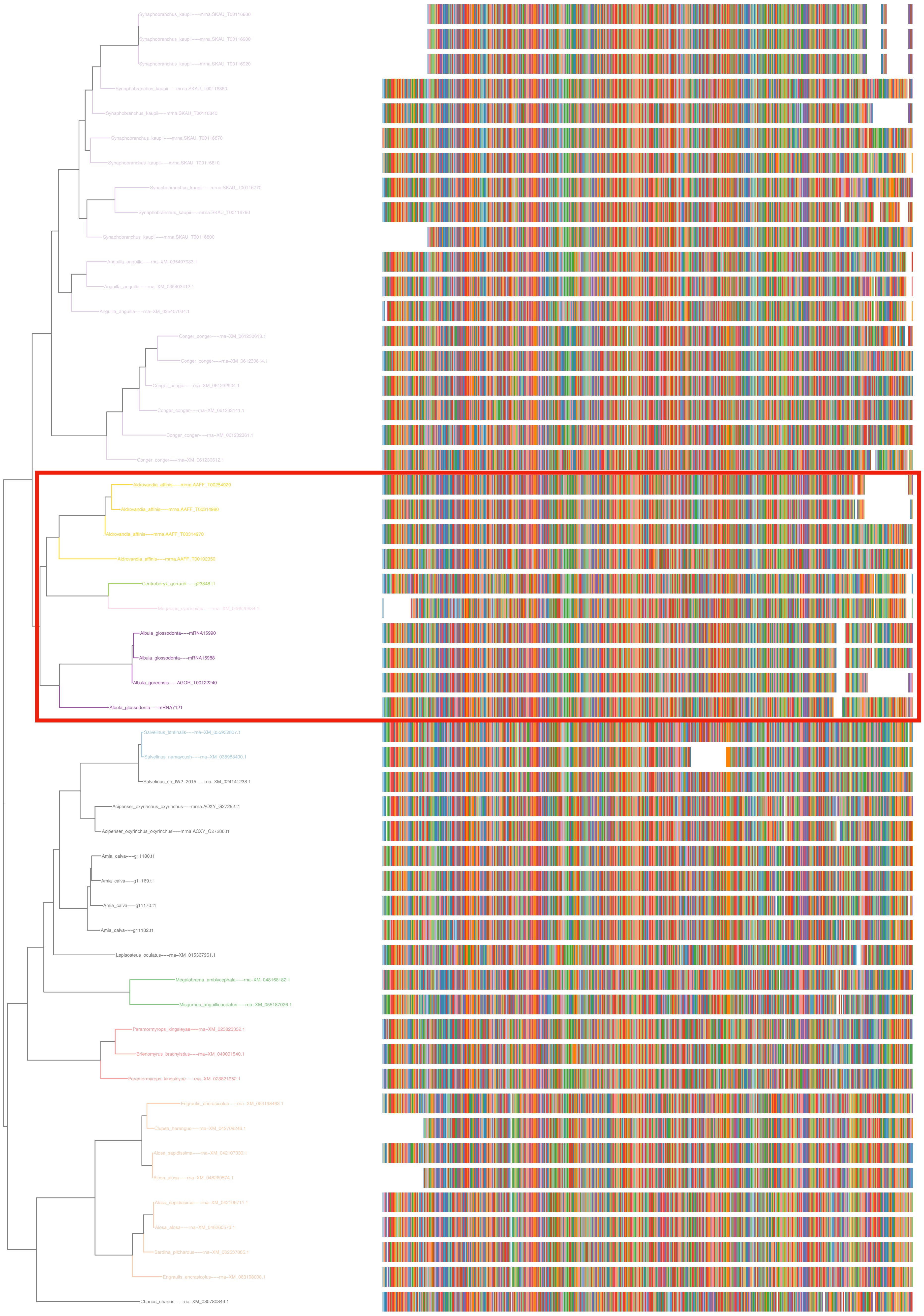

### N5.HOG0004451

#### dS distribution

dS quantiles (0.1, 0.2, 0.3, 0.4, 0.5)
Mean dS
dS values between genes in the orthogroup N5.HOG0004451

##### Inter-species dS distribution

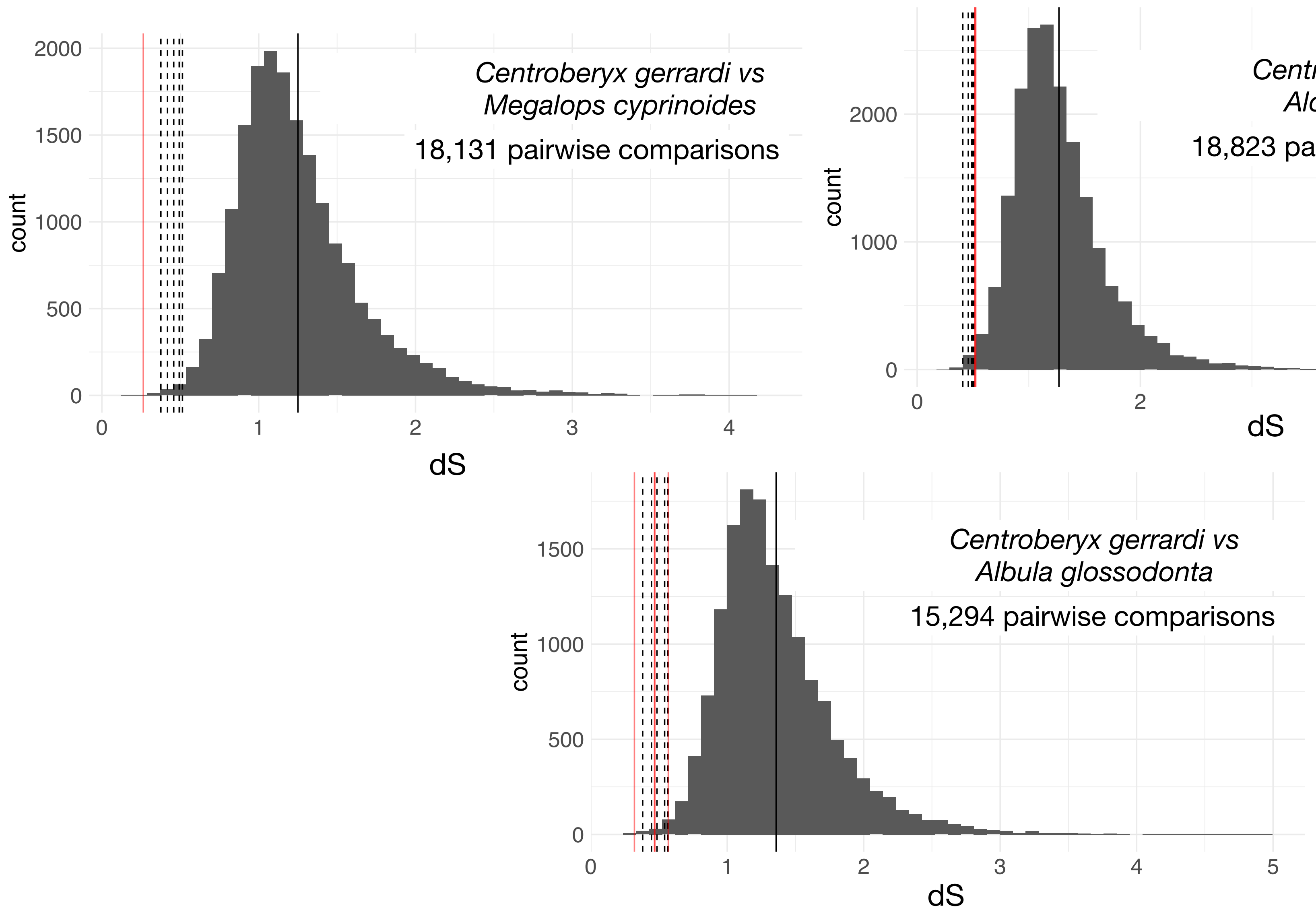


##### Intra-orthogroup dS

All pairwise comparisons
dS values between *Centroberyx gerrardi* and *Megalops cyprinoides* genes
dS values between *Centroberyx gerrardi*and *Aldrovandia affinis* genes
dS values between *Centroberyx gerrardi* and *Albula glossodonta* genes

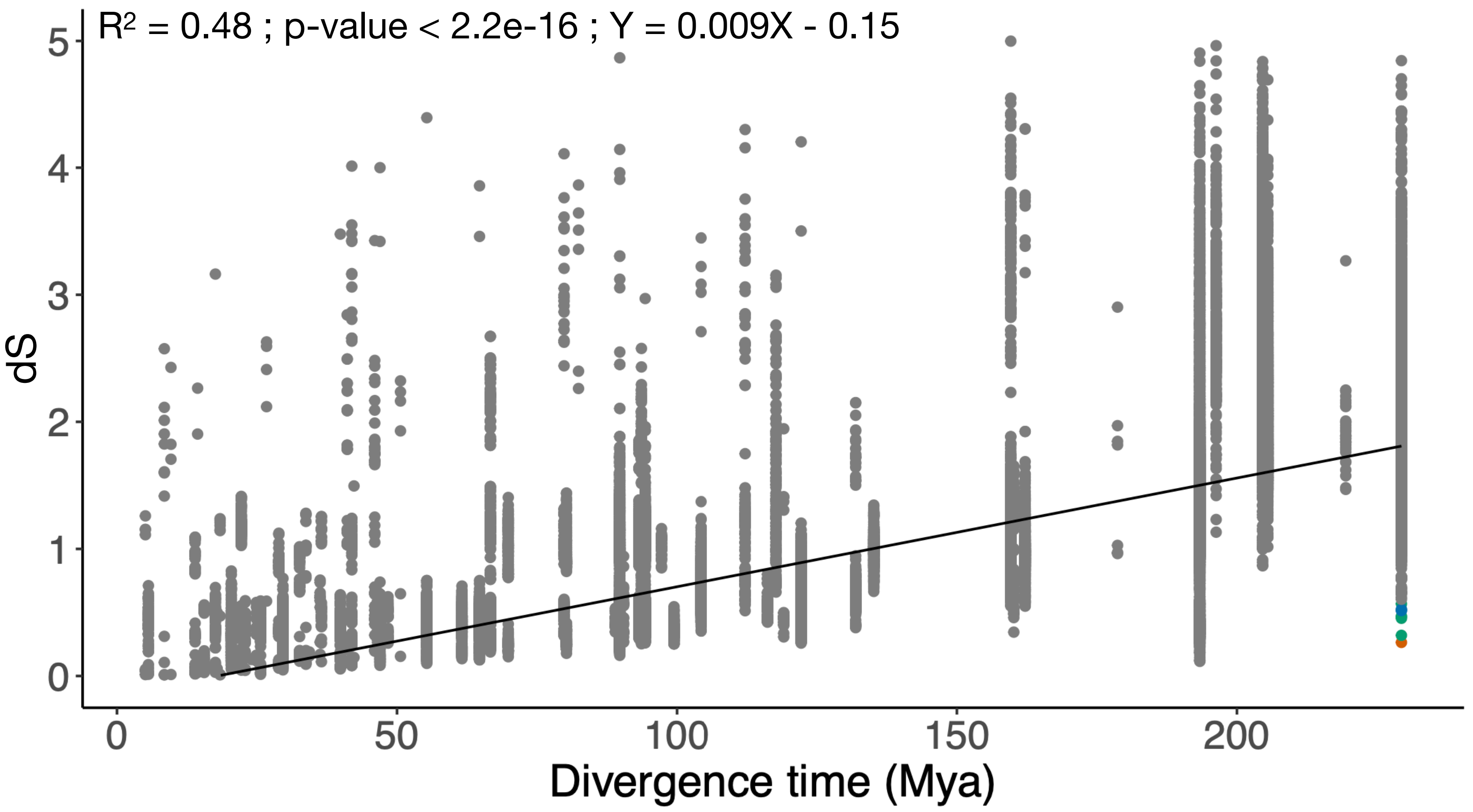

### N5.HOG0004451

#### Micro-synteny

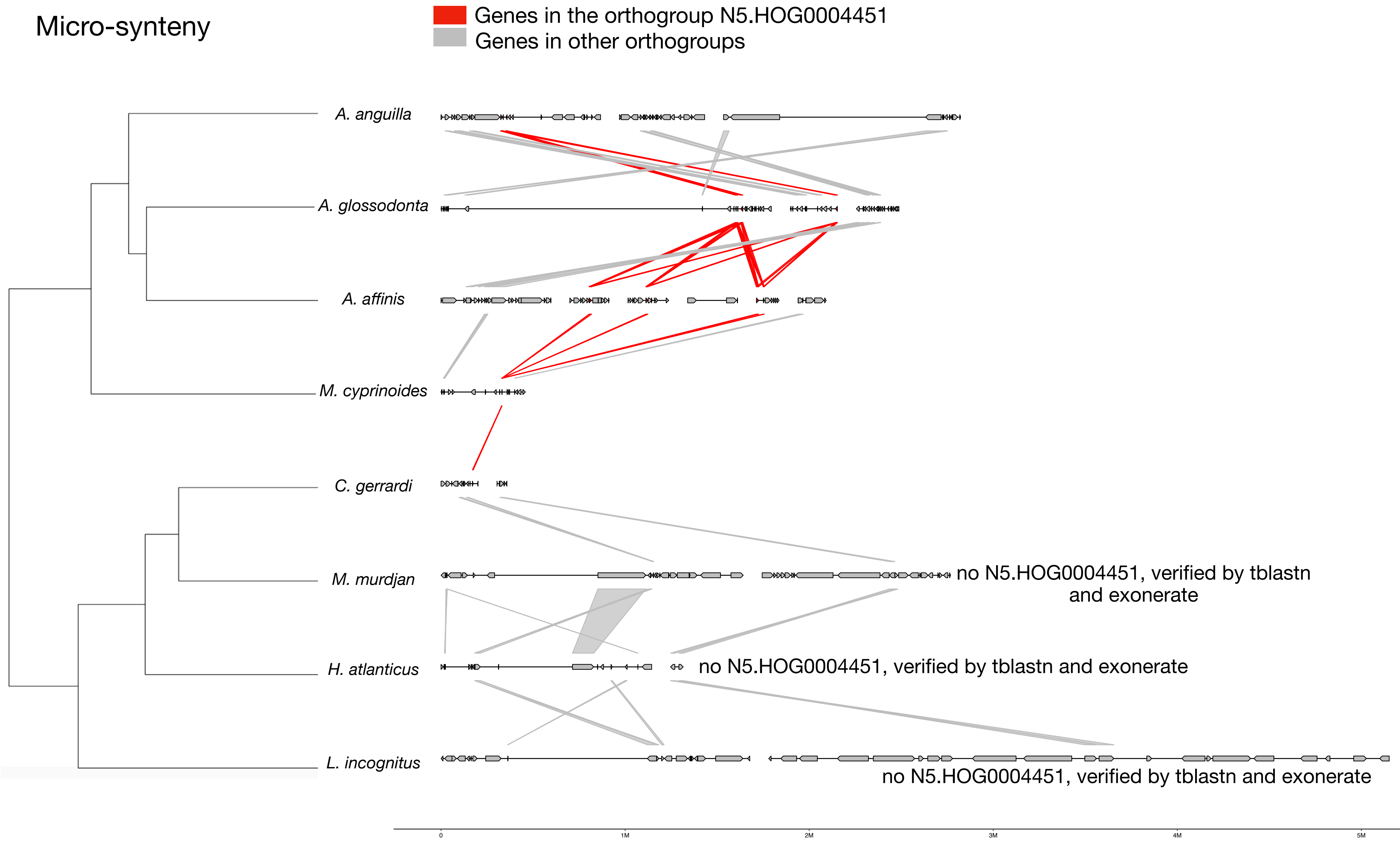


#### Zoom on *M. cyprinoides* and *C. gerrardi*

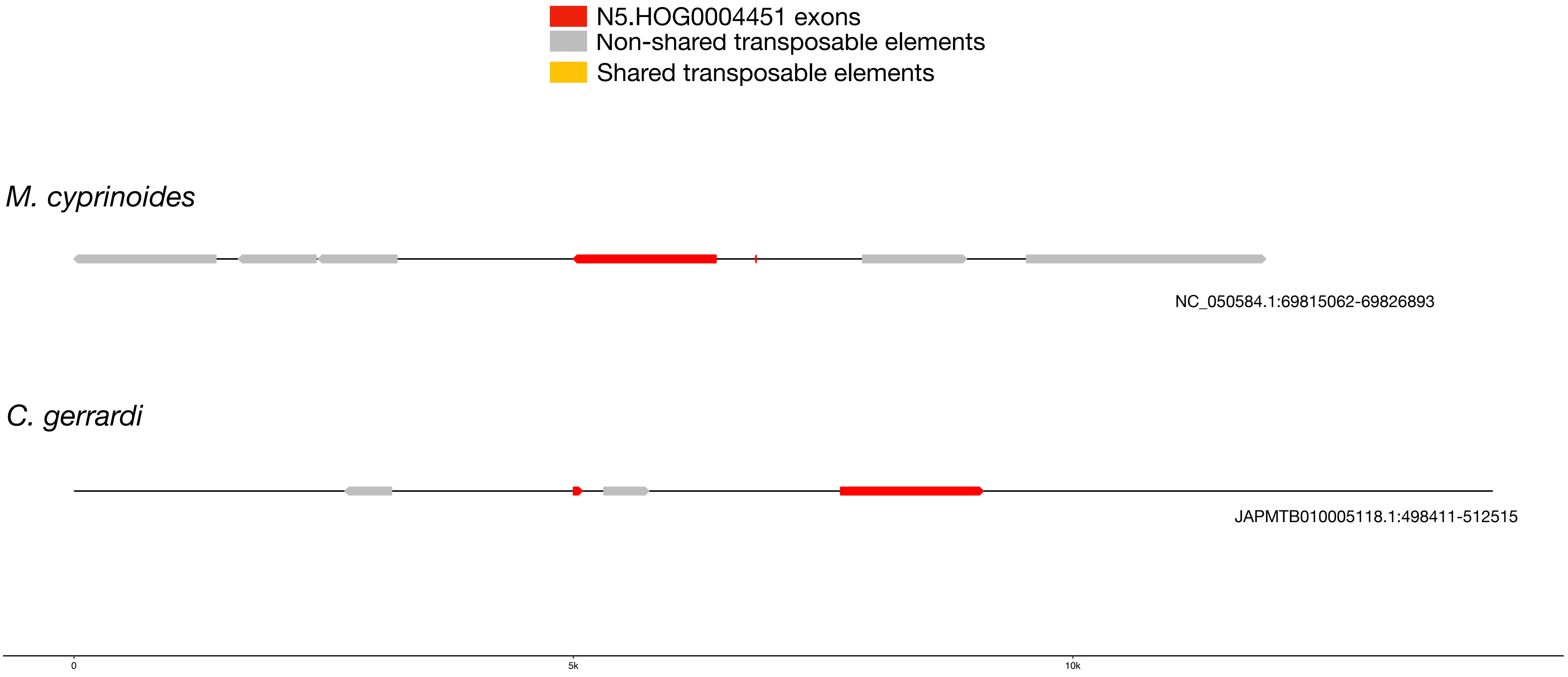

N5.HOG0004451

N5.HOG0004451 can be retrieved on the genome of a *Centroberyx gerrardi* closely related species (*Beryx splendens*)

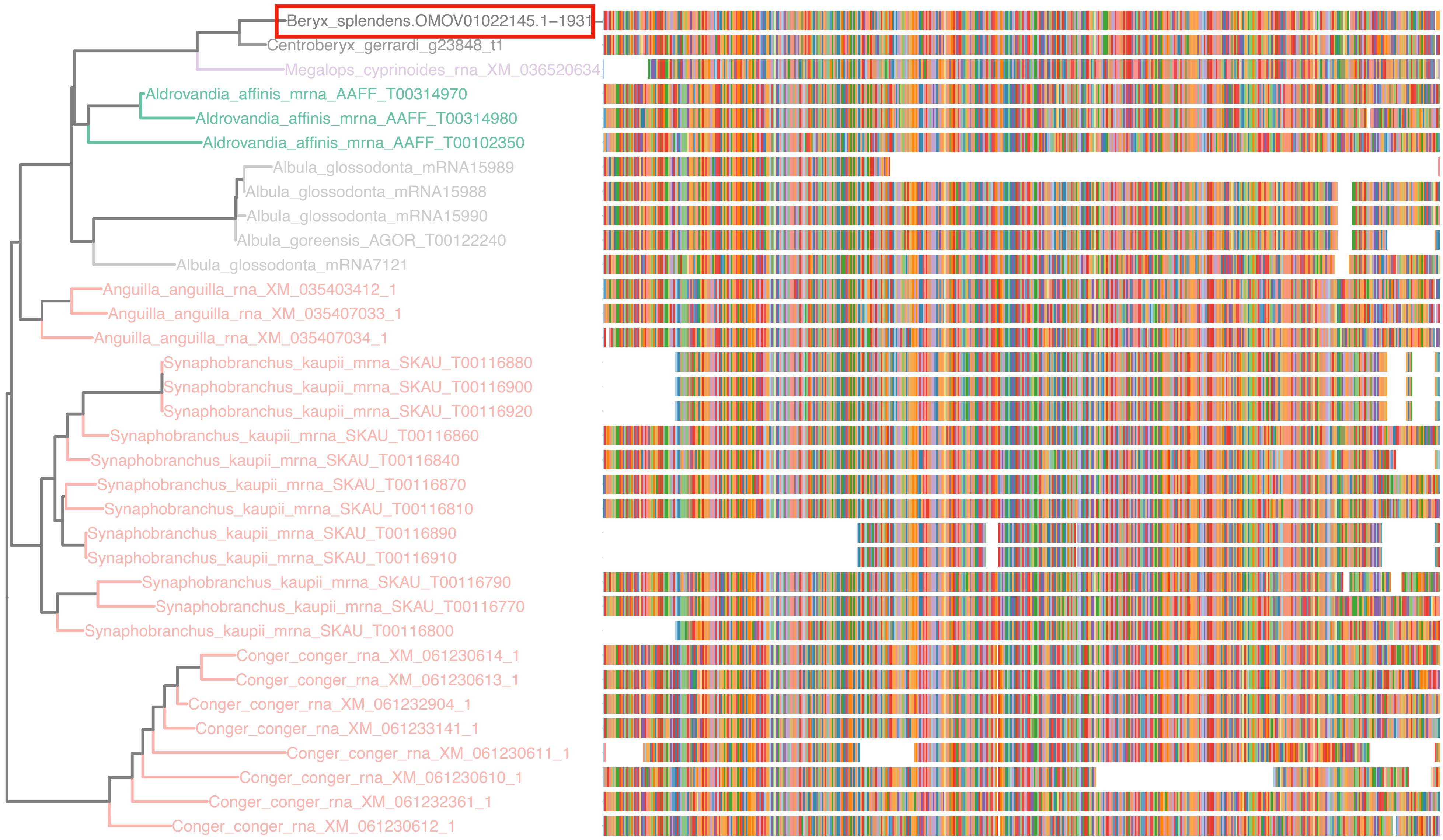

### N5.HOG0004451

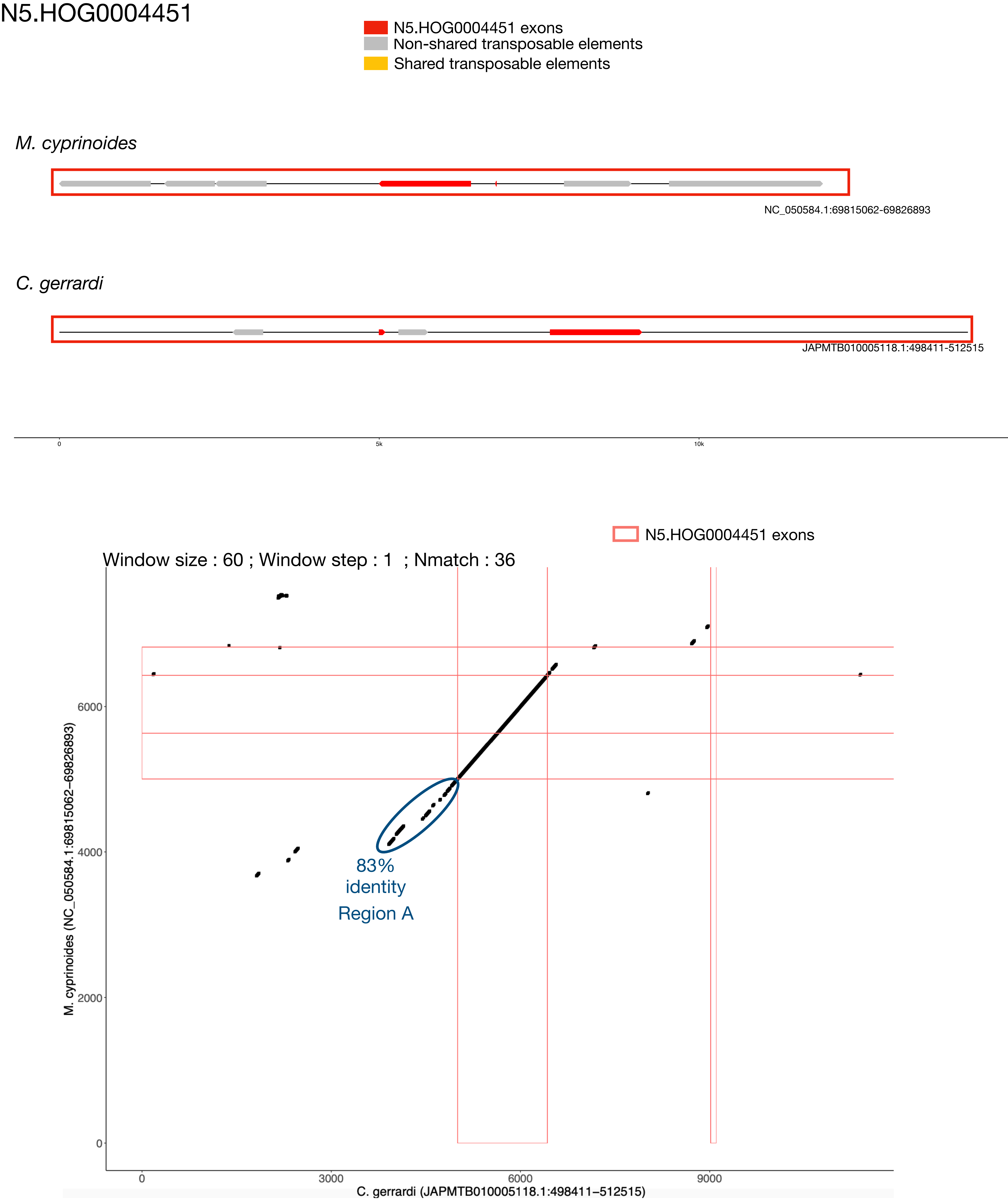

N5.HOG0004451 Blastn of conserved regions against all genome assemblies

#### Region A

| sseqid | pident | length | mismatch | gapopen | qstart | qend | sstart | send | evalue | bitscore |
|---|---|---|---|---|---|---|---|---|---|---|
| Centroberyx_gerrardi-JAPMTB010005118.1 | 100.000 | 1201 | 0 | 0 | 1 | 1201 | 508716 | 507516 | 0.0 | 2218 |
| Megalops_cyprinoides-NC_050584.1 | 84.737 | 224 | 0 | 0 | 42 | 265 | 69819837 | 69820061 | 2.22e-41 | 1642 |
| Centroberyx_gerrardi-JAPMTB010008026.1 | 91.597 | 119 | 10 | 0 | 1 | 119 | 2073 | 1955 | 4.41e-36 | 165 |
| Centroberyx_gerrardi-JAPMTB010007340.1 | 93.636 | 110 | 7 | 0 | 1 | 110 | 62288 | 62397 | 4.41e-36 | 165 |
| Centroberyx_gerrardi-JAPMTB010002041.1 | 88.889 | 135 | 14 | 1 | 1 | 134 | 158600 | 158466 | 4.41e-36 | 165 |
| Centroberyx_gerrardi-JAPMTB010000749.1 | 93.636 | 110 | 7 | 0 | 1 | 110 | 11941 | 11832 | 4.41e-36 | 165 |
| Centroberyx_gerrardi-JAPMTB010007530.1 | 90.833 | 120 | 10 | 1 | 1 | 119 | 20251 | 20132 | 2.05e-34 | 159 |
| Centroberyx_gerrardi-JAPMTB010004607.1 | 88.148 | 135 | 15 | 1 | 1 | 134 | 450597 | 450731 | 2.05e-34 | 159 |
| Centroberyx_gerrardi-JAPMTB010004570.1 | 88.462 | 130 | 13 | 2 | 8 | 136 | 36117 | 35989 | 2.65e-33 | 156 |
| Centroberyx_gerrardi-JAPMTB010006088.1 | 89.076 | 119 | 13 | 0 | 1 | 119 | 41026 | 40908 | 4.44e-31 | 148 |
| Centroberyx_gerrardi-JAPMTB010003731.1 | 91.000 | 100 | 9 | 0 | 20 | 119 | 3691 | 3592 | 3.46e-27 | 135 |
| Centroberyx_gerrardi-JAPMTB010003670.1 | 87.611 | 113 | 13 | 1 | 1 | 112 | 290831 | 290719 | 1.61e-25 | 130 |
| Centroberyx_gerrardi-JAPMTB010005852.1 | 89.000 | 100 | 10 | 1 | 1 | 99 | 375996 | 376095 | 2.69e-23 | 122 |
| Centroberyx_gerrardi-JAPMTB010007661.1 | 90.217 | 92 | 8 | 1 | 29 | 119 | 332012 | 332103 | 3.48e-22 | 119 |
| Centroberyx_gerrardi-JAPMTB010004740.1 | 82.963 | 135 | 17 | 3 | 2 | 136 | 281596 | 281468 | 1.25e-21 | 117 |
| Centroberyx_gerrardi-JAPMTB010003279.1 | 89.362 | 94 | 9 | 1 | 1 | 93 | 286827 | 286734 | 1.25e-21 | 117 |
| Centroberyx_gerrardi-JAPMTB010000932.1 | 83.898 | 118 | 18 | 1 | 3 | 119 | 172710 | 172827 | 5.83e-20 | 111 |
| Centroberyx_gerrardi-JAPMTB010000844.1 | 83.051 | 118 | 19 | 1 | 3 | 119 | 70461 | 70344 | 2.71e-18 | 106 |

### N5.HOG0004451

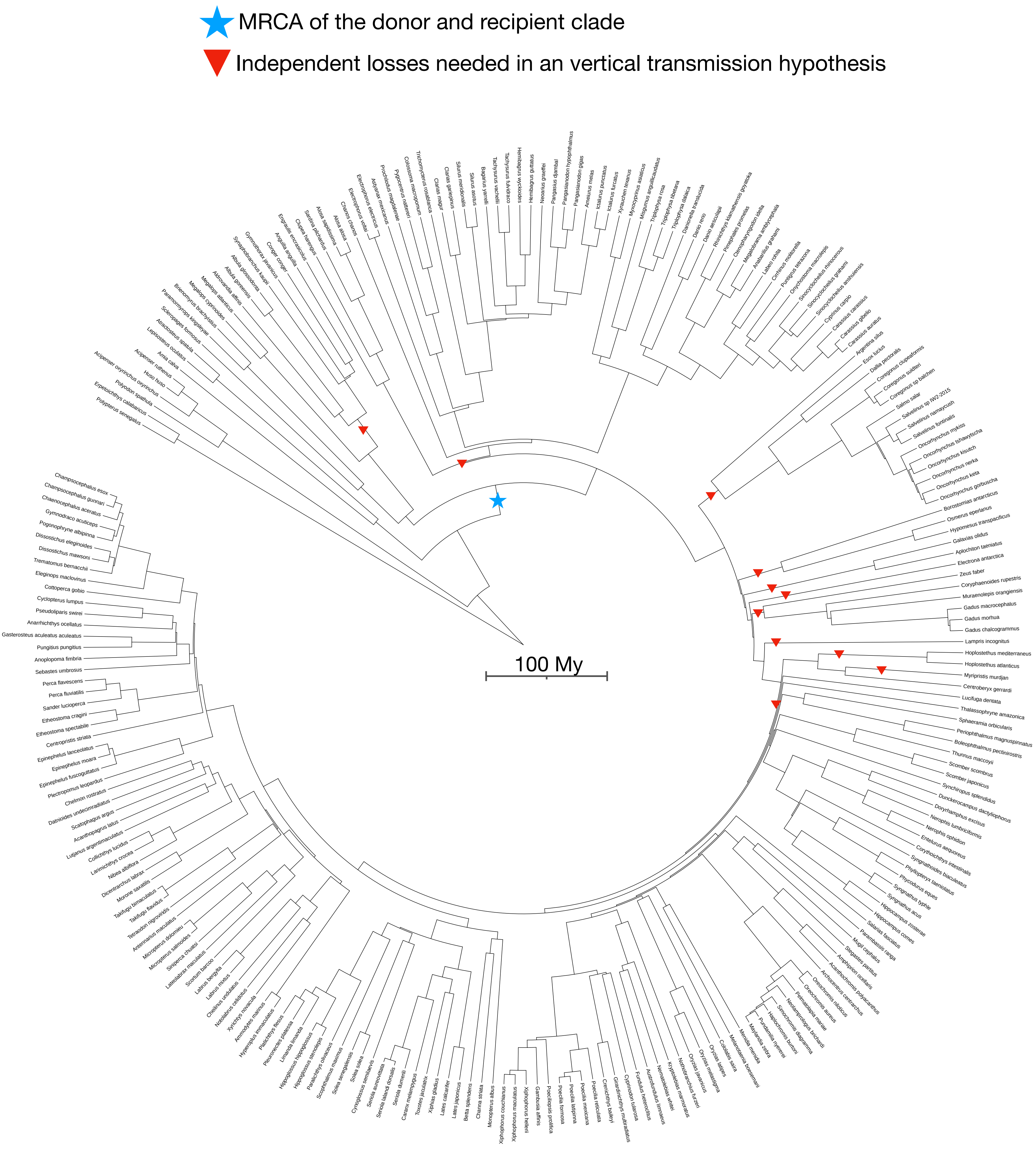

N5.HOG0030200

### N5.HOG0030200

Species tree

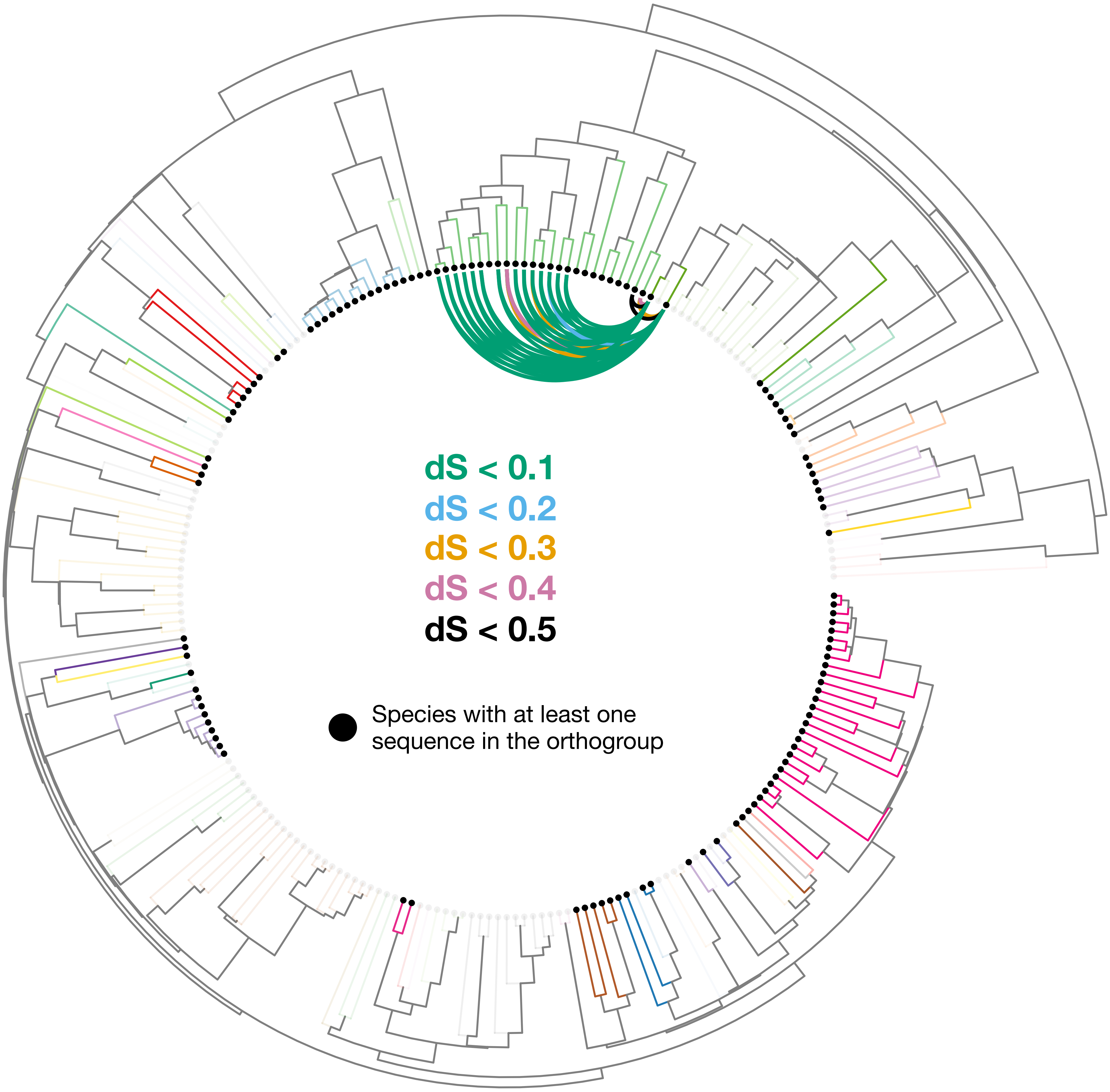

### N5.HOG0030200

Orthogroup phylogeny and protein alignment

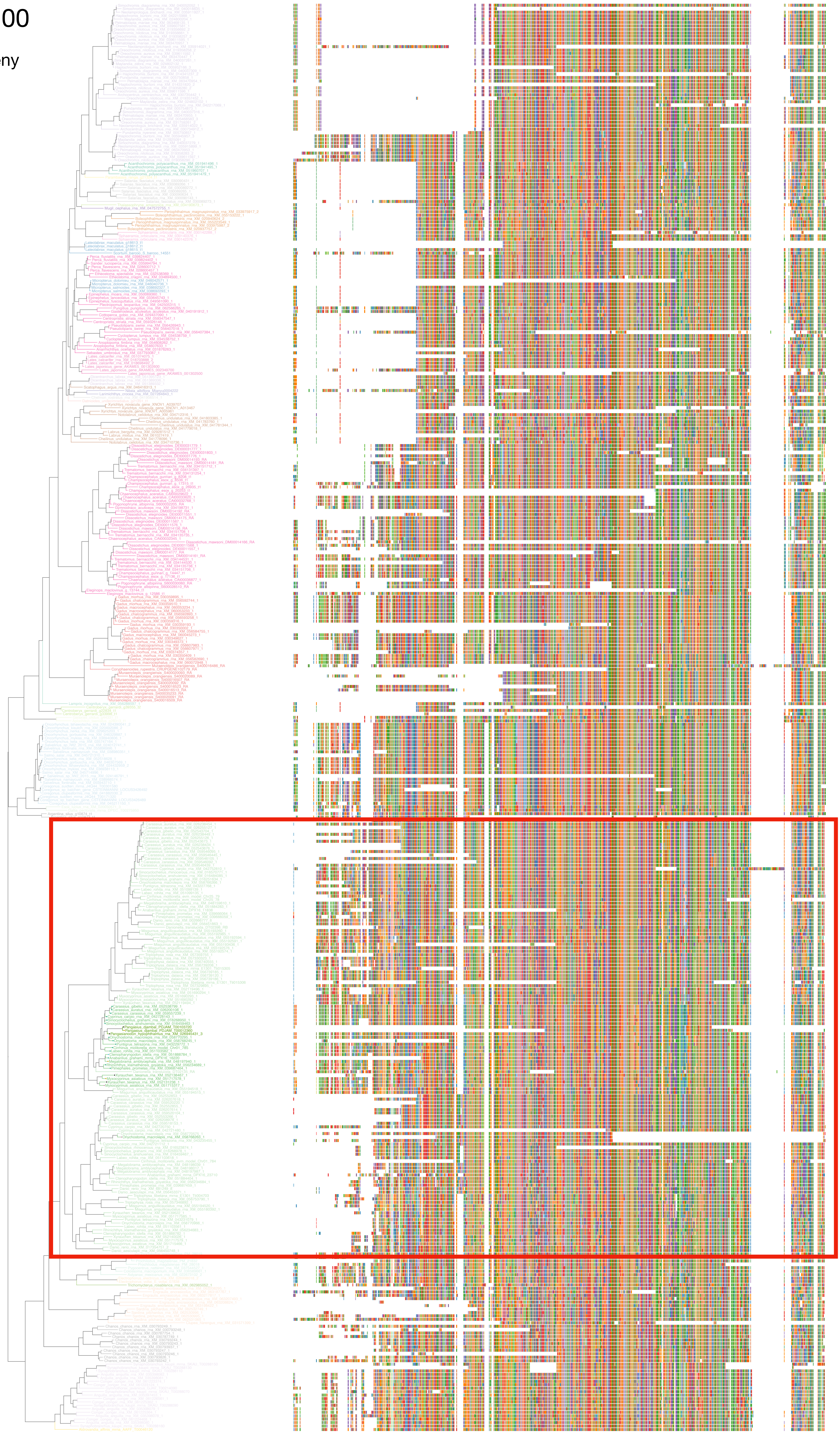

N5.HOG0030200

Zoom on HGT clade with bootstrap values

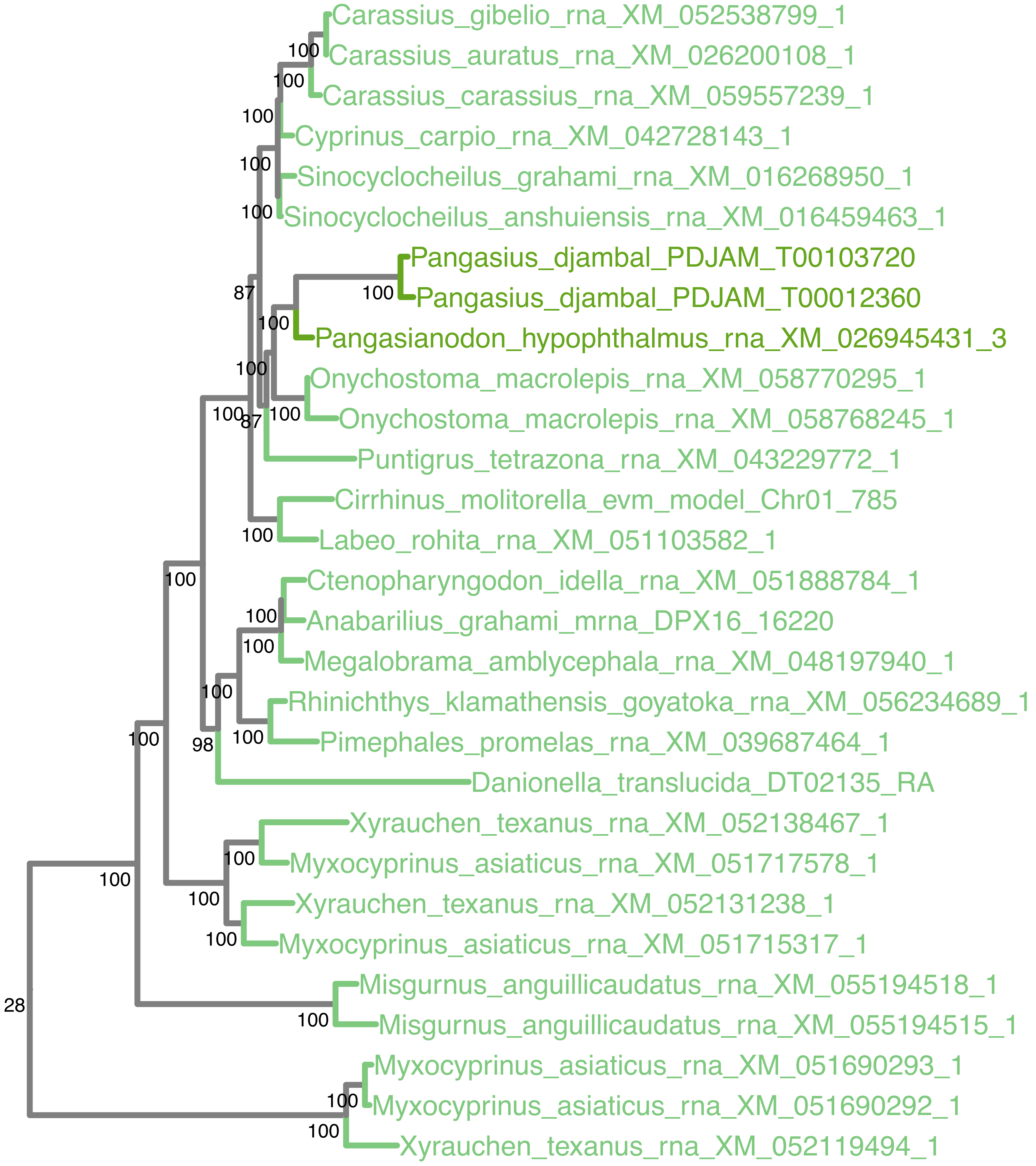

### N5.HOG0030200

Orthogroup phylogeny
and
trimmed
protein alignment

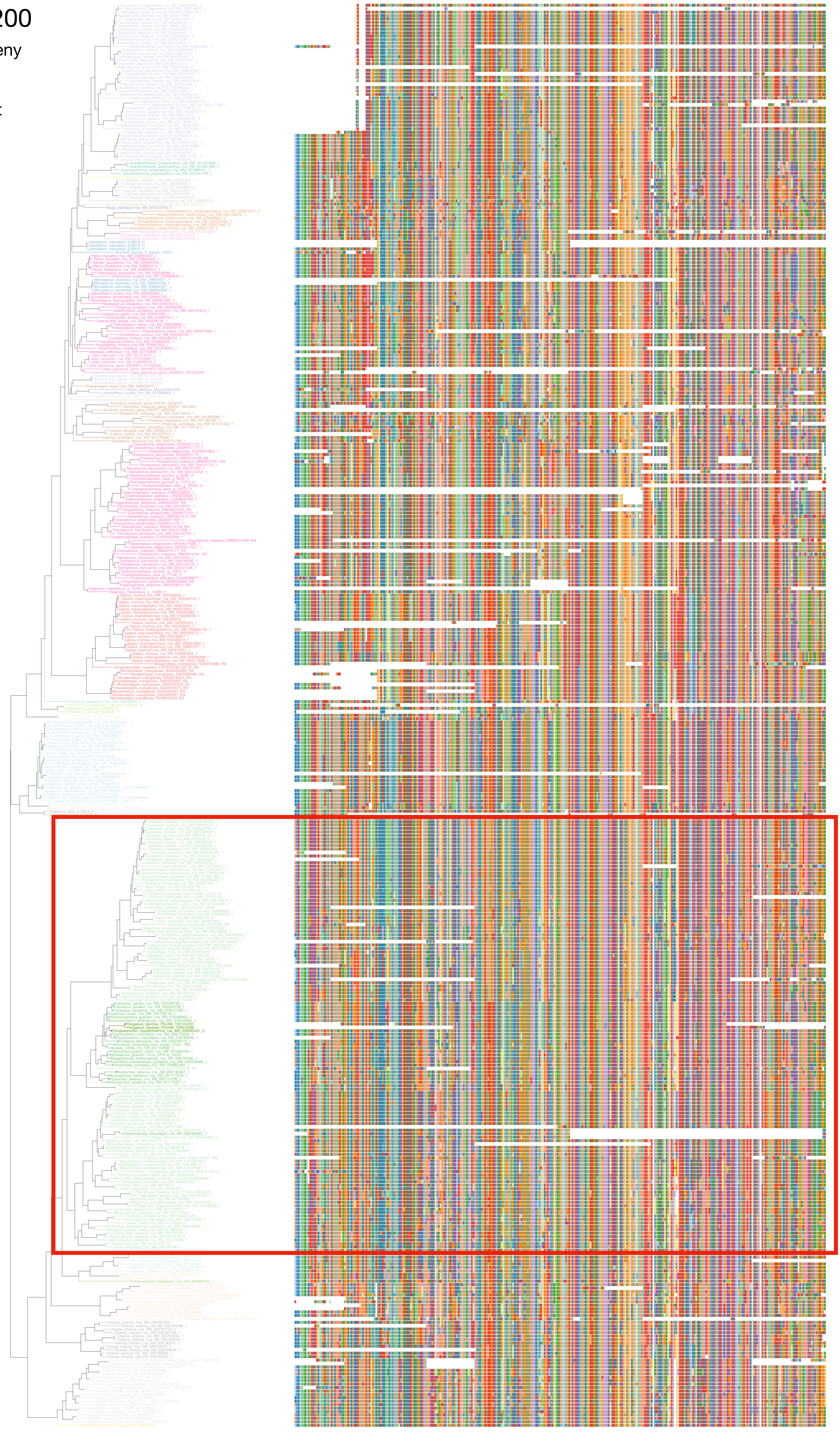

### N5.HOG0030200

« Best-match » phylogeny
and
protein alignment

50 best match against ray-finned fishes protein databases
10 best match against non-fish proteins from UniProt

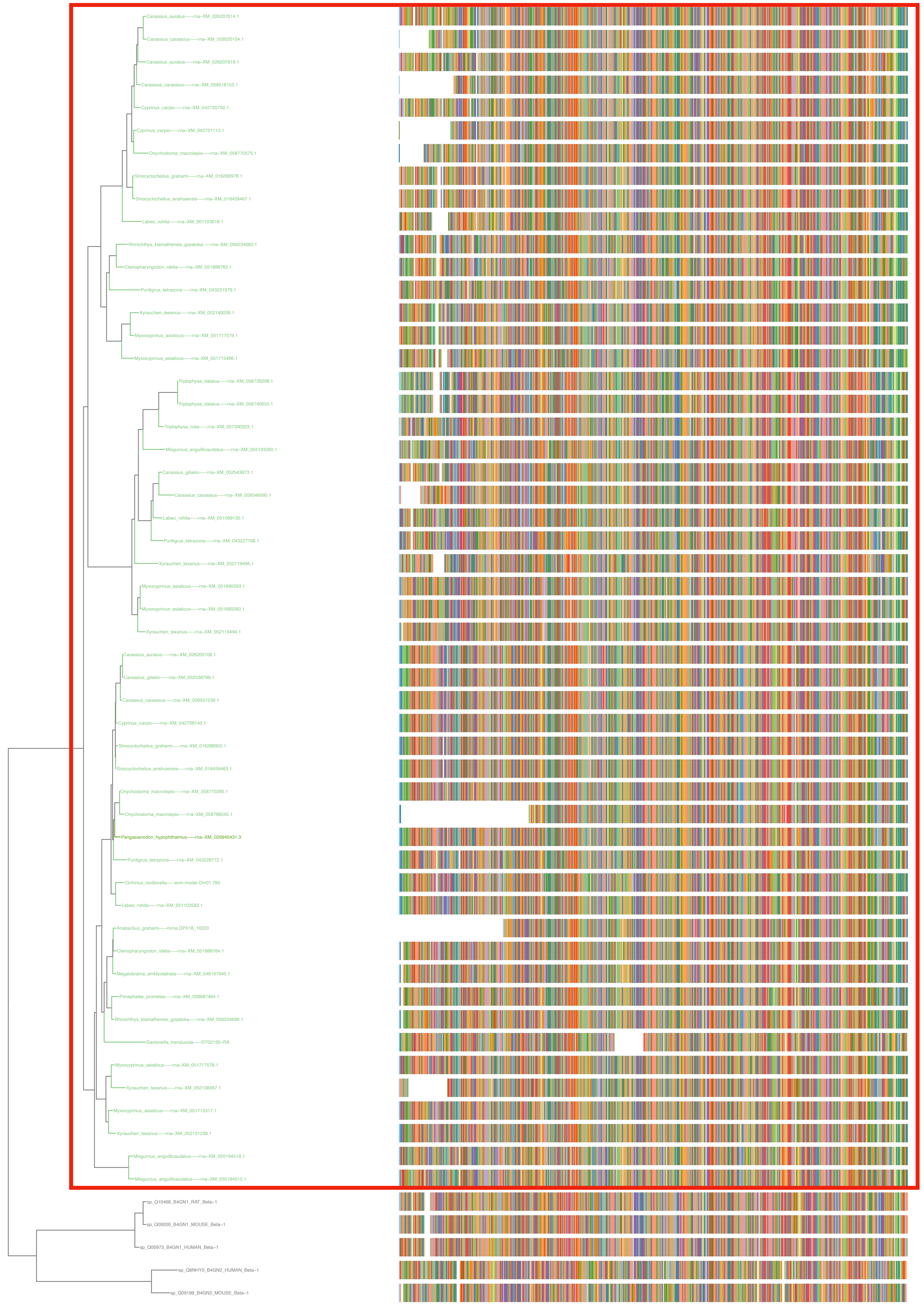

### N5.HOG0030200

#### dS distribution

##### Inter-species dS distribution

dS quantiles (0.1, 0.2, 0.3, 0.4, 0.5)
Mean dS
dS values between genes in the orthogroup N5.HOG0030200

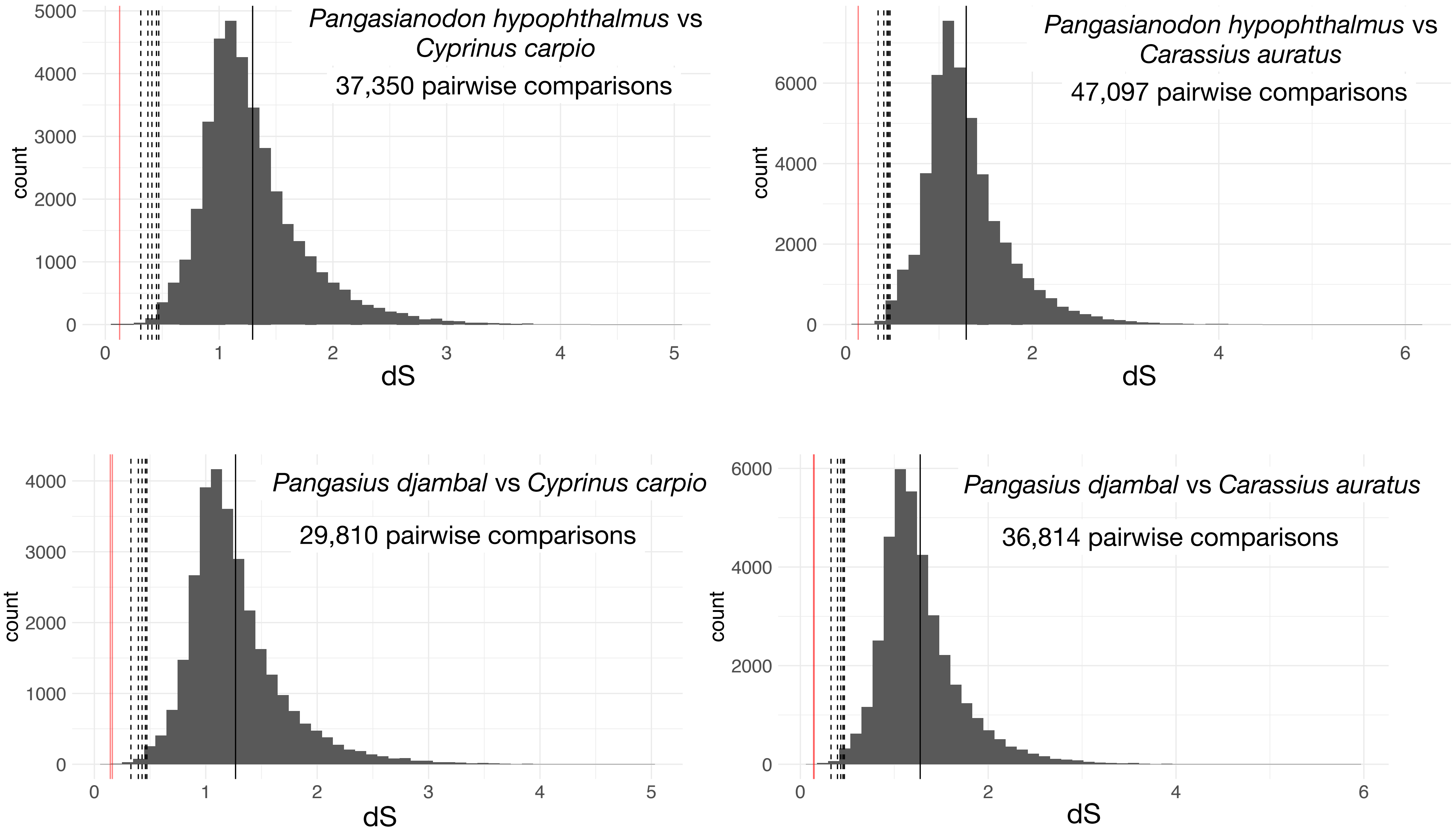


##### Intra-orthogroup dS

All pairwise comparisons
dS values between Pangasiidae and Cypriniformes genes

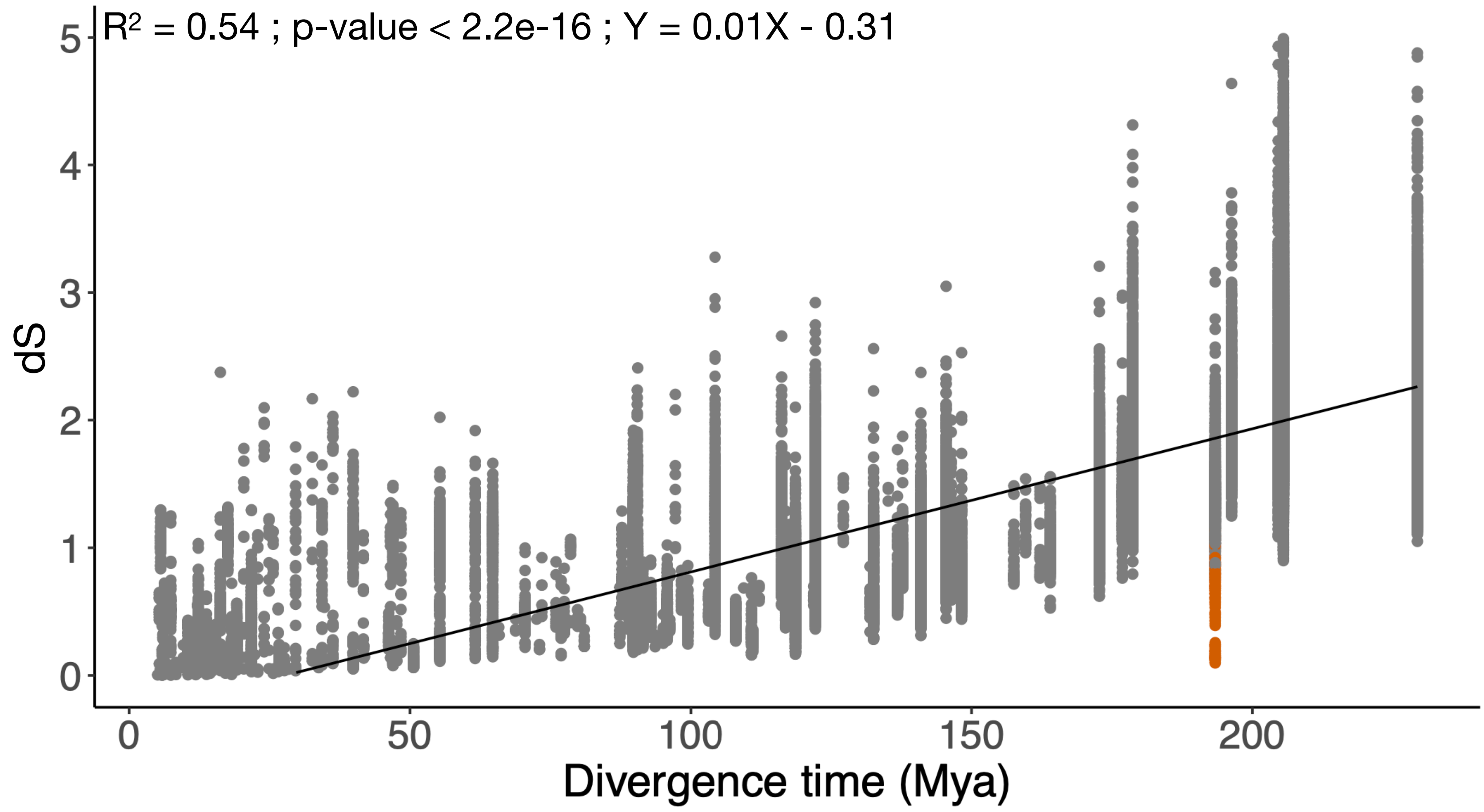

### N5.HOG0030200

#### Micro-synteny

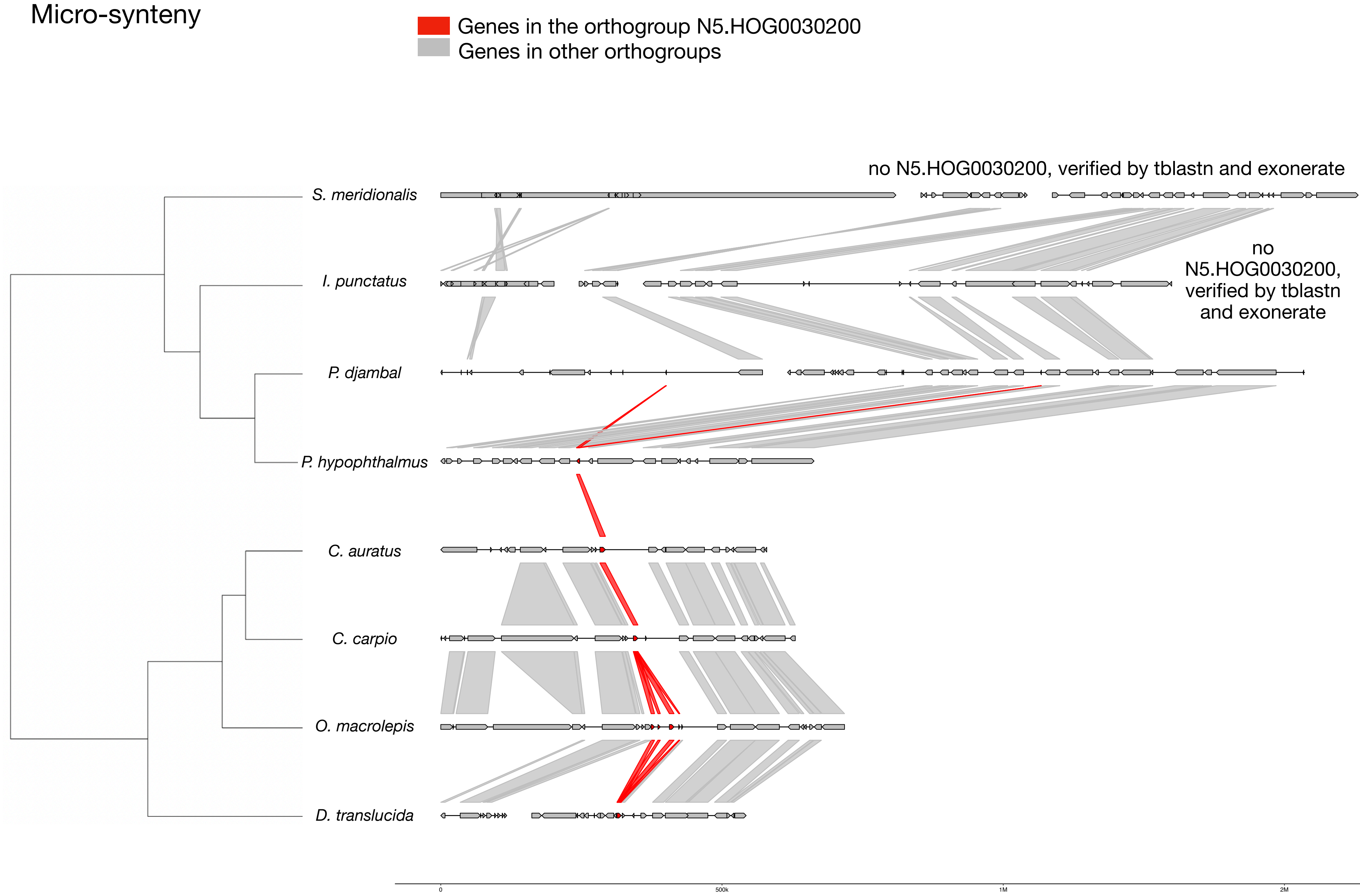


#### Zoom on *C. auratus* and *P. hypophthalmus*

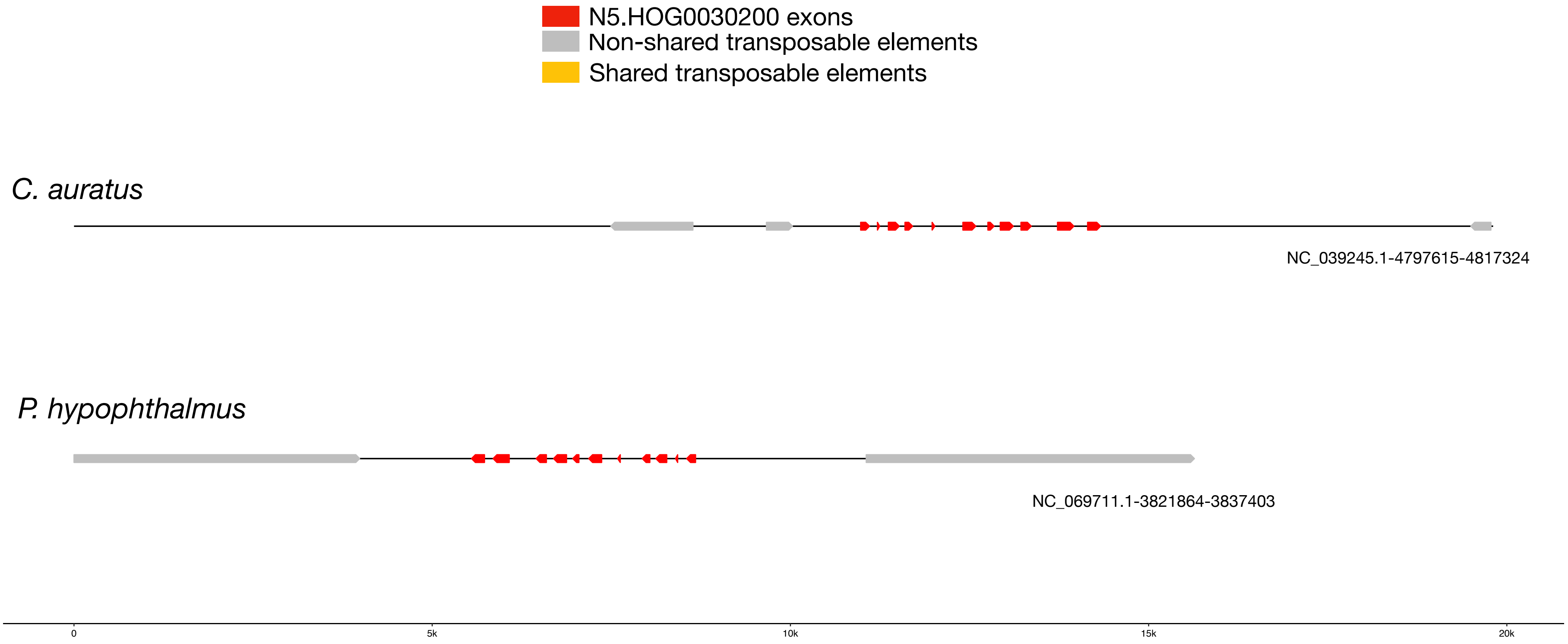

### N5.HOG0030200

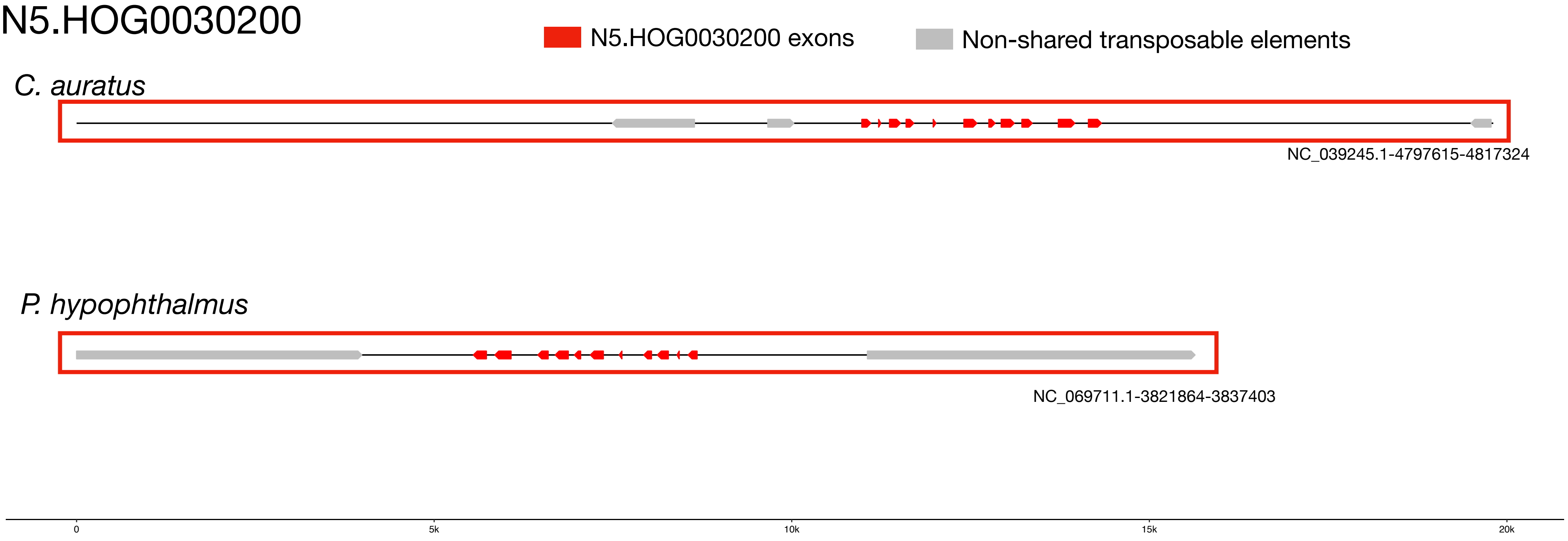


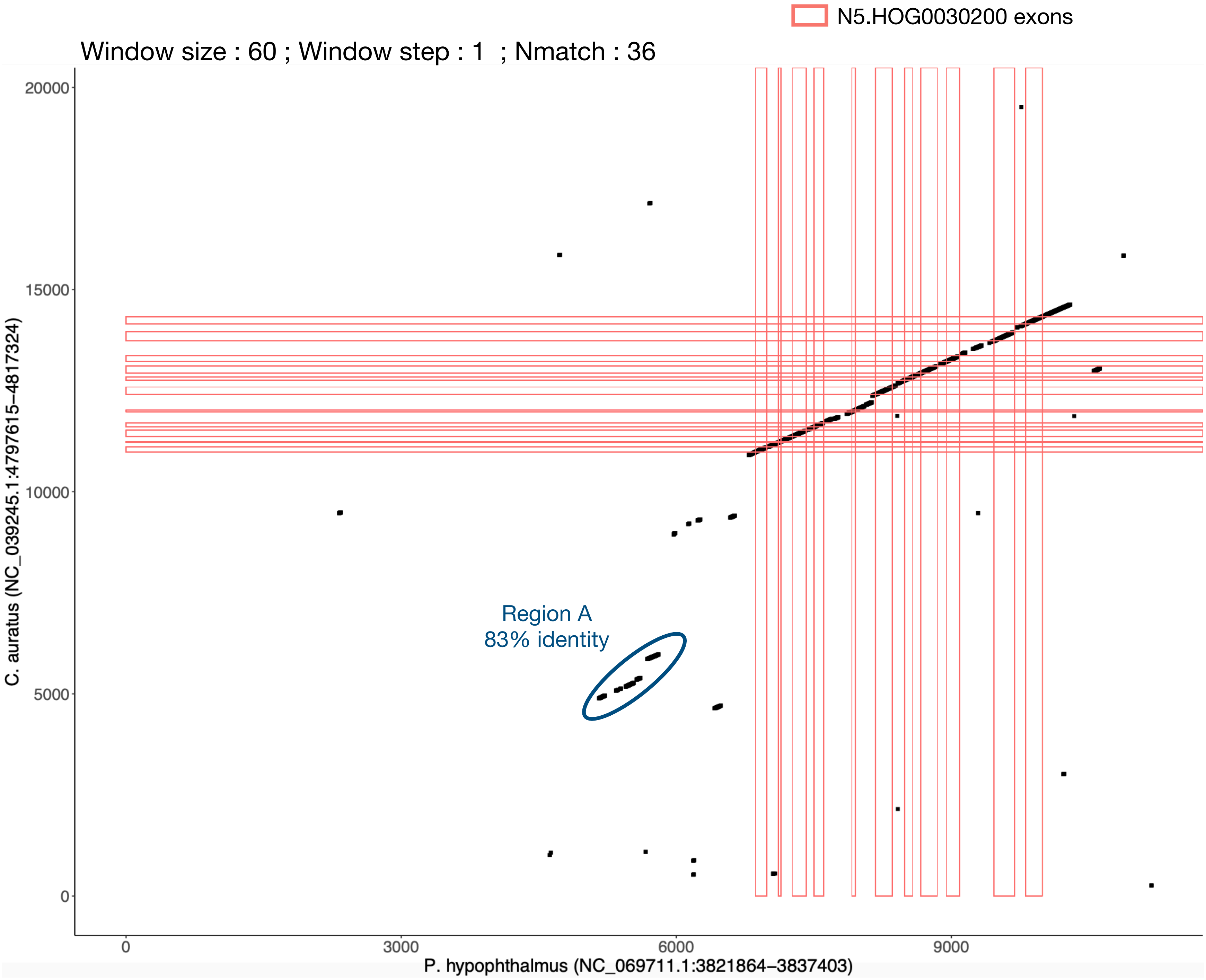

### N5.HOG0030200 Blastn of conserved regions against all genome assemblies

#### Region A

| sseqid | pident | length | mismatch | gapopen | qstart | qend | sstart | send | evalue | bitscore |
|---|---|---|---|---|---|---|---|---|---|---|
| Pangasianodon_hypophthalmus-NC_069711.1 | 100.000 | 801 | 0 | 0 | 1 | 801 | 3832304 | 3831504 | 0.0 | 1480 |
| Pangasius_djambal-CM040976.1 | 83.807 | 352 | 23 | 10 | 20 | 368 | 3497123 | 3496803 | 5.89e-78 | 303 |
| Pangasius_djambal-CM040976.1 | 95.139 | 144 | 7 | 0 | 591 | 734 | 3496779 | 3496636 | 3.65e-55 | 228 |
| Onychostoma_macrolepis-NC_081157.1 | 86.607 | 112 | 11 | 1 | 26 | 137 | 42941240 | 42941347 | 6.36e-23 | 121 |

### N5.HOG0030200

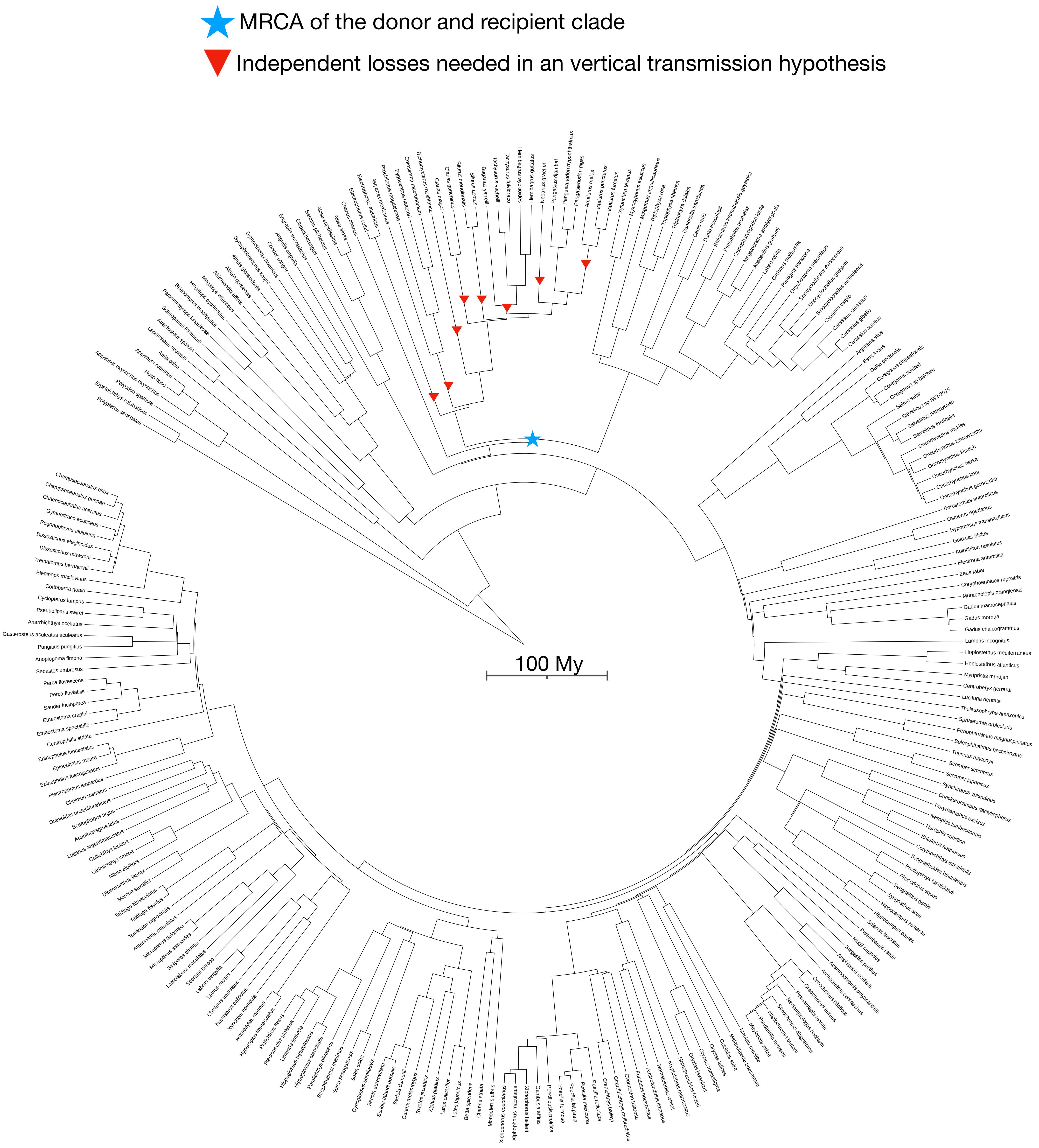

N5.HOG0034670

### N5.HOG0034670

Species tree

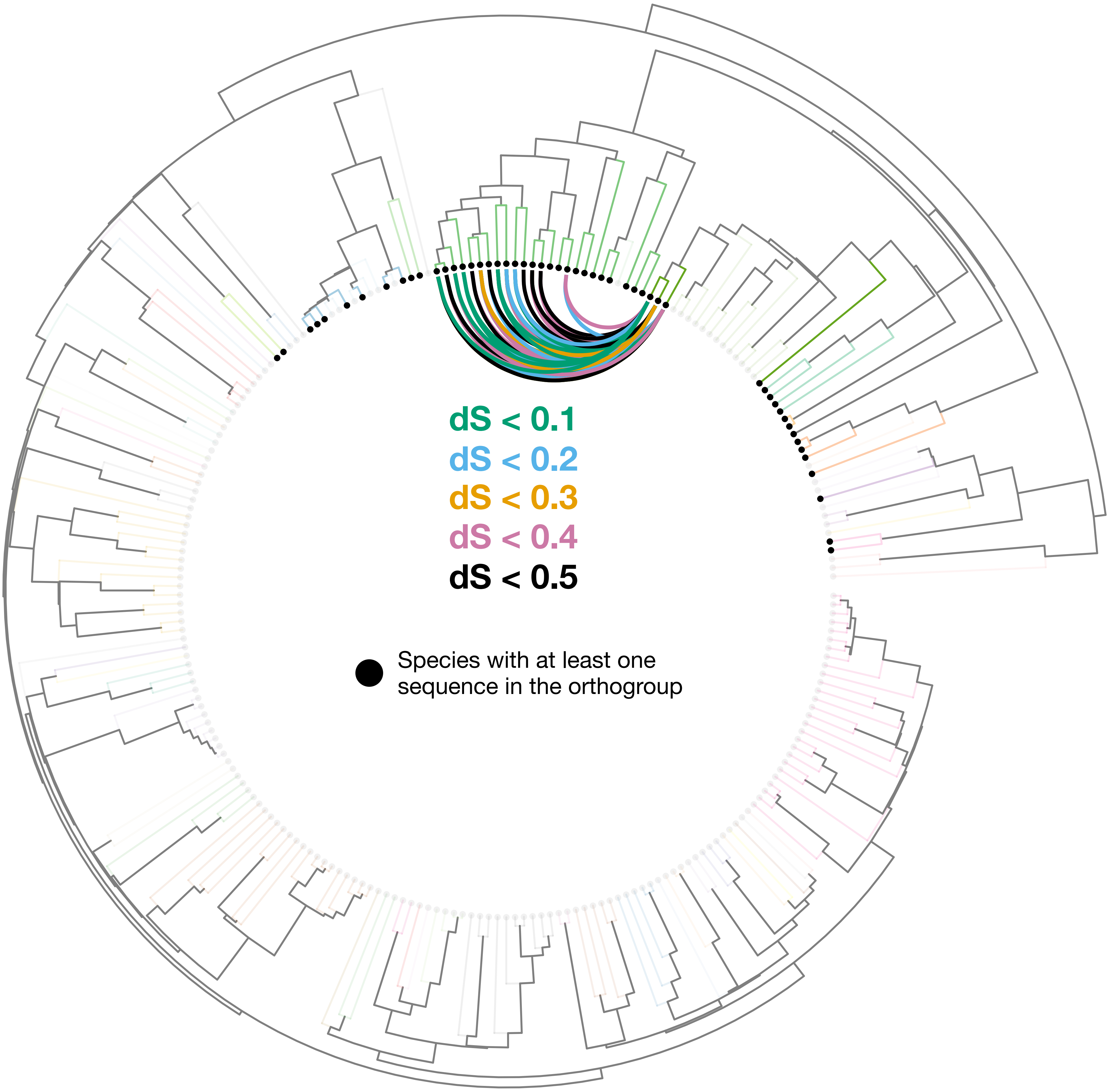

### N5.HOG0034670

Orthogroup phylogeny and protein alignment

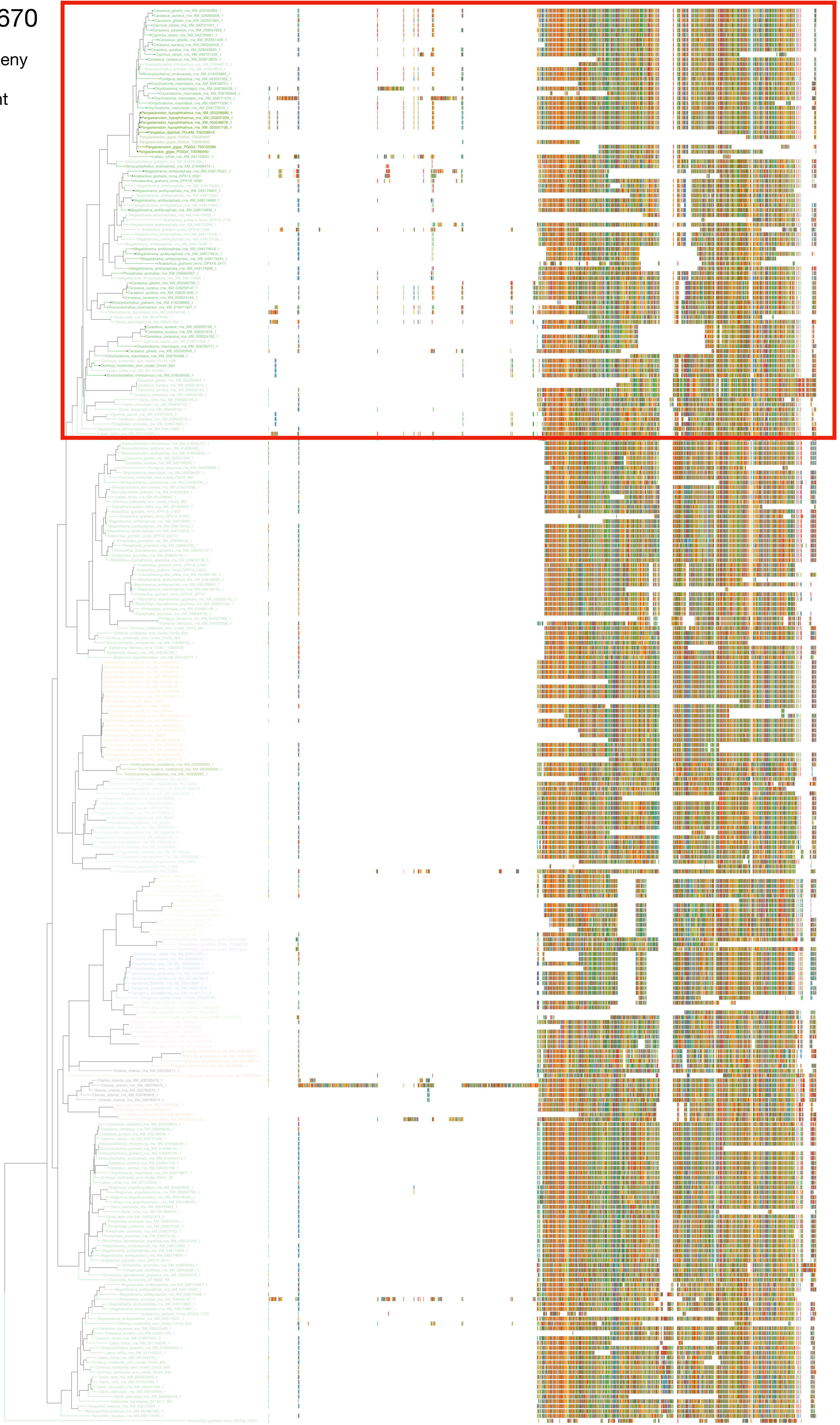

### N5.HOG0034670

Zoom on HGT clade with bootstrap values

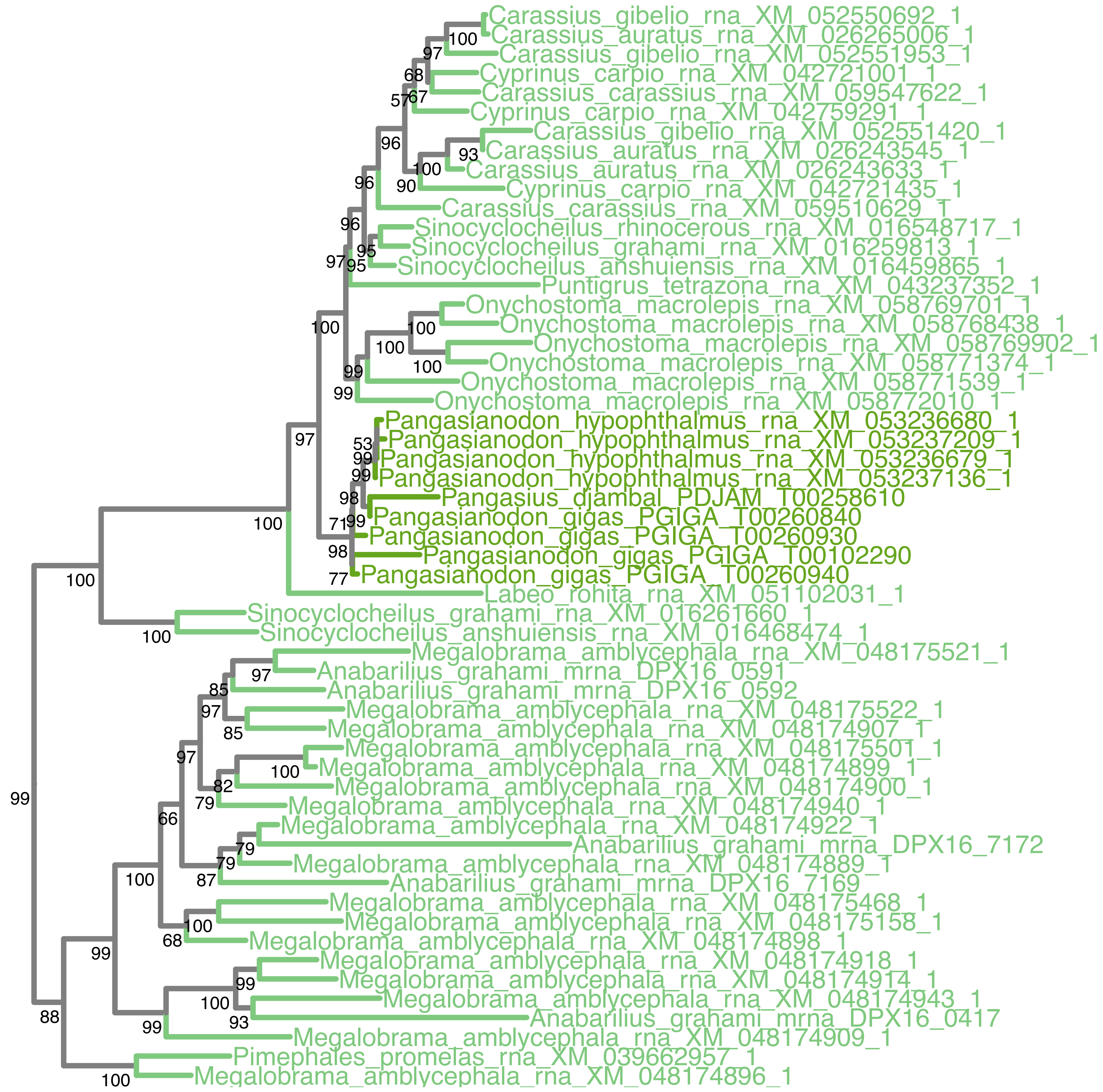

### N5.HOG0034670

Orthogroup phylogeny
and
trimmed
protein alignment

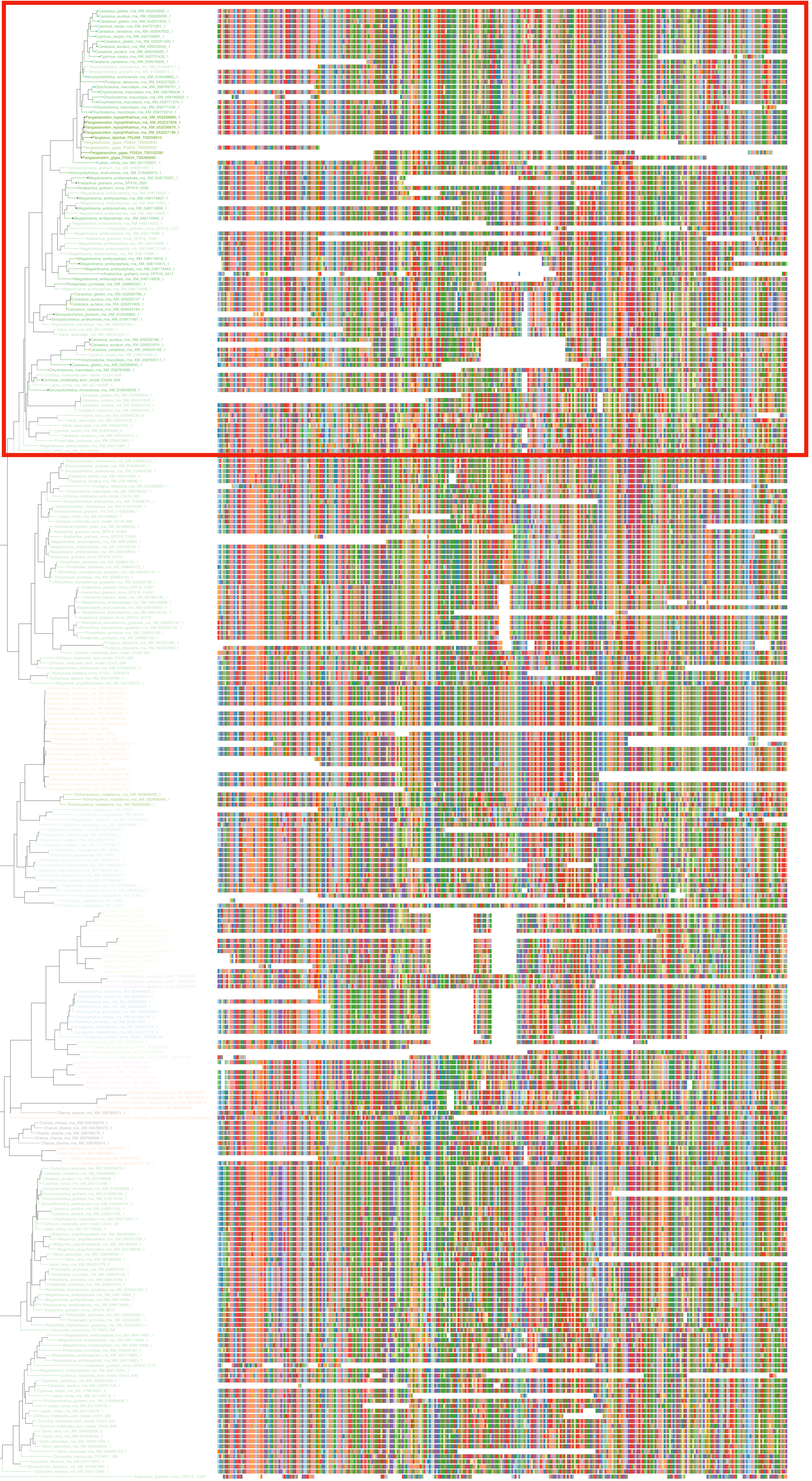

### N5.HOG0034670

« Best-match » phylogeny
and
protein alignment

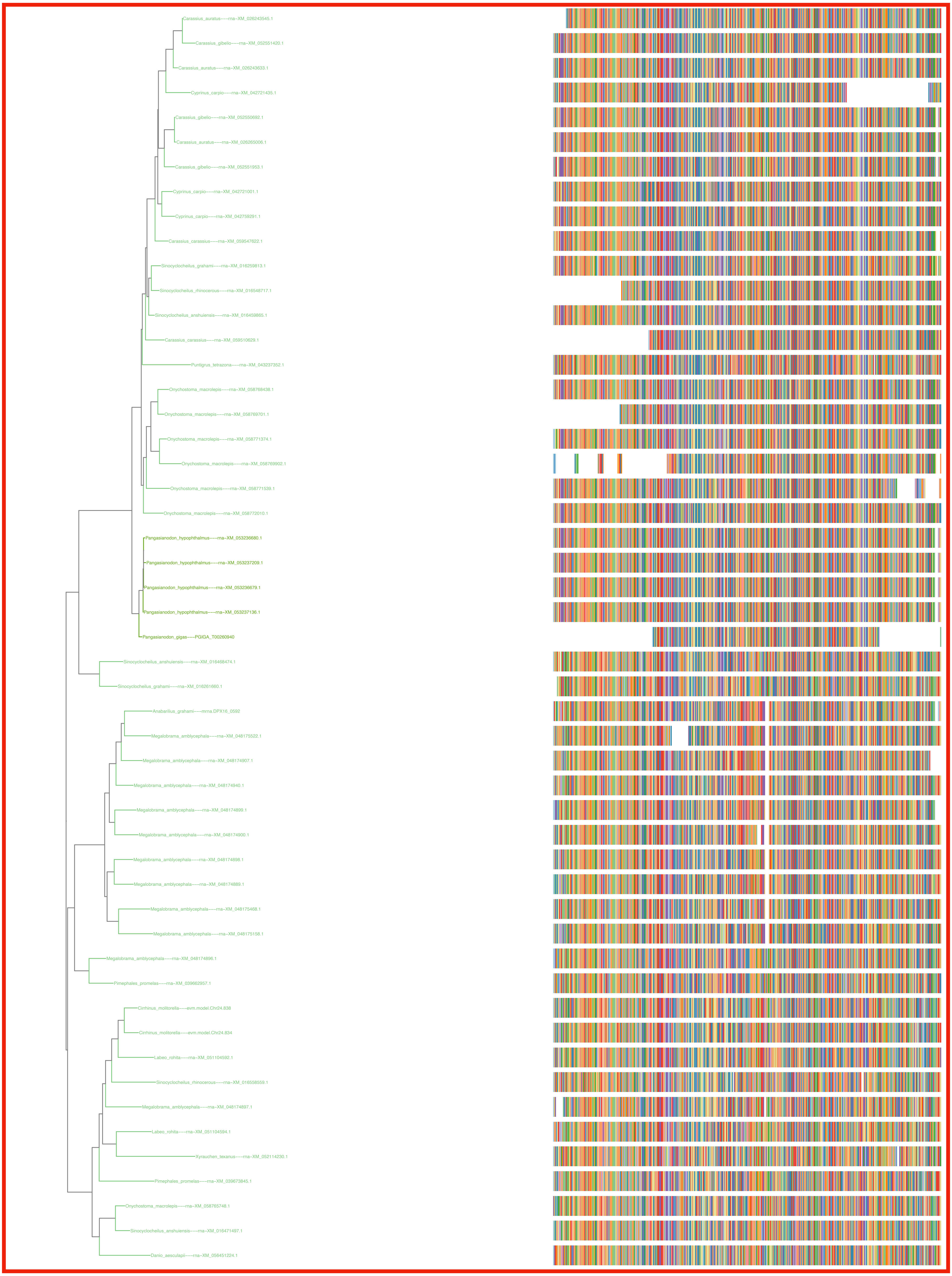

### N5.HOG0034670
dS distribution

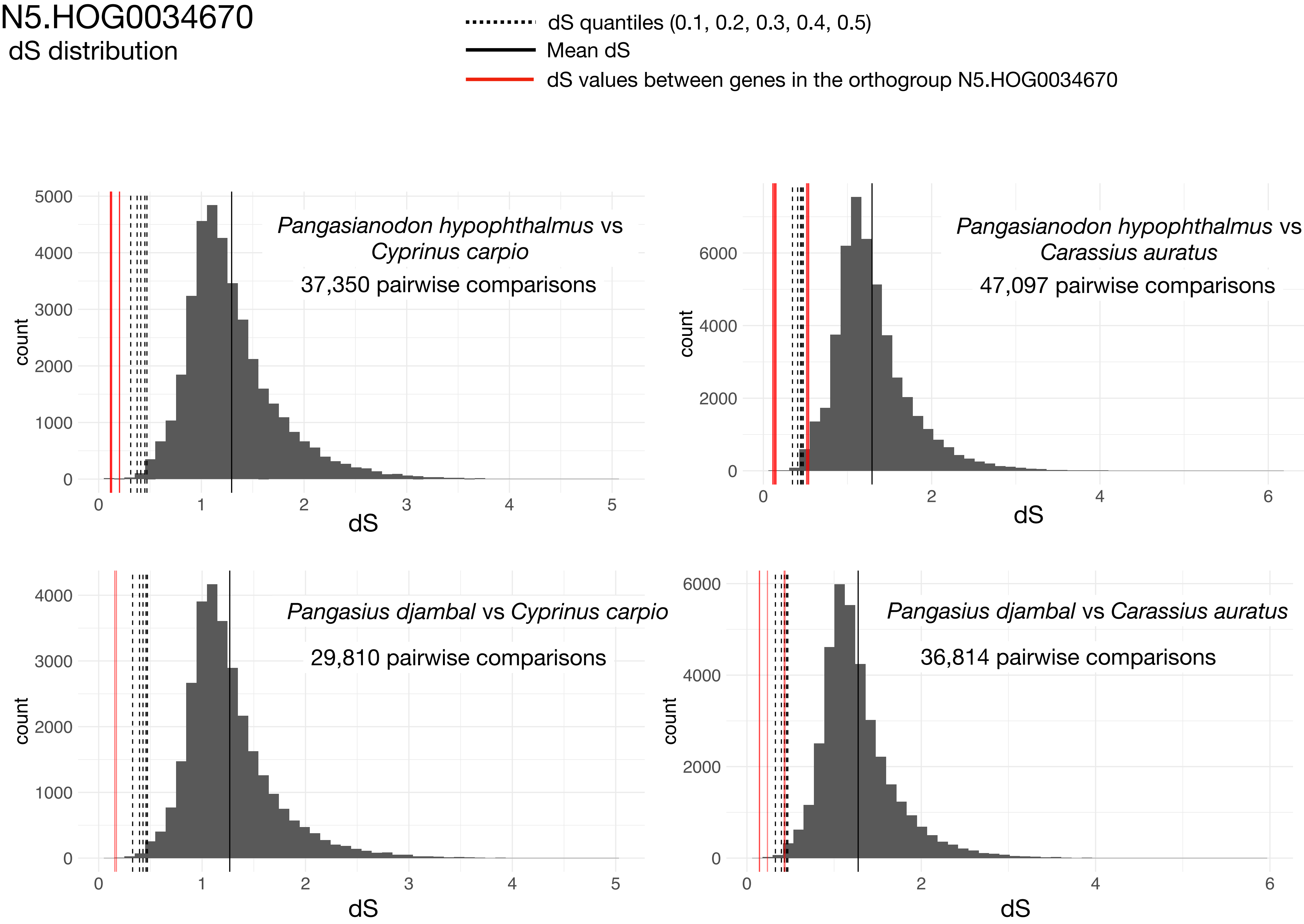


Intra-orthogroup dS

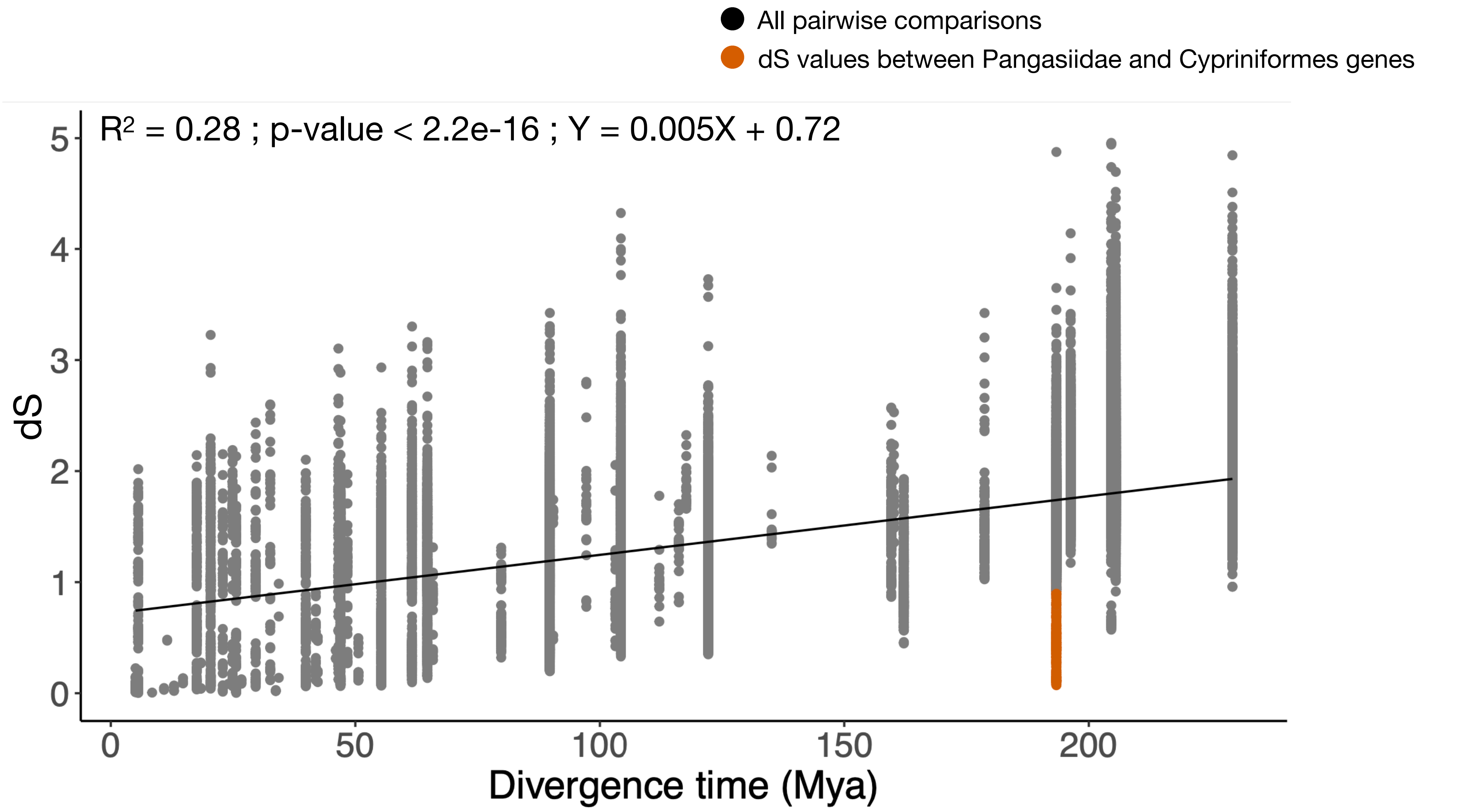

### N5.HOG0034670

#### Micro-synteny

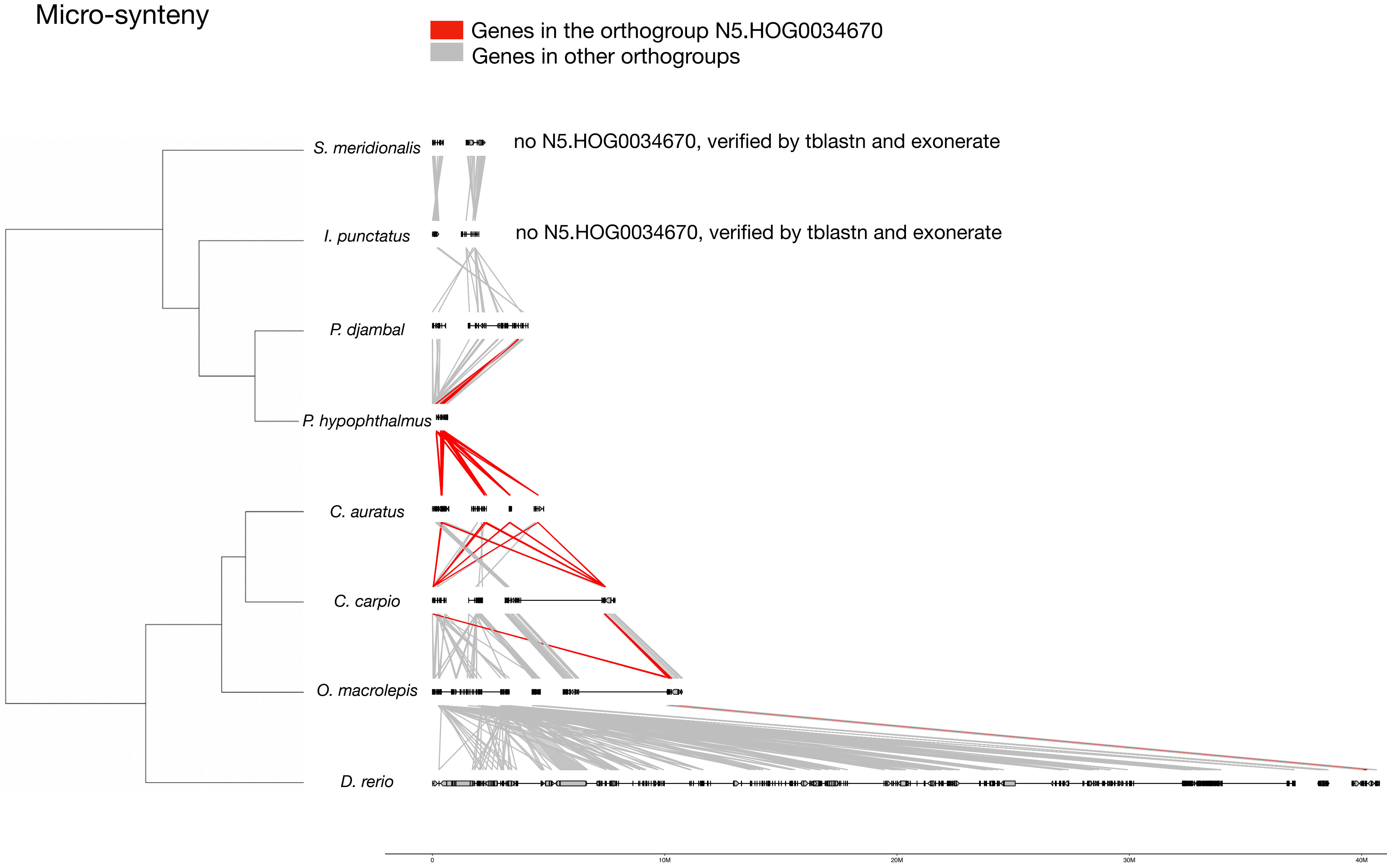


#### Zoom on *C. auratus* and *P. hypophthalmus*

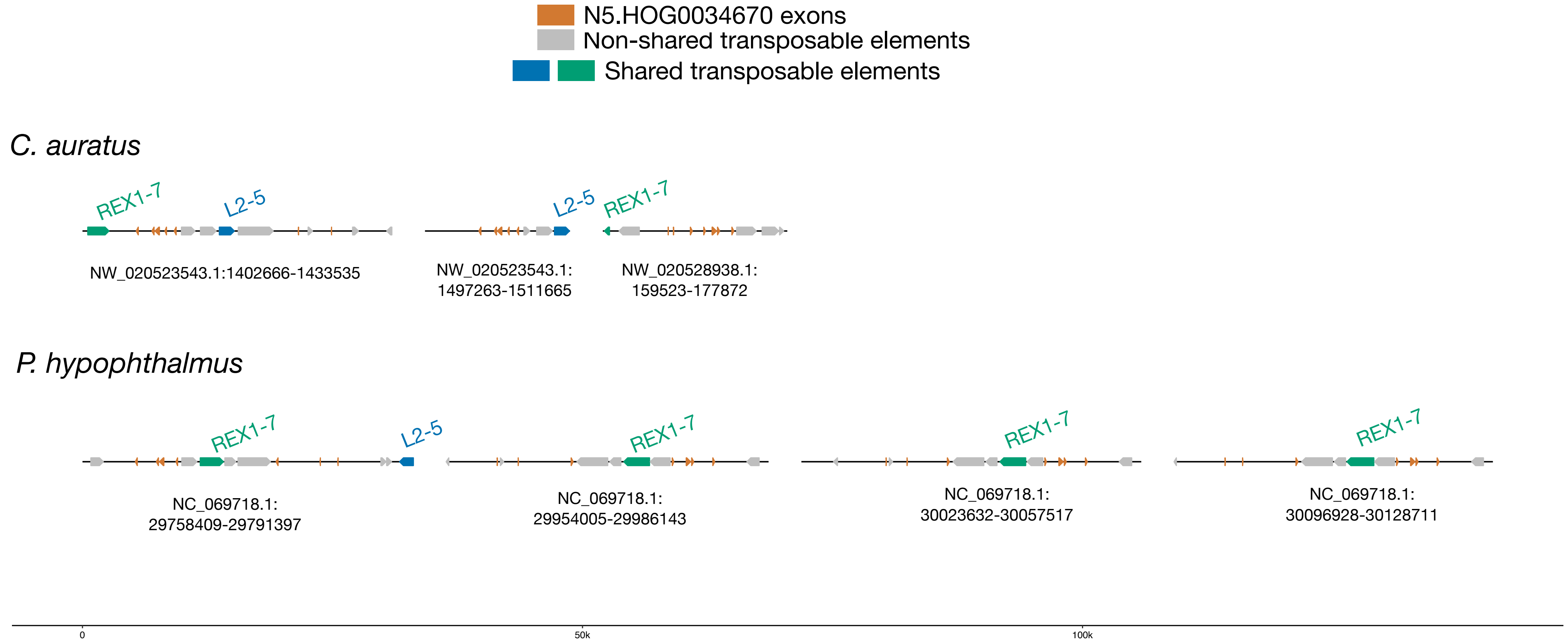


Both shared TEs are in many copies in all the genomes above

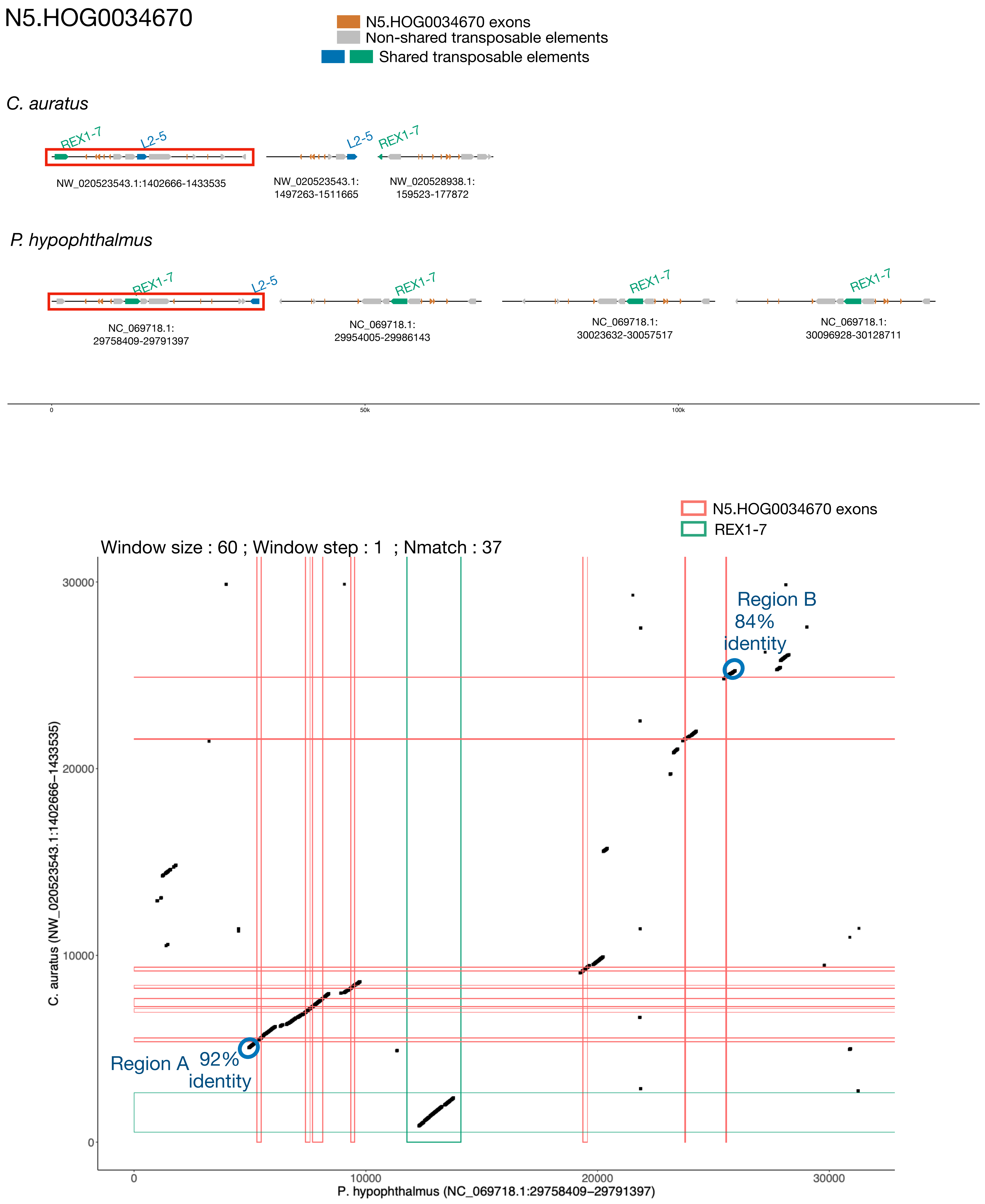
N5.HOG0034670
N5.HOG0034670 exons
Non-shared transposable elements
Shared transposable elements
C. auratus
REX1-7
L2-5
NW_020523543.1:1402666-1433535
NW_020523543.1:
1497263-1511665
NW_020528938.1:
159523-177872
P. hypophthalmus
NC_069718.1:
29758409-29791397
NC_069718.1:
29954005-29986143
NC_069718.1:
30023632-30057517
NC_069718.1:
30096928-30128711
0
50k
100k
REX1-7
Window size : 60 ; Window step : 1 ; Nmatch : 37
Region A 92% identity
Region B 84% identity
C. auratus (NW_020523543.1:1402666−1433535)
P. hypophthalmus (NC_069718.1:29758409−29791397)
0
10000
20000
30000

### N5.HOG0034670 Blastn of conserved regions against all genome assemblies

#### Region A

| sseqid | pident | length | mismatch | gapopen | qstart | qend | sstart | send | evalue | bitscore |
|---|---|---|---|---|---|---|---|---|---|---|
| Pangasianodon_hypophthalmus-[illegible] | 100.000 | 295 | 0 | 0 | 1 | 295 | 29763408 | 29763702 | 3.02e-151 | 545 |
| Pangasianodon_hypophthalmus-[illegible] | 100.000 | 295 | 0 | 0 | 1 | 295 | 30052518 | 30052224 | 3.02e-151 | 545 |
| Pangasianodon_hypophthalmus-[illegible] | 99.661 | 295 | 1 | 0 | 1 | 295 | 30123717 | 30123423 | 1.40e-149 | 540 |
| Pangasianodon_hypophthalmus-[illegible] | 99.322 | 295 | 1 | 1 | 1 | 295 | 29981149 | 29980856 | 2.35e-147 | 532 |
| Pangasianodon_gigas-CM040462.1 | 97.966 | 295 | 5 | 1 | 1 | 295 | 28633531 | 28633824 | 1.10e-140 | 510 |
| Sinocyclocheilus_anshuiensis-[illegible] | 96.259 | 294 | 10 | 1 | 2 | 295 | 5312 | 5604 | 8.64e-132 | 481 |
| Pangasius_djambal-[illegible] | 95.932 | 295 | 11 | 1 | 1 | 295 | 548 | 255 | 1.12e-130 | 477 |
| Pangasius_djambal-[illegible] | 95.593 | 295 | 12 | 1 | 1 | 295 | 357 | 64 | 5.20e-129 | 472 |
| Pangasius_djambal-CM040983.1 | 95.593 | 295 | 12 | 1 | 1 | 295 | 29742576 | 29742869 | 5.20e-129 | 472 |
| Pangasius_djambal-CM040983.1 | 95.254 | 295 | 13 | 1 | 1 | 295 | 29740224 | 29739931 | 2.42e-127 | 466 |
| Pangasius_djambal-CM040983.1 | 93.515 | 293 | 9 | 3 | 2 | 294 | 29733221 | 29732939 | 1.14e-115 | 427 |
| Pangasius_djambal-CM040983.1 | 92.491 | 293 | 10 | 4 | 2 | 294 | 29726134 | 29725854 | 4.13e-110 | 409 |
| Pangasius_djambal-CM040983.1 | 90.847 | 295 | 12 | 5 | 1 | 295 | 29724337 | 29724616 | 9.01e-102 | 381 |
| Pangasius_djambal-CM040983.1 | 95.708 | 233 | 9 | 1 | 63 | 295 | 29762506 | 29762275 | 1.51e-99 | 374 |
| Pangasius_djambal-CM040983.1 | 100.000 | 36 | 0 | 0 | 1 | 36 | 29753589 | 29753554 | 2.87e-07 | 67.6 |
| Pangasius_djambal-[illegible] | 94.949 | 297 | 12 | 2 | 1 | 295 | 387 | 92 | 3.13e-126 | 462 |
| Onychostoma_macrolepis-[illegible] | 94.576 | 295 | 13 | 2 | 1 | 295 | 7710782 | 7710491 | 1.88e-123 | 453 |
| Onychostoma_macrolepis-[illegible] | 93.220 | 295 | 16 | 2 | 1 | 295 | 6651600 | 6651310 | 8.82e-117 | 431 |
| Onychostoma_macrolepis-[illegible] | 90.604 | 298 | 18 | 7 | 1 | 295 | 3091562 | 3091852 | 1.94e-103 | 387 |
| Onychostoma_macrolepis-[illegible] | 90.169 | 295 | 21 | 6 | 1 | 295 | 9722092 | 9721806 | 1.17e-100 | 377 |
| Onychostoma_macrolepis-[illegible] | 89.933 | 298 | 20 | 7 | 1 | 295 | 6897612 | 6897902 | 4.19e-100 | 375 |
| Onychostoma_macrolepis-[illegible] | 89.796 | 294 | 15 | 5 | 2 | 295 | 6317665 | 6317943 | 3.26e-96 | 363 |
| Onychostoma_macrolepis-[illegible] | 90.323 | 279 | 16 | 8 | 1 | 276 | 2517702 | 2517972 | 5.46e-94 | 355 |
| Onychostoma_macrolepis-[illegible] | 93.162 | 234 | 12 | 2 | 39 | 272 | 7264157 | 7263928 | 1.53e-89 | 340 |
| Onychostoma_macrolepis-[illegible] | 86.441 | 295 | 19 | 3 | 1 | 295 | 436439 | 436712 | 2.01e-78 | 303 |
| Onychostoma_macrolepis-[illegible] | 91.327 | 196 | 11 | 2 | 1 | 196 | 2680866 | 2681055 | 3.41e-66 | 263 |
| Pangasius_djambal-[illegible] | 95.802 | 262 | 10 | 1 | 1 | 262 | 261 | 1 | 5.31e-114 | 422 |
| Cyprinus_carpio-NC_056599.1 | 91.611 | 298 | 20 | 4 | 1 | 295 | 40710039 | 40710334 | 1.49e-109 | 407 |
| Cyprinus_carpio-NC_056599.1 | 89.933 | 298 | 25 | 4 | 1 | 295 | 41934594 | 41934889 | 3.24e-101 | 379 |
| Cyprinus_carpio-NC_056599.1 | 89.073 | 302 | 24 | 9 | 1 | 295 | 40604969 | 40605268 | 2.52e-97 | 366 |
| Cyprinus_carpio-NC_056599.1 | 86.195 | 297 | 22 | 8 | 2 | 295 | 41997505 | 41997225 | 2.01e-78 | 303 |
| Cyprinus_carpio-NC_056578.1 | 91.582 | 297 | 22 | 3 | 1 | 295 | 1343122 | 1343417 | 1.49e-109 | 407 |
| Carassius_auratus-NW_020523543.1 | 91.216 | 296 | 21 | 4 | 1 | 293 | 1407765 | 1408058 | 8.95e-107 | 398 |
| Pangasius_djambal-[illegible] | 90.909 | 297 | 10 | 5 | 1 | 295 | 438 | 157 | 2.51e-102 | 383 |
| Carassius_gibelio-NC_068398.1 | 90.714 | 280 | 24 | 2 | 16 | 295 | 46399663 | 46399386 | 5.42e-99 | 372 |
| Carassius_gibelio-NC_068398.1 | 88.814 | 295 | 30 | 2 | 1 | 295 | 45499146 | 45499437 | 4.22e-95 | 359 |
| Carassius_gibelio-NC_068398.1 | 85.127 | 316 | 23 | 7 | 1 | 293 | 43410593 | 43410279 | 7.22e-78 | 302 |
| Sinocyclocheilus_rhinocerous-[illegible] | 89.527 | 296 | 26 | 3 | 2 | 295 | 32743 | 33035 | 1.95e-98 | 370 |
| Sinocyclocheilus_rhinocerous-[illegible] | 95.238 | 42 | 2 | 0 | 152 | 193 | 32930 | 32889 | 2.87e-07 | 67.6 |
| Pangasius_djambal-[illegible] | 96.035 | 227 | 8 | 1 | 69 | 295 | 1 | 226 | 7.01e-98 | 368 |
| Pangasius_djambal-[illegible] | 95.982 | 224 | 8 | 1 | 72 | 295 | 558 | 336 | 3.26e-96 | 363 |
| Carassius_auratus-NW_020528938.1 | 88.070 | 285 | 27 | 3 | 16 | 295 | 172895 | 172613 | 9.20e-87 | 331 |
| Pangasius_djambal-[illegible] | 95.631 | 206 | 8 | 1 | 63 | 268 | 205 | 1 | 3.31e-86 | 329 |
| Labeo_rohita-NW_026128143.1 | 87.000 | 300 | 27 | 11 | 1 | 295 | 56200 | 55908 | 1.19e-85 | 327 |
| Carassius_carassius-NC_081755.1 | 86.869 | 297 | 28 | 6 | 1 | 295 | 7880896 | 7880609 | 5.54e-84 | 322 |
| Pangasius_djambal-[illegible] | 95.699 | 186 | 8 | 0 | 1 | 186 | 186 | 1 | 2.60e-77 | 300 |
| Pangasius_djambal-[illegible] | 95.699 | 186 | 8 | 0 | 1 | 186 | 109 | 294 | 2.60e-77 | 300 |
| Pangasius_djambal-[illegible] | 93.714 | 175 | 9 | 1 | 1 | 173 | 175 | 1 | 1.22e-65 | 261 |
| Pangasius_djambal-[illegible] | 93.714 | 175 | 9 | 1 | 1 | 173 | 175 | 1 | 1.22e-65 | 261 |
| Pangasius_djambal-[illegible] | 96.350 | 137 | 4 | 1 | 159 | 295 | 390 | 255 | 1.61e-54 | 224 |
| Pangasius_djambal-[illegible] | 96.350 | 137 | 4 | 1 | 159 | 295 | 399 | 264 | 1.61e-54 | 224 |
| Megalobrama_amblycephala-[illegible] | 82.772 | 267 | 27 | 13 | 1 | 264 | 2072438 | 2072688 | 2.08e-53 | 220 |
| Megalobrama_amblycephala-[illegible] | 82.022 | 267 | 29 | 12 | 1 | 264 | 2026620 | 2026870 | 4.50e-50 | 209 |
| Sinocyclocheilus_grahami-[illegible] | 93.662 | 142 | 7 | 2 | 146 | 286 | 2462 | 2602 | 1.25e-50 | 211 |
| Pangasius_djambal-[illegible] | 97.248 | 109 | 2 | 1 | 187 | 295 | 264 | 157 | 2.73e-42 | 183 |
| Pangasius_djambal-[illegible] | 96.703 | 91 | 3 | 0 | 1 | 91 | 91 | 1 | 7.69e-33 | 152 |
| Pangasius_djambal-[illegible] | 96.667 | 90 | 3 | 0 | 1 | 90 | 90 | 1 | 2.77e-32 | 150 |
| Labeo_rohita-NW_026129689.1 | 78.039 | 255 | 35 | 17 | 13 | 258 | 326852 | 327094 | 1.66e-29 | 141 |
| Sinocyclocheilus_rhinocerous-[illegible] | 96.471 | 85 | 3 | 0 | 146 | 230 | 88 | 4 | 1.66e-29 | 141 |
| Sinocyclocheilus_grahami-[illegible] | 95.000 | 60 | 2 | 1 | 195 | 254 | 770461 | 770519 | 4.73e-15 | 93.5 |
| Sinocyclocheilus_rhinocerous-[illegible] | 93.651 | 63 | 3 | 1 | 198 | 260 | 252909 | 252848 | 4.73e-15 | 93.5 |
| Pangasius_djambal-[illegible] | 98.000 | 50 | 1 | 0 | 246 | 295 | 487 | 438 | 2.20e-13 | 87.9 |
| Pangasius_djambal-[illegible] | 100.000 | 43 | 0 | 0 | 253 | 295 | 273 | 231 | 3.68e-11 | 80.5 |
| Pangasius_djambal-[illegible] | 100.000 | 36 | 0 | 0 | 1 | 36 | 38 | 3 | 2.87e-07 | 67.6 |

### N5.HOG0034670 Blastn of conserved regions against all genome assemblies

#### Region B

| sseqid | pident | length | mismatch | gapopen | qstart | qend | sstart | send | evalue | bitscore |
|---|---|---|---|---|---|---|---|---|---|---|
| Pangasianodon_hypophthalmus-NC_069718.1 | 100.000 | 301 | 0 | 0 | 1 | 301 | 29783966 | 29784266 | 1.43e-154 | 556 |
| Pangasianodon_hypophthalmus-NC_069718.1 | 100.000 | 56 | 0 | 0 | 1 | 56 | 29959110 | 29959055 | 2.23e-18 | 104 |
| Pangasianodon_hypophthalmus-NC_069718.1 | 100.000 | 56 | 0 | 0 | 1 | 56 | 30032106 | 30032051 | 2.23e-18 | 104 |
| Pangasianodon_hypophthalmus-NC_069718.1 | 100.000 | 56 | 0 | 0 | 1 | 56 | 30102032 | 30101977 | 2.23e-18 | 104 |
| Pangasianodon_gigas-JAKMWG010000680.1 | 99.336 | 301 | 2 | 0 | 1 | 301 | 466 | 766 | 3.09e-151 | 545 |
| Pangasius_djambal-CM040997.1 | 96.013 | 301 | 8 | 3 | 5 | 301 | 383488 | 383188 | 1.90e-133 | 486 |
| Pangasius_djambal-CM040983.1 | 95.724 | 304 | 8 | 3 | 1 | 299 | 30057217 | 30057520 | 6.83e-133 | 484 |
| Pangasius_djambal-CM040983.1 | 96.429 | 56 | 2 | 0 | 1 | 56 | 29883558 | 29883503 | 4.84e-15 | 93.5 |
| Pangasianodon_gigas-JAKMWG010001669.1 | 98.519 | 270 | 1 | 2 | 35 | 301 | 1 | 270 | 1.48e-129 | 473 |
| Pangasianodon_gigas-JAKMWG010000751.1 | 97.778 | 270 | 3 | 2 | 35 | 301 | 1 | 270 | 3.20e-126 | 462 |
| Pangasianodon_gigas-JAKMWG010021885.1 | 99.142 | 233 | 1 | 1 | 1 | 232 | 233 | 1 | 7.03e-113 | 418 |
| Sinocyclocheilus_rhinocerous-NW_015635605.1 | 94.043 | 235 | 11 | 2 | 2 | 233 | 124 | 358 | 2.01e-93 | 353 |
| Sinocyclocheilus_rhinocerous-NW_015623876.1 | 92.797 | 236 | 11 | 4 | 2 | 233 | 16103 | 16336 | 2.02e-88 | 337 |
| Onychostoma_macrolepis-NC_081157.1 | 89.796 | 245 | 11 | 4 | 2 | 232 | 442925 | 443169 | 7.38e-78 | 302 |
| Onychostoma_macrolepis-NC_081157.1 | 87.395 | 238 | 14 | 10 | 1 | 233 | 7215785 | 7216011 | 4.51e-65 | 259 |
| Onychostoma_macrolepis-NC_081157.1 | 86.900 | 229 | 20 | 8 | 2 | 227 | 6333846 | 6334067 | 9.75e-62 | 248 |
| Onychostoma_macrolepis-NC_081157.1 | 85.957 | 235 | 22 | 9 | 2 | 233 | 6983720 | 6983946 | 1.63e-59 | 241 |
| Onychostoma_macrolepis-NC_081157.1 | 85.408 | 233 | 16 | 4 | 2 | 233 | 3064745 | 3064530 | 4.57e-55 | 226 |
| Onychostoma_macrolepis-NC_081157.1 | 85.845 | 219 | 19 | 10 | 2 | 216 | 7242758 | 7242548 | 5.91e-54 | 222 |
| Onychostoma_macrolepis-NC_081157.1 | 90.533 | 169 | 7 | 3 | 67 | 233 | 3433363 | 3433202 | 9.89e-52 | 215 |
| Onychostoma_macrolepis-NC_081157.1 | 100.000 | 38 | 0 | 0 | 264 | 301 | 6334080 | 6334117 | 2.27e-08 | 71.3 |
| Onychostoma_macrolepis-NC_081157.1 | 100.000 | 37 | 0 | 0 | 265 | 301 | 3433194 | 3433158 | 8.15e-08 | 69.4 |
| Onychostoma_macrolepis-NC_081157.1 | 100.000 | 37 | 0 | 0 | 265 | 301 | 7216019 | 7216055 | 8.15e-08 | 69.4 |
| Sinocyclocheilus_rhinocerous-NW_015643588.1 | 92.417 | 211 | 11 | 4 | 2 | 207 | 7402 | 7612 | 3.43e-76 | 296 |
| Sinocyclocheilus_anshuiensis-NW_015537967.1 | 95.187 | 187 | 8 | 1 | 2 | 187 | 34676 | 34862 | 1.24e-75 | 294 |
| Sinocyclocheilus_grahami-NW_015507105.1 | 94.706 | 170 | 8 | 1 | 2 | 170 | 18239 | 18408 | 3.48e-66 | 263 |
| Cyprinus_carpio-NC_056599.1 | 93.296 | 179 | 9 | 3 | 4 | 181 | 41947585 | 41947761 | 1.25e-65 | 261 |
| Cyprinus_carpio-NC_056599.1 | 90.164 | 183 | 14 | 4 | 2 | 182 | 40720781 | 40720961 | 7.59e-58 | 235 |
| Cyprinus_carpio-NC_056599.1 | 88.710 | 186 | 18 | 3 | 2 | 186 | 42450232 | 42450049 | 1.64e-54 | 224 |
| Cyprinus_carpio-NC_056578.1 | 91.803 | 183 | 10 | 4 | 2 | 181 | 1352718 | 1352898 | 2.71e-62 | 250 |
| Carassius_auratus-NW_020528938.1 | 92.090 | 177 | 12 | 2 | 7 | 182 | 166040 | 165865 | 9.75e-62 | 248 |
| Carassius_gibelio-NC_068398.1 | 92.353 | 170 | 11 | 2 | 7 | 175 | 46392855 | 46392687 | 1.63e-59 | 241 |
| Labeo_rohita-NC_066872.1 | 87.195 | 164 | 10 | 8 | 2 | 158 | 18470780 | 18470939 | 4.67e-40 | 176 |
| Carassius_carassius-NC_081755.1 | 83.237 | 173 | 16 | 6 | 2 | 173 | 7854648 | 7854488 | 3.66e-31 | 147 |
| Triplophysa_rosa-NC_079905.1 | 93.258 | 89 | 4 | 1 | 215 | 301 | 8636968 | 8636880 | 3.69e-26 | 130 |
| Triplophysa_rosa-NC_079905.1 | 97.619 | 42 | 1 | 0 | 170 | 211 | 4165702 | 4165743 | 6.30e-09 | 73.1 |
| Triplophysa_rosa-NC_079892.1 | 83.221 | 149 | 8 | 11 | 170 | 301 | 9140144 | 9140292 | 2.22e-23 | 121 |
| Triplophysa_rosa-NC_079890.1 | 89.130 | 92 | 5 | 2 | 215 | 301 | 25168210 | 25168301 | 4.80e-20 | 110 |
| Triplophysa_rosa-NC_079890.1 | 83.051 | 118 | 10 | 7 | 179 | 286 | 29522847 | 29522730 | 1.04e-16 | 99.0 |
| Triplophysa_dalaica-NC_079566.1 | 88.506 | 87 | 5 | 1 | 215 | 301 | 7593569 | 7593650 | 2.89e-17 | 100 |
| Triplophysa_rosa-NC_079900.1 | 87.500 | 88 | 6 | 3 | 215 | 300 | 2426546 | 2426630 | 3.74e-16 | 97.1 |
| Triplophysa_dalaica-NC_079542.1 | 86.517 | 89 | 10 | 1 | 215 | 301 | 13972143 | 13972055 | 3.74e-16 | 97.1 |
| Triplophysa_rosa-NC_079895.1 | 86.747 | 83 | 8 | 1 | 222 | 301 | 15934989 | 15934907 | 6.26e-14 | 89.8 |
| Triplophysa_tibetana-CM017895.1 | 83.146 | 89 | 12 | 2 | 215 | 301 | 25122156 | 25122243 | 1.35e-10 | 78.7 |
| Triplophysa_tibetana-CM017895.1 | 89.831 | 59 | 4 | 2 | 230 | 286 | 28838482 | 28838424 | 1.75e-09 | 75.0 |
| Triplophysa_rosa-NC_079897.1 | 78.667 | 150 | 7 | 9 | 167 | 299 | 17972654 | 17972513 | 4.87e-10 | 76.8 |
| Sinocyclocheilus_rhinocerous-NW_015627234.1 | 78.519 | 135 | 11 | 9 | 174 | 291 | 89252 | 89385 | 6.30e-09 | 73.1 |
| Sinocyclocheilus_rhinocerous-NW_015627234.1 | 78.519 | 135 | 12 | 8 | 174 | 291 | 90631 | 90765 | 6.30e-09 | 73.1 |
| Triplophysa_rosa-NC_079893.1 | 78.322 | 143 | 7 | 10 | 167 | 292 | 22408195 | 22408330 | 2.27e-08 | 71.3 |
| Misgurnus_anguillicaudatus-NC_073338.1 | 100.000 | 37 | 0 | 0 | 216 | 252 | 54993697 | 54993661 | 8.15e-08 | 69.4 |
| Carassius_carassius-NC_081788.1 | 85.938 | 64 | 7 | 1 | 230 | 291 | 10989698 | 10989635 | 2.93e-07 | 67.6 |

### N5.HOG0034670

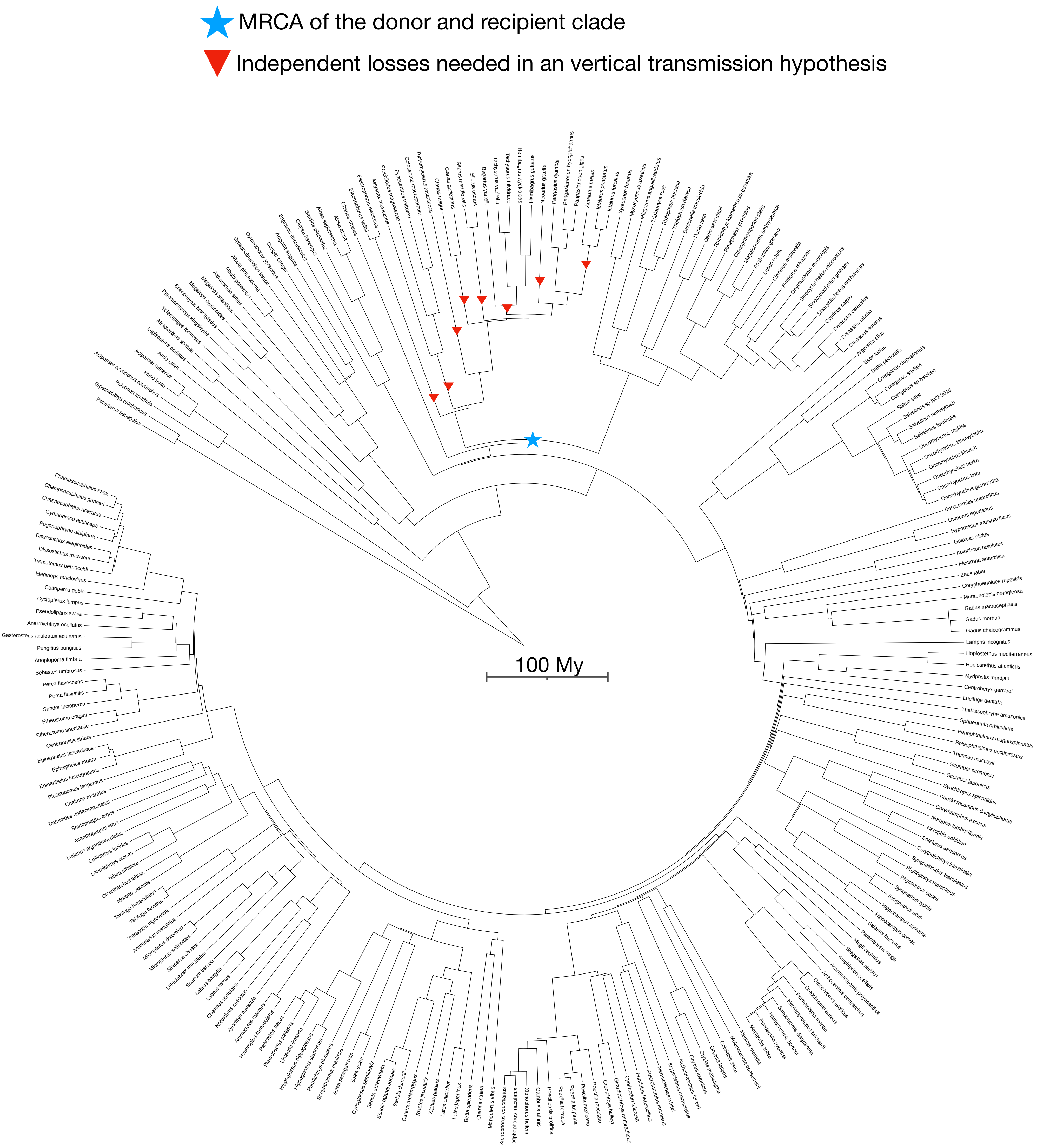

N5.HOG0019111

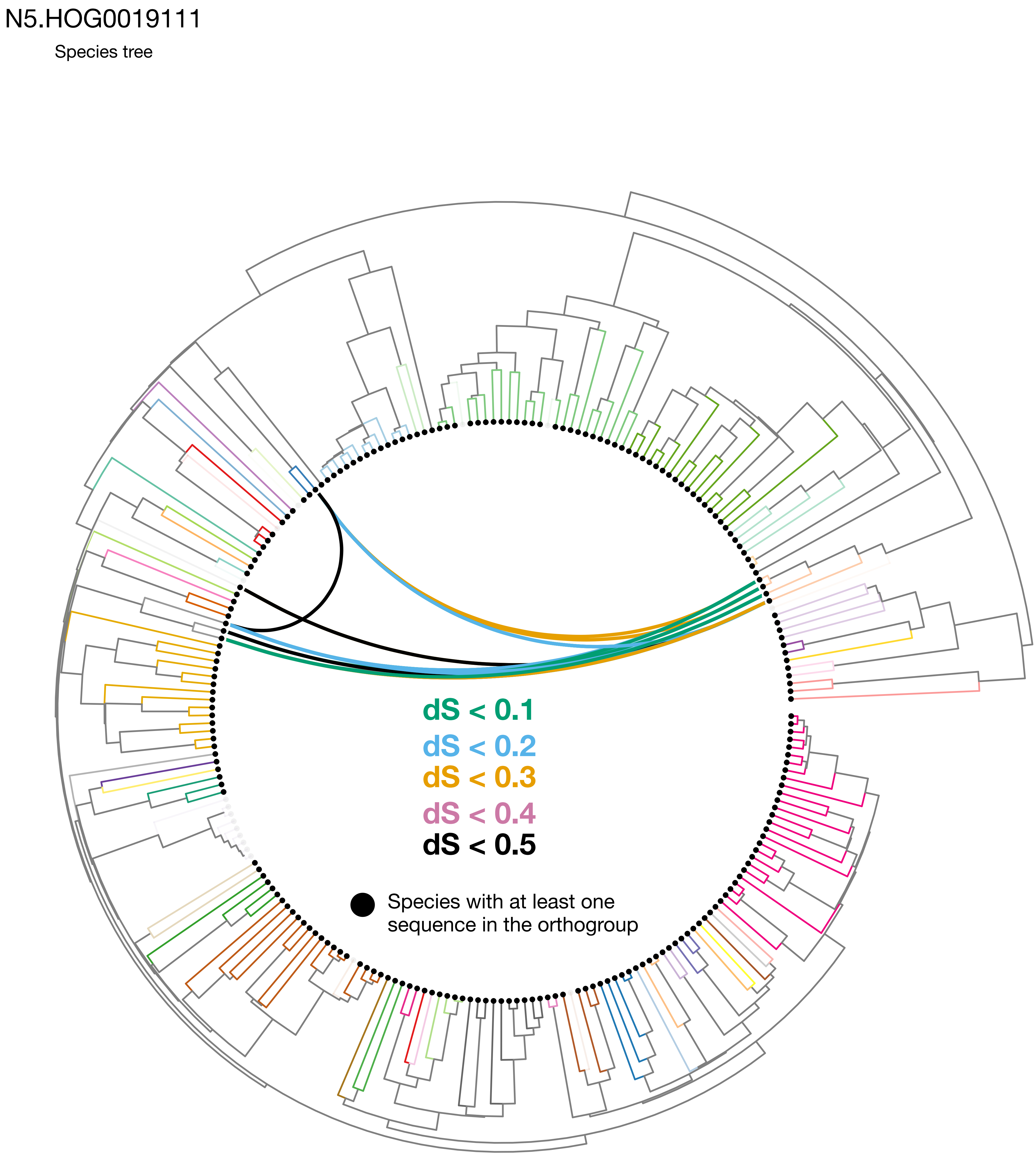


Expected pattern under a double HGT

### N5.HOG0019111

Orthogroup phylogeny
and
protein alignment

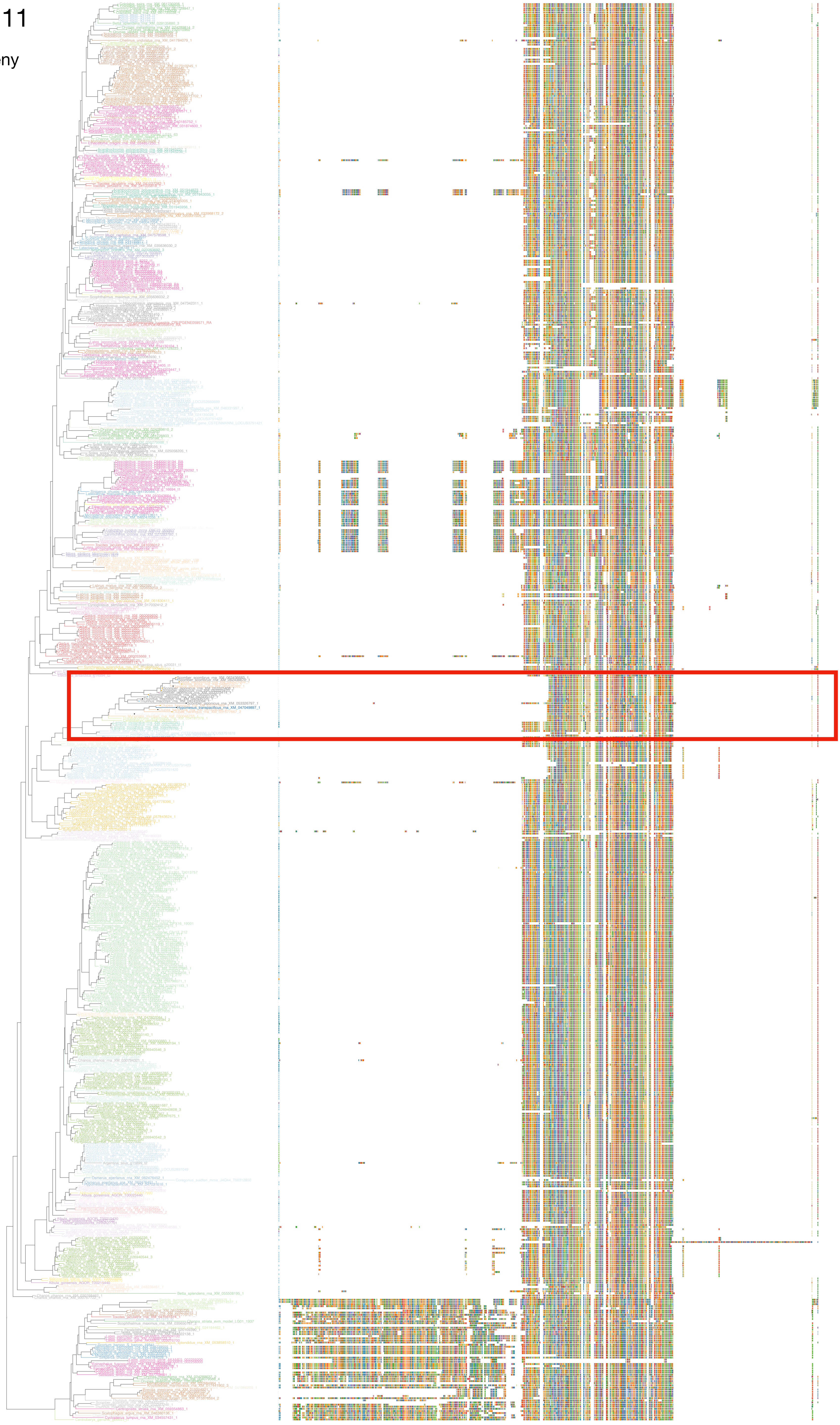

### N5.HOG0019111

Zoom on HGT clade with bootstrap values

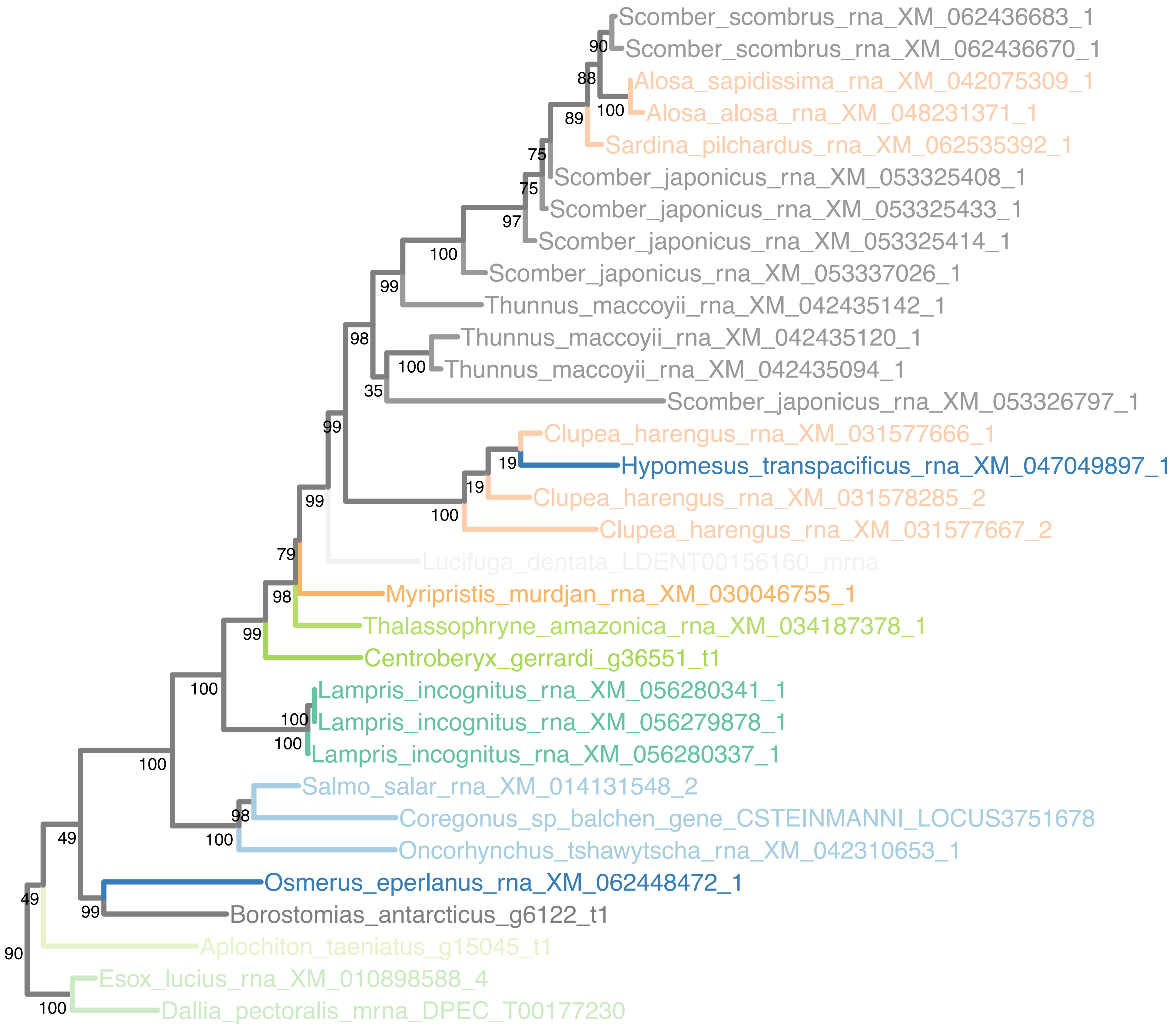

### N5.HOG0019111

Orthogroup phylogeny
and
trimmed
protein alignment

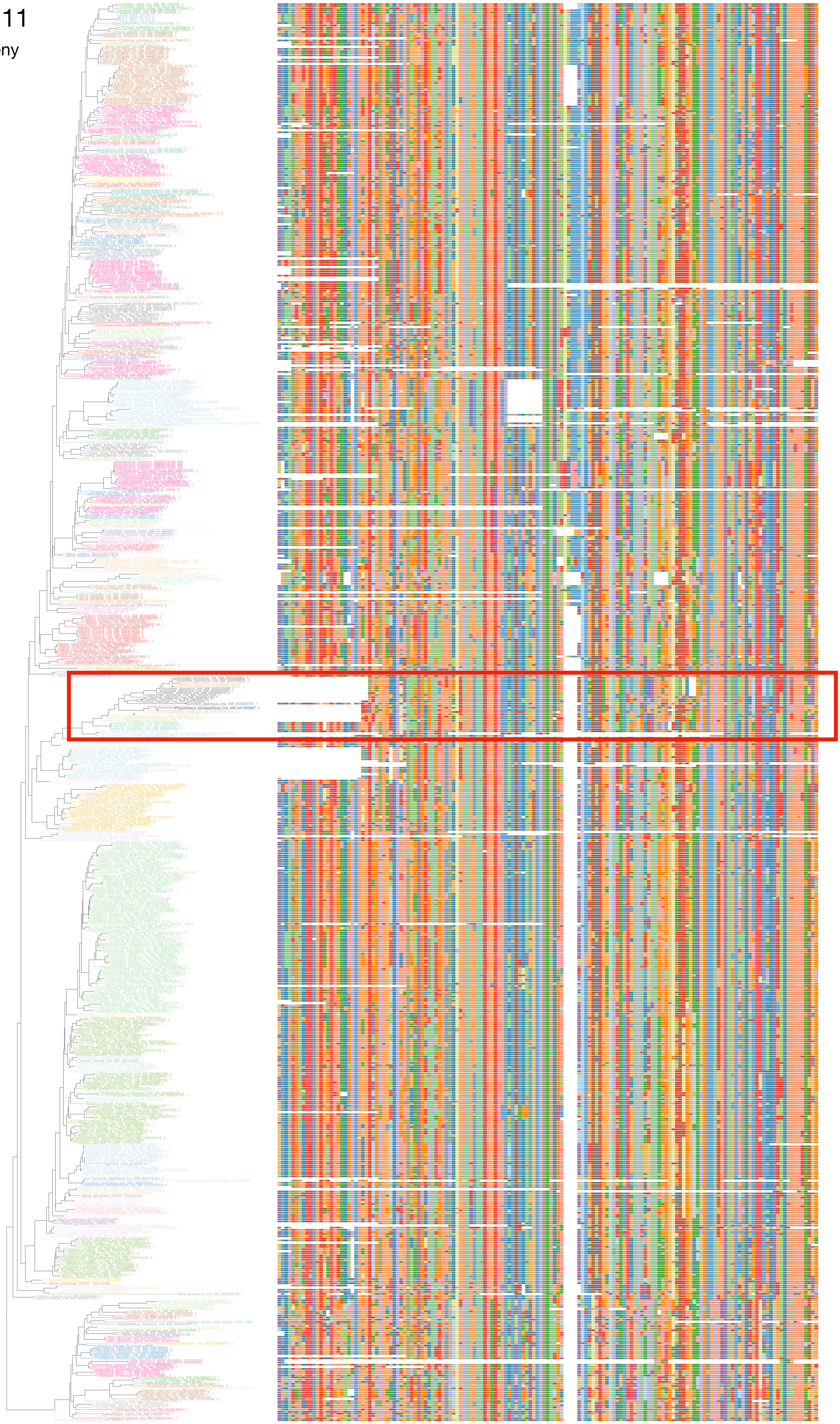

### N5.HOG0019111

« Best-match »
phylogeny
and
protein alignment

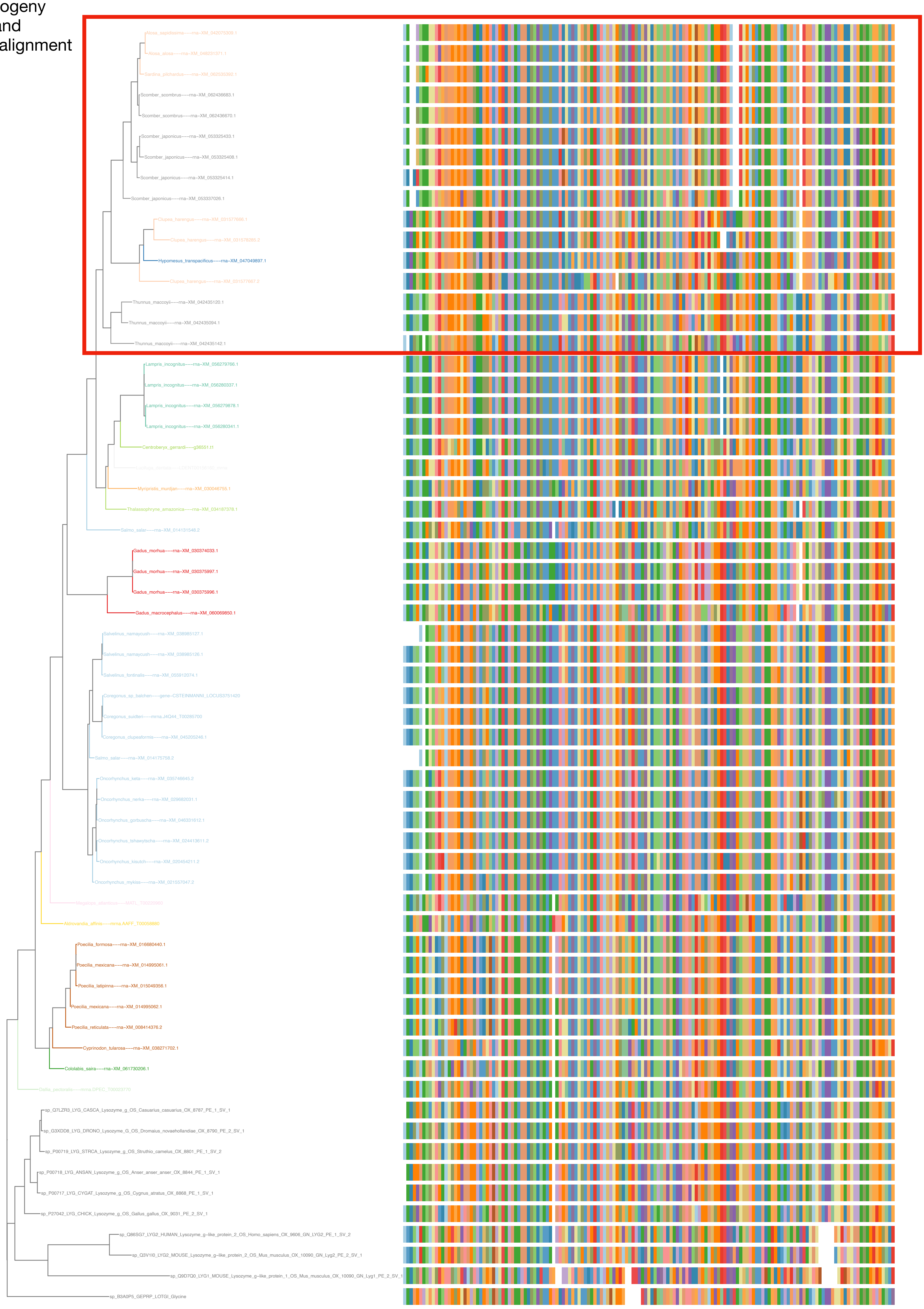

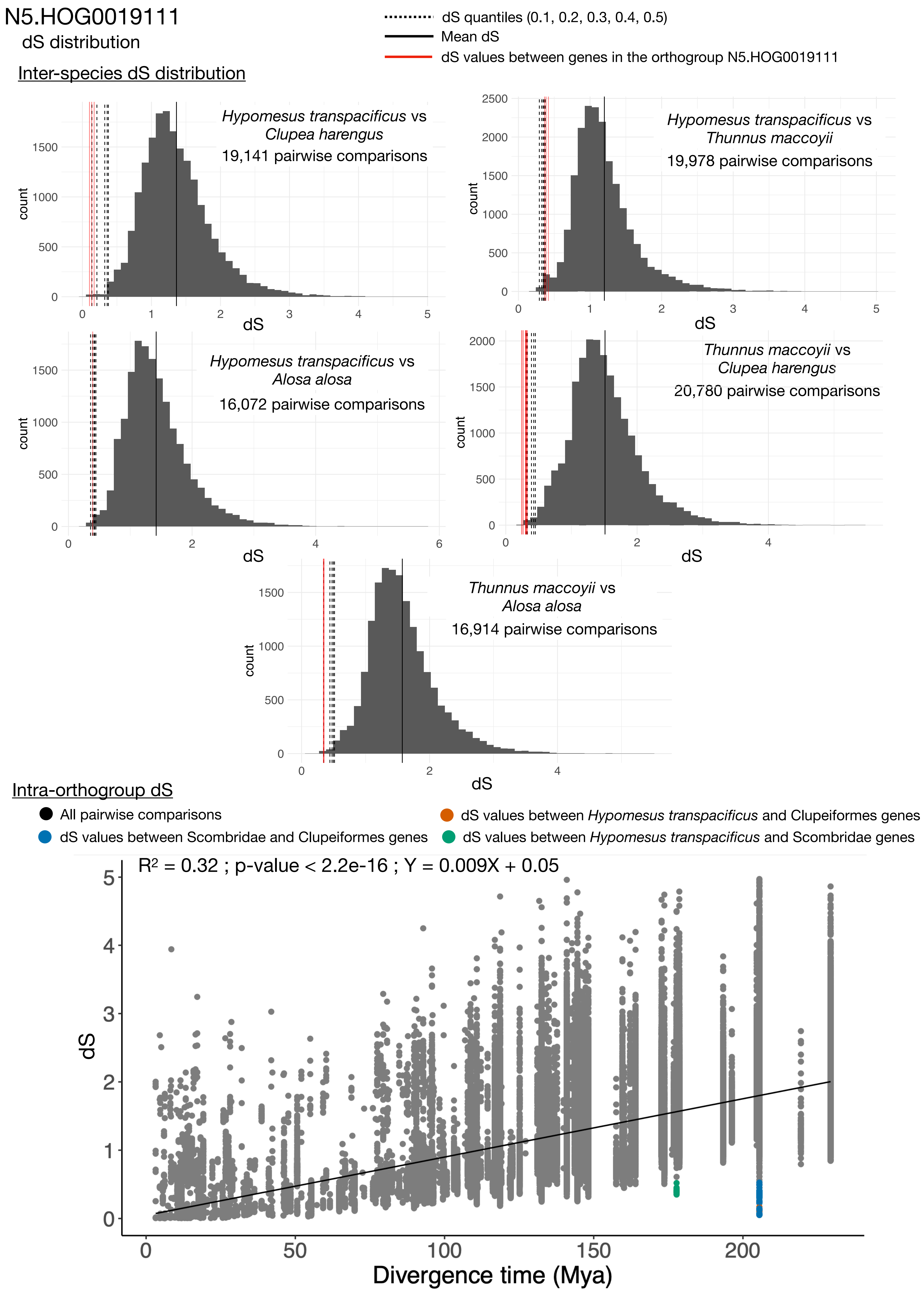

N5.HOG0019111
dS distribution
dS quantiles (0.1, 0.2, 0.3, 0.4, 0.5)
Mean dS
dS values between genes in the orthogroup N5.HOG0019111
Inter-species dS distribution
Hypomesus transpacificus vs Clupea harengus
19,141 pairwise comparisons
Hypomesus transpacificus vs Thunnus maccoyii
19,978 pairwise comparisons
Hypomesus transpacificus vs Alosa alosa
16,072 pairwise comparisons
Thunnus maccoyii vs Clupea harengus
20,780 pairwise comparisons
Thunnus maccoyii vs Alosa alosa
16,914 pairwise comparisons
count
dS
Intra-orthogroup dS
All pairwise comparisons
dS values between Scombridae and Clupeiformes genes
dS values between Hypomesus transpacificus and Clupeiformes genes
dS values between Hypomesus transpacificus and Scombridae genes
R² = 0.32 ; p-value < 2.2e-16 ; Y = 0.009X + 0.05
dS
Divergence time (Mya)

### N5.HOG0019111

#### Micro-synteny

##### #1 : Clupeiformes, Osmeriformes and close speices

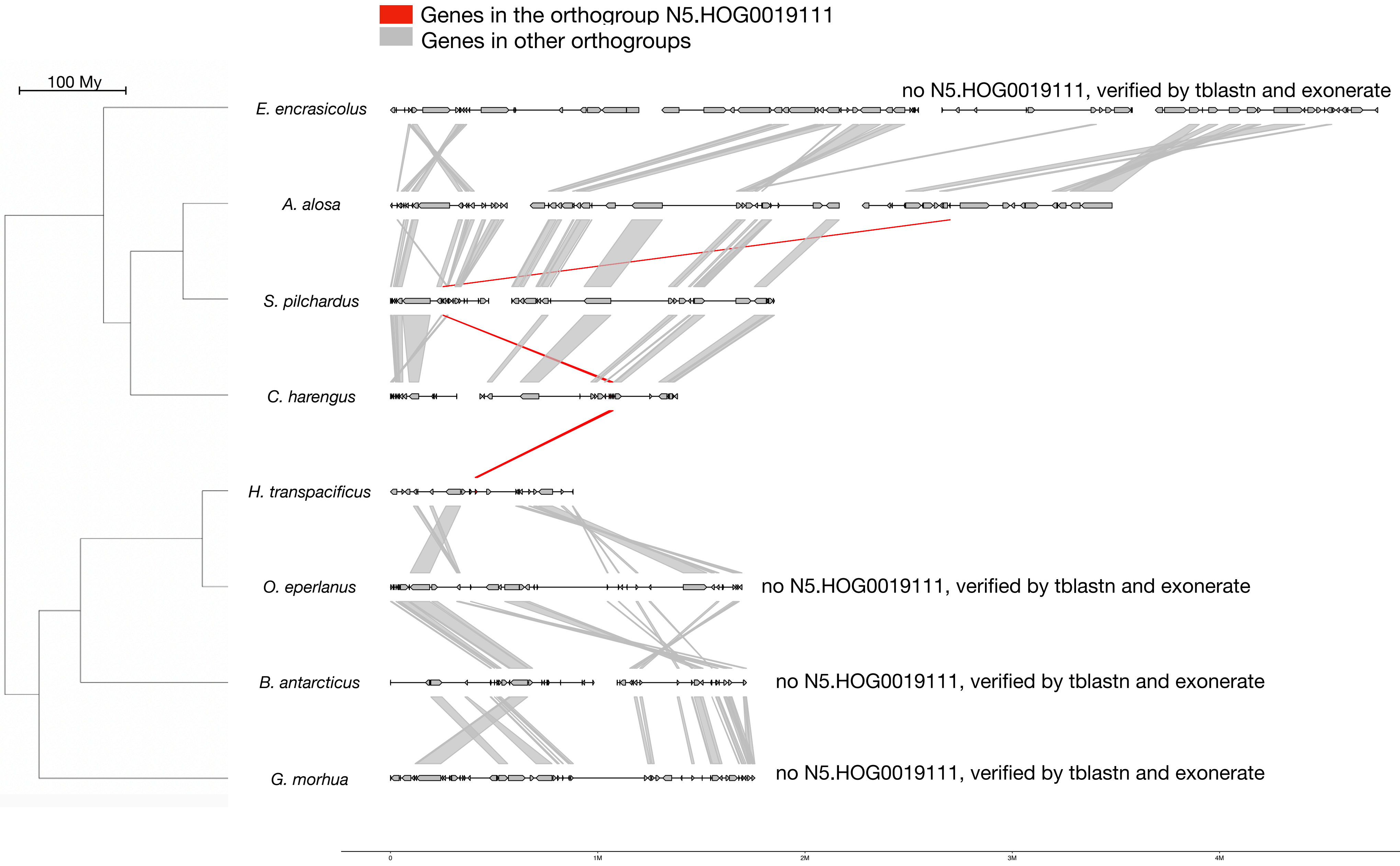


##### #2 : Clupeiformes, Scombridae and close speices

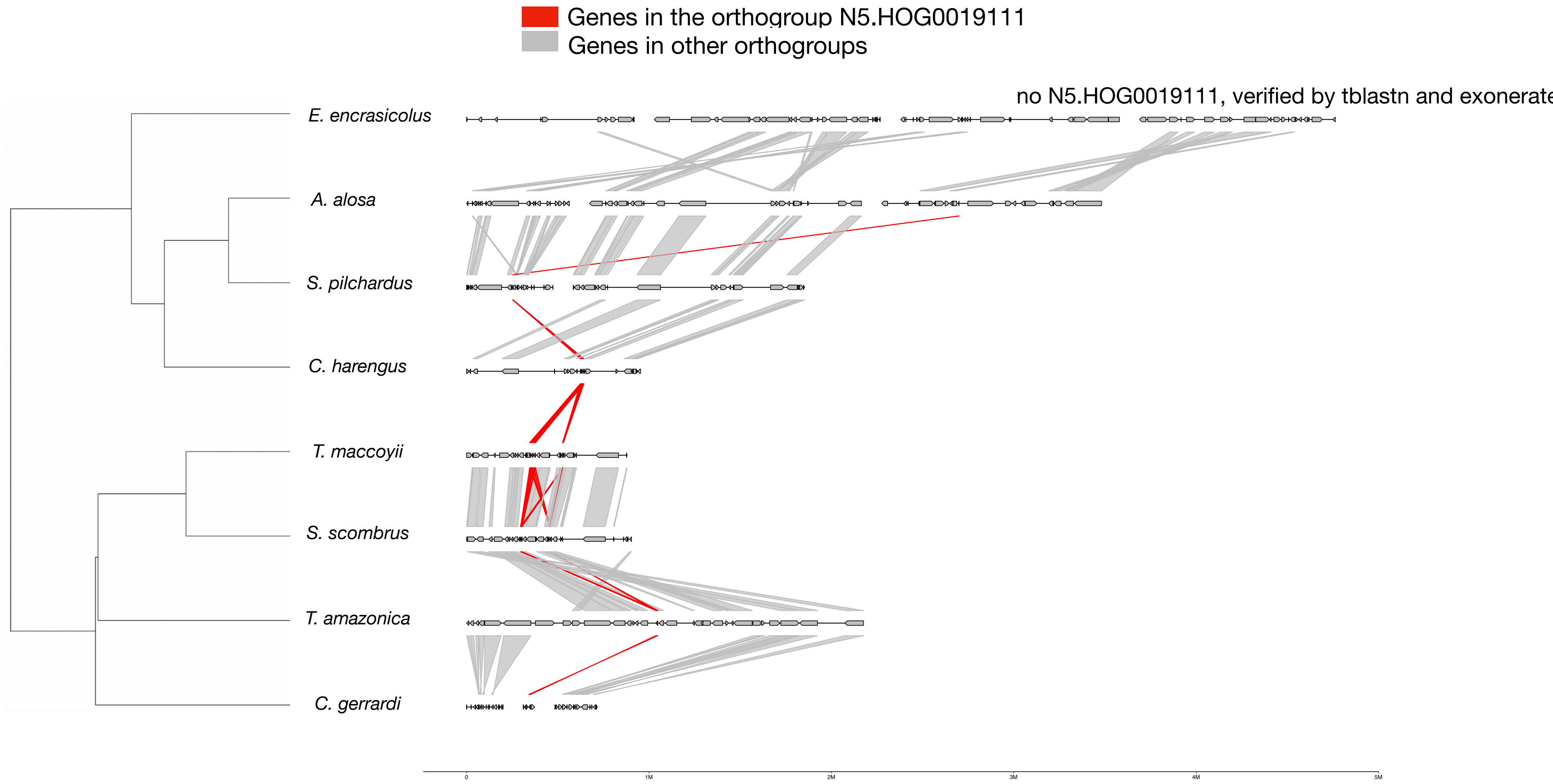

### N5.HOG0019111

#### Micro-synteny

##### #3 : Osmeriformes, Scombridae and close speices

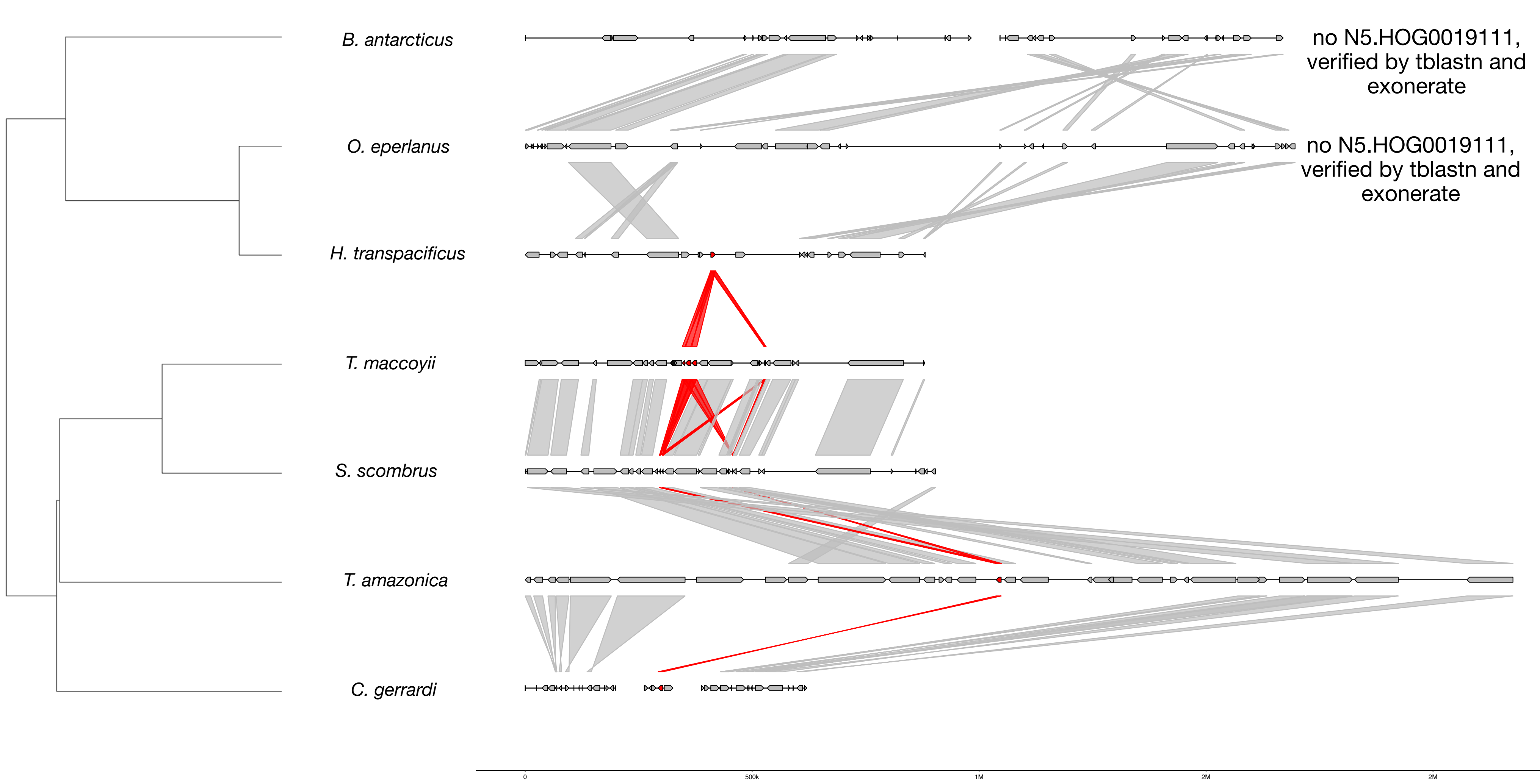


#### Zoom on *C. harengus, H. transpacificus* and *T. maccoyii*

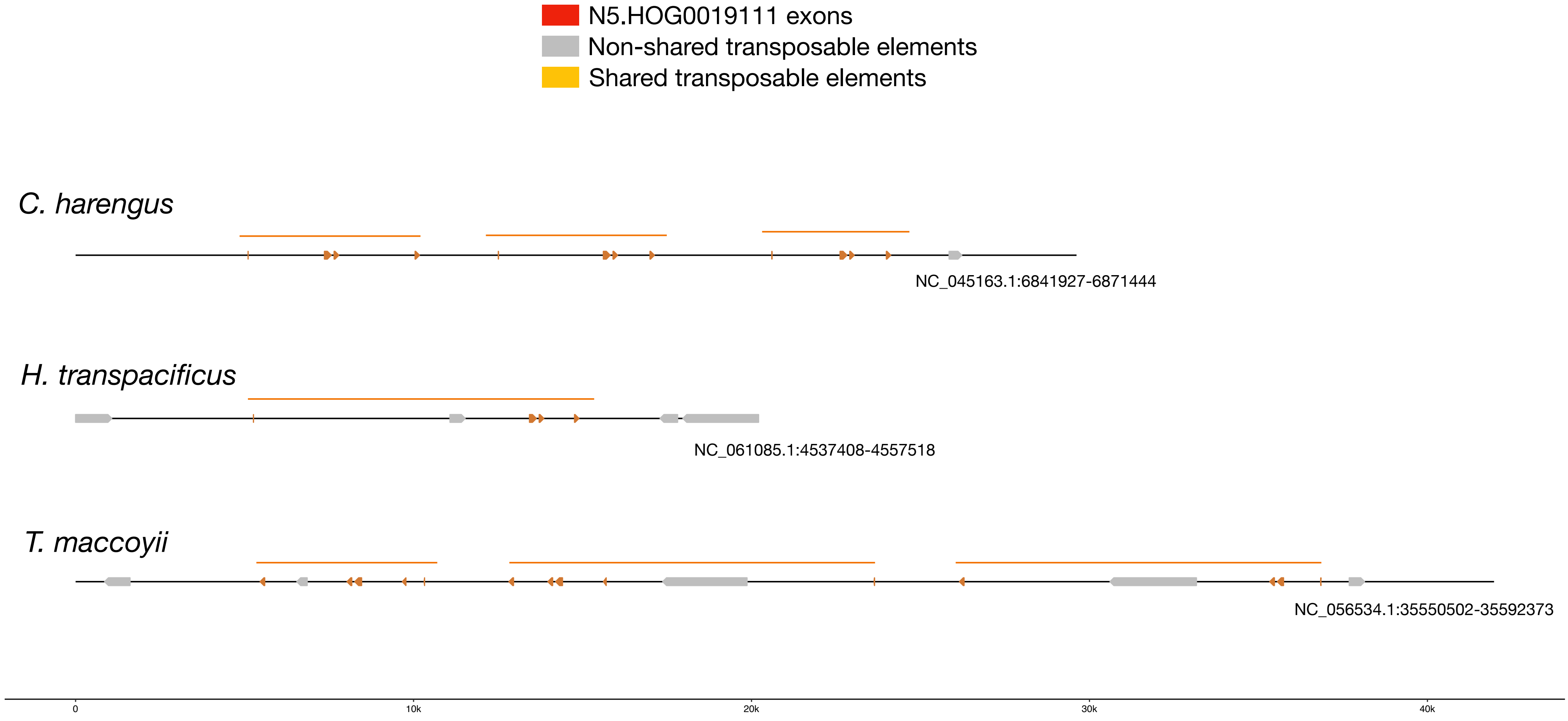

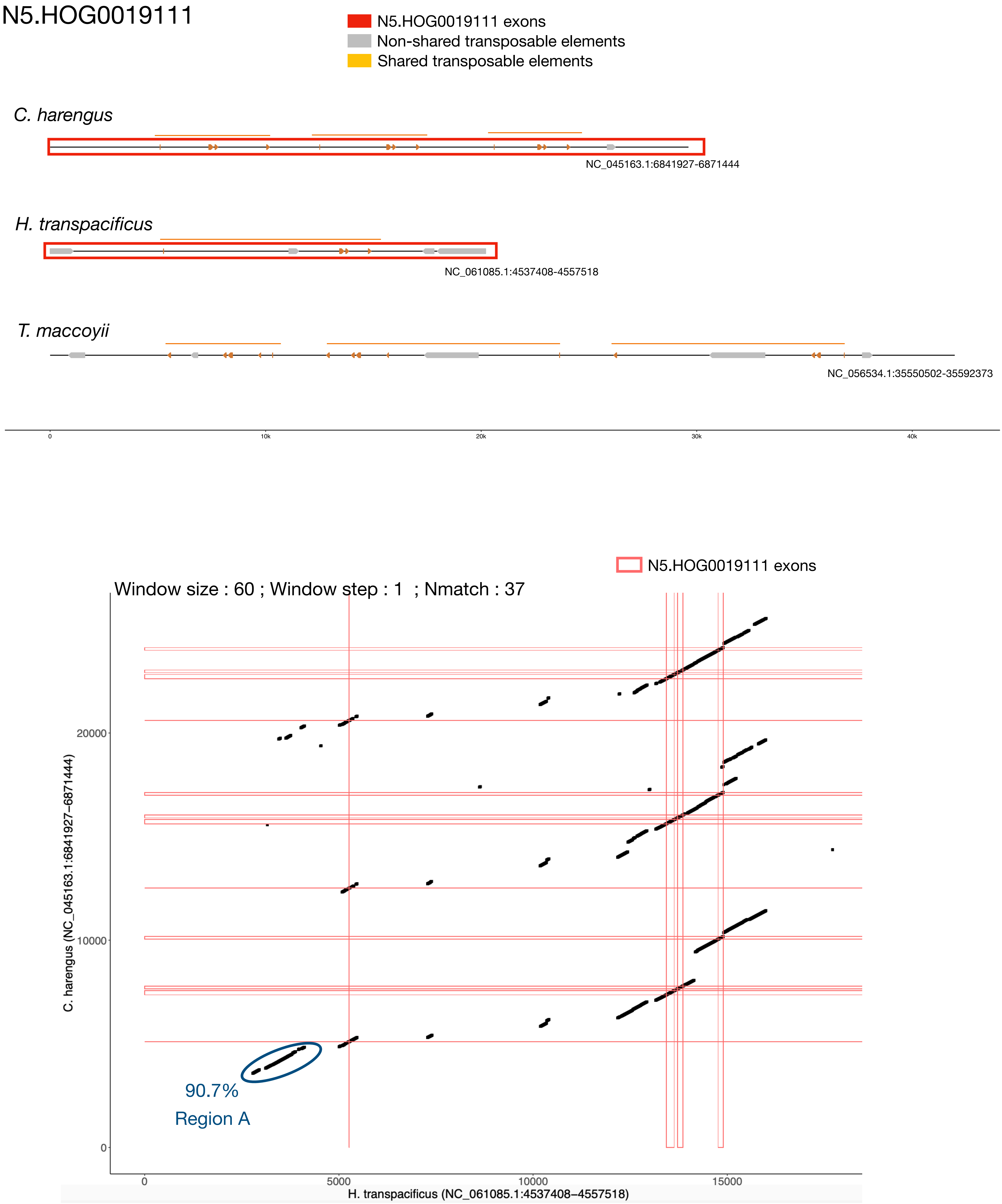
N5.HOG0019111
N5.HOG0019111 exons
Non-shared transposable elements
Shared transposable elements
C. harengus
NC_045163.1:6841927-6871444
H. transpacificus
NC_061085.1:4537408-4557518
T. maccoyii
NC_056534.1:35550502-35592373
0
10k
20k
30k
40k
N5.HOG0019111 exons
Window size : 60 ; Window step : 1 ; Nmatch : 37
C. harengus (NC_045163.1:6841927–6871444)
20000
10000
0
90.7%
Region A
0
5000
10000
15000
H. transpacificus (NC_061085.1:4537408–4557518)

### N5.HOG0019111 Blastn of conserved regions against all genome assemblies

#### Region A

| sseqid | pident | length | mismatch | gapopen | qstart | qend | sstart | send | evalue | bitscore |
|---|---|---|---|---|---|---|---|---|---|---|
| Hypomesus_transpacificus-NC_061085.1 | 100.000 | 1501 | 0 | 0 | 1 | 1501 | 4540107 | 4541607 | 0.0 | 2772 |
| Hypomesus_transpacificus-NC_061085.1 | 90.734 | 572 | 37 | 6 | 427 | 993 | 4798712 | 4798152 | 0.0 | 749 |
| Hypomesus_transpacificus-NC_061085.1 | 88.197 | 466 | 26 | 8 | 986 | 1430 | 4796922 | 4796465 | 1.71e-145 | 529 |
| Hypomesus_transpacificus-NC_061085.1 | 95.588 | 204 | 9 | 0 | 115 | 318 | 4798918 | 4798715 | 6.68e-85 | 327 |
| Clupea_harengus-NC_045163.1 | 88.811 | 715 | 45 | 15 | 436 | 1134 | 6845772 | 6846467 | 0.0 | 845 |
| Clupea_harengus-NC_045163.1 | 90.047 | 211 | 15 | 4 | 114 | 318 | 6845534 | 6845744 | 4.11e-67 | 268 |
| Clupea_harengus-NC_045163.1 | 91.549 | 142 | 4 | 2 | 961 | 1094 | 6861701 | 6861842 | 3.29e-43 | 189 |
| Clupea_harengus-NC_045163.1 | 88.095 | 126 | 12 | 3 | 1316 | 1441 | 6862176 | 6862298 | 2.01e-30 | 147 |
| Thunnus_maccoyii-NC_056534.1 | 86.826 | 167 | 17 | 3 | 1121 | 1282 | 35561219 | 35561053 | 5.51e-41 | 182 |
| Thunnus_maccoyii-NC_056534.1 | 86.228 | 167 | 18 | 3 | 1121 | 1282 | 35574538 | 35574372 | 2.56e-39 | 176 |
| Thunnus_maccoyii-NC_056534.1 | 86.228 | 167 | 18 | 3 | 1121 | 1282 | 35587747 | 35587581 | 2.56e-39 | 176 |
| Thunnus_maccoyii-NC_056534.1 | 89.855 | 138 | 8 | 1 | 962 | 1093 | 35567520 | 35567383 | 3.31e-38 | 172 |
| Thunnus_maccoyii-NC_056534.1 | 89.855 | 138 | 8 | 1 | 962 | 1093 | 35574809 | 35574672 | 3.31e-38 | 172 |
| Thunnus_maccoyii-NC_056534.1 | 89.855 | 138 | 8 | 1 | 962 | 1093 | 35580770 | 35580633 | 3.31e-38 | 172 |
| Thunnus_maccoyii-NC_056534.1 | 88.406 | 138 | 10 | 1 | 962 | 1093 | 35561490 | 35561353 | 7.17e-35 | 161 |
| Thunnus_maccoyii-NC_056534.1 | 84.524 | 168 | 20 | 4 | 1121 | 1282 | 35567249 | 35567082 | 7.17e-35 | 161 |
| Thunnus_maccoyii-NC_056534.1 | 84.524 | 168 | 20 | 4 | 1121 | 1282 | 35580499 | 35580332 | 7.17e-35 | 161 |
| Thunnus_maccoyii-NC_056534.1 | 85.443 | 158 | 11 | 9 | 113 | 261 | 35593497 | 35593343 | 1.20e-32 | 154 |
| Scomber_scombrus-NC_084971.1 | 83.439 | 157 | 17 | 7 | 113 | 261 | 14428648 | 14428493 | 1.21e-27 | 137 |
| Scomber_japonicus-NC_070579.1 | 82.911 | 158 | 17 | 8 | 113 | 261 | 25623098 | 25623254 | 1.56e-26 | 134 |
| Hypomesus_transpacificus-NW_025813826.1 | 100.000 | 64 | 0 | 0 | 53 | 116 | 2056926 | 2056863 | 4.38e-22 | 119 |
| Osmerus_eperlanus-NC_085019.1 | 95.833 | 72 | 2 | 1 | 53 | 123 | 9581668 | 9581739 | 5.66e-21 | 115 |

### N5.HOG0019111

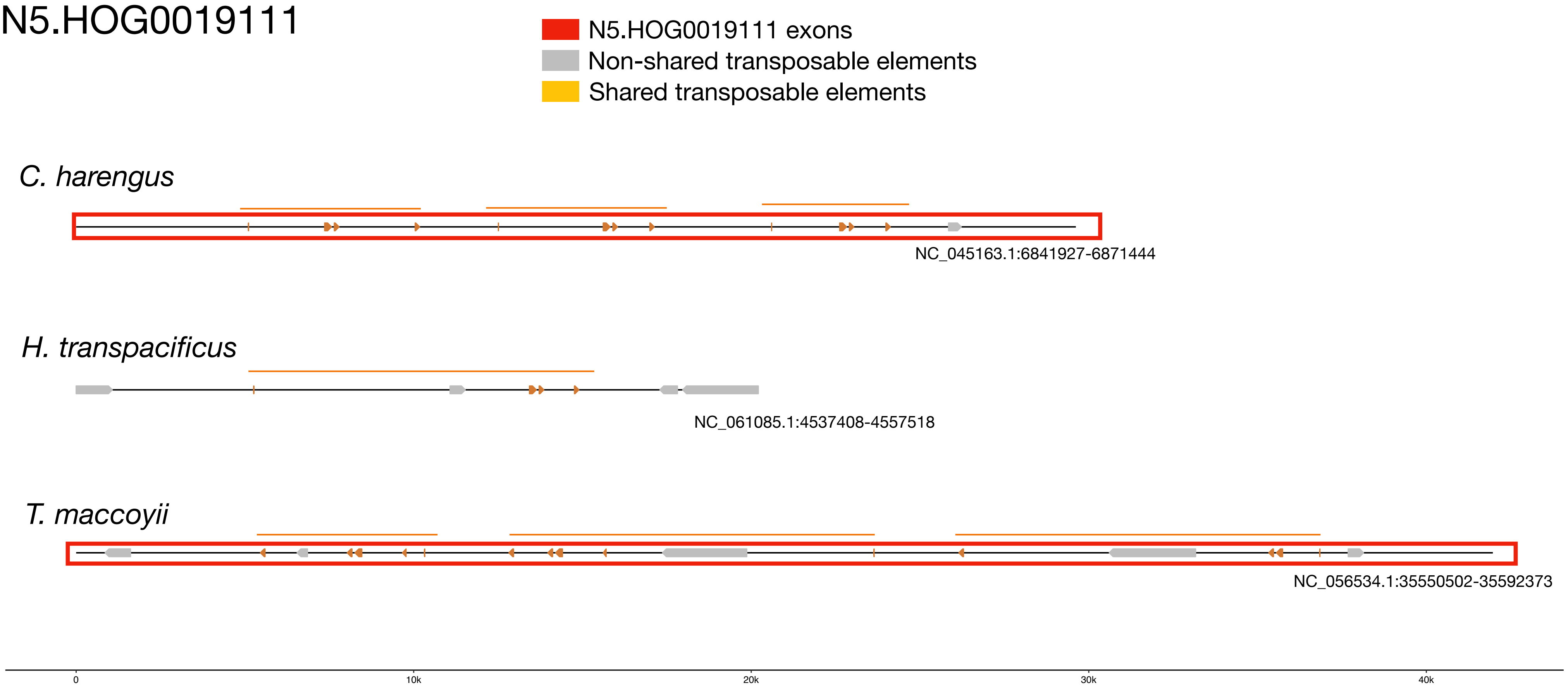


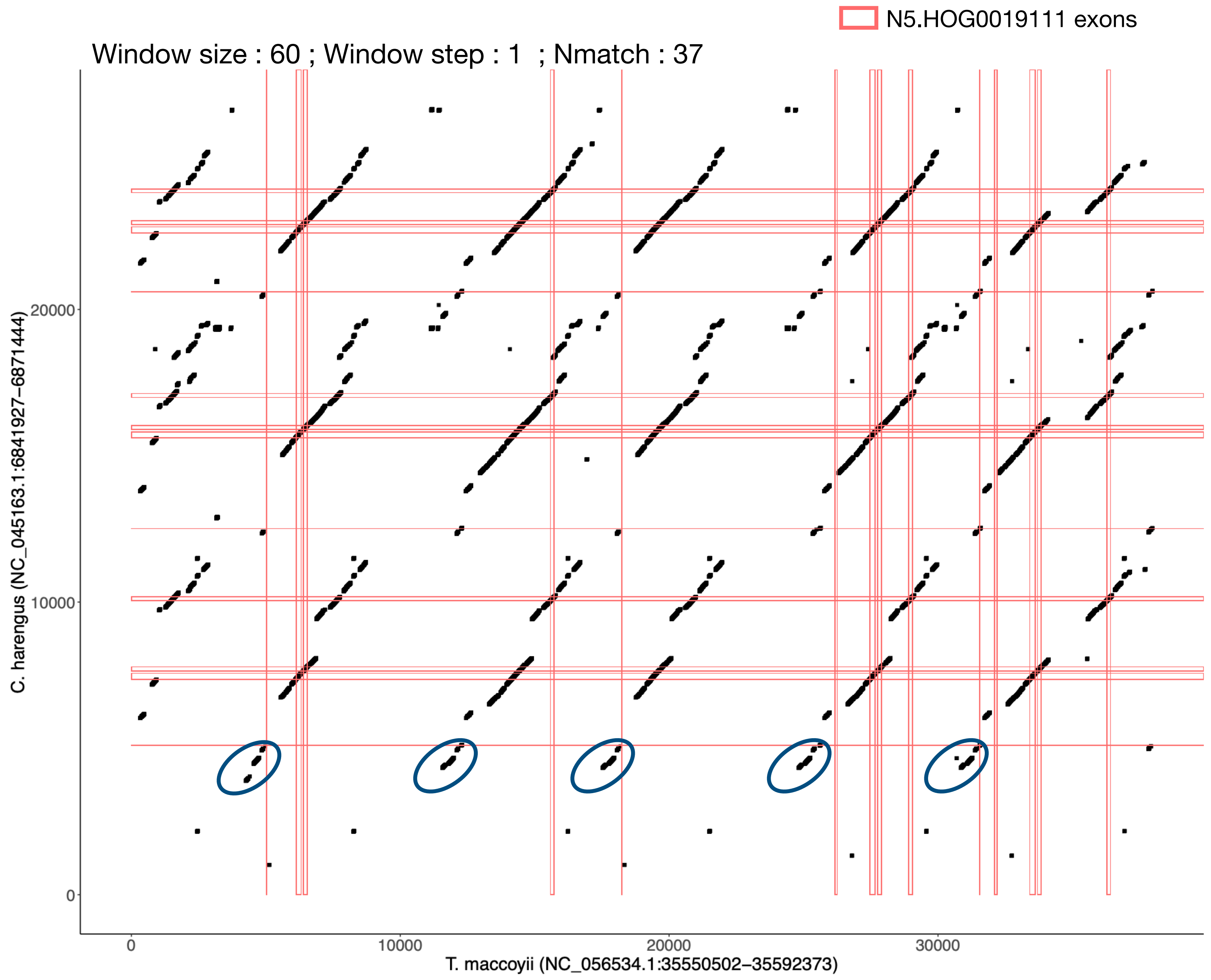

### N5.HOG0019111

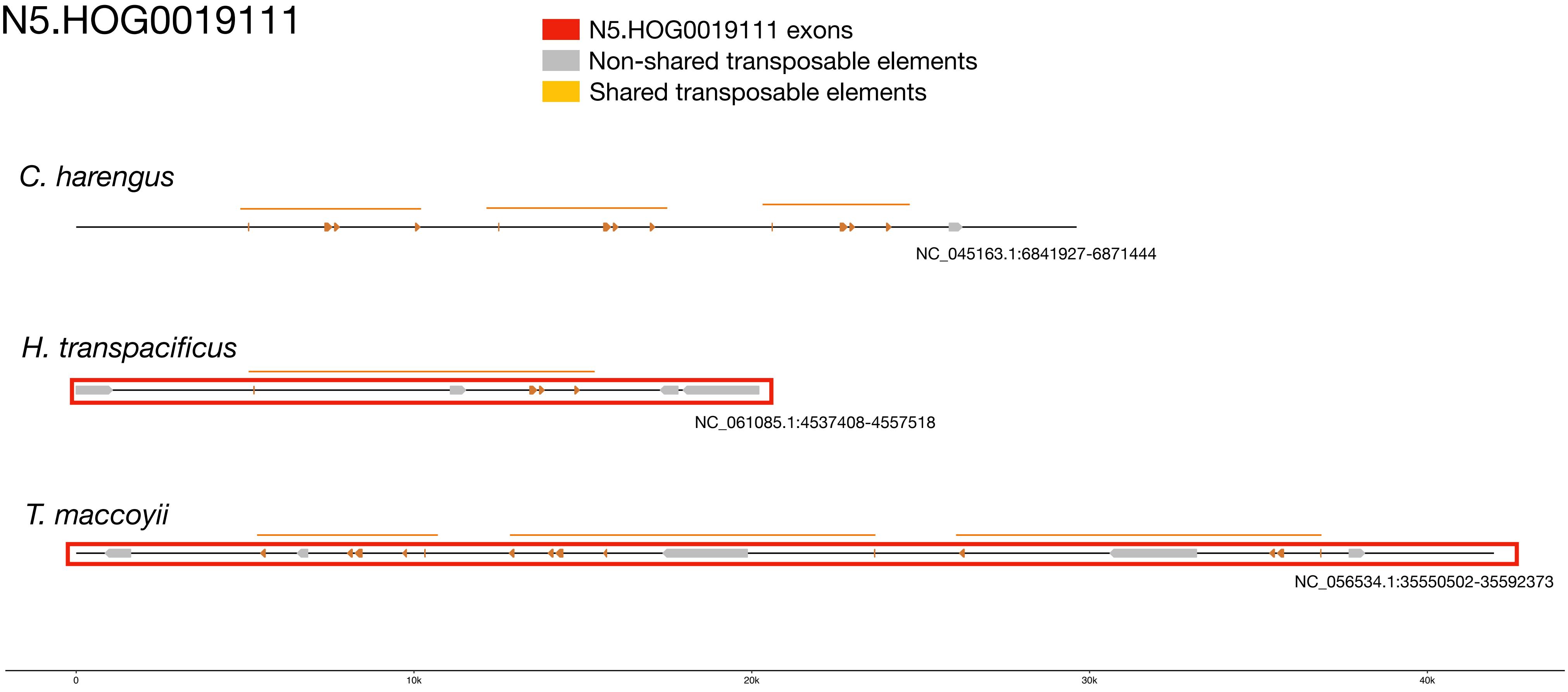


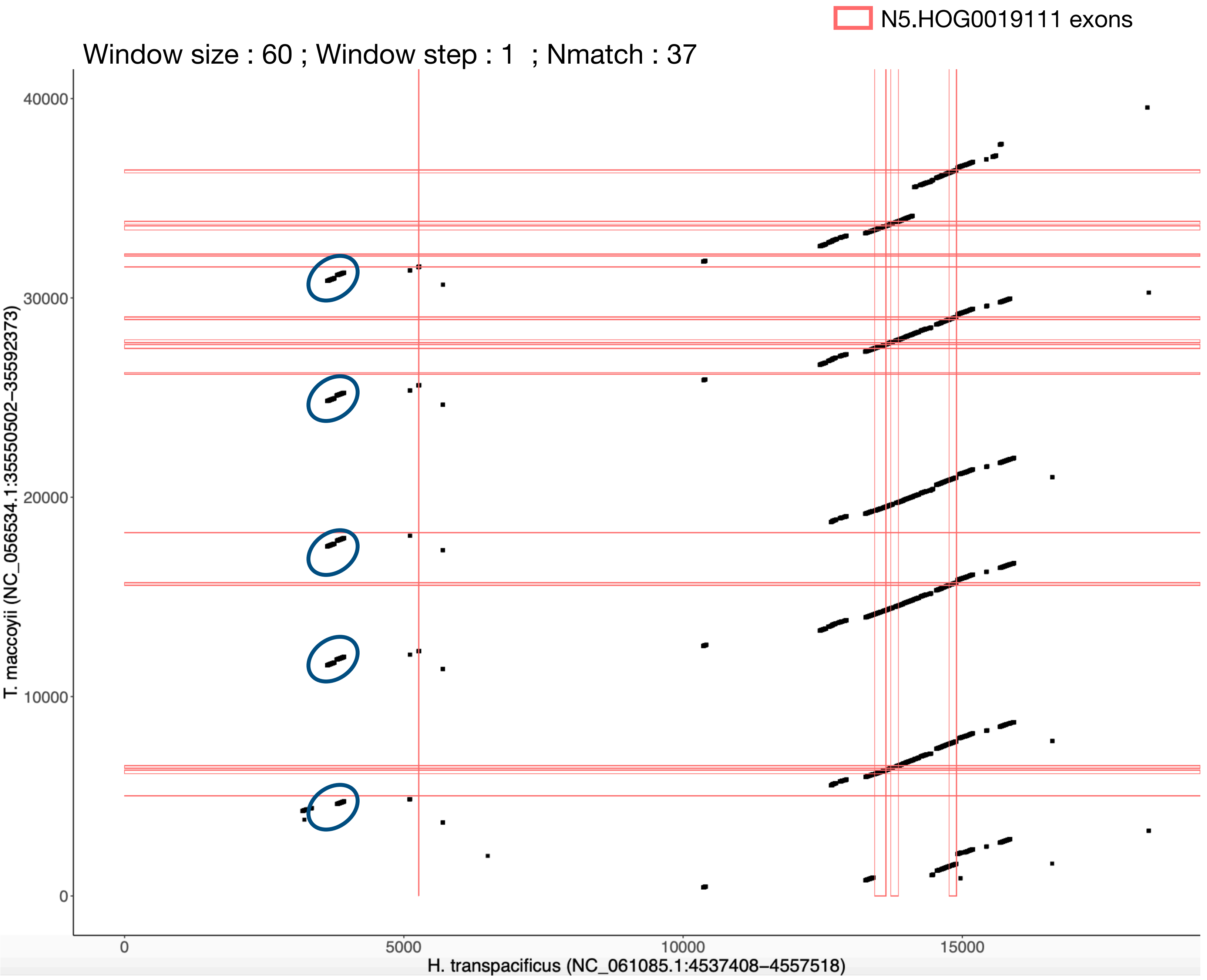

N5.HOG0019111

The gene is found in an alernative assembly of *H. transpacificus*

( 99.8% DNA identity)

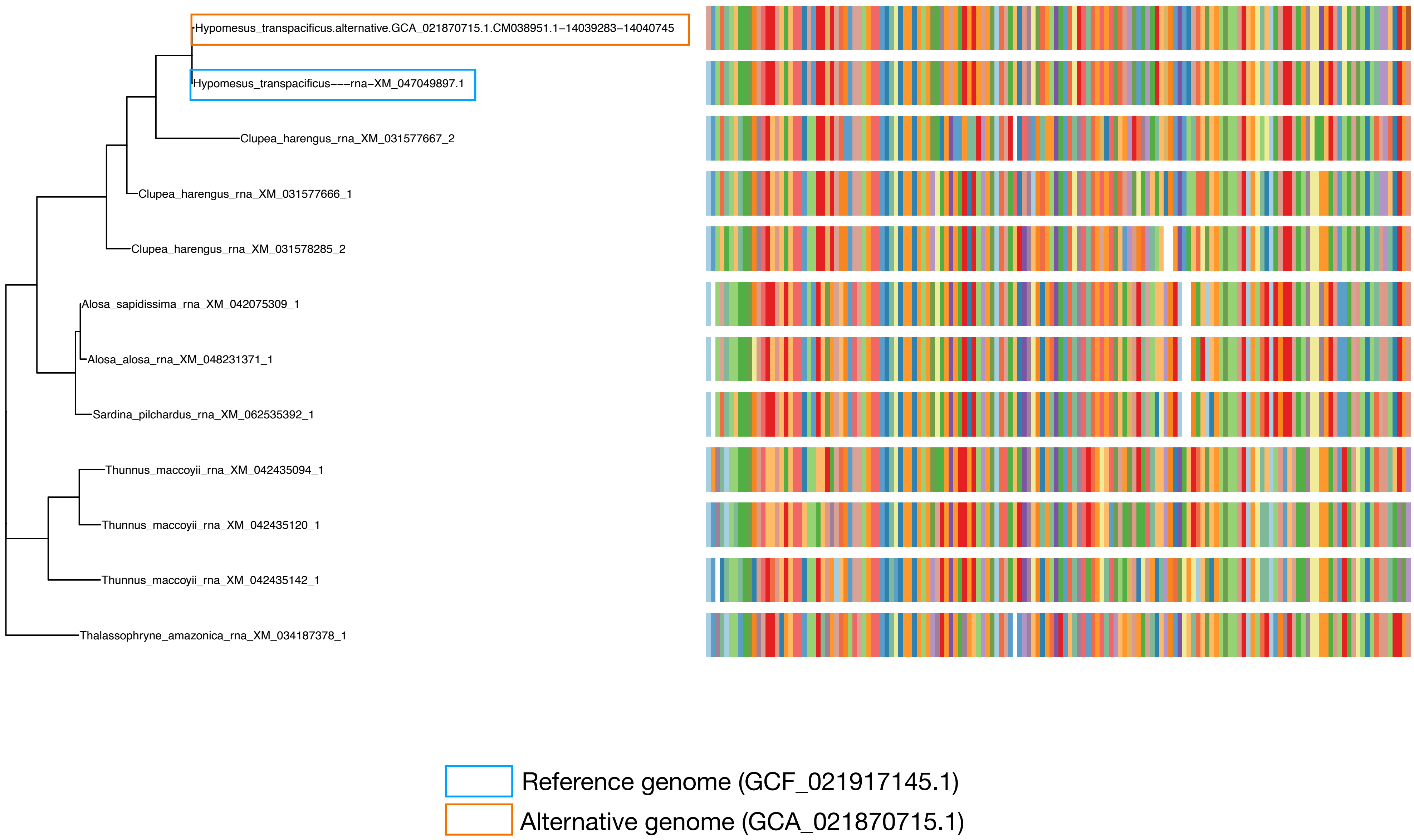

N5.HOG0019111

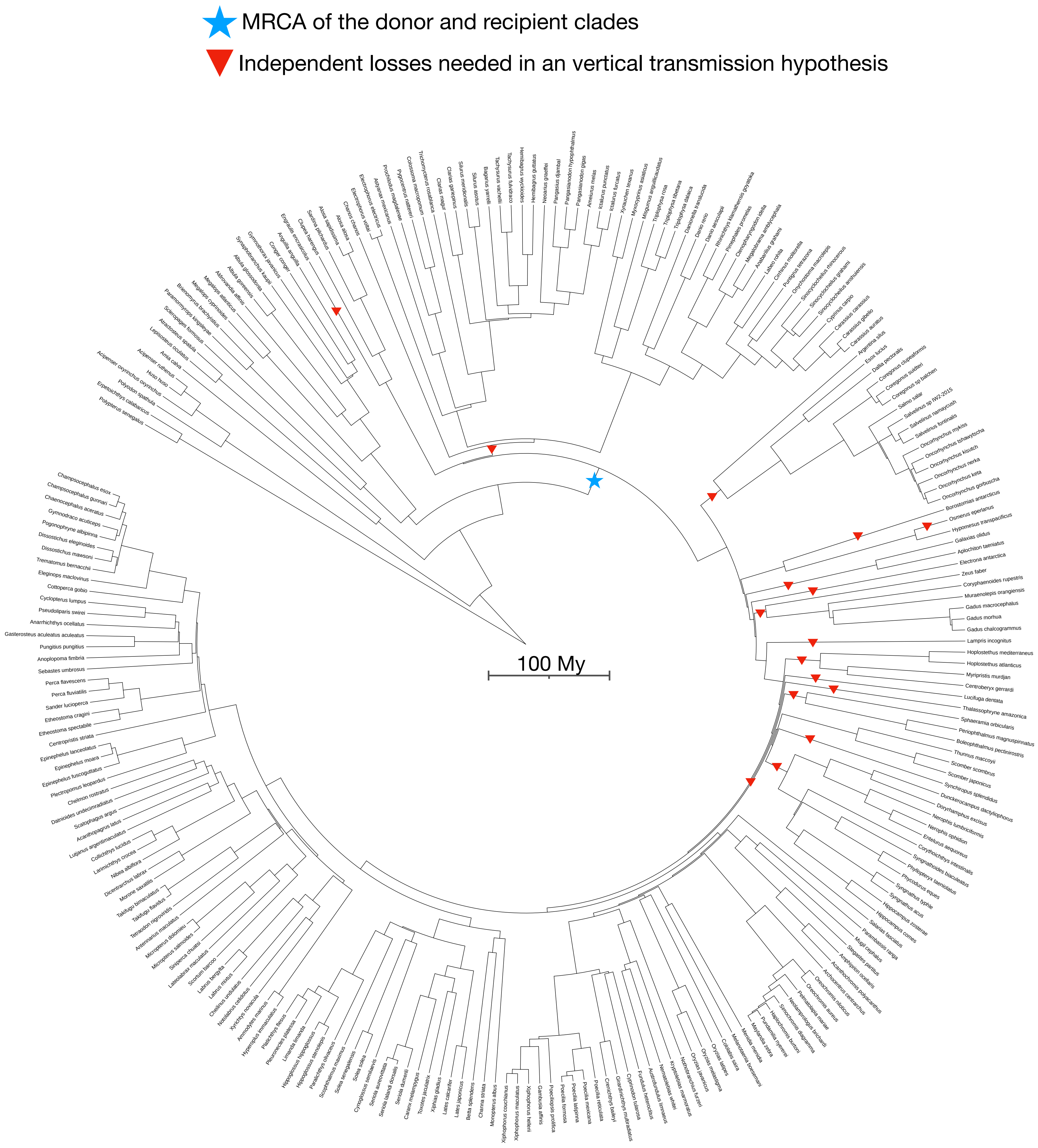
